# How robust are genomic offset predictions to methodological choices? Insights from perennial ryegrass

**DOI:** 10.64898/2026.03.02.708178

**Authors:** Marie Pegard, Susanne Lachmuth, Jean-Paul Sampoux, José Blanco-Pastor, Philippe Barre, Matthew C Fitzpatrick

## Abstract

Genomic offsets are increasingly used to quantify the mismatch between a population’s current genetic composition and the composition predicted under changed environmental conditions. While genomic offset is a promising tool for assessing climate maladaptation, the sensitivity of predictions to different methodological choices is not well understood.

In this study, we compared two fundamentally different approaches to detect outliers before predicting genomic offsets: Gradient Forest (GF, non-linear, non-parametric) and Canonical Correlation Analysis (CANCOR, linear, parametric). To do so, we used 457 natural populations of perennial ryegrass (*Lolium perenne* L.), an important agricultural forage species throughout Europe. Using a data set of 189,968 SNPs and 75 climatic variables, we experimentally validated genomic offsets against 105 phenotypic traits measured across three common gardens during multiple years. We also assessed the sensitivity of outlier detection and genomic offset predictions to the number and spatial distribution of sampled populations.

Both GF and CANCOR detected a substantial number of outlier loci associated with environmental gradients (2,113 and 653, respectively), with 429 loci identified by both approaches. When used to model spatial variation in genetic adaptation and estimate genomic offsets, the different outlier sets produce spatially congruent projections. We also found significant correlations between experienced genomic offset in the common garden predicted by both outlier sets and phenotypic traits, identifying traits that could serve as good fitness proxies for assessing climate risk. Analyses based on different population subsamples revealed that GF was less sensitive to sample size and geographic biases than CANCOR. Our findings provide practical guidance for designing genomic offset studies in both agricultural and natural systems and suggest that non-linear, non-paramateric methods like GF may be less sensitive to sampling design and therefore potentially more robust for predicting climate maladaptation.

## Introduction

Identifying the genetic basis of adaptation is essential for understanding and predicting how species will tolerate changing environments. Adaptive loci offer critical insights into stress tolerance and other key traits, enabling researchers to anticipate species’ responses to climate change, identify valuable genetic resources, and develop proactive conservation strategies (Gain et al., 2023; Lind et al., 2024; Wang et al., 2023). These genomic signatures also can inform the prediction of “genomic offset,” a metric used to estimate the potential risk of population maladaptation. Genomic offset quantifies the disruption of current genotype-environment associations by estimating the mismatch between a population’s contemporary genetic composition and the genetic composition expected in another environment (Capblancq et al., 2020a; Fitzpatrick et al., 2021; Fitzpatrick and Keller, 2015; Rellstab et al., 2021). Genomic offsets are increasingly used to assess population-level vulnerability, inform management decisions such as assisted gene flow or population translocation, and guide breeding programs for climate-resilient crops and forages (Borrell et al., 2020; Shryock et al., 2021; Wang et al., 2023).

The field of landscape genomics provides a diverse toolbox for predicting genomic offset, which involves a two-step process (Capblancq et al., 2020a): (1) identifying a set of adaptive loci using genotype-environment association (GEA) methods and (2) using those loci to model genetic turnover along environmental gradients and predict genomic offsets. Adaptive loci identification has evolved from agnostic FST outlier tests (Colli et al., 2014; Eckert et al., 2010) to more mechanistic association models like LFMM (Frichot et al., 2012; Frichot and François, 2015) and Bayenv (Günther and Coop, 2013) that account for population structure. More recently, multivariate methods have gained prominence, including linear, constrained ordination approaches like Redundancy Analysis (RDA)

(Capblancq and Forester, 2021) and Canonical Correlation Analysis (CANCOR) (Blanco-Pastor et al., 2020), which can effectively detect multilocus adaptation signatures. Concurrently, non-linear machine learning approaches, such as Gradient Forests (GF) (Ellis et al., 2012; Fitzpatrick and Keller, 2015), an extension of Random Forest, have been shown to capture complex, non-linear relationships between genetic variation and environmental variables. While the process of predicting genomic offset is conceptually divided into two steps, certain machine learning and multivariate approaches, such as Gradient Forests (GF) and Redundancy Analysis (RDA), integrate these steps.

Once a set of candidate loci is identified, statistical methods such as GF, Generalized Dissimilarity Modeling (GDM) (Fitzpatrick and Keller, 2015), RDA (Capblancq and Forester, 2021), LFMM (Frichot et al., 2012), and the Risk of Non-Adaptedness (RONA) approach (Borrell et al., 2020) can be used to predict genomic offsets. Despite their growing application, these methods have limitations and there remains a need for robust validation (Archambeau et al., 2026; Fitzpatrick et al., 2026; Láruson et al., 2022; Lind and Lotterhos, 2025). Several recent studies (Fitzpatrick et al., 2021; Láruson et al., 2022; Lachmuth et al., 2023; Lind et al., 2024; Francisco et al., 2024) have shown, counterintuitively, that loci identified through GEAs do not consistently improve offset predictions relative to randomly selected. Furthermore, the robustness of genomic offset estimates can be sensitive to spatial sampling strategies. For instance, sampling a geographic subset of a species’ range may fail to capture the full environmental niche, potentially leading to truncated adaptive gradients and introducing a major source of uncertainty in offset predictions (Aguirre-Liguori et al., 2023). Finally, validation is needed to determine the extent to which genomic offsets predict fitness-related phenotypes, a necessary step before predictions are applied in management contexts (Lachmuth et al., 2023; Lind and Lotterhos, 2024).

To address these gaps, we used perennial ryegrass (*Lolium perenne* L.) as study system. Beyond its status as a ubiquitous and economically important forage species, perennial ryegrass is cultivated worldwide and faces significant abiotic stress from climate change (Cullen et al., 2014; Miao et al., 2022; Rogers et al., 2024). What makes this system ideal for evaluating genomic offset models is the availability of a comprehensive suite of traits linked to species development that have been measured over several years and across several hundred populations in multiple common gardens - a depth and breadth rarely matched in genomic offset studies. Perennial ryegrass also offers an opportunity to explore how methodological decisions influence genomic offset predictions given that a previous comprehensive analysis identified 633 adaptive loci using the linear CANCOR method (Blanco-Pastor et al., 2020) and linked these loci to major climatic gradients and adaptive phenotypic traits. The wide distribution of its natural populations of ryegrass across diverse climatic conditions further makes it an ideal system for studying local adaptation (Carlier et al., 2009; Petermann and Buzhdygan, 2021; Tamburini et al., 2022).

Here, we build upon the study by Blanco-Pastor et al., 2020 to assess how methodological choices impact genomic offset models. Specifically, we aim to: (1) compare genomic offset projections derived from two sets of outlier loci, one set identified using CANCOR (linear) and a second set from GF (non-linear); (2) evaluate how the choice of GEA method (linear vs. non-linear) for outlier detection influences the relationship between predicted genomic offsets and a suite of key adaptive phenotypic traits, some of which may serve as proxies for population fitness; and (3) investigate how the spatial sampling of populations—using either the full range of the species versus geographic subsets—affects inferences obtained from these analyses. As such, this study provides practical guidance on how methodological choices in detecting adaptive loci, sampling populations, and modeling genomic offset influence the prediction of climate maladaptation for this important grassland agricultural species and other taxa more generally.

## Material and methods

### Methodological Framework

To address the complexity of genomic adaptation to climate change, we conducted four key analyses using a unique dataset of 457 perennial ryegrass populations, combining genomic, environmental, and phenotypic data. First, we used CANCOR and Gradient Forests (GF) to identify outlier loci across 189,968 SNPs and select relevant climate variables.

Second, we fitted two sets of GF models, one using the CANCOR-identified outliers and a second using the GF-identified outliers, along with their respective environmental variables. We refer to these models as GF_CANCOR_ and GF_GF_ respectively. These models were parameterized to explore relationships between SNPs and environmental predictors, with performance evaluated using out-of-bag R² and variable importance metrics.

Third, we experimentally validated the GF_CANCOR_ and GF_GF_ models by comparing predicted offsets e to development traits measured in three common gardens. We assessed how well genomic offsets predicted population performance under contrasted climatic conditions, benchmarking against geographic and environmental distances.

Fourth, we used the GF_CANCOR_ and GF_GF_ models to map genomic variation under current and future climate scenarios (RCP 4.5 and RCP 8.5, 2041–2070) from which we calculated genomic offsets. We assessed the similarity of these spatial predictions and visualized genetic composition shifts via PCA and Procrustes analysis, comparing past and future adaptive patterns. Additionally, we evaluated the robustness of our results to sampling effects by fitting models to subsets of populations selected using different sampling strategies (random subsets, geographic biases, and core collections). We also conducted functional enrichment analyses to explore the biological significance of outlier SNPs.

Collectively, our multi-faceted approach allowed us to comprehensively explore the influence of methodological choices, sampling strategies, and environmental gradients on genomic offset predictions and their reliability.

### Genomic material and data

We used 457 populations of perennial ryegrass (**Figure 1** and **Supplementary Table S1**), selected to capture the existing natural genetic diversity of perennial ryegrass across its native distribution range in Europe and the Near East. See Blanco-Pastor et al., (2020) for additional details.

**Figure 1:**
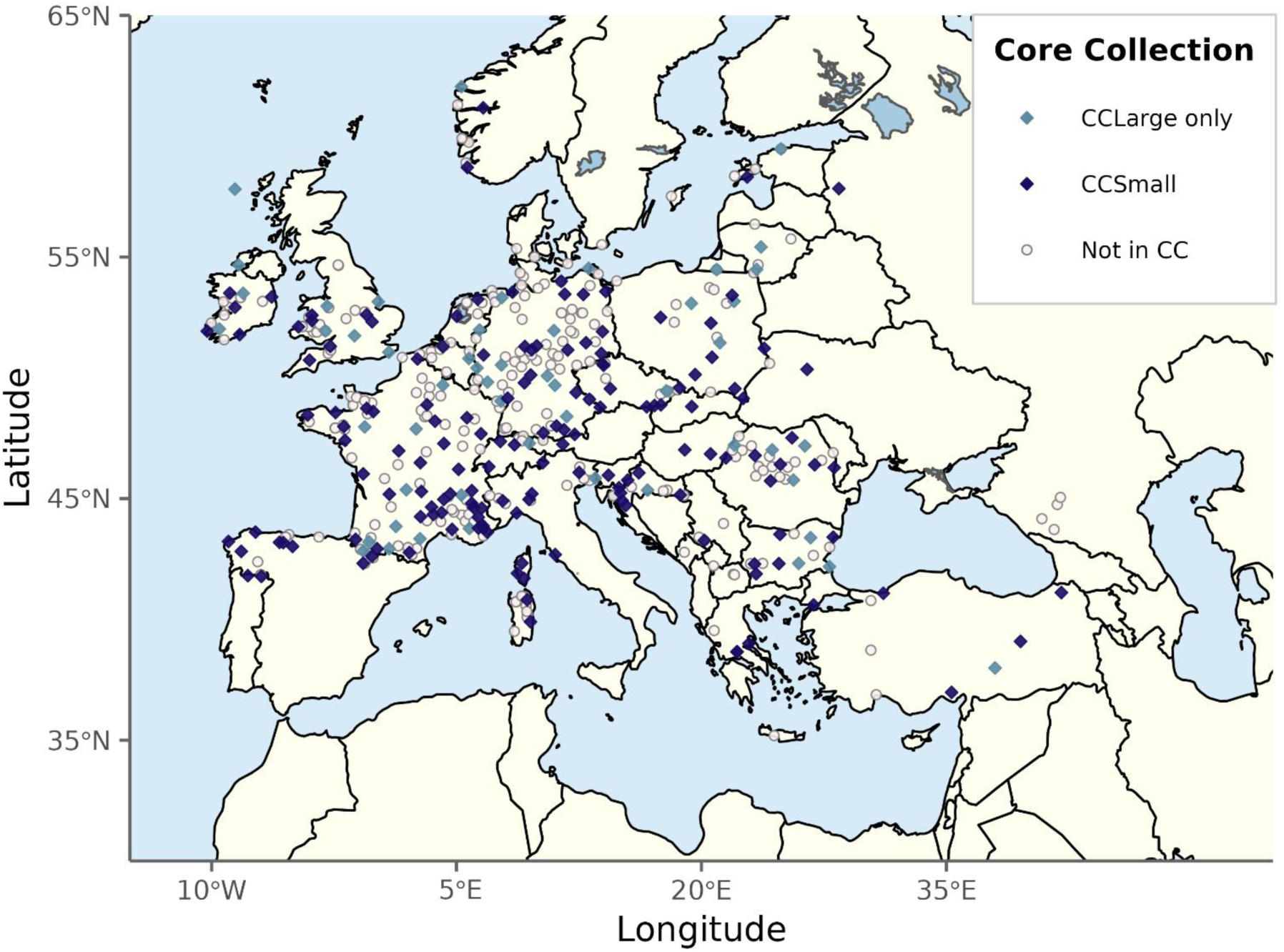
Geographic distribution of the 457 *Lolium perenne* populations used in this study, showing the two Core Collection cross-validation scenarios. Dark-blue diamonds represent populations included in the small core collection (CCSmall, n = 153), which is a subset of the large core collection. Light-blue diamonds indicate populations belonging to the large core collection only (CCLarge, n = 205). Grey circles represent the remaining populations not selected in any core collection. Both core collections were designed to maximise the genetic and environmental diversity represented within the dataset.

The genomic data were obtained using a GBS pool-Seq protocol based on Byrne et al., (2013). Briefly, leaf material from approximately 300 individuals per population was pooled for DNA extraction. Variants were called using the draft reference genome sequence of Byrne et al., 2015. Variant’s positions were compared with the last assembly (Nagy et al., 2022) and are reported in the supplementary material (**Supplementary Table S2**). LD pruning was performed by scaffold. Only one SNP was kept when two loci were linked with a kinship-corrected correlation exceeding 0.4 (Keep et al., 2020). SNP loci with a MAF greater than 5% in at least 10 populations were retained. The final dataset comprised alternative allele frequencies (AAFs) of 189,968 SNP in the 457 natural populations. Missing values were imputed using the mean allele frequency across populations.

### SNP Position Conversion, Gene Annotation, and Functional Enrichment Analysis

The SNP positions from the genome assembly v1.4 (Byrne et al., 2015) were converted to the corresponding positions in the genome assembly v2.6.1 (Nagy et al., 2022) using a BLAST search of the v1.4 scaffolds against the chromosome assembly v2.6.1. The positions of SNPs were then compared with gene positions in the genome. Gene annotations, including Gene Ontology terms, were retrieved from the Plaza Monocots database (available at https://bioinformatics.psb.ugent.be/plaza.dev/instances/monocots_05/download).

Each outlier SNP (see below) in v2.6.1 was mapped to candidate gene sequences, and the corresponding Gene Ontology terms were extracted. A SNP was considered linked to a gene if it was located either within the gene or within 1,000 base pairs (bp) of the gene’s start or end. SNPs located beyond this threshold were classified as not linked to any gene. This systematic annotation of outlier SNPs enabled the assessment of their potential functional implications based on gene proximity and associated Gene Ontology terms.

To further explore the functional significance of these genes, Gene Ontology terms enrichment and comparative analysis of gene clusters were performed using the clusterProfiler package (Yu et al., 2012) in the R environment. Genomic annotations were accessed via AnnotationHub and AnnotationDbi (Morgan et al., 2005; Pagès H et al., 2025), ensuring the use of standardized and up-to-date annotation data. Statistical testing for over-representation of Gene Ontology terms was conducted, with multiple testing correction applied to control the false discovery rate (FDR).

### Environmental variables

We characterized the environmental conditions at the sites of origin of the 457 perennial ryegrass populations using 75 environmental variables that described aspects of climate (**Supplementary Table S3**). This dataset included the 19 standard bioclimatic variables (Bio1–Bio19) for the period 1989–2010 (Fick and Hijmans, 2017) as well as an additional 8 bioclimatic variables that describe aspects of evapotranspiration in every season (Blanco-Pastor et al., 2020). We also included 48 additional variables across three broad categories: ecophysiological indices (e.g., cumulative and daily water stress, growth period lengths, heat/drought stress durations, and growing-degree-days), seasonal climate descriptors (e.g., seasonal evapotranspiration, precipitation, water balance, average temperatures, solar radiation, and extremes), and ETCCDI-derived indices (e.g., precipitation intensity, frequency of rainfall days ≥1mm, and maximum 1-day precipitation per season). To describe future conditions using our set of 75 variables, we obtained future climate data from the regional climate model CCLM4-8-17 (Bucchignani et al., 2016) under two Representative Concentration Pathways (RCP 4.5 and RCP 8.5), maintaining the same spatial resolution (0.05°) as the historical data.

### Phenotypic traits

For phenotyping, 377 of the 457 perennial ryegrass populations were sown in experimental gardens in three locations: Poel Island (PO) in Germany (53.990°N, 11.468°E) on 8 April 2015, Melle (ME) in Belgium (50.976°N, 3.780°E) on 2 October 2015, and Lusignan (LU) in France (46.402°N, 0.082°E) on 9 April 2015. Each population was sown in three replicated blocks, in a plot of 1 m^2^ micro-swards each. Trials were monitored until the end of 2017 at PO and ME and until the end of 2018 at LU. On the basis of their adaptive value, a set of 105 phenotypic traits were selected from the 145 phenotypic traits of Blanco-Pastor et al., 2020. This set included traits related to vigor after sowing, plant morphology, sward density, phenology, reproductive investment, vegetative growth dynamics, regrowth after cutting, susceptibility to abiotic and biotic stresses, persistence dynamics, chemical composition of aerial biomass, and leaf lamina traits. Phenotypic trait measurements are available in **Supplementary Table S4**. To account for missing phenotypic data, we performed imputation using a Genomic Best Linear Unbiased Prediction (GBLUP) model (Whittaker et al. 2000; Meuwissen et al. 2001). This genomic-based mixed model allowed us to estimate missing values for each trait based on the genetic relationships between populations:

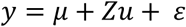

where y represents the phenotypes, μ the global mean, u the vector of random additive effects following 𝑁(0, 𝐺𝜎_𝑎_^2^) with 𝜎_𝑎_^2^ being the additive variance and G the genomic relationship matrix between accessions, ε is the vector of residual effects following 𝑁(0, 𝐼𝜎_𝑒_^2^) with 𝜎_𝑒_^2^ being the residual variance.

Z represents identity matrices linking the plots to the random effects. The genomic relationship matrix (G) was based on VanRaden (2008), adapted to use allele frequencies instead of allele dosage (Ashraf et al., 2014). The genotyping matrix with AAF (M) was standardized by the minimum allele frequency (P) to obtain the genotyping matrix (Z), which was then used to compute 𝐺, as follows:

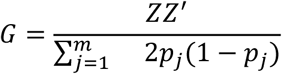

The denominator serves as a scaling parameter, representing the sum of the expected SNP variance across genotypes (Ashraf et al., 2014). Here, 𝑚 denotes the number of markers, 𝑝_𝑗_ the frequency of the 𝑗^𝑡ℎ^ marker.

### Canonical Correlation Analysis (CANCOR)

Canonical Correlation Analysis (CANCOR) is a multivariate statistical method designed to identify linear associations between two sets of variables. In this study, we applied CANCOR to simultaneously analyze genomic loci, environmental predictors, and phenotypic traits (e.g., fitness and adaptive traits measured in common gardens). By extracting canonical variates that maximize the correlation between these datasets, CANCOR identifies loci whose allele frequencies co-vary with both environmental gradients and phenotypic responses. CANCOR assumes linear and continuous relationships between genomic, environmental, and phenotypic variables, making it particularly effective for detecting broad-scale adaptive patterns where linear trends dominate.

### Outliers detection with CANCOR

We followed the methodology of Blanco-Pastor et al., 2020 to select outlier loci using CANCOR. Briefly, to investigate adaptive diversity, we examined the association of genomic polymorphisms with environmental variables (GEA) and phenotypic traits (GWAS). First, the slopes describing the relationships between genomic polymorphisms and environmental variability, and between genomic polymorphisms and phenotypic variability, were estimated using linear mixed models. We then analyzed the correlation between these two sets of slopes using CANCOR. Finally, the significance of outlier loci was tested using a Χ^2^ test on Mahalanobis distances, following the method of Luu et al., (2017) and Capblancq et al., (2018). The outlier SNPs (hereafter called CANCOR outliers) were selected with a threshold of FDR = 0.1. Following the CANCOR analysis, we selected environmental variables based on their contribution to the canonical axes. Specifically, we retained variables with a projection norm ≥ 0.95 on the first two environmental canonical planes. Additionally, these variables had to exhibit a correlation of ±0.5 with at least one SNP.

### Gradient Forests

Gradient Forests (GF) is an extension of Random Forests (Breiman, 2001), a nonparametric, machine learning approach of designed to model complex, nonlinear relationships. GF inherits the robust statistical framework of Random Forests, providing reliable measures of model performance and variable importance (Ellis et al., 2012; Fitzpatrick and Keller, 2015) for relationships between allele frequencies and environmental predictors. By constructing an ensemble of regression trees, GF aggregates fitted patterns of allele frequency changes into non-linear, monotonic cumulative turnover functions for each environmental gradient, enabling the detection of environmental thresholds and statistical outliers in genomic data.

### Selection of Outlier SNPs

We followed the methodology of Fitzpatrick et al., 2021 to identify outlier SNPs. GF was fitted to alternative allele frequencies for all 189,968 SNPs across all environmental variables, yielding an R² value for each locus-variable pair (**Supplementary Table S6**). Empirical p-values were calculated by ranking these R² values against a null distribution derived from a set of 931 random SNPs. This set was used to establish an R² threshold corresponding to a p-value of 0.01, which was applied to select SNP outliers (hereafter “GF outliers”). This set was first randomly selected, and then we calculated the correlation matrix (r²) between them to retain only those with an r² value lower than 0.35.

### Model Fitting and Parameterization

GF models were fitted to explore relationships between SNP outliers and environmental variables. Two sets of models were constructed: (1) GF_CANCOR_, using the CANCOR-identified outliers and the subset of environmental variables selected by CANCOR; and (2) GF_GF_:, using the GF-identified outliers and the full set of environmental variables. For both models, we used 1,000 regression trees, a variable correlation threshold of 0.5 (Ellis et al., 2012), and an *mtry* parameter of 10. All other parameters were set to default values.

### Evaluation of Model Performance and Variable Importance

The predictive performance of GF for each SNP was evaluated using the out-of-bag R², which estimates generalization error. SNPs were ranked based on how well environmental gradients explained allele frequency changes. Predictor importance was quantified by measuring the decrease in model performance upon random permutation of each predictor, with a conditional approach for correlated variables (Ellis et al., 2012). GF also calculates “raw split importance” values, which were aggregated to construct nonlinear turnover functions for each environmental variable. These functions highlight gradients strongly associated with genetic variation and describe rate and magnitude of allele frequency changes along these gradients.

### Projection of Genomic Variation

To project current and future genomic variation, we used cumulative importance curves derived from GF to transform environmental data into units of genetic turnover. GF models (GF_CANCOR_ and GF_GF_) were trained using current (1989-2010) climatic conditions and applied to future climate projections from the regional climate model CCLM4-8-17 (RCP 4.5 and RCP 8.5, 2041–2070) using the *predict* function in the gradientForest R package.

### Adaptive Genetic Composition

We visualized spatial patterns of adaptive genetic composition using Principal Components Analysis (PCA) on transformed environmental variables. To preserve the importance of each gradient, we centered but did not scale the data. The resulting adaptive compositional variation was displayed using an RGB color palette, where the first three principal components correspond to red, green, and blue channels, respectively. To compare past and future genetic patterns, we employed Procrustes superimposition on PCA ordinations, rotating matrices to minimize squared distances between sites in genetic space (Peres-Neto and Jackson, 2001). Procrustes residuals ensured comparability of colors in genetic space inside a model (either GF_GF_ or GF_CANCOR_), enabling visualization of predicted genetic composition over time.

### Local Genomic offset

We quantified the genetic difference between current and future conditions, i.e., the genomic offset, as the Euclidean distance between current (1989–2010) and future (2041-2070) transformed environments, focusing on local offset (Gougherty et al., 2021; Lachmuth et al., 2023) to assess potential *in situ* climate maladaptation. The resulting values were mapped across the study area to visualize spatial variation in predicted maladaptation under each emissions scenario (Fut4.5 and Fut8.5) and for each GF model (GF_GF_ and GF_CANCOR_).

### Map Comparison Analysis

We conducted a comprehensive comparison of the GO maps predicted by the two GF models (GF_GF_ and GF_CANCOR_) between current (1989-2010) and future climatic conditions for both emissions scenarios (Fut4.5 and Fut8.5). To quantitatively assess the similarity between these spatial predictions, we used the Spatial Index of Pattern (SIP; Wiederholt et al., 2019). This statistic ranges from 0 (no similarity) to 1 (identical) patterns and calculates the local correlation between two raster maps using a moving-window approach, even when the magnitudes between the two raster differ.

### Experimental Validation of Genetic Offset Predictions

#### Experienced genomic offset estimation

To experimentally evaluate the link between genomic offset (GO) and population fitness, we analyzed the relationship between field performance under common garden climatic conditions (LU, ME, PO) and the GO values predicted between the sites of origin and the common garden locations. Specifically, we used our fitted GF models (GF_GF_ and GF_CANCOR_) to transform the climatic conditions experienced by each population during the field trial. This “experienced” genomic offset was calculated as the Euclidean distance between the transformed climatic conditions at the population’s site of origin and those at the common garden site.

#### Estimation of predictive power

The predictive power was assessed by computing Pearson correlation coefficients between the experienced GO values and the phenotypic performance measured at each common garden site. Statistical significance of each correlation was evaluated using the associated p-values. To benchmark the predictive power of the experienced GO against null expectation, we compared it against two standard distances: (1) geographic distance, calculated as the geodesic distance in kilometers between the site of origin and the common garden; and (2) “naïve” environmental distance, calculated as the Mahalanobis distance using the untransformed versions of the same climatic variables used in the GF_GF_ and GF_CANCOR_ models.

### Impact of Population Sampling on Outlier Detection and Genomic Offset Estimation

The number, spatial distribution, and geographic bias of sampled populations could influence the selection of outlier loci, fitted relationships between environmental gradients and genomic variation, and predictions of genomic offset. To assess the effects of sample size, geographic bias, and population representation on model performance, we repeated the analyses described above on different subsamples of our ryegrass study populations.

### Population Subsampling Design?

#### Impact of the number of sampled populations

Randomized subsets (Data composition effects): To evaluate the robustness of outlier detection and genomic offset predictions relative to sample size, we generated 50 permutations for each of four subset sizes of 100, 150, 200 and 300 randomly selected populations, hereafter referred to as ‘random’ (**Supplementary Table S1**).

#### Impact of the sampling geographic representation

Geographically Biased Subsets (Spatial sampling effects): To investigate the impact of spatial sampling bias, i.e., sampling of populations from a subset of the range, which could result in truncation of environmental gradients, we created four geographically restricted subsets (**Supplementary Table S1**). These subsets were designed to reflect varying degrees of environmental heterogeneity: (1) UnbalW (Western Europe): 197 populations from France, Spain, Italy, England, Ireland, and Belgium; (2) UnbalE (Eastern Europe): 260 populations located in Eastern Europe; (3) UnbalN (Northern Europe): 255 populations located at latitudes north of 47.5°N; and (4) UnbalS (Southern Europe): 202 populations located at latitudes south of 47.5°N.

Core Collections (Diversity effects): Additionally, we utilized two core collections representative of the phenotypic and neutral genetic diversity of the FDset of 457 perennial ryegrass populations. The large core collection (CCLarge) consisted of 205 populations, while the small core collection (CCSmall) comprised 153 populations (**Figure 1** and **Supplementary Table S1**).

#### Comparative Analysis

To assess model robustness and variability, we compared the results obtained from all subset-based evaluations (random, geographic subsets, and core collections) against those derived from the FDset. This comparison provided a quantitative estimate of how sampling strategy influences the stability of outlier detection and the reliability of genomic offset predictions.

#### Precision and sensitivity of outlier detection

SNPs detected with the FDset either with GF or CANCOR served as the respective reference set of true positives (TP) for each method. For each subset, we then computed: (i) precision, defined as the proportion of detected SNPs that were also identified in the FDset (TP / n_detected); (ii) sensitivity, defined as the proportion of FDset detected SNPs recovered by the replicate (TP / n_global); and (iii) the false discovery rate (FDR), defined as the proportion of detected SNPs absent from the FDset (FP / n_detected). These metrics were computed separately for GF and CANCOR method and sampling configuration.

#### Sampling effects on local genomic offset prediction

We used the SIP to assess the sensitivity of the mapped GO predictions to population sampling design. We compared the maps derived from the FDset containing all populations to those generated using different subsets of the data.

#### Sampling effects on experimental validation

Correlations between experienced genomic offset and corrected phenotypes were then recomputed for each subset. These subset-derived correlations were compared to those obtained with the full dataset. The coefficient of determination (R²) of this relationship was used to quantify the stability of GO–phenotype associations across sampling configurations.

### Statistical Analysis

All statistical analyses were conducted in R (version 4.4.1) (R Core Team, 2024). The detection of biological composition patterns along environmental gradients was performed with the package gradientForest (version 0.1-34; Ellis et al., 2012), which calculates turnover functions from split-point importances provided by auxiliary package extendedForest (version 1.6.1; Ellis et al., 2012), based on randomForest (Liaw and Wiener, 2002). We used the extendedForest package and the gradientForest package to fit Gradient Forests models, estimate genomic offsets, and calculate variable importance. Outlier detection was performed using custom scripts based on the methodology described by Fitzpatrick et al., 2021, leveraging functions from the gradientForest package for model fitting and performance evaluation. The CANCOR analysis was performed using the R package vegan (version 2.6-8, Oksanen et al., 2001).

Principal Component Analysis (PCA) was performed using the stats package (version 4.4.1) and Procrustes superimposition was conducted with the vegan package (version 2.6-8; Oksanen et al., 2001) to compare genetic composition across time points.

Pearson correlation analyses were carried out using the base stats package in R. All visualizations were generated using ggplot2 (version 3.5.1; Wickham, 2016).

Functional annotation of genes was performed using the topGO (version 2.58.0; Alexa A and Rahnenführer J, 2025), clusterProfiler (version 4.14.4; Yu et al., 2012), AnnotationHub (version 3.14.0) and AnnotationDbi (version 1.68.0) packages, enabling the enrichment of Gene Ontology (GO) terms and the integration of InterPro annotations (Morgan et al., 2005; Pagès H et al., 2025). SNP positions were mapped to genes using GFF3 files, with the microseq (version 2.1.6; Snipen and Liland, 2016) package used to extract genomic features.

## Results

### Outlier Detection and Climatic Variable Selection: CANCOR vs GF

#### Comparison of the SNP detected as outliers by GF and CANCOR

Out of the 189,968 SNPs analyzed, GF identified 2,113 as candidates for climate adaptation (hereafter “GF outliers”). The model R^2^ values for these loci ranged from 0.39 to 0.73 (mean R^2^ = 0.47). For CANCOR, we found that the distribution of p-values followed theoretical expectations (uniform with enrichment only at low values) solely when considering the first two canonical dimensions (K = 2). Consequently, we restricted our analysis K = 2. At a false discovery rate (FDR) of 0.1, CANCOR identified 653 outlier loci (hereafter “CANCOR outliers”; **Supplementary Table S5**). The majority of these loci (555) were identical to those reported in Blanco-Pastor et al., 2020, with only 98 SNPs differing from the previous study. This discrepancy is likely attributable to the reduced set of climatic and phenotypic variables used for detection in the current analysis. Among these 653 outliers, 429 were also detected by GF. This shared set represents 65.7% of the CANCOR outliers and 20.3% of the GF outliers. In total, the two methods combined identified 2,337 unique candidate SNPs.

#### Genomic distribution and gene ontology enrichment of detected outliers

The genomic distribution of the identified outliers largely reflected the composition of the initial genotyping array, which was enriched for genic regions. Of the 189,968 total SNPs analyzed, 61% were located within genes and 9% within 1,000 base pairs (bp) of a gene. Similarly, among the 2,113 GF outliers, (**Supplementary Table S6**), 1,361 (64%) were located within a gene, 161 (8%) were within 1,000 bp of a gene, and 591 (28%) were intergenic. The CANCOR outliers showed a nearly identical distribution, with 420 (64%) being located within a gene, 48 (7%) within 1,000 bp of a gene, and 185 (28%) in intergenic regions. In contrast, the subset of 429 outlier SNPs detected by both methods showed slight enrichment for coding regions compared to the background, with 299 (70%) being located within a gene, 21 (5%) within 1,000 bp of a gene, and 109 (25%) in intergenic regions.

Gene ontology (GO) enrichment analysis revealed distinct biological signatures between GF and CANCOR. A total of 128 Gene Ontology terms were significantly enriched for SNPs detected exclusively by GF (**Supplementary Table S7**). These terms were primarily associated with growth and morphogenesis (e.g., cell development, unidimensional cell growth, pollen tube development) and lipid metabolism (e.g., icosanoid secretion, fatty acid transport). In contrast, CANCOR outliers were enriched for 57 Gene Ontology terms, while the SNPs detected by both methods showed enrichment for 38 Gene Ontology terms. Notably, cell development was the only biological process enriched across all three sets of outliers (GF-specific, CANCOR-specific, and common outliers). These findings indicate that while both methods capture signals related to core developmental processes, they otherwise highlight distinct aspects of underlying biological processes, with limited but meaningful overlap in core cellular functions.

#### Comparison of the Climatic Variables Selected by GF and CANCOR

GF_GF_ ranked only a few predictor variables with relatively high importance, revealing that only a small subset of variables contributed strongly to the model (**Figure 2**). The top-ranked predictors were dominated by measures of temperature (Bio4, Bio7, Bio9, Bio11), solar radiation (sis_au, sis_wi, sis_su), and evapotranspiration (bio.ad.25, bio.ad.26, bio.ad.27). We also found some concordance between GF and CANCOR. CANCOR, which only selects variables and does not provide an estimate of importance, preferentially included predictors ranked as highly important by GF_GF_, capturing all 5 of the top 5 variables and 6 of the top 10 (**Figure 2**, asterisks). Notably, the variables deemed least important by GF were almost universally not selected by CANCOR.

**Figure 2:**
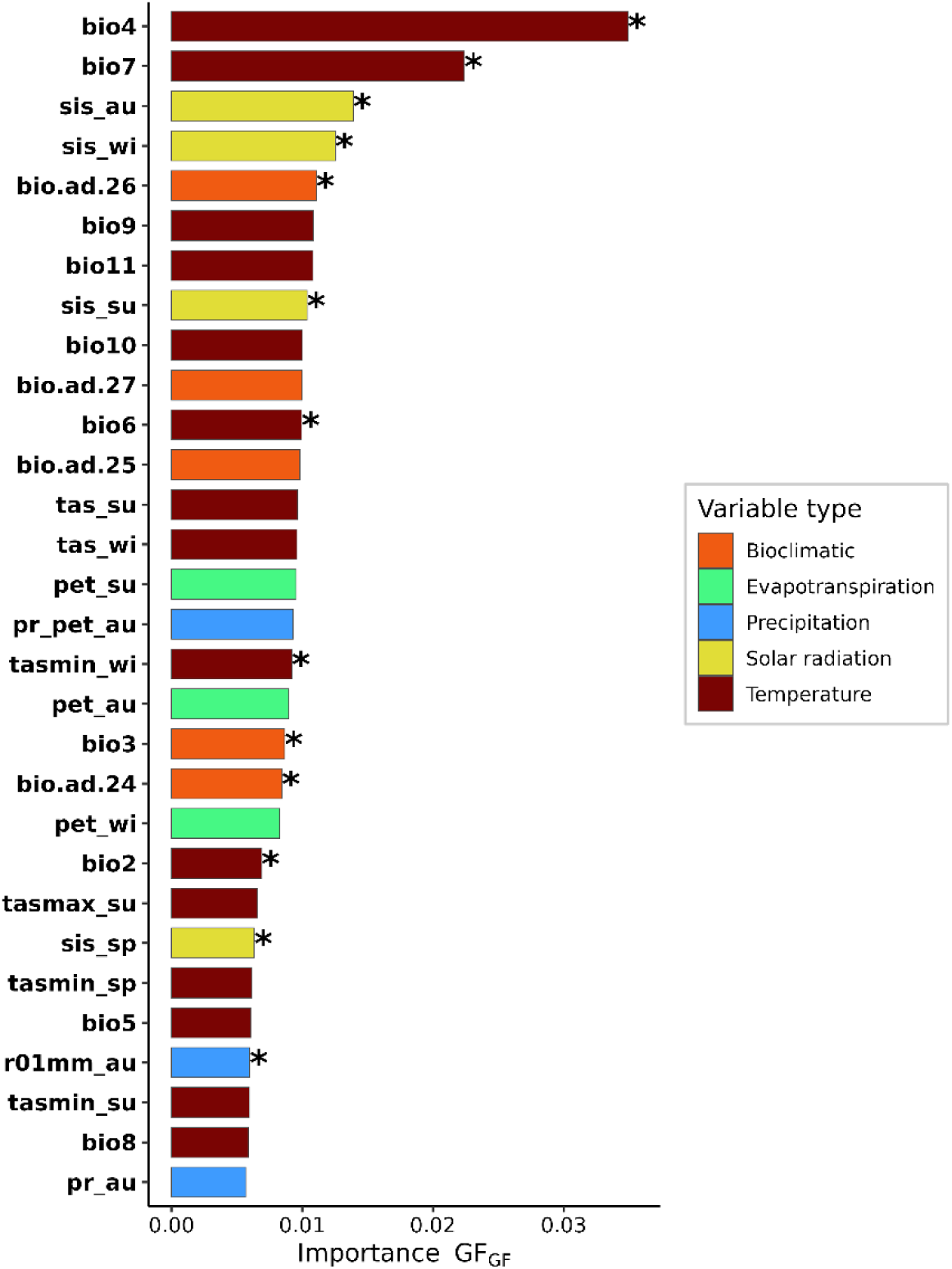
Ranked importance of the top 30 most important climatic variables from GF_GF_ fit using the Full Data set of outlier loci. Bars are colored by variable type: temperature (dark red), precipitation (blue), solar radiation (yellow), evapotranspiration (green), and bioclimatic indices (orange). Asterisks (*) indicate variables also selected by CANCOR..

### Impact of Loci Selection and Climatic Variables on Mapped Current and Future Genomic Patterns and Offset Projections

#### Genomic Composition Maps

We visualized the adaptive genetic composition for each time period and climate scenario (Current, Fut4.5, and Fut8.5) using RGB maps (**Figure 3**). Across GF models fit using different outlier sets (GF_GF_ or GF_CANCOR_), there is a shift in mapped genetic composition between the current and projected future conditions (Fut4.5 and Fut8.5). Genetic gradients are predicted to become less distinct in the future, with some regions of genetic similarity being replaced or homogenized. For example, northwestern regions with pink shading under current climate are progressively replaced in the future by purple shading, while orange areas in the south transition to green. In contrast, changes in eastern Europe appear less pronounced.

**Figure 3:**
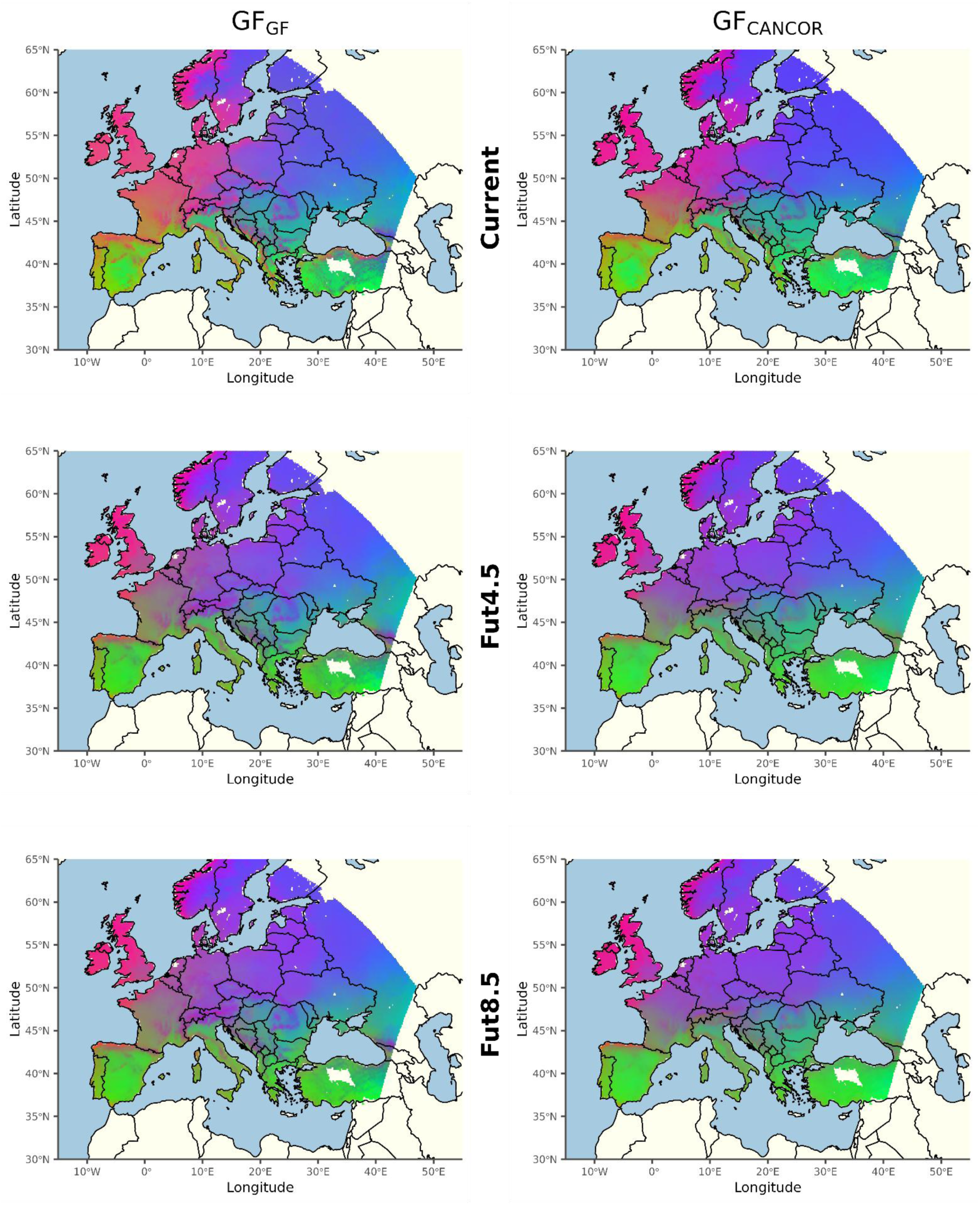
Predicted spatial variation in adaptive genetic composition from GF fit using (columns) different sets of outlier loci and applied to (rows) contemporary climate (1989-2010) and two future climate scenarios (Fut4.5 and Fut8.5). Shading represents gradients in genetic turnover derived from transformed climatic predictors that were subjected to Principal Components Analysis (PCA) to reduce the transformed climatic variables into three factors, which were then assigned to a RGB color palette. The resulting color similarity corresponds to the similarity of expected patterns of adaptive genetic composition. The PCA was centered but not scale transformed to preserve differences in the magnitude of genetic importance among the environmental variables.

While the absolute color scales of the two models are not strictly equivalent - due to differences in the number of input variables and outlier SNPs - the spatial structure and boundaries of regions predicted to have similar genetic composition (and therefore similar shading) are comparable. Visual inspection suggests that shifts in genetic composition occur at similar geographic locations in GF_GF_ and GF_CANCOR_. This spatial congruence is supported by the SIP statistics, which measure the covariance between maps. We observed high SIP values (and therefore high similarity) between GF_GF_ or GF_CANCOR_ across all climatic contexts: 0.945 for the current comparison, 0.936 for Fut4.5, and 0.926 for Fut8.5. These results confirm that despite methodological differences in outlier SNPs and variable selection, both approaches project similar landscapes of adaptive genetic variation.

#### Local Genomic Offset Projections

Regardless of the model (GF_GF_ or GF_CANCOR_), a pronounced region of high offset was evident spanning southwest to northeast from southern Spain to southern Sweden (**Figure 4**). In contrast, Eastern Europe exhibited relatively lower offset, while the United Kingdom and Ireland (with the exception of southern England) were projected to have the lowest offset. This pattern suggests that perennial ryegrass populations in the UK and Eastern Europe are at relatively lower risk under these climate change scenarios, requiring less genomic turnover to maintain current gene-environment relationships compared to populations in the high-offset band.

**Figure 4:**
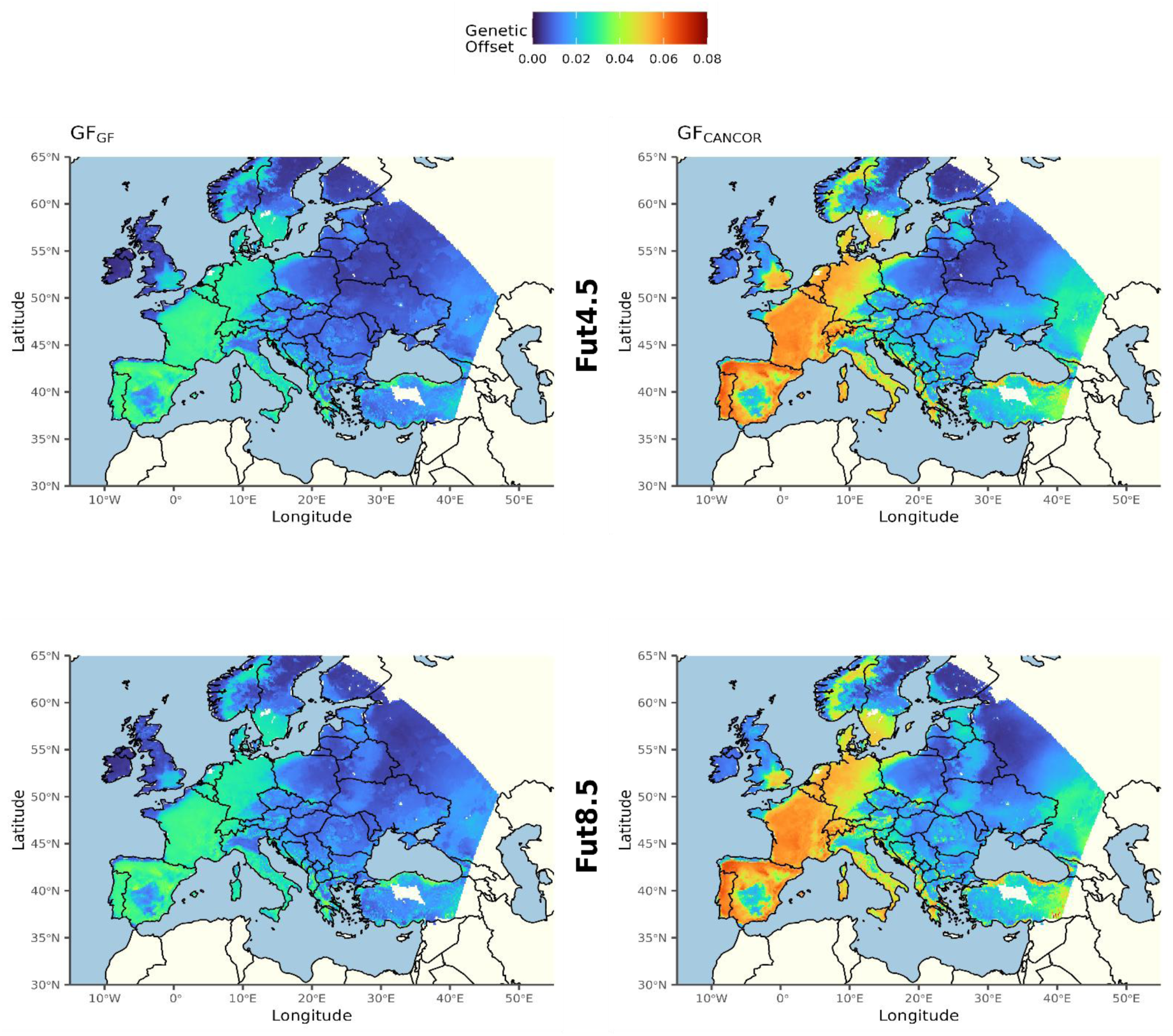
Local genomic offset from GF fit using (rows) different sets of outlier loci and applied to (columns) two future climate scenarios. Warmer colors represent relatively high offset and cooler colors represent relatively low offset.

While the magnitude of projected offsets differed between methods due to data input differences, the spatial patterns showed strong congruence (**Figure 4**). The SIP statistics indicate comparable levels of similarity both between offset projections from GF_GF_ and GF_CANCOR_ models within the same future climate scenario (SIP = 0.721 for Fut4.5; SIP = 0.754 for Fut8.5) and between the two climate scenarios within a single model (SIP = 0.722 for GF_GF_; SIP = 0.770 for GF_CANCOR_). This suggests that choice of GEA method and climate scenario contribute similarly to variation in offset spatial patterns. Notably, the lowest SIP values were observed when both the outlier set and the climate scenario differed (SIP = 0.591 for GF_GF_ Fut4.5 vs. GF_CANCOR_ Fut8.5; SIP = 0.562 for GF_CANCOR_ Fut4.5 vs. GF_GF_ Fut8.5), indicating that the two sources of variation compound rather than compensate each other. Overall, spatial congruence was slightly higher under the more severe climate scenario (Fut8.5), suggesting that stronger environmental shifts may produce similar spatial patterns of offset predictions across methods.

### Is the Relationship Between experienced Genomic Offset and Phenotype Affected by Loci Selection?

#### Concordance between GF_GF_ and GF_CANCOR_ in predicting phenotypic variation

To experimentally test how well genetic offset predicts performance of populations in new environments, we calculated the experienced genomic offset, defined as the offset between the climate of a population’s site of origin and the climate experienced during the common garden experiment. We assessed the ability of experienced offset to predict maladaptation by correlating it with a suite of phenotypic traits linked to species development (**Figure 5**, **Supplementary Table S8, Supplementary Figure S1 and S2**). Of the 105 phenotypic traits analyzed, 42 (40%) showed significant correlations with experienced genomic offset estimated by GF_GF_, while 53 (50%) were significantly correlated with experienced genomic offset from GF_CANCOR_ (p < 0.05 after Bonferroni correction). The mean absolute correlation was 0.18 for GF_GF_ and 0.21 for GF_CANCOR_, with correlations ranging from-0.40 (WSC_10_me17, water-soluble carbohydrates) to 0.61 (RAS_po15, vigour after sowing) for GF_GF_, and from-0.44 (resDST_lu17, description) to 0.50 (NSL_lu17, investment in sexual reproduction) for GF_CANCOR_. Notably, 36 traits (34%) were significant in both models, while 6 traits were uniquely significant for GF_GF_ and 17 for GF_CANCOR_. Out of 105 trait–site combinations, 59 (56%) showed a significant correlation with genomic offset in at least one model. The strongest correlations were observed for traits related to seedling vigour (RAS: r = 0.61), spring growth dynamics (NSL_lu17: r = 0.52; LSP_lu17: r = 0.47; resDST_lu17: r = 0.39–0.48), and biomass biochemistry (PRT_10_me17: r = 0.46; ADL_10_me17: r = 0.41). Overall, GF_GF_ and GF_CANCOR_ produced concordant results: traits that were significantly correlated with experienced offset in one model generally showed similar trends in the other (**Figure 5**).

**Figure 5:**
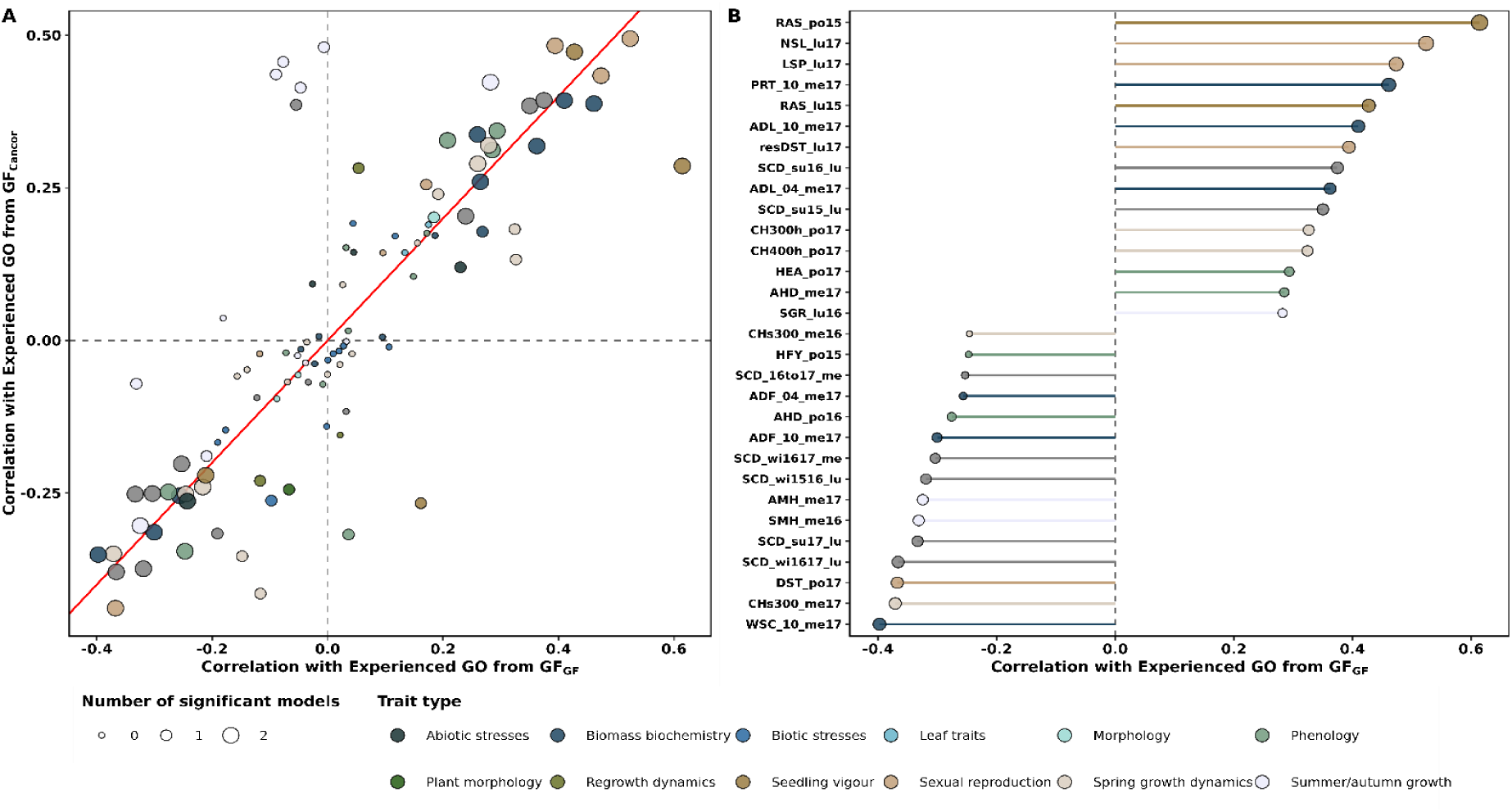
Correlation between phenotypic traits and the predicted genomic offset experienced in the common garden experiments for GF_GF_ and GF_CANCOR_. The scatter plot (A) compares the correlation between phenotypic traits and genomic offset as estimated by GF_GF_ (x-axis) versus GF_CANCOR_ (y-axis). The red line indicates the 1:1 relationship. Each point represents a phenotypic trait, with color indicating the type of trait (e.g., abiotic stresses, biochemistry of aerial biomass, morphology, etc.). Point size represents the number of methods (0, 1, or 2) that yielded a significant correlation between the predicted genomic offset (GF_GF_ or GF_CANCOR_) and the respective trait. The bar plot (B) represents the 30 strongest absolute correlation values between phenotypic traits and genomic offset as estimated by GF_GF_, with its length and direction indicating the strength and sign of the correlation (positive to the right, negative to the left).

Genotype–environment associations alone are sufficient to predict phenotypic maladaptation This concordance is notable given that GF_GF_ identifies candidate loci based solely on genotype–environment associations, without any phenotypic input, whereas GF_CANCOR_ selects outlier loci using both genomic and phenotypic data. Despite this fundamental methodological difference, the two approaches converged on the same major trait–offset associations, particularly for traits that can be interpreted as fitness proxies (e.g., seedling vigour, persistence, biomass biochemistry). The 17 traits uniquely significant for GF_CANCOR_ — concentrated in summer/autumn growth variables and spring growth dynamics at the PO site — tended to show weak or near-zero correlations with GF_GF_ but moderate to strong correlations with GF_CANCOR_, sometimes with reversed signs (**Figure 5**). This pattern is consistent with potential circularity in the GF_CANCOR_ approach, where phenotype-guided outlier selection may inflate correlations for traits closely related to the selection criteria.

#### Offset–trait relationships are context-dependent across trait categories

Among traits significant in both models, the strength and direction of correlations varied by trait category (**Figure 5**), reflecting the biological complexity of fitness in perennial species. For persistence traits, higher experienced offset was associated with reduced performance in most cases, e.g., winter soil coverage decline at ME (SCD_wi1617_me) was negatively correlated with experienced offset, indicating that populations with greater offset were less able to maintain cover during winter. Conversely, summer soil coverage at LU (SCD_su15_lu) showed a positive correlation, suggesting that populations originating from continental climates with high experienced offset outperformed locals under dry summer conditions. In the biomass biochemistry category, both models detected increased lignin content (ADL_10_me17) in high-offset populations, consistent with elevated stress responses, while water-soluble carbohydrate content (WSC_10_me17) was negatively correlated with experienced offset, suggesting reduced winter hardening capacity. These contrasting patterns illustrate that offset–trait relationships are context-dependent: depending on trait function and environmental conditions, significant correlations may reflect maladaptation, compensatory responses, or pre-adaptation to extreme events.

#### Climatic — not geographic — distance drives experienced genomic offset

PCA on the GF_GF_-transformed current climatic conditions in the study region, including at the common gardens and sites of origin (a representation of the adaptive genetic composition; **Figure 6**; see **supplementary Figure S3** for GF_CANCOR_ results), revealed that the climate conditions during the common garden experiments (represented by site-year combinations in **Figure 6**) occupied a relatively narrow portion of the adaptive genetic composition space. Only the climate of PO in 2016 diverged substantially, occupying central position in the predicted adaptive space. The variation in adaptive compositions was primarily driven by temperature seasonality (*bio4*), annual temperature range (*bio7*), and seasonal precipitation patterns (*pr_pet_au*), as indicated by the vector loadings on the PCA axes (**Figure 6**).

**Figure 6:**
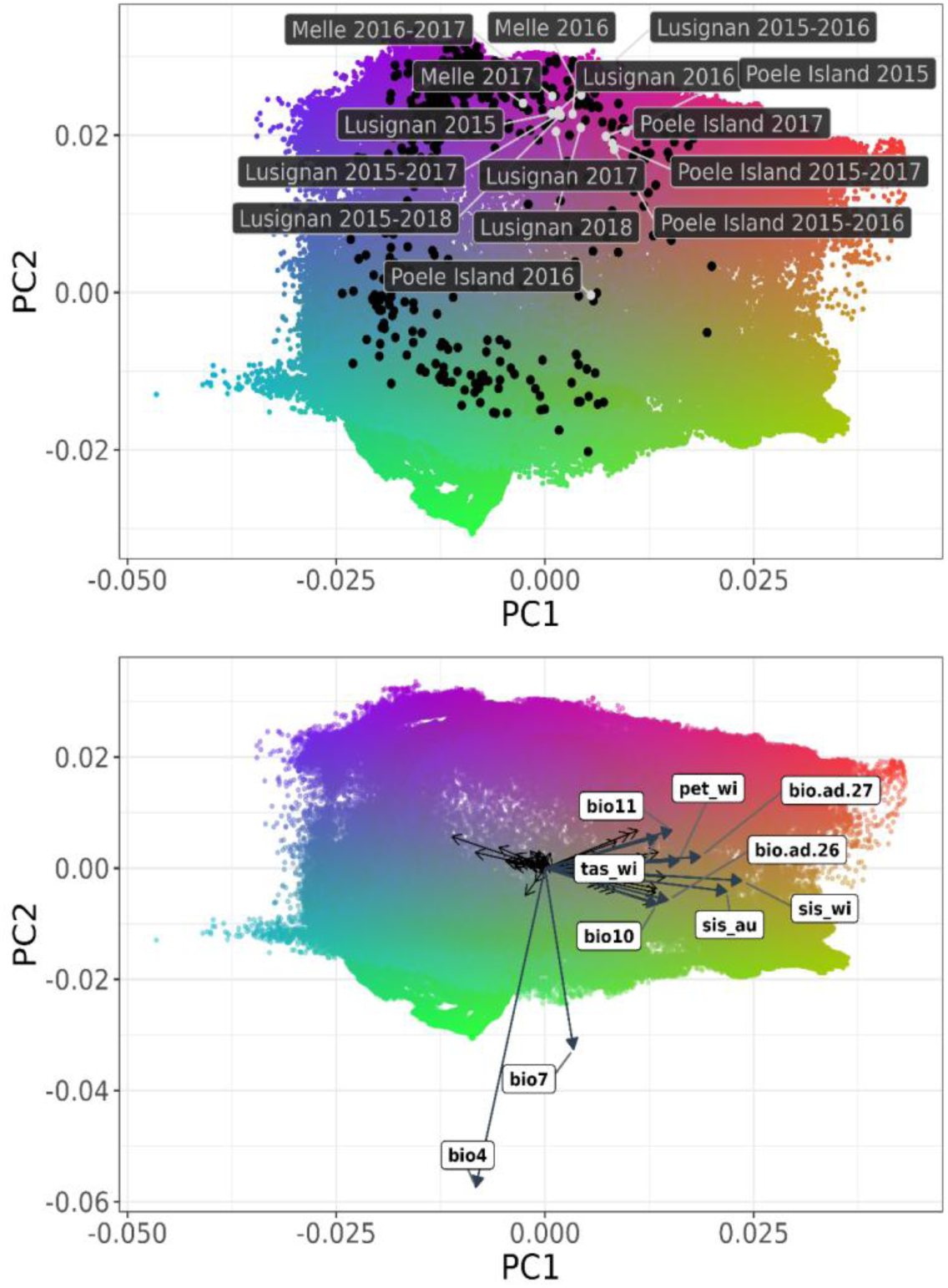
Biplots showing predicted variation in adaptive genomic composition from GF_GF_. Shading represents gradients in genetic turnover derived from transformed climatic predictors. These transformed variables were subjected to Principal Components Analysis (PCA) to reduce the data into three factors, which were then assigned to a RGB color palette. The PCA was centered but not scaled to preserve differences in the magnitude of genetic importance among the climatic variables. Color similarity corresponds to the similarity of expected patterns of genetic composition; locations with similar colors are expected to harbor populations with similar genetic composition. Each colored point is a climate grid cell within the study region. In the top panel, black points represent the 457 sampled populations, while white points indicate experimental gardens, with year labels denoting average climate in that location for those periods. The right panel displays the relative contribution of the top 10 climatic variables to the PCA axes. Primary variables are highlighted, while remaining variables are shown as black arrows. Arrow direction and length represent the correlation and influence of variables, such as seasonal temperature (e.g., *tas_su*, *tas_au*), solar radiation (e.g., *sis_wi*, *sis_au*), and bioclimatic indices (e.g., *bio4*, *bio7*, *bio11*), on PC1 and PC2.

To further disentangle the drivers of experienced offset, we compared it against both environmental distance (Mahalanobis) and geographic distance (**Figure 7**). We found a significant positive correlation between Mahalanobis distance and experienced genomic offset for both models (p <0.001), though the relationship was stronger for GF_CANCOR_ (r = 0.58) than for GF_GF_ (r = 0.27). The variance in this relationship was largely driven by the specific climatic conditions of each site-year combination (Figure 6, left panels). In contrast, there was no clear relationship between geographic distance and experienced offset (**Figure 7**, right panels). Populations geographically closest to a common garden did not necessarily experience the lowest genomic offset. This underscores that “local” populations in a geographic sense are not always “local” in a climatic sense and that temporal variation can represent an important driver of offset, both of which should be considered when interpreting experienced offset and changes in fitness related traits.

**Figure 7:**
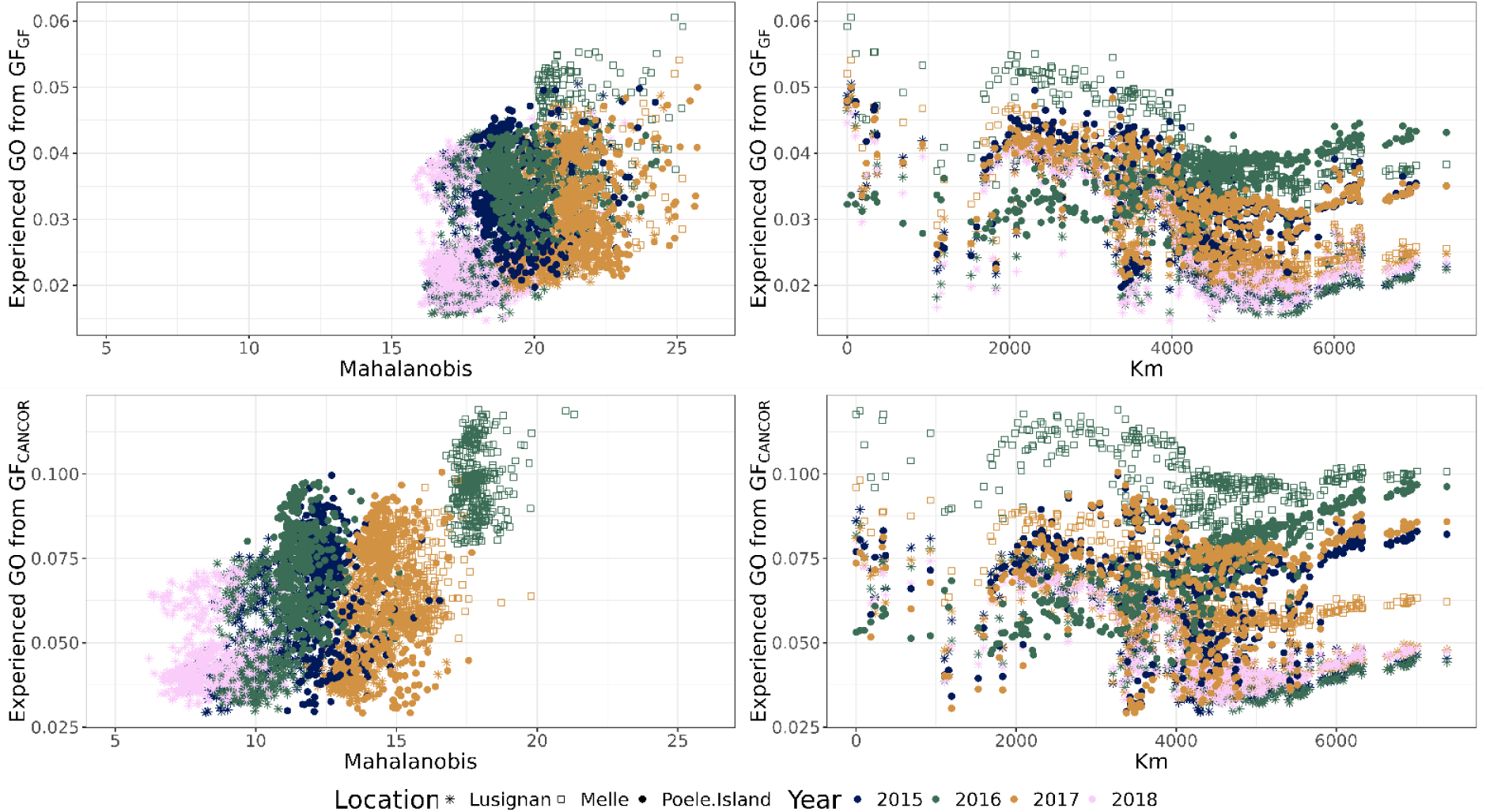
Scatter plots of Mahalanobis distance (left) and geographic distance (right) between population site-of-origin and common gardens versus experienced genomic offset predicted using different sets of outlier loci (rows, top=GF_CANCOR_, bottom=GF_GF_).Experimental years are indicated using a color ramp, while different symbols represent the different garden locations. Mahalanobis distances were estimated using the same climatic variables as those used to estimate experienced genomic offset.

### Model sensitivities to sampling

#### Outlier detection performance across sampling designs

To assess the sensitivity of the GEA methods to sample size and sampling bias, we compared the outlier sets detected in subsets of varying sizes and composition against those detected using the FDset.

SNP-level sensitivity and precision depended on both method and sampling design. At the SNP level, GF consistently outperformed CANCOR in both sensitivity and precision across most sampling configurations (**Figure 8A**). GF achieved sensitivity values typically ranging from 50% to over 90%, with precision between 40% and 75%, whereas CANCOR showed lower and more variable performance, with sensitivity generally between 10% and 75% and precision rarely exceeding 50%. Importantly, CANCOR also tended to have substantially higher false discovery rates (FDR), particularly for geographically unbalanced and small subsets, where FDR frequently exceeded 0.75. In contrast, GF maintained lower FDR values across most sampling designs. Core collections (CCLarge and CCSmall) performed well under both methods, though GF still yielded higher precision at comparable sensitivity levels. Among geographically unbalanced subsets, the Southern (UnBalS) and Eastern (UnBalE) designs achieved moderate sensitivity under both methods but with markedly reduced precision, especially for CANCOR. The Western (UnBalW) and Northern (UnBalN) subsets performed poorly regardless of the method, with both sensitivity and precision below 25–30%, confirming that excluding these regions leads to a substantial loss of adaptive signal.

**Figure 8:**
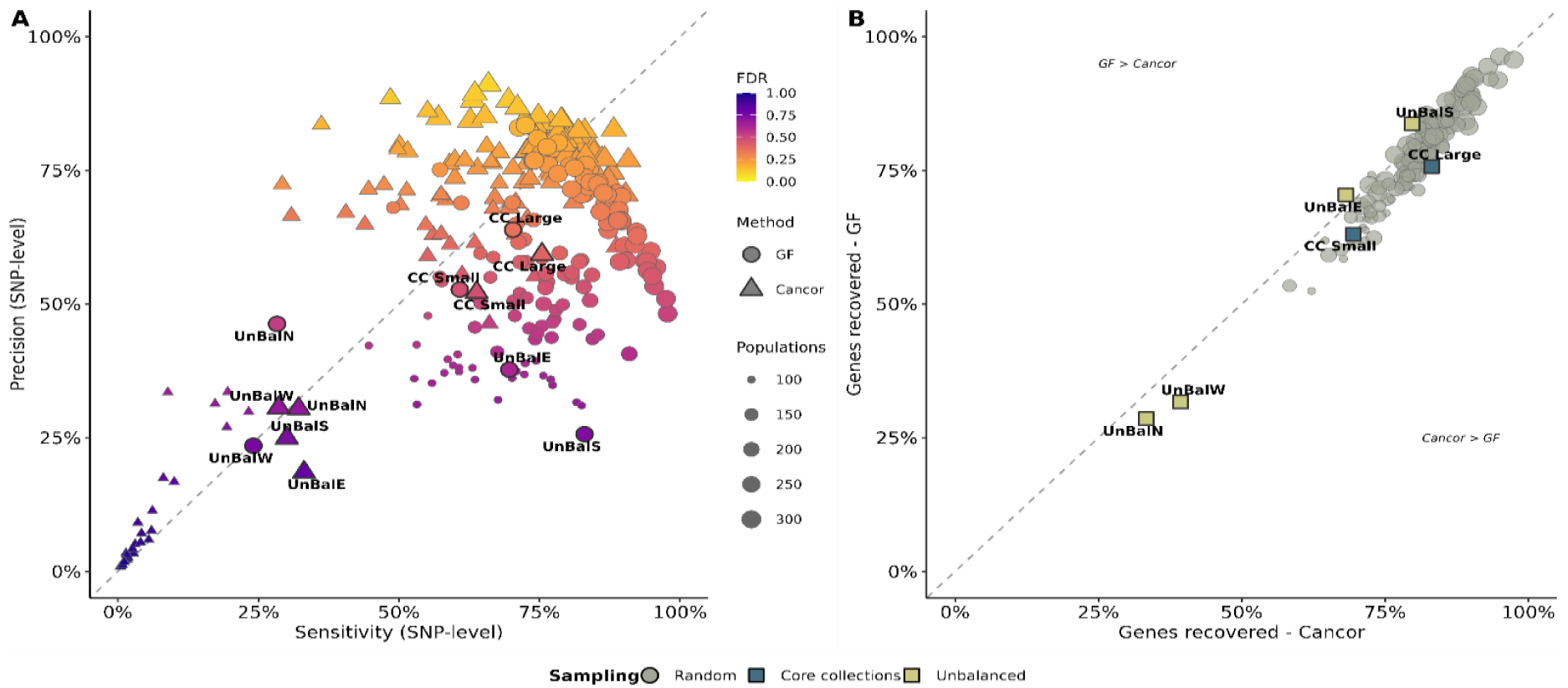
Robustness of candidate loci detection across sampling strategies and methods. (A) Precision-sensitivity trade-off at the SNP level for Gradient Forest (GF, circles) and Canonical Correlation Analysis (CANCOR, triangles). Each point represents an individual sampling replicate. Point size indicates the number of populations in the random subsample, while the color gradient indicates the False Discovery Rate (FDR); yellow represents low FDR (≤0.25) and purple represents high FDR (≥0.75). (B) Gene-level recovery and consistency between methods. Each point represents the proportion of genes identified in the full dataset that were recovered by a given subsample, plotted for GF (y-axis) versus CANCOR (x-axis). Colored squares indicate targeted sampling strategies: core collections (blue) and unbalanced geographic sampling (yellow).

Gene-level recovery is more robust but reveals method-specific differences. When the comparison was extended to the gene level — asking whether the same functional regions were flagged even if specific SNPs differed — recovery rates were substantially higher and more stable than at the SNP level (**Figure 8B**). For most sampling designs, GF recovered a higher proportion of the full-dataset genes than CANCOR, with most points falling above the 1:1 diagonal. Core collections and the UnBalS design recovered 60–85% of genes under both methods, with GF showing a slight advantage in most cases. The CC Large design yielded the most balanced performance, with both methods recovering approximately 75% of genes. The most striking discrepancy was observed for the UnBalW and UnBalN subsets, which recovered only ∼25–30% of genes under GF — comparable to or lower than CANCOR— confirming that these geographic regions help to discover unique adaptive information that cannot be compensated for if absent from the sampling. Overall, these results indicate that GF provides more reliable and less inflated outlier detection across a wider range of sampling conditions than CANCOR, and that gene-level analyses are substantially more robust to sampling variation than SNP-level comparisons.

#### Spatial congruence between genomic offset maps across climate scenarios

To assess how sampling strategy, model choice, and climate scenario affect the spatial projections of genomic offset, we compared the maps generated from each population subset against the maps generated from the FDset using SIP statistics. An ANOVA confirmed that among all tested factors, model (GF_GF_ and GF_CANCOR_), number of populations, and sampling strategy, only climate scenario (Fut4.5 and Fut8.5) did not significantly influence the similarity of the genomic offset maps (p < 0.05).

The spatial similarity (SIP) between genomic offset maps computed under the two future climate scenarios (Fut4.5 and Fut8.5) differed markedly between the two GF-based methods (**Figure 9**). Overall, SIP values were higher for GF_GF_ than for GF_Cancor_, as most points fell above the 1:1 diagonal line in both panels, indicating that genomic offset maps derived from GF_GF_ were more consistent across climate scenarios than those derived from GF_Cancor_.

**Figure 9:**
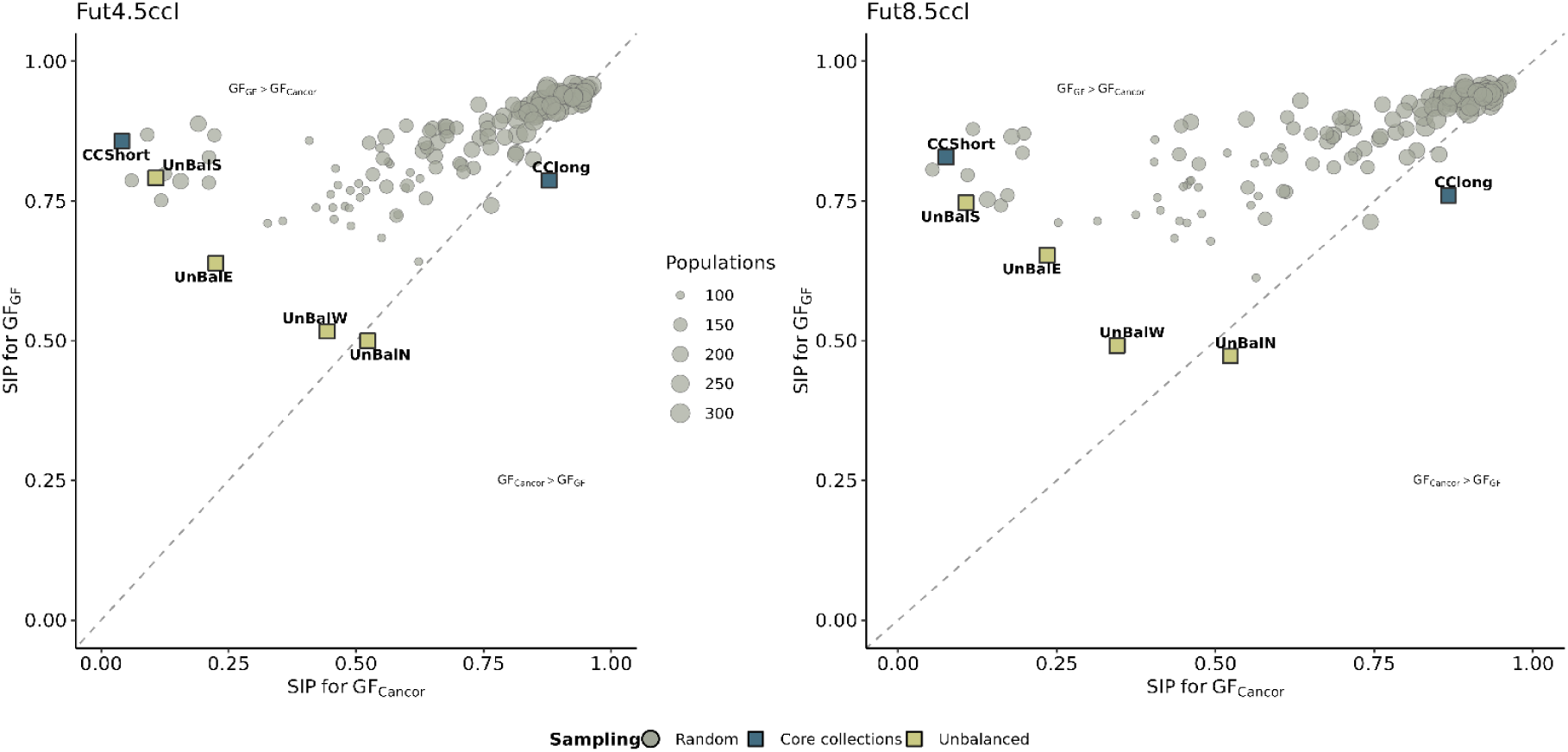
Sensitivity of genomic offset projections to population sampling design. The similarity index in pattern (SIP) measures the concordance between spatial projections derived from subsets of populations versus the full dataset, for RCP 4.5 (left) and RCP 8.5 (right) future climate scenarios. Grey circles represent random subsampling replicates, while colored squares represent different geographic sampling strategies: core collections (blue; CCLarge, CCSmall) and unbalanced sampling (yellow; UnBalE, UnBalN, UnBalS, UnBalW). Circle size is proportional to the number of populations in the subset. The dashed line indicates equal performance between GF_GF_ and GF_CANCOR_.

Under the moderate climate scenario (Fut4.5), GF_Cancor_ SIP values were highly variable, ranging from near 0 to ∼1.0, whereas GF_GF_ SIP values were generally high (>0.75) for most subset replicates. Under the more severe scenario (Fut8.5), SIP values for GF_Cancor_ increased, and the two methods showed greater agreement, with points clustering closer to the diagonal — though GF_GF_ still tended to yield more spatially consistent predictions.

Sampling design strongly influenced these patterns. Balanced or well-connected designs (CCLarge) yielded high SIP values for both methods under both scenarios, whereas unbalanced designs with geographic bias (UnBalW, UnBalN) produced lower SIP values, particularly for GF_GF_ (∼0.50). Short connectivity designs (CCSmall) and southern-biased unbalanced designs (UnBalS) showed a pronounced discrepancy between the two methods, with high SIP for GF_GF_ and low SIP for GF_Cancor_, especially for Fut4.5. Population sample size did not appear to have a strong systematic effect on spatial congruence.

These results suggest that GF_GF_ produces more robust and spatially stable genomic offset predictions across climate scenarios than CANCOR, but that the degree of concordance between methods is modulated by both the severity of projected climate change and the spatial configuration of the sampling design.

#### Sensitivity of the relation of experienced GO and phenotypes

Lastly, we evaluated the robustness of the relationship between experienced genomic offset and phenotype performance under different sampling schemes. We calculated Pearson correlation coefficients between experienced offset and phenotypic traits for each subsample and regressed them against the coefficients derived from the FDset. GF_GF_ showed higher stability than GF_CANCOR_, as indicated by the higher R^2^ between the subset-derived correlations and those from the FDset (**Figure 10**). This suggests that the biological signal detected by GF is less sensitive to sampling noise than CANCOR. In contrast, GF_CANCOR_ showed greater scatter around the regression line, indicating that associations between offset and phenotype derived from this method are more variable and dependent on the specific composition of the training set.

**Figure 10:**
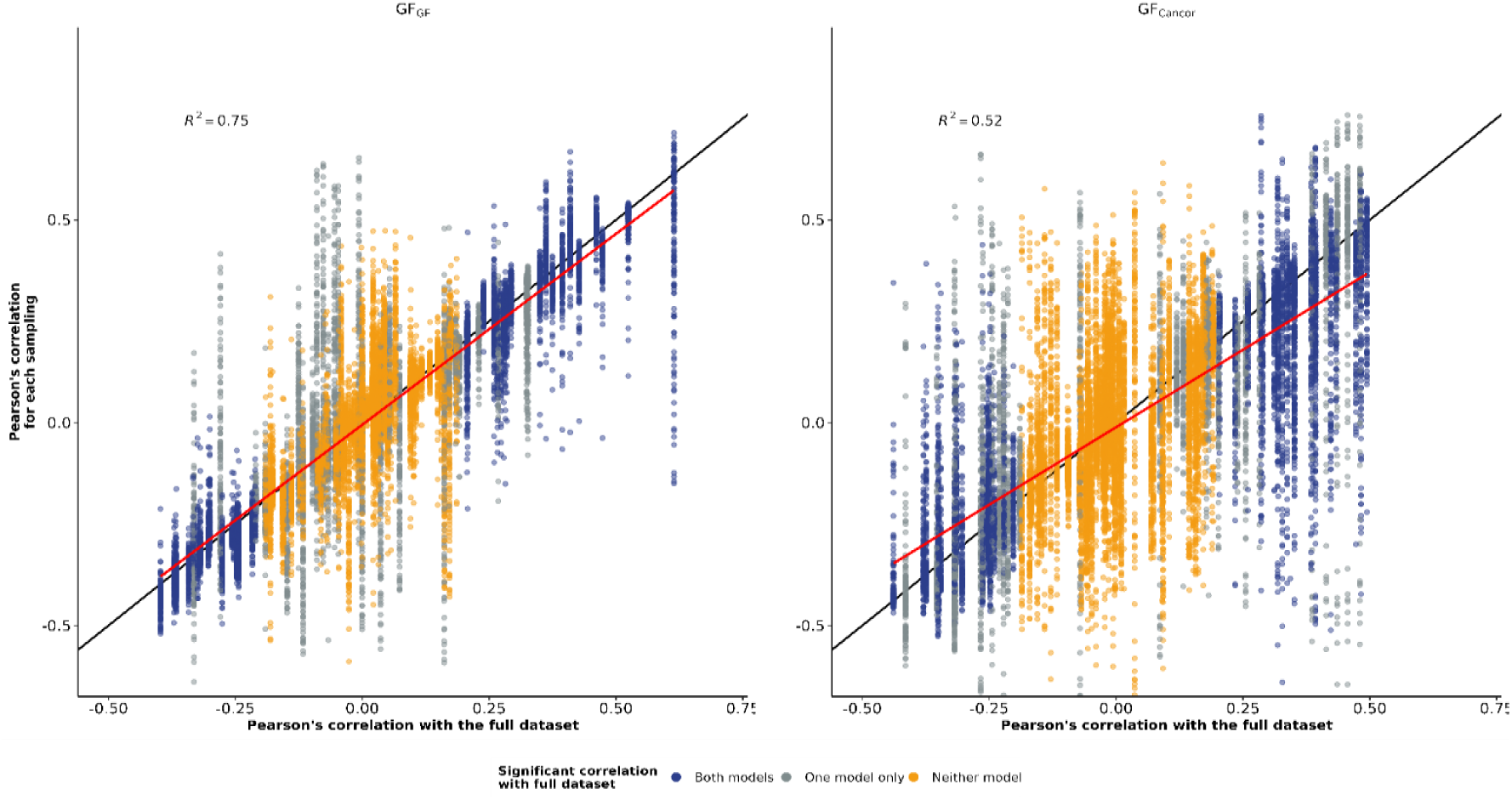
Sensitivity of the correlation between experienced genomic offset and phenotypic traits to population sampling design for (left) GF_GF_ and (right) GF_Cancor_. The y-axis represents Pearson’s correlation coefficient (*r*) for each population sampling replicate, plotted against the corresponding coefficient derived from the full dataset (x-axis). Data points are color-coded by the significance in the full dataset: orange denotes non-significance for both models, gray signifies significance for only one model, and blue represents significance for both models. The red line represents linear regression (with associated *R*^2^), while the black line represents the 1:1 identity slope.

## Discussion

This study provides a comprehensive evaluation of how methodological choices, namely outlier detection methods, predictor variable set, different sampling schemes, and modeling frameworks - influence genomic offset models. By comparing two fundamentally different methods - Gradient Forest (GF), a non-linear, machine learning method and Canonical Correlation Analysis (CANCOR), a linear constrained ordination approach - we show that while these two methods identify an overlapping set of candidate loci and tend to produce spatially congruent projections, they exhibit distinct sensitivities to input data and possess complementary strengths and distinct limitations.

### Complementarity of linear and nonlinear approaches

Despite fundamental differences in their analytical frameworks, GF and CANCOR identified meaningful overlap of 429 outlier SNPs among the 2,337 detected in total. This degree of convergence is notable, as previous studies often report minimal overlap between genome scan methods (Archambeau et al., 2026; Fitzpatrick et al., 2021; Forester et al., 2018; Lazic et al., 2024). Obtaining similar results from these contrasting statistical approaches suggests this shared set of loci likely represents a robust core of adaptive variation detectable regardless of statistical assumptions. Gene ontology enrichment analysis further corroborated the biological relevance of these loci, showing significant enrichment for processes critical to adaptive responses to climate, including cell development and growth, particularly among GF-detected outliers. Nonetheless, the contrasting assumptions underlying GF and CANCOR contributed to differences in the number and type of outliers detected, as well as their associated climatic variables.

The methods also showed complementary strengths. GF, with its capacity to model interactions and thresholds, excelled at identifying loci associated with complex, non-linear responses as might be expected for loci involved in adaptations to extreme climatic conditions (Láruson et al., 2022). In contrast, CANCOR proved effective at detecting loci associated with gradual environmental gradients, such as precipitation or temperature clines (Blanco-Pastor et al., 2020). However, the set of climatic variables deemed important by both methods exhibited considerable overlap, indicating that they capture similar environmental drivers of adaptive genetic variation. This alignment suggests that while GF and CANCOR emphasize different aspects of the adaptive landscape, they ultimately converge on key environmental axes shaping local adaptation-– even if the functional shape of the fitted relationships differ. Consequently, combining these approaches could provide a more holistic view of climate adaptation than either method in isolation.

The relationship between GO and phenotypic traits proved to be highly trait-and context-dependent, reflecting the complexity of adaptive responses in perennial ryegrass. For instance, soil coverage traits, such as description (SCD_wi1617_me), exhibited negative correlations with GO despite apparent increases in coverage (**Supplementary material S2 and S3**), suggesting a capacity to maintain growth during winter when conditions allow, provided the genotypes are adapted to the environment. Biochemical traits, including lignin content (ADL_10_me17) and water-soluble carbohydrates (WSC_10_me17), further illustrated this complexity: while lignin content correlated positively with GO, supporting the hypothesis that high-GO populations experience increased stress; water-soluble carbohydrates (WSC_10_me17) showed a negative correlation, indicating a reduced capacity for winter hardening and potential fitness costs (Keep et al., 2021; Volaire et al., 2023).

Despite the spatial congruence between the two methods, a clear divergence emerged during experimental validation: 17 traits, primarily summer/autumn growth and spring growth dynamics at the PO site, were uniquely significant for GF_CANCOR_. These traits tended to show weak or near-zero correlations with GF_GF_ but moderate to strong correlations with GF_CANCOR_, sometimes with reversed signs. Several non-exclusive mechanisms may explain this pattern. First, CANCOR’s integration of phenotypic information during outlier detection may allow inferences of adaptive signals that are not captured by genotype–environment associations alone. Second, the concentration of these uniquely significant traits at the PO site, which experienced atypical climatic conditions during the experiment, suggests that year-and site-specific environmental anomalies were modeled differently by each method, likely due to differences in climatic variable selection or the non-linear nature of the adaptive gradient. Regardless of the underlying cause, the key finding is that GF_GF_, which relies exclusively on genotype–environment associations in isolation of phenotypic data successfully recovered all 36 trait–offset associations detected by both methods, including traits with the strongest fitness-related signals (seedling vigour, persistence, biomass biochemistry). This highlights the utility of GF in capturing meaningful adaptive signals with closest links to measurable fitness outcomes.

### Genomic offset projections and sources of uncertainty

A key finding of our study is the spatial concordance (SIP = 0.72–0.77) between projected geographic patterns of genomic offset produced by the two methods despite using different sets of loci (Rellstab and Keller, 2024). Our analyses revealed that the choice of outlier detection method and climate scenario each contributed to variation in offset spatial patterns. The comparable SIP statistic between methods under the same scenario or within the same model shows that no single source of uncertainty dominates. These two sources probably have a compoundable effect since the lowest SIP values were observed when both method and climate scenario differed. This additive pattern of uncertainty has practical implications: when projecting genomic offset for conservation or breeding purposes, both the choice of method and climate scenario should be treated as equally important sources of prediction variability.

To assess whether the predictive power of genomic offset merely reflects naive climatic distance, we compared experienced offset against both Mahalanobis environmental distance and Euclidean geographic distance. Our findings were consistent with previous studies demonstrating that genomic offset outperforms naive environmental distance metrics in predicting fitness-related traits in common gardens (Capblancq and Forester, 2021; Fitzpatrick et al., 2021; Láruson et al., 2022; Lind et al., 2024), presumably because GF appropriately weights and scales environmental gradients to reflect their relative genetic importance rather than treating all climatic axes equally. The weaker correlation between Mahalanobis distance and GF_GF_ offset compared to GF_CANCOR_ further supports this interpretation: GF_GF_ transforms the environmental space more extensively through its non-linear framework, producing offset values that diverge more substantially from a simple linear distance metric. In contrast, geographic distance showed no clear relationship with genomic offset, confirming that geographically proximate populations are not necessarily the best candidates for climate change mitigation, especially in regions where climatic gradients are shallow. Temporal climatic variation across experimental years also contributed substantially to variance in genomic offset, underscoring that both spatial and temporal components of environmental heterogeneity should be considered when interpreting offset–fitness relationships in common garden experiments. This might also suggest that anomalous weather events in a given year can influence the adaptive response of a population. Consequently, future genomic offset models may need to incorporate inter-annual climatic variability rather than relying solely on long-term averages. and that

### Influence of sampling schemes on genomic offset models

Given the size of our dataset, we were able to explore how population sampling design can affect genomic offset projections, and found that they can be sensitive to the number of populations that are used to fit models (Aguirre-Liguori et al., 2023). Our results show that CANCOR tends to be more data hungry and more sensitive to sampling bias than GF. CANCOR required relatively large sample sizes to stabilize the number of outliers detected and recover signals apparent in the FDset. Similarly, mapped patterns from CANCOR exhibited greater sensitivity to sampling schemes. In contrast, GF remained relatively robust even at small sample sizes. CANCOR, by comparison, showed lower and more variable performance, particularly for small or geographically biased subsets. This corroborates findings by Aguirre-Liguori et al., 2023, who demonstrated that GF can achieve stable offset estimates with as few as 15–20 populations, provided the environmental space is adequately covered. The sensitivity of outlier detection and GO estimation to sampling strategies emerged as a critical consideration in this study. At the SNP level, GF consistently outperformed CANCOR in both sensitivity and precision across most sampling configurations. Core collections performed well under both methods, though GF still yielded higher precision at comparable sensitivity levels, emphasizing the value of strategic sampling for capturing adaptive variation. Surprisingly, when the comparison was extended to the gene level, recovery rates were substantially higher and more stable, suggesting that while specific SNP calls may fluctuate, both methods consistently flag the same functional regions. This finding increases confidence in the biological relevance of detected signals, despite method-specific differences at the marker level. Furthermore, it suggests that future research could transition from single-SNP analyses to the use of short-distance haplotypes to better capture adaptive variation. This finding warrant confirmation in other species, given that perennial ryegrass exhibits rapid linkage disequilibrium (LD) decay. Comparisons of GO maps generated under different sampling regimes further revealed that random sampling produced projections most similar to the FDset, while unbalanced sampling introduced greater divergence. Taken together, these findings can help guide study design. When logistical or funding constraints limit the number of populations that can be sampled, GF may be a more robust choice for analysis. Furthermore, our results demonstrate that randomized geographic sampling outperformed targeted or geographically biased sampling designs, producing genomic offset projections that were most consistent with those obtained from the FDset. Collectively, our findings suggest that maximizing genetic and environmental representation should be prioritized over the total number of populations to adequately cover a species’ environmental range. This conclusion reinforces recommendations to prioritize spatial replication over individual re-sequencing depth to minimize uncertainty in genomic offset estimations.

However, we showed that the relationship between GO and phenotypic traits varied depending on the outlier detection method and sampling strategy, highlighting how methodological choice can alter interpretations of genetic maladaptation, especially regarding so-called “fitness” traits (Archambeau et al., 2026). For instance, correlations between GO and potential fitness proxies, such as seedling growth, were often stronger than those observed for adult growth traits, underscoring the importance of selecting phenotypic indicators sensitive to climate-driven stress and early life stages (Verrico et al., 2025). These findings align with previous studies demonstrating that GO can either constrain or enhance adaptive capacity, depending on the trait and environmental context (Capblancq et al., 2020b; Fitzpatrick and Keller, 2015).

#### Genomic offset projection implications

The identification of adaptive loci and GO projections holds important implications for conservation and breeding programs. In conservation, GO maps can inform assisted migration strategies by pinpointing regions where populations are most vulnerable to future climate scenarios (Lachmuth et al., 2023). For example, the southwest to northeast pattern of high GO observed from southern Spain to Sweden identifies populations that may require priority intervention, such as genetic rescue or assisted gene flow, to mitigate maladaptation risks. In breeding programs, GO projections enable the proactive incorporation of adaptive alleles from wild or landrace populations into elite cultivars, thereby enhancing their resilience to climate change (Hansen et al., 2025; Hanson et al., 2024). Integrating GO metrics into breeding pipelines (Beck et al., 2025) could accelerate the development of climate-resilient crops, and combining genomic offset with genomic prediction of adaptive traits may further enhance selection accuracy by jointly targeting both climate-adaptive alleles and phenotypic performance (Pégard et al., 2023b, 2023a). However, realizing this potential requires empirical validation across species and environments to confirm the predictive utility of these complementary approaches (Galaretto et al., 2026). Current methods for estimating individual-level GO remain underdeveloped, particularly in quantifying uncertainty, which limits their immediate operational application (Verrico et al., 2025).

### Perspectives

All statistical methods are subject to method-specific biases that may affect their predictive accuracy. GF’s reliance on non-linear relationships renders it susceptible to overfitting in regions with sparse environmental coverage or underrepresented climatic extremes (Lind et al., 2024). Furthermore, Lind and Lotterhos, (2025) demonstrated through extensive simulations that offset prediction accuracy degrades under increasingly novel climates, underscoring the need for caution when projecting offset to future conditions substantially different from the training environment. Conversely, while CANCOR is robust to multicollinearity, it may fail to detect adaptive signals characterized by non-linear responses or complex interactions between variables. Additionally, our study assumed that the selected environmental variables adequately captured the climatic niche of perennial ryegrass, yet consideration of important factors that are difficult to quantify, such as microclimatic conditions or biotic interactions (e.g., soil microbiota, competition), could further influence adaptive patterns.

To advance genomic forecasting, future research should prioritize temporal replication through multi-year phenotypic trials to improve the robustness of GO-phenotype correlations, particularly for traits with high interannual variability. Experimental validation, via reciprocal transplant experiments or controlled environment studies, could empirically test the adaptive significance of detected outliers and their associations with fitness-related traits. The development of hybrid models combining GF’s non-linear flexibility with CANCOR’s multivariate framework may offer a more balanced approach for capturing the full complexity of adaptive landscapes. Finally, translating GO insights into actionable conservation and breeding strategies will require collaboration with stakeholders to ensure effective implementation in real-world contexts (Verrico et al., 2025).

An important conceptual caveat concerns the interpretation of our common garden validation. Common garden experiments evaluate “local-foreign” fitness offsets — comparing populations of different origins within a shared test environment — whereas the intended application of genomic offset is to predict “home-away” responses, i.e., how a population will perform when its home environment changes (Lotterhos, 2024). These two quantities are related but not equivalent, and the success of genomic offset in predicting local-foreign patterns does not guarantee equal accuracy for home-away predictions under future climate change. Our study provides empirical support for the genomic offset framework at the local-foreign level, but extrapolation to substantially different future climates warrants caution.

## Conclusion

The goal of this study was to assess how methodological choices influence genomic offset models. Despite fundamental differences in their analytical frameworks, GF and CANCOR identified a large shared proportion of candidate loci and produced spatially congruent offset projections. Both methods projected a consistent diagonal band of high genomic offset from southern Spain to southern Sweden, with eastern Europe and the British Isles at comparatively lower risk. Critically, the choice of GEA method and climate scenario contributed similarly to variation in spatial patterns, and these two sources of uncertainty compounded rather than compensated each other. This additive pattern implies that both factors should be treated as equally important when projecting genomic offset for conservation or breeding purposes. We also found that both GF_GF_ and GF_CANCOR_ detected significant correlations between experienced genomic offset and phenotypic traits, and GF_GF_ was able to find the strongest fitness proxies among a large number of traits. Lastly, GF proved substantially more robust than CANCOR across population sampling designs, maintaining high sensitivity and precision with low false discovery rates even at reduced sample sizes. In contrast, CANCOR showed greater instability and inflated FDR, particularly for small or geographically biased subsets. Importantly, gene-level recovery was far more stable than SNP-level comparisons for both methods, indicating that the functional adaptive signal is resilient to analytical variation at the marker level. Random geographic sampling consistently outperformed targeted strategies in reproducing full-dataset projections, and GF_GF_ produced more spatially stable predictions across climate scenarios and sampling designs than GF_CANCOR_. These findings recommend GF as the more robust framework for applied contexts where prediction stability is paramount, and highlight that maximizing environmental coverage through spatial replication should take priority over sequencing depth.

Finally, taken together, our results demonstrate that: (1) the genomic offset framework captures meaningful adaptive signals in perennial ryegrass, (2) this signal is robust to substantial methodological variation, and (3) GF provides a conservative yet reliable baseline for predicting climate maladaptation — even without phenotypic calibration. Nonetheless, important caveats remain. For example, common garden validation provides a test of local-foreign fitness differences, a related but not identical quantity to the home-away offsets that genomic offset is intended to predict under future climate change (Lotterhos, 2024), and prediction accuracy may degrade under increasingly novel climates (Lind and Lotterhos, 2025). Translating these findings into operational conservation and breeding strategies will require further validation across a wider range of environmental transfers, multi-year replication, and explicit quantification of prediction uncertainty.

## Acknowledgements

We thank the curators from European genebanks and staff from agronomic research institutes who contributed to the *in situ* collections in 2015. We are grateful to the staff from IPK, ILVO, and INRAE who established the common garden experiments and recorded the phenotypic data. We thank Tom Ruttink for his assistance with the alignment of SNP positions to the chromosome-scale genome assembly v2.6.1 (Nagy et al., 2022). Climate data were processed by CERFACS (https://cerfacs.fr) from EURO4M-MESAN and EUMETSAT CM SAF grids for observed data (1989–2010). *Lolium perenne* is a species covered under the Multilateral System of the International Treaty on Plant Genetic Resources for Food and Agriculture. All genetic materials used in this study were made available to the authors following the signature of a Standard Material Transfer Agreement (SMTA) in compliance with the Nagoya Protocol. MCF acknowledges funding support from NSF grant PGR-1856450.

