## SupplementaryFigS3 for "How robust are genomic offset predictions to methodological choices? Insights from perennial ryegrass"

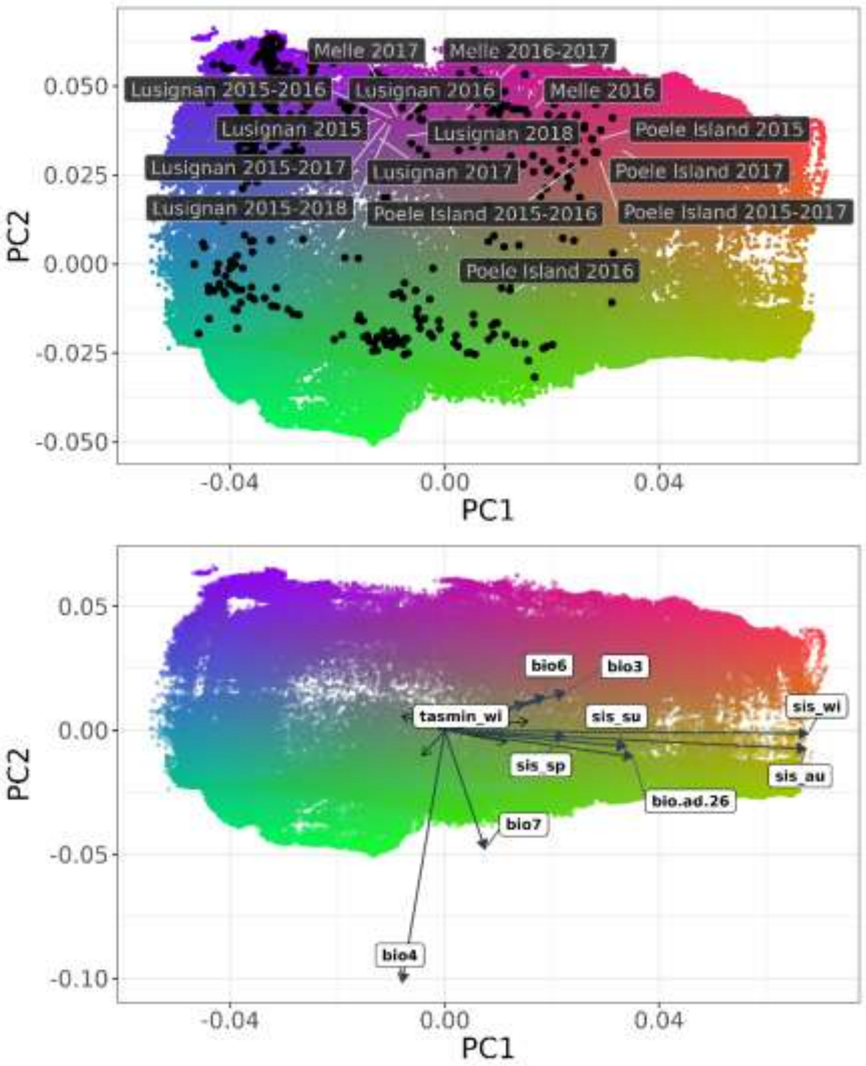


Figure S1 : Biplots showing predicted variation in adaptive genomic composition from $GF_{CANCOR}$. Shading represents gradients in genetic turnover derived from transformed climatic predictors. These transformed variables were subjected to Principal Components Analysis (PCA) to reduce the data into three factors, which were then assigned to a RGB color palette. The PCA was centered but not scaled to preserve differences in the magnitude of genetic importance among the climatic variables. Color similarity corresponds to the similarity of expected patterns of genetic composition; locations with similar colors are expected to harbor populations with similar genetic composition. Each colored point is a climate grid cell within the study region. In the top panel, black points represent the 457 sampled populations, while white points indicate experimental gardens, with year labels denoting average climate in that location for those periods. The right panel displays the relative contribution of the top 10 climatic variables to the PCA axes. Primary variables are highlighted, while remaining variables are shown as black arrows. Arrow direction and length represent the correlation and influence of variables, such as seasonal temperature (e.g., *tas_su*, *tas_au*), solar radiation (e.g., *sis_wi*, *sis_au*), and bioclimatic indices (e.g., *bio4*, *bio7*, *bio11*), on PC1 and PC2.
