## SupplementaryFigS1 for "How robust are genomic offset predictions to methodological choices? Insights from perennial ryegrass"

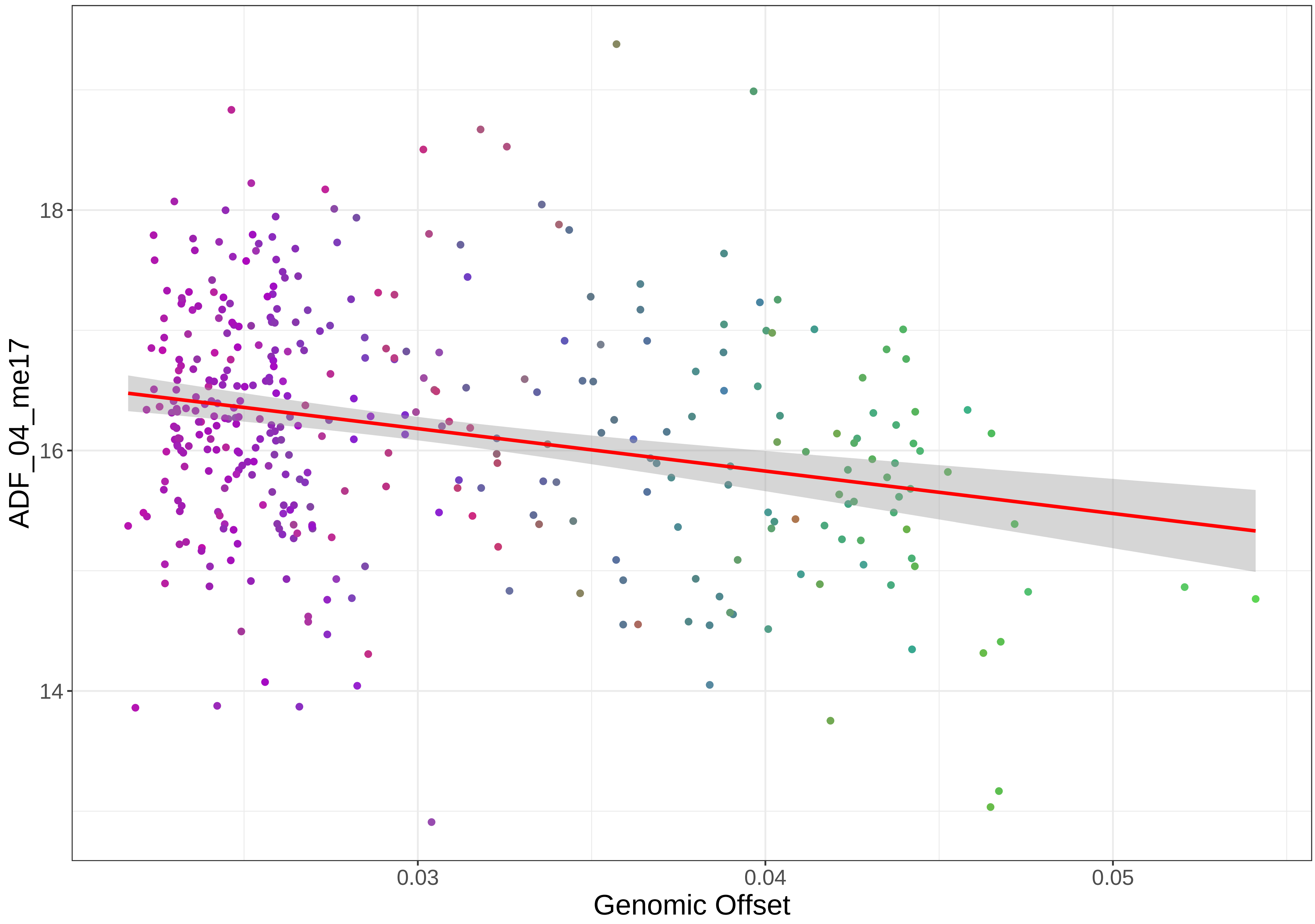

18
16
14
ADF_04_me17
0.03
0.04
0.05
Genomic Offset

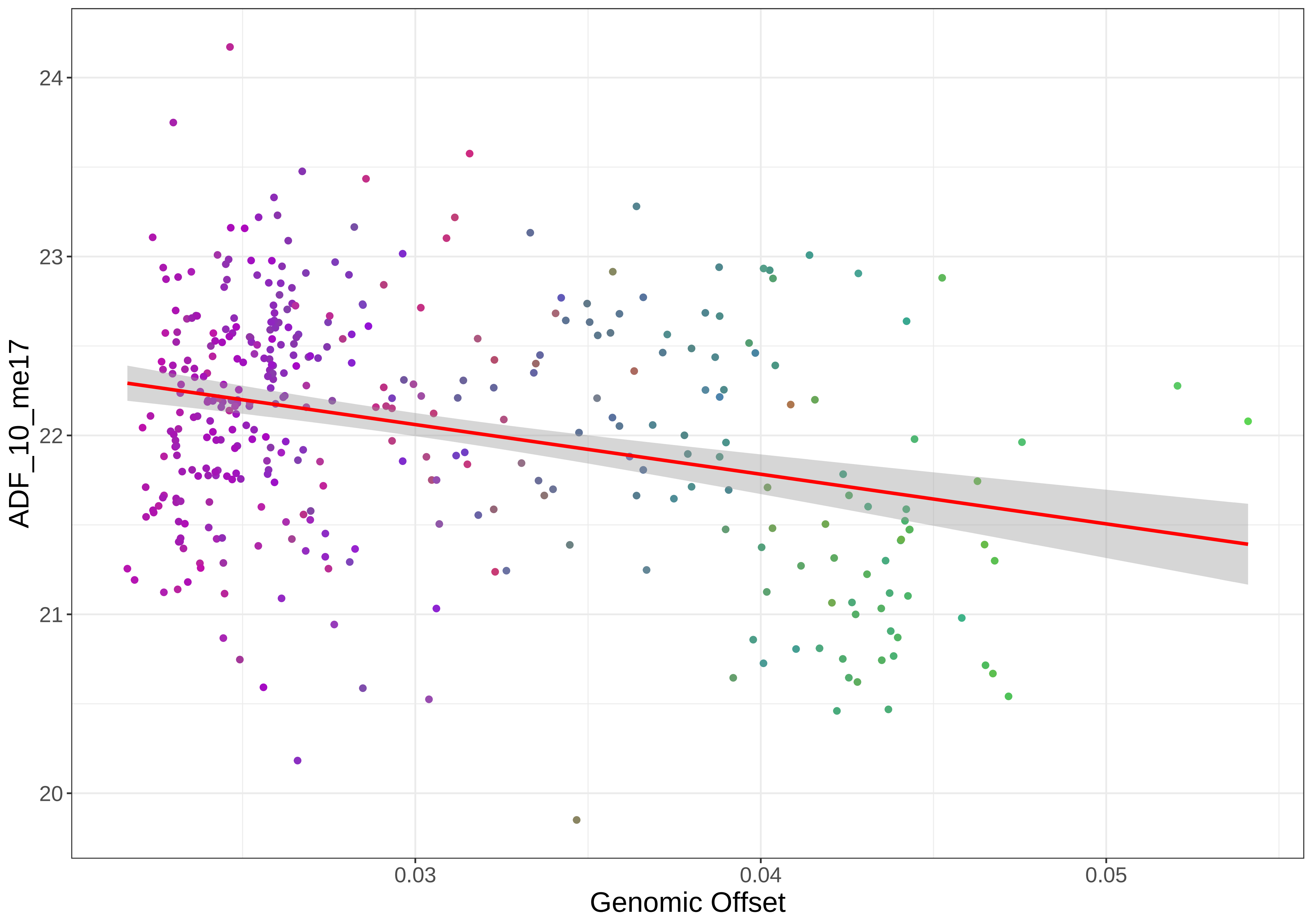

ADF_10_me17
24
23
22
21
20
0.03
0.04
0.05
Genomic Offset

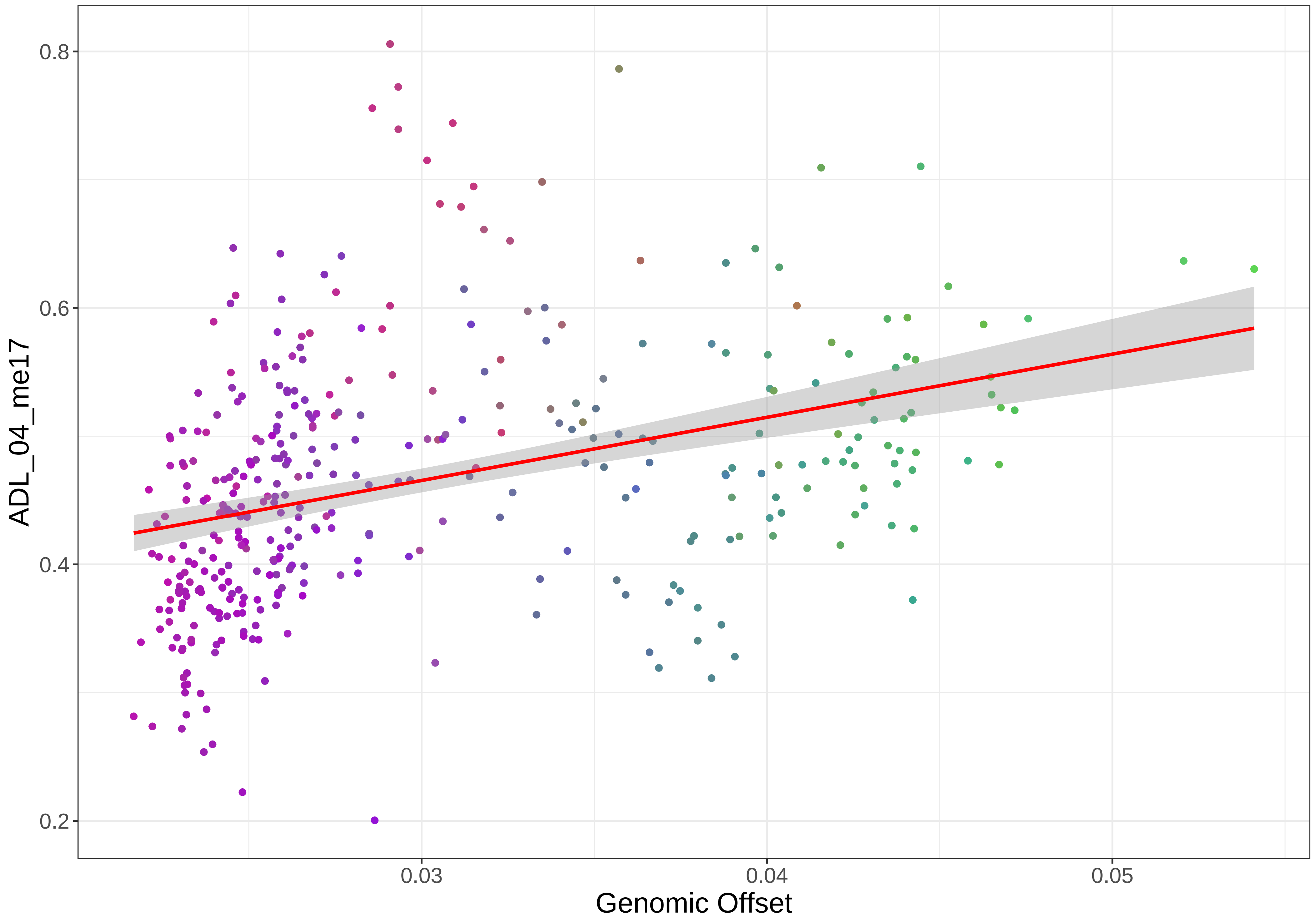
0.8
0.6
0.4
0.2
ADL_04_me17
0.03
0.04
0.05
Genomic Offset

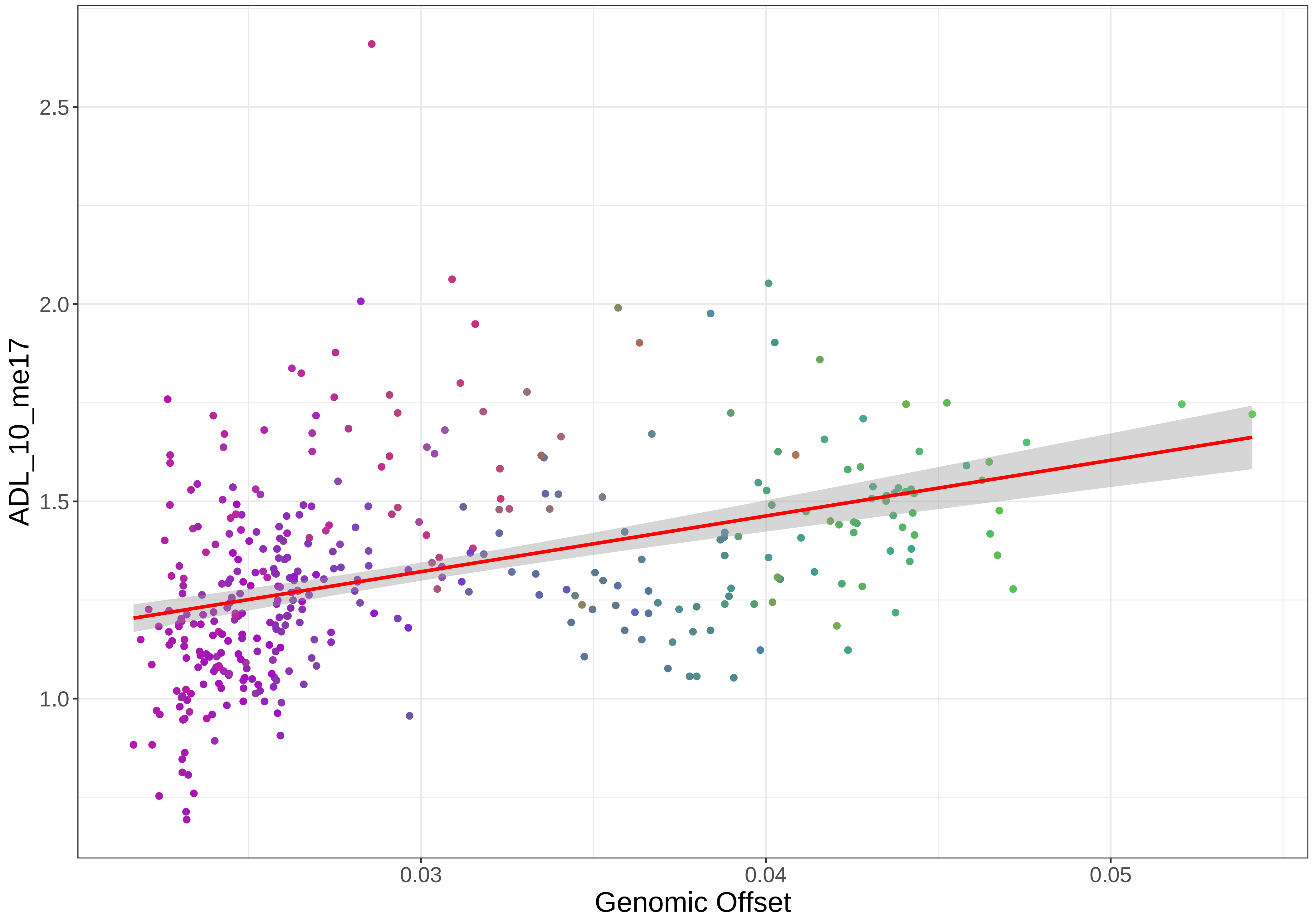
2.5
2.0
1.5
1.0
ADL_10_me17
0.03
0.04
0.05
Genomic Offset

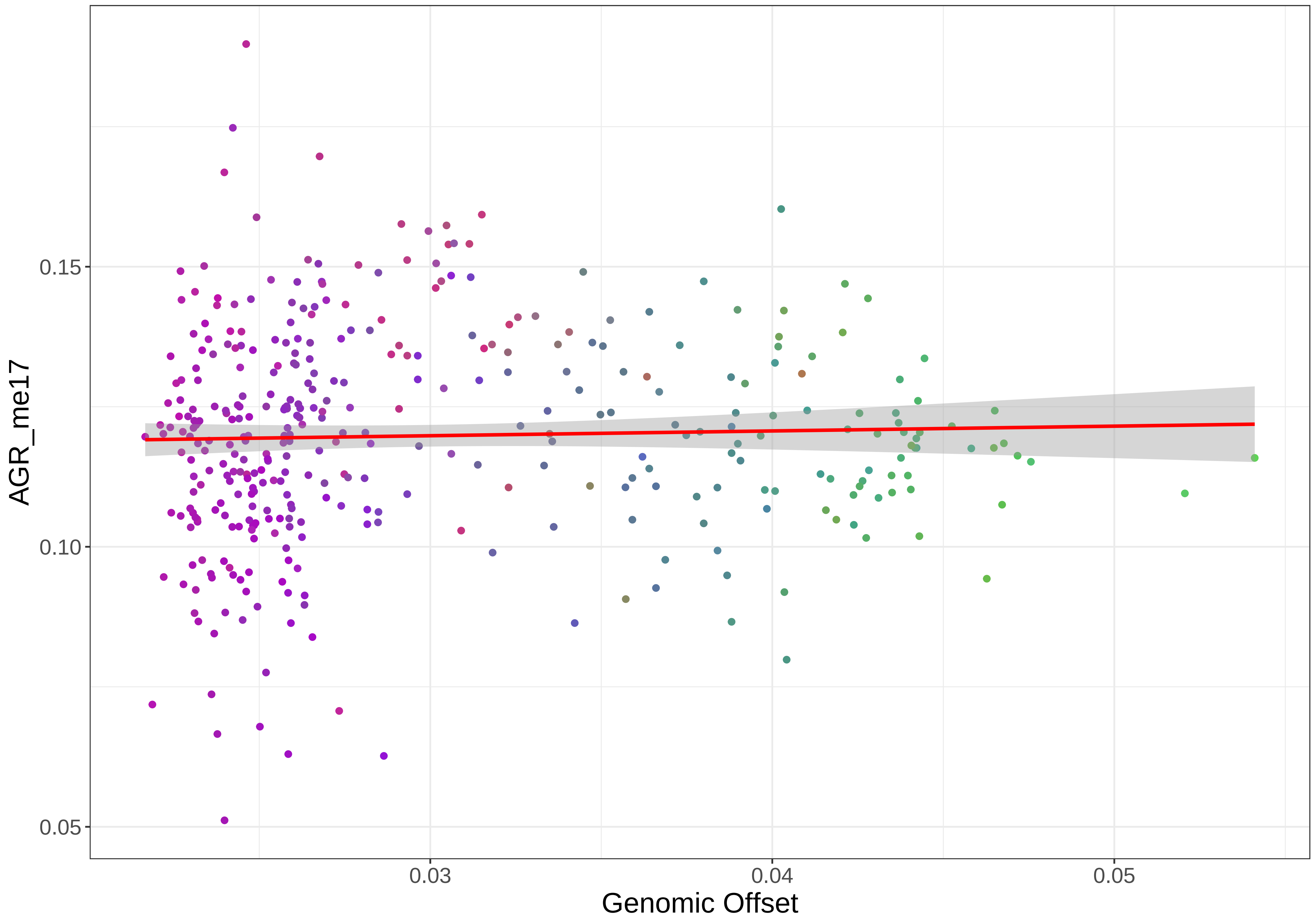

0.15
0.10
0.05
AGR_me17
0.03
0.04
0.05
Genomic Offset

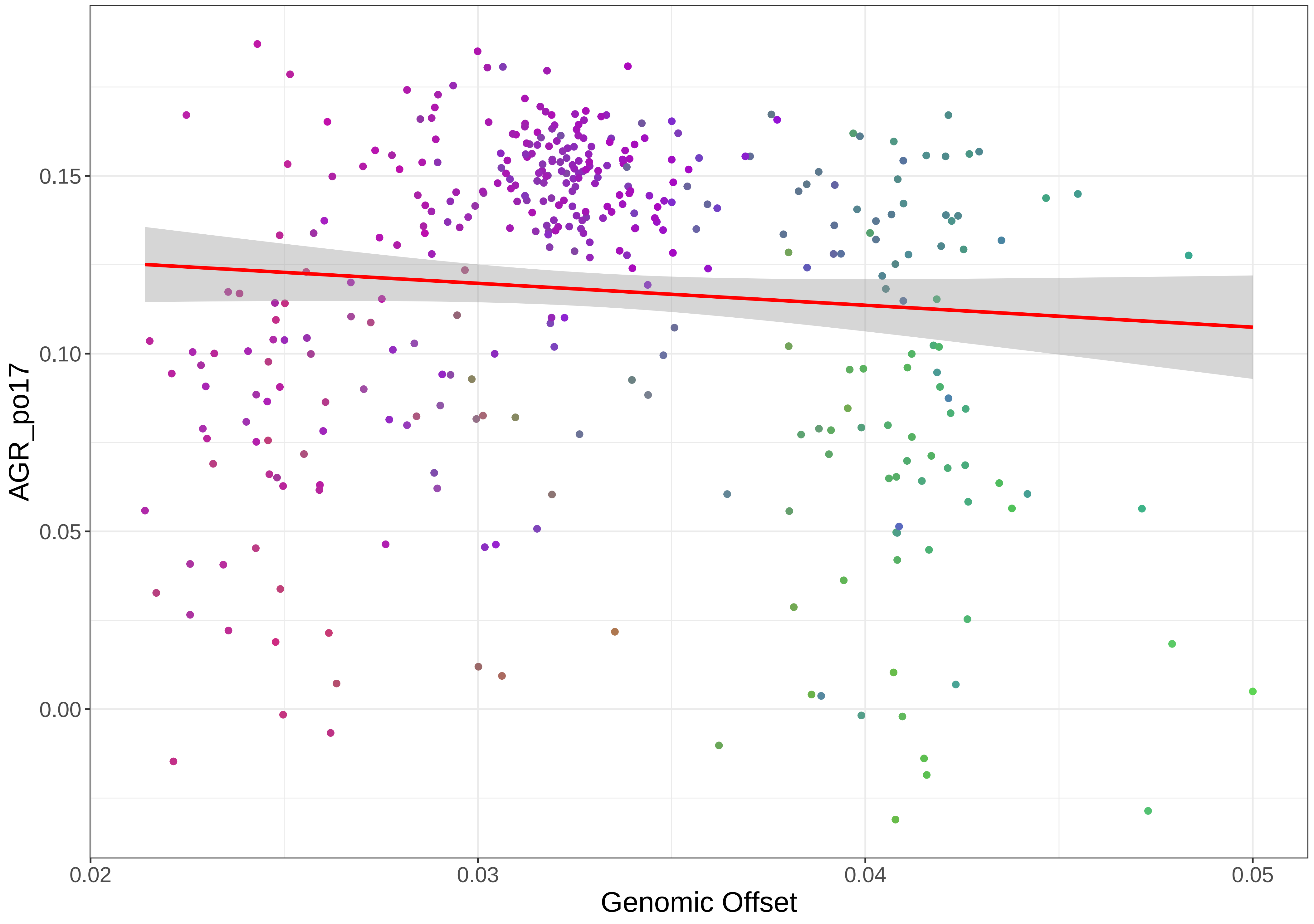
0.15
0.10
0.05
0.00
AGR_po17
0.02
0.03
0.04
0.05
Genomic Offset

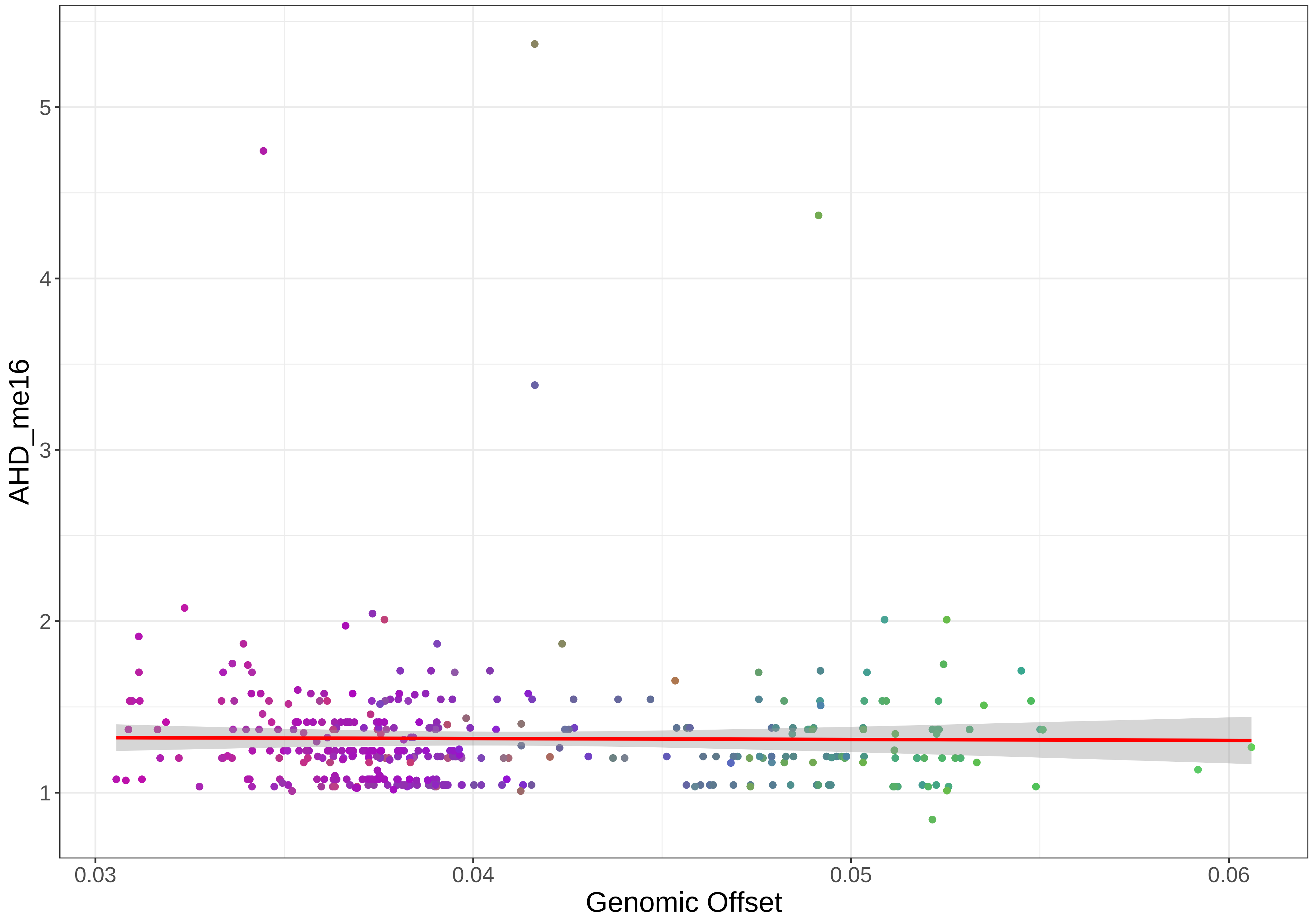

AHD_me16
5
4
3
2
1
0.03
0.04
0.05
0.06
Genomic Offset

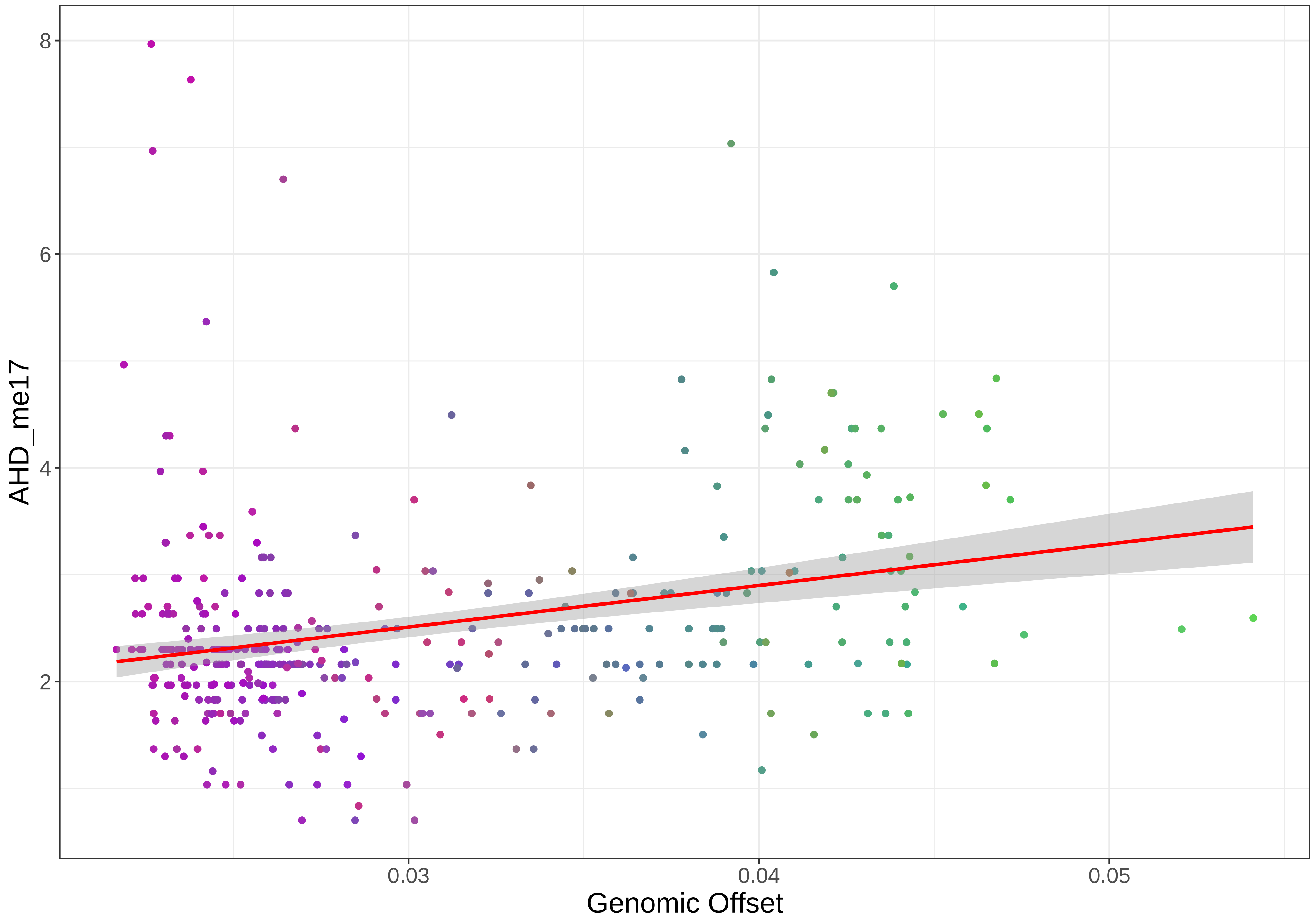

AHD_me17
8
6
4
2
0.03
0.04
0.05
Genomic Offset

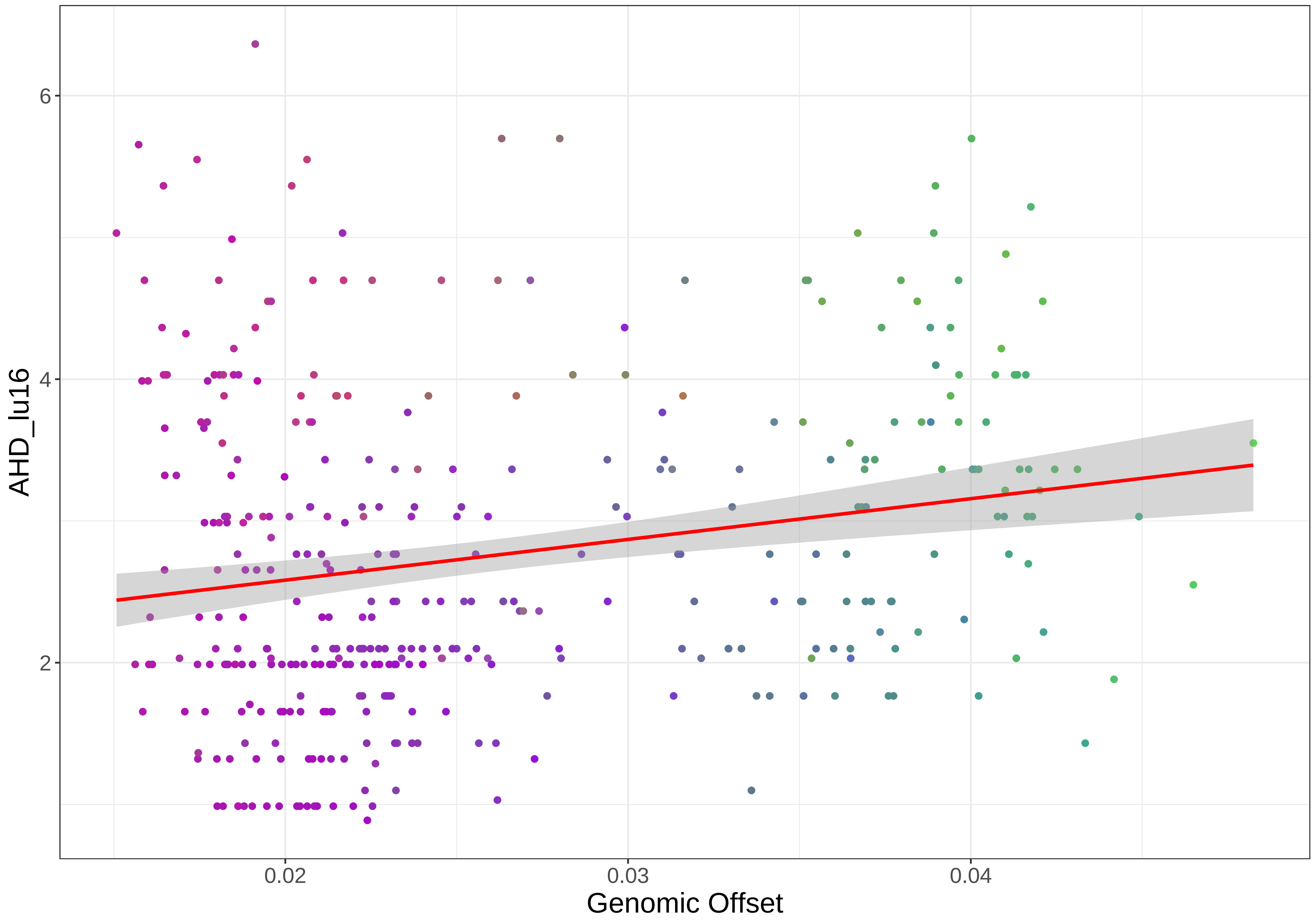

AHD_lu16
6
4
2
0.02
0.03
0.04
Genomic Offset

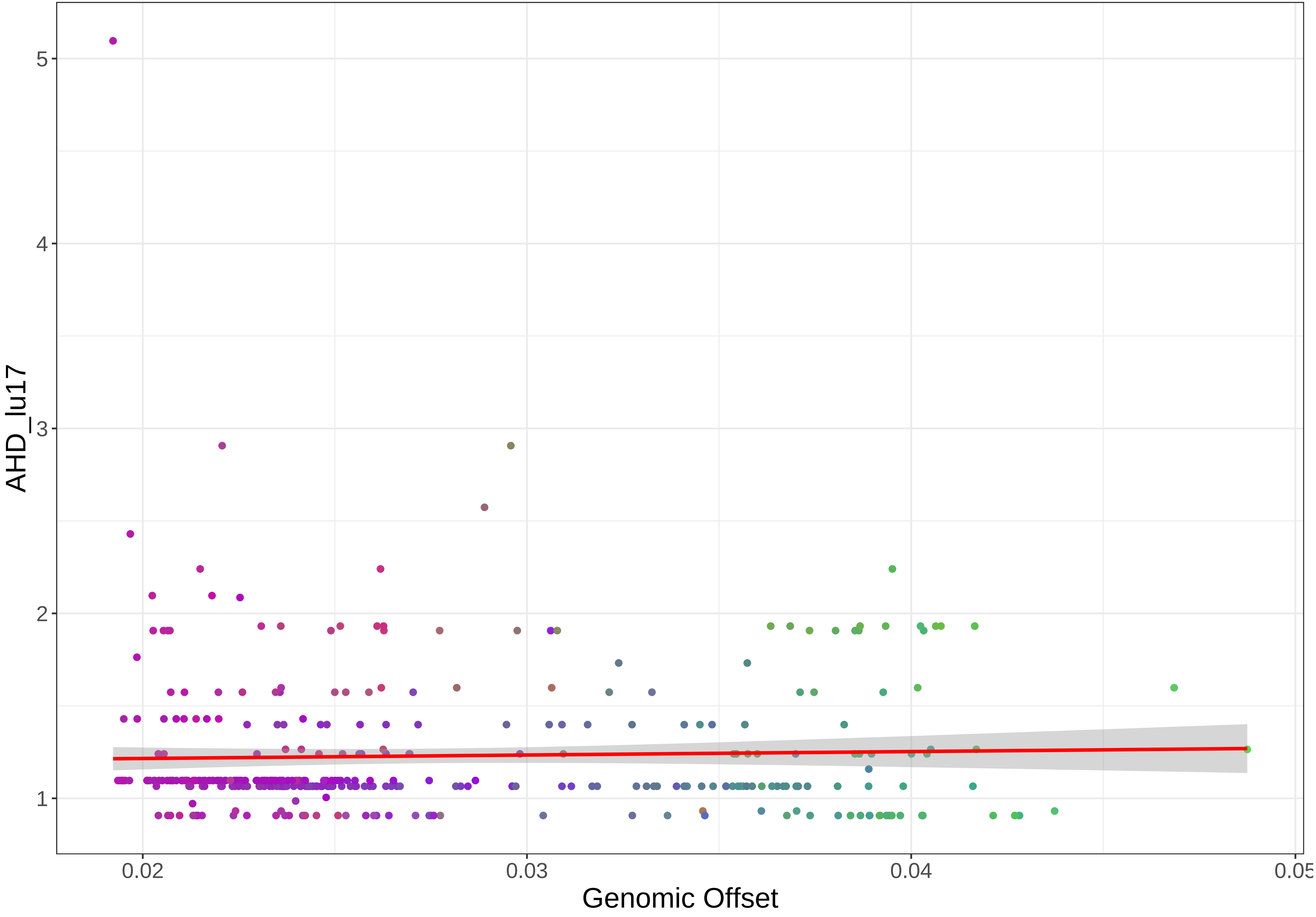

AHD_lu17
5
4
3
2
1
0.02
0.03
0.04
0.05
Genomic Offset

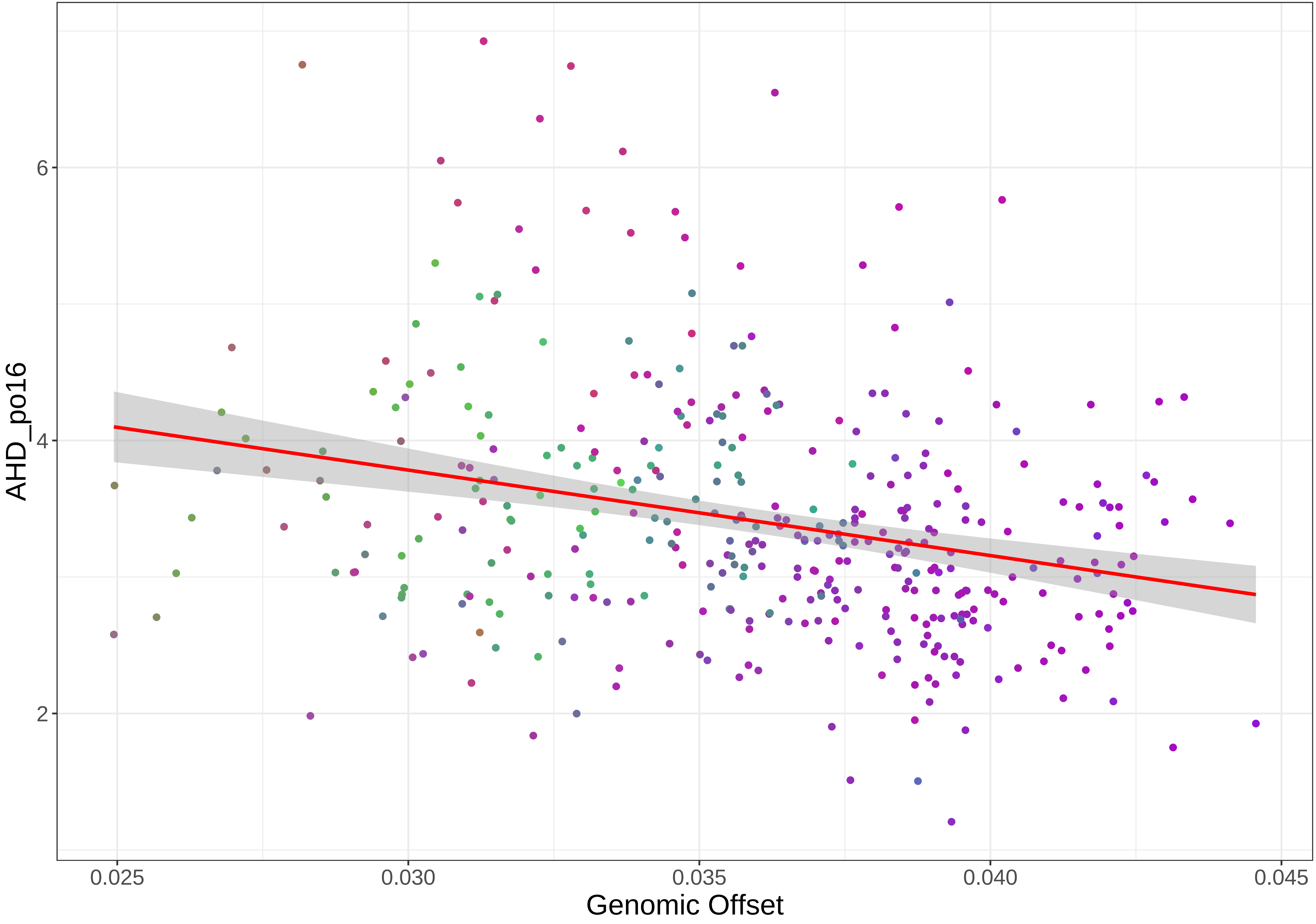

6
4
2
AHD_po16
0.025
0.030
0.035
0.040
0.045
Genomic Offset

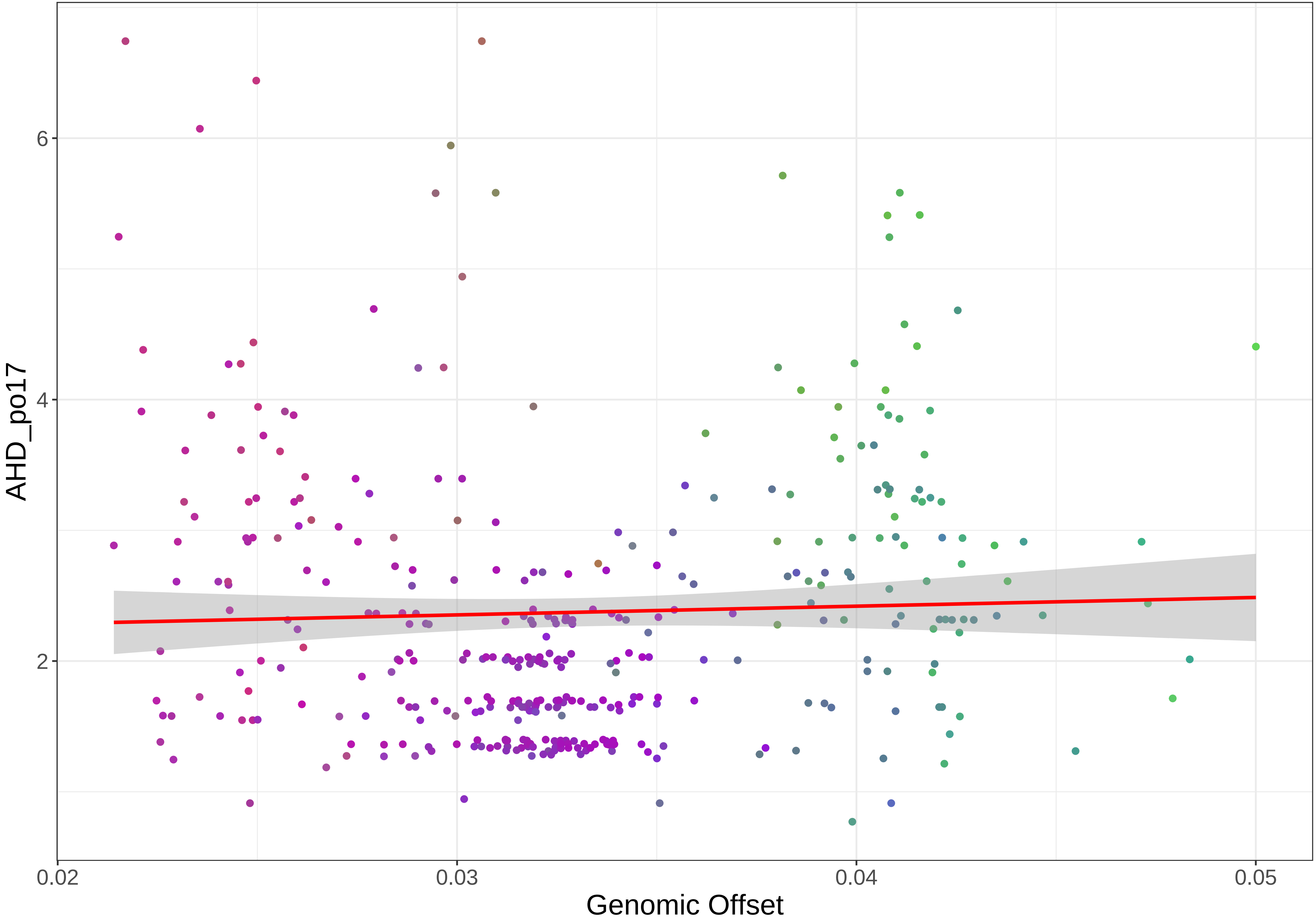
AHD_po17
6
4
2
0.02
0.03
0.04
0.05
Genomic Offset

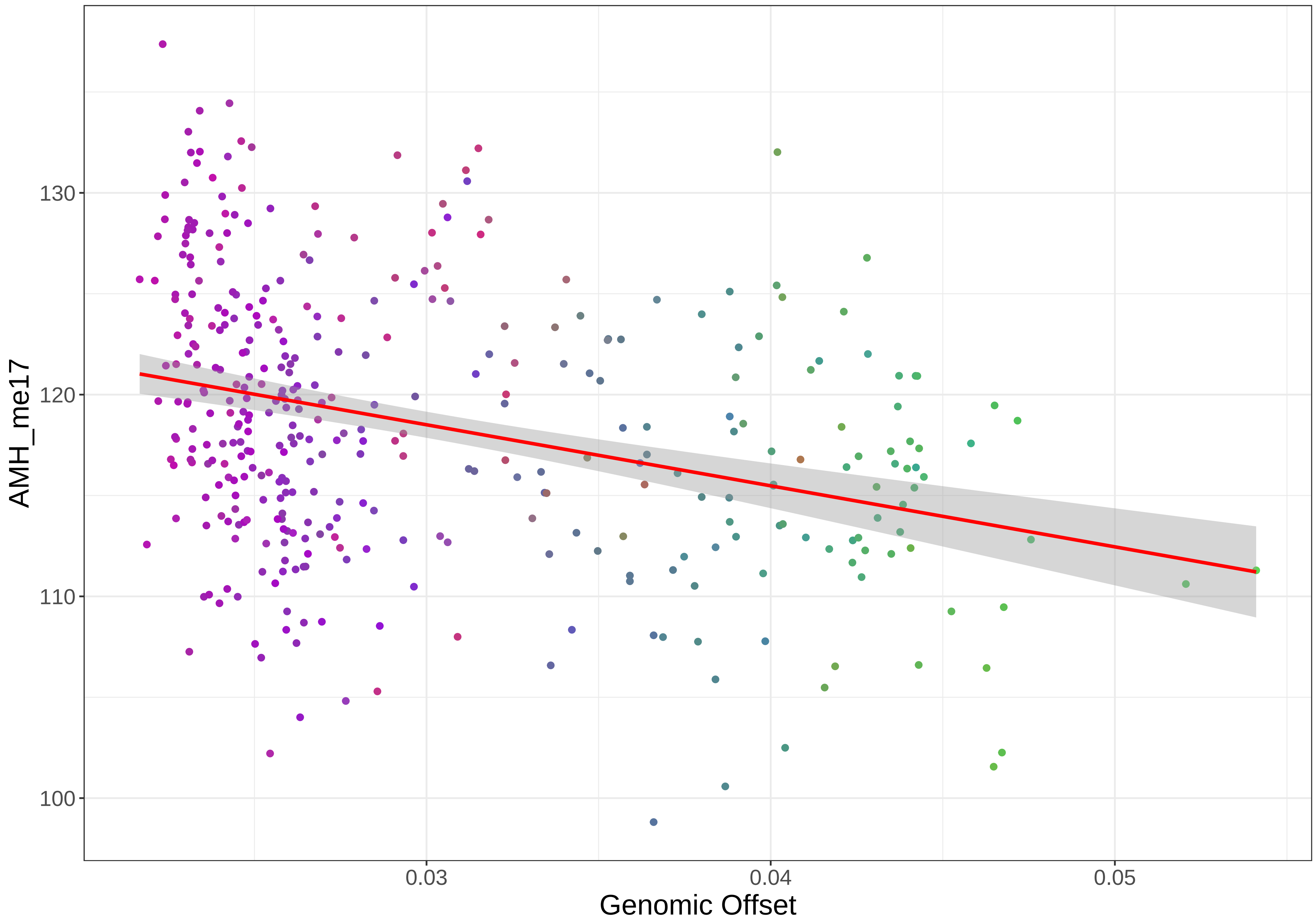

AMH_me17
130
120
110
100
0.03
0.04
0.05
Genomic Offset

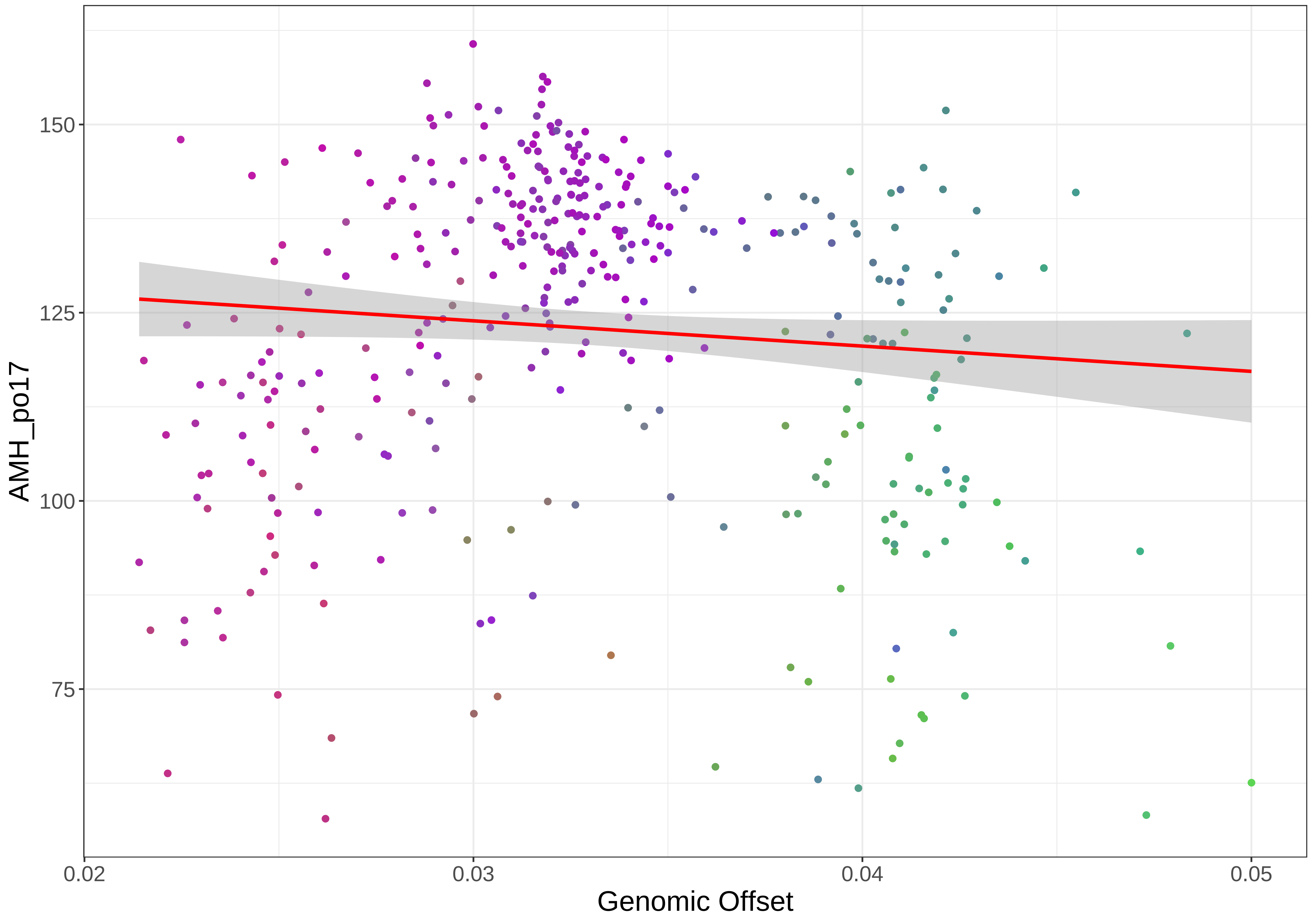

AMH_po17
150
125
100
75
0.02
0.03
0.04
0.05
Genomic Offset

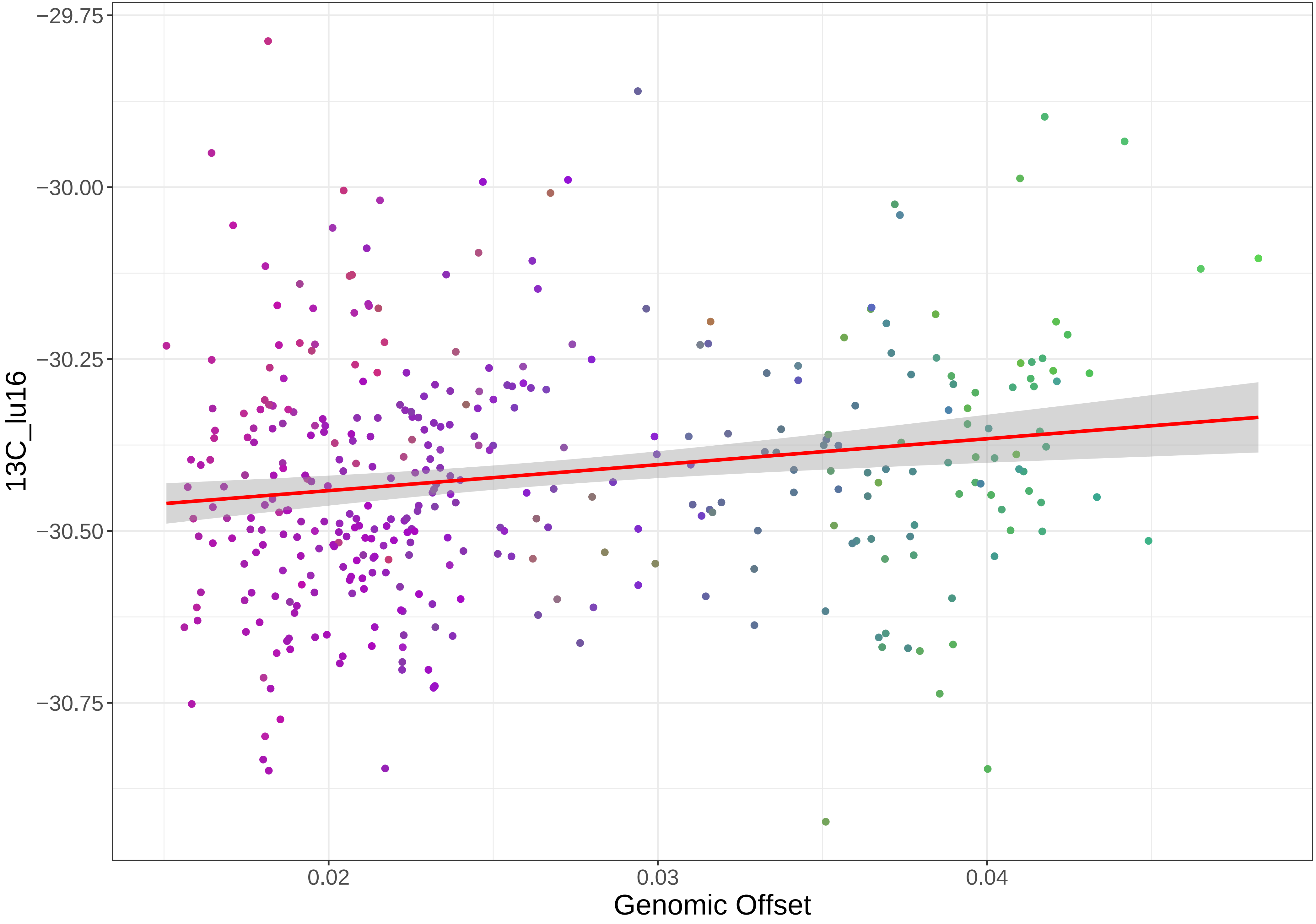

−29.75
−30.00
−30.25
−30.50
−30.75
13C_lu16
0.02
0.03
0.04
Genomic Offset

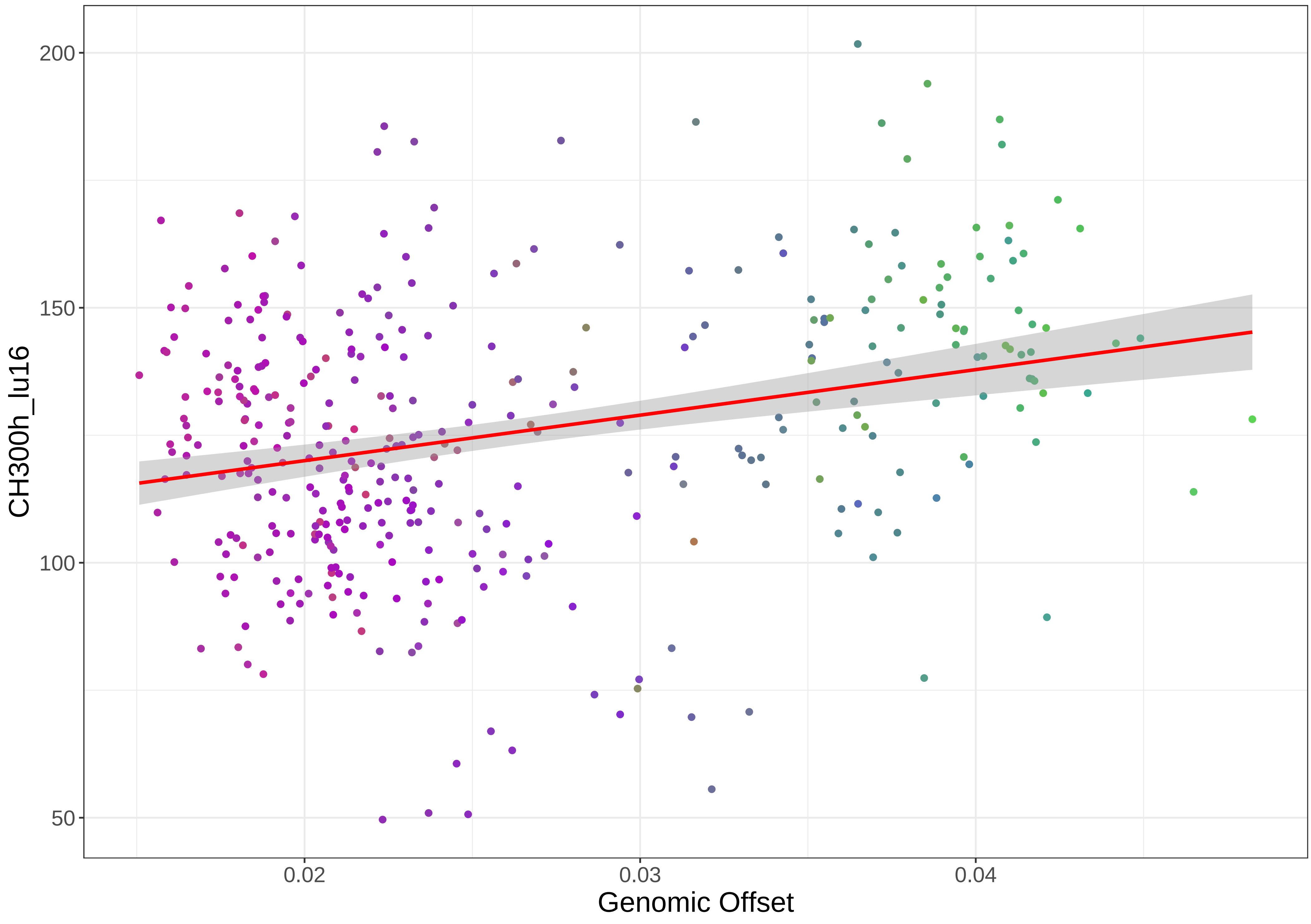

200
150
100
50
CH300h_lu16
0.02
0.03
0.04
Genomic Offset

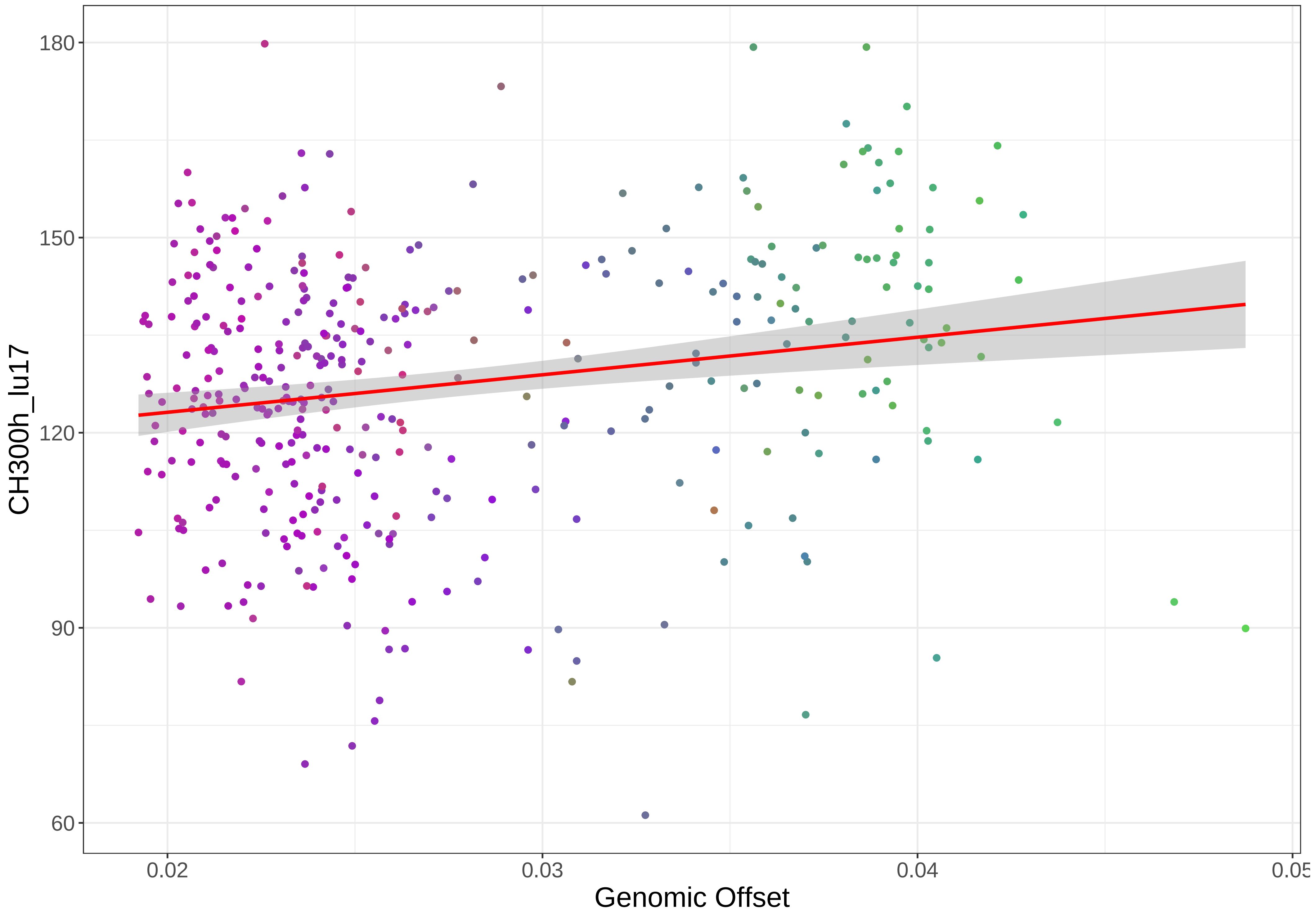
180
150
120
90
60
CH300h_lu17
0.02
0.03
0.04
0.05
Genomic Offset

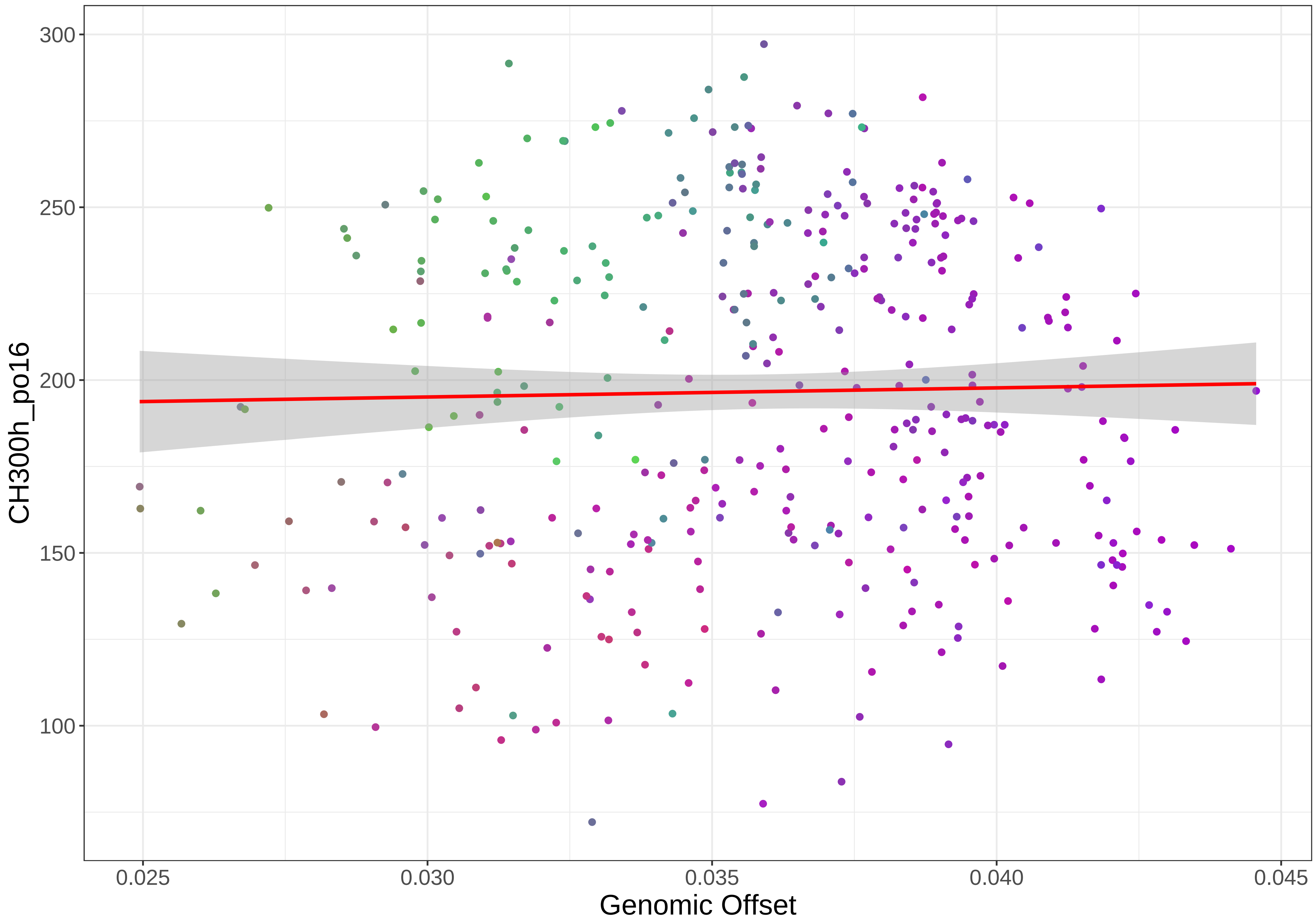

300
250
200
150
100
CH300h_po16
0.025
0.030
0.035
0.040
0.045
Genomic Offset

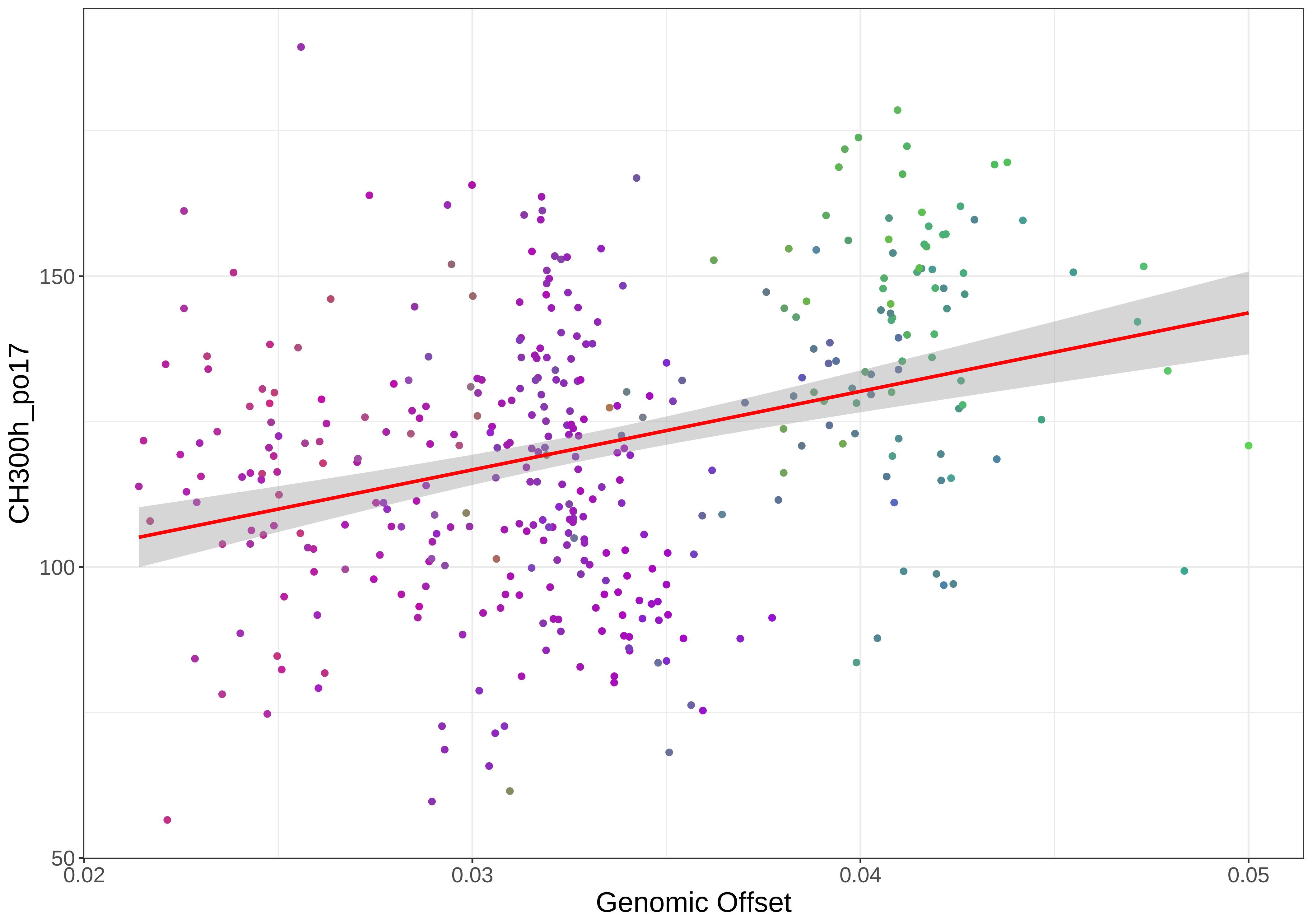

CH300h_po17
150
100
50
0.02
0.03
0.04
0.05
Genomic Offset

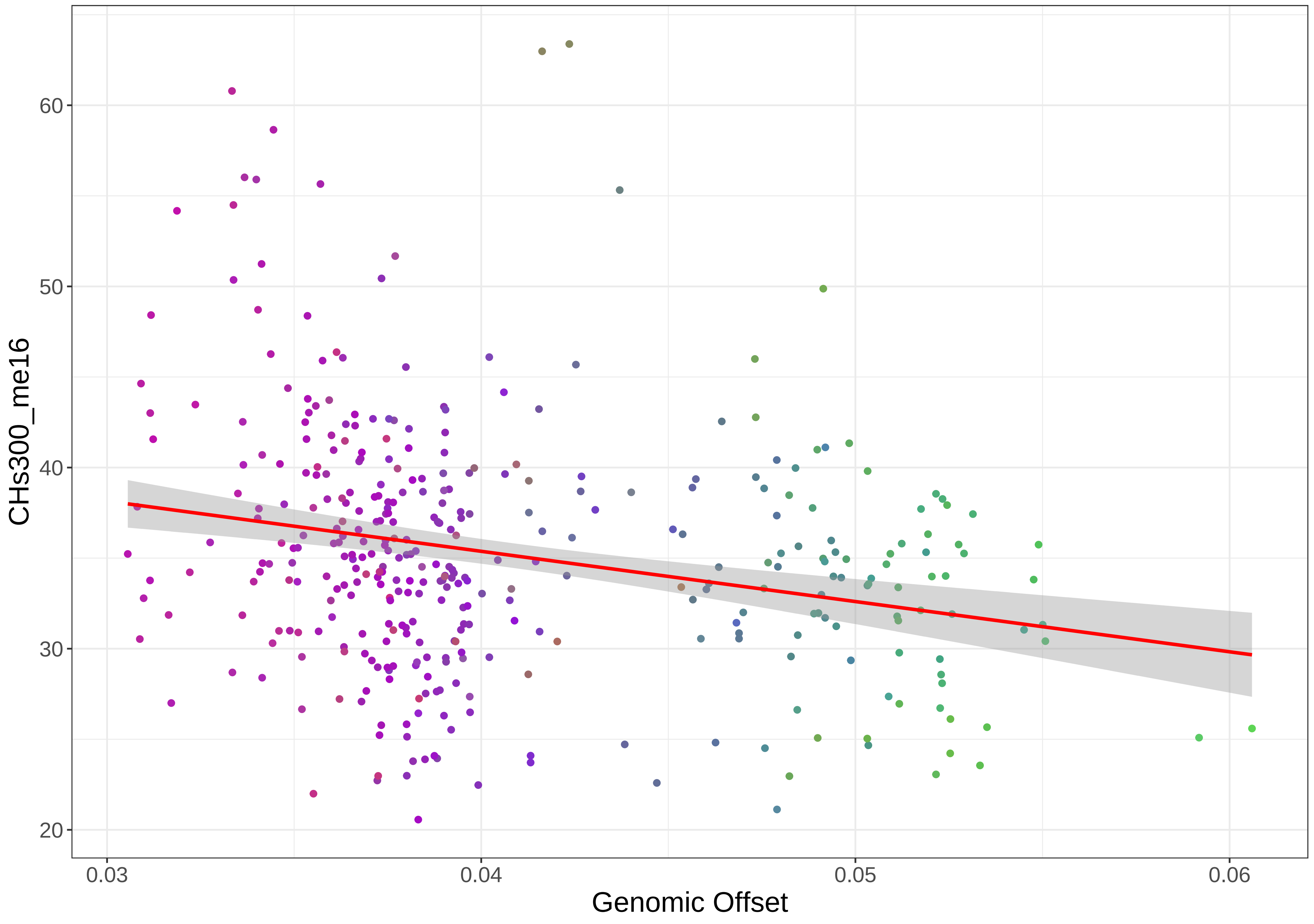

60
50
40
30
20
CHs300_me16
0.03
0.04
0.05
0.06
Genomic Offset

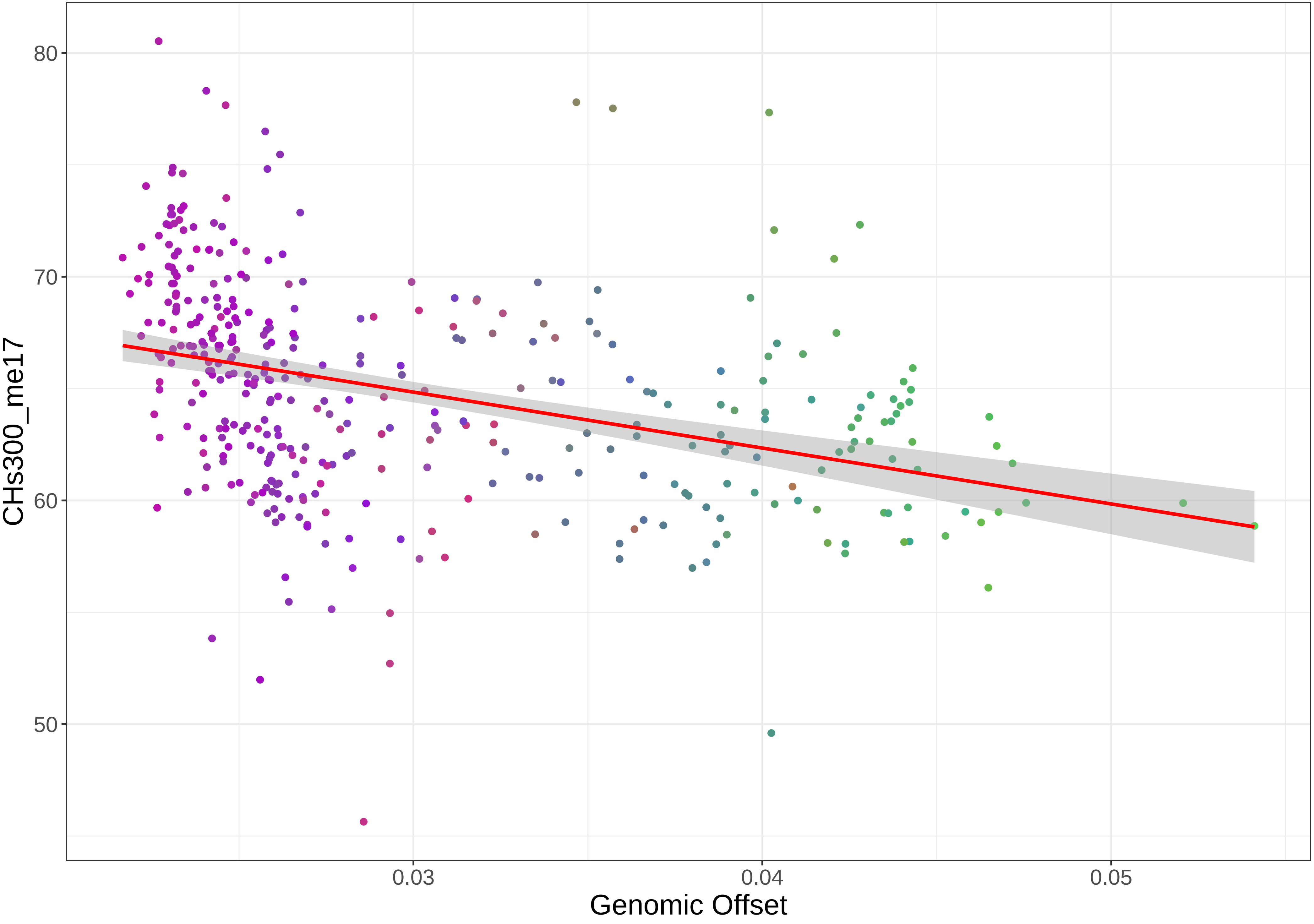

80
70
60
50
CHs300_me17
0.03
0.04
0.05
Genomic Offset

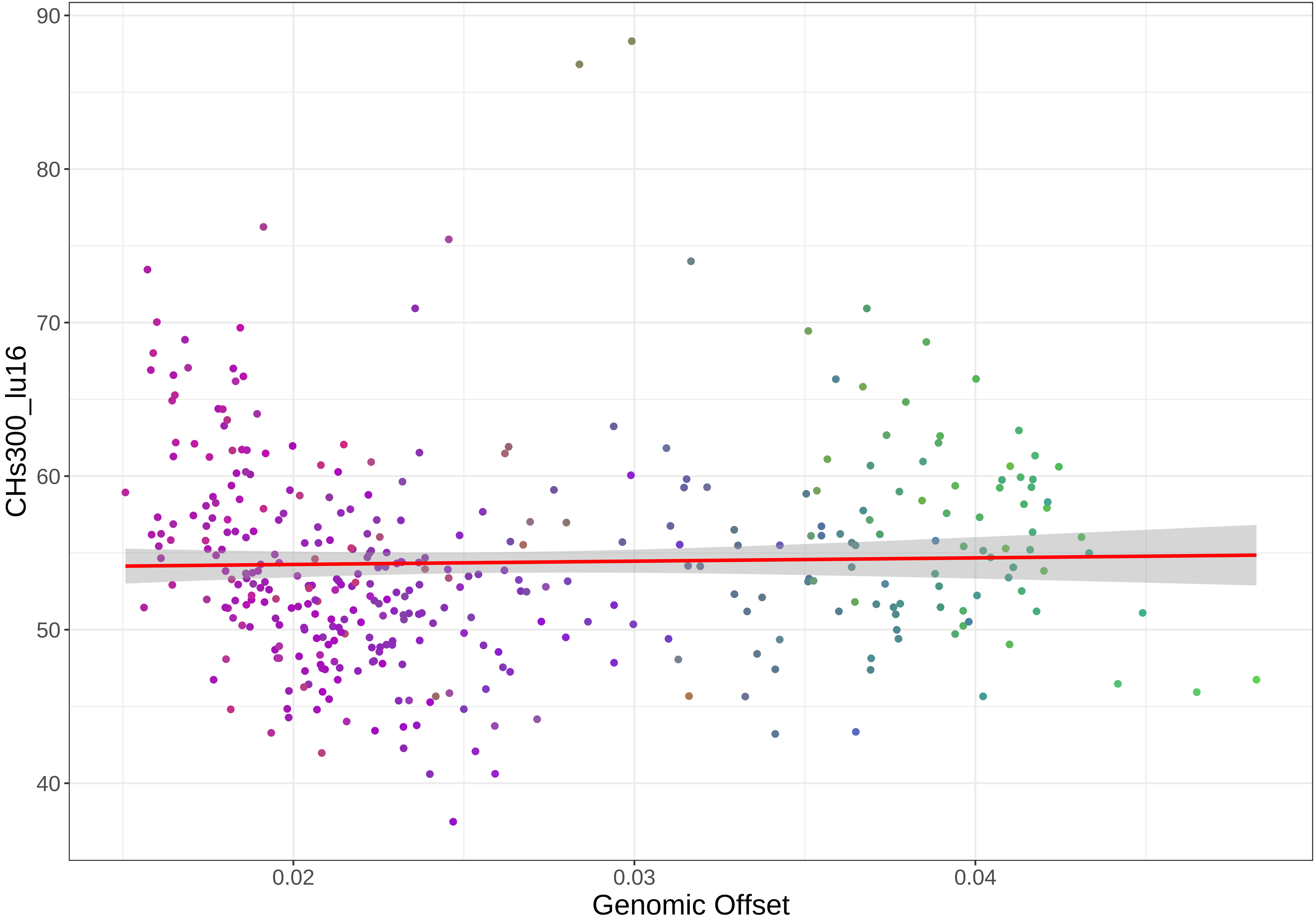
CHs300_lu16
90
80
70
60
50
40
0.02
0.03
0.04
Genomic Offset

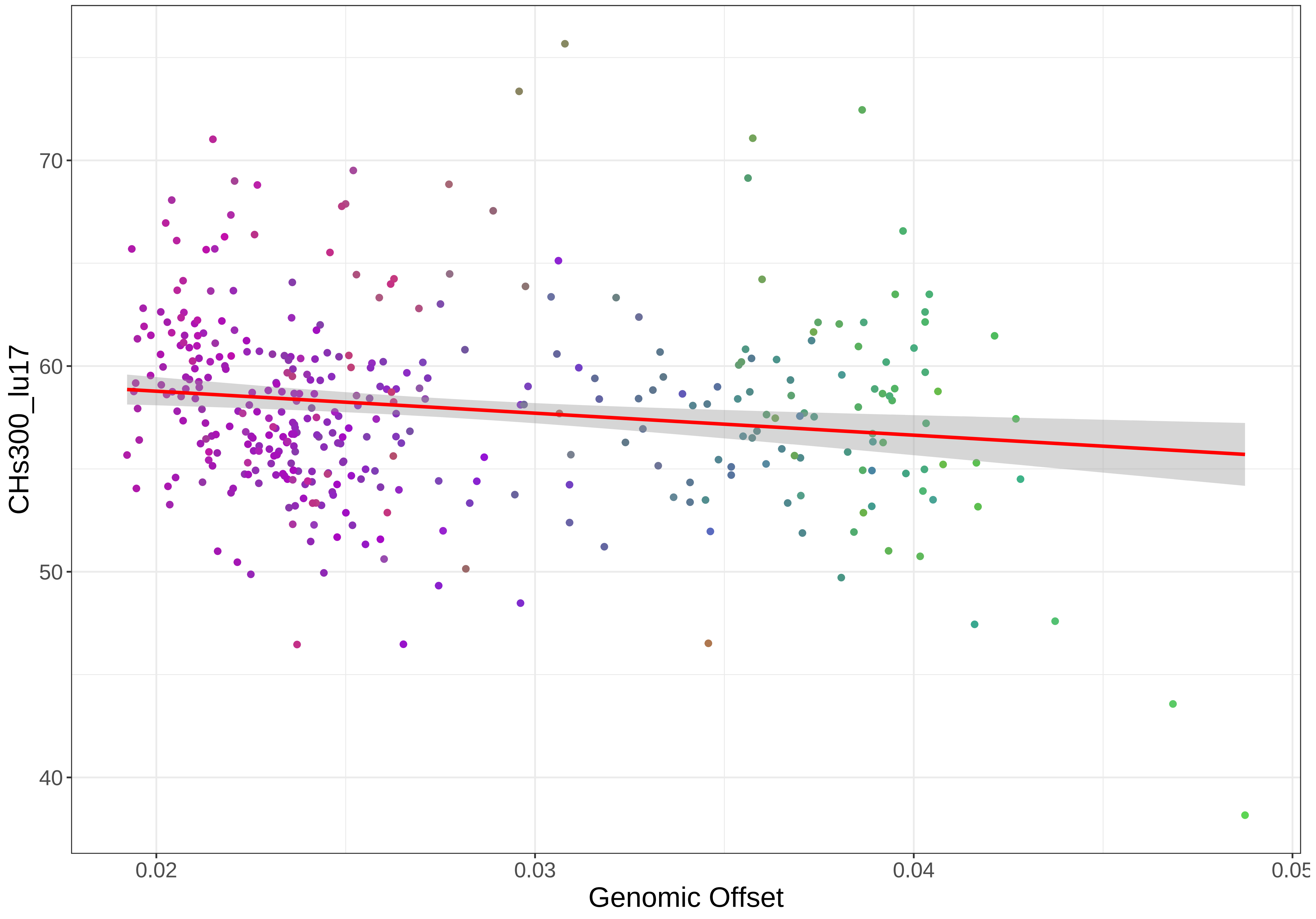
70
60
50
40
CHs300_lu17
0.02
0.03
0.04
0.05
Genomic Offset

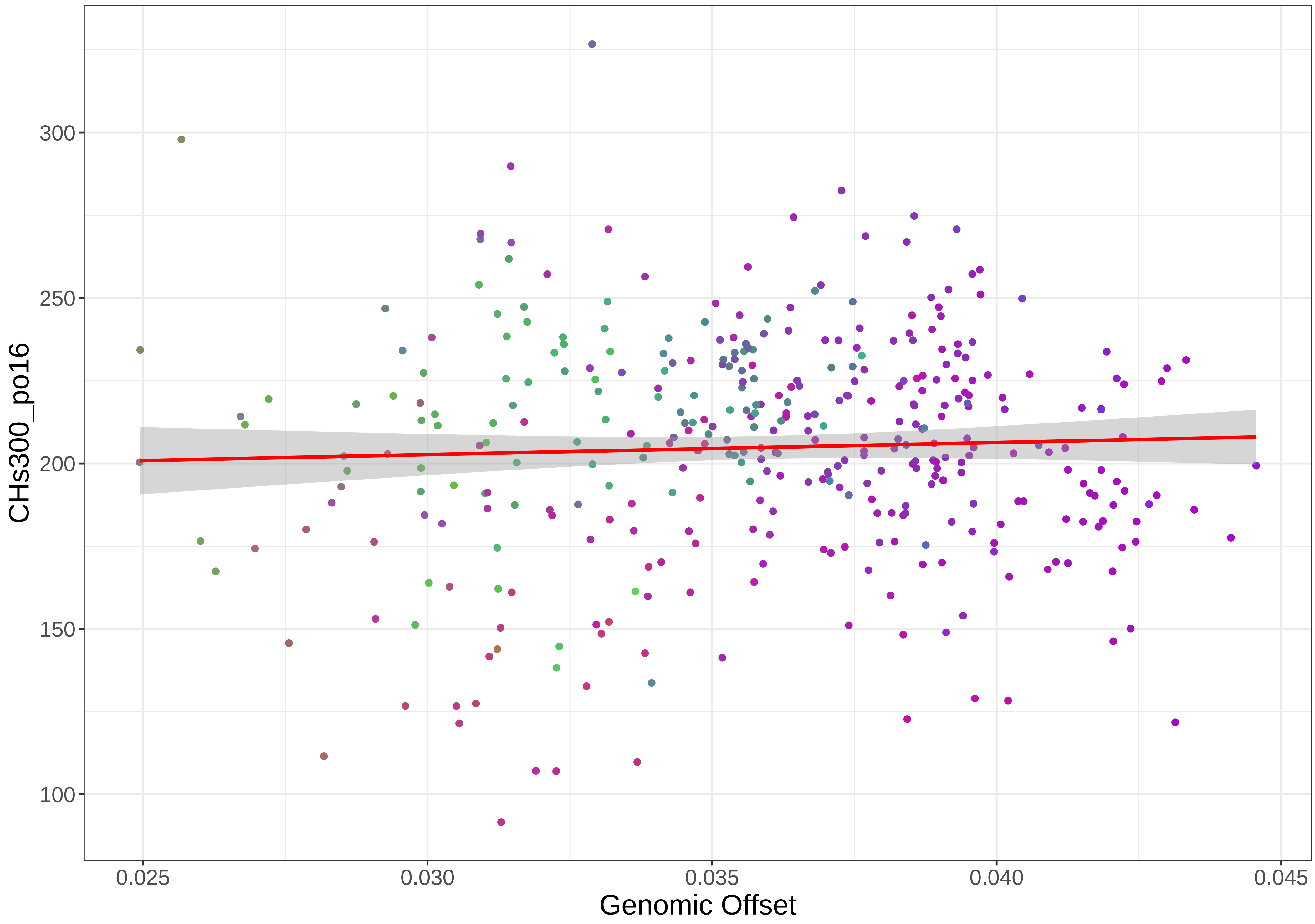

CHs300_po16
300
250
200
150
100
0.025
0.030
0.035
0.040
0.045
Genomic Offset

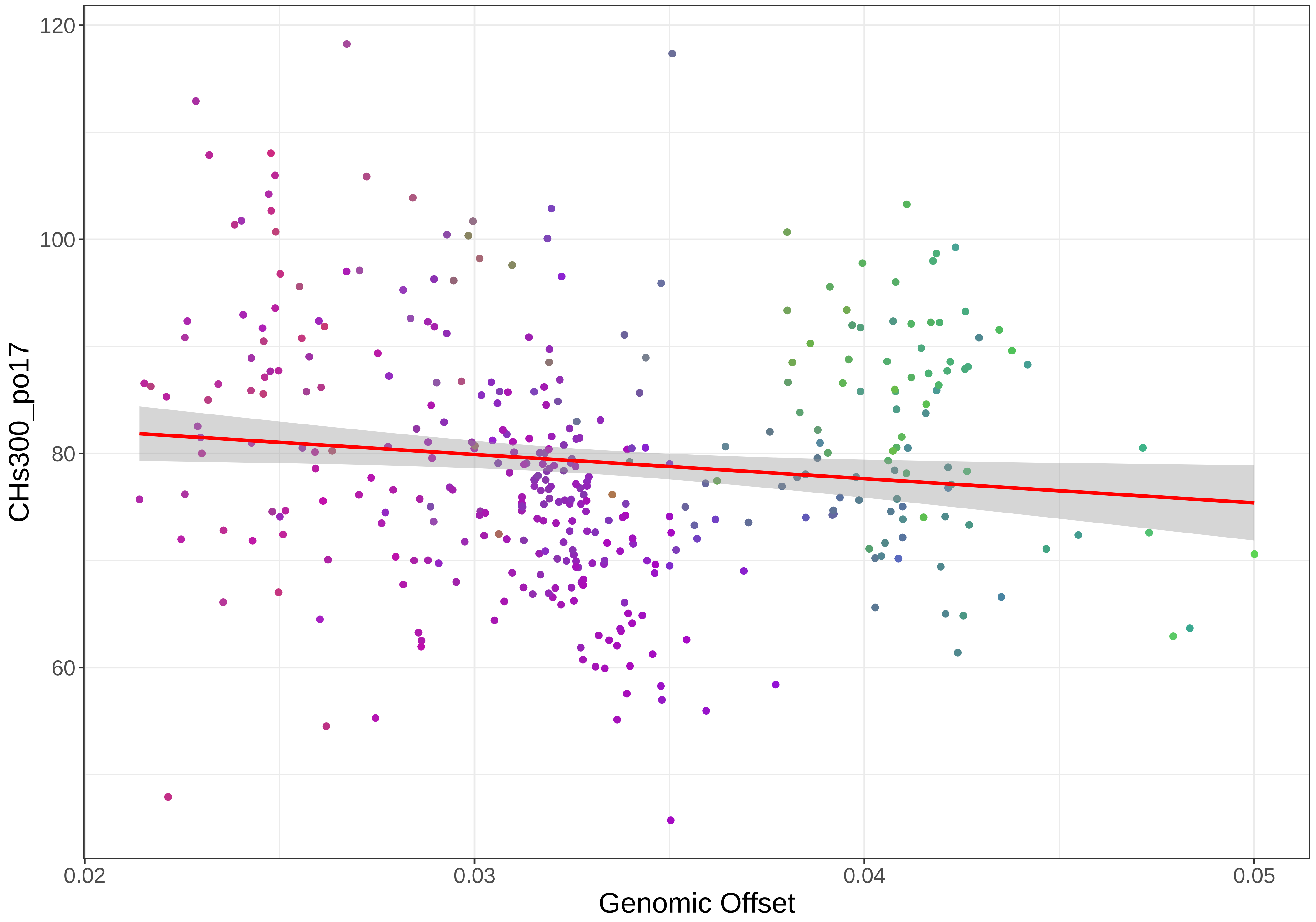

120
100
80
60
CHs300_po17
0.02
0.03
0.04
0.05
Genomic Offset

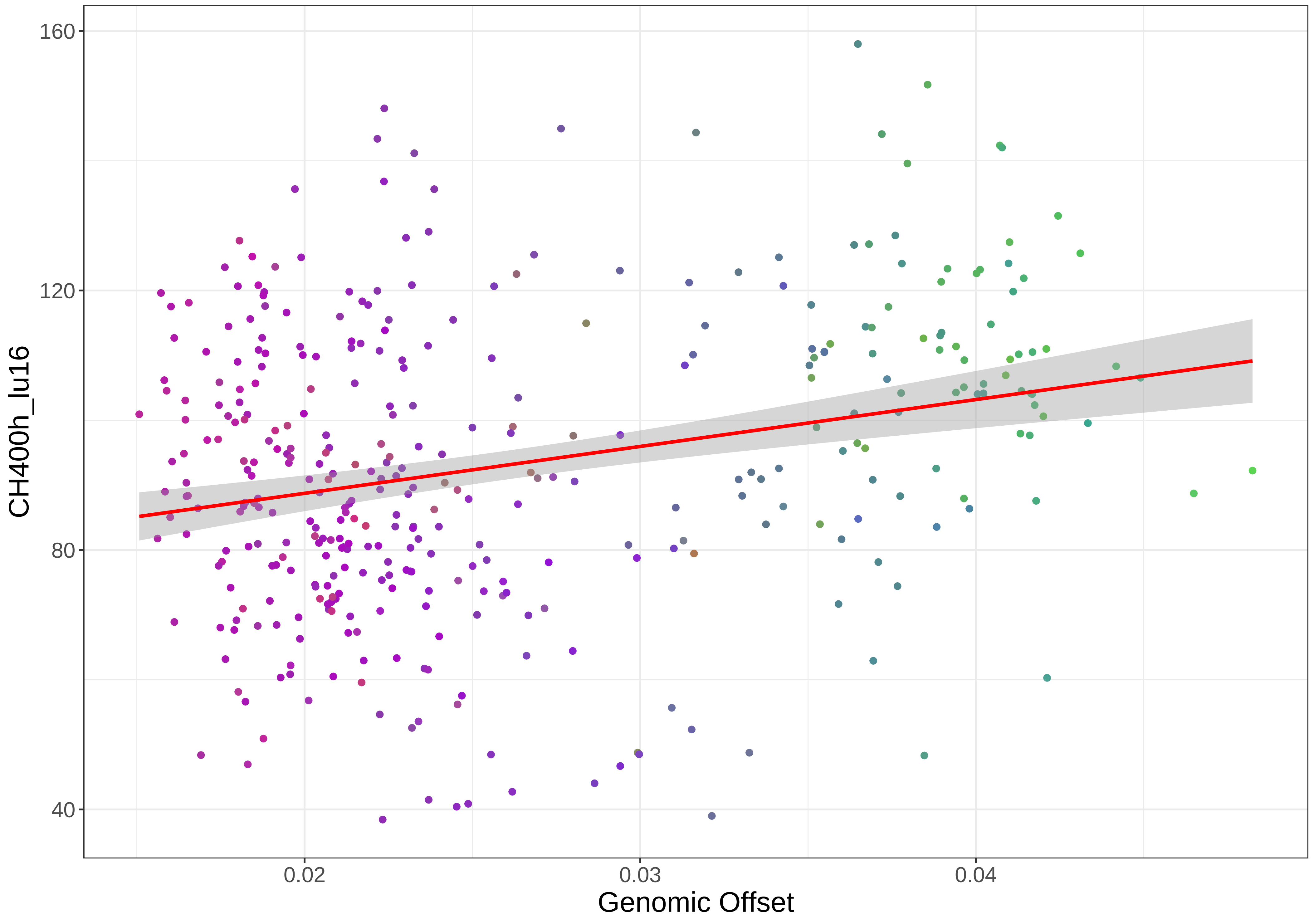

160
120
80
40
0.02
0.03
0.04
CH400h_lu16
Genomic Offset

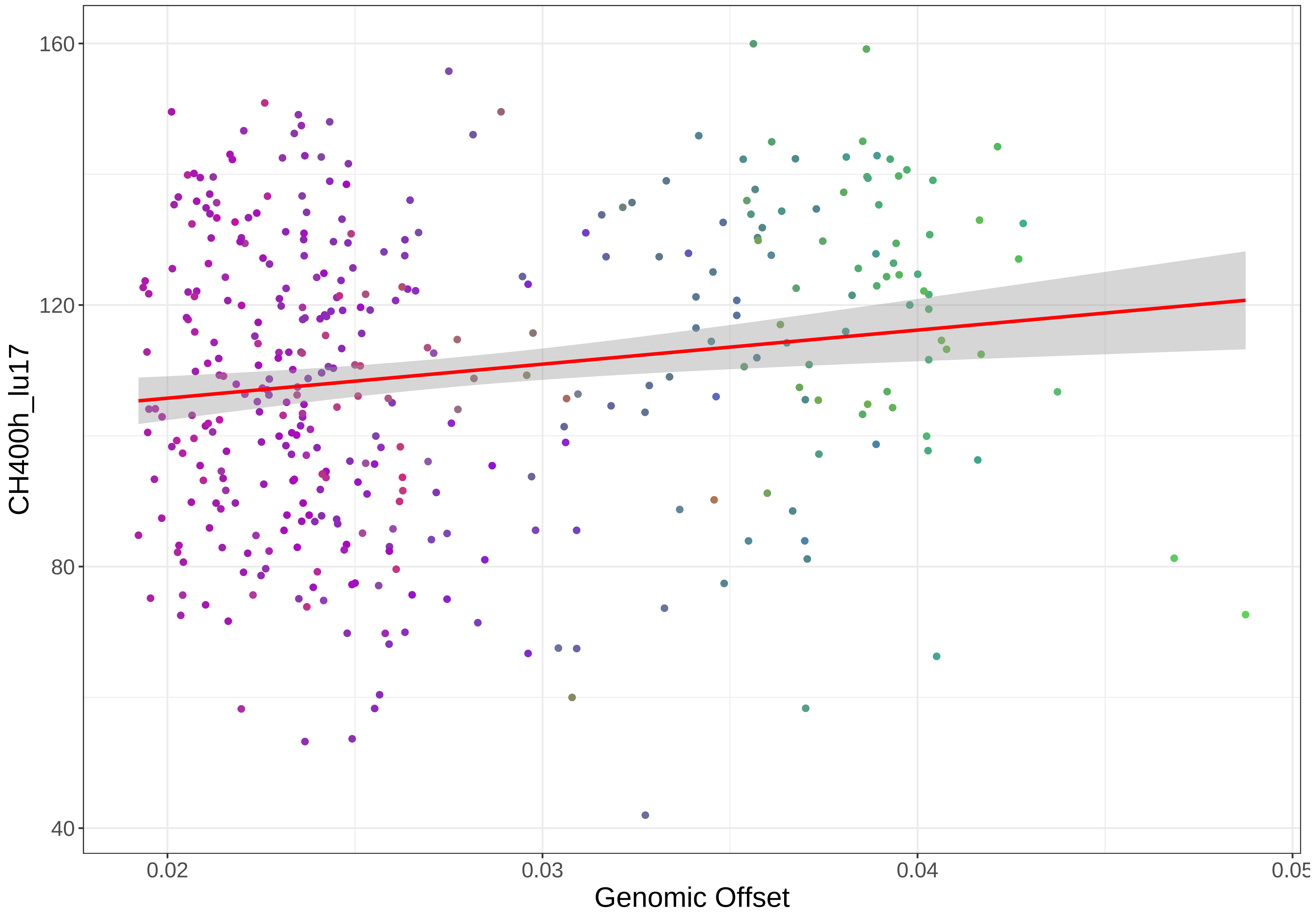
160
120
80
40
CH400h_lu17
0.02
0.03
0.04
0.05
Genomic Offset

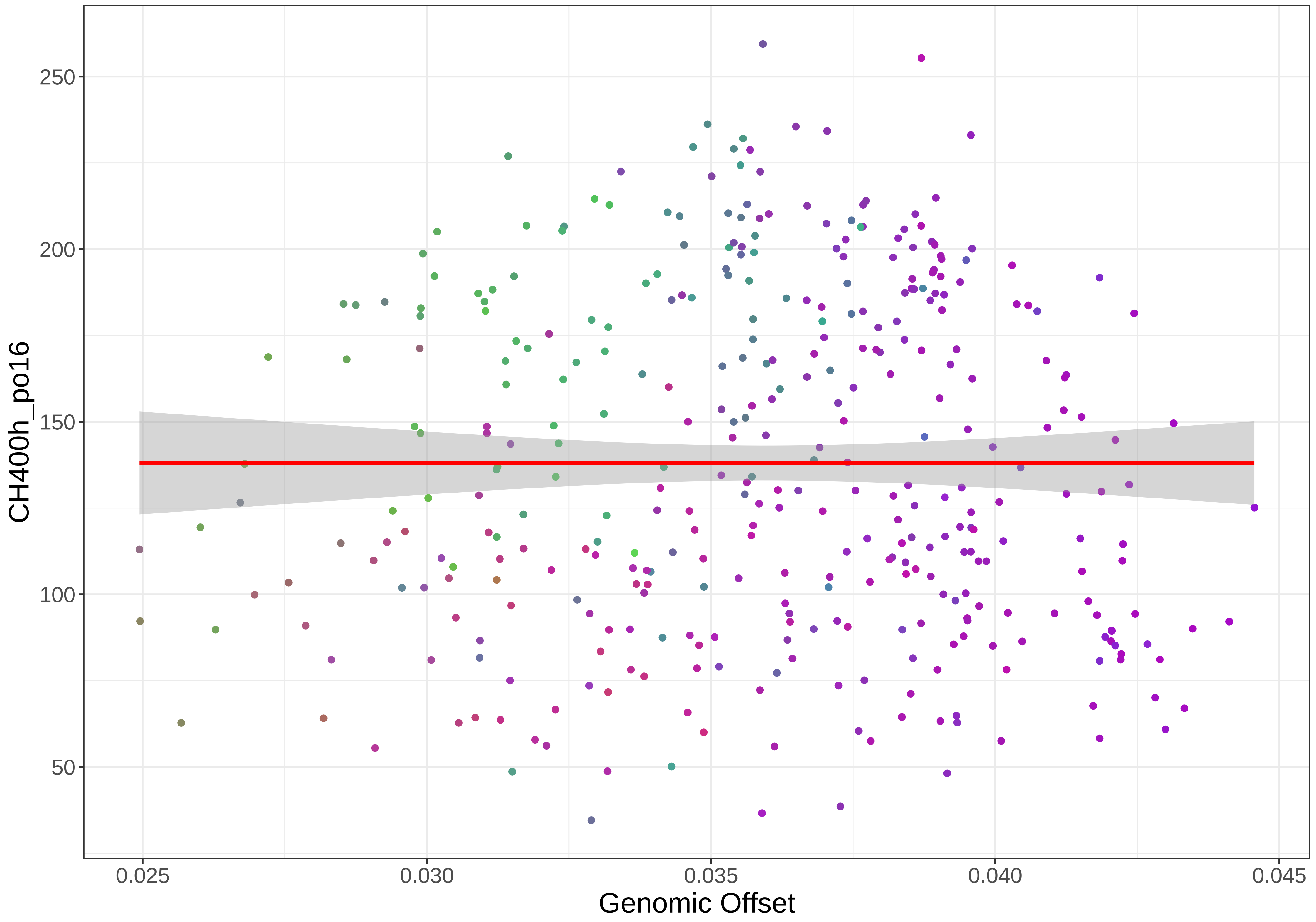

CH400h_po16
250
200
150
100
50
0.025
0.030
0.035
0.040
0.045
Genomic Offset

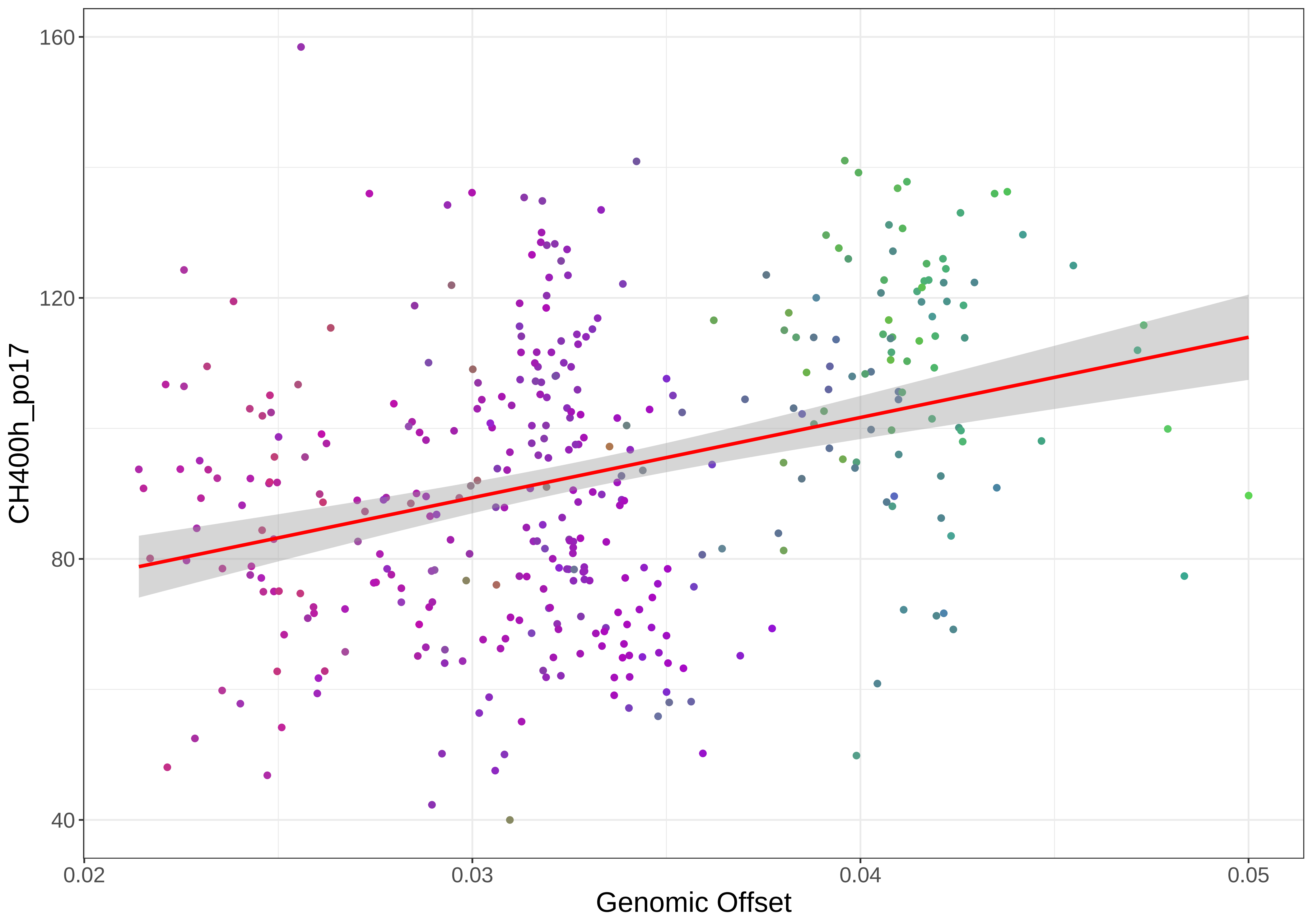

160
120
80
40
CH400h_po17
0.02
0.03
0.04
0.05
Genomic Offset

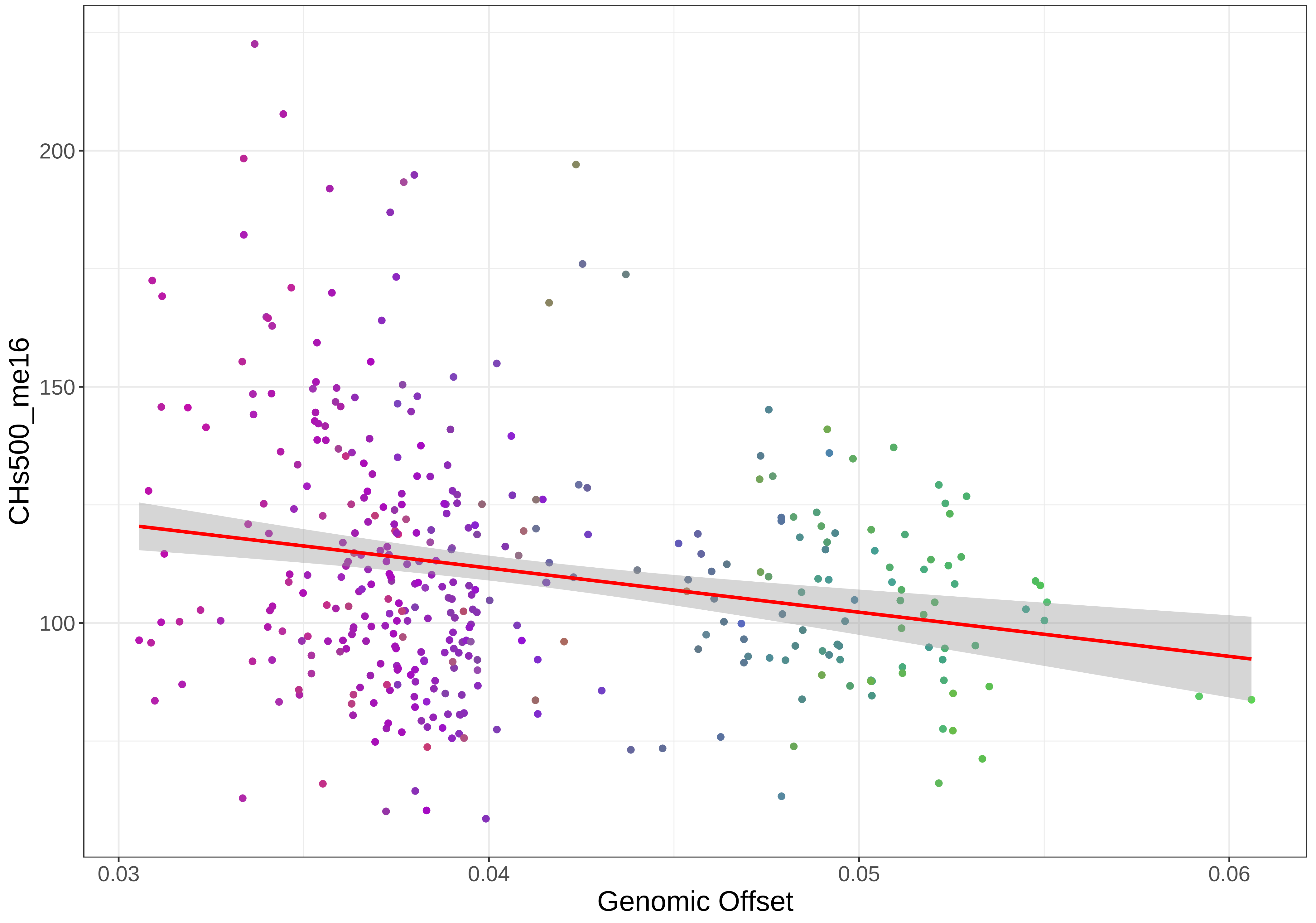

200
150
100
0.03
0.04
0.05
0.06
CHs500_me16
Genomic Offset

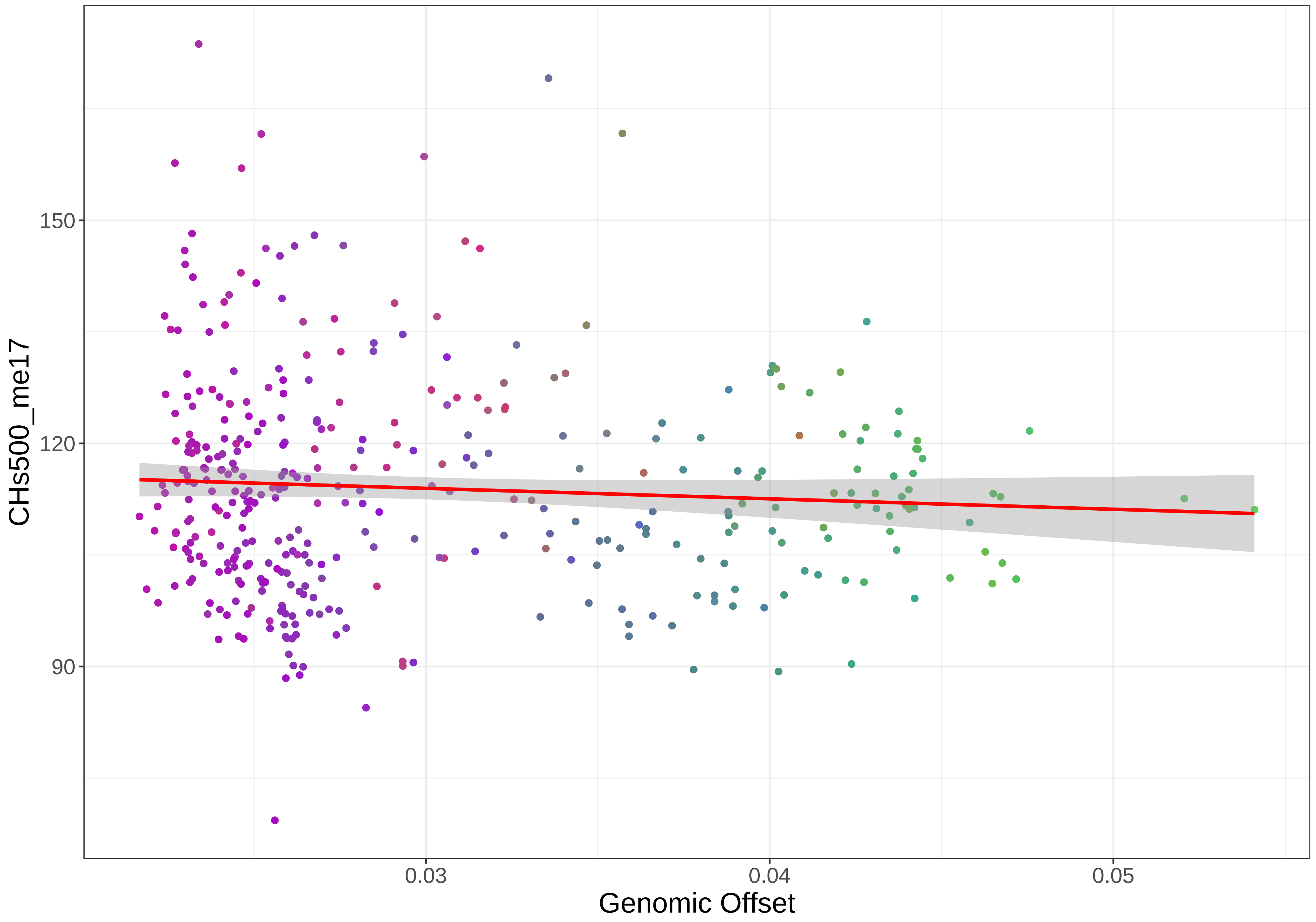

150
120
90
CHs500_me17
0.03
0.04
0.05
Genomic Offset

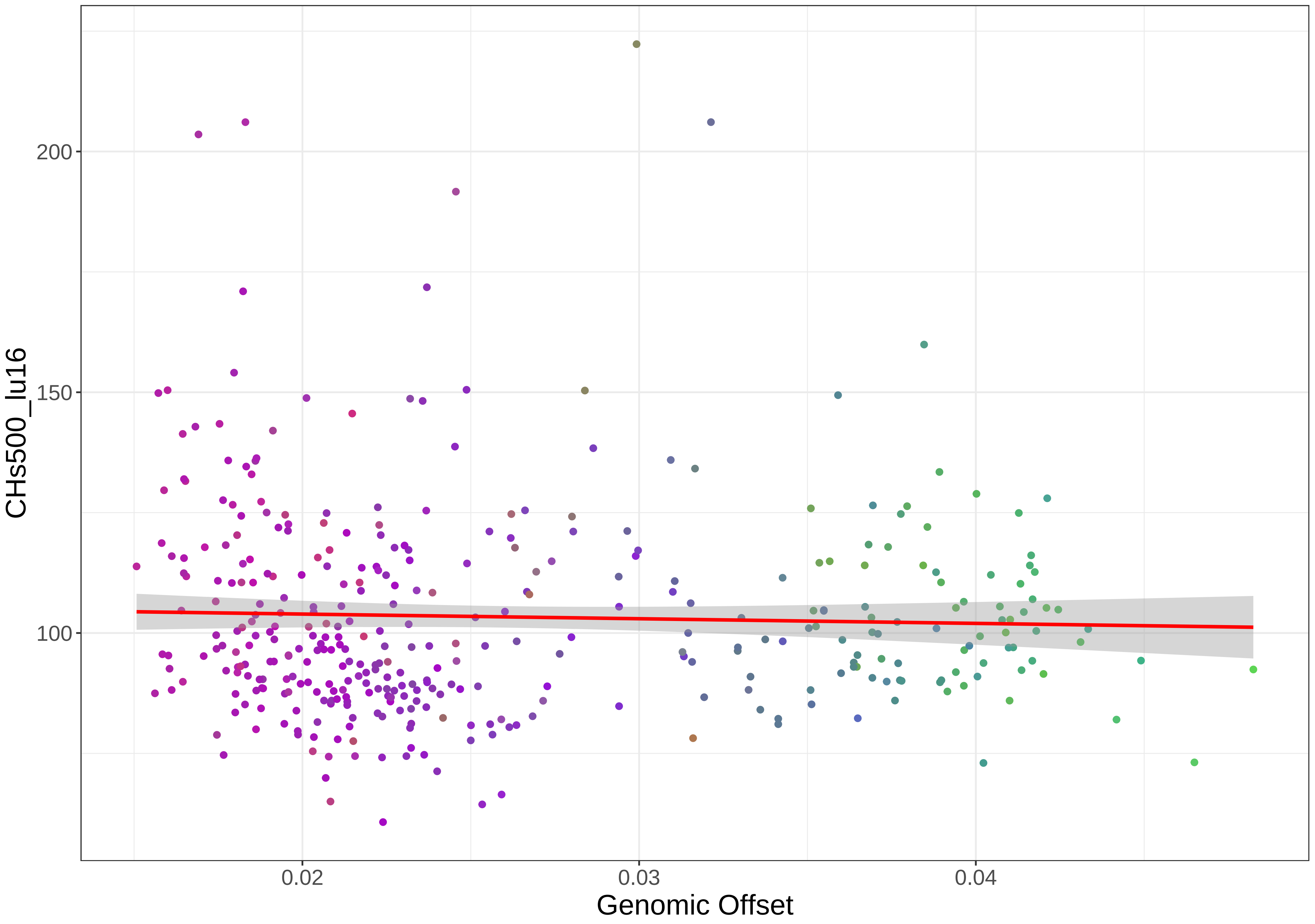

200
150
100
CHs500_lu16
0.02
0.03
0.04
Genomic Offset

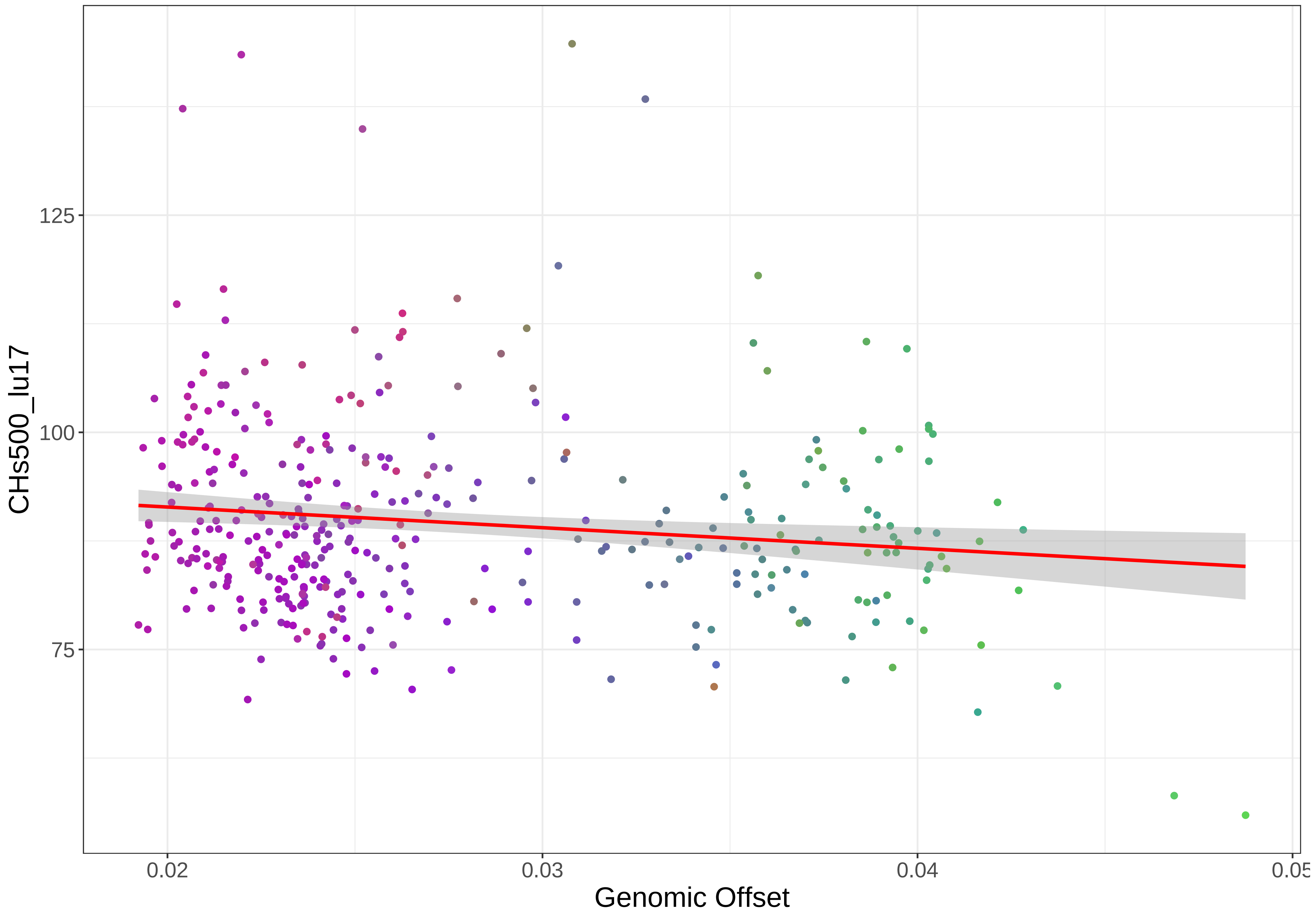

CHs500_lu17
125
100
75
0.02
0.03
0.04
0.05
Genomic Offset

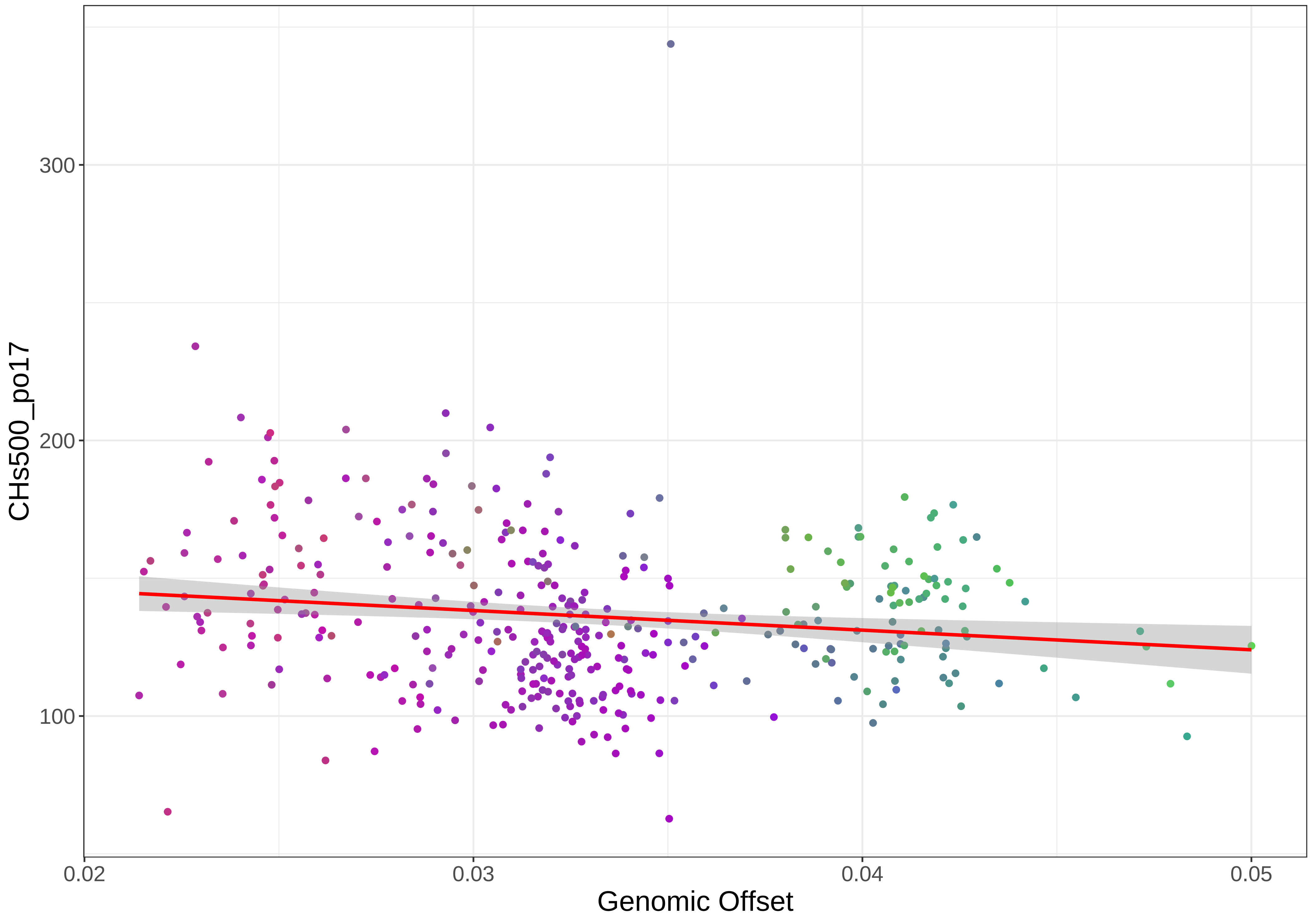
CHs500_po17
300
200
100
0.02
0.03
0.04
0.05
Genomic Offset

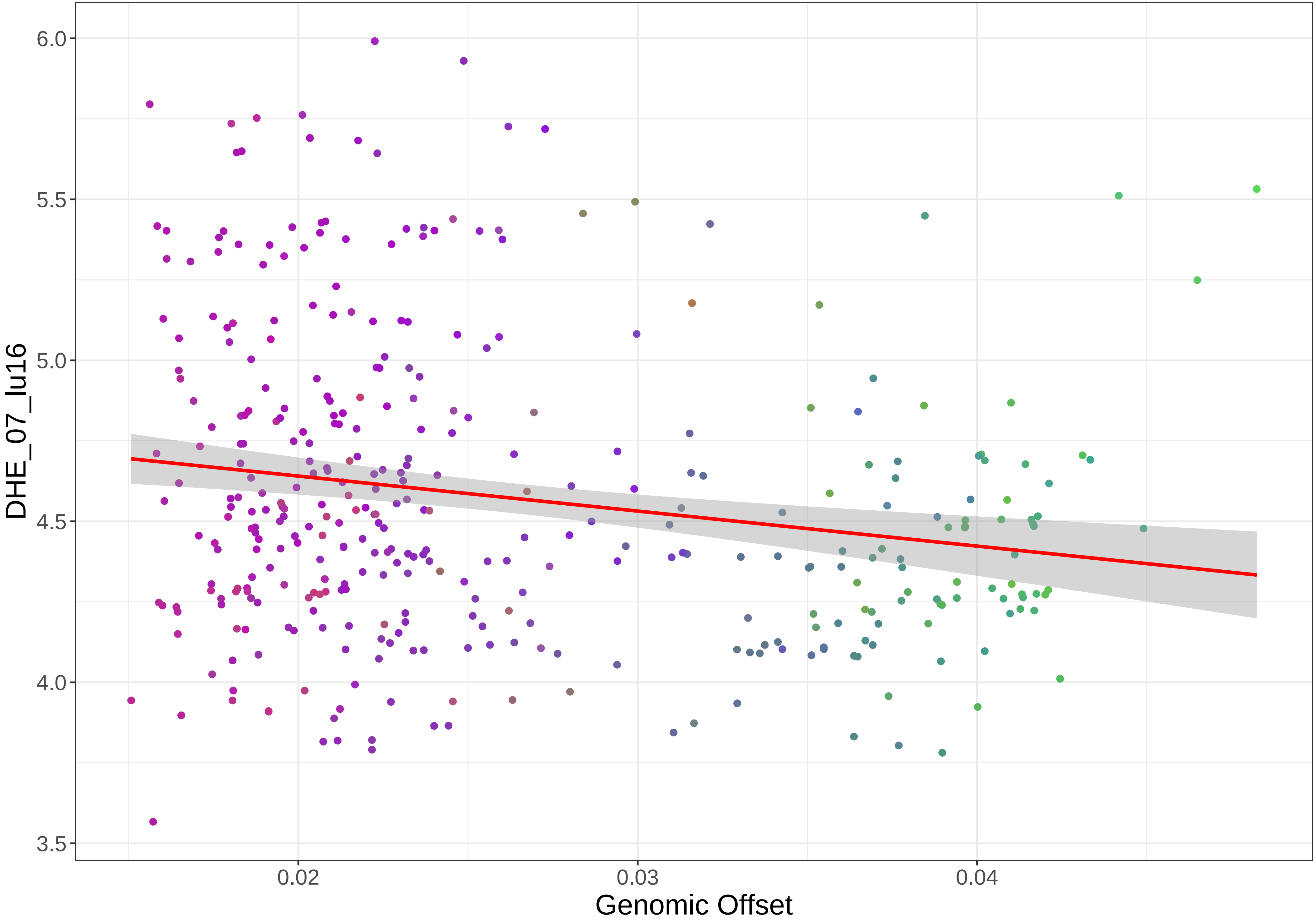

6.0
5.5
5.0
4.5
4.0
3.5
DHE_07_lu16
0.02
0.03
0.04
Genomic Offset

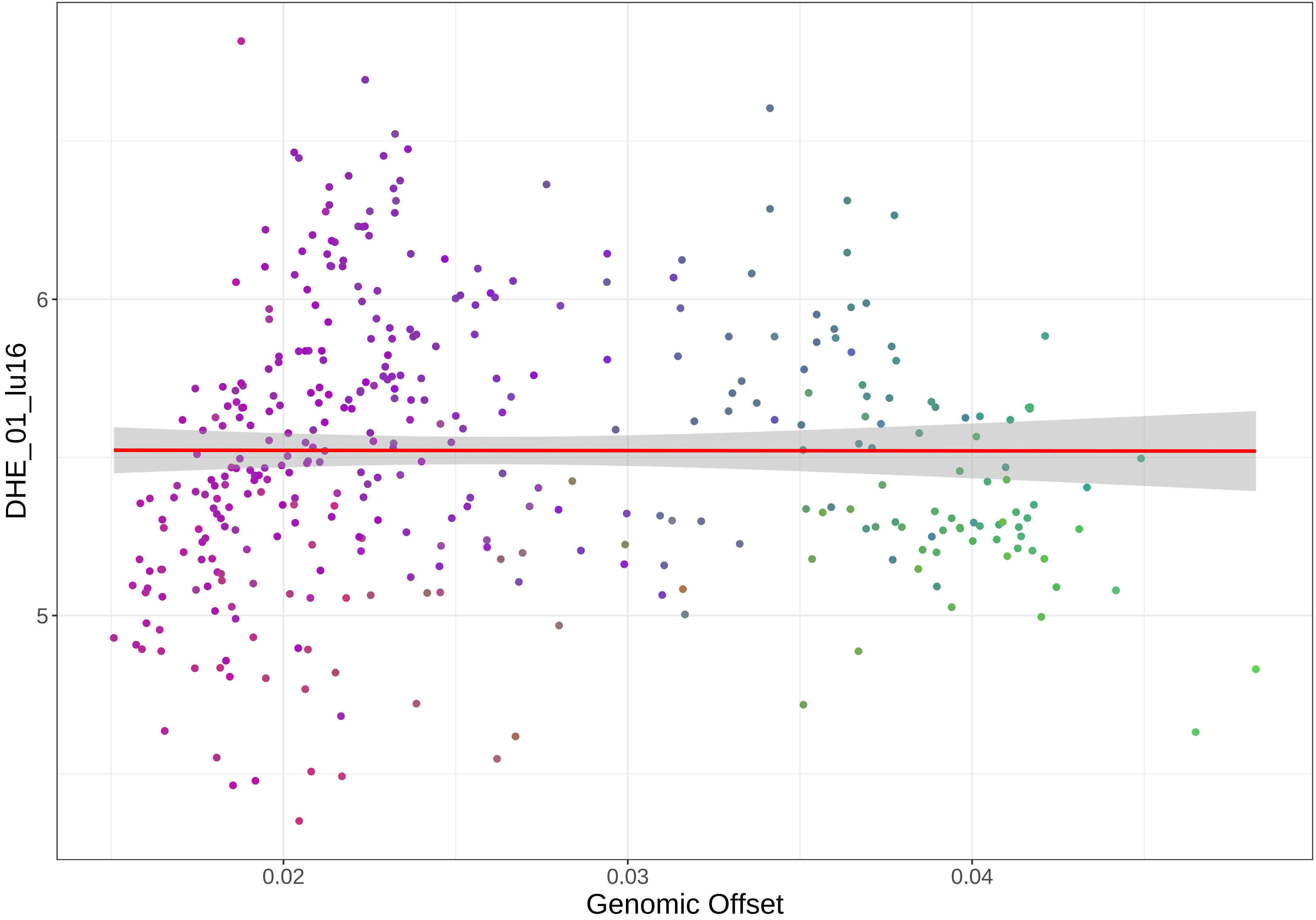

DHE_01_lu16
6
5
0.02
0.03
0.04
Genomic Offset

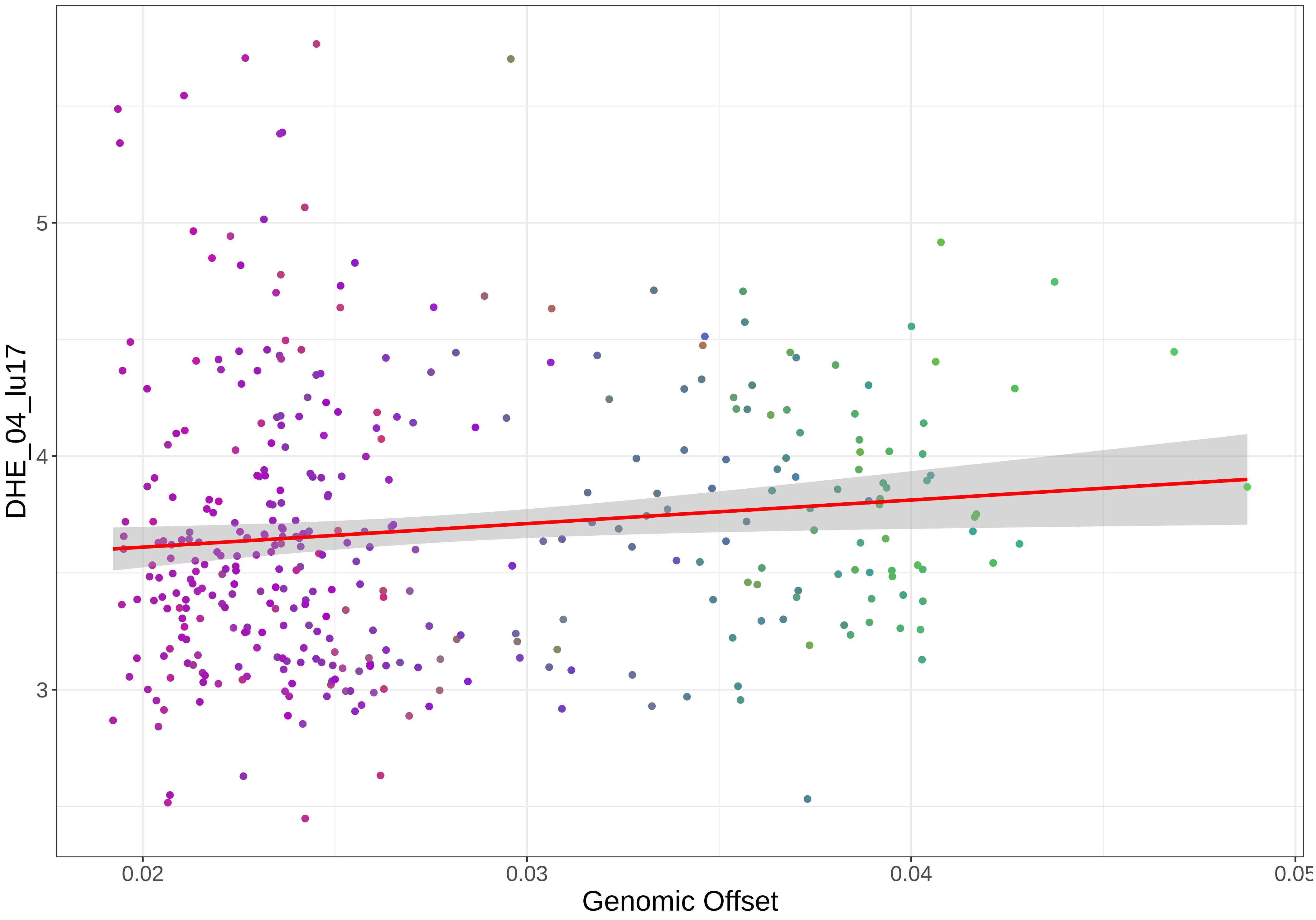
DHE_04_lu17
5
4
3
0.02
0.03
0.04
Genomic Offset

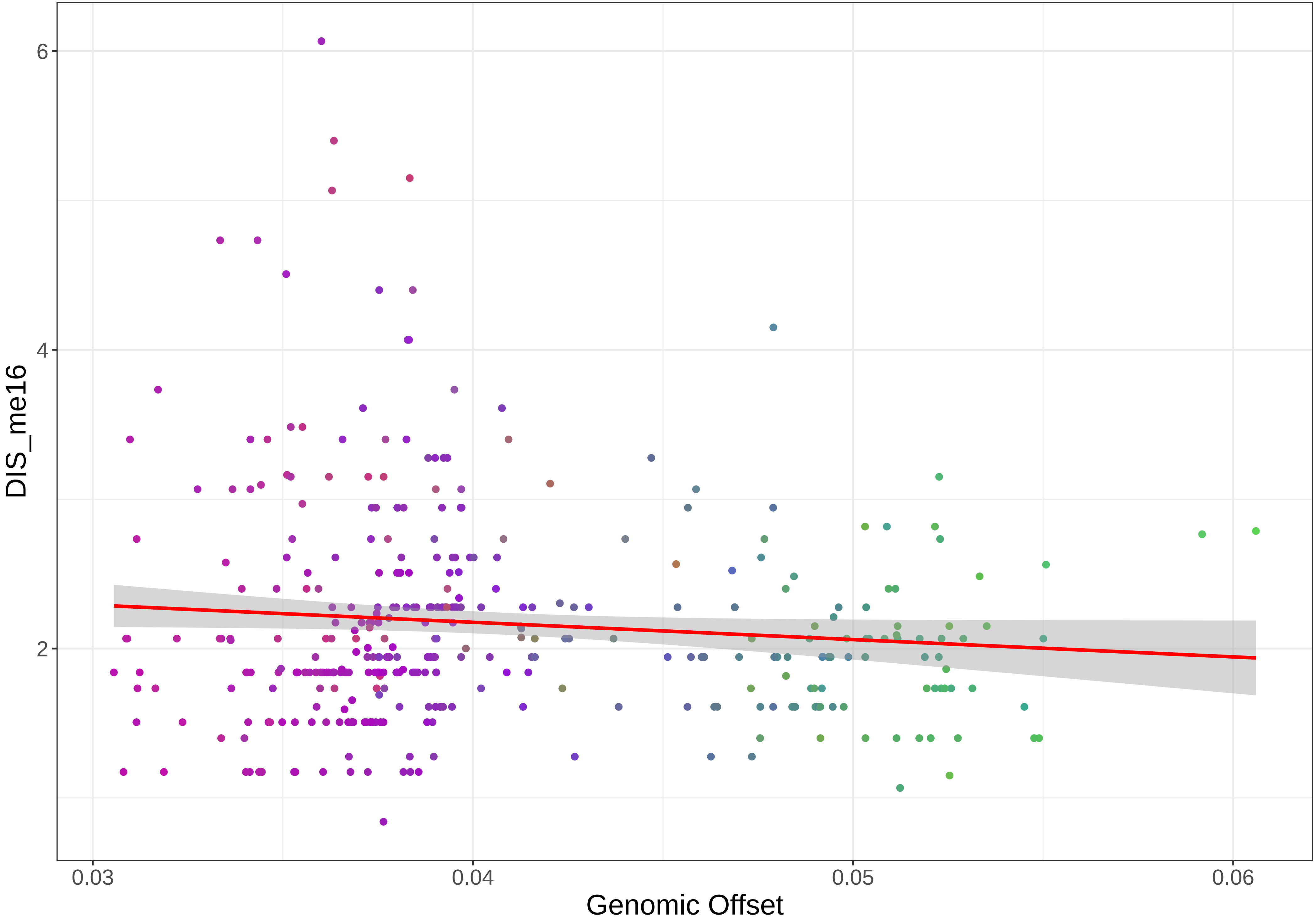

DIS_me16
6
4
2
0.03
0.04
0.05
0.06
Genomic Offset

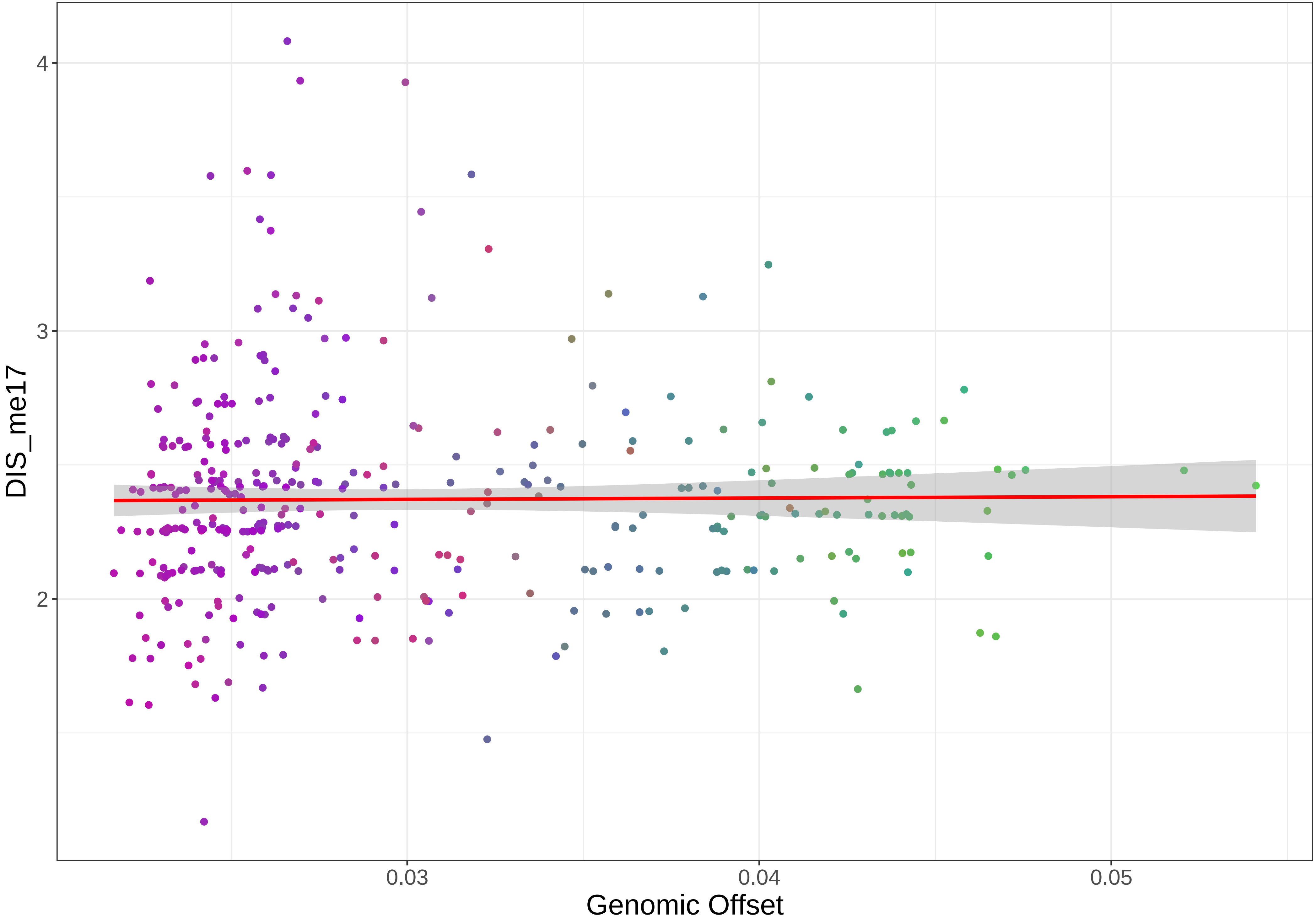
DIS_me17
4
3
2
0.03
0.04
0.05
Genomic Offset

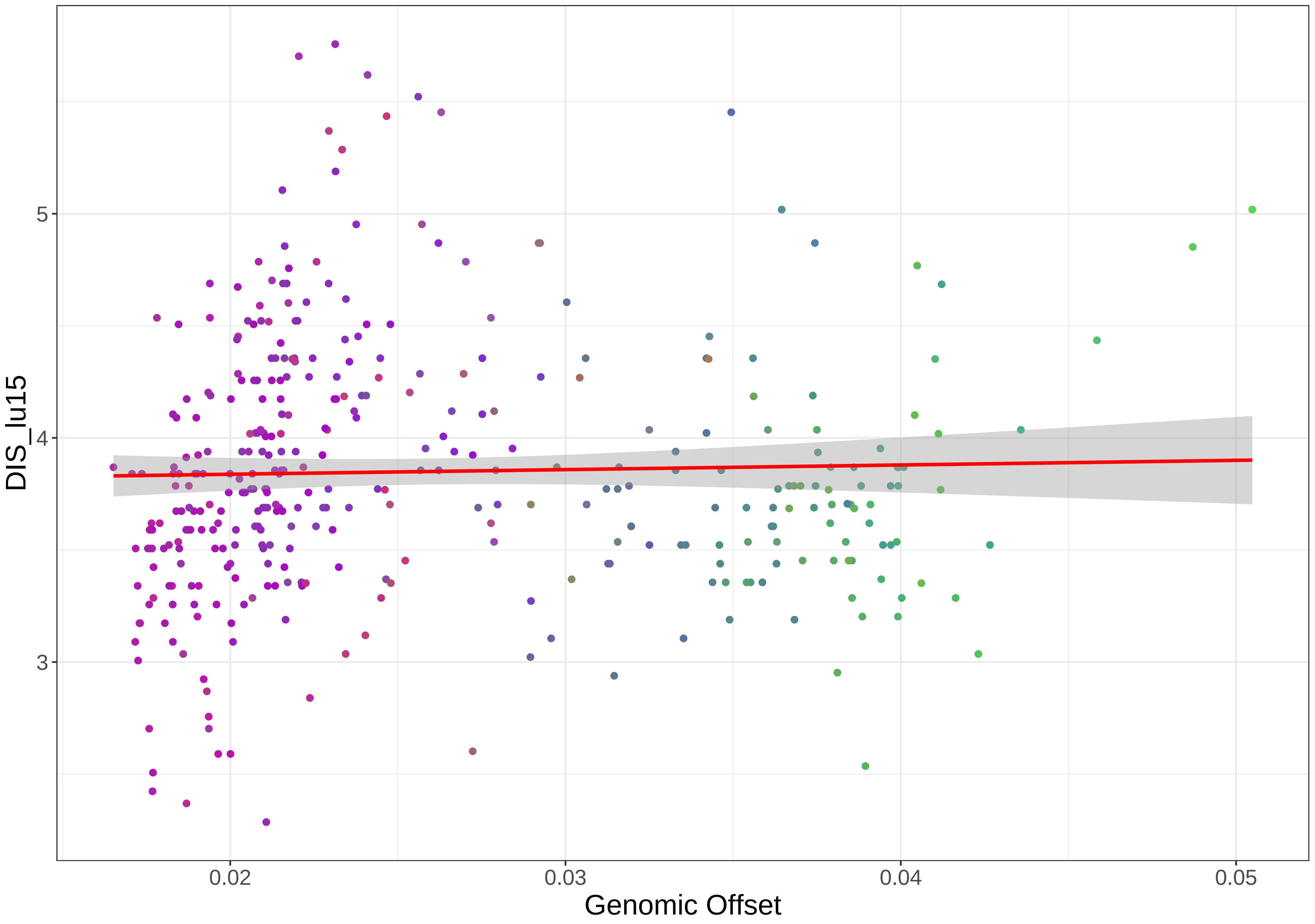
DIS_lu15
5
4
3
0.02
0.03
0.04
0.05
Genomic Offset

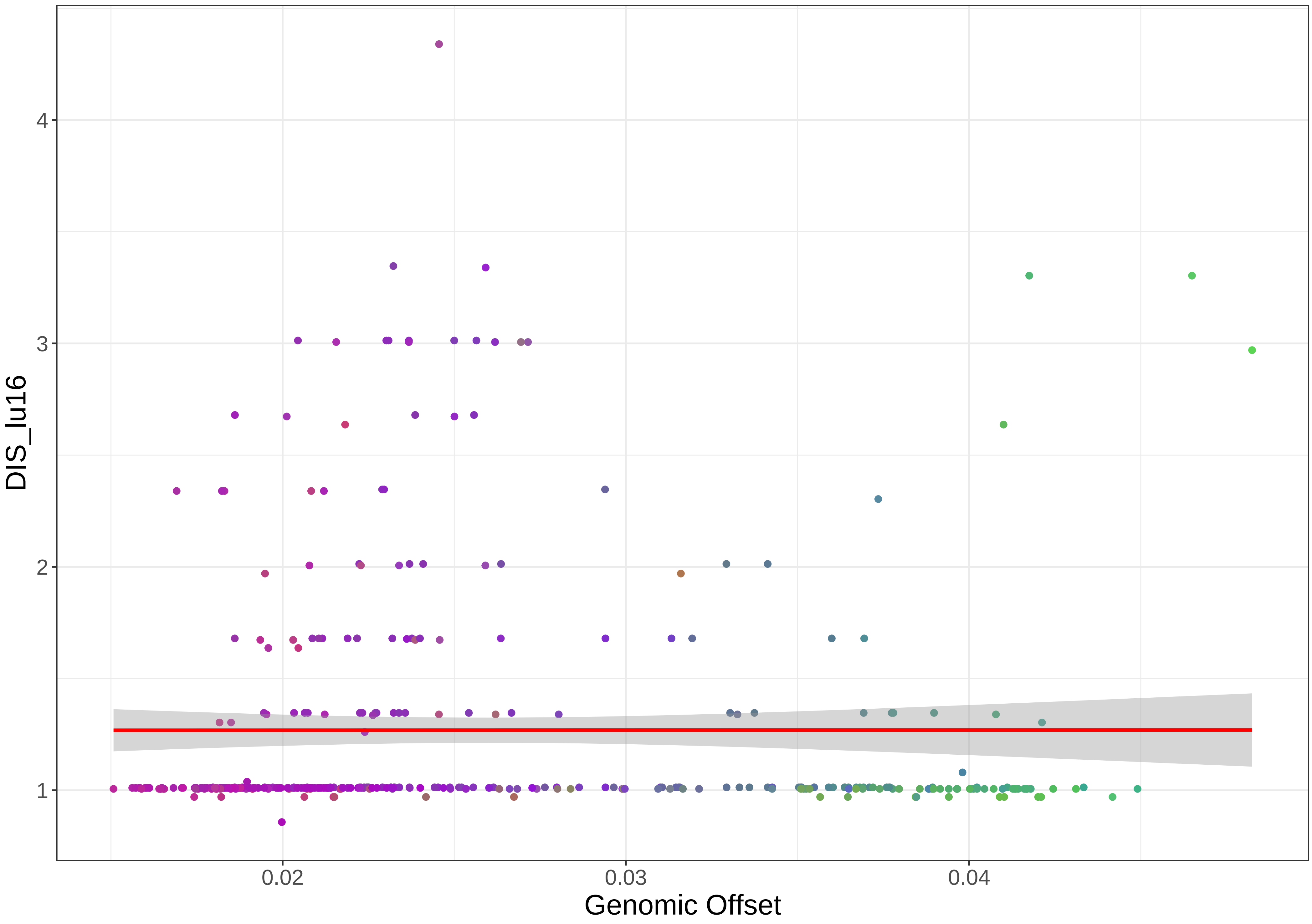

DIS_lu16
4
3
2
1
0.02
0.03
0.04
Genomic Offset

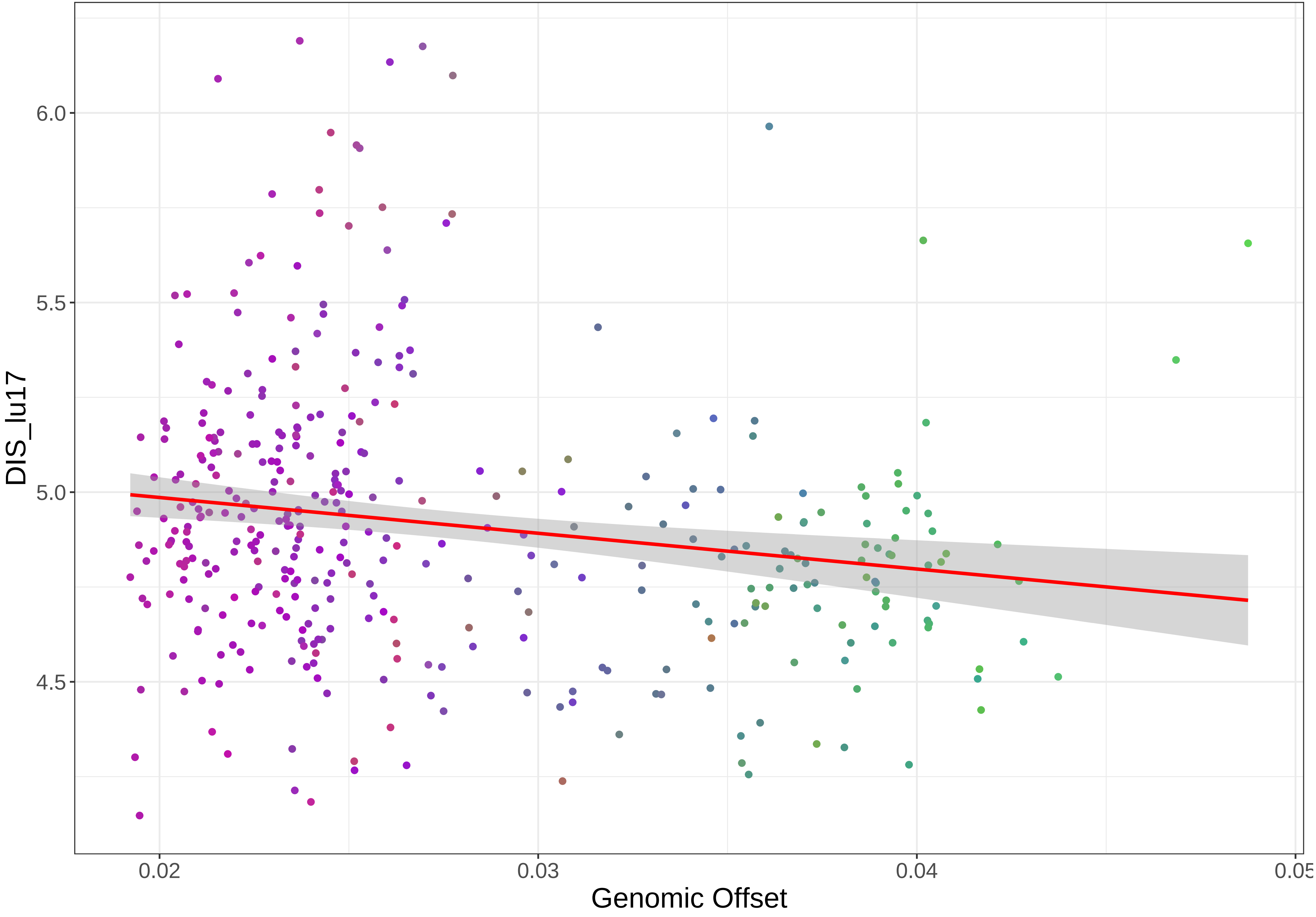

DIS_lu17
6.0
5.5
5.0
4.5
0.02
0.03
0.04
0.05
Genomic Offset

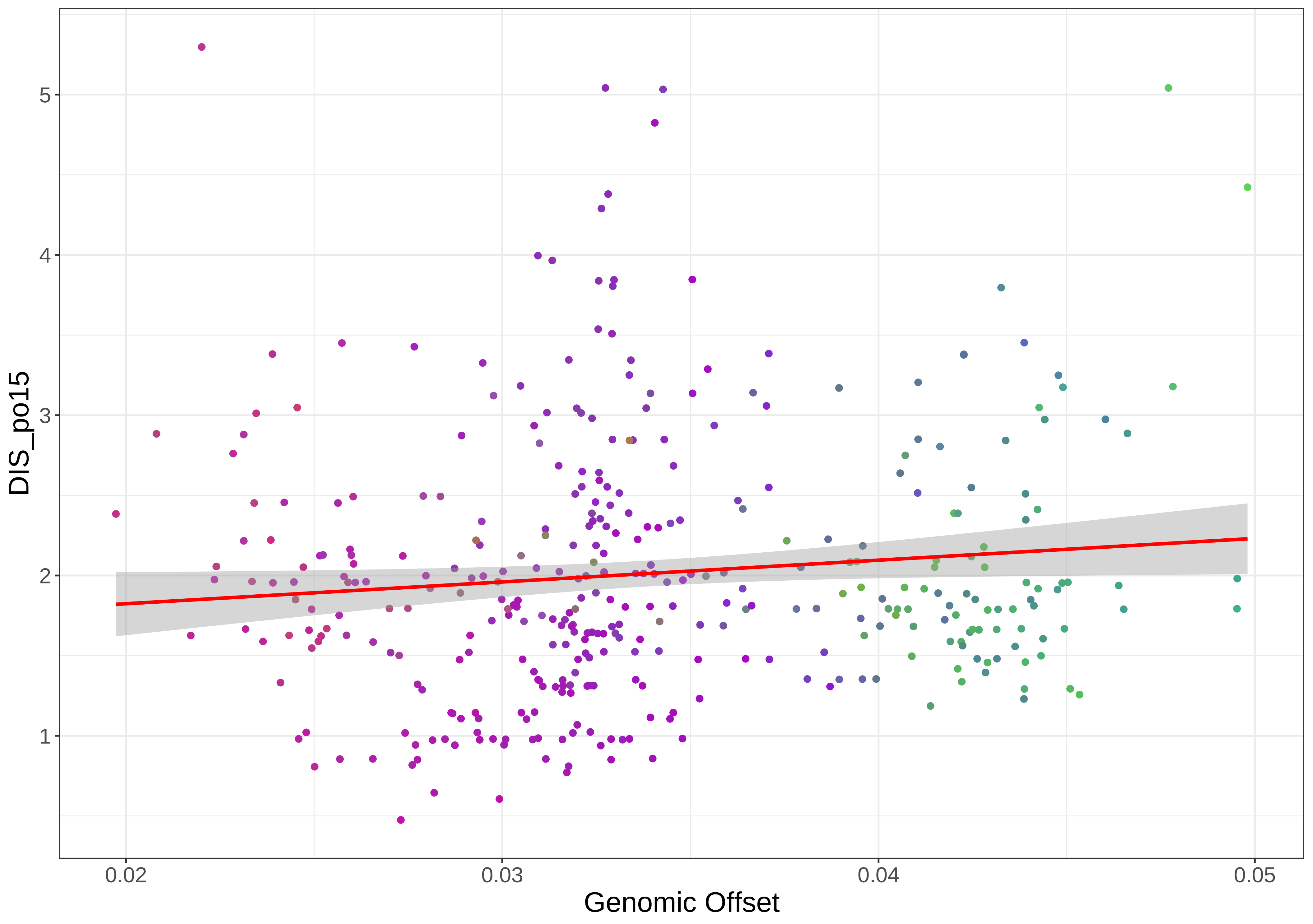

DIS_po15
Genomic Offset
5
4
3
2
1
0.02
0.03
0.04
0.05

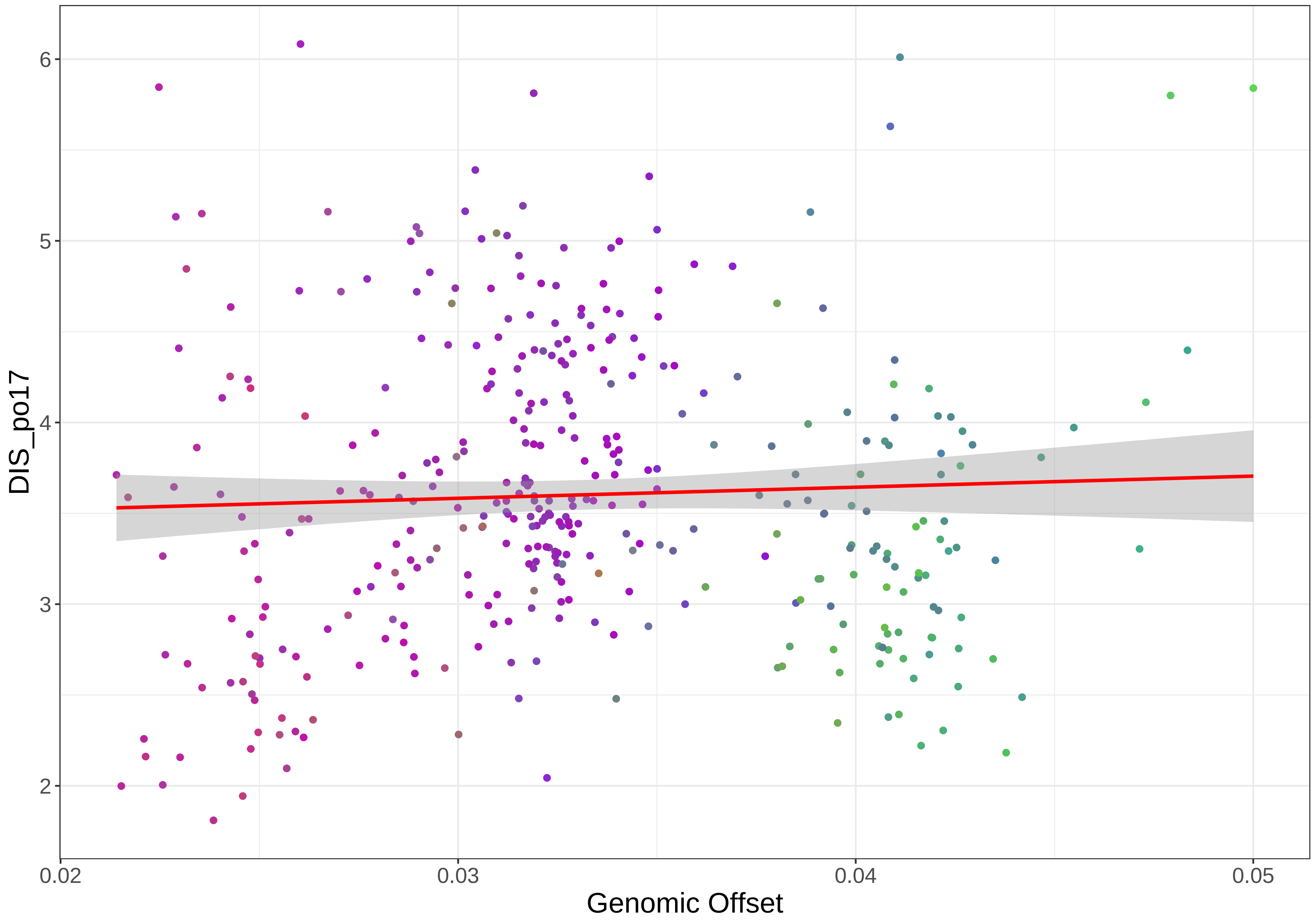

6
5
4
3
2
DIS_po17
0.02
0.03
0.04
0.05
Genomic Offset

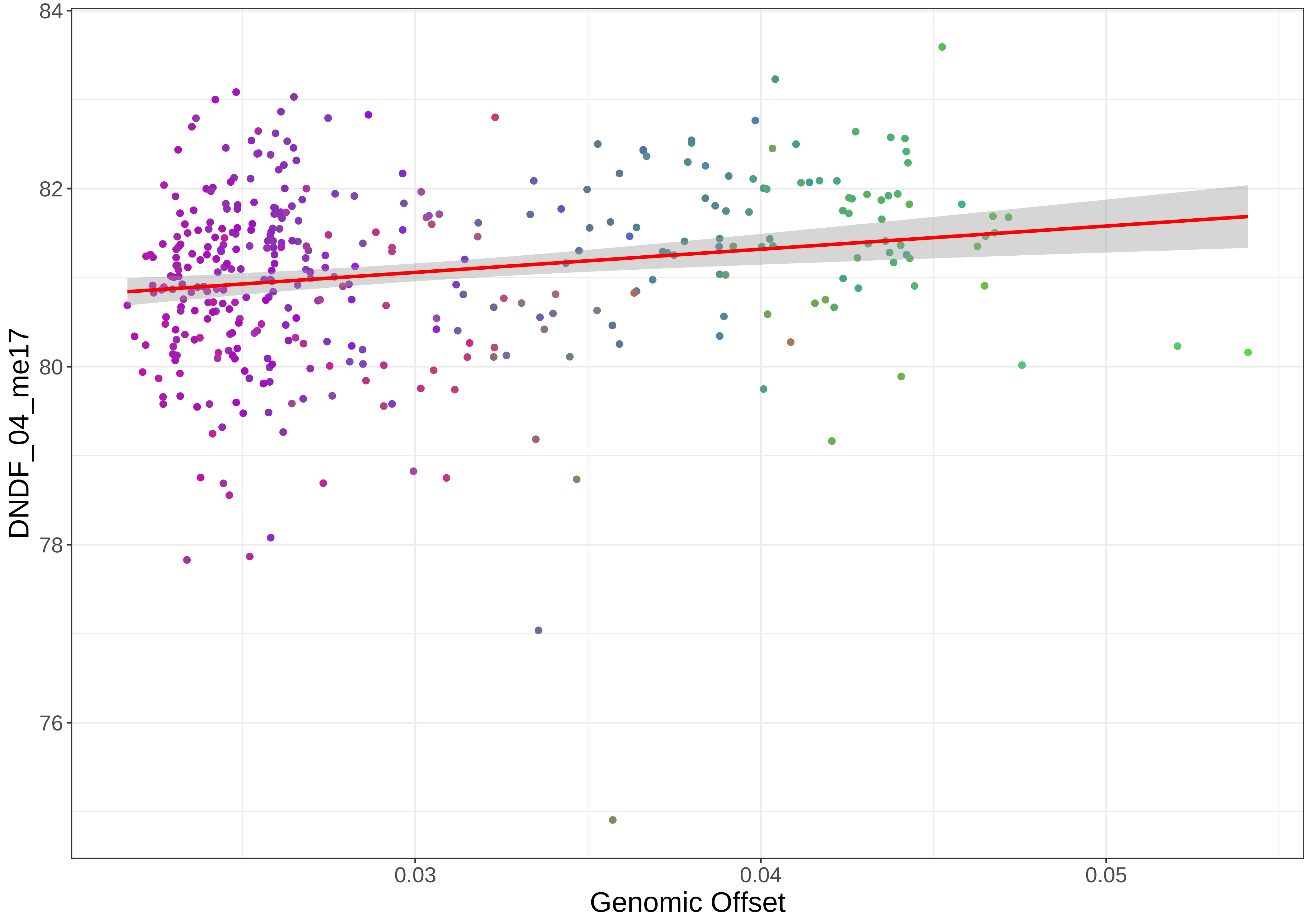

84
82
80
78
76
DNDF_04_me17
0.03
0.04
0.05
Genomic Offset

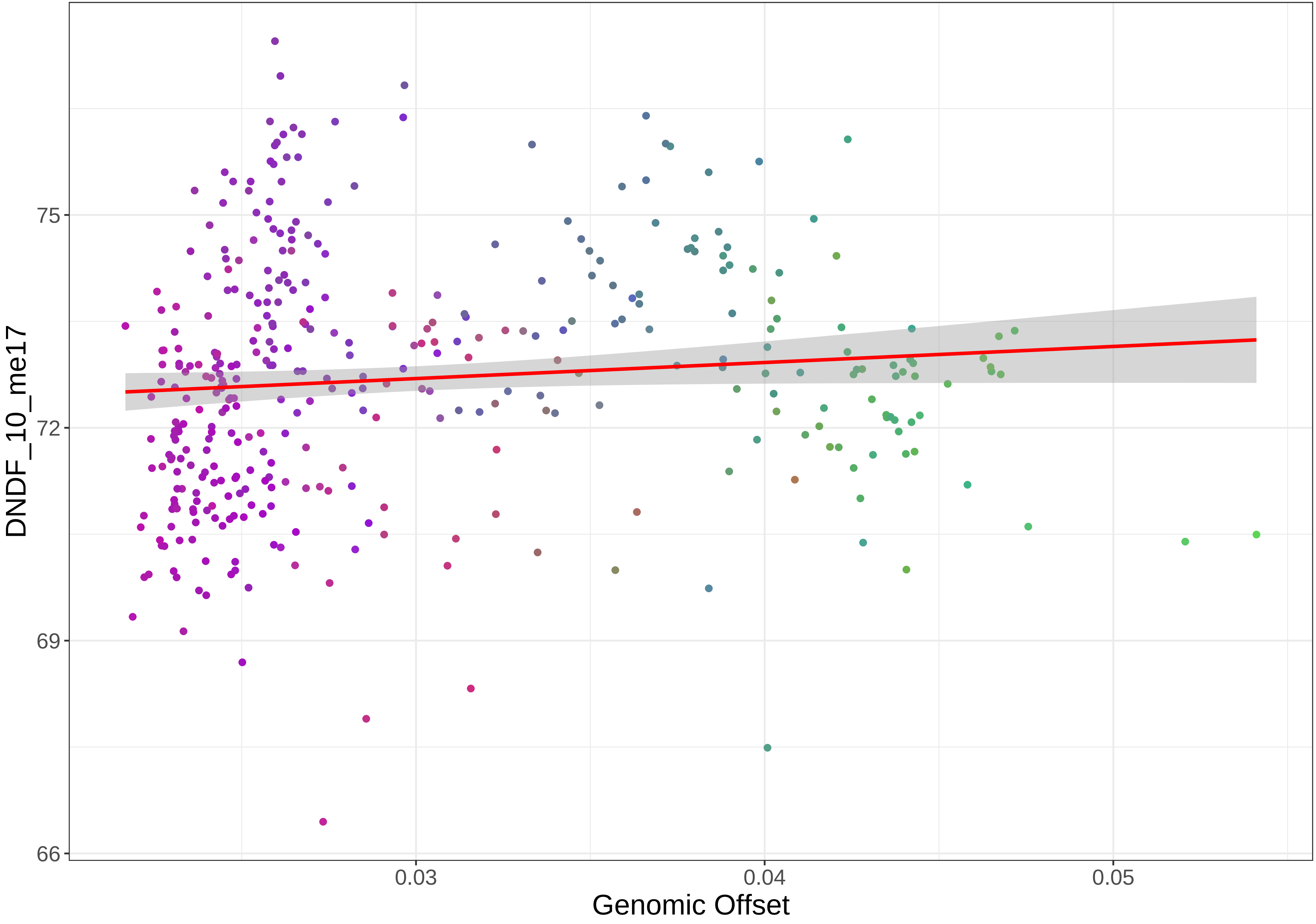

DNDF_10_me17
75
72
69
66
0.03
0.04
0.05
Genomic Offset

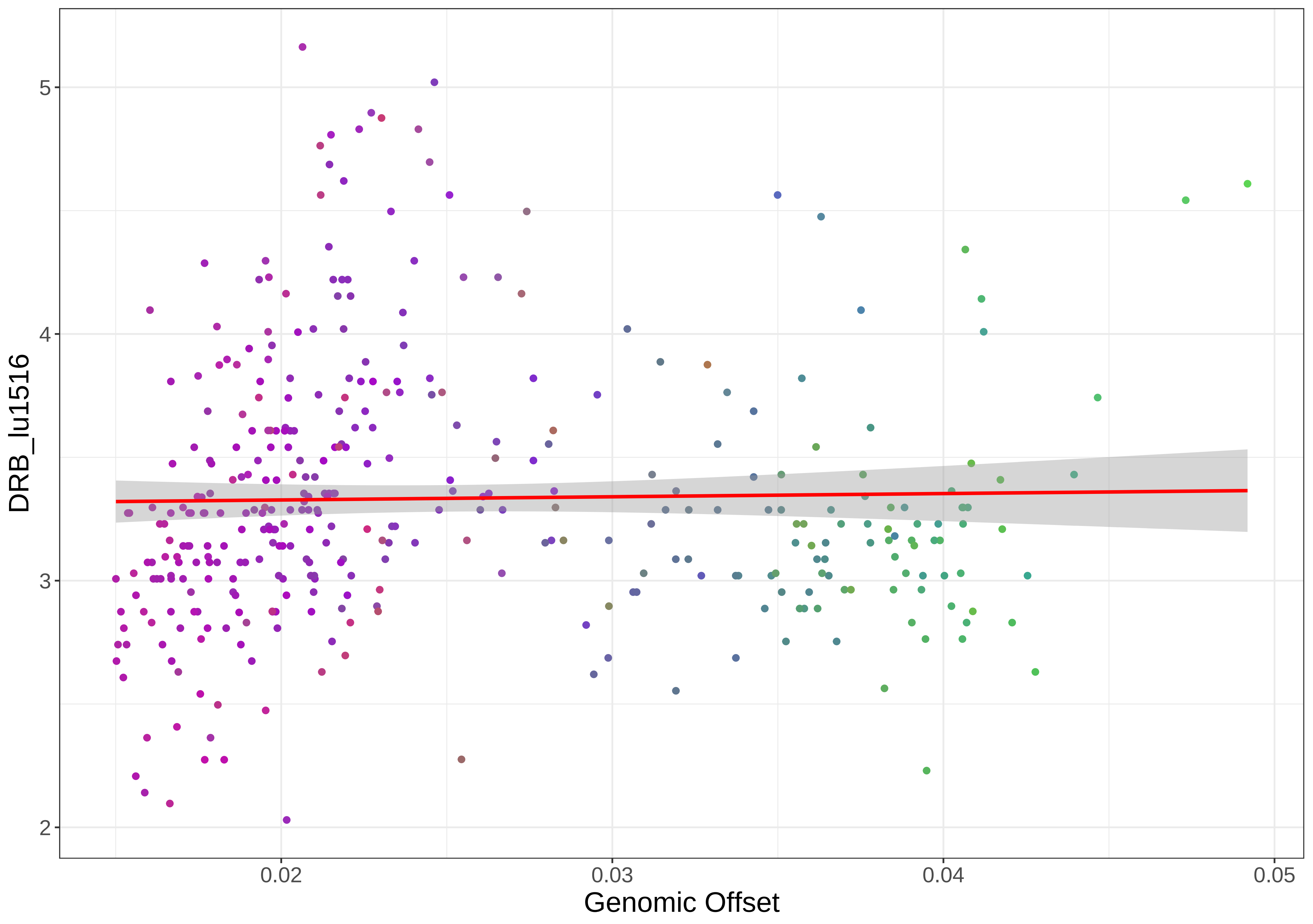

DRB_lu1516
5
4
3
2
0.02
0.03
0.04
0.05
Genomic Offset

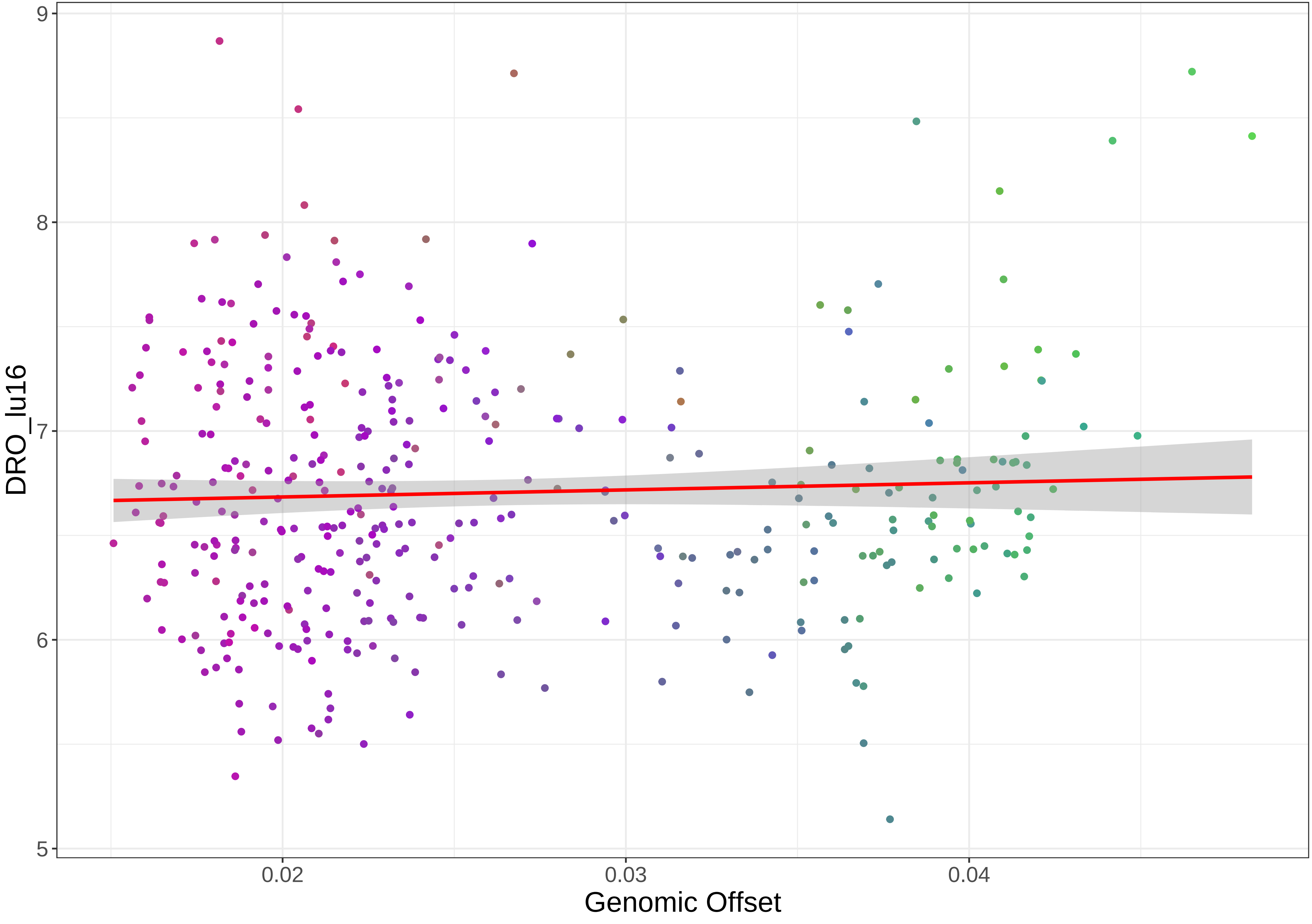

DRO_lu16
9
8
7
6
5
0.02
0.03
0.04
Genomic Offset

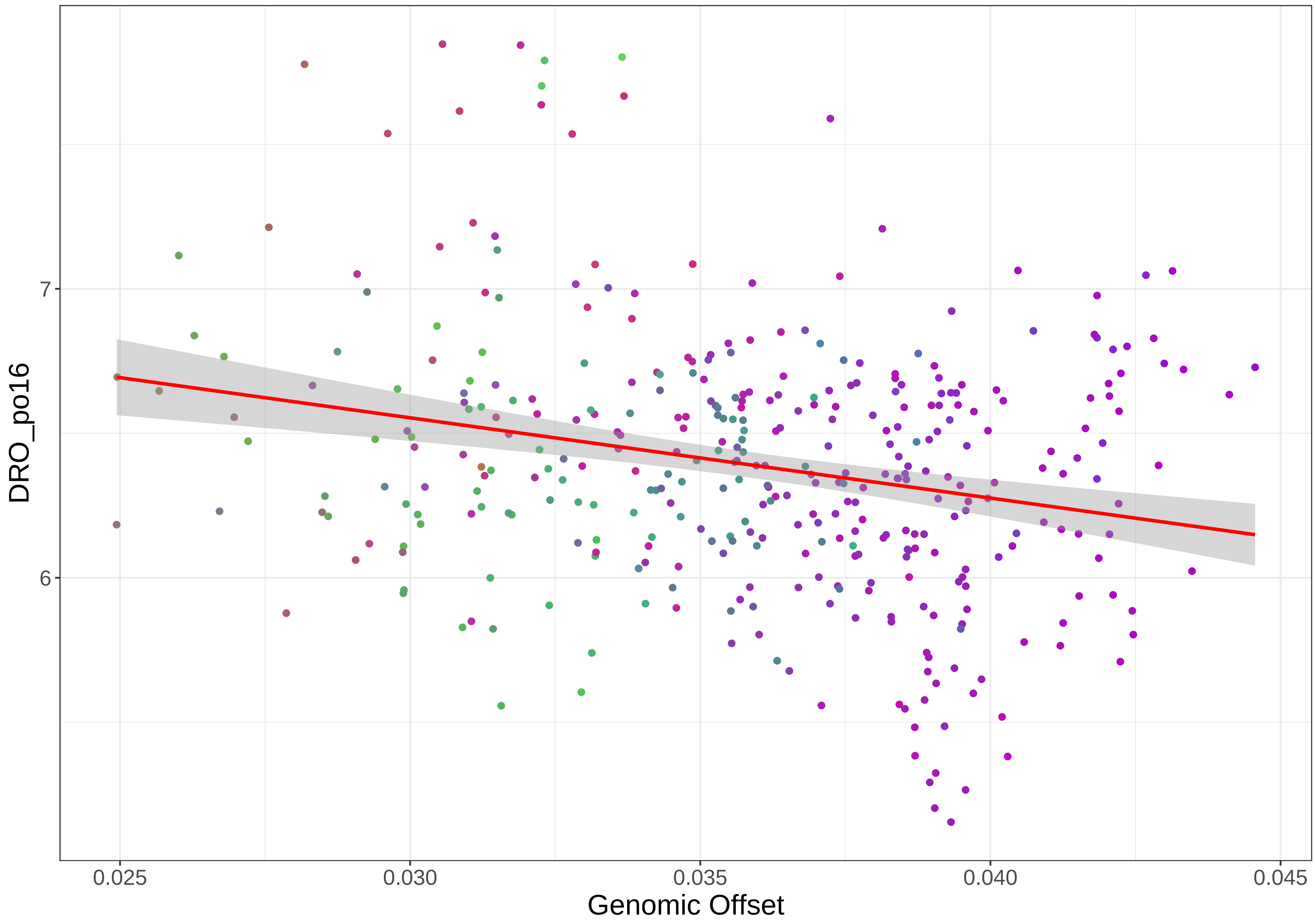
7
6
DRO_po16
0.025
0.030
0.035
0.040
0.045
Genomic Offset

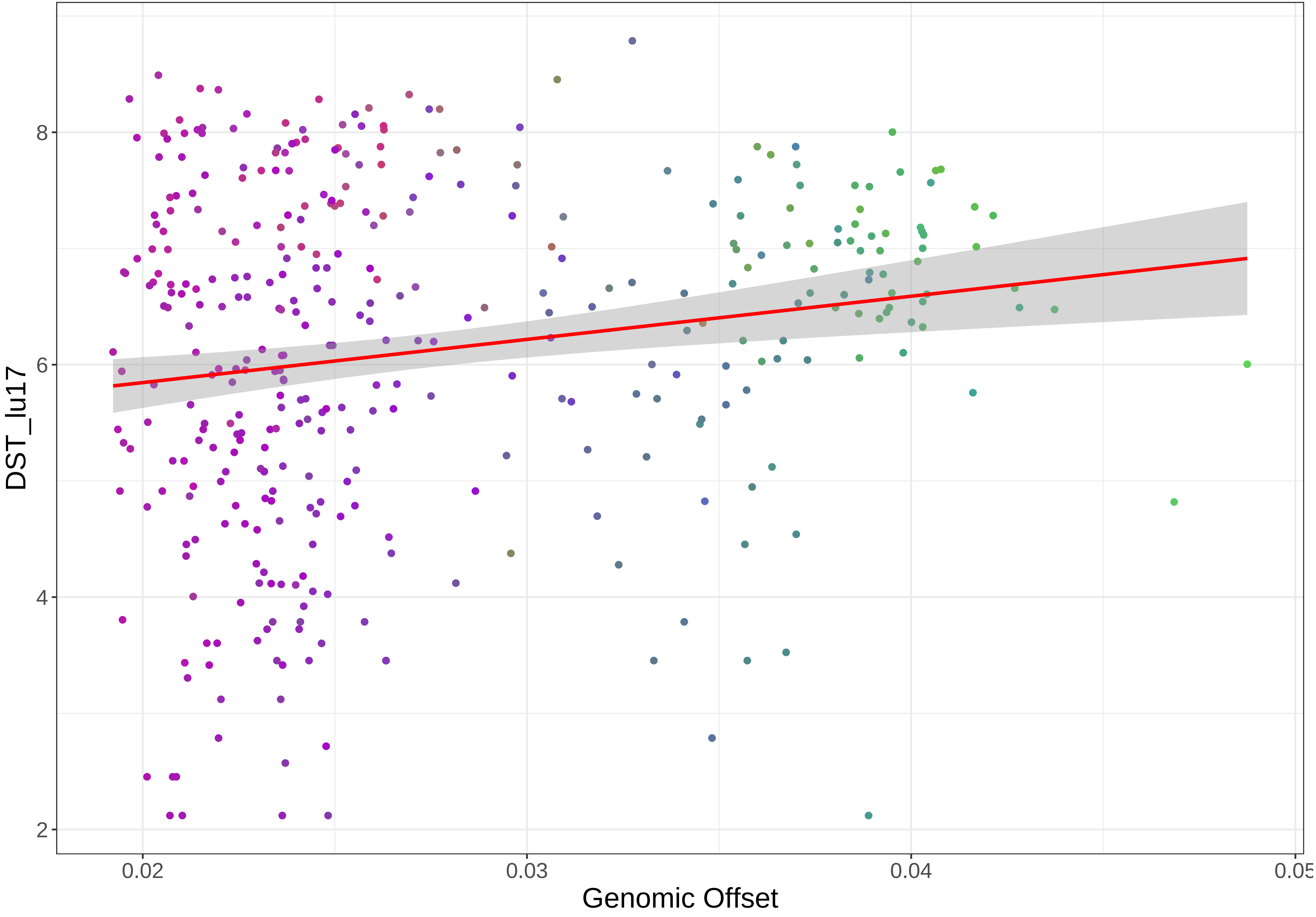

DST_lu17
8
6
4
2
0.02
0.03
0.04
0.05
Genomic Offset

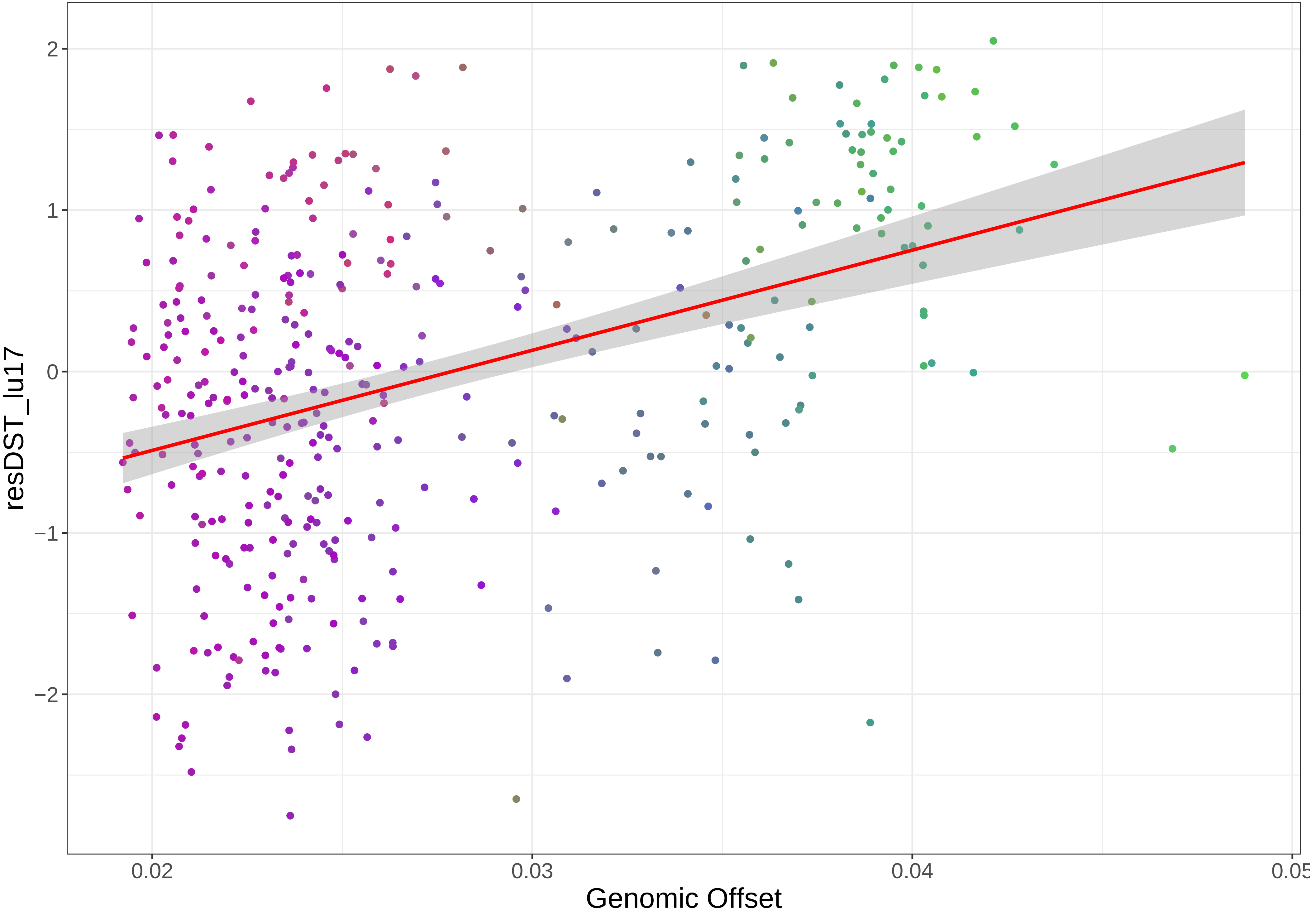

2
1
0
−1
−2
resDST_lu17
0.02
0.03
0.04
0.05
Genomic Offset

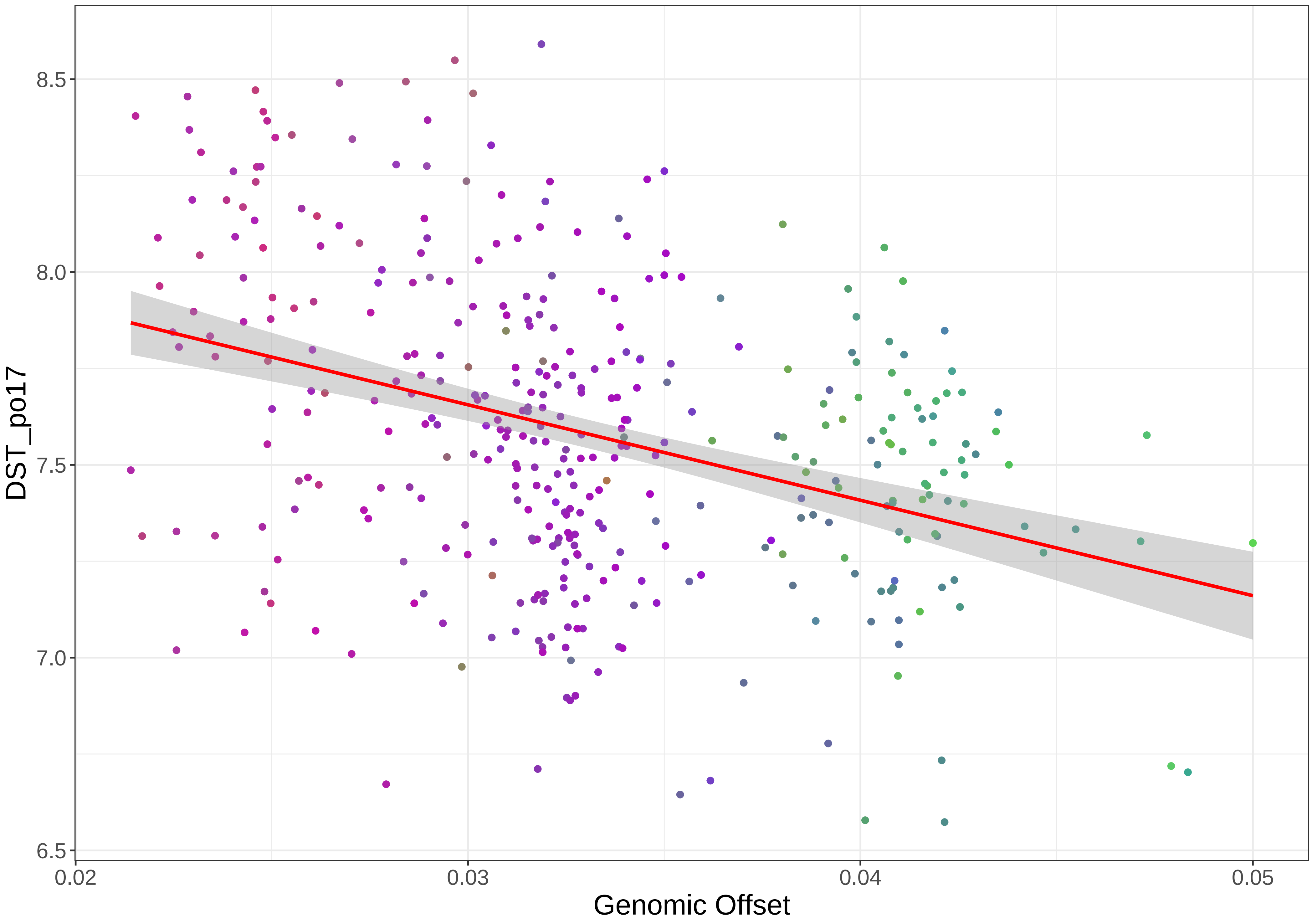

8.5
8.0
7.5
7.0
6.5
DST_po17
0.02
0.03
0.04
0.05
Genomic Offset

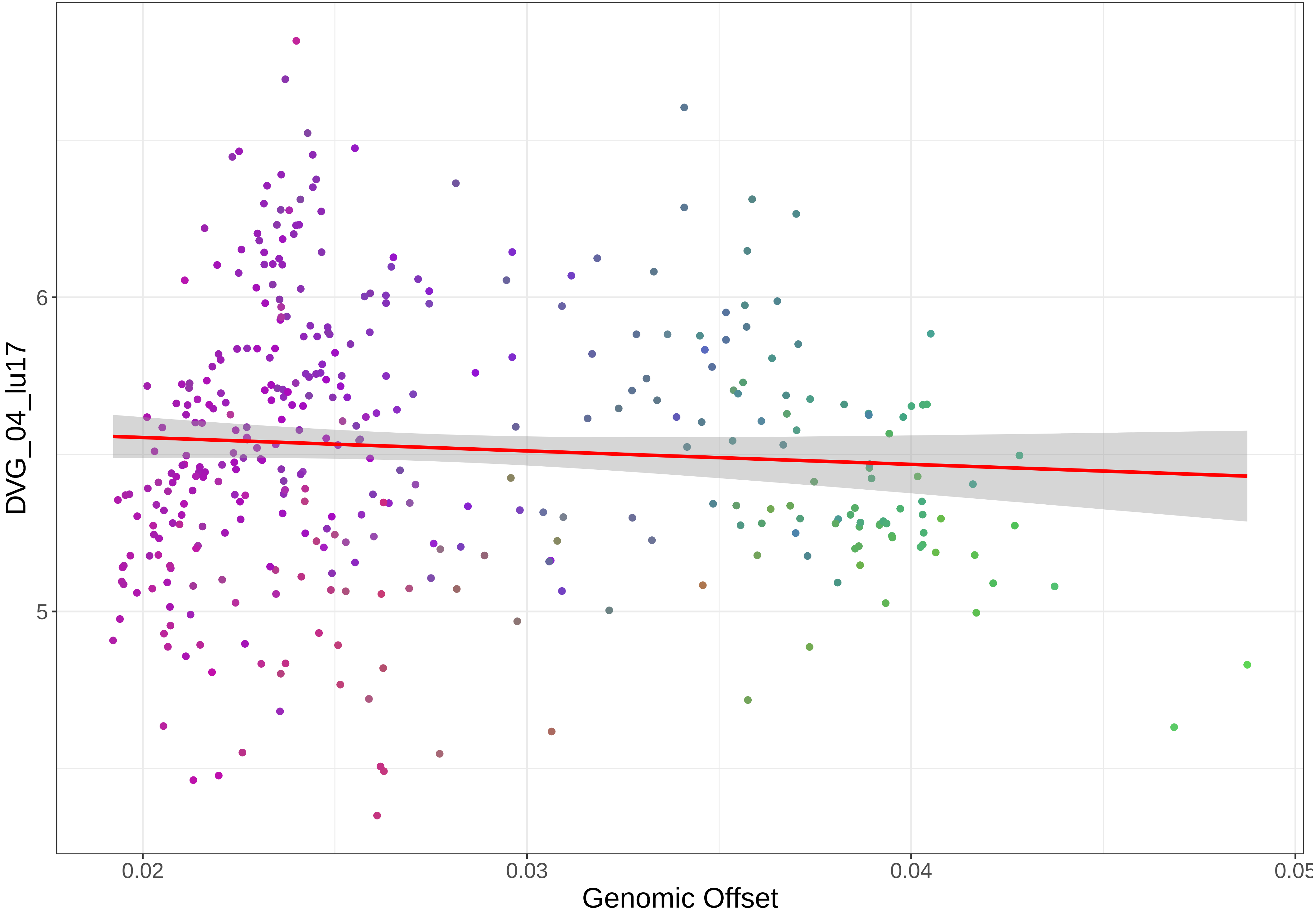

DVG_04_lu17
6
5
0.02
0.03
0.04
0.05
Genomic Offset

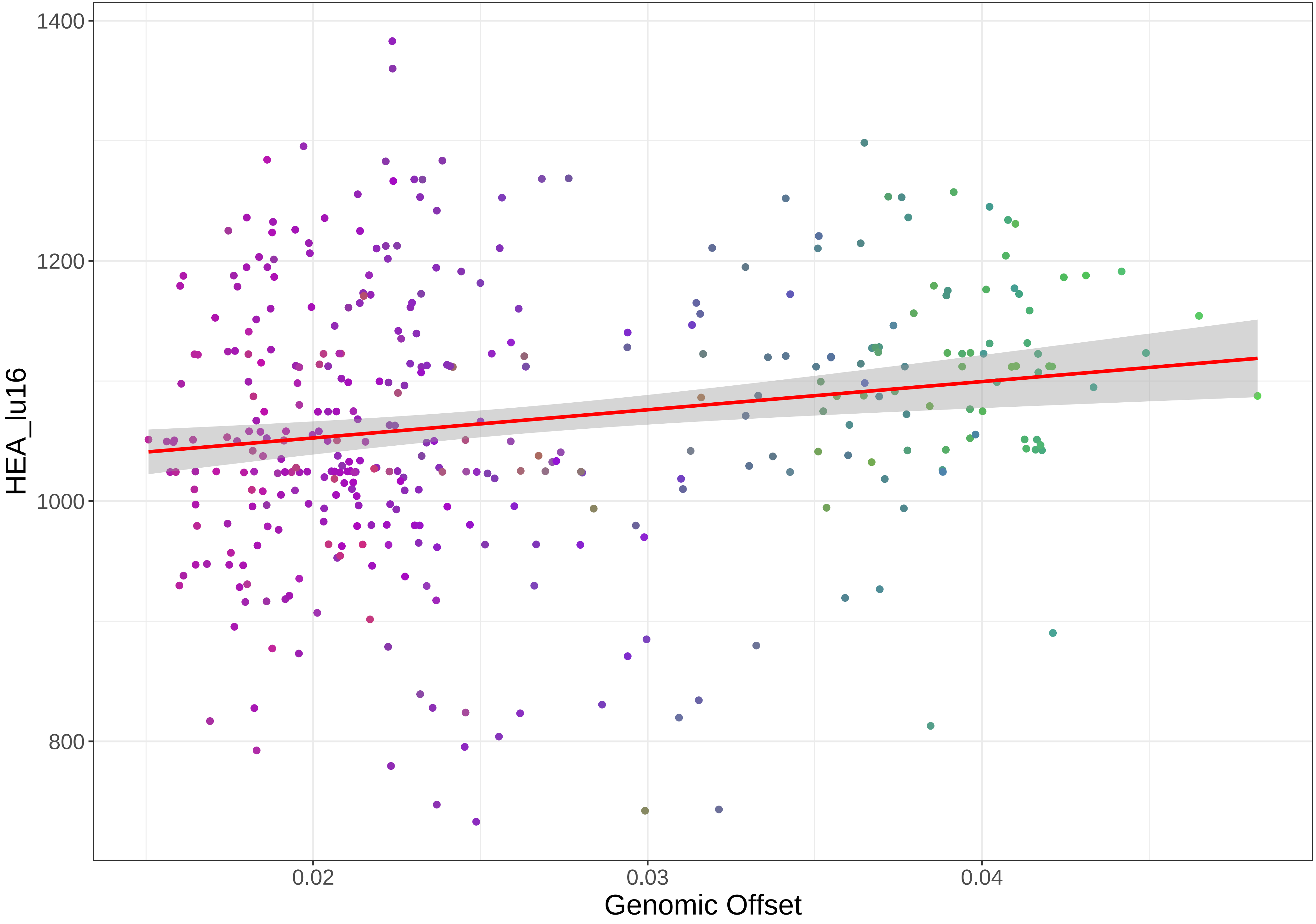

1400
1200
1000
800
HEA_lu16
0.02
0.03
0.04
Genomic Offset

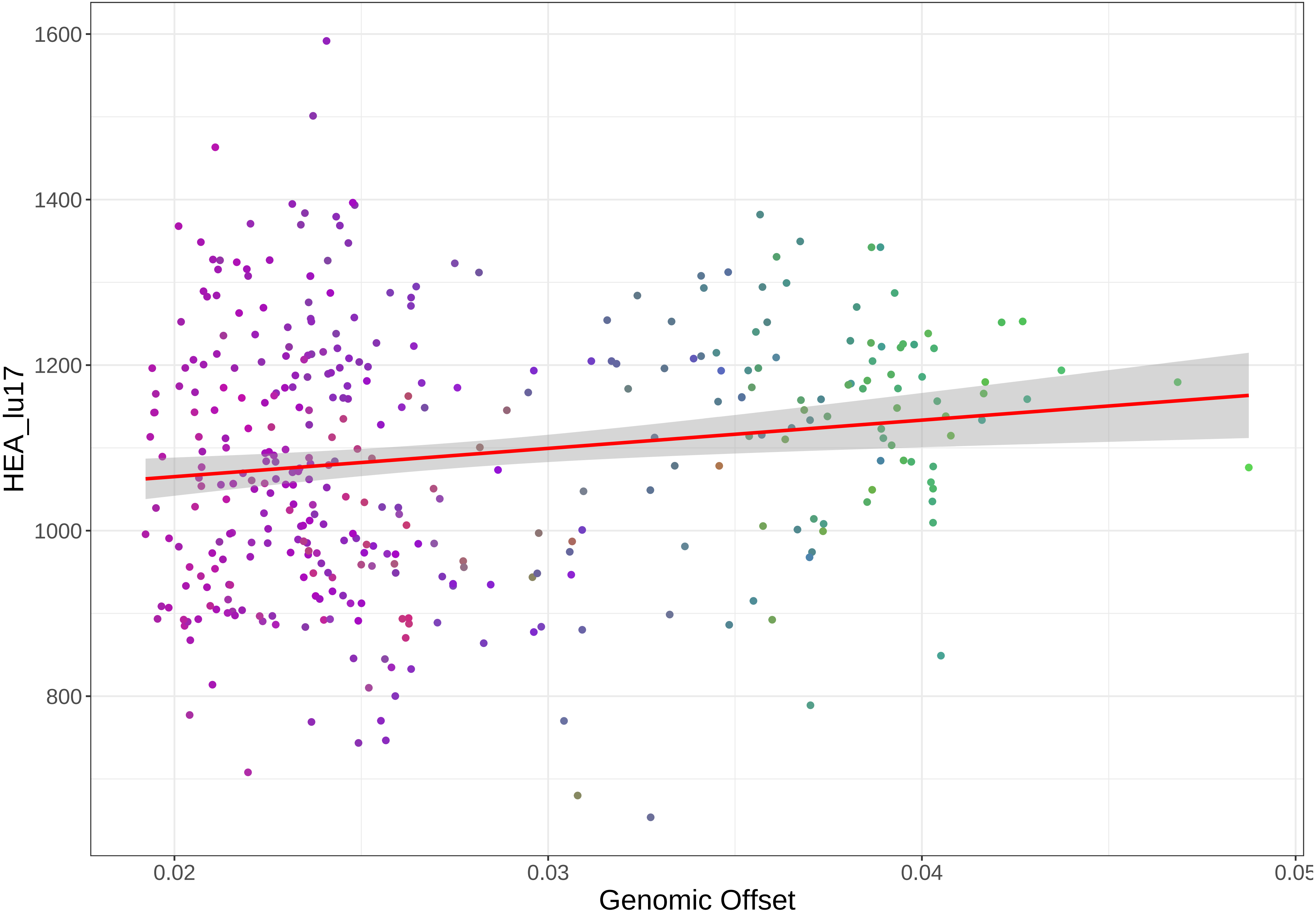
HEA_lu17
1600
1400
1200
1000
800
0.02
0.03
0.04
0.05
Genomic Offset

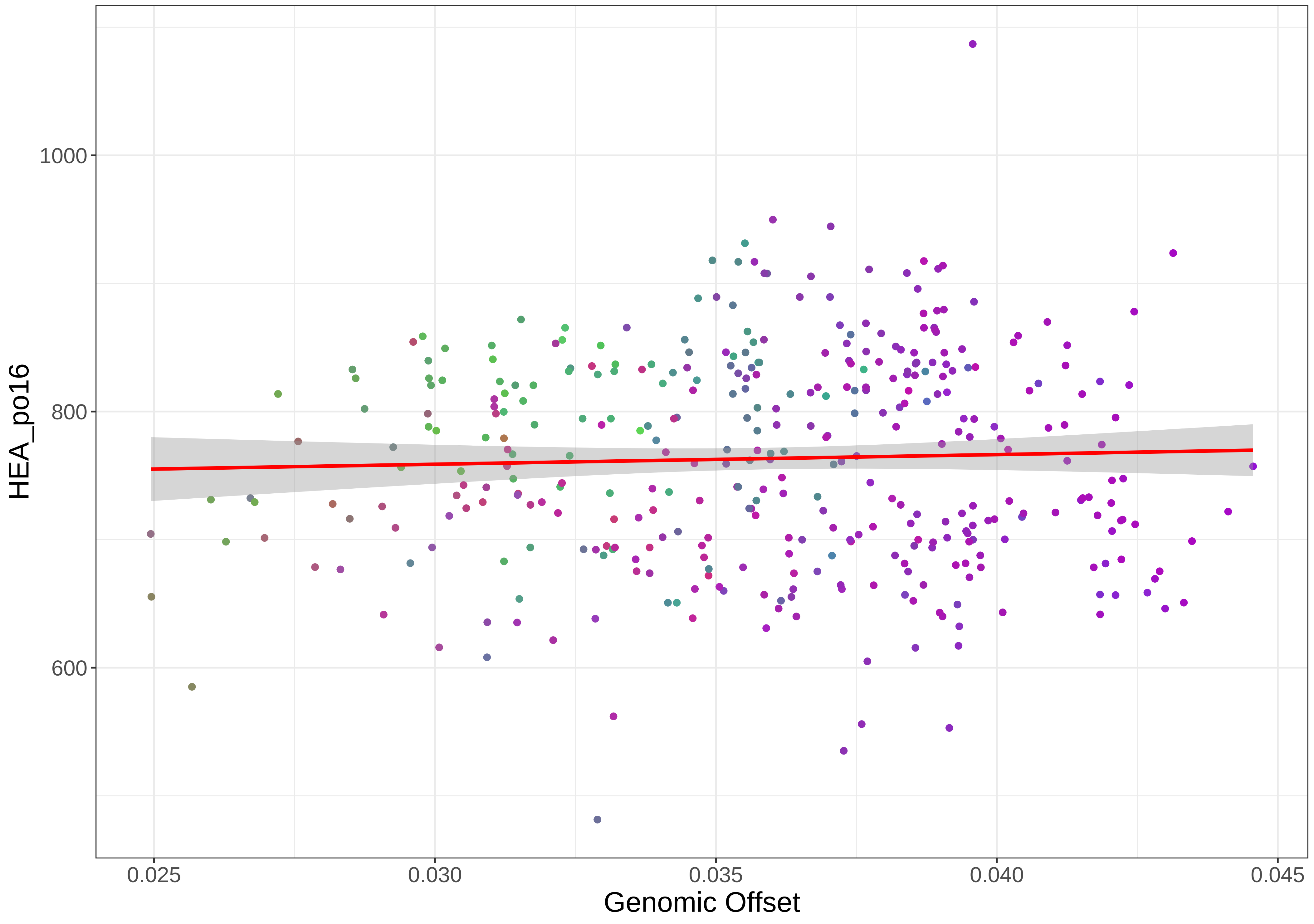
1000
800
600
HEA_po16
0.025
0.030
0.035
0.040
0.045
Genomic Offset

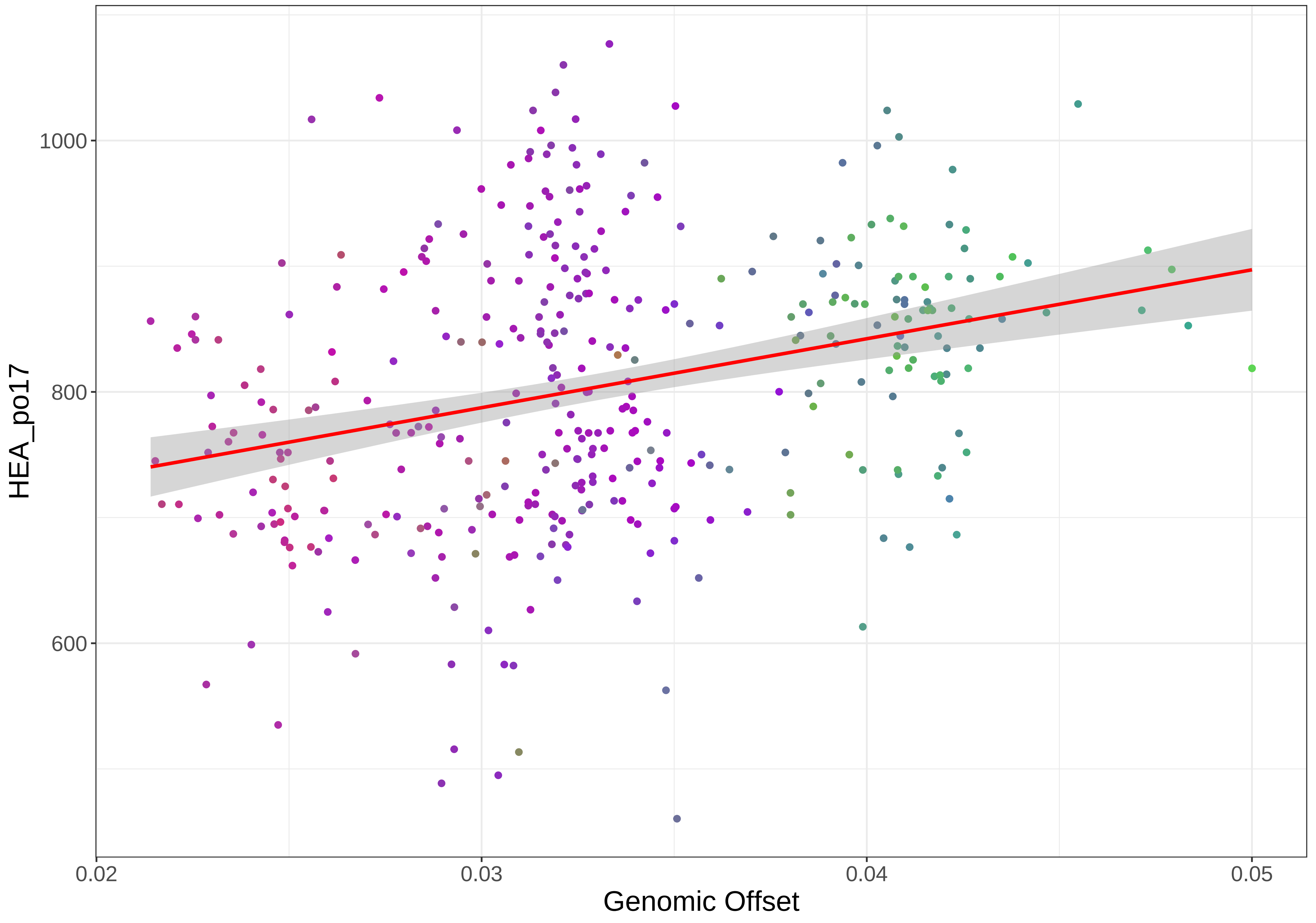
HEA_po17
1000
800
600
0.02
0.03
0.04
0.05
Genomic Offset

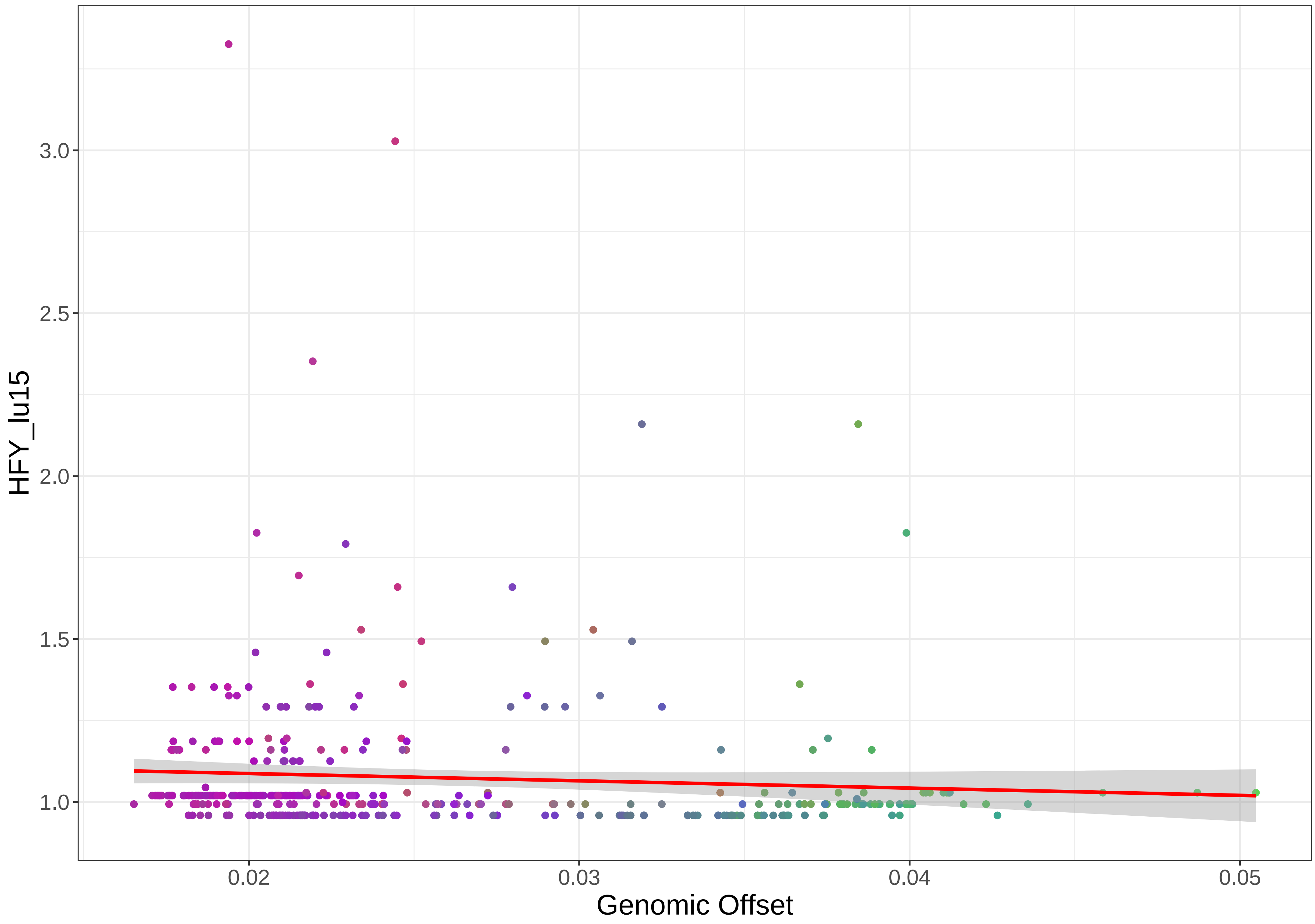

HFY_lu15
3.0
2.5
2.0
1.5
1.0
0.02
0.03
0.04
0.05
Genomic Offset

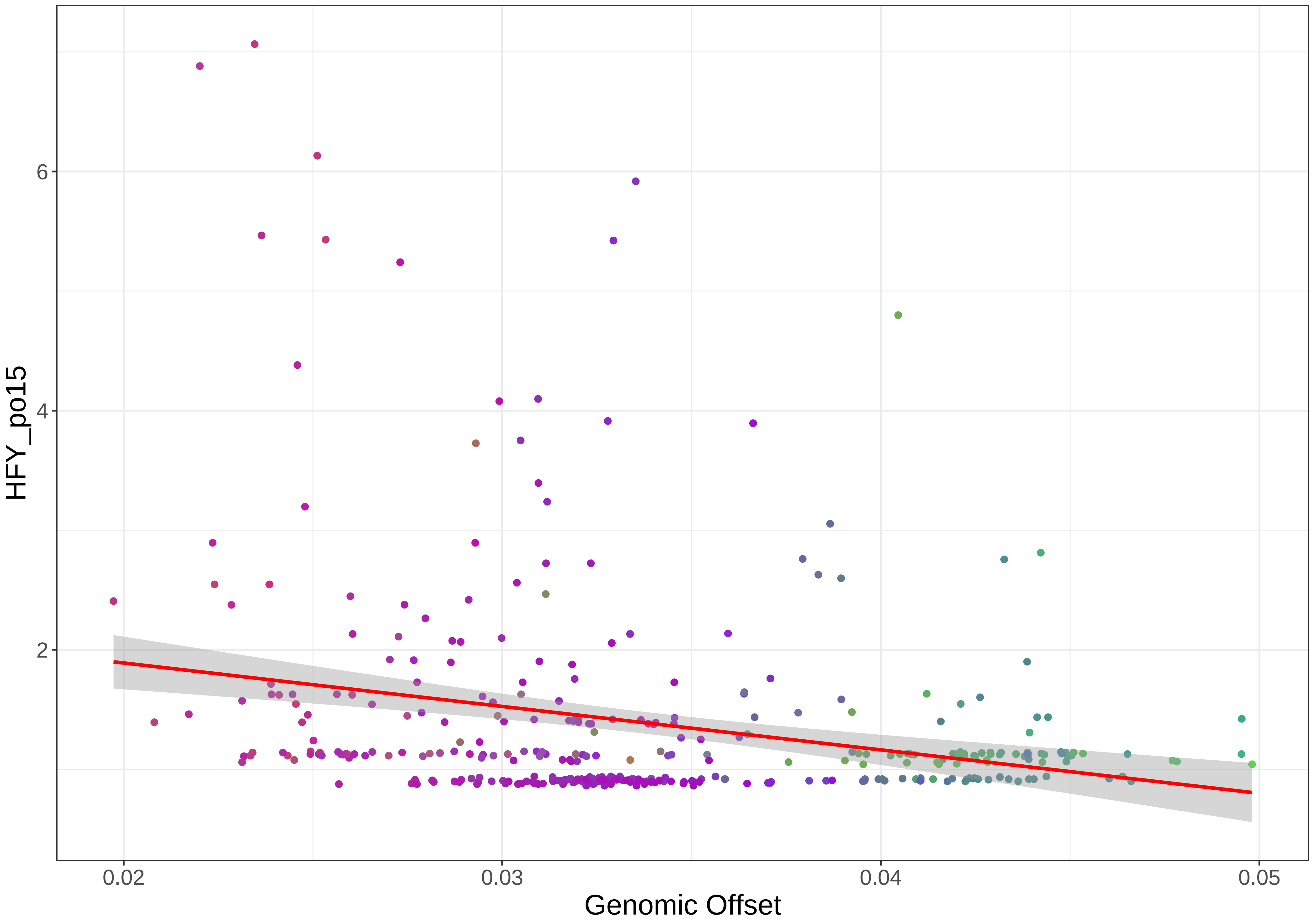

HFY_po15
6
4
2
0.02
0.03
0.04
0.05
Genomic Offset

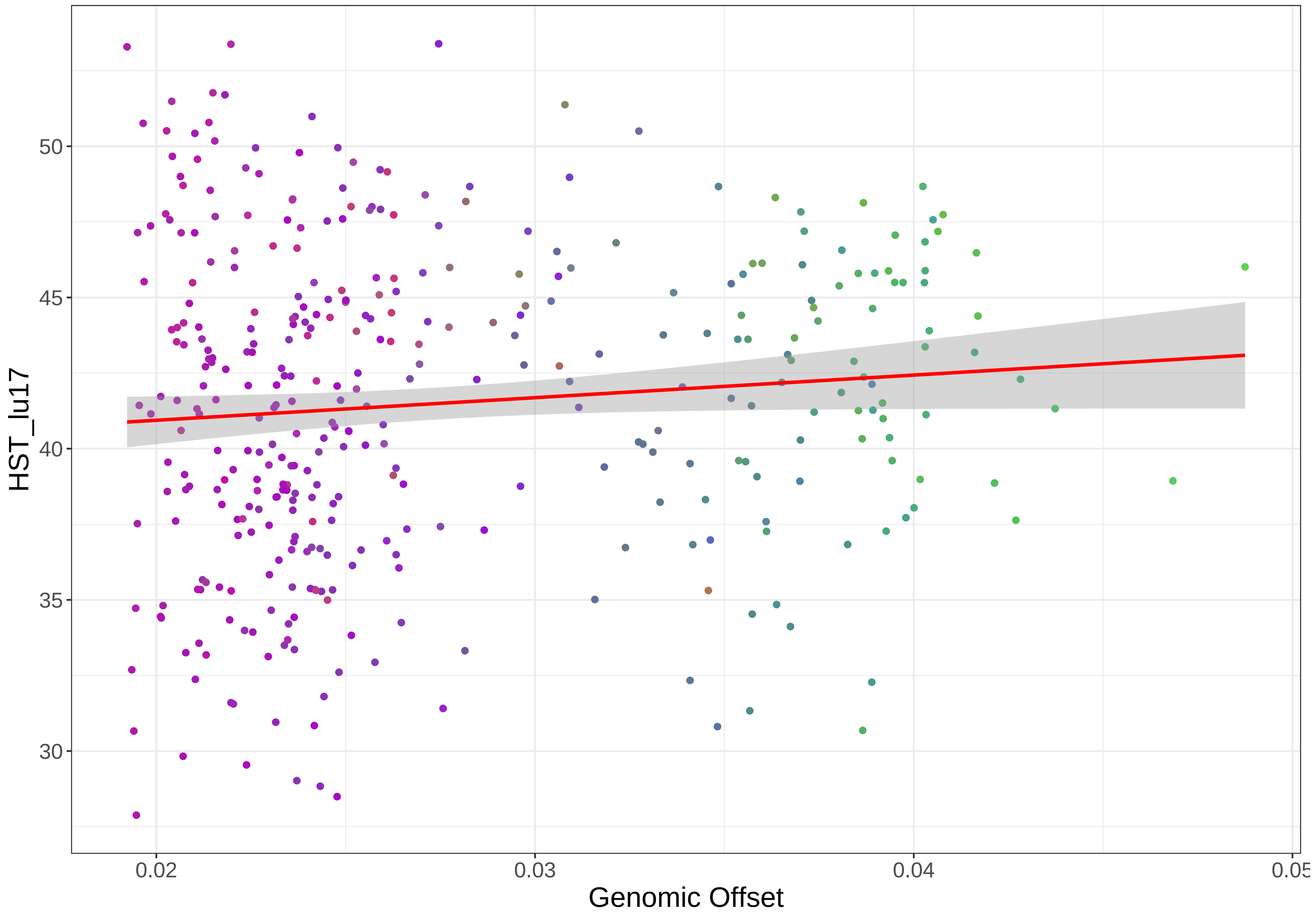
50
45
40
35
30
HST_lu17
0.02
0.03
0.04
0.05
Genomic Offset

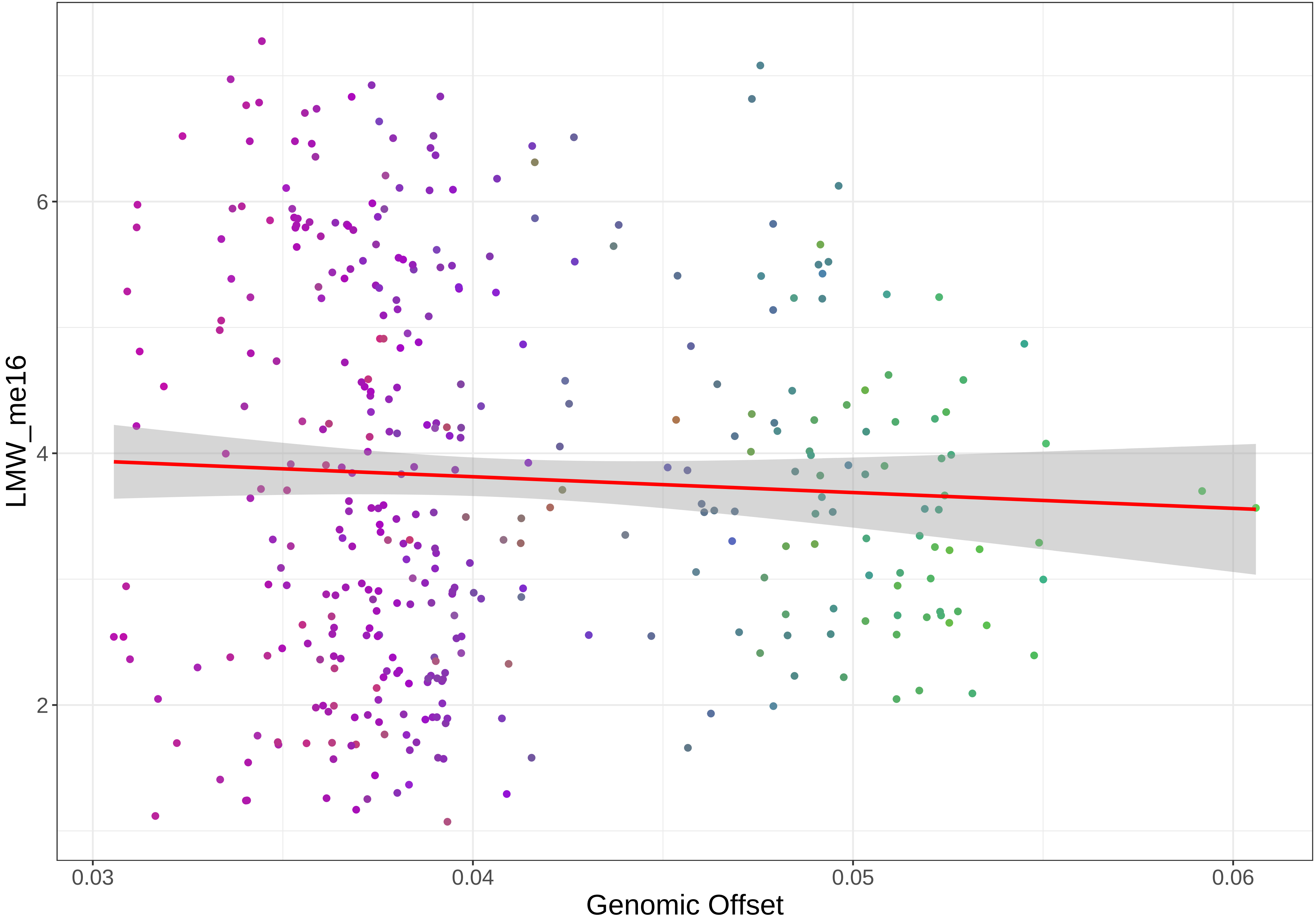

LMW_me16
6
4
2
0.03
0.04
0.05
0.06
Genomic Offset

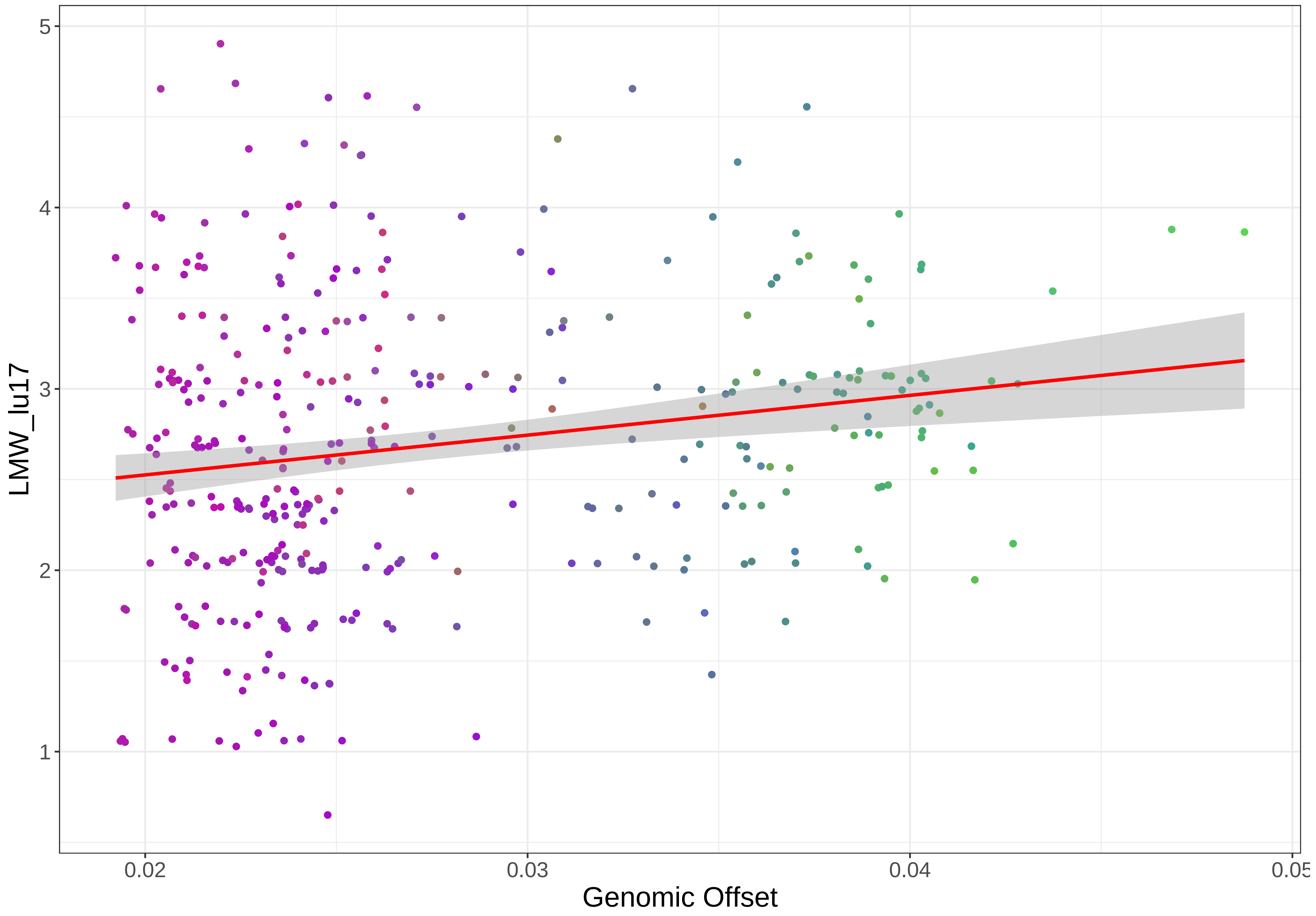

LMW_lu17
Genomic Offset
5
4
3
2
1
0.02
0.03
0.04
0.05

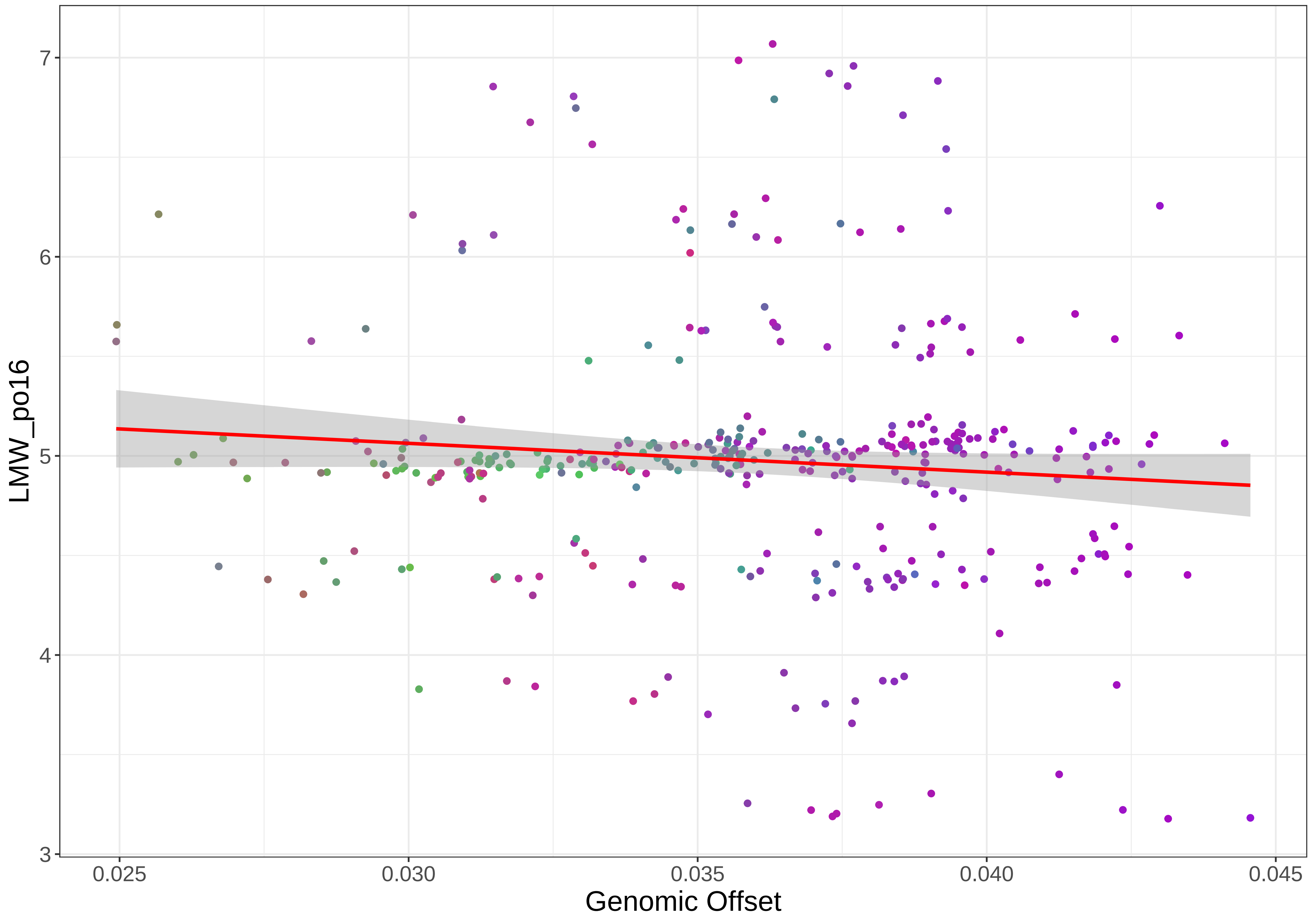

7
6
5
4
3
LMW_po16
0.025
0.030
0.035
0.040
0.045
Genomic Offset

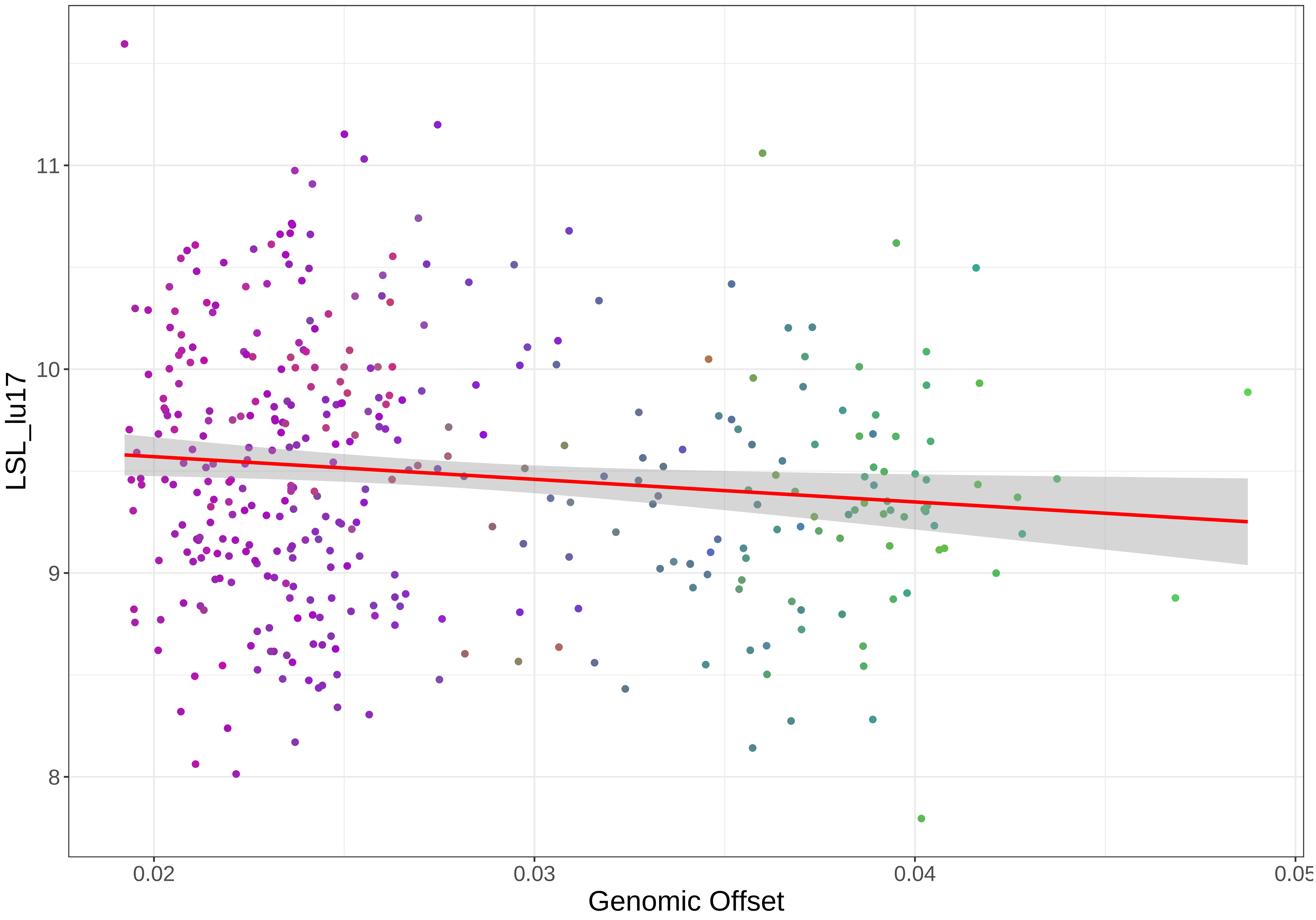
LSL_lu17
11
10
9
8
0.02
0.03
0.04
0.05
Genomic Offset

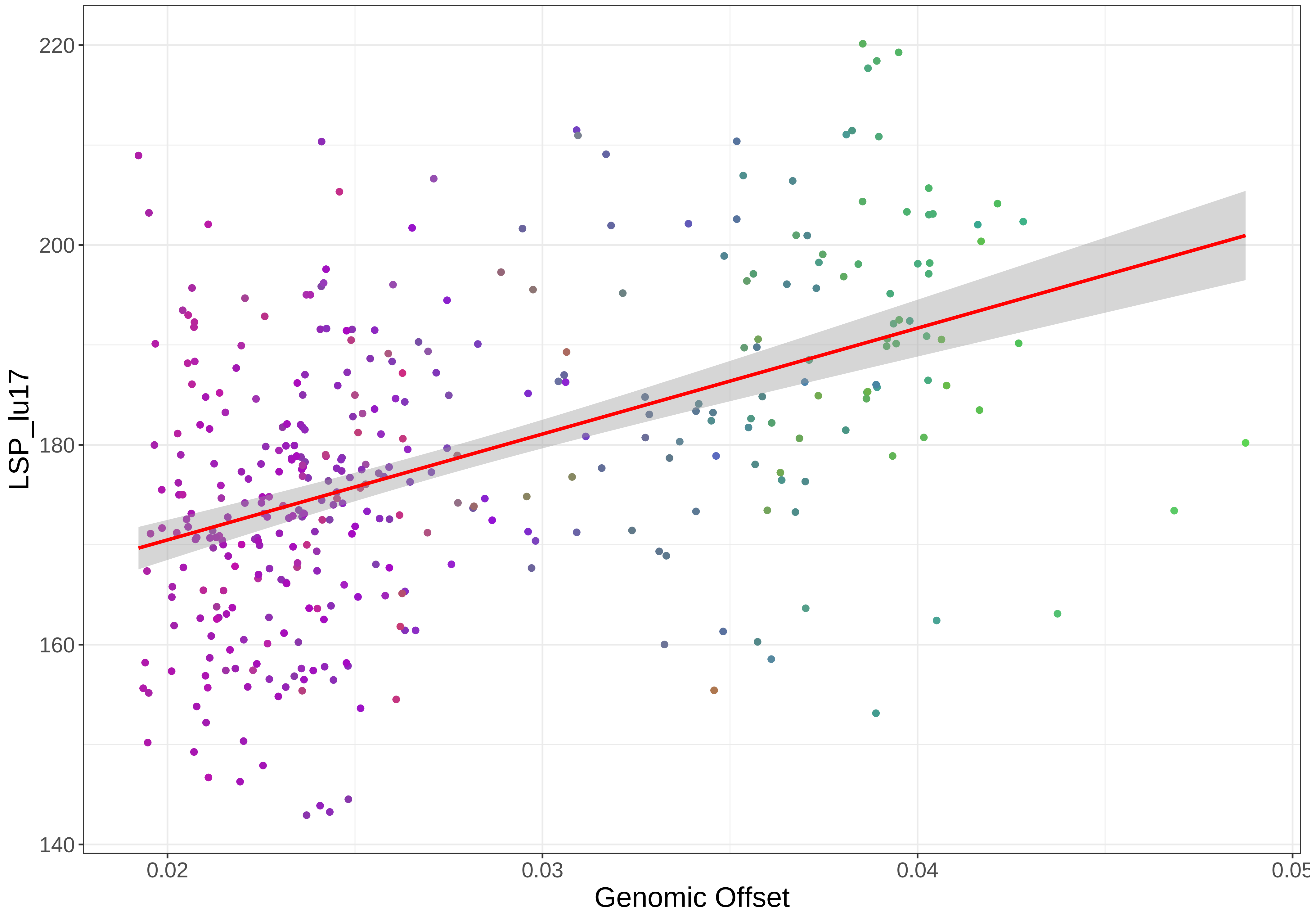

220
200
180
160
140
LSP_lu17
0.02
0.03
0.04
0.05
Genomic Offset

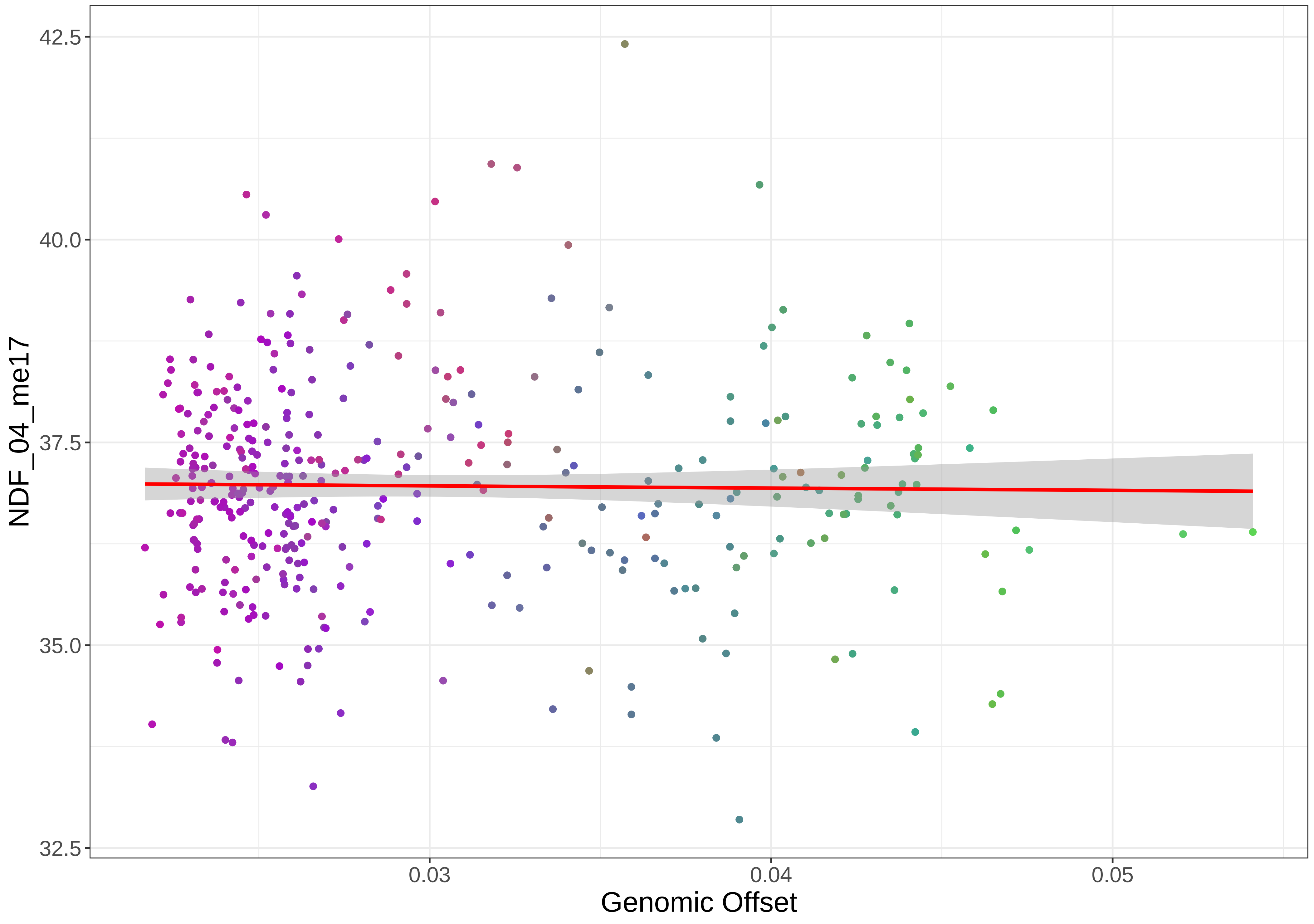

42.5
40.0
37.5
35.0
32.5
NDF_04_me17
0.03
0.04
0.05
Genomic Offset

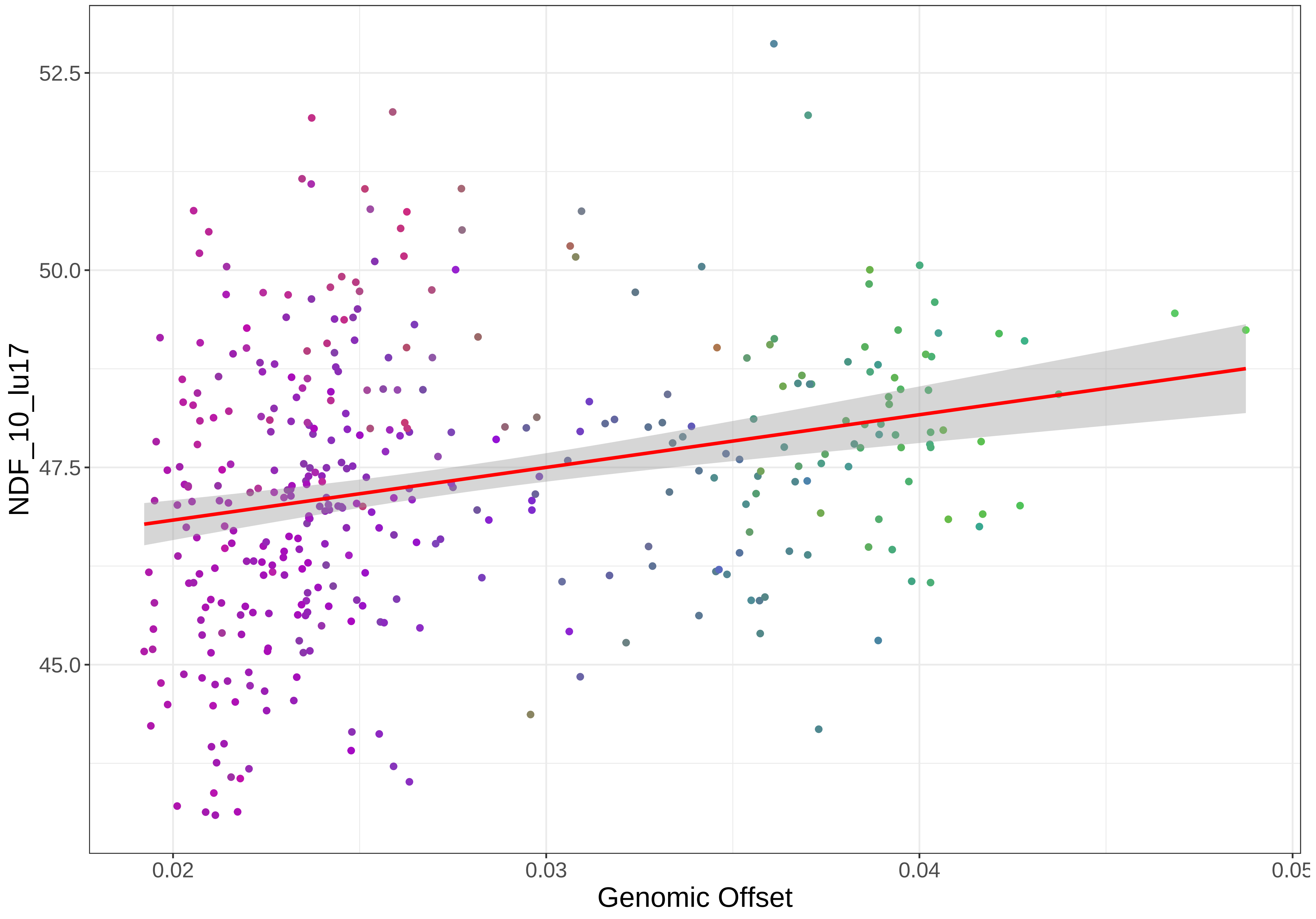
52.5
50.0
47.5
45.0
NDF_10_lu17
0.02
0.03
0.04
0.05
Genomic Offset

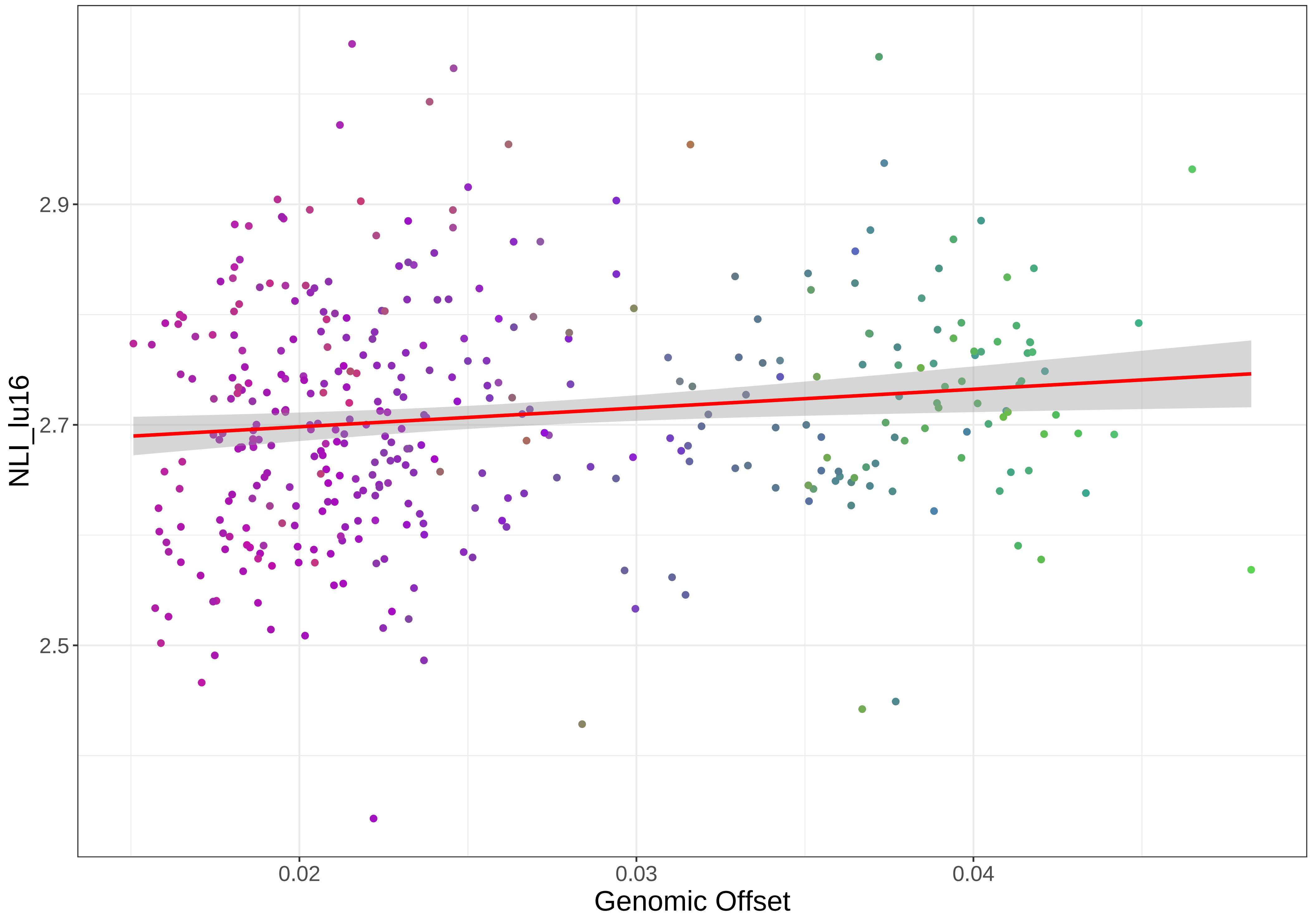

NLI_lu16
2.9
2.7
2.5
0.02
0.03
0.04
Genomic Offset

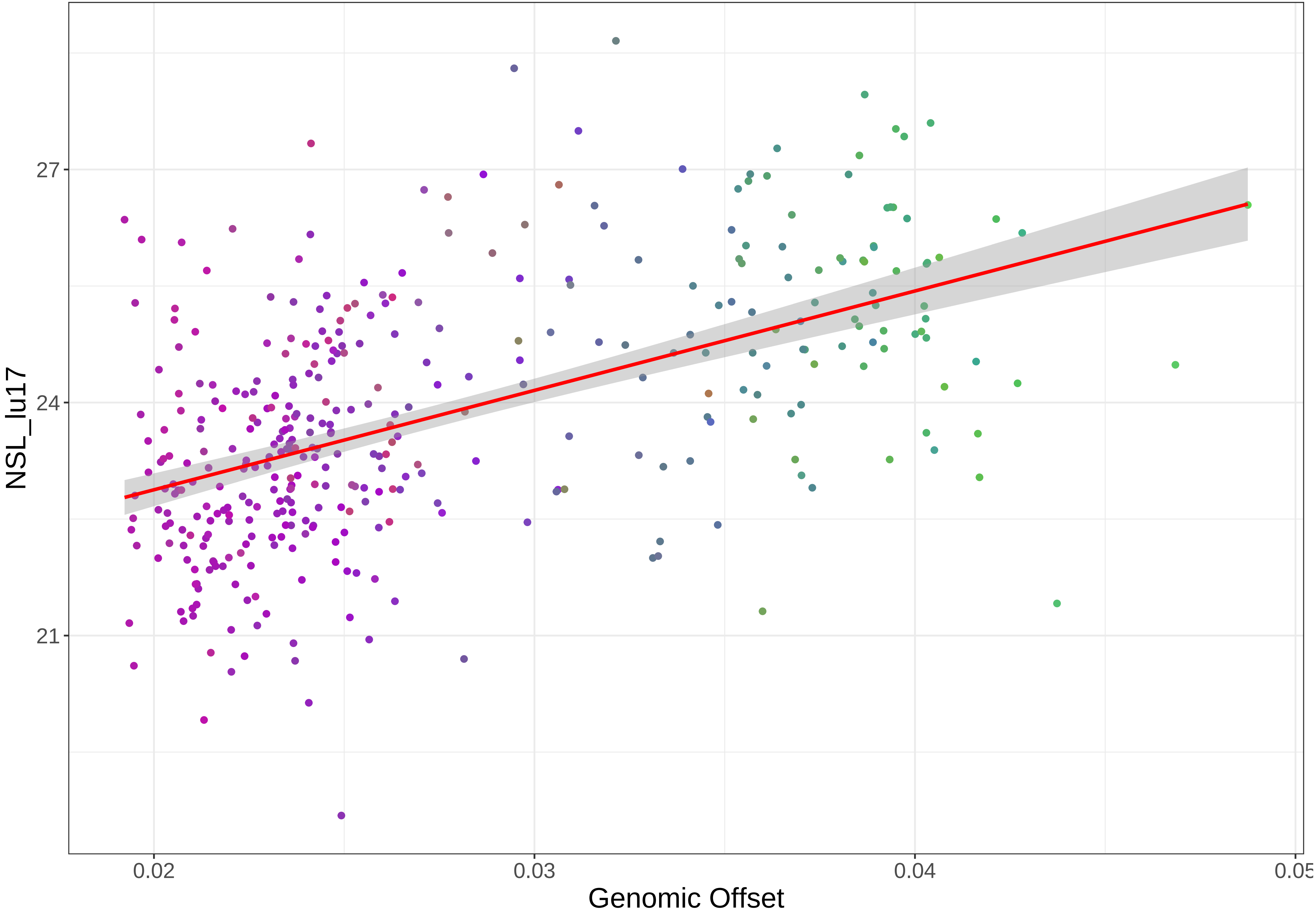

27
24
21
NSL_lu17
0.02
0.03
0.04
0.05
Genomic Offset

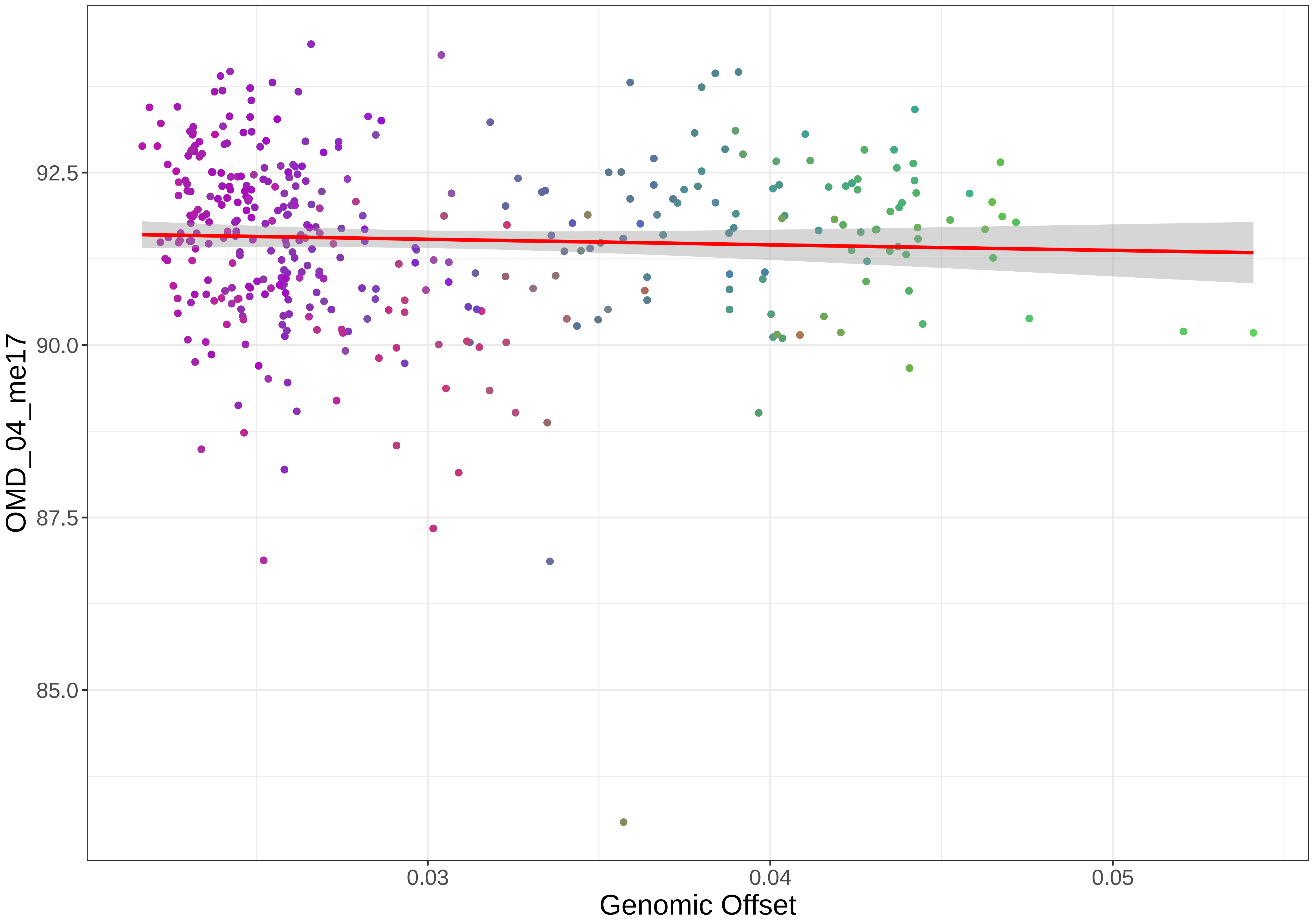

OMD_04_me17
92.5
90.0
87.5
85.0
0.03
0.04
0.05
Genomic Offset

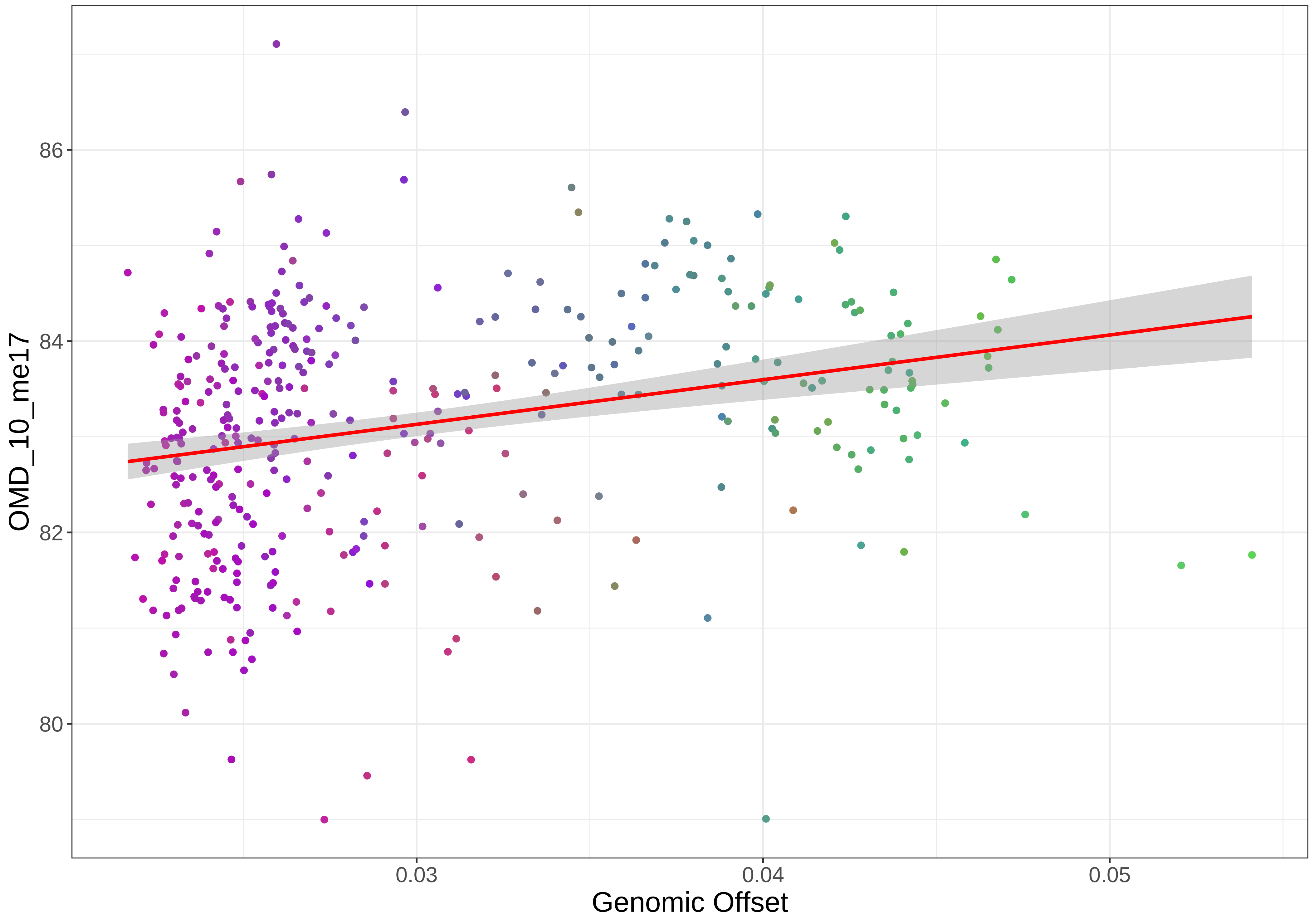

86
84
82
80
OMD_10_me17
0.03
0.04
0.05
Genomic Offset

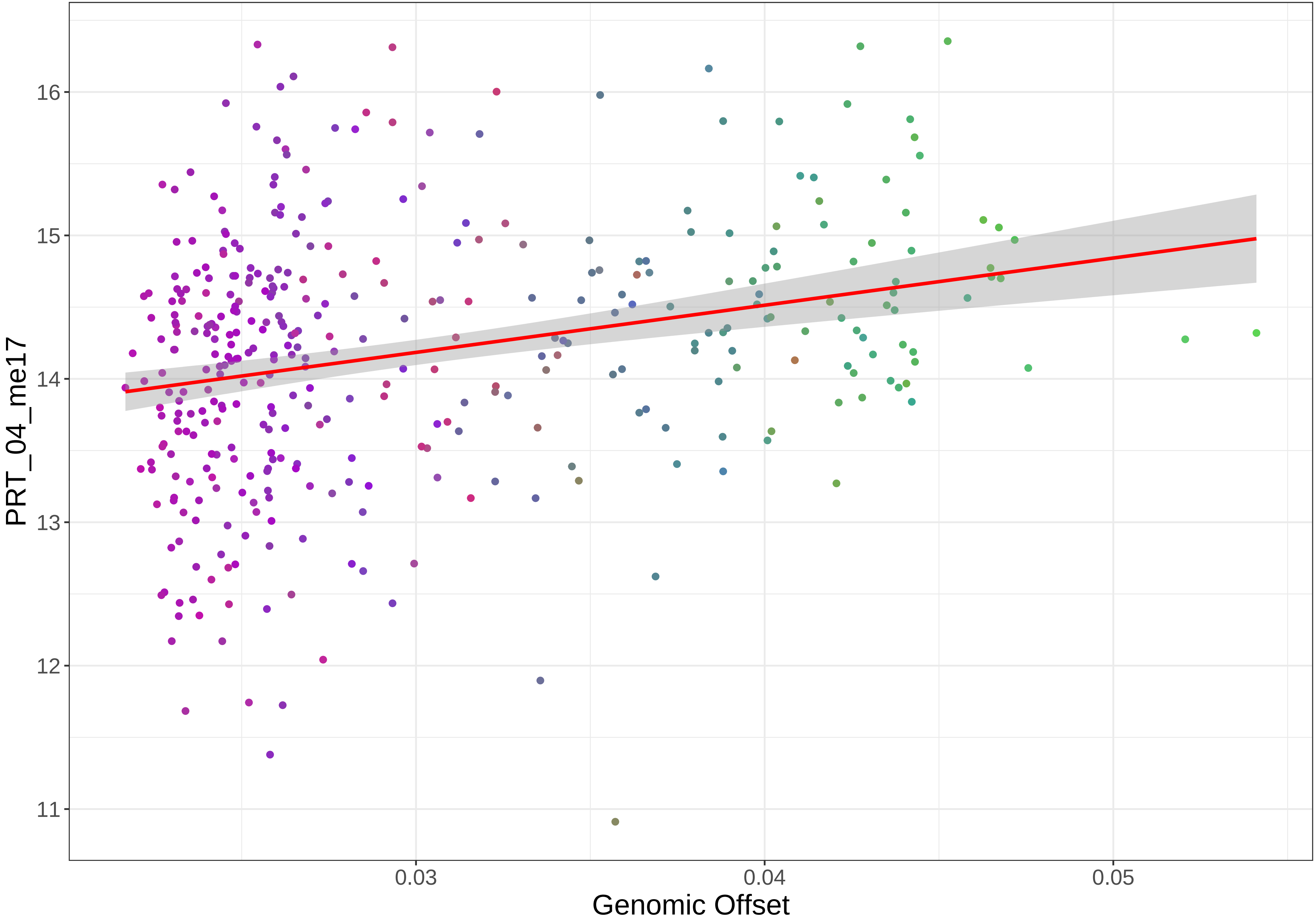

PRT_04_me17
16
15
14
13
12
11
0.03
0.04
0.05
Genomic Offset

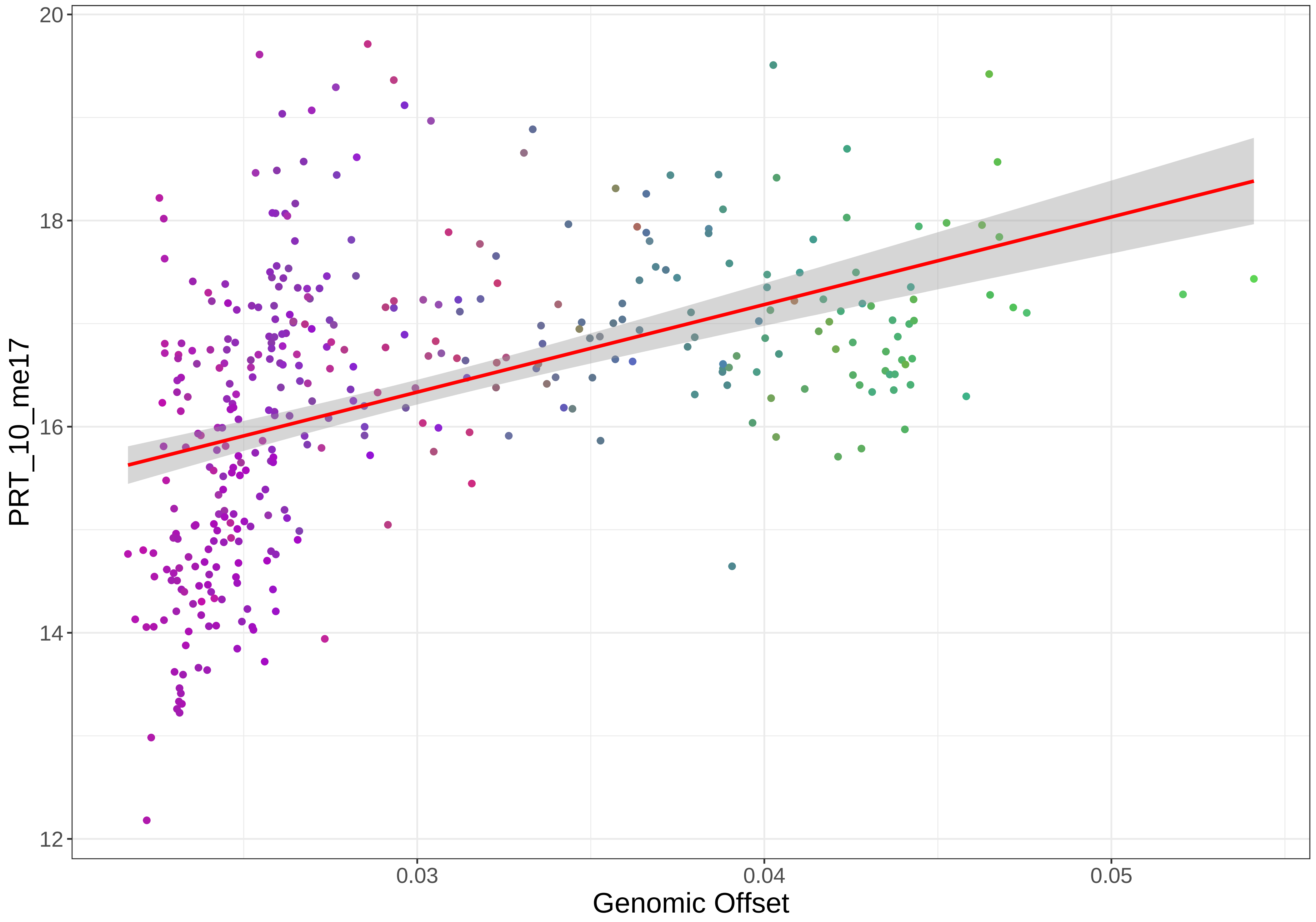

20
18
16
14
12
PRT_10_me17
0.03
0.04
0.05
Genomic Offset

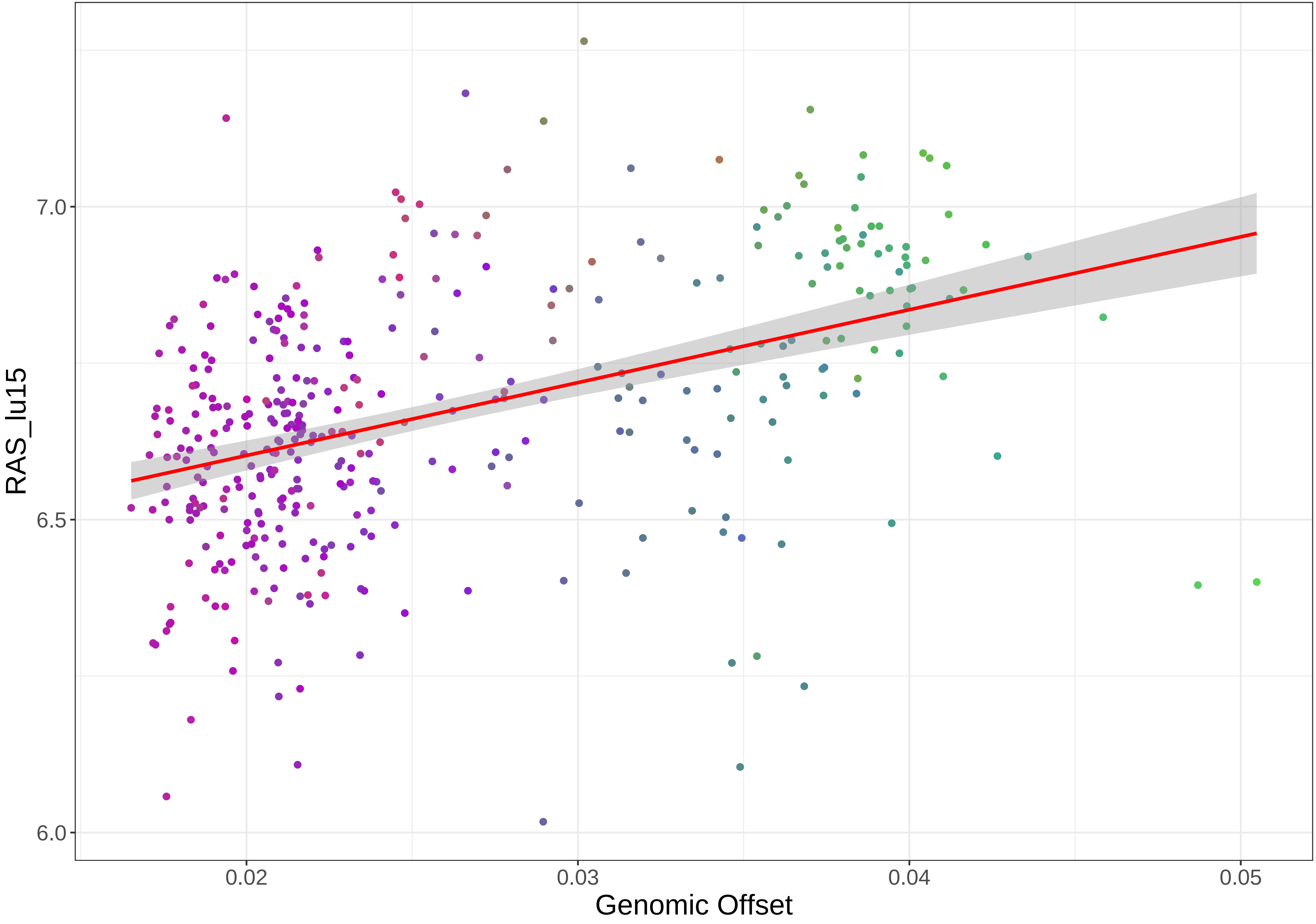

RAS_lu15
7.0
6.5
6.0
0.02
0.03
0.04
0.05
Genomic Offset

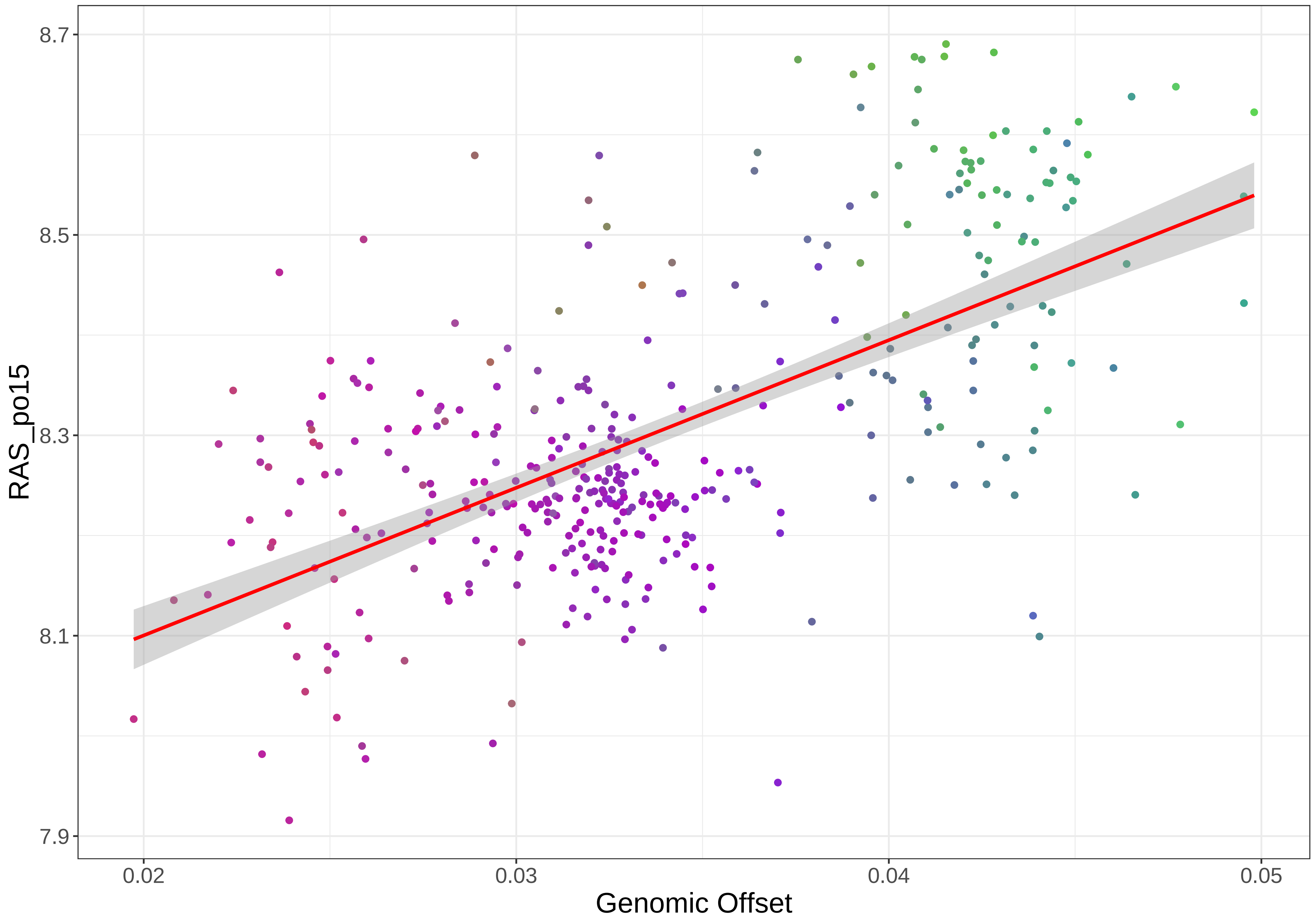

RAS_po15
8.7
8.5
8.3
8.1
7.9
0.02
0.03
0.04
0.05
Genomic Offset

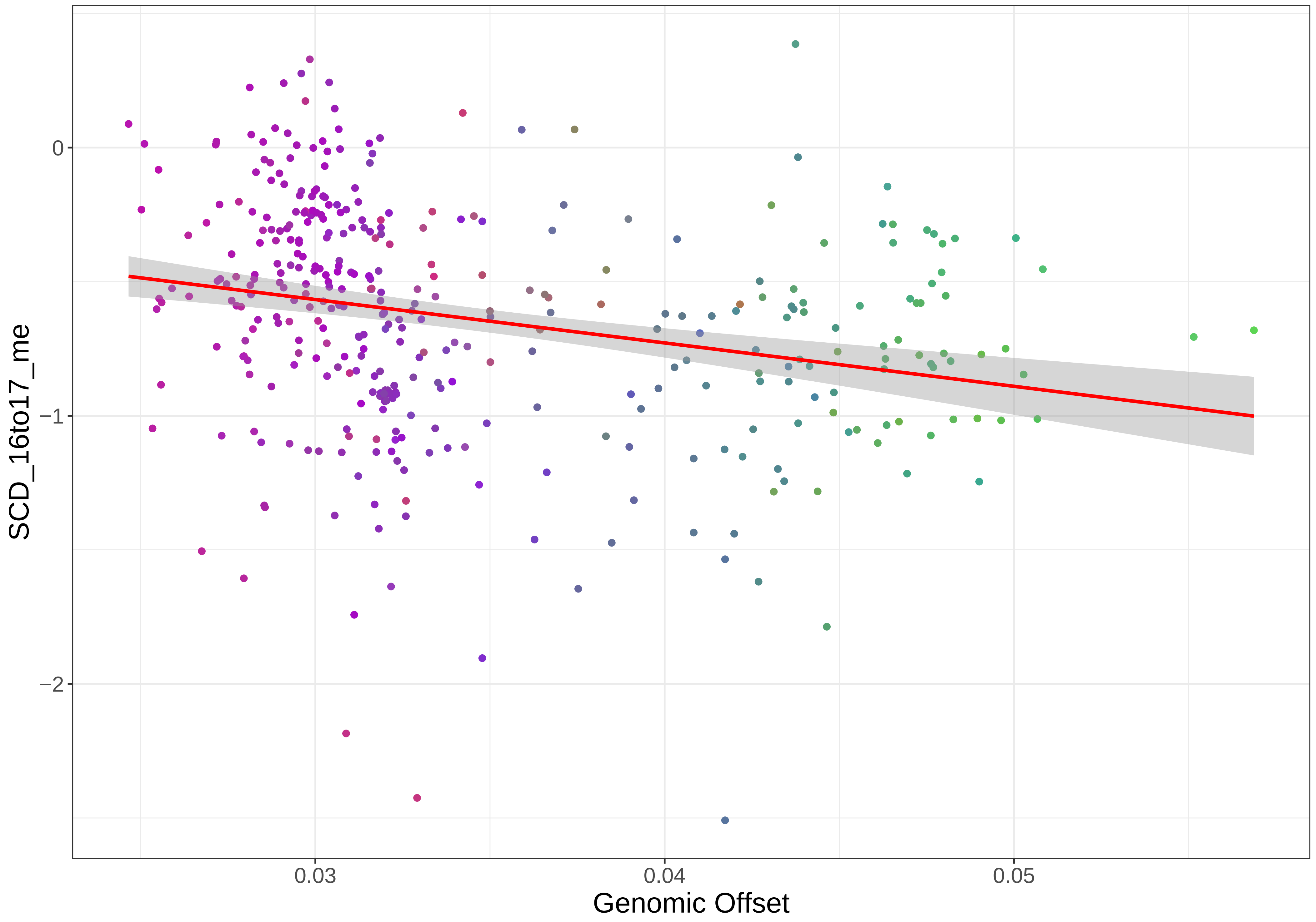
0
−1
−2
SCD_16to17_me
0.03
0.04
0.05
Genomic Offset

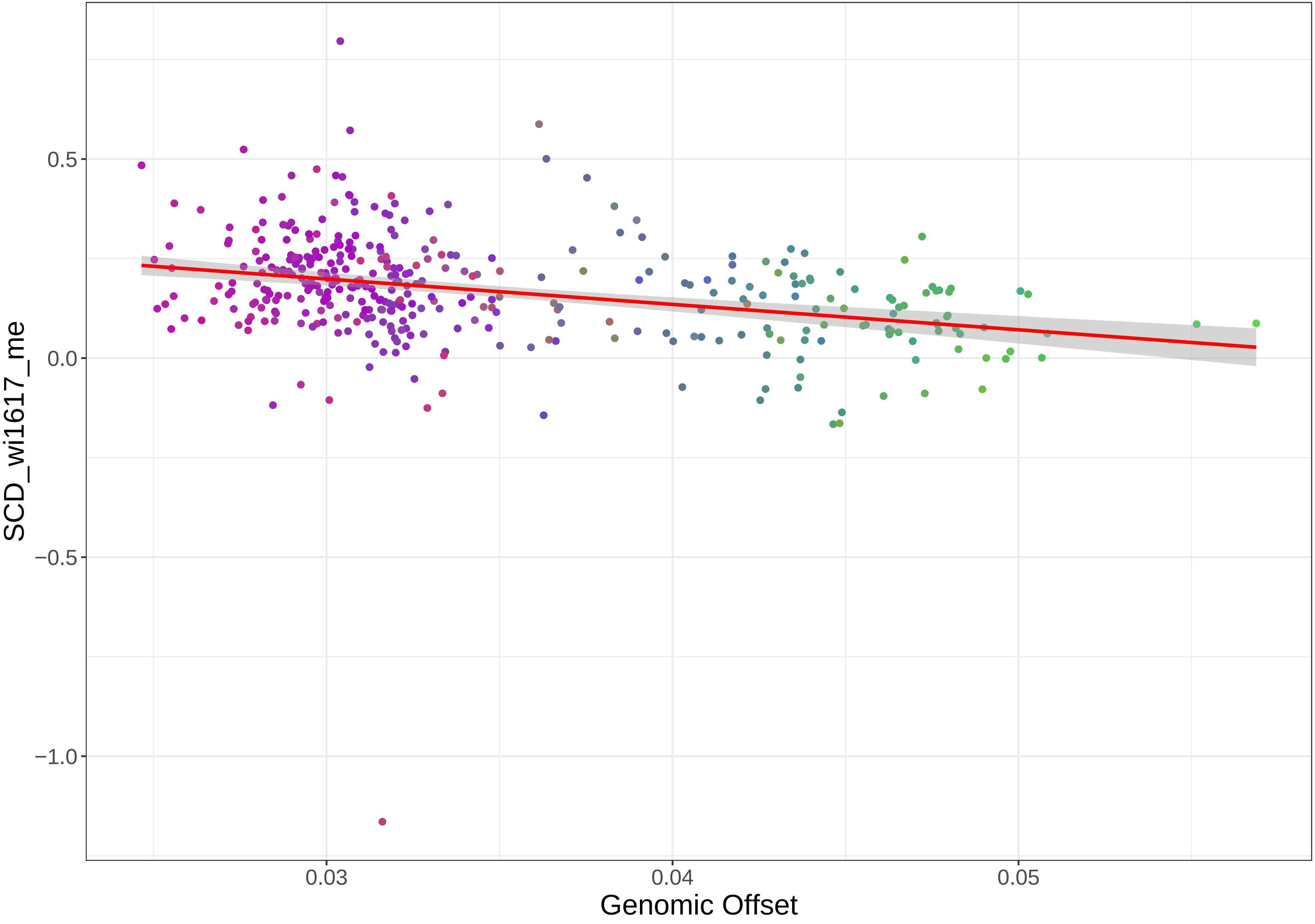
SCD_wi1617_me
0.5
0.0
−0.5
−1.0
0.03
0.04
0.05
Genomic Offset

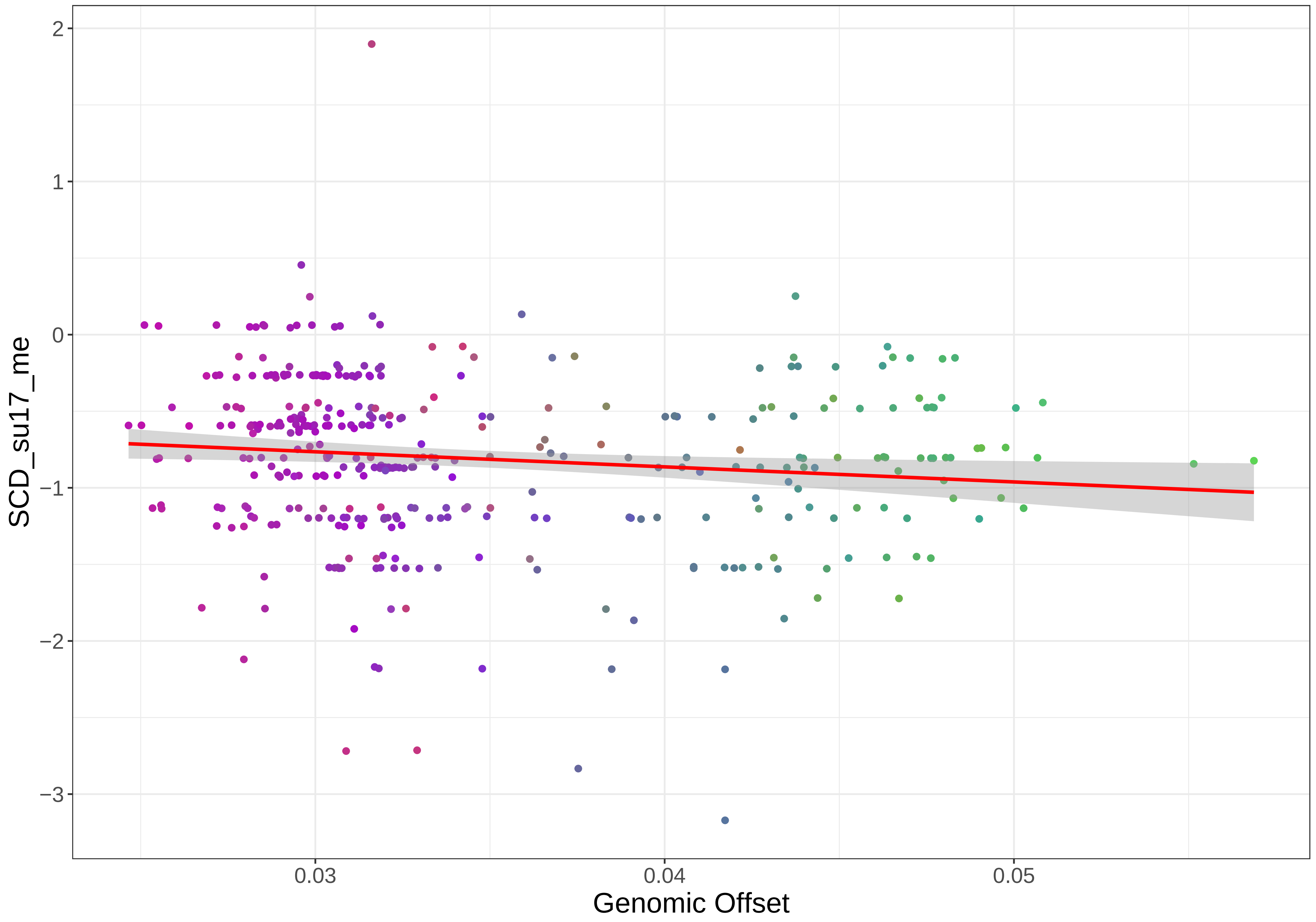

2
1
0
−1
−2
−3
SCD_su17_me
0.03
0.04
0.05
Genomic Offset

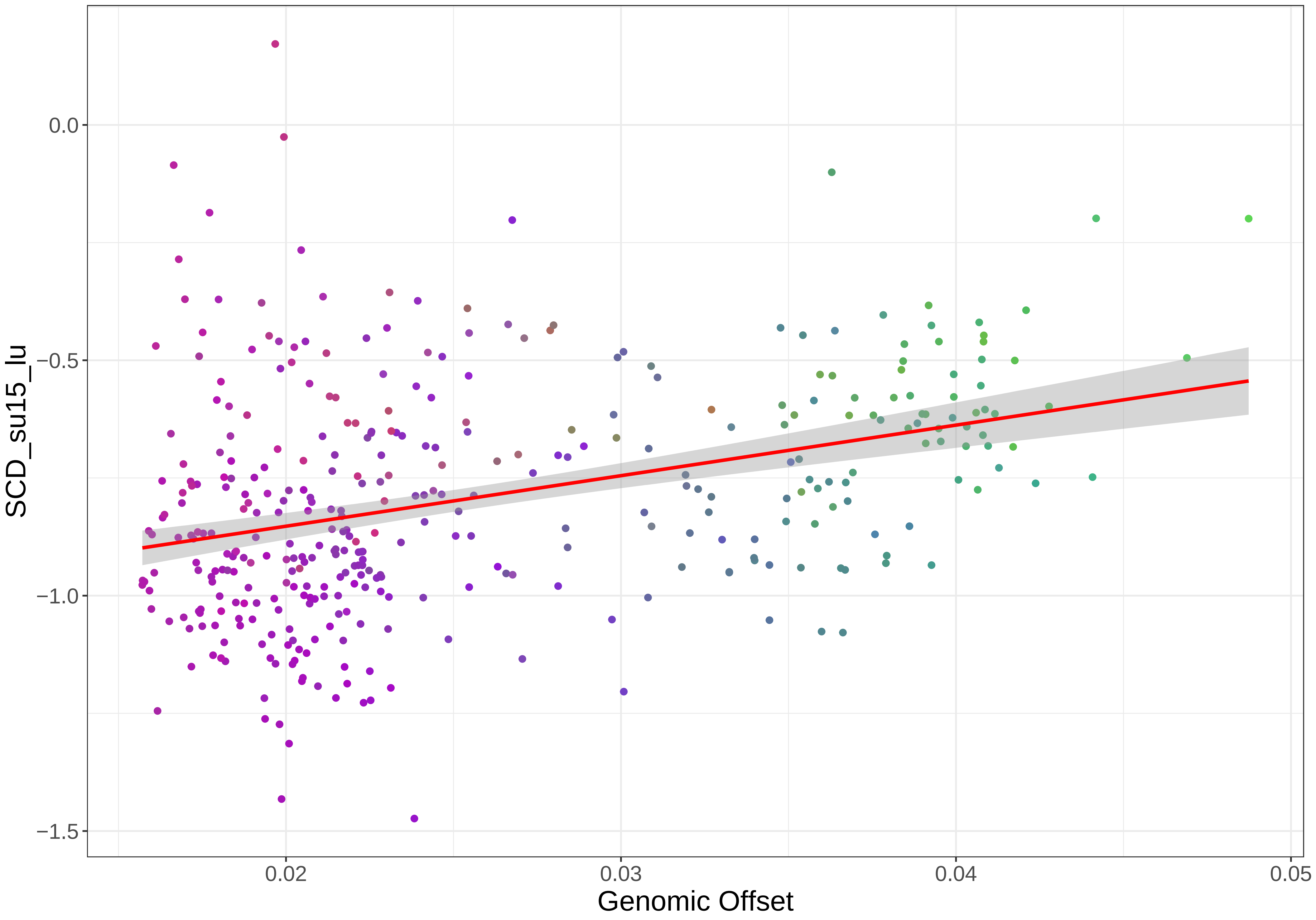

SCD_su15_lu
0.0
−0.5
−1.0
−1.5
0.02
0.03
0.04
0.05
Genomic Offset

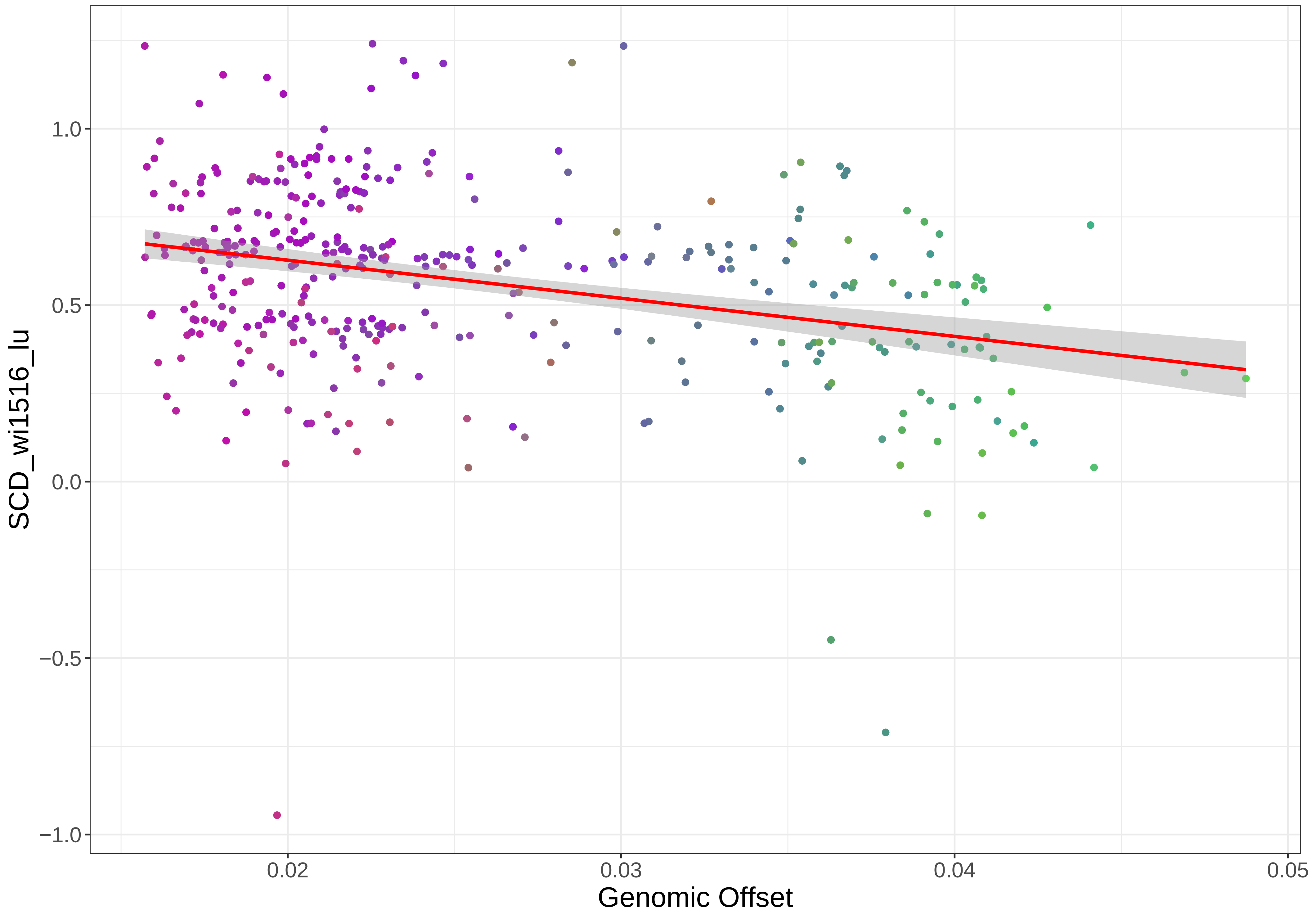

1.0
0.5
0.0
−0.5
−1.0
0.02
0.03
0.04
0.05
SCD_wi1516_lu
Genomic Offset

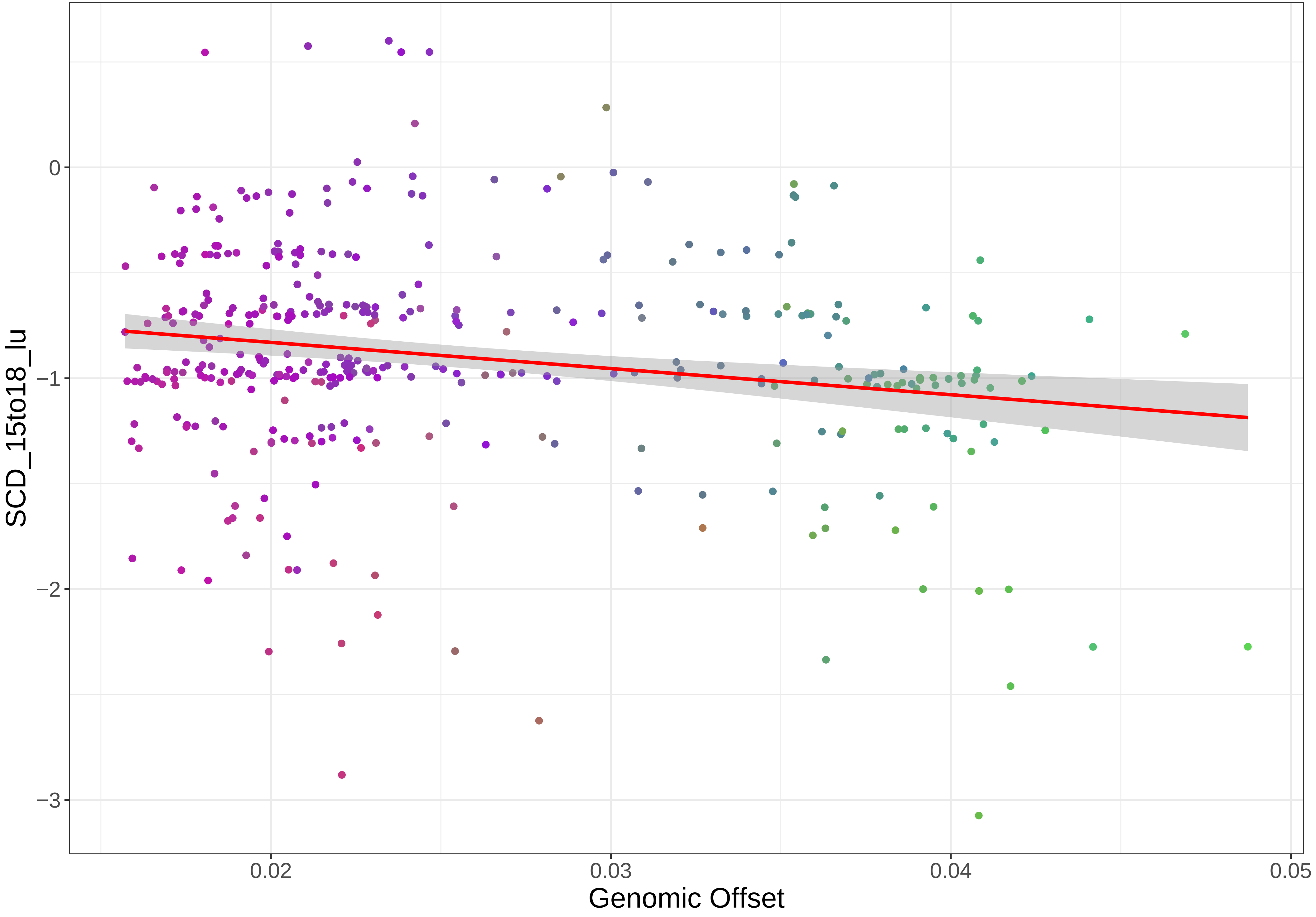

SCD_15to18_lu
0
−1
−2
−3
0.02
0.03
0.04
0.05
Genomic Offset

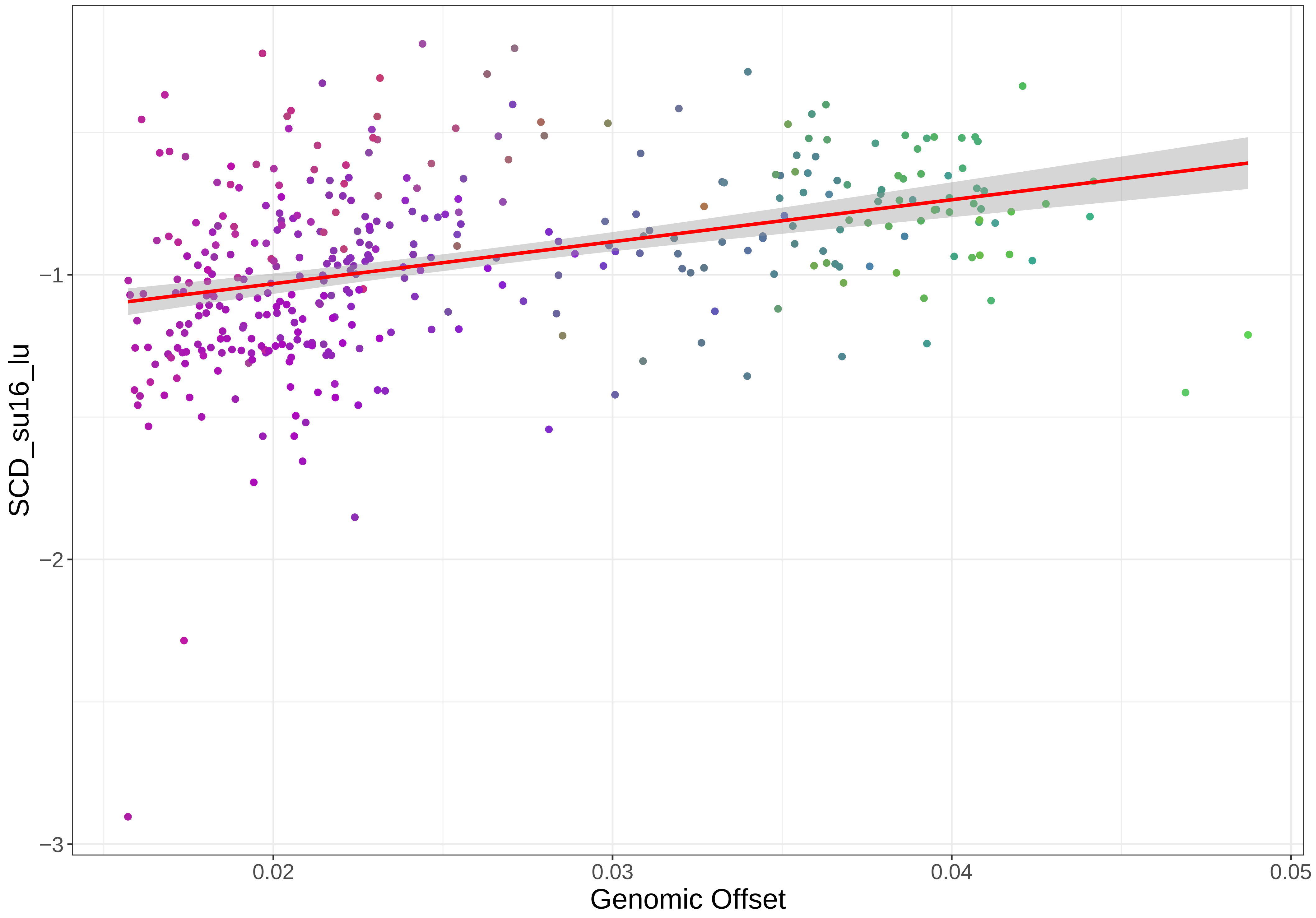
SCD_su16_lu
−1
−2
−3
0.02
0.03
0.04
0.05
Genomic Offset

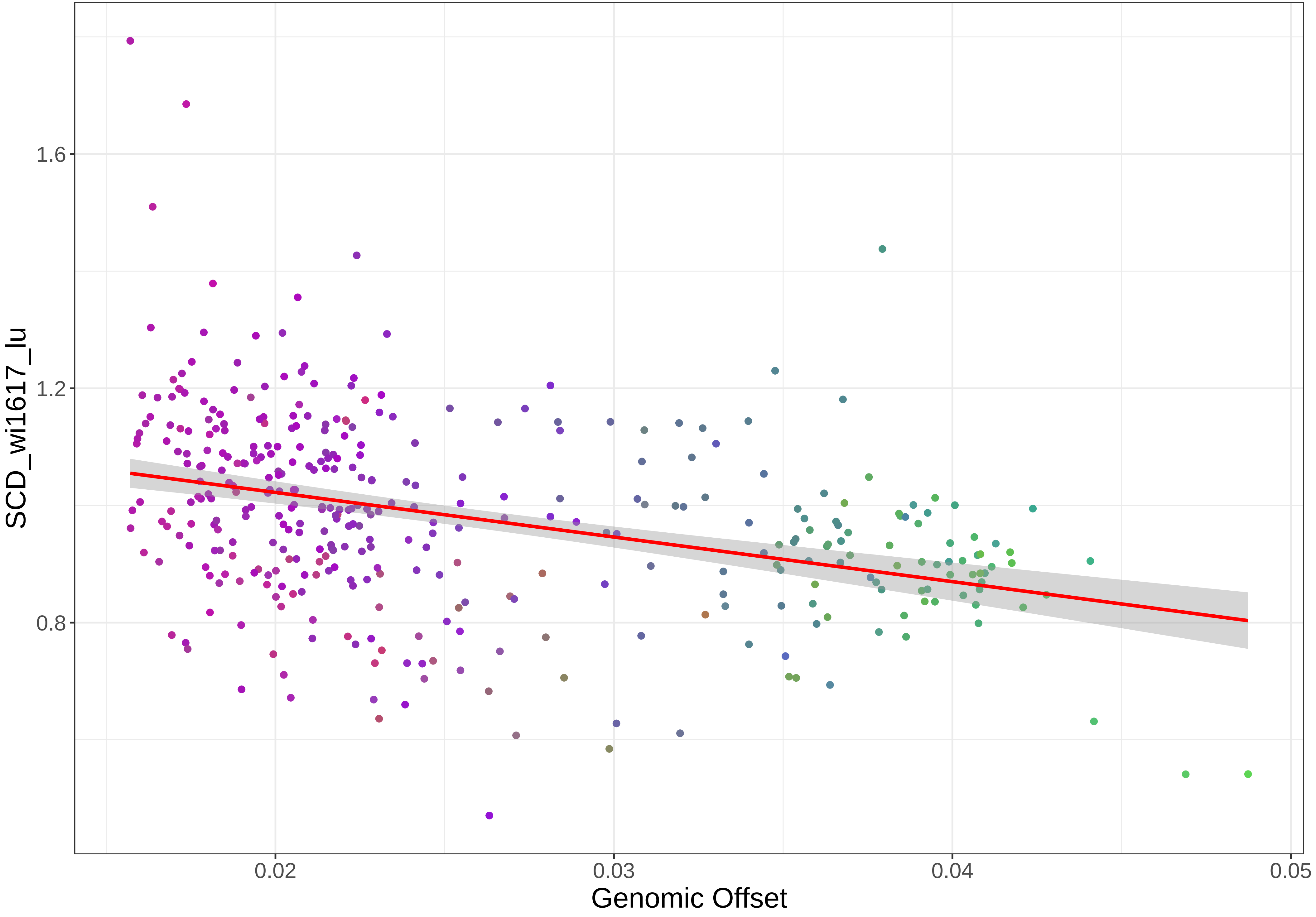

1.6
1.2
0.8
SCD_wi1617_lu
0.02
0.03
0.04
0.05
Genomic Offset

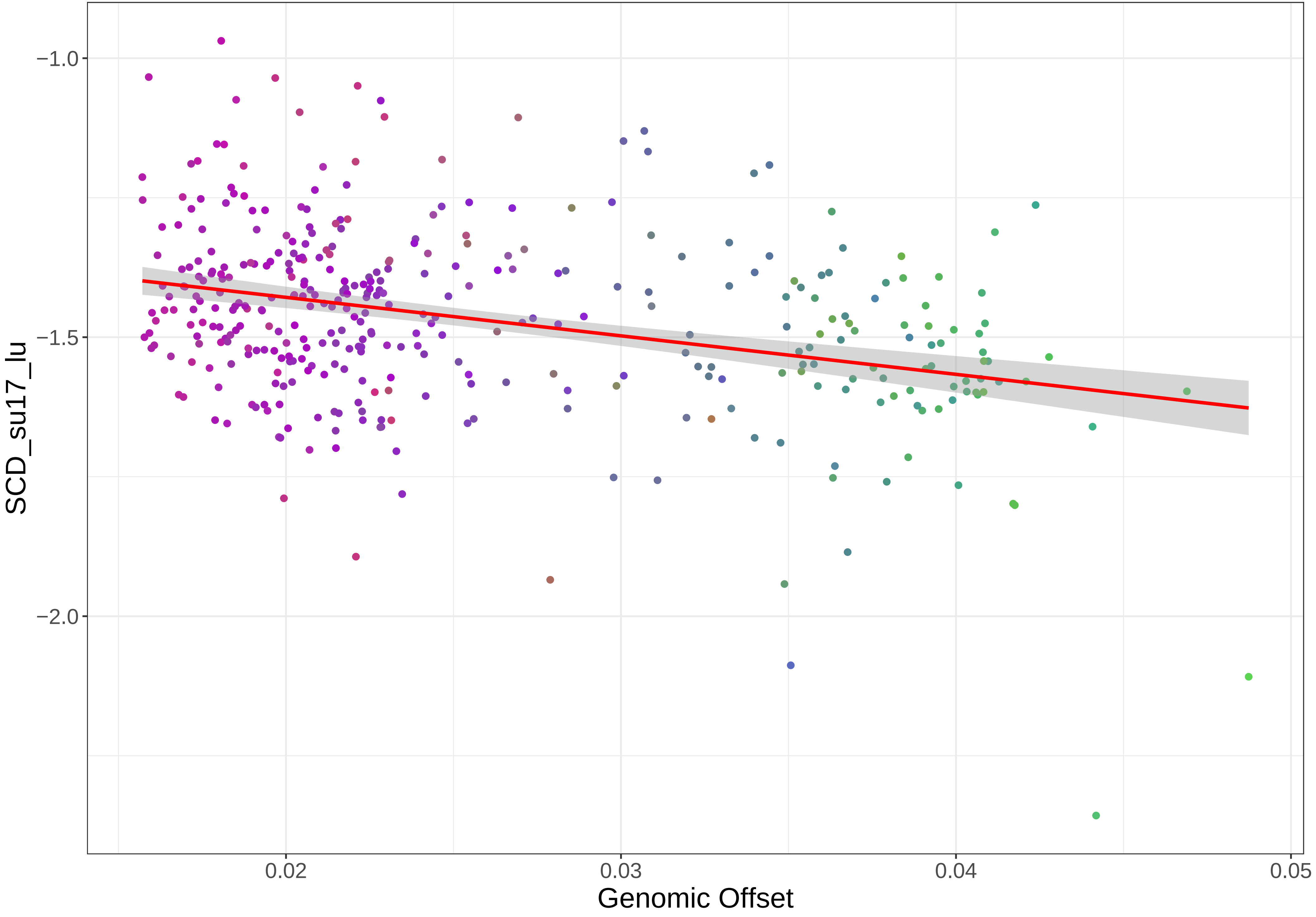

SCD_su17_lu
−1.0
−1.5
−2.0
Genomic Offset
0.02
0.03
0.04
0.05

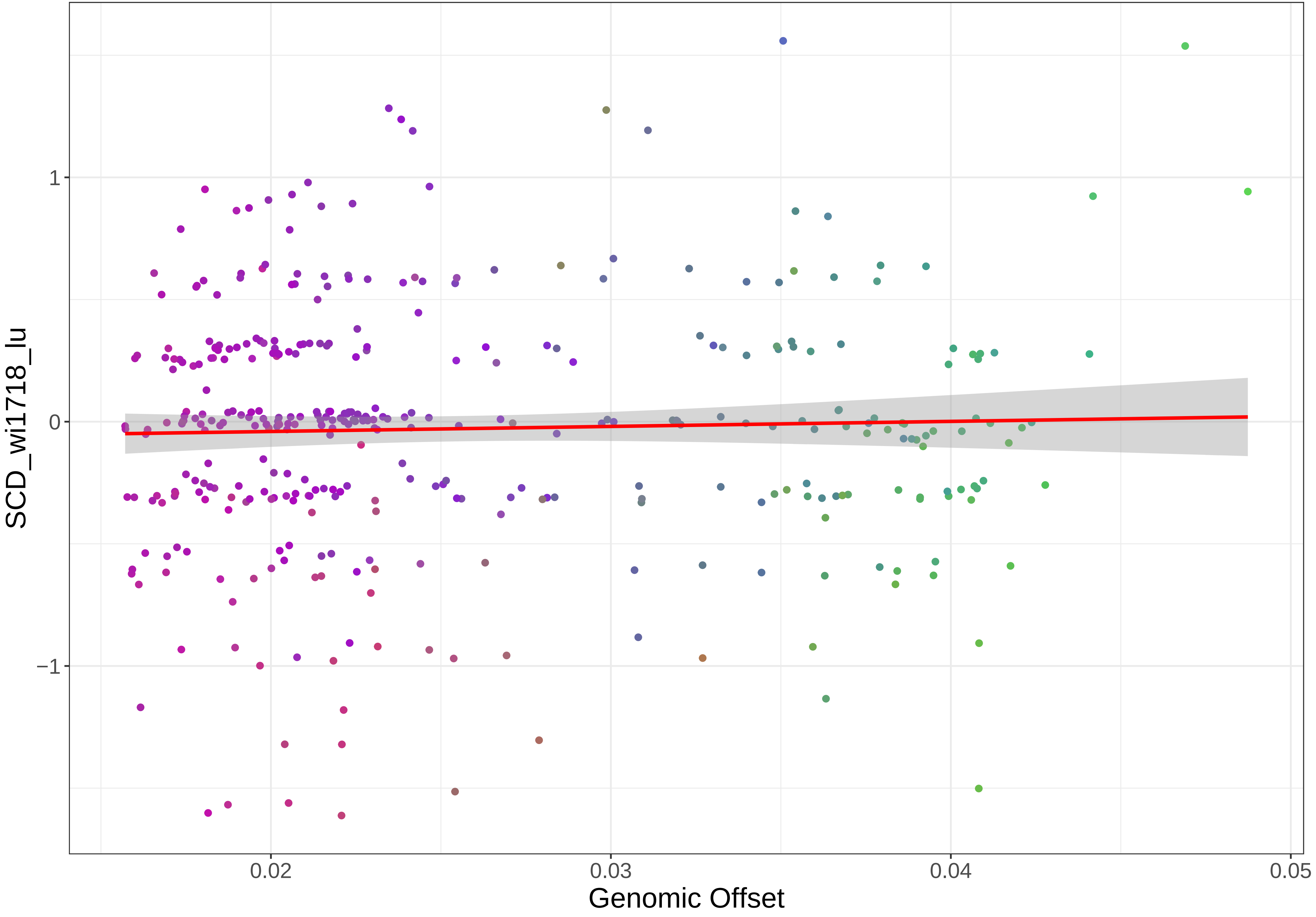
SCD_wi1718_lu
1
0
−1
0.02
0.03
0.04
0.05
Genomic Offset

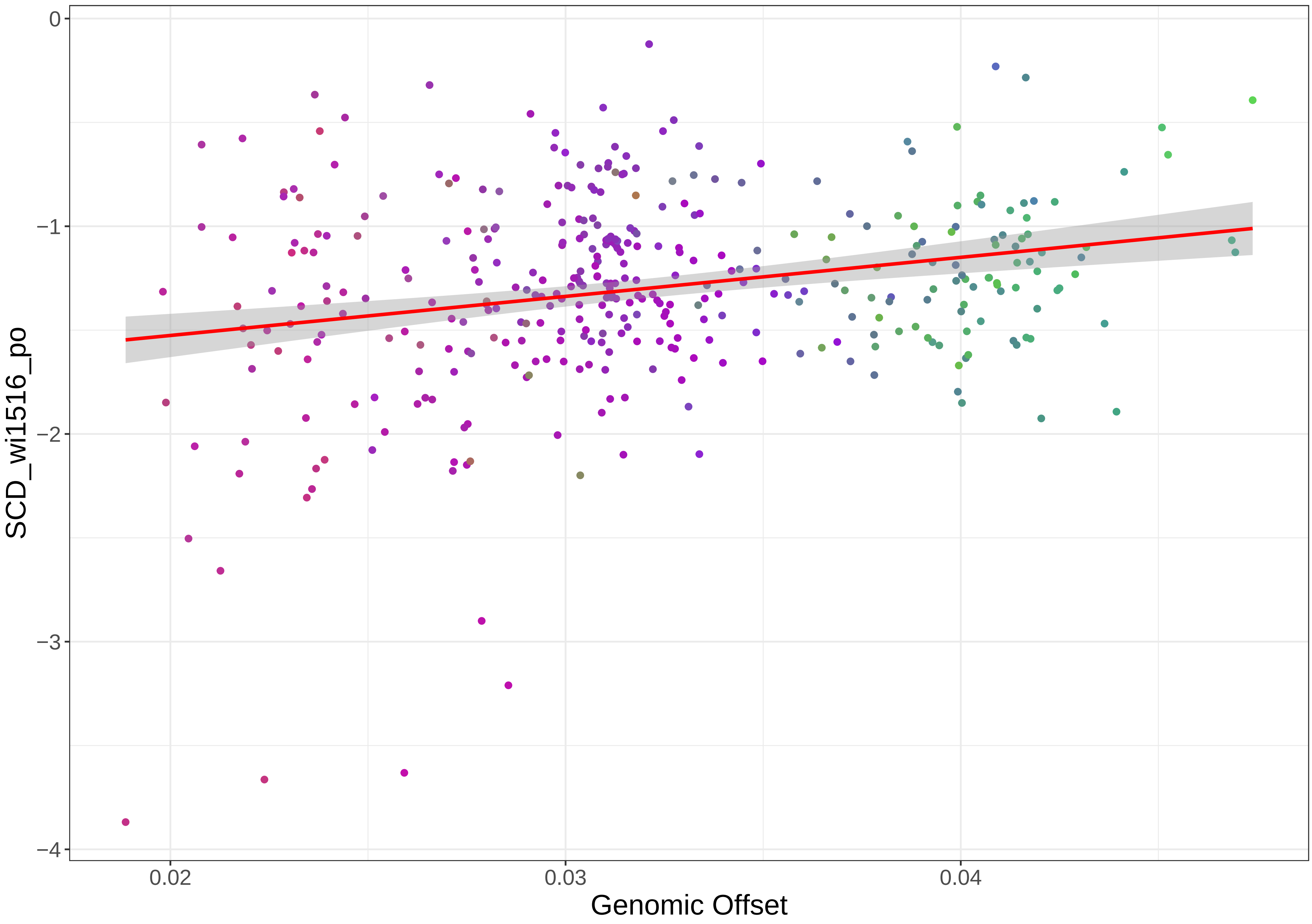

0
−1
−2
−3
−4
SCD_wi1516_po
0.02
0.03
0.04
Genomic Offset

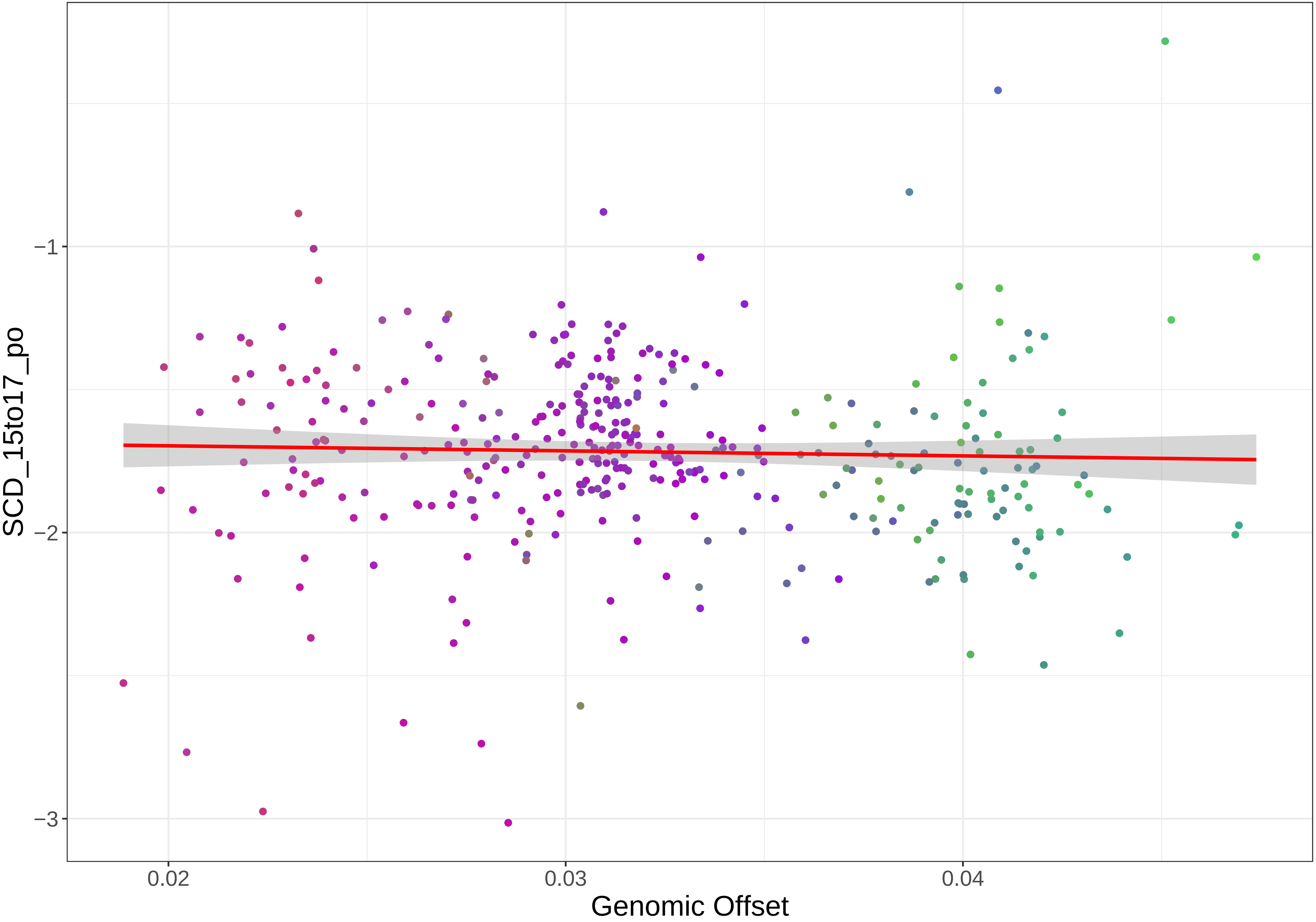

SCD_15to17_po
−1
−2
−3
0.02
0.03
0.04
Genomic Offset

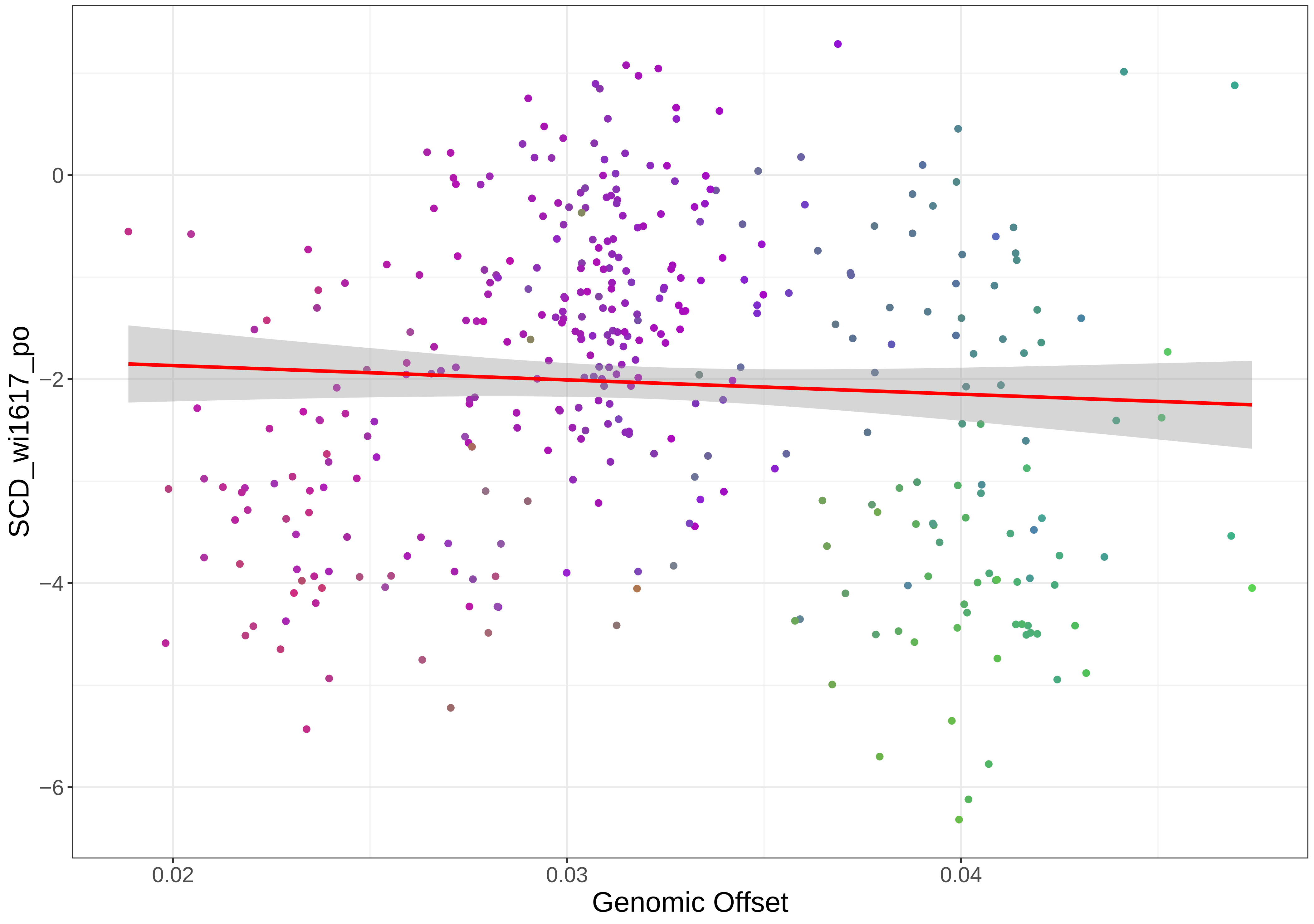

0
−2
−4
−6
SCD_wi1617_po
0.02
0.03
0.04
Genomic Offset

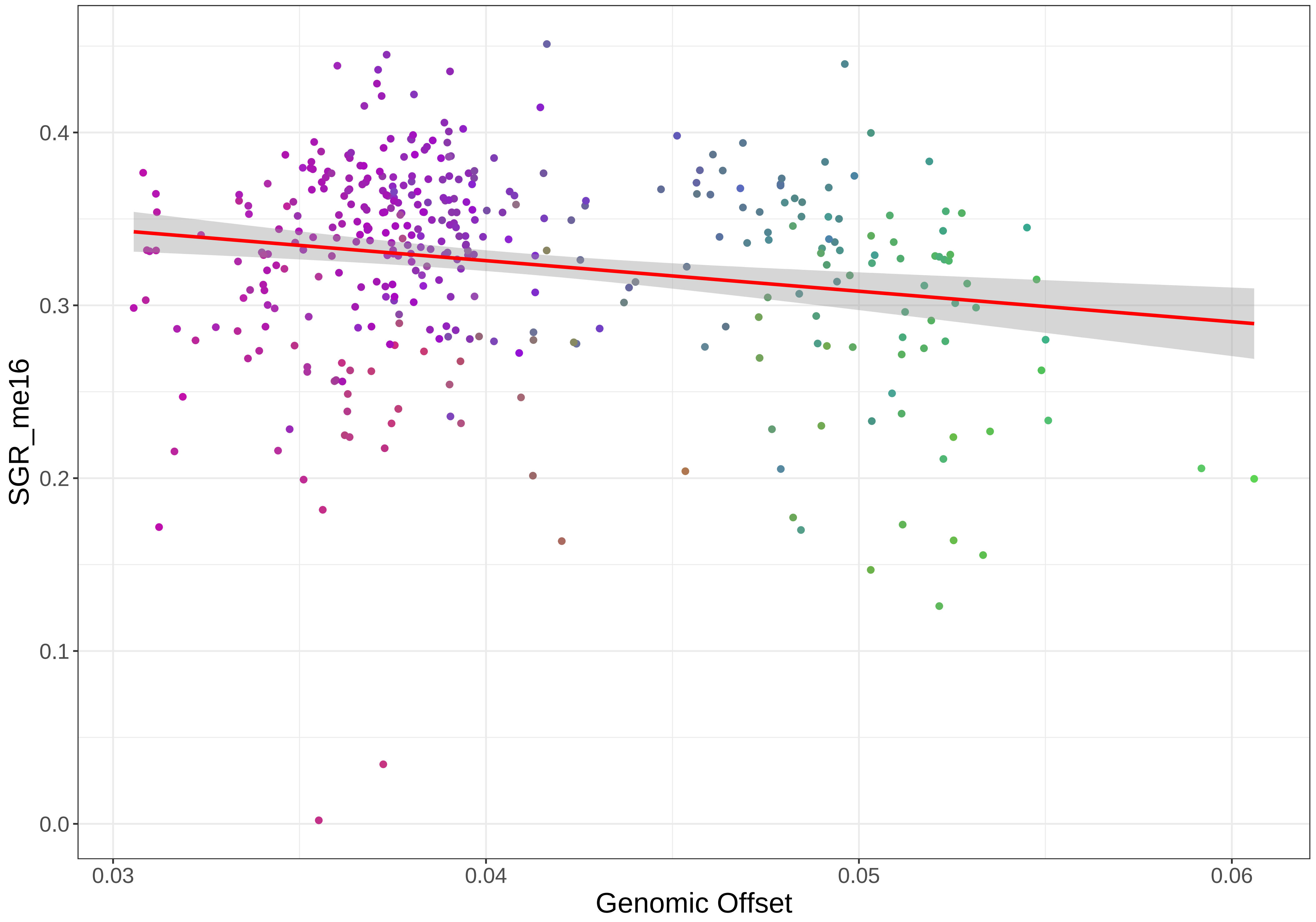
SGR_me16
0.0
0.1
0.2
0.3
0.4
0.03
0.04
0.05
0.06
Genomic Offset

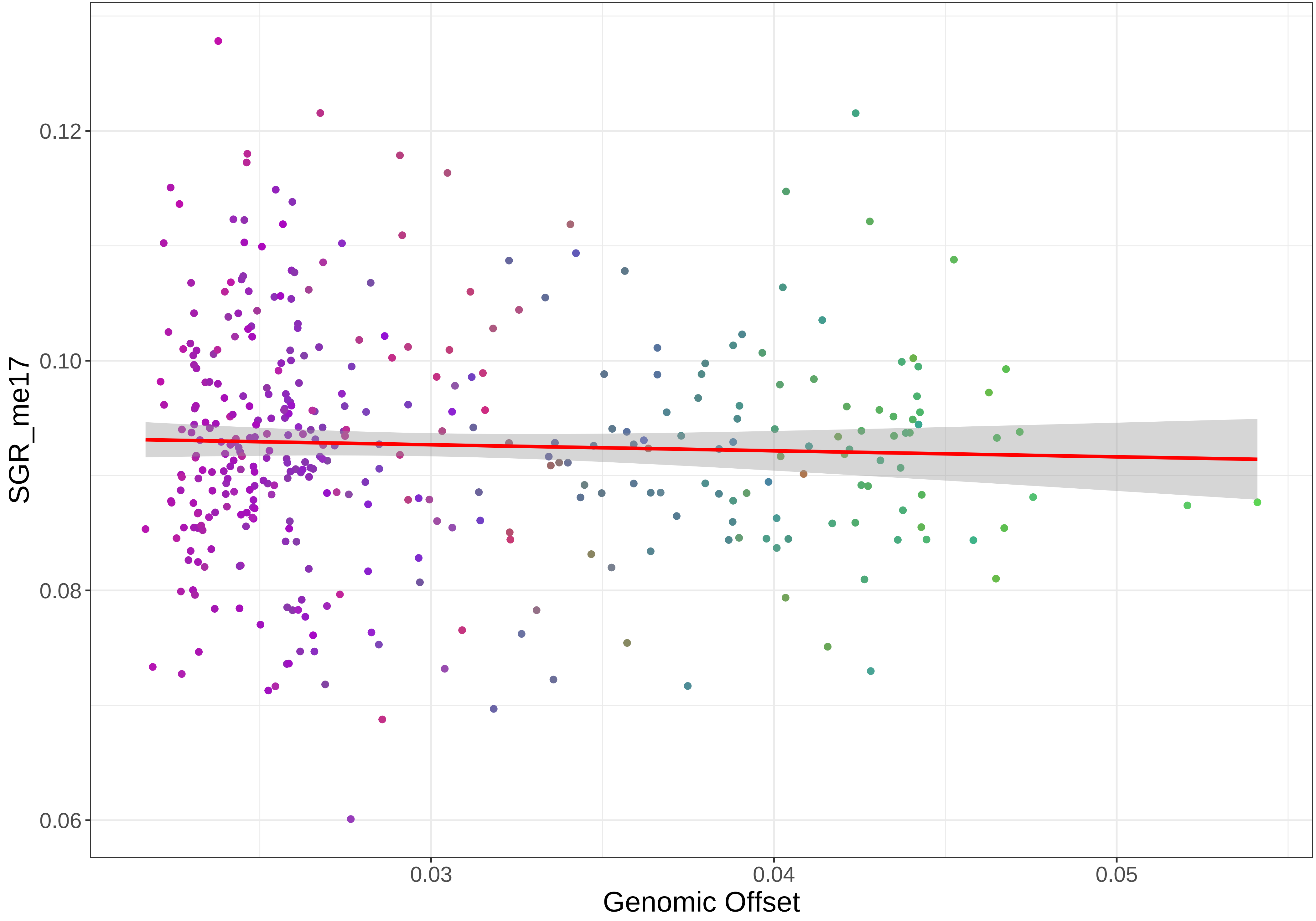

SGR_me17
0.12
0.10
0.08
0.06
0.03
0.04
0.05
Genomic Offset

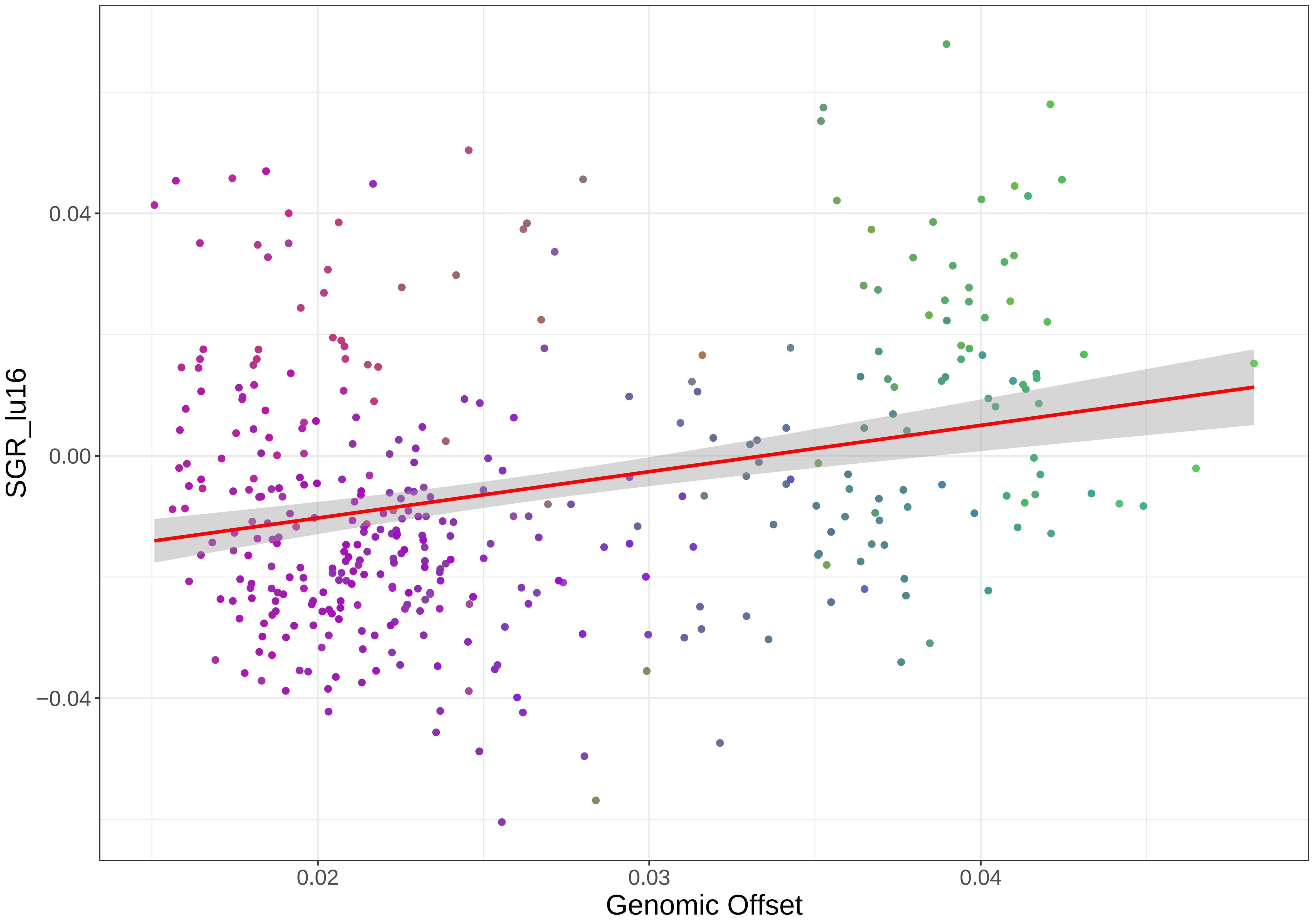
SGR_lu16
0.04
0.00
−0.04
0.02
0.03
0.04
Genomic Offset

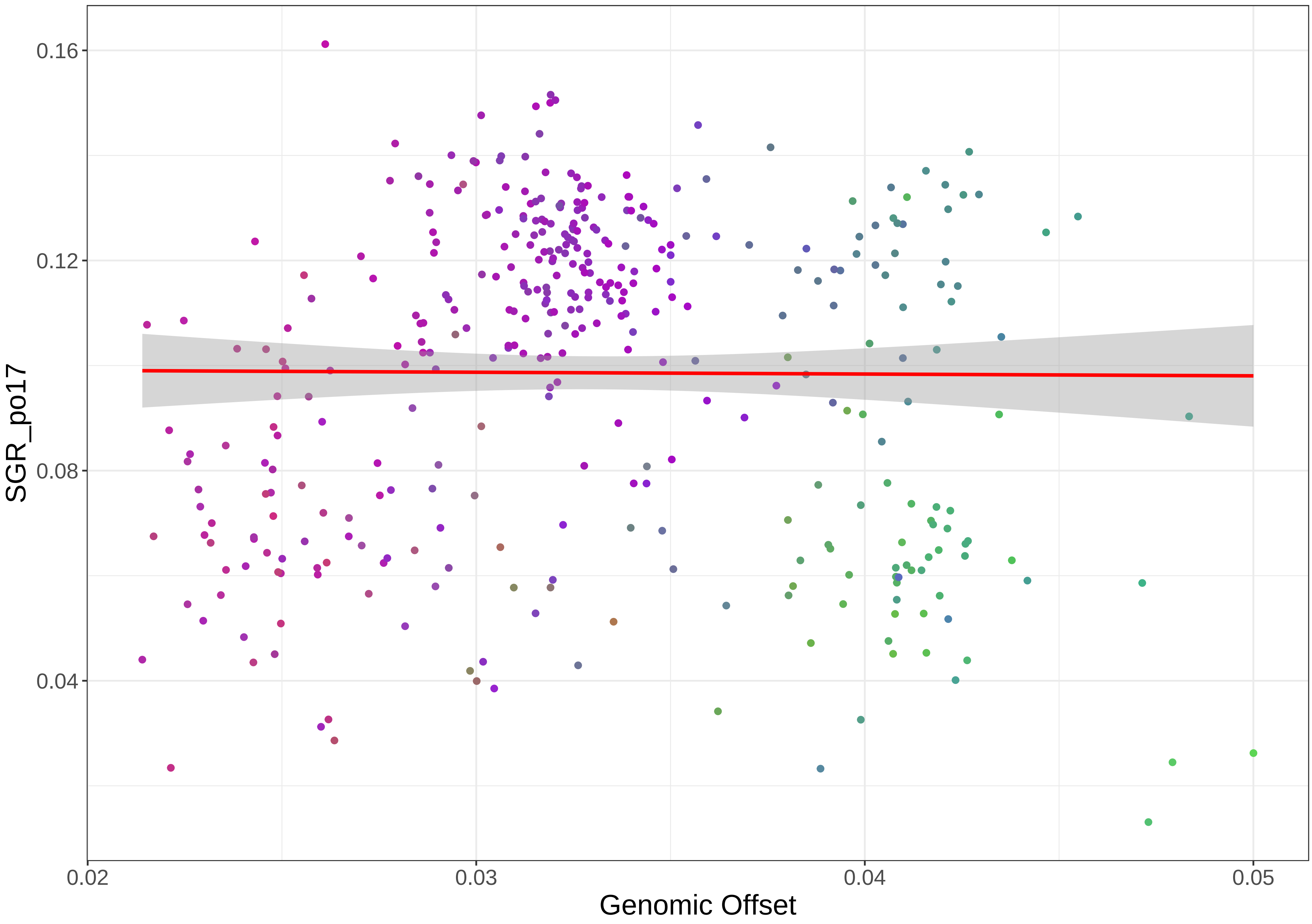
0.16
0.12
0.08
0.04
SGR_po17
0.02
0.03
0.04
0.05
Genomic Offset

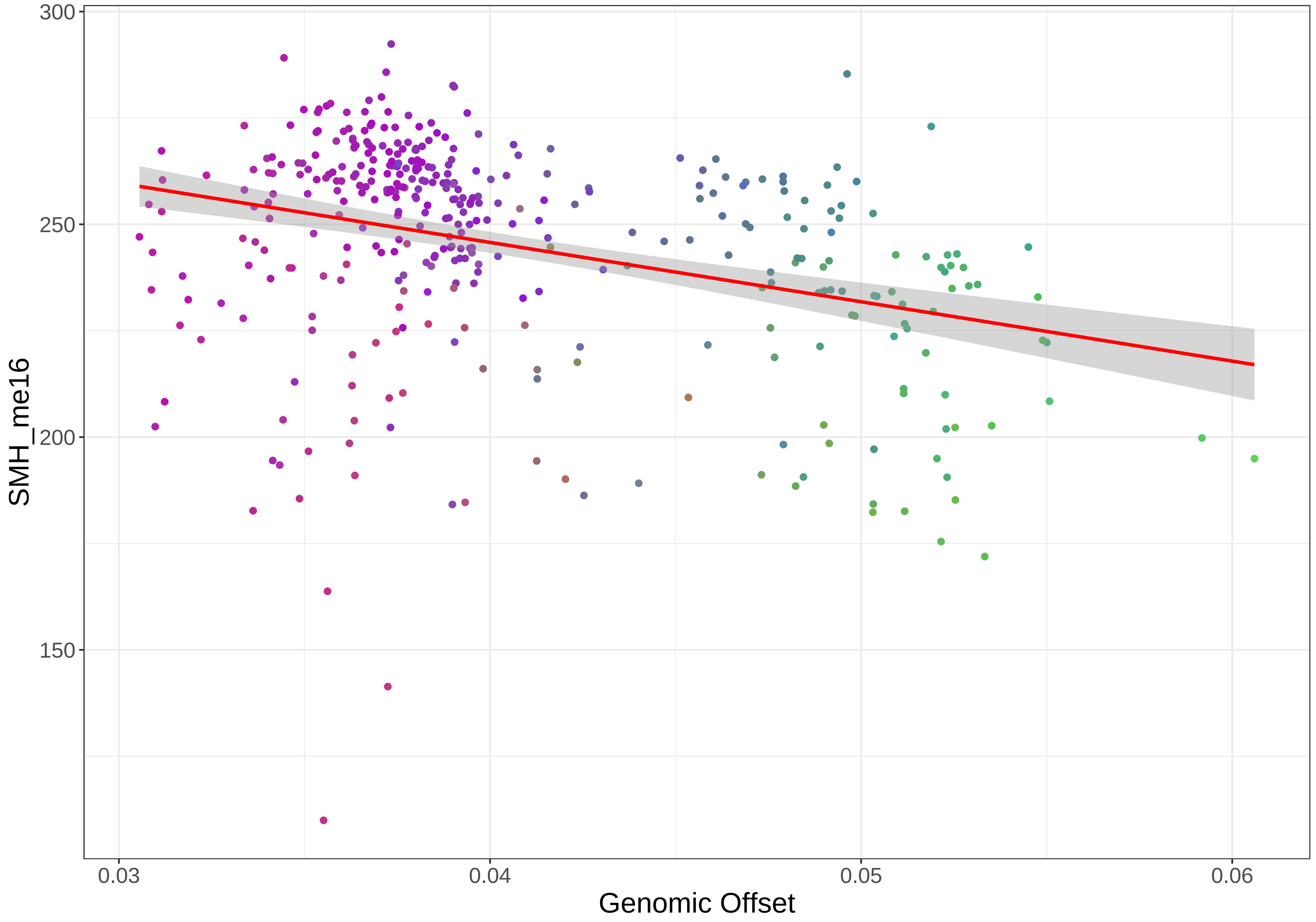

300
250
200
150
SMH_me16
0.03
0.04
0.05
0.06
Genomic Offset

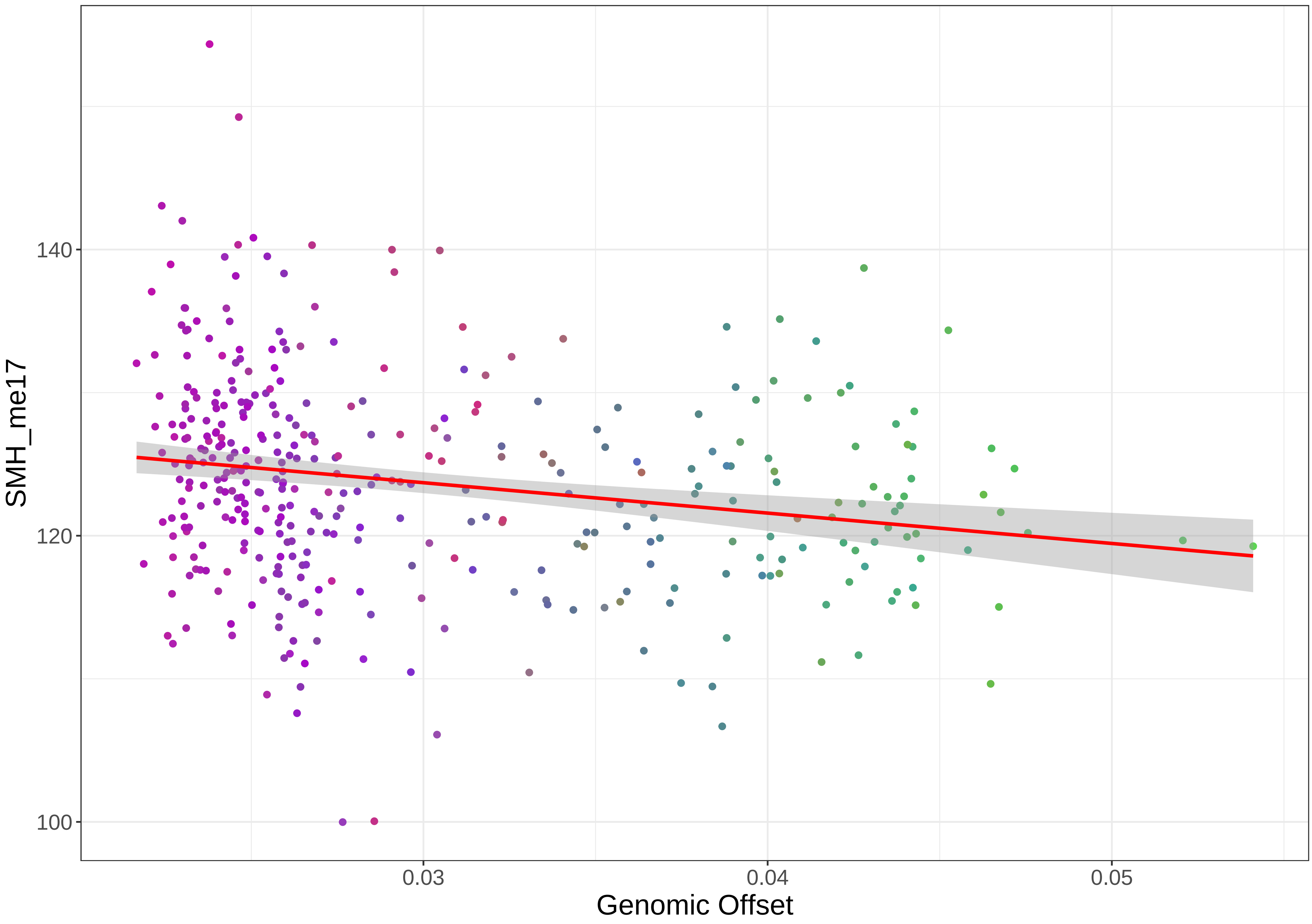

SMH_me17
140
120
100
0.03
0.04
0.05
Genomic Offset

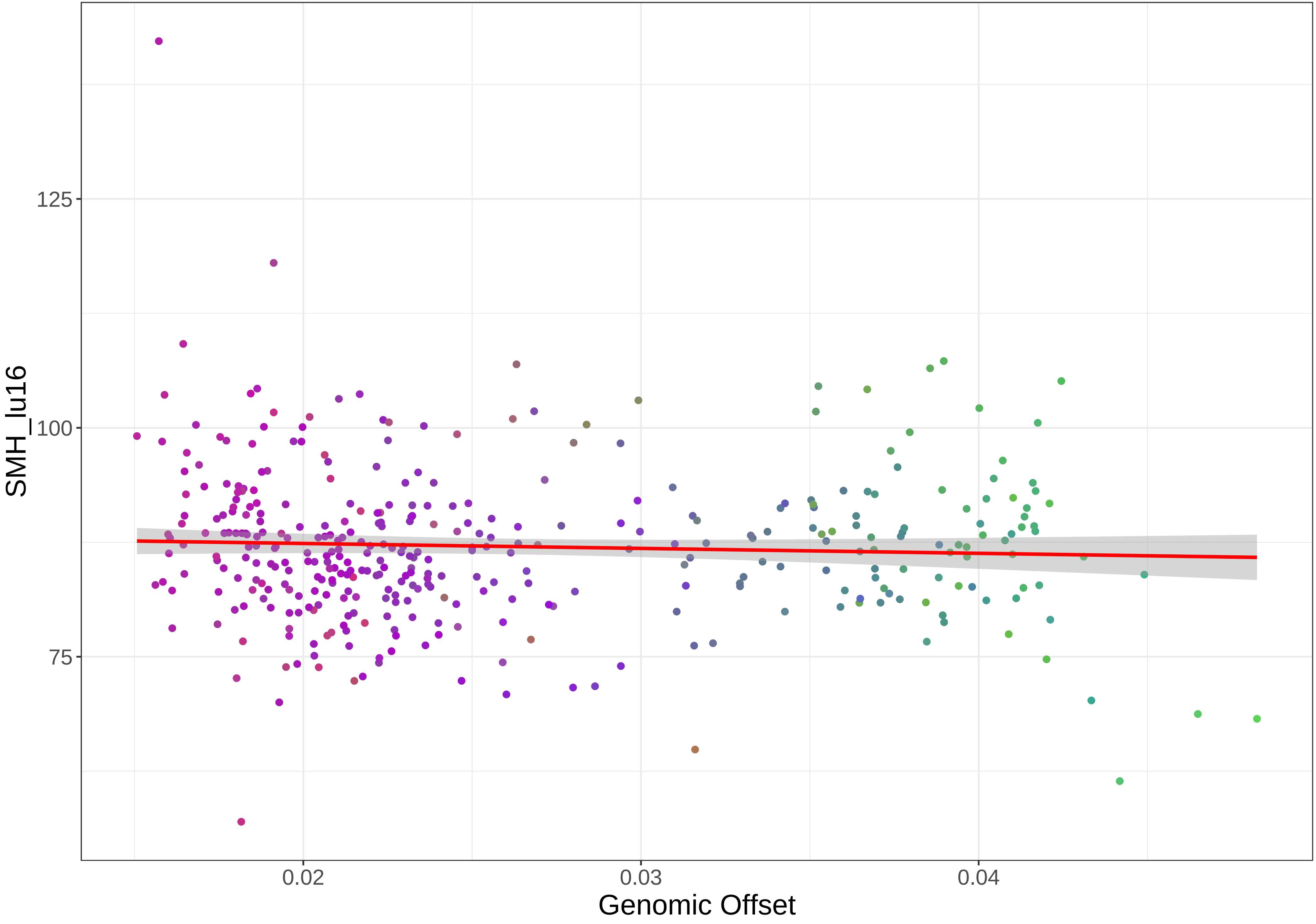
SMH_lu16
125
100
75
0.02
0.03
0.04
Genomic Offset

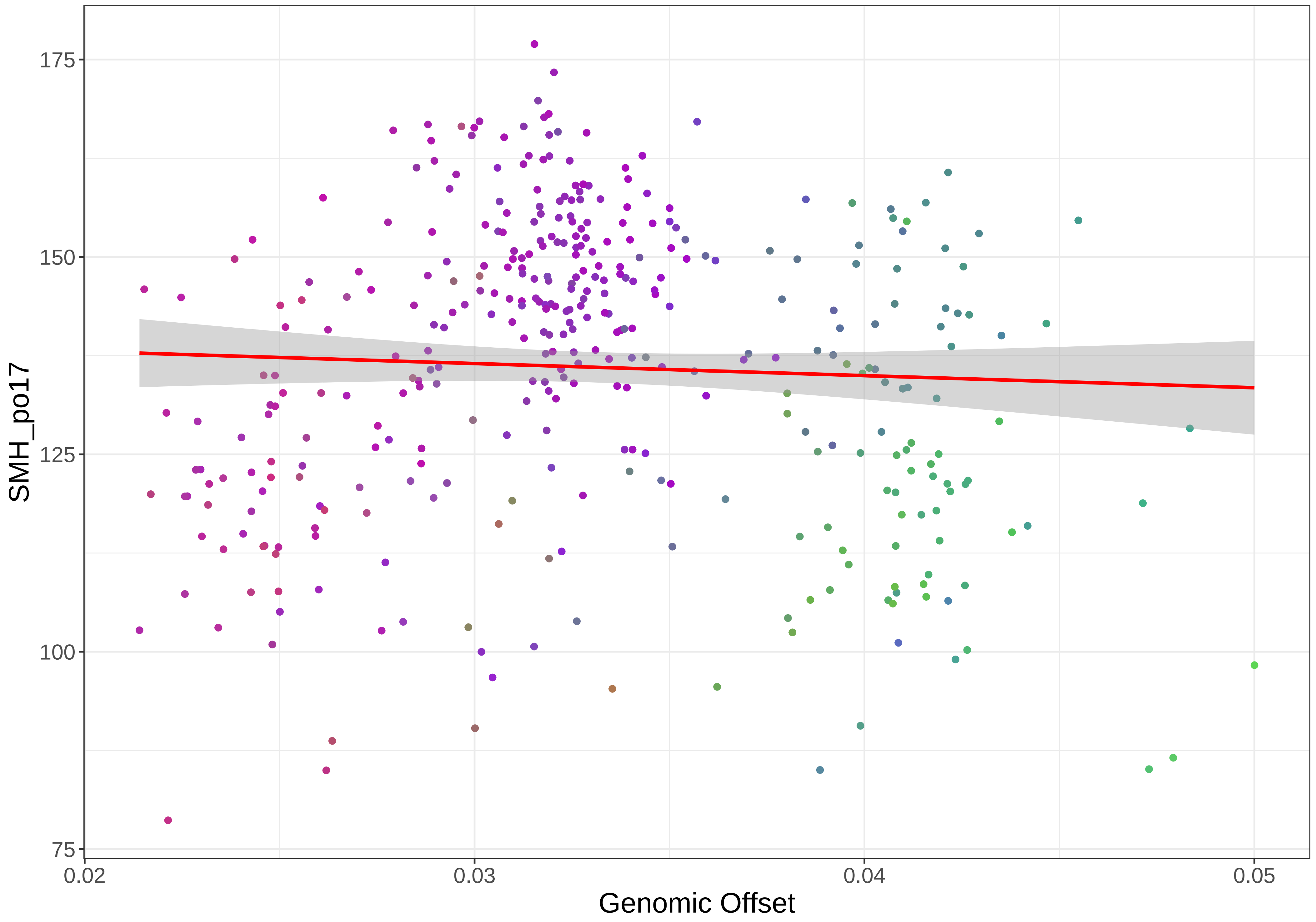
175
150
125
100
75
SMH_po17
0.02
0.03
0.04
0.05
Genomic Offset

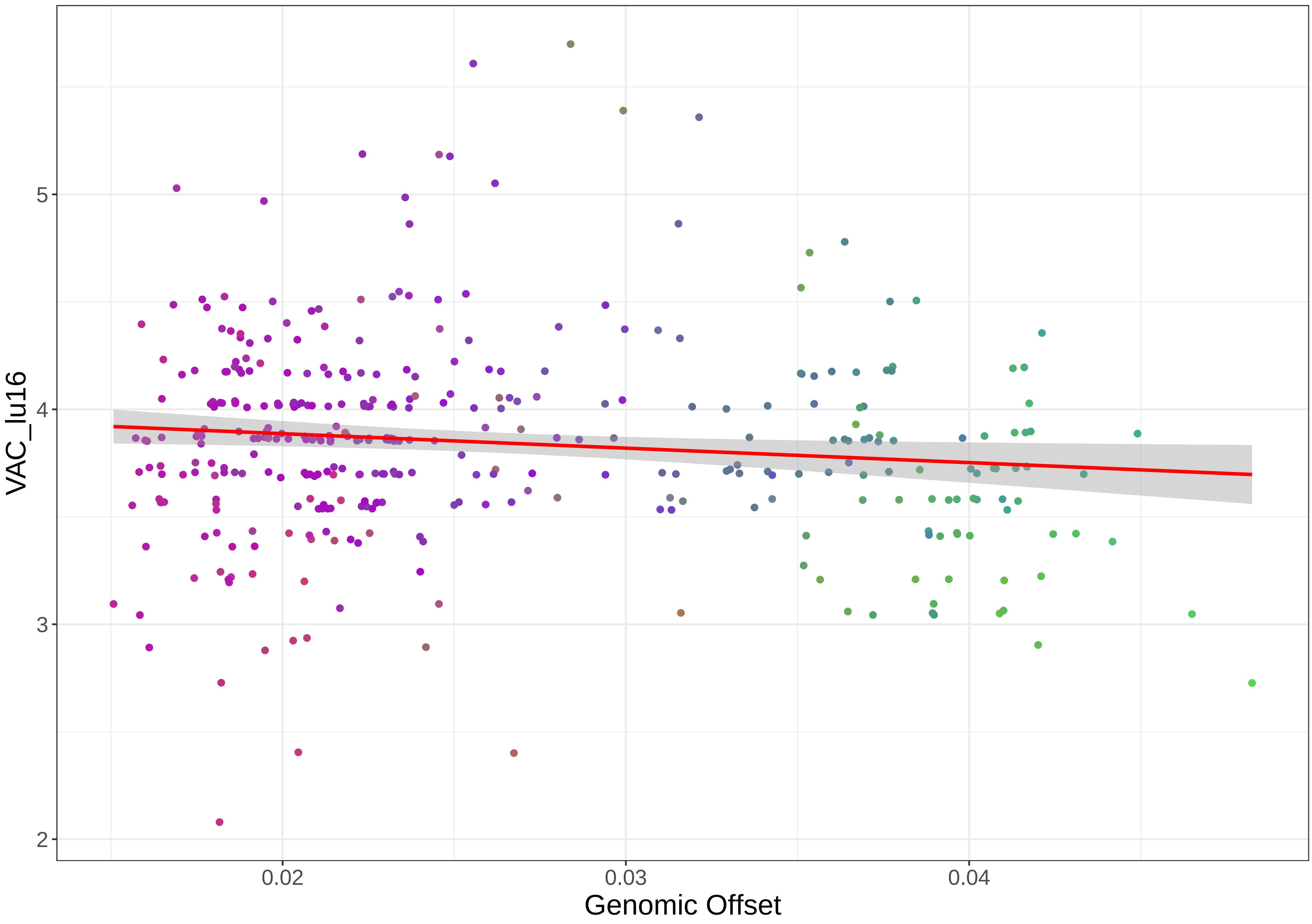
VAC_lu16
5
4
3
2
0.02
0.03
0.04
Genomic Offset

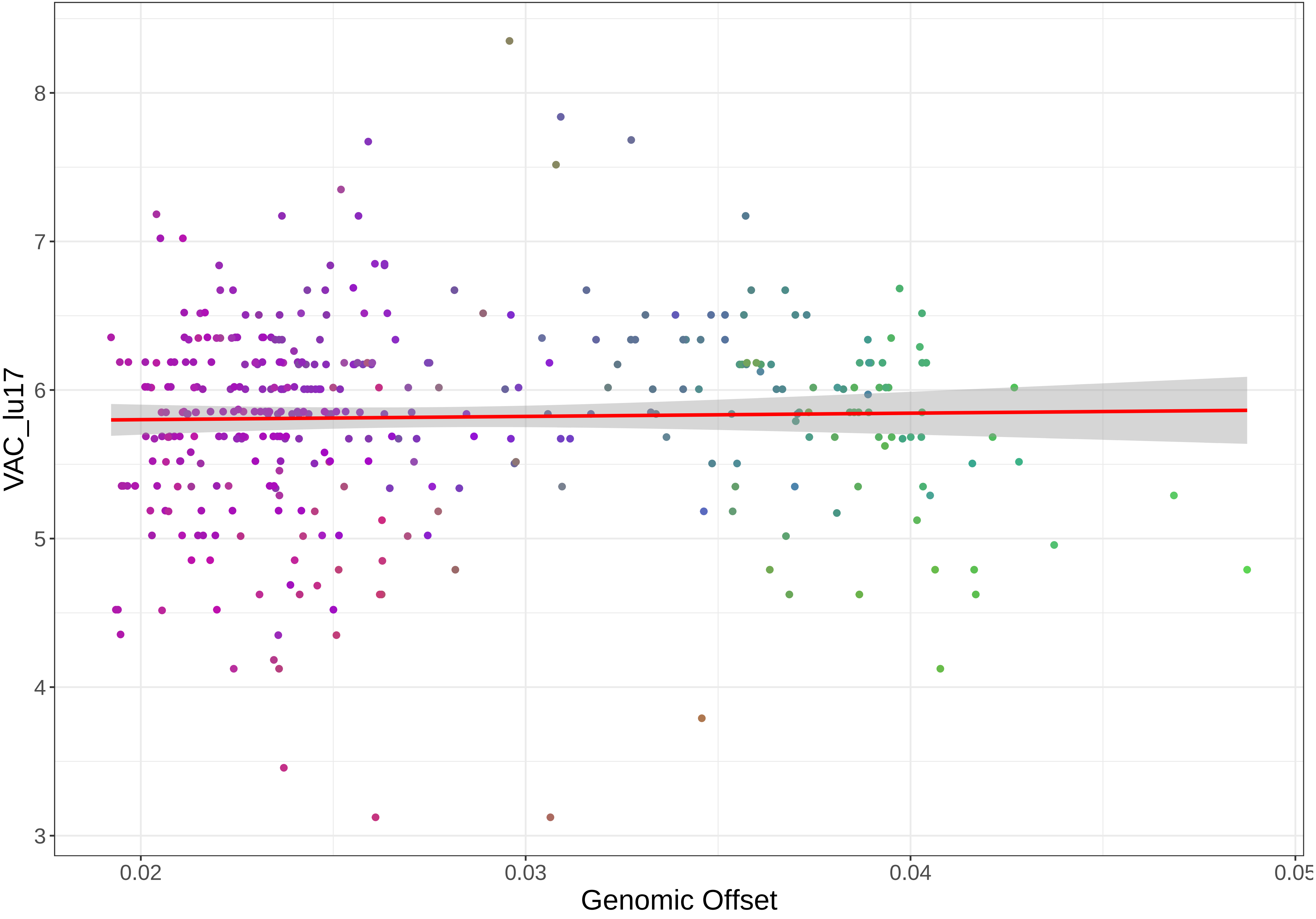

VAC_lu17
8
7
6
5
4
3
0.02
0.03
0.04
0.05
Genomic Offset

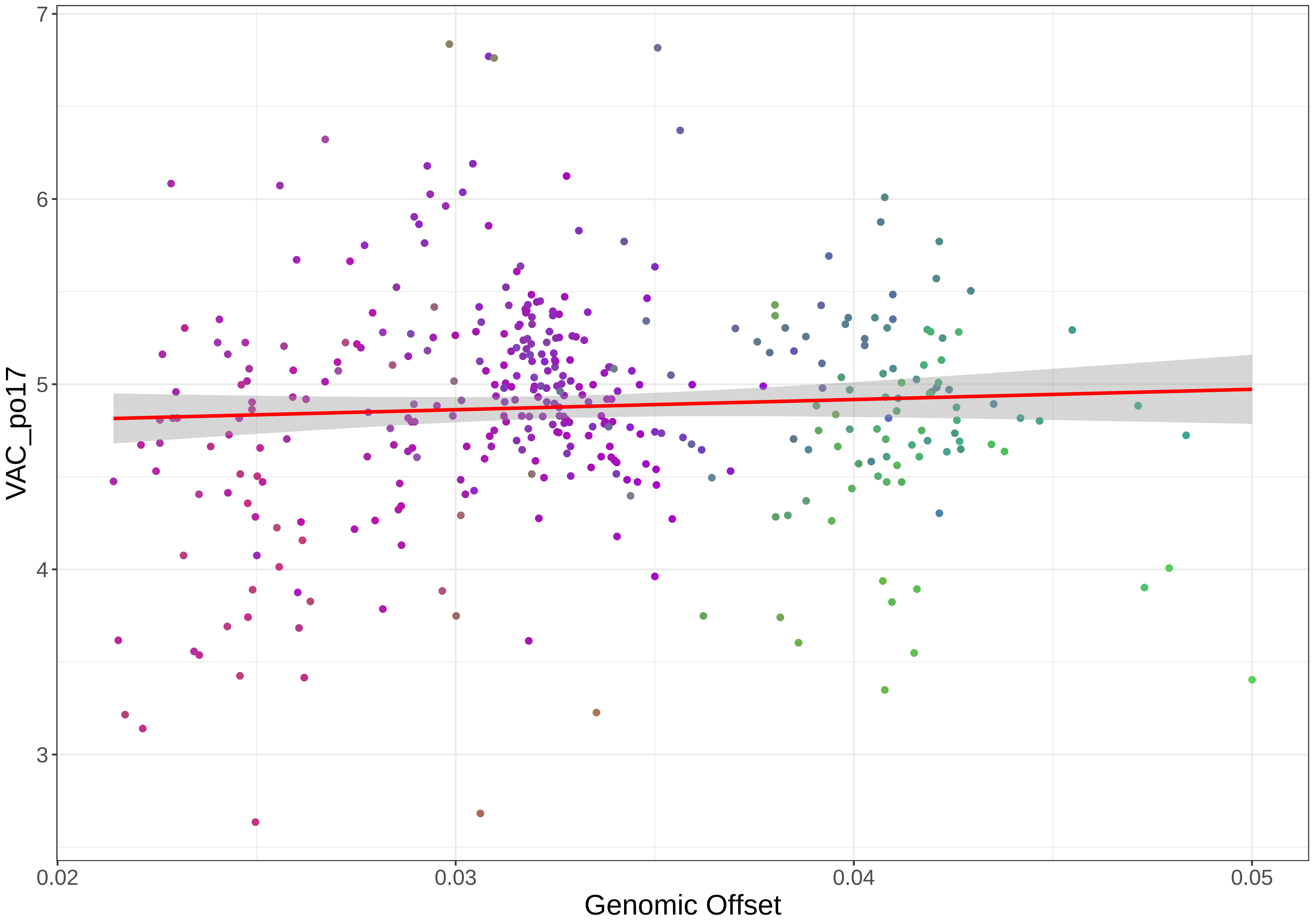

7
6
5
4
3
VAC_po17
0.02
0.03
0.04
0.05
Genomic Offset

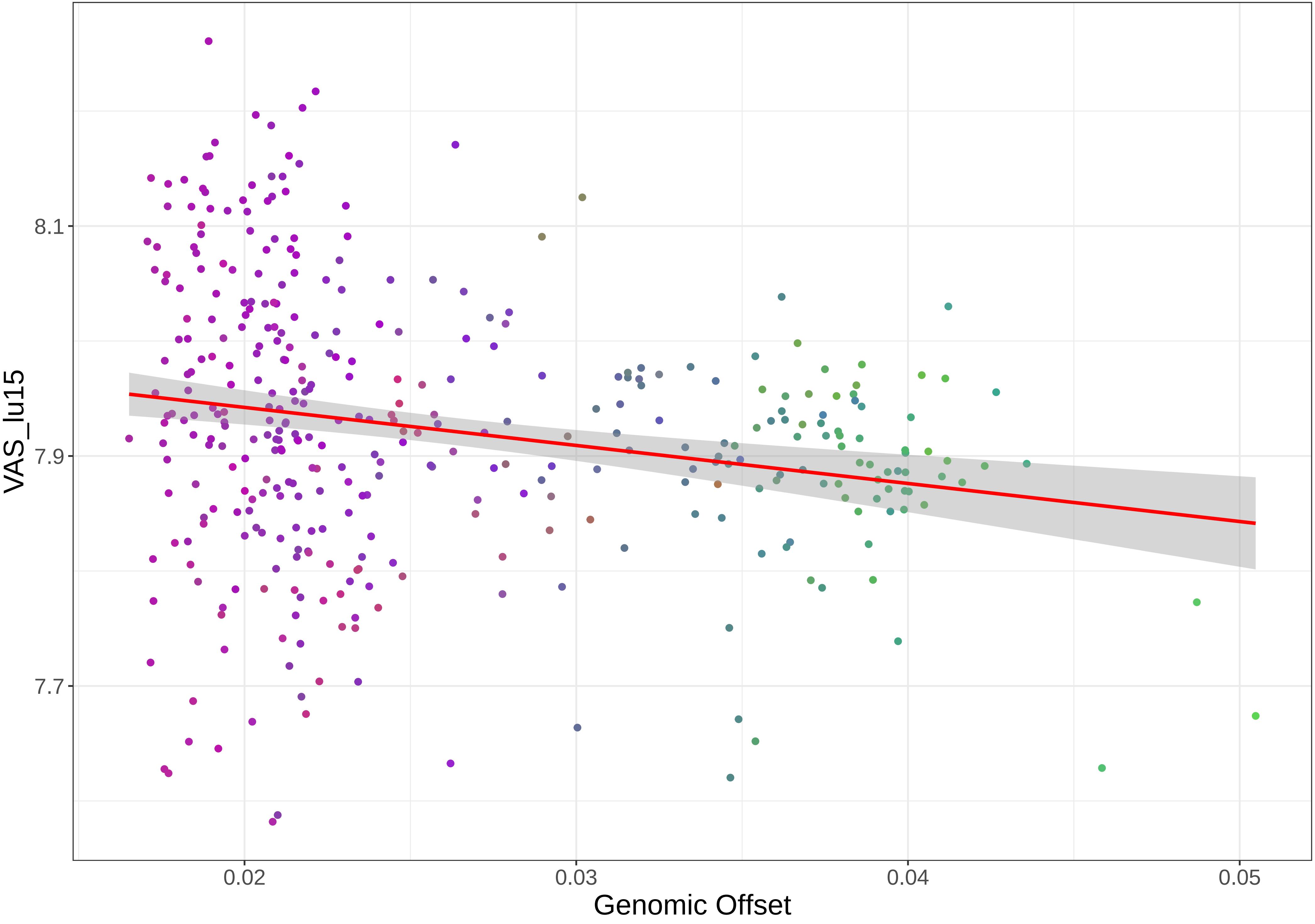
VAS_lu15
8.1
7.9
7.7
0.02
0.03
0.04
0.05
Genomic Offset

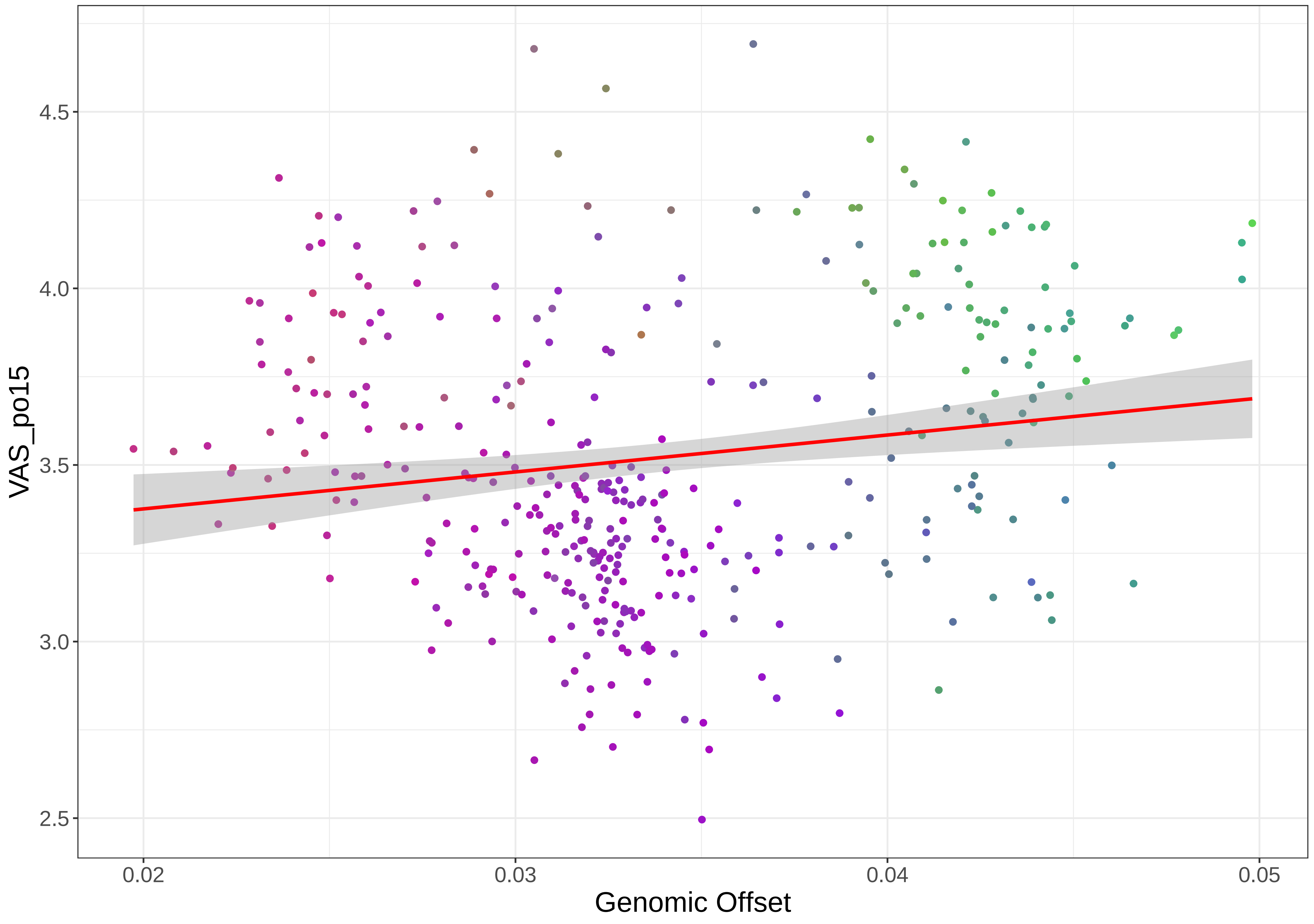

VAS_po15
4.5
4.0
3.5
3.0
2.5
0.02
0.03
0.04
0.05
Genomic Offset

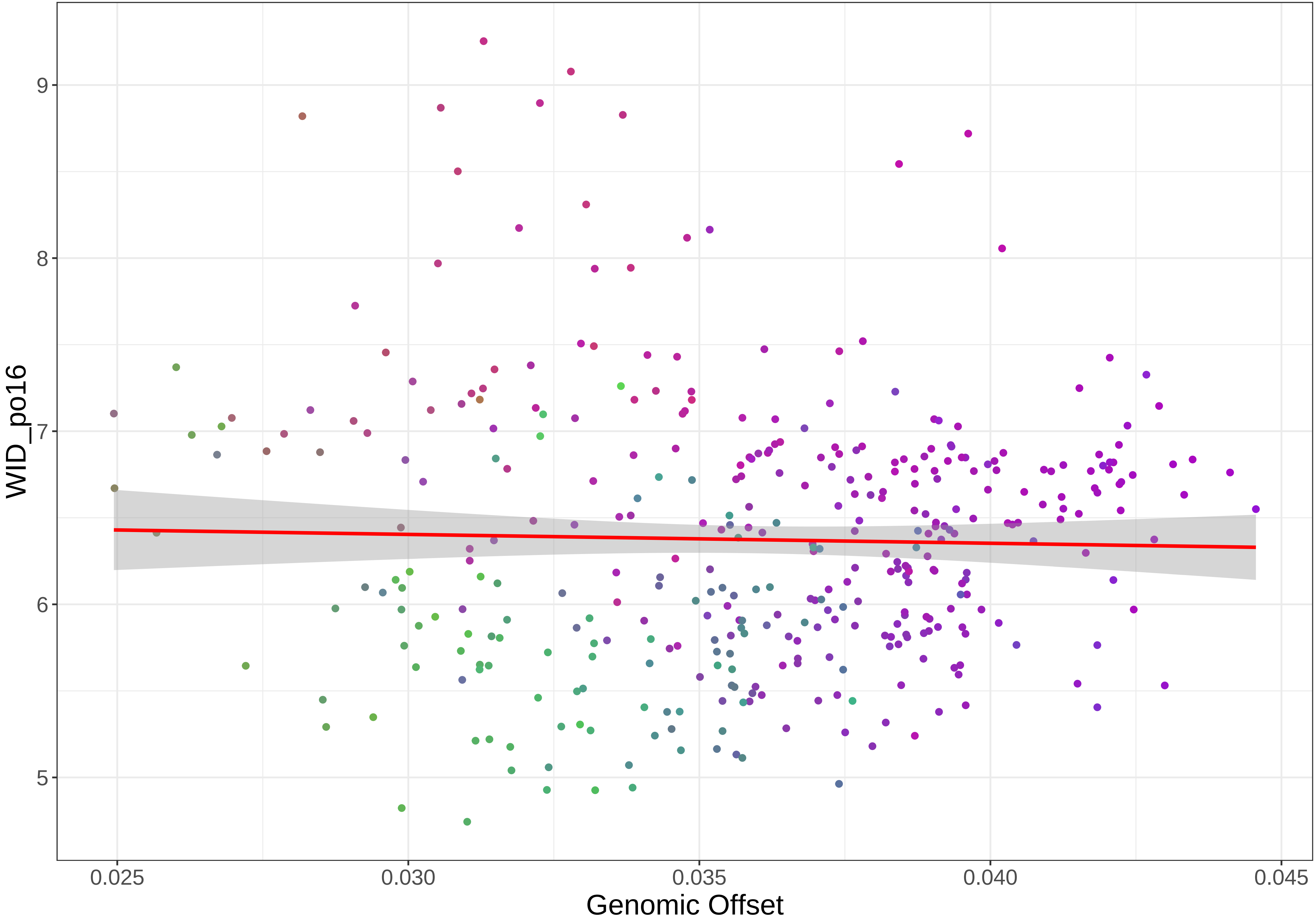

WID_po16
9
8
7
6
5
0.025
0.030
0.035
0.040
0.045
Genomic Offset

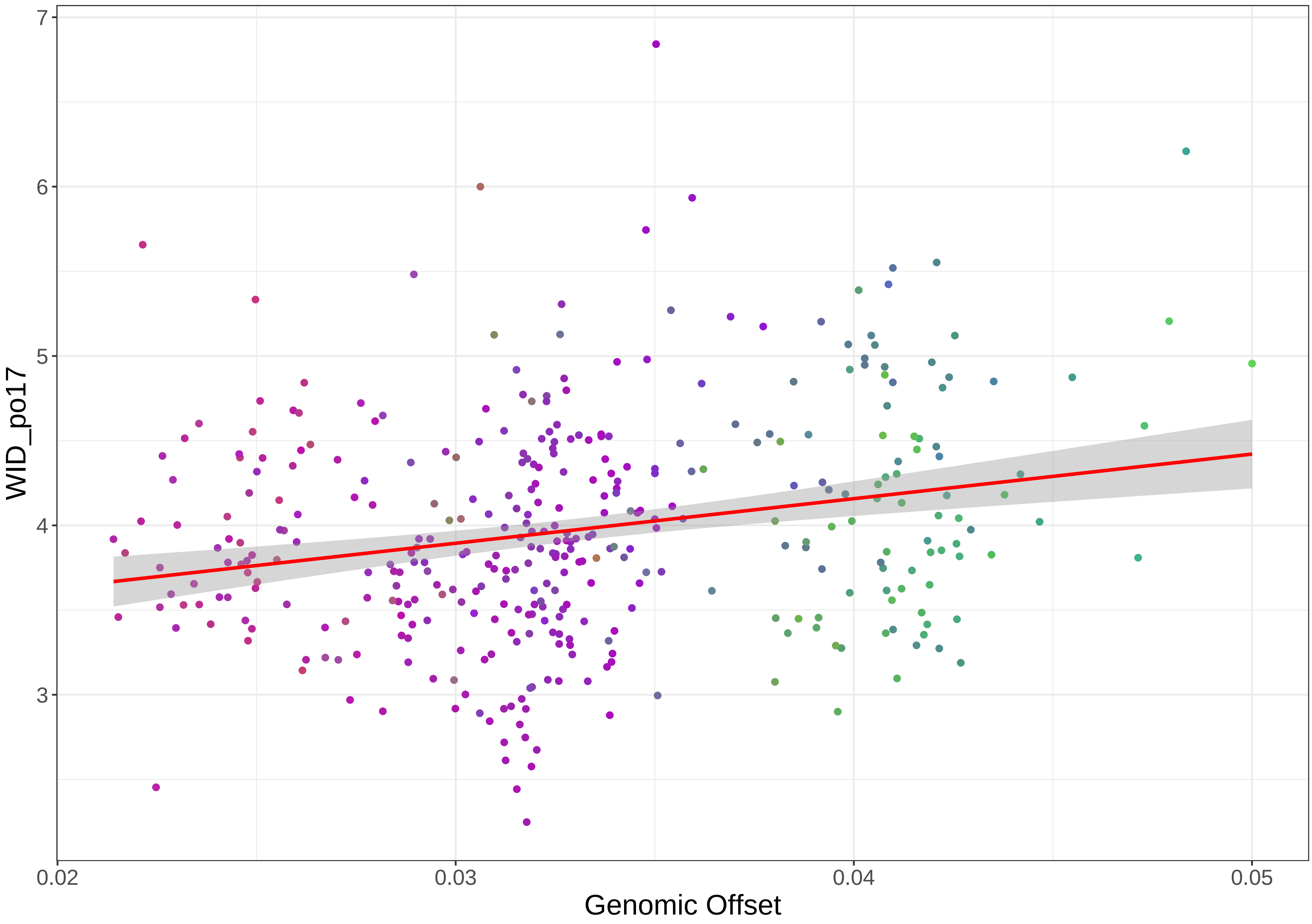

WID_po17
Genomic Offset
0.02
0.03
0.04
0.05
3
4
5
6
7

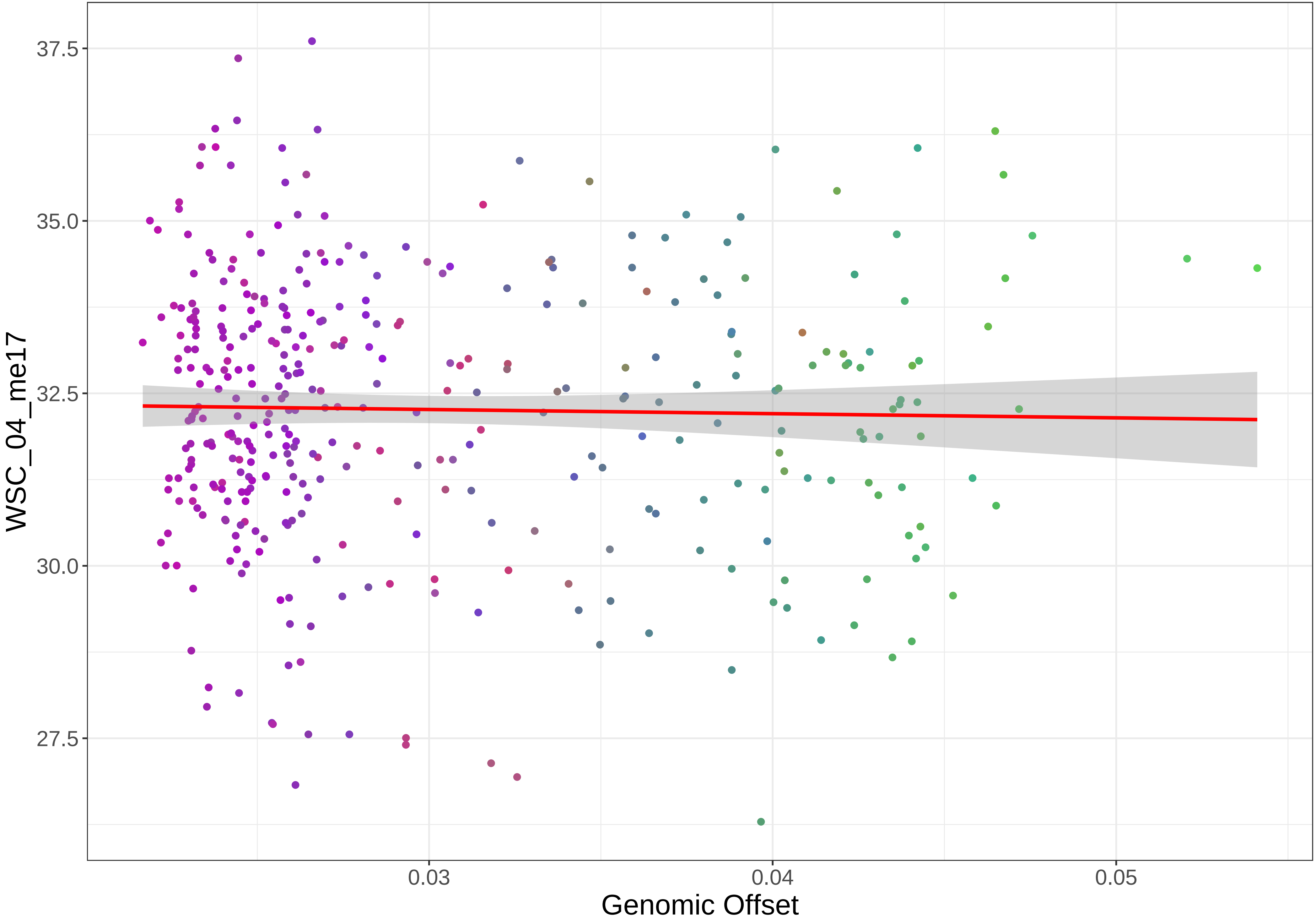

37.5
35.0
32.5
30.0
27.5
WSC_04_me17
0.03
0.04
0.05
Genomic Offset

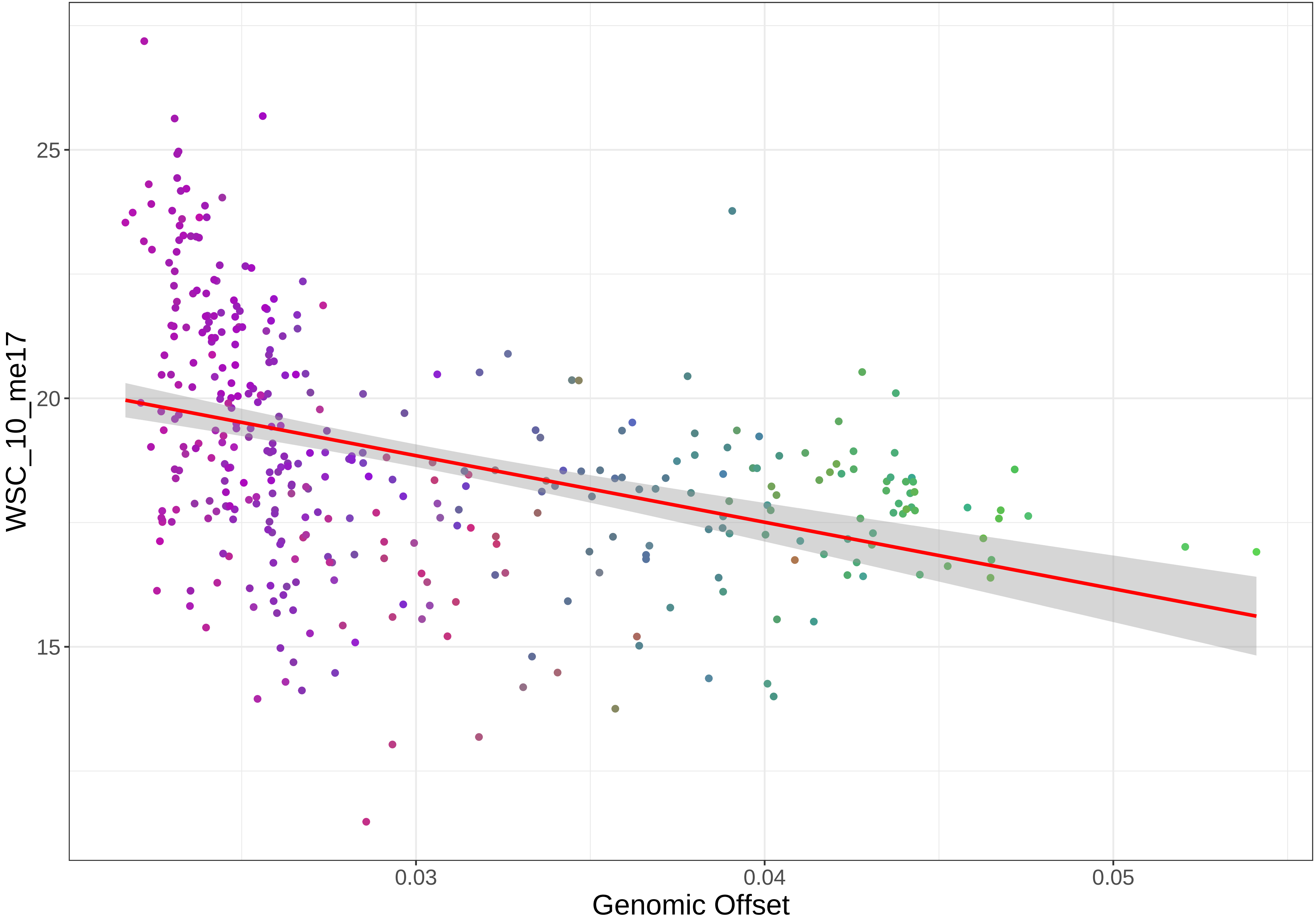

WSC_10_me17
25
20
15
0.03
0.04
0.05
Genomic Offset
