## SupplementaryFigS2 for "How robust are genomic offset predictions to methodological choices? Insights from perennial ryegrass"

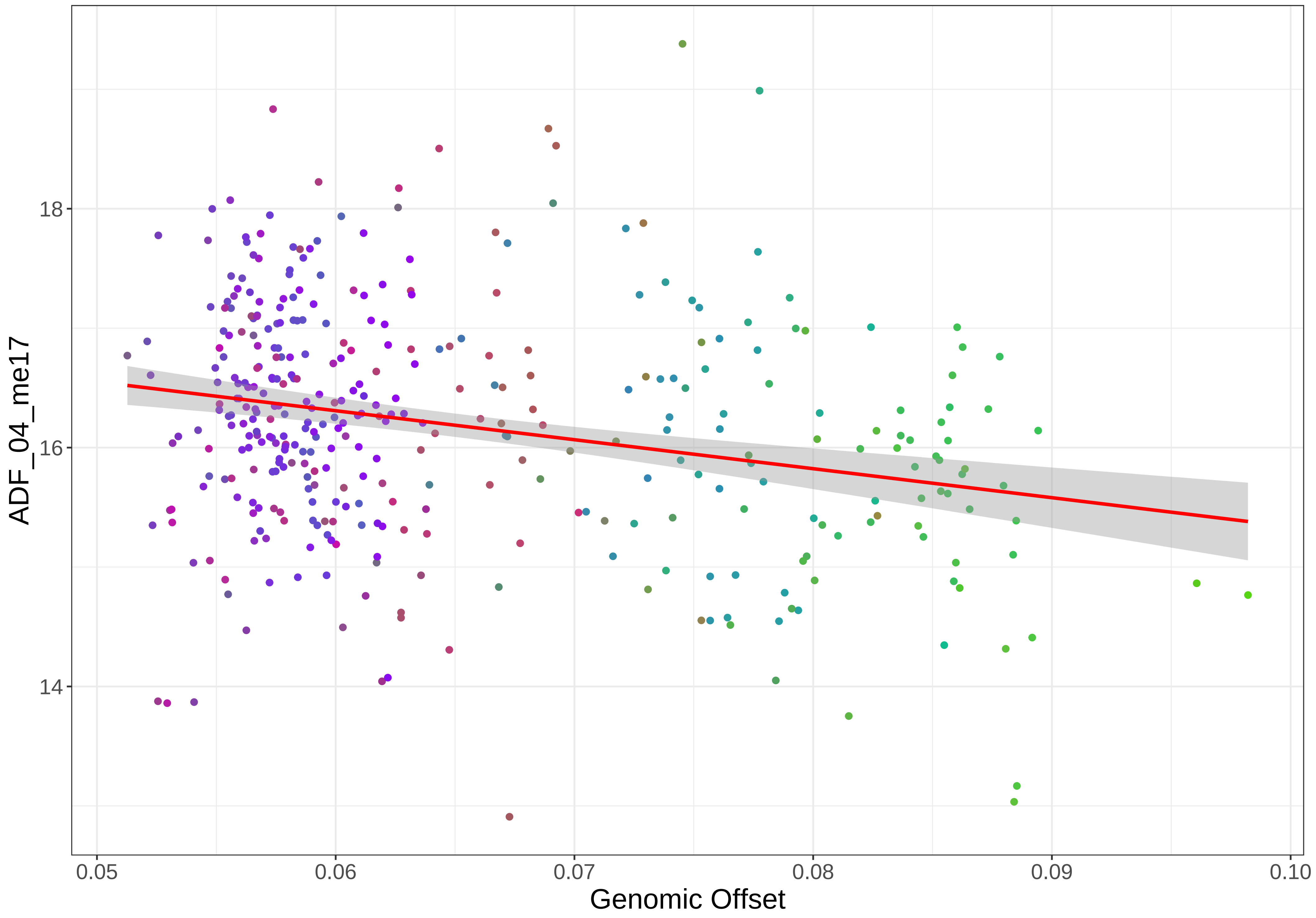

ADF_04_me17
18
16
14
0.05
0.06
0.07
0.08
0.09
0.10
Genomic Offset

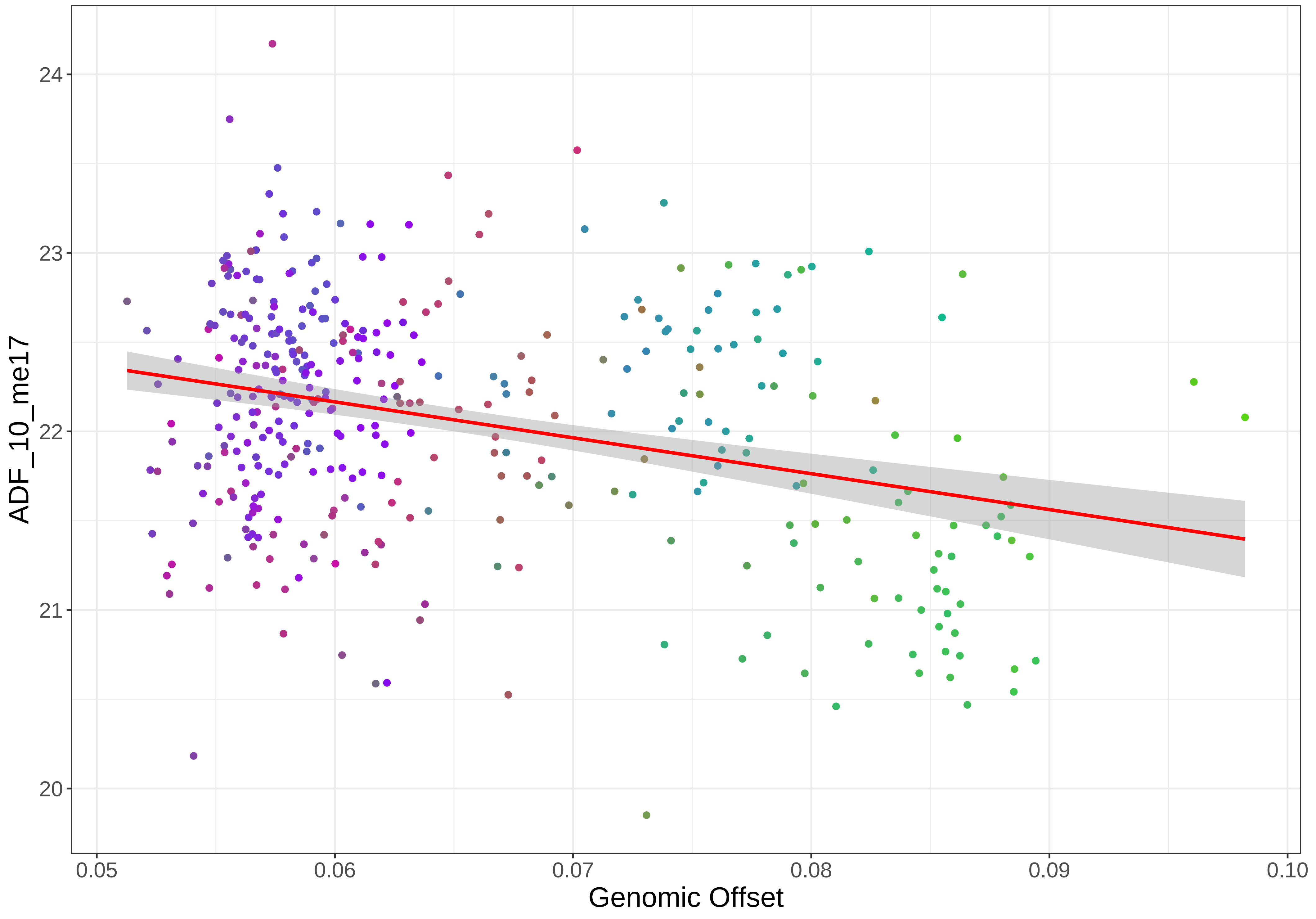

ADF_10_me17
24
23
22
21
20
0.05
0.06
0.07
0.08
0.09
0.10
Genomic Offset

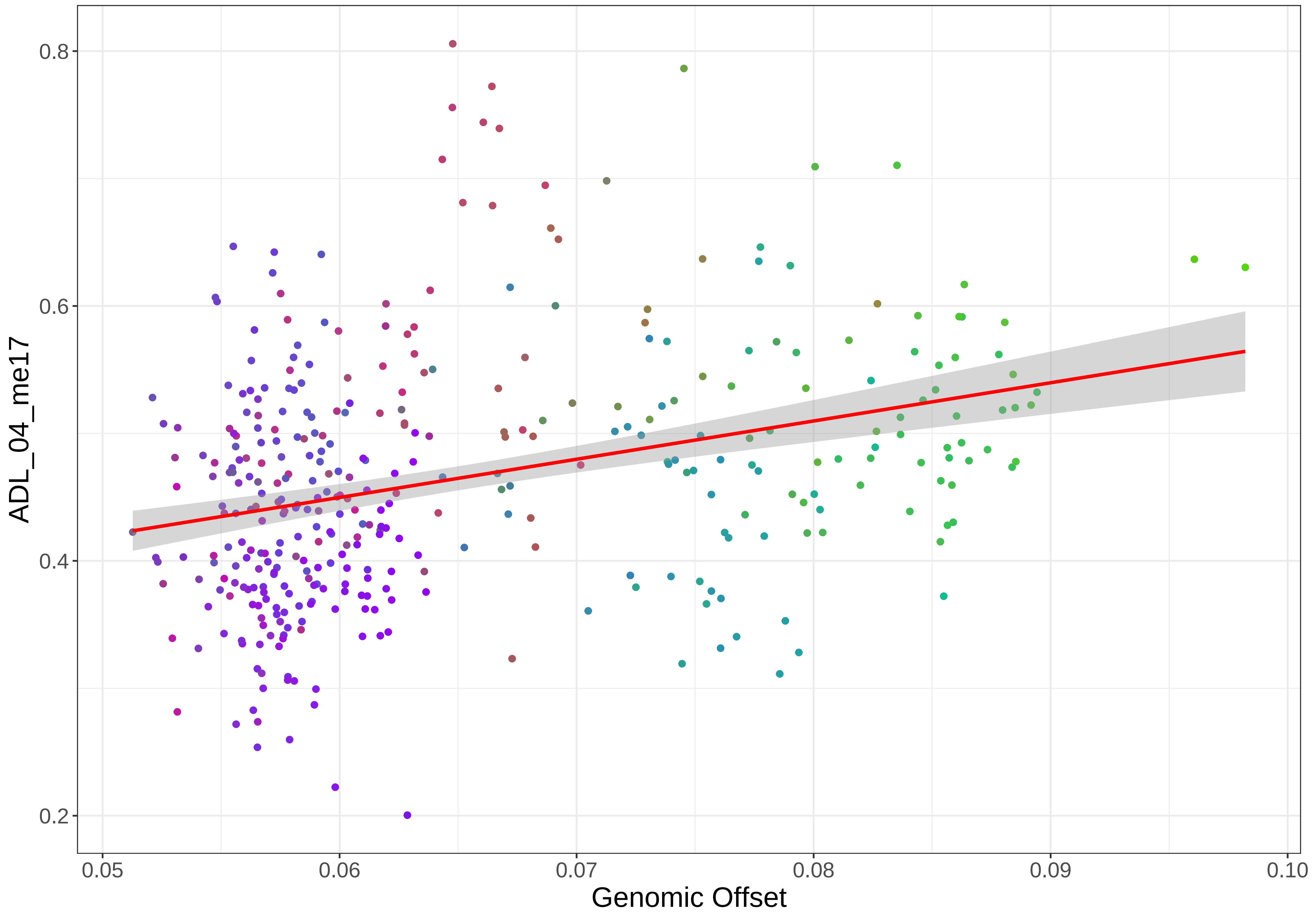

0.8
0.6
0.4
0.2
ADL_04_me17
0.05
0.06
0.07
0.08
0.09
0.10
Genomic Offset

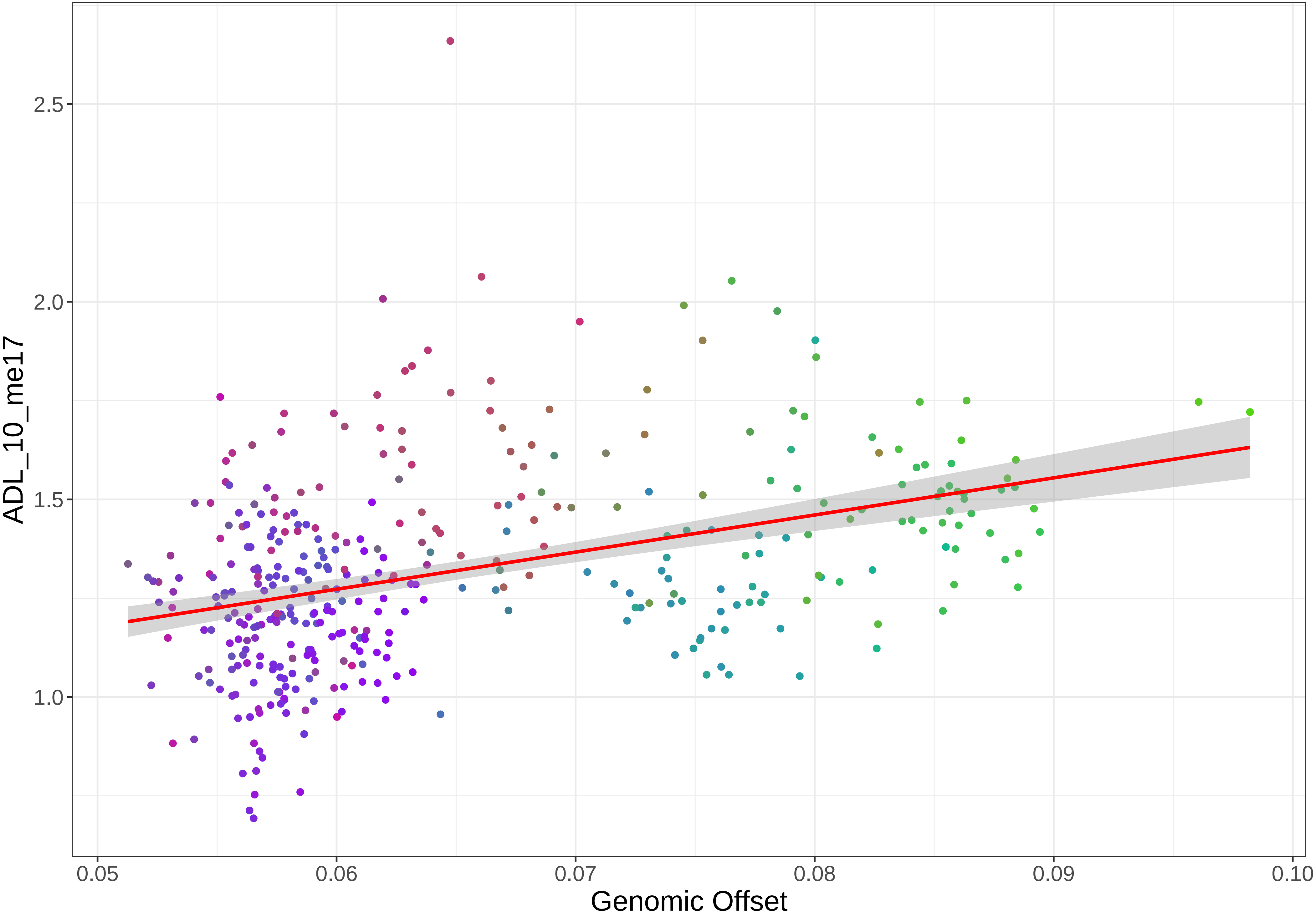

ADL_10_me17
2.5
2.0
1.5
1.0
0.05
0.06
0.07
0.08
0.09
0.10
Genomic Offset

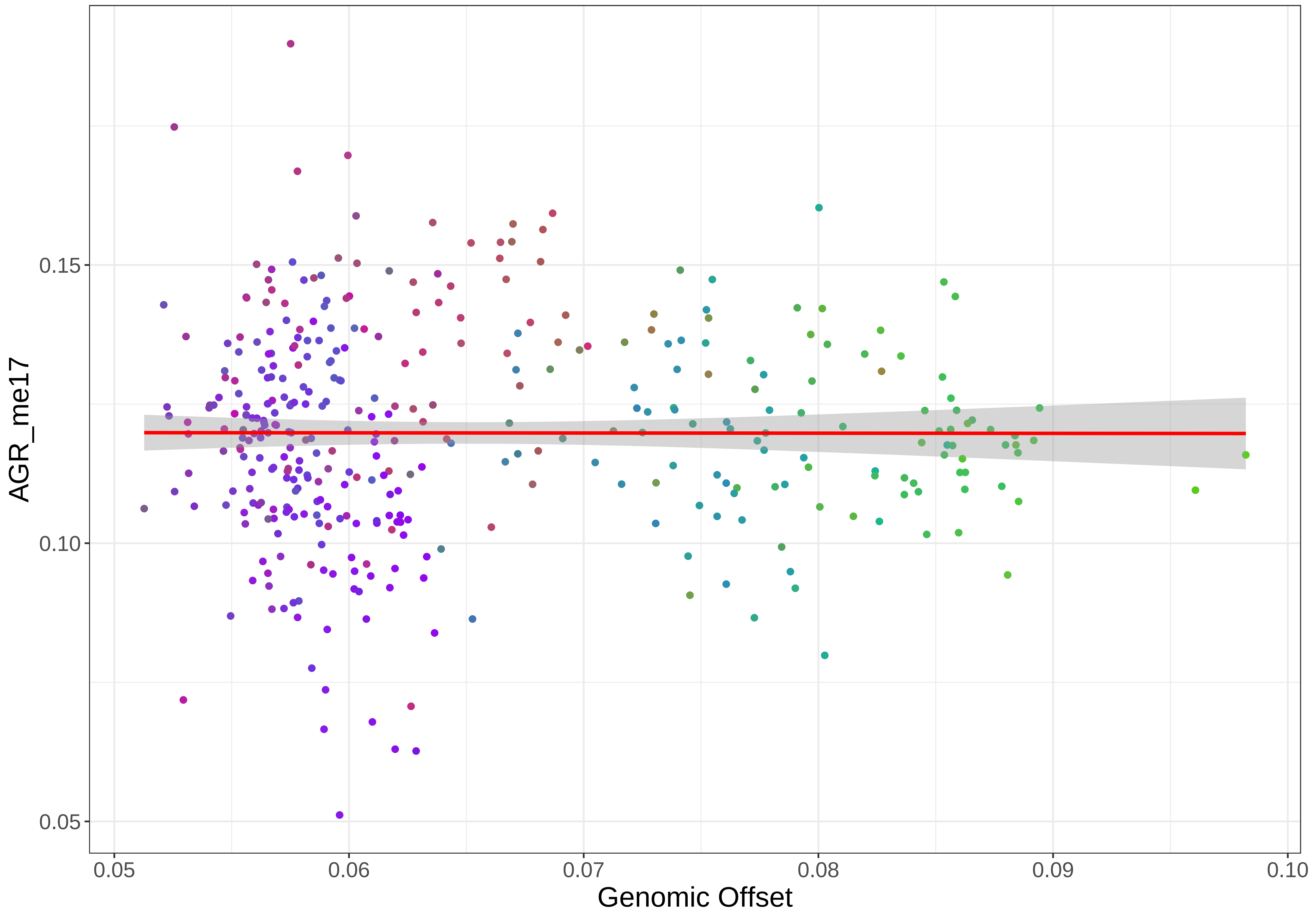

AGR_me17
0.15
0.10
0.05
0.05
0.06
0.07
0.08
0.09
0.10
Genomic Offset

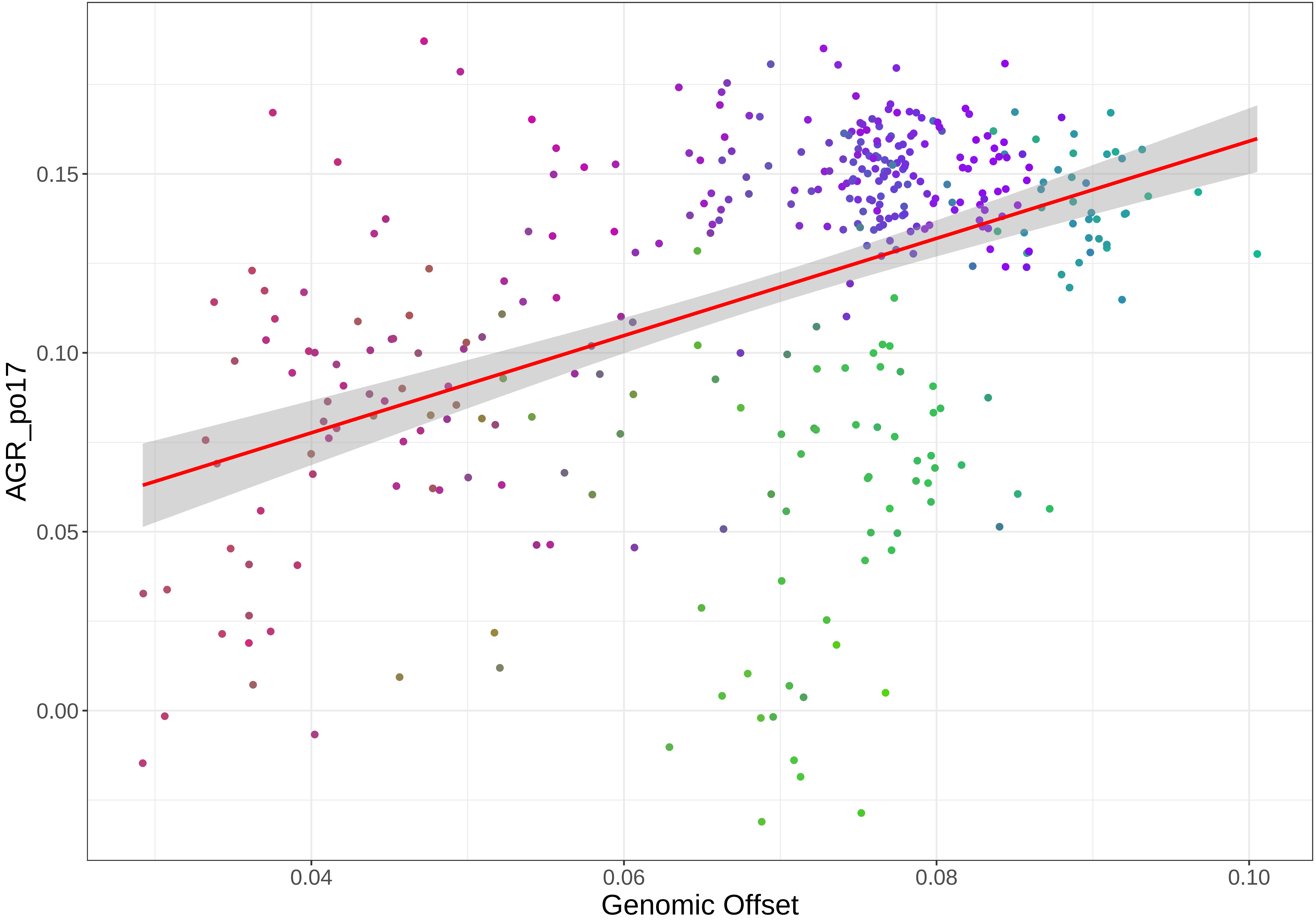

AGR_po17
0.15
0.10
0.05
0.00
0.04
0.06
0.08
0.10
Genomic Offset

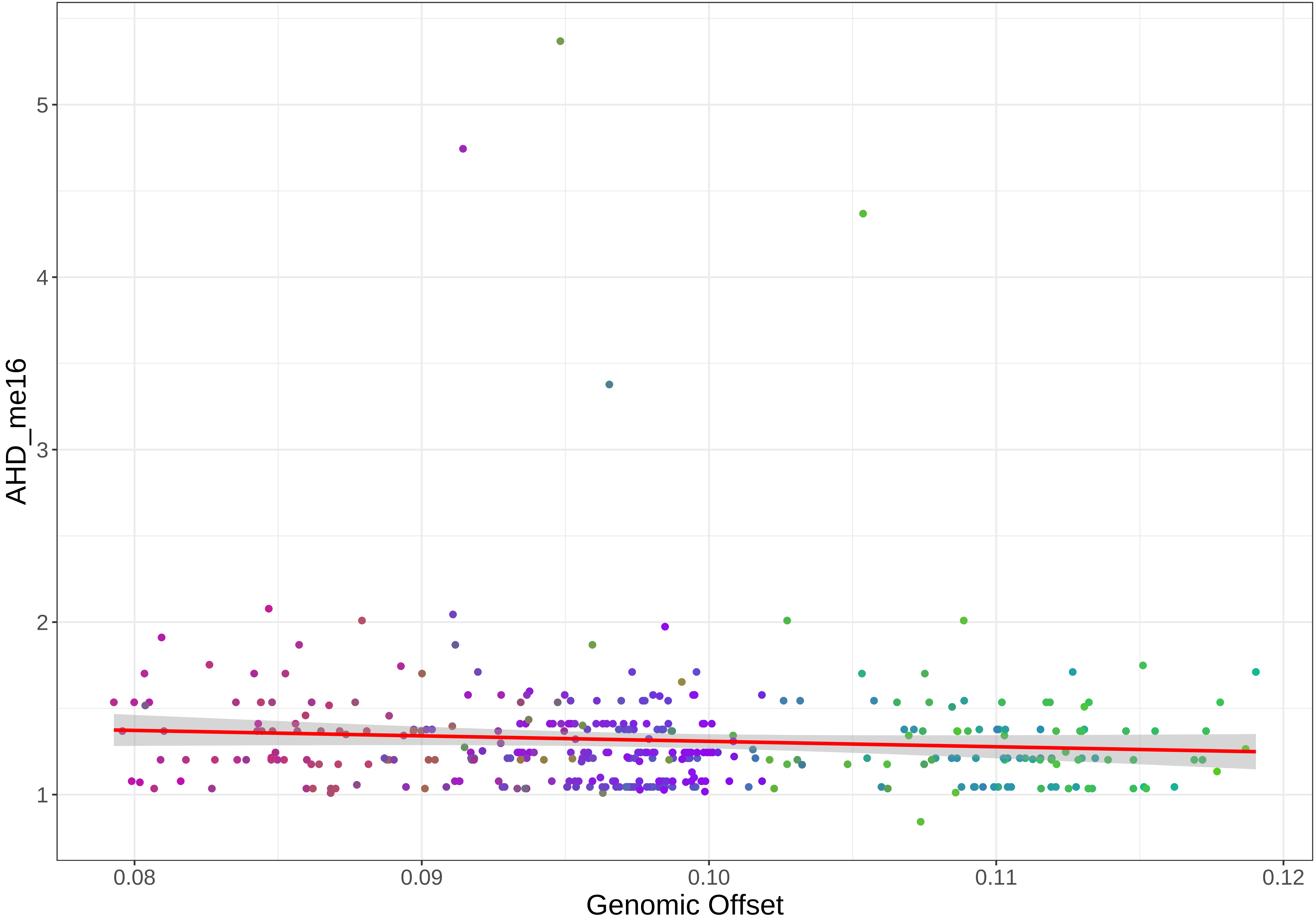

AHD_me16
1
2
3
4
5
0.08
0.09
0.10
0.11
0.12
Genomic Offset

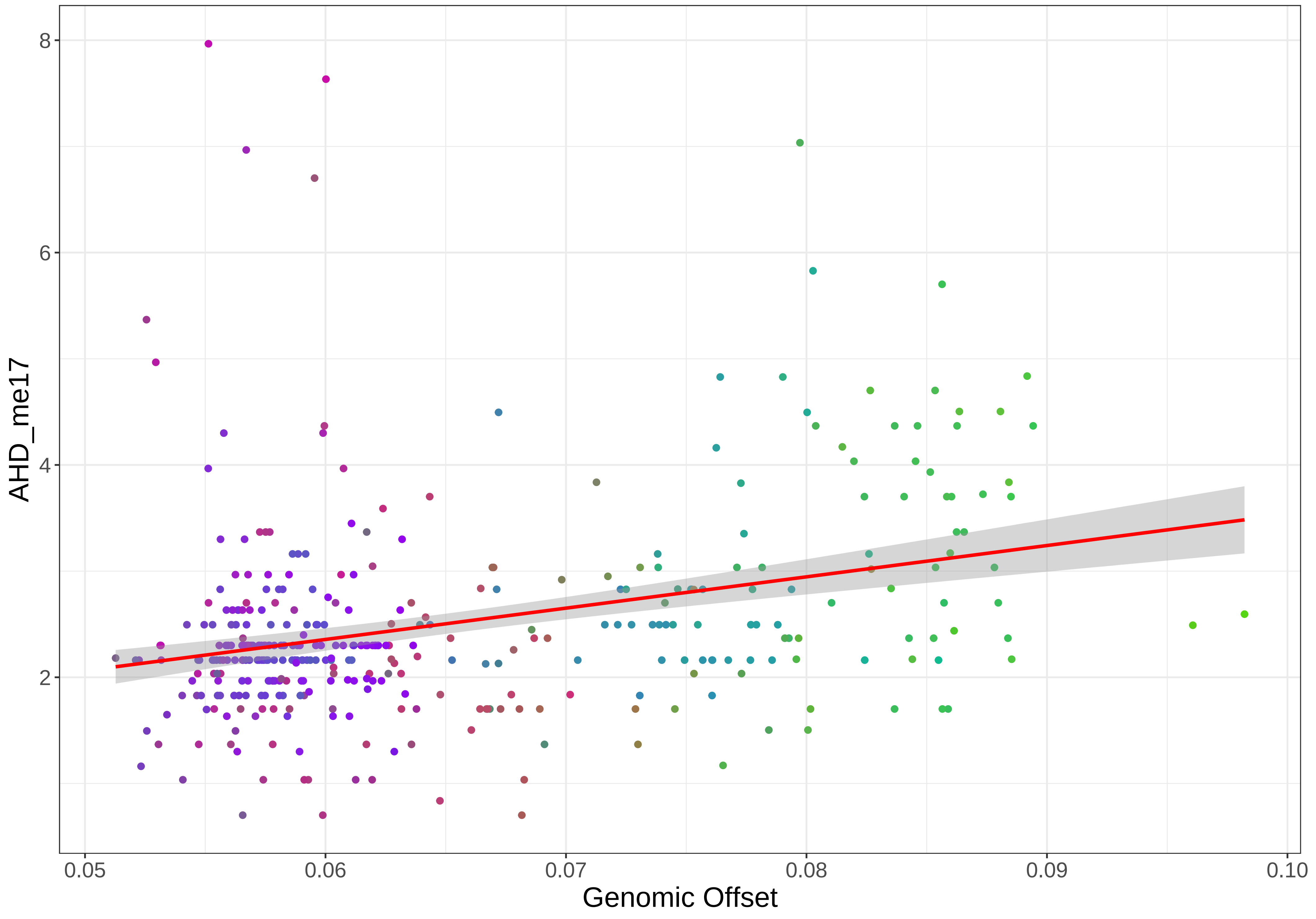

8
6
4
2
AHD_me17
0.05
0.06
0.07
0.08
0.09
0.10
Genomic Offset

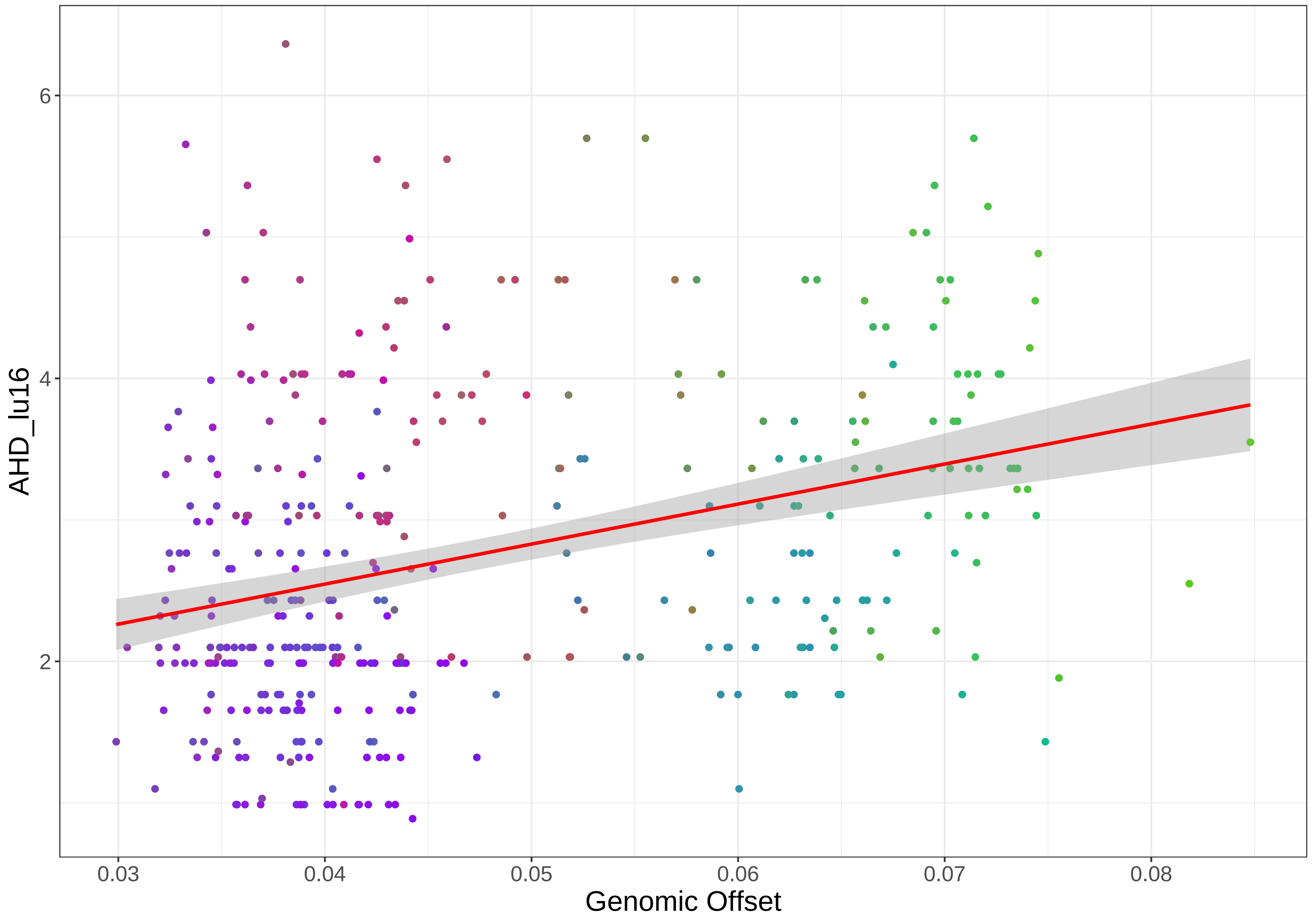

AHD_lu16
Genomic Offset
6
4
2
0.03
0.04
0.05
0.06
0.07
0.08

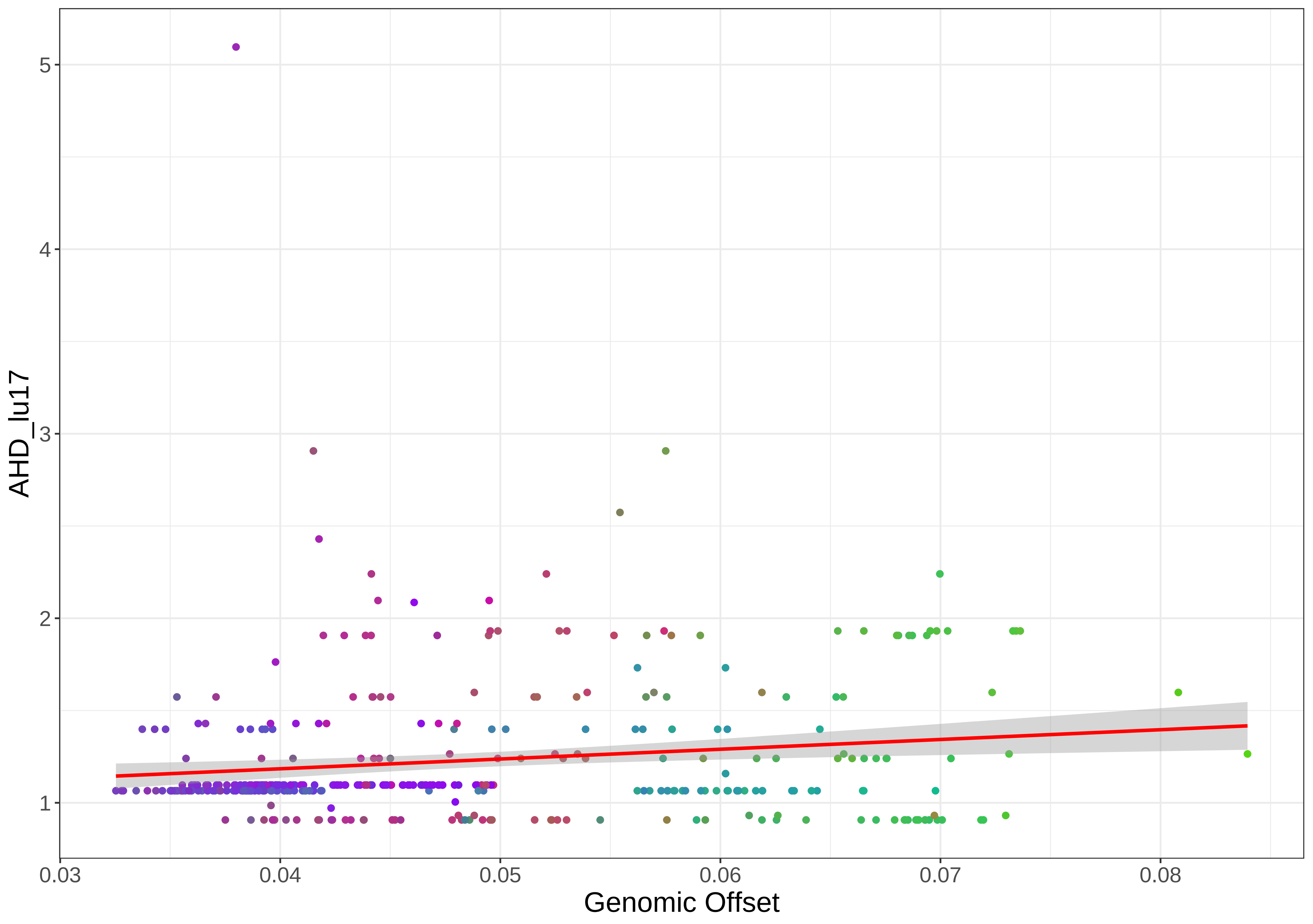

AHD_lu17
5
4
3
2
1
0.03
0.04
0.05
0.06
0.07
0.08
Genomic Offset

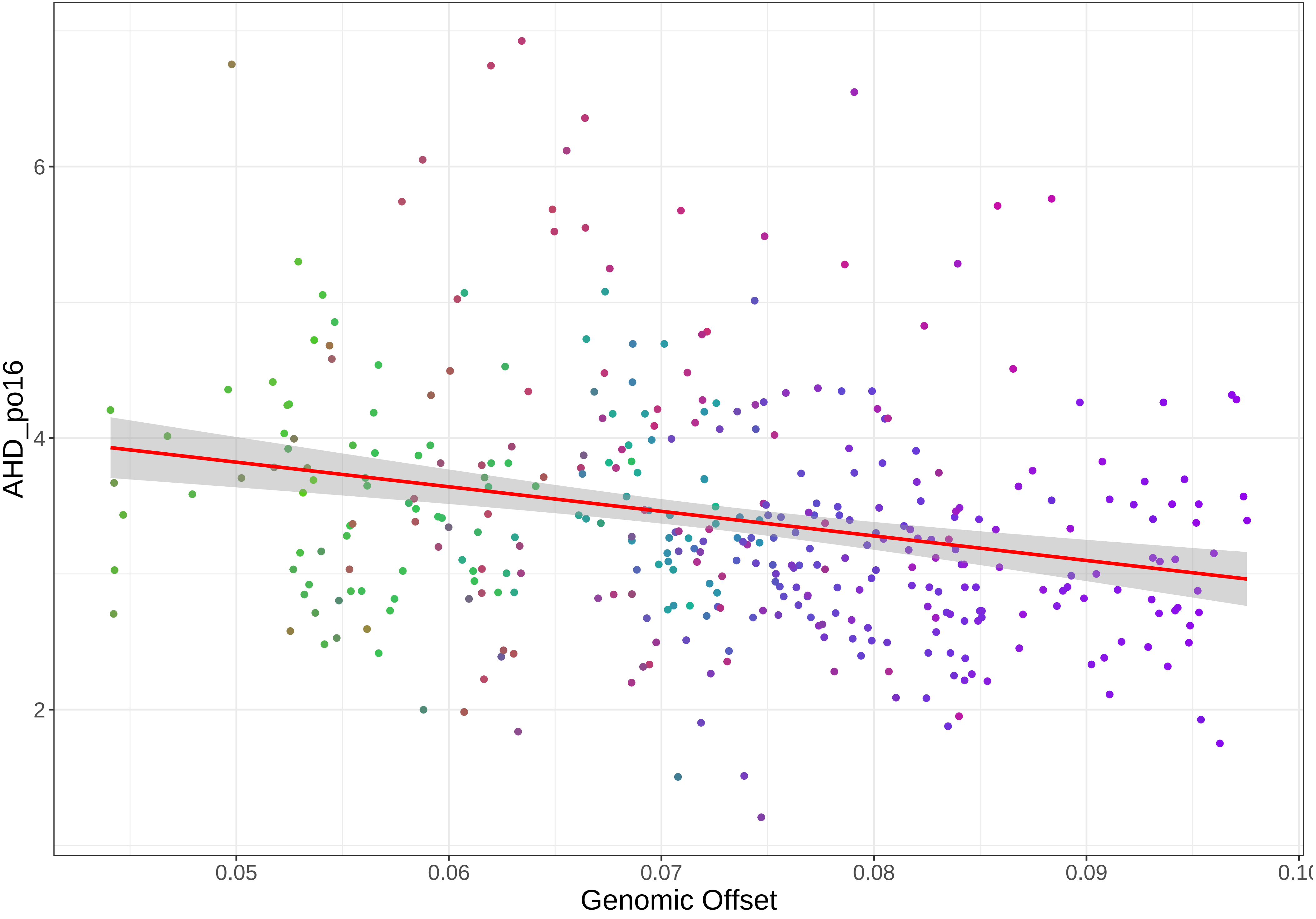

AHD_po16
6
4
2
0.05
0.06
0.07
0.08
0.09
Genomic Offset

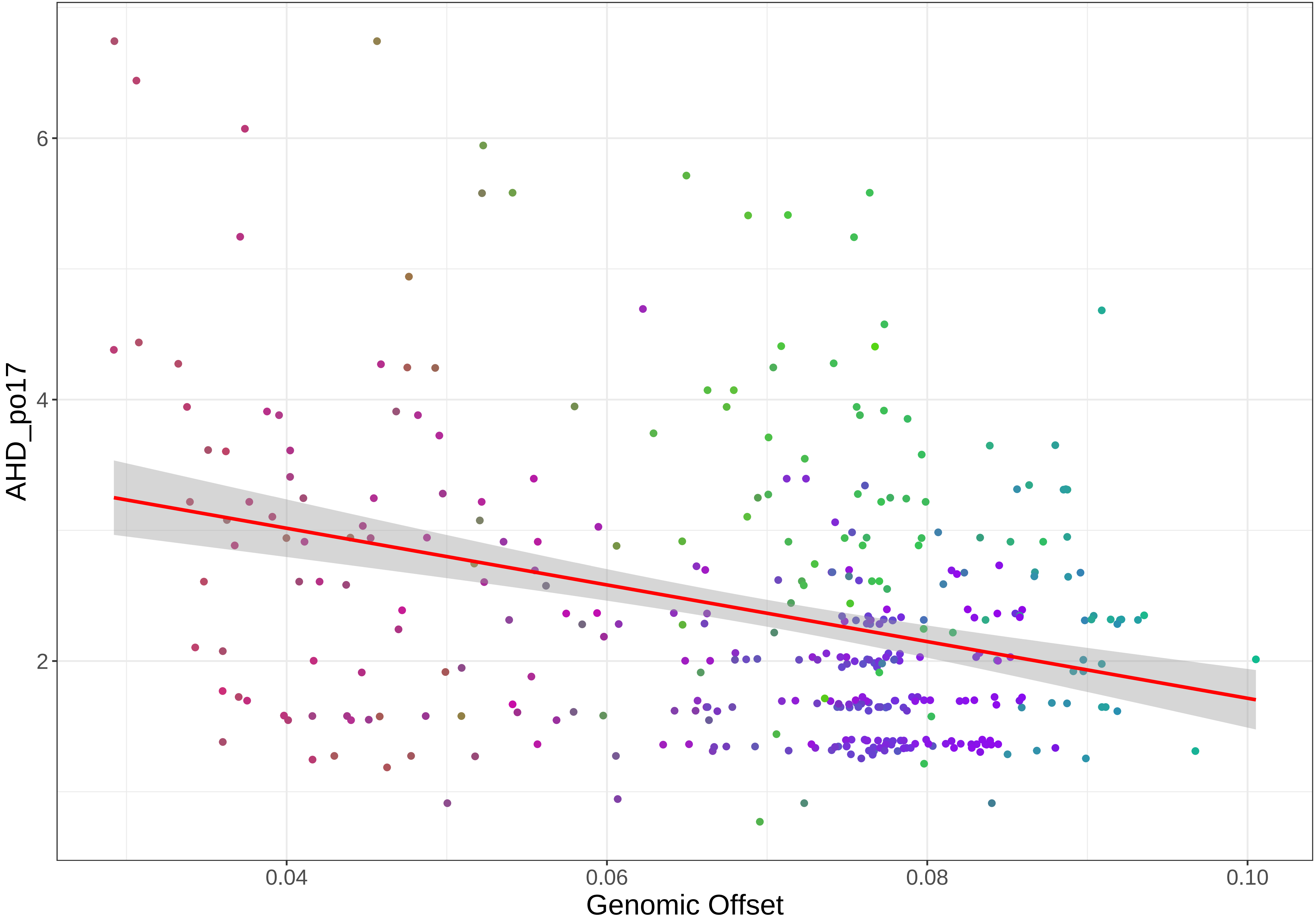

AHD_po17
6
4
2
0.04
0.06
0.08
0.10
Genomic Offset

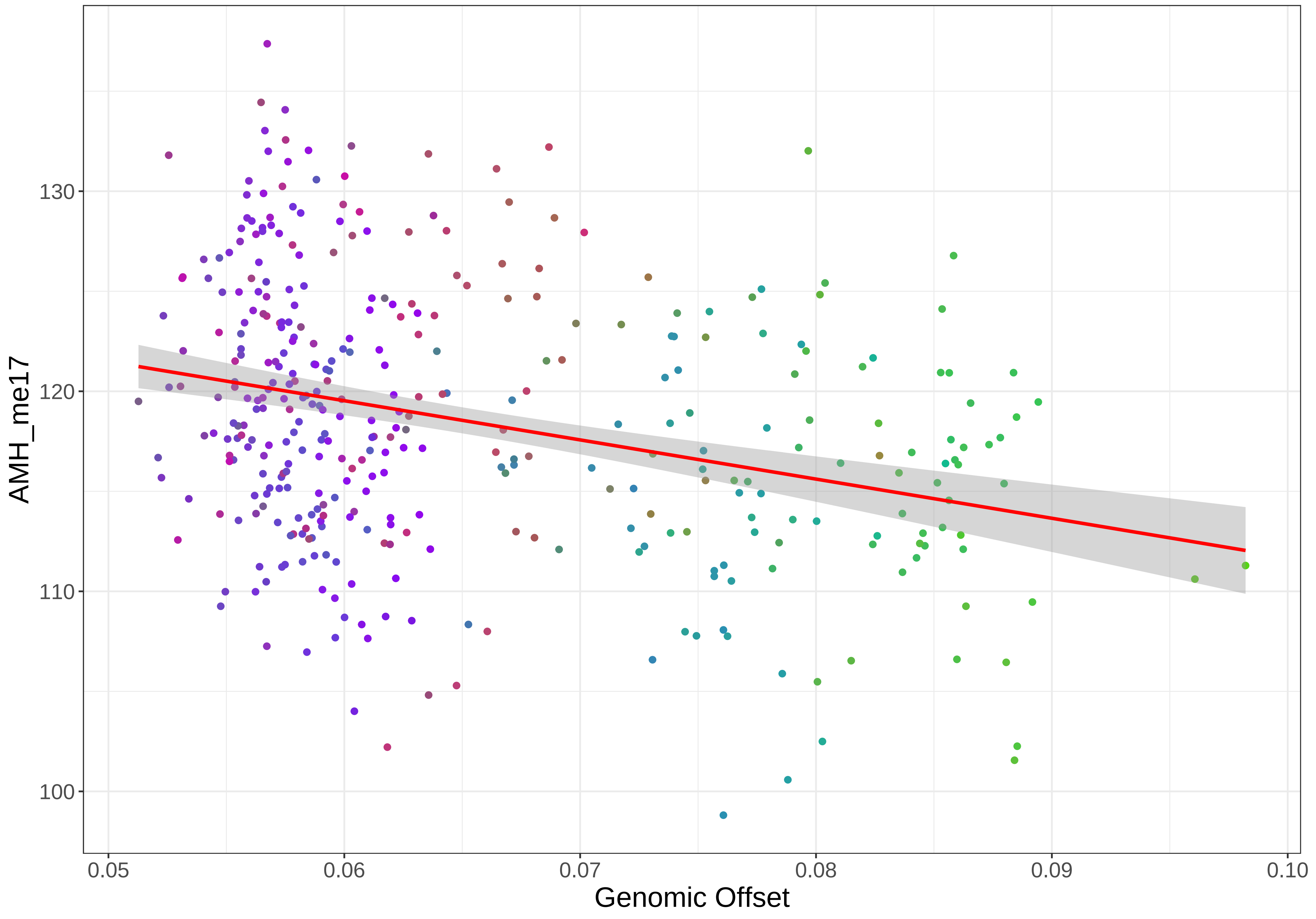
AMH_me17
130
120
110
100
0.05
0.06
0.07
0.08
0.09
0.10
Genomic Offset

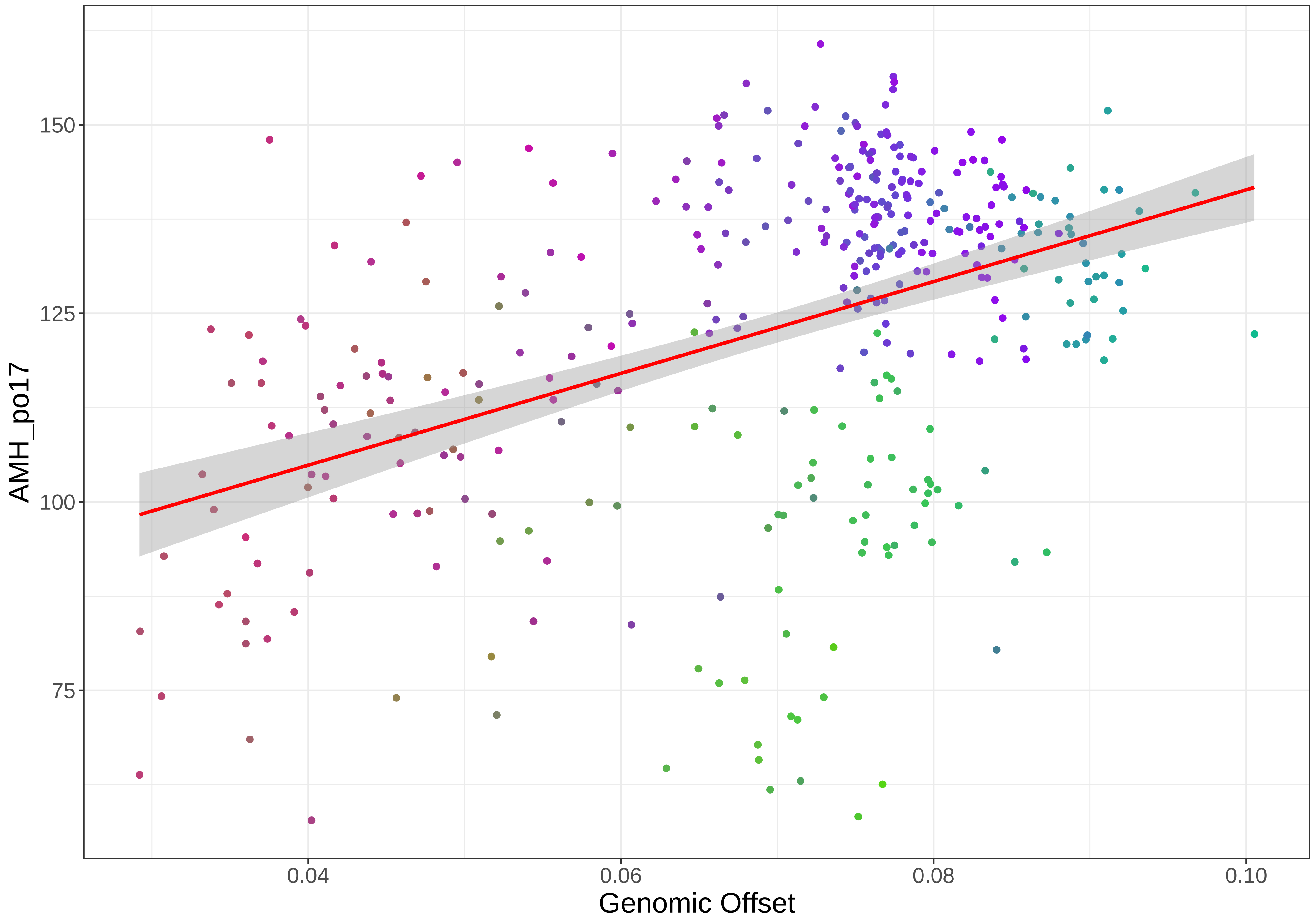

150
125
100
75
AMH_po17
0.04
0.06
0.08
0.10
Genomic Offset

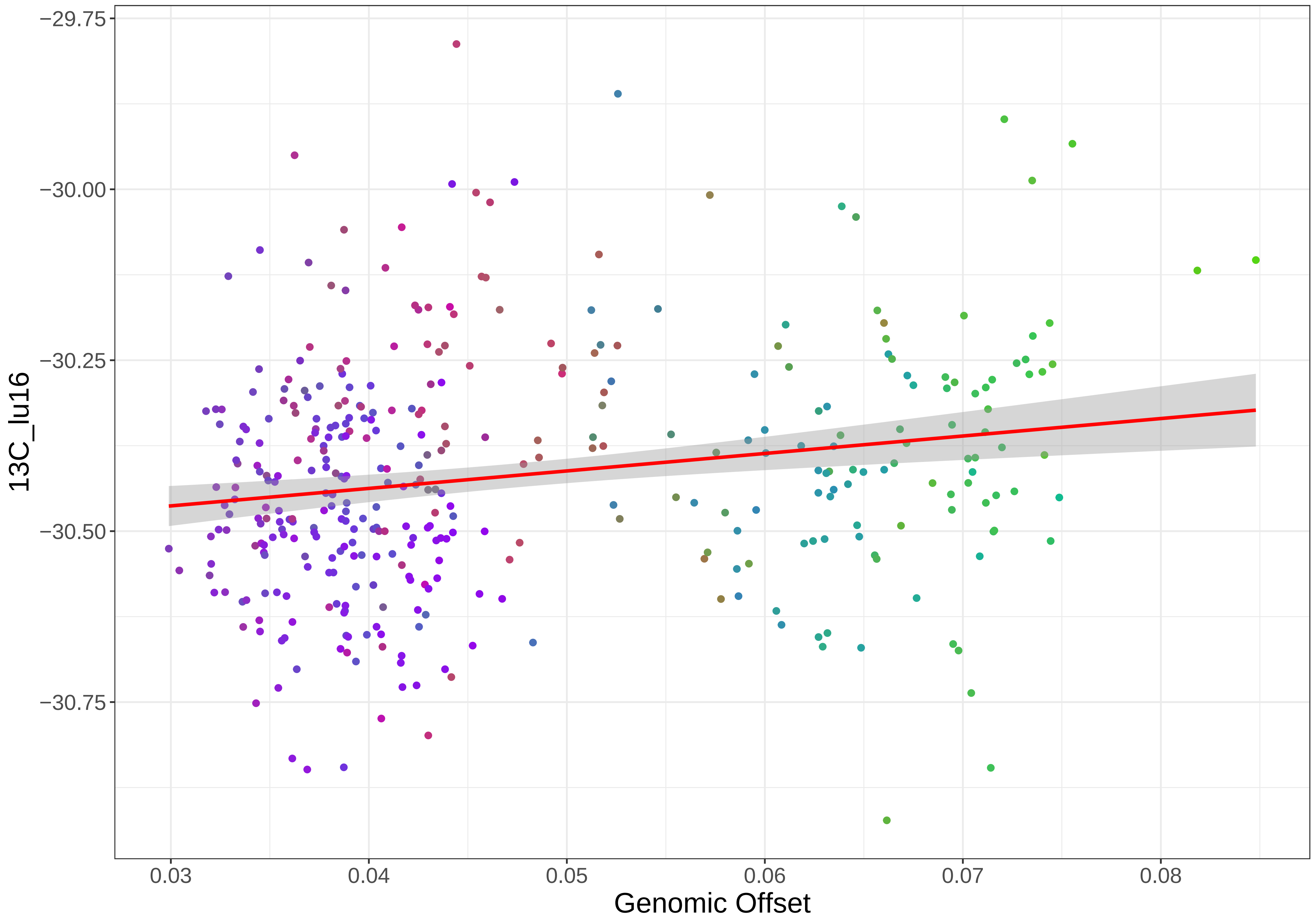

−29.75
−30.00
−30.25
−30.50
−30.75
13C_lu16
0.03
0.04
0.05
0.06
0.07
0.08
Genomic Offset

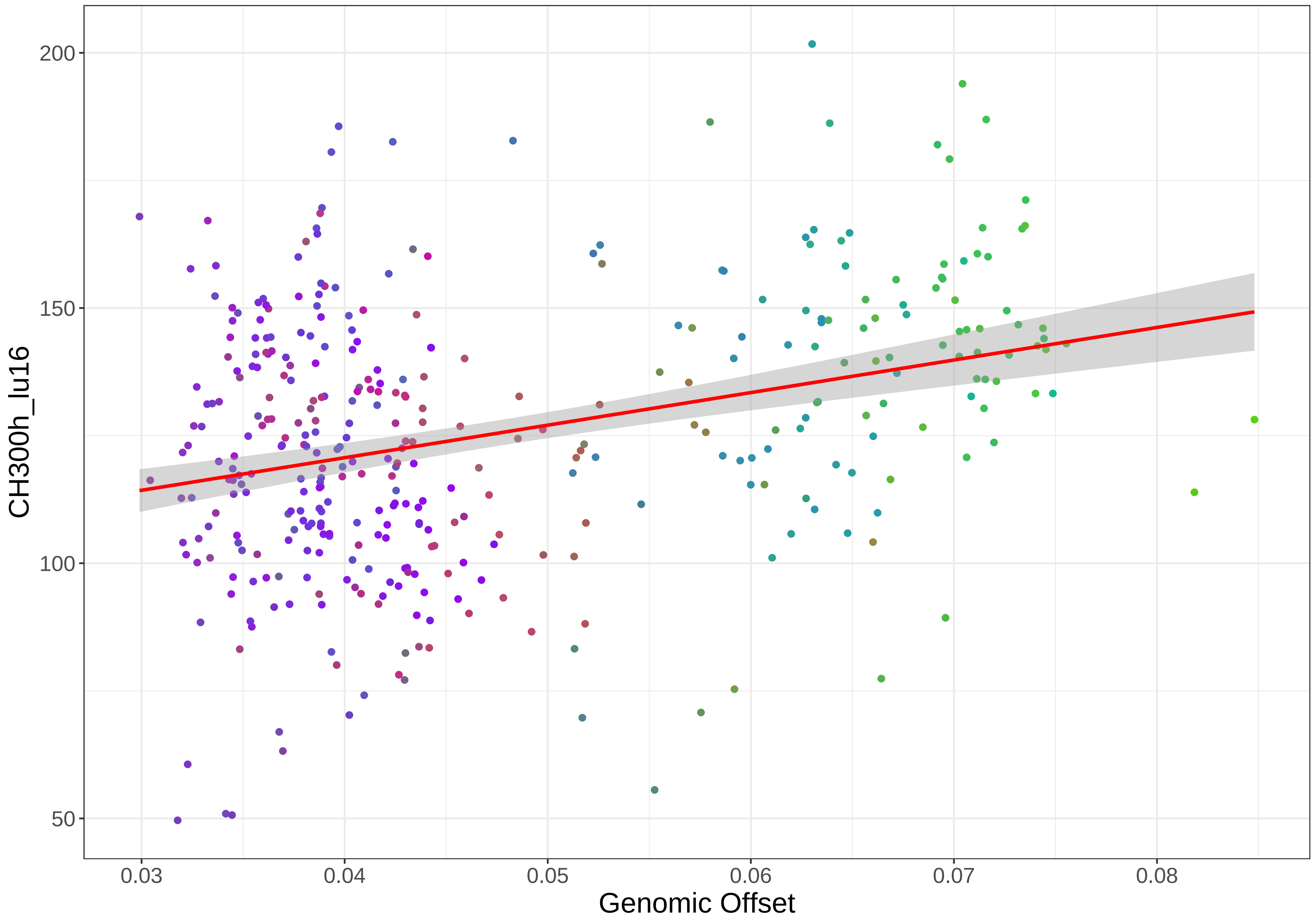

CH300h_lu16
200
150
100
50
0.03
0.04
0.05
0.06
0.07
0.08
Genomic Offset

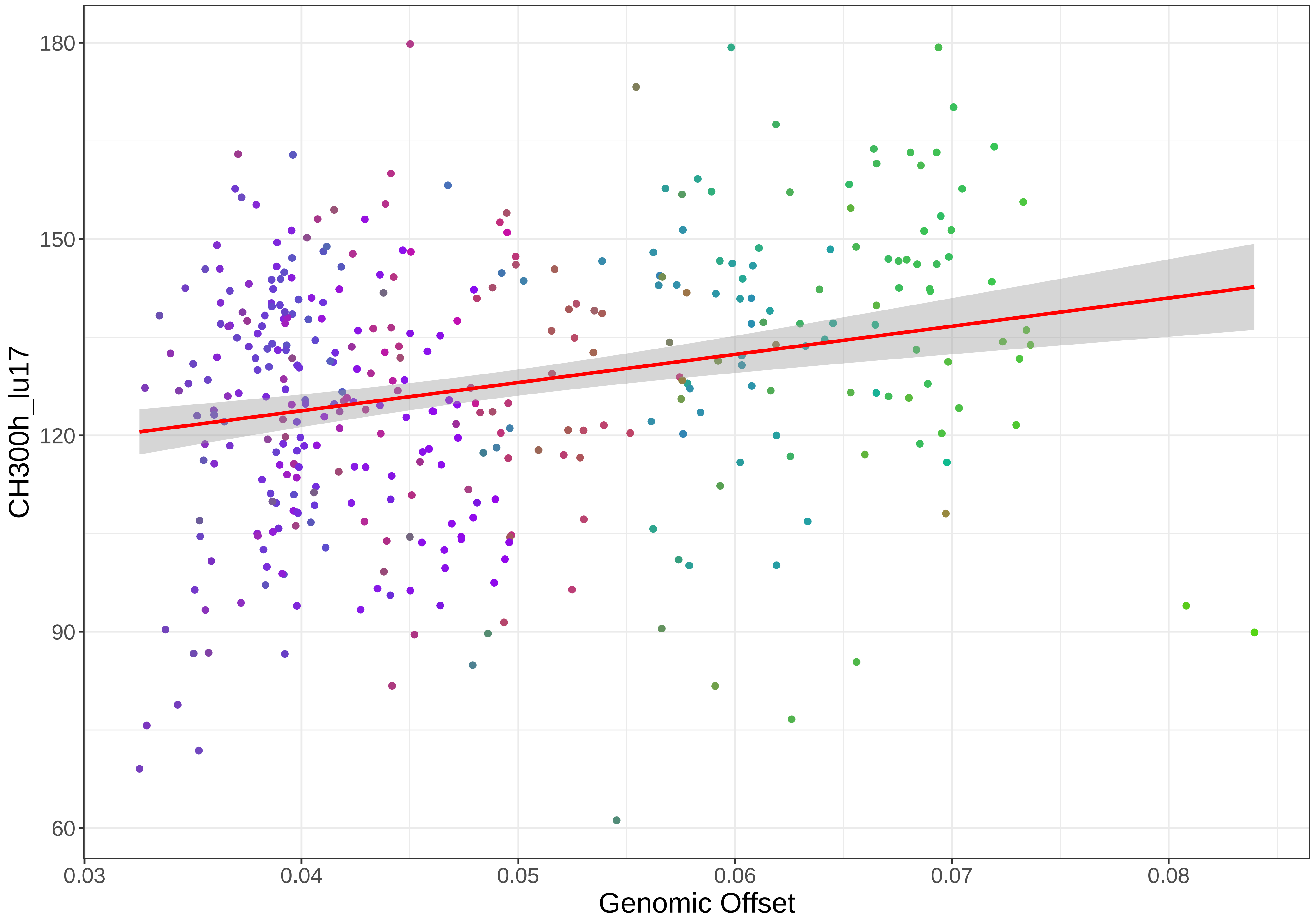

CH300h_lu17
180
150
120
90
60
0.03
0.04
0.05
0.06
0.07
0.08
Genomic Offset

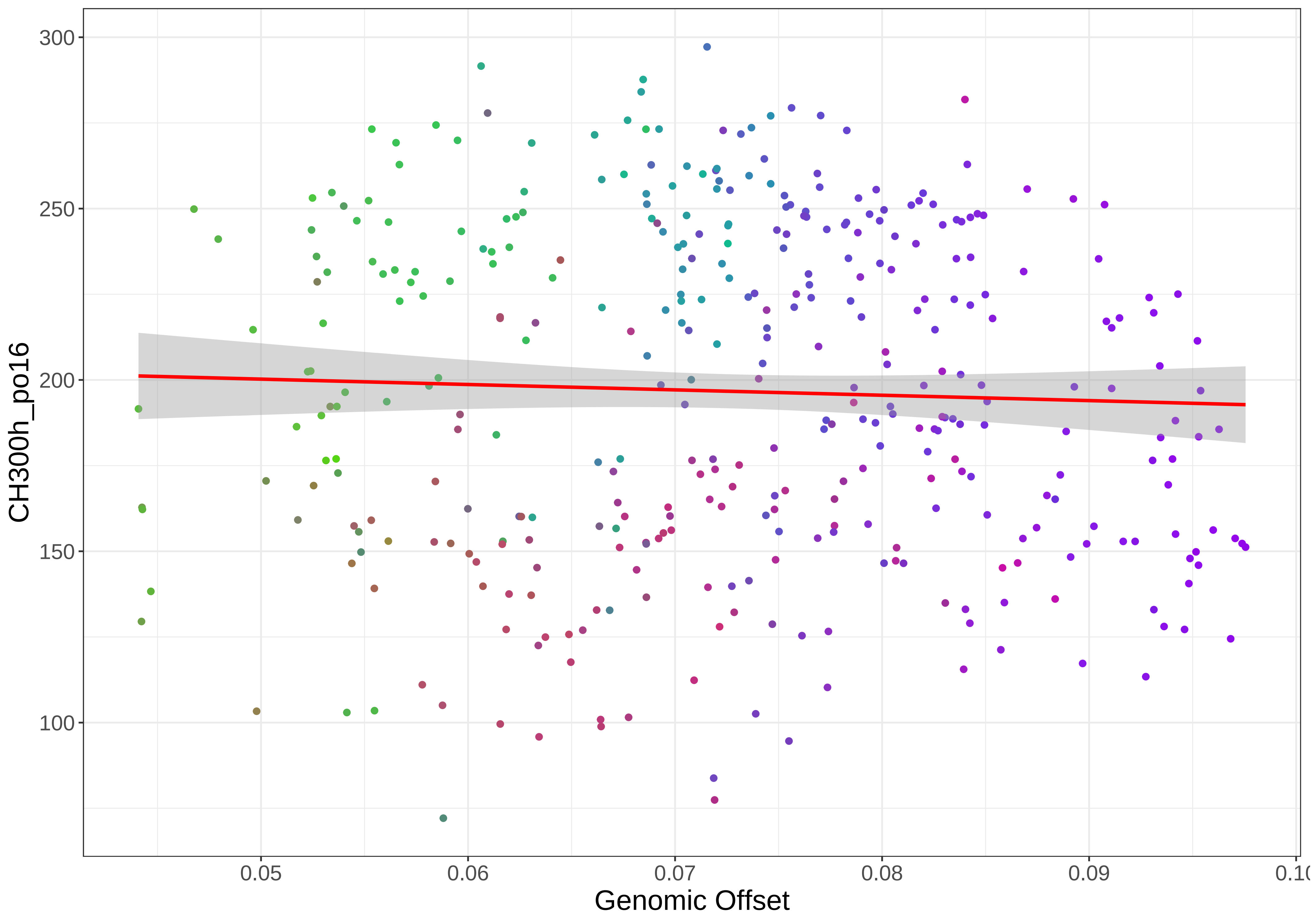

CH300h_po16
300
250
200
150
100
0.05
0.06
0.07
0.08
0.09
Genomic Offset

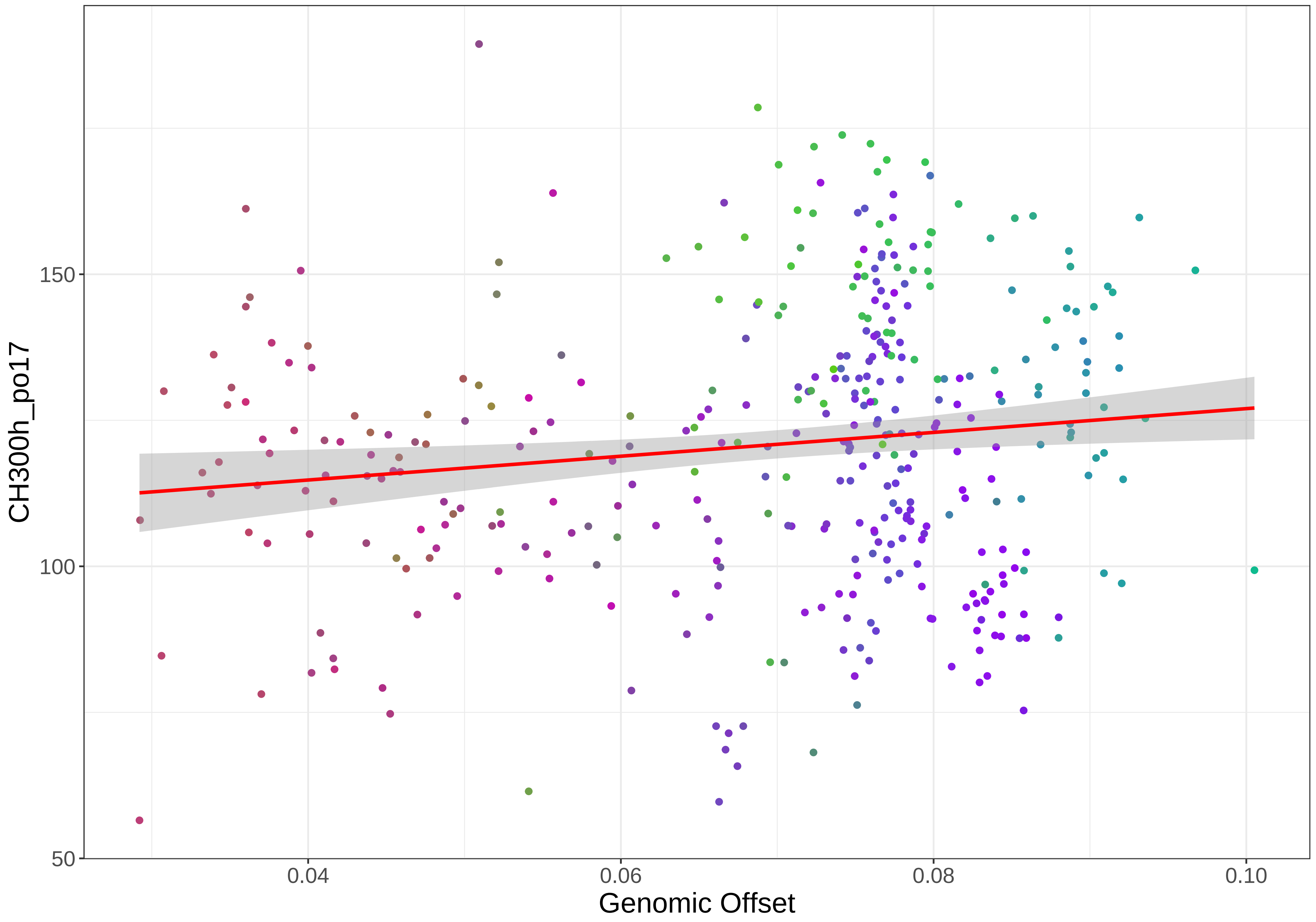

CH300h_po17
150
100
50
0.04
0.06
0.08
0.10
Genomic Offset

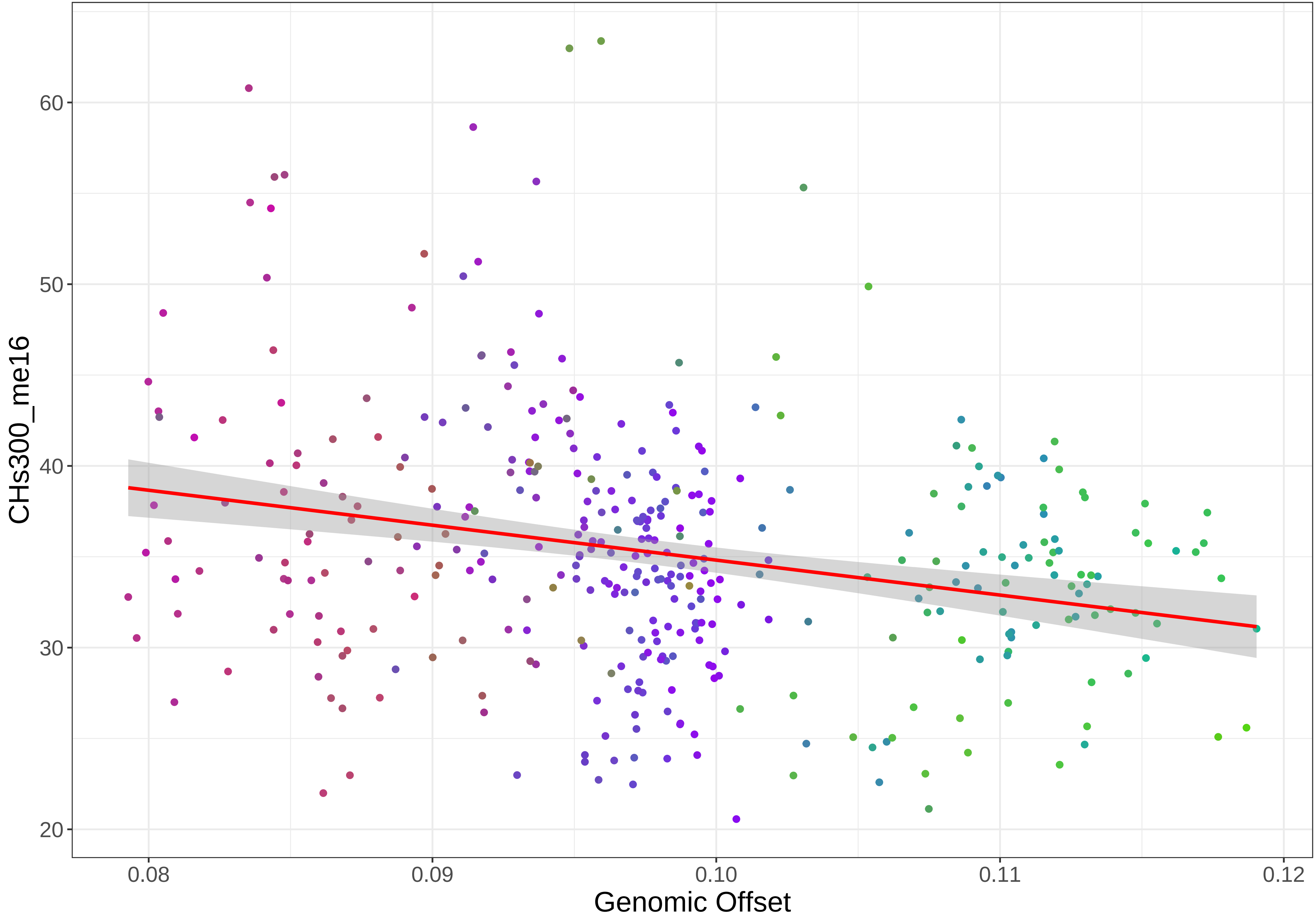

CHs300_me16
60
50
40
30
20
0.08
0.09
0.10
0.11
0.12
Genomic Offset

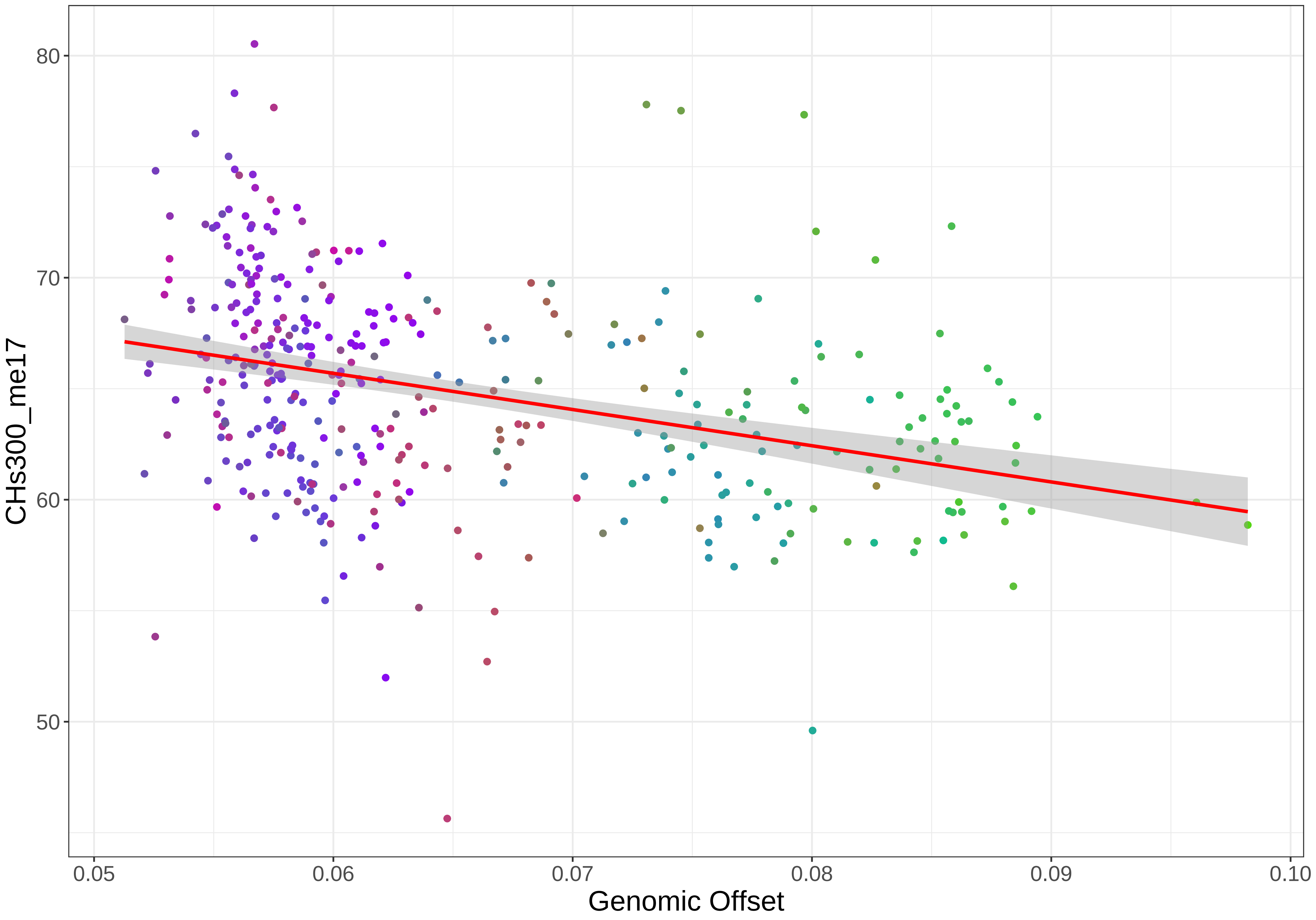

80
70
60
50
CHs300_me17
0.05
0.06
0.07
0.08
0.09
0.10
Genomic Offset

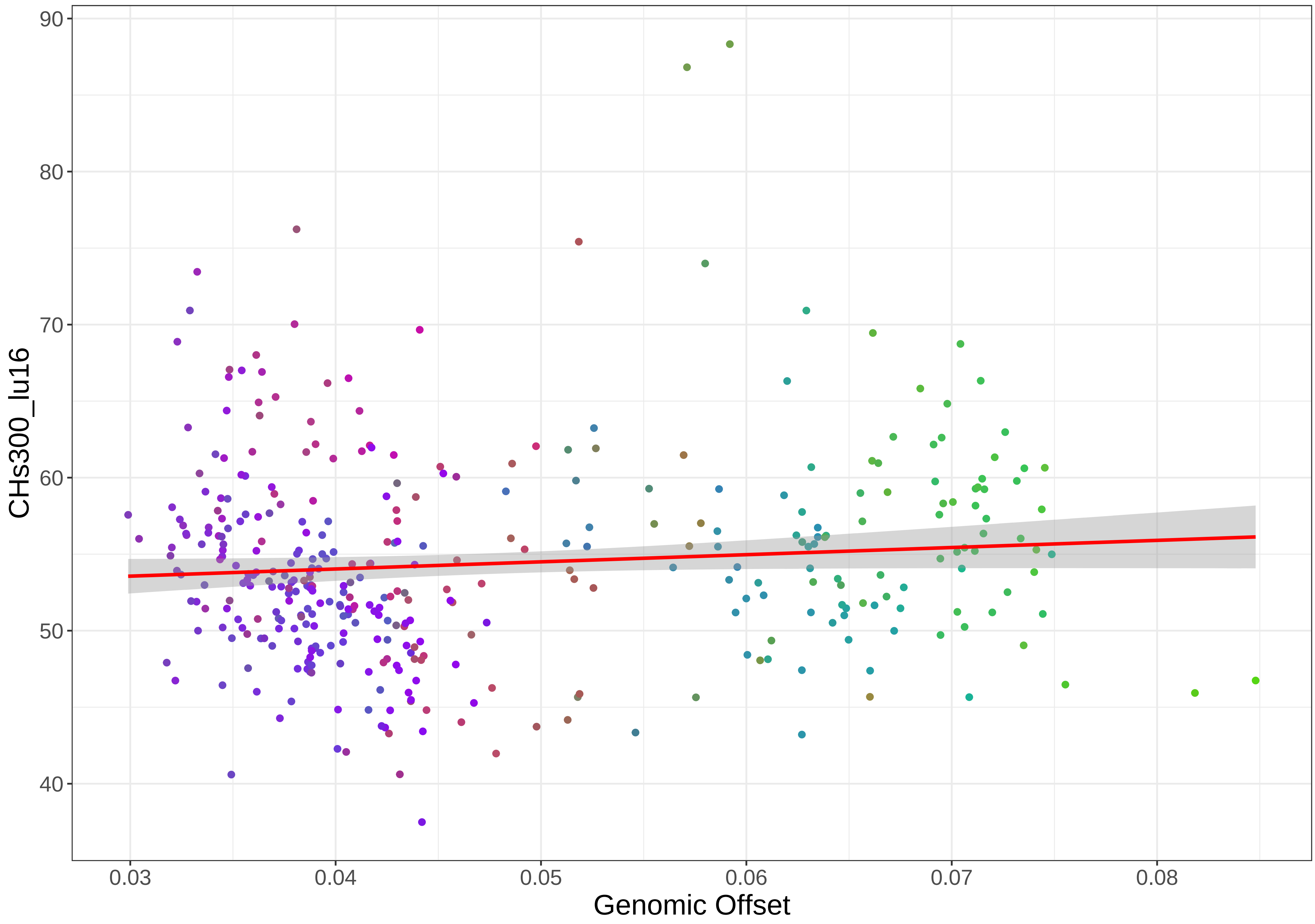

90
80
70
60
50
40
CHs300_lu16
0.03
0.04
0.05
0.06
0.07
0.08
Genomic Offset

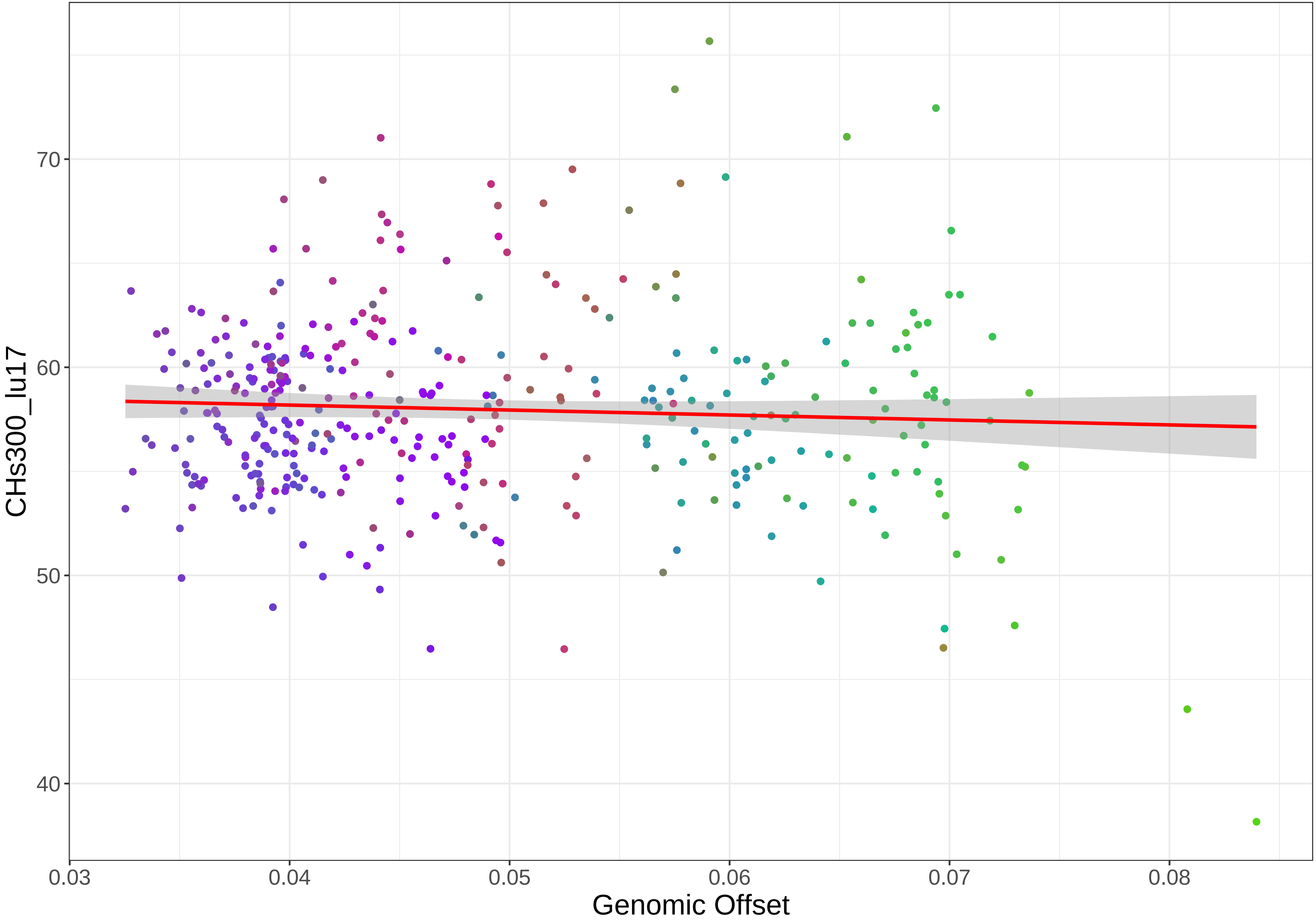
CHs300_lu17
70
60
50
40
0.03
0.04
0.05
0.06
0.07
0.08
Genomic Offset

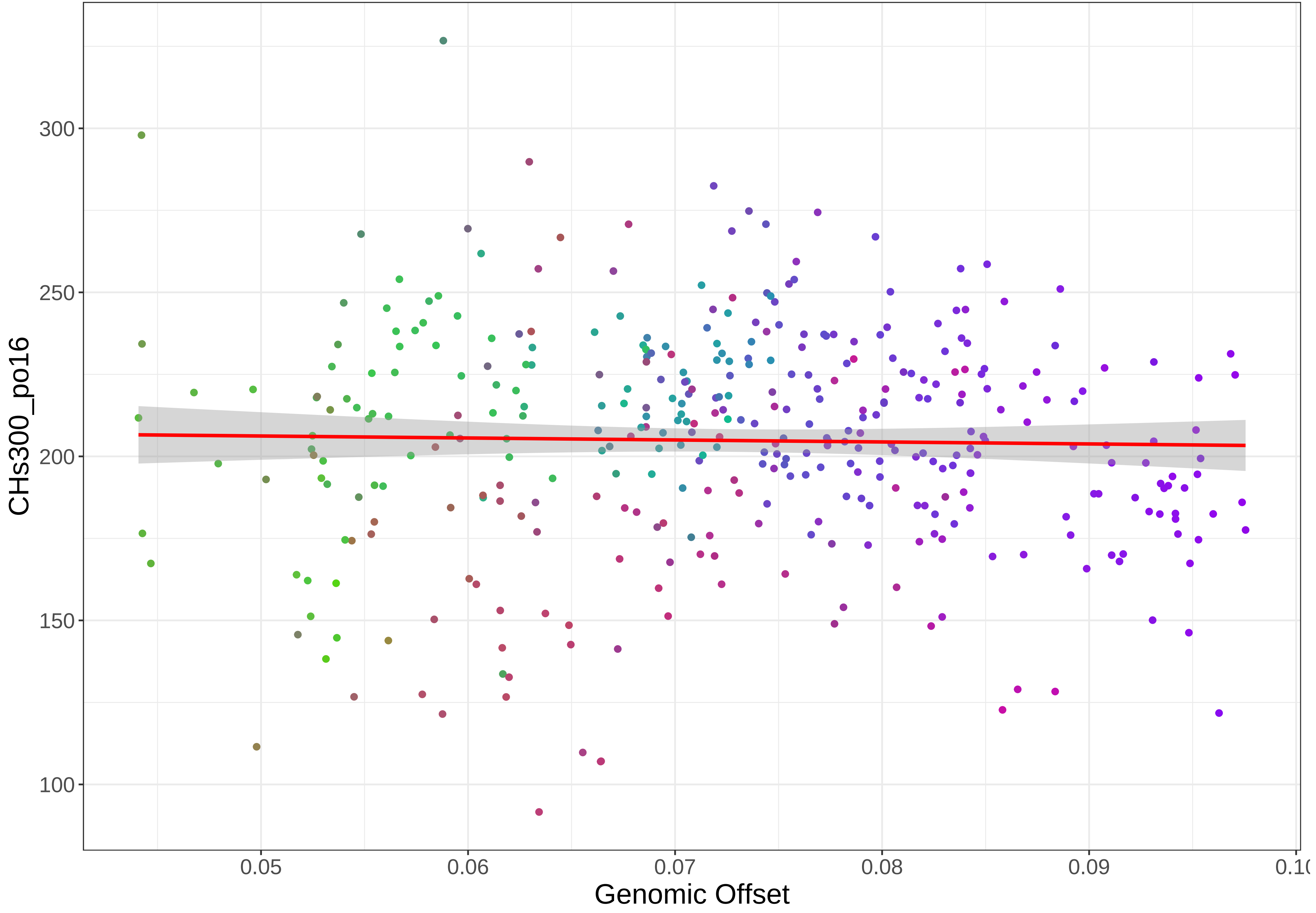
CHs300_po16
300
250
200
150
100
0.05
0.06
0.07
0.08
0.09
0.10
Genomic Offset

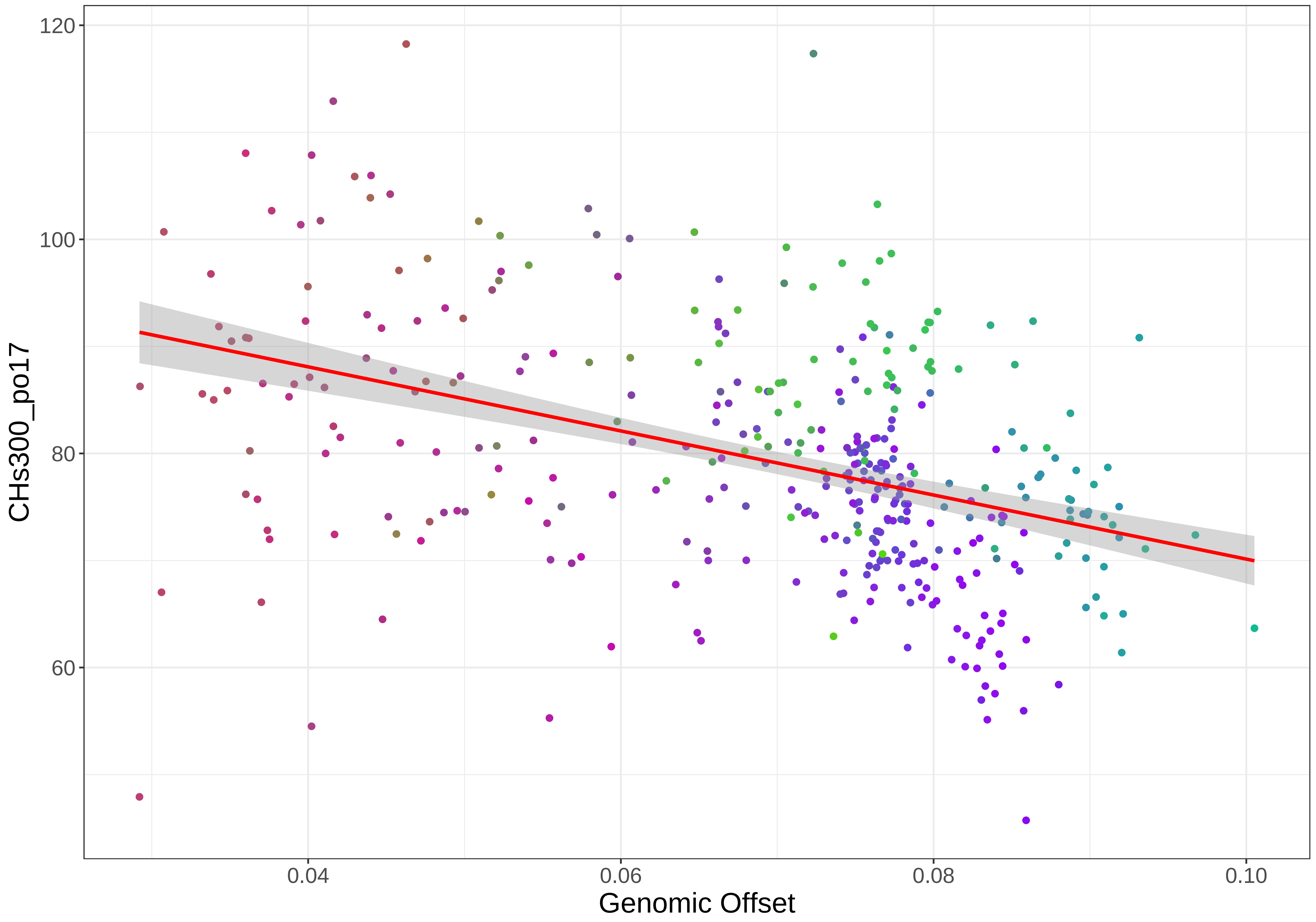

120
100
80
60
CHs300_po17
0.04
0.06
0.08
0.10
Genomic Offset

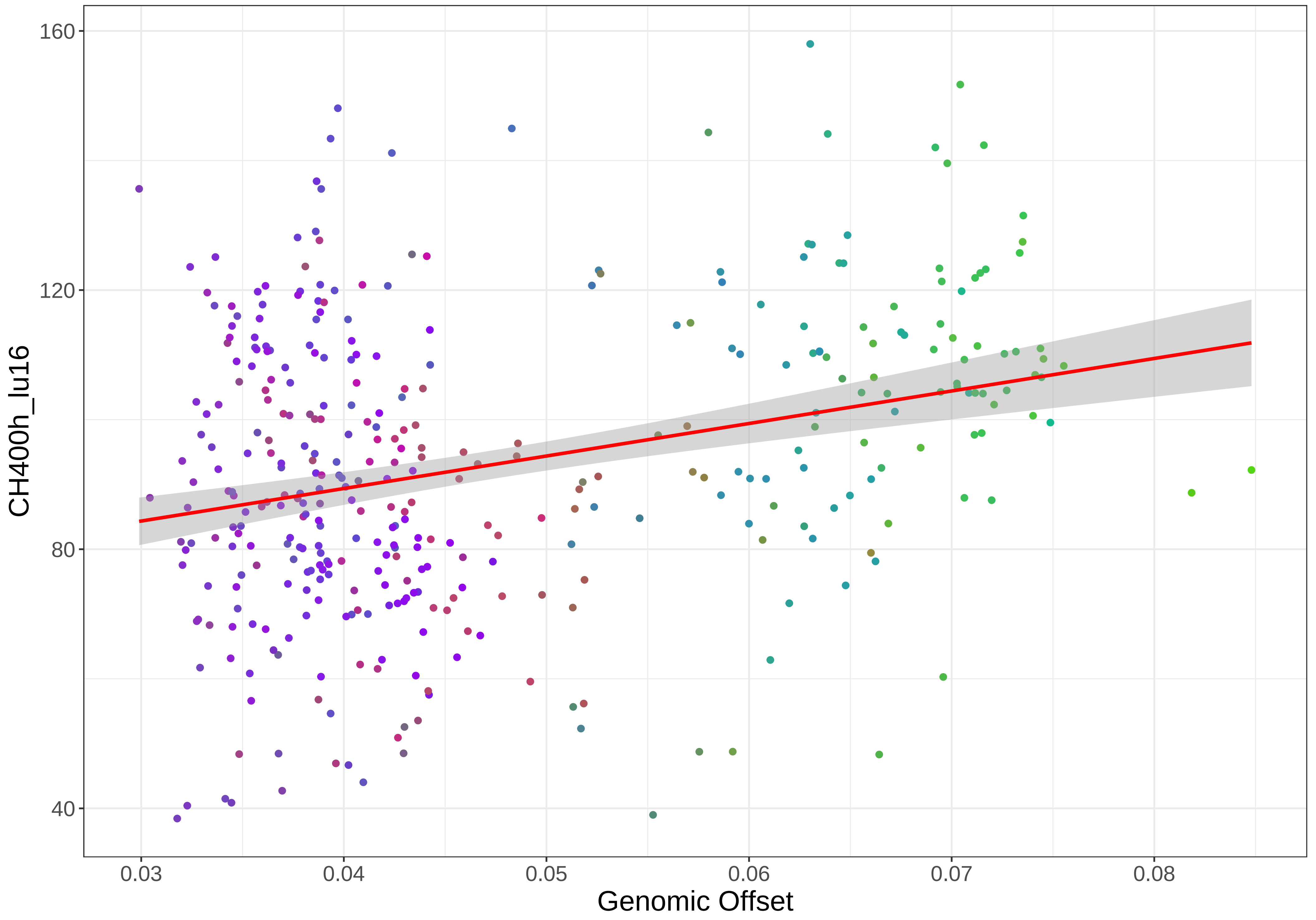

160
120
80
40
CH400h_lu16
0.03
0.04
0.05
0.06
0.07
0.08
Genomic Offset

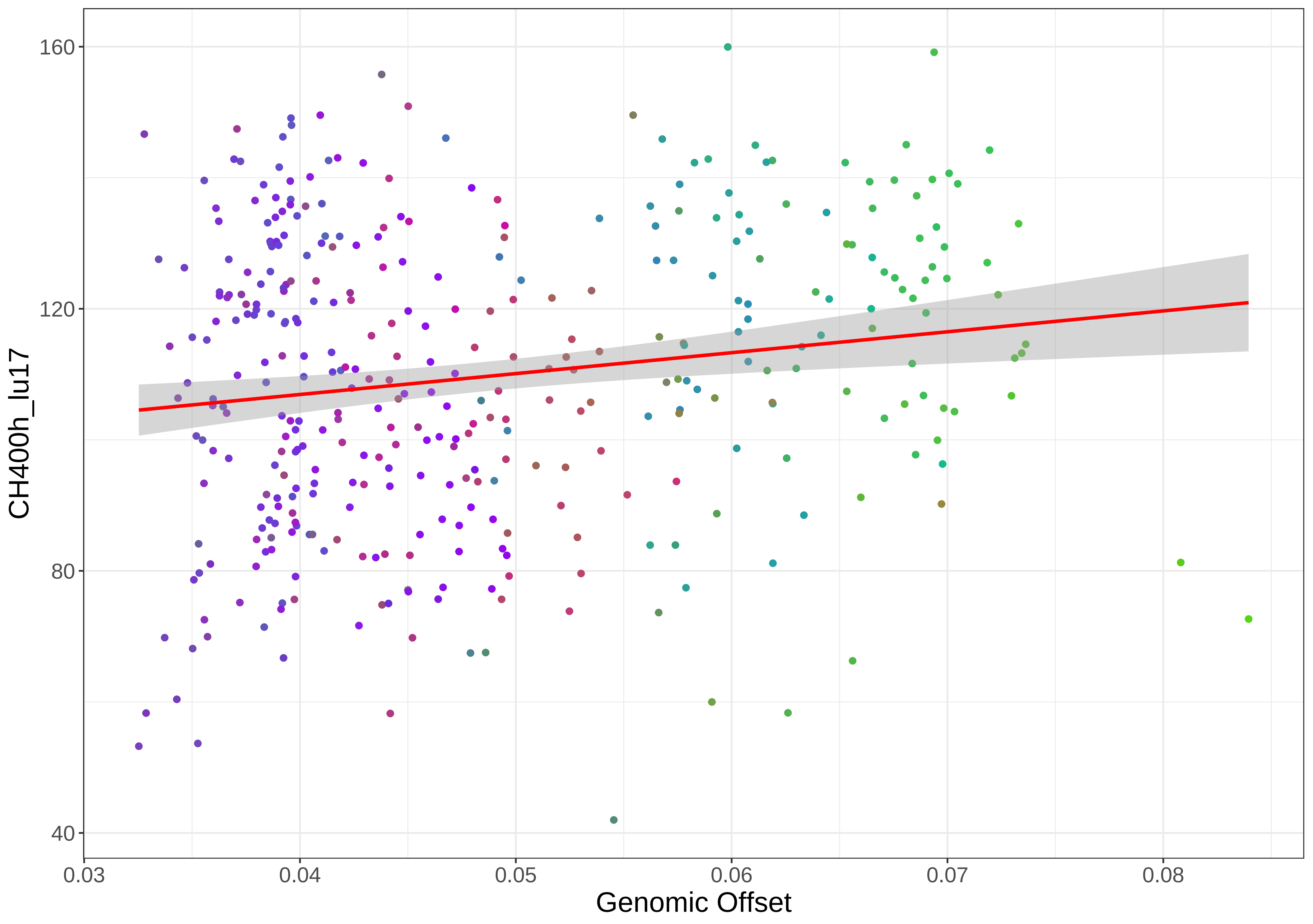

CH400h_lu17
160
120
80
40
0.03
0.04
0.05
0.06
0.07
0.08
Genomic Offset

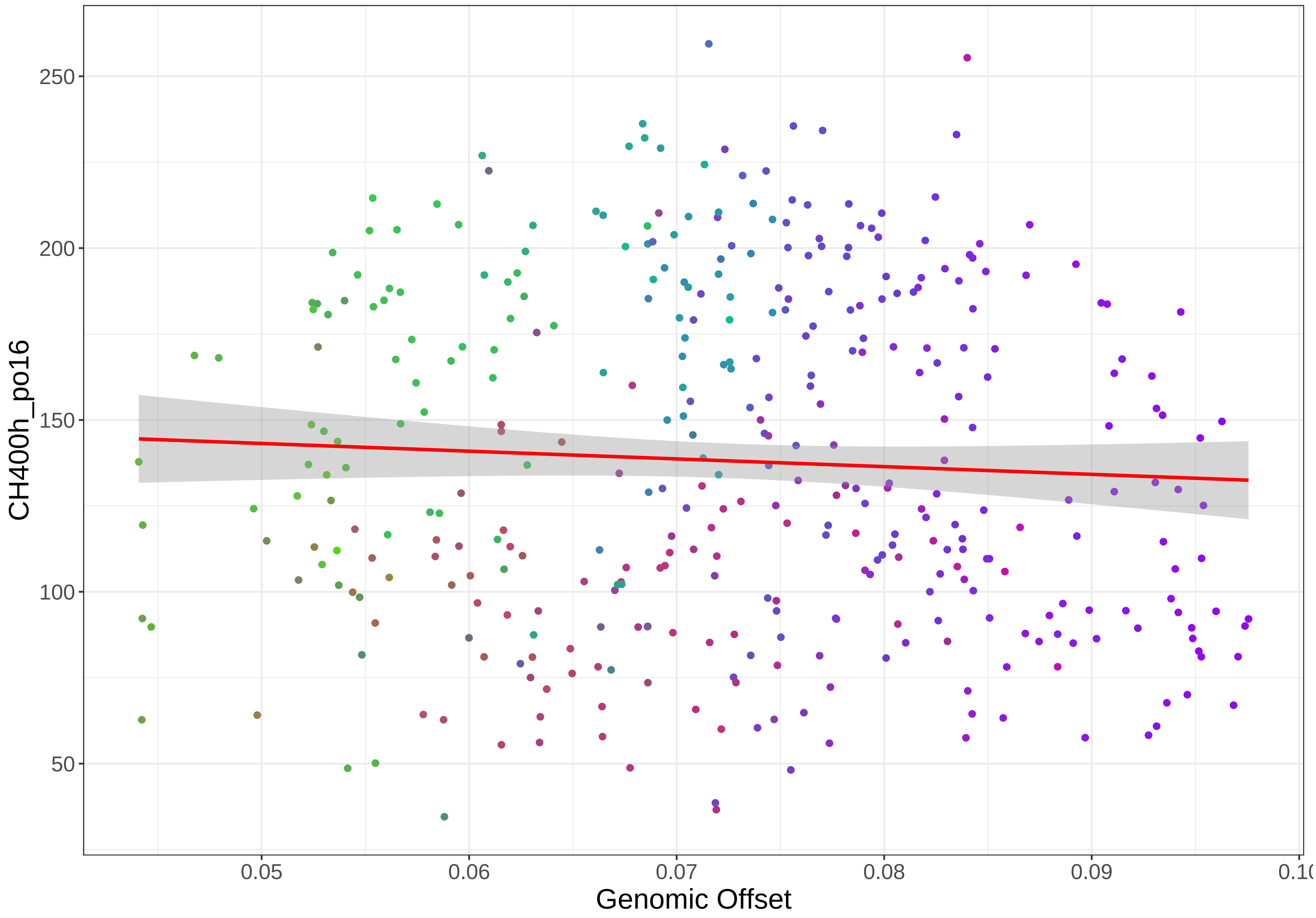

250
200
150
100
50
CH400h_po16
0.05
0.06
0.07
0.08
0.09
Genomic Offset

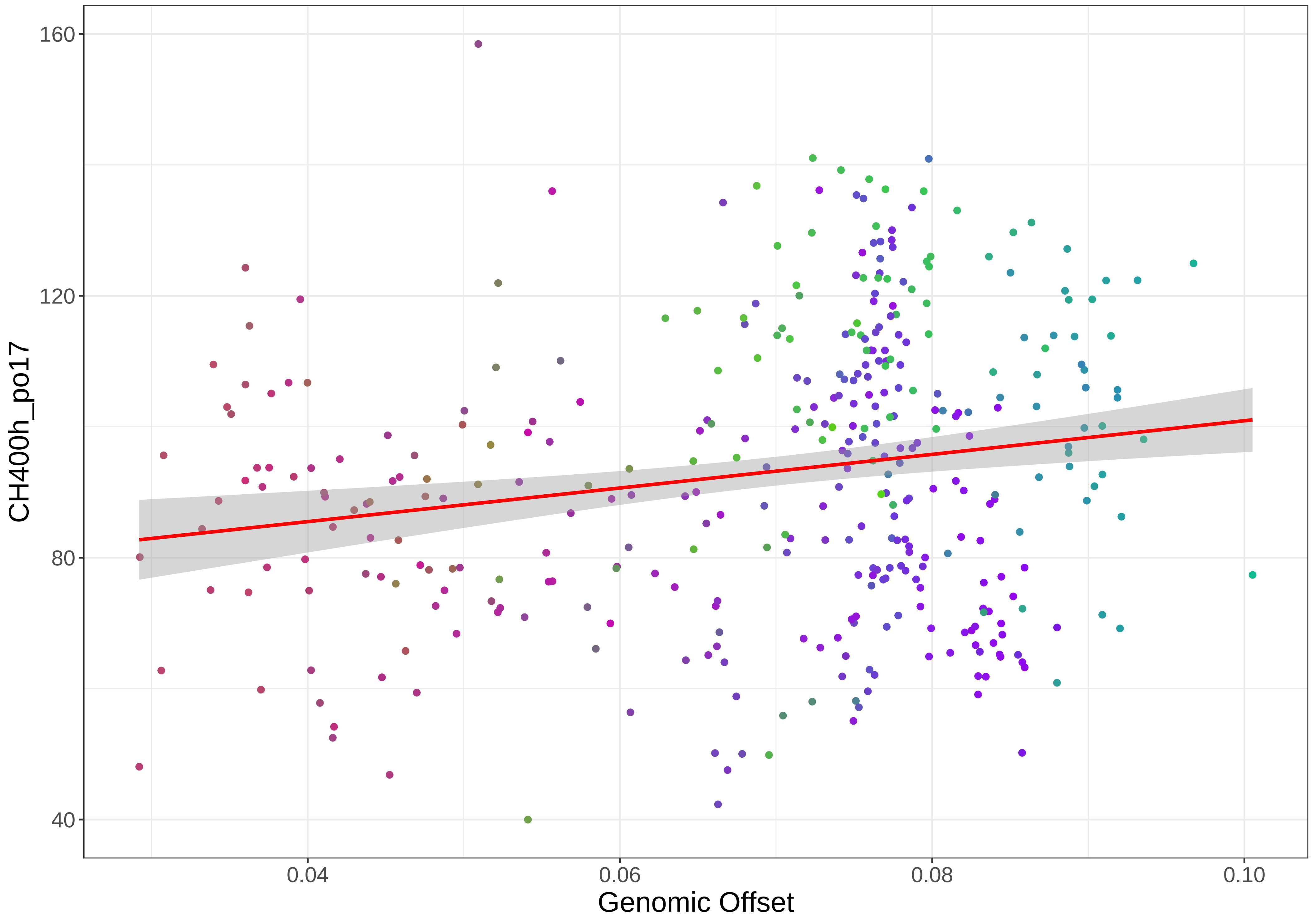

160
120
80
40
CH400h_po17
0.04
0.06
0.08
0.10
Genomic Offset

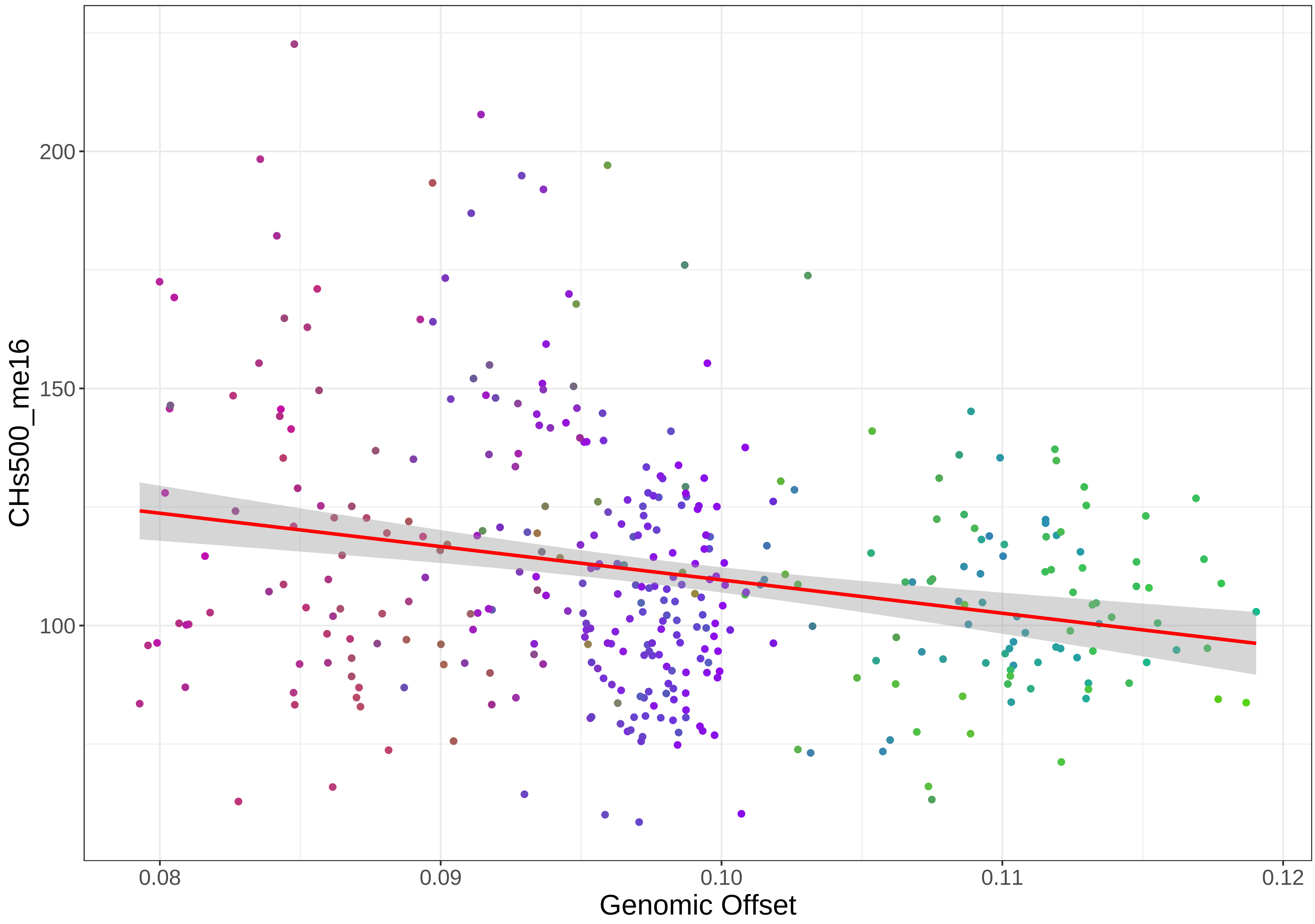

CHs500_me16
200
150
100
0.08
0.09
0.10
0.11
0.12
Genomic Offset

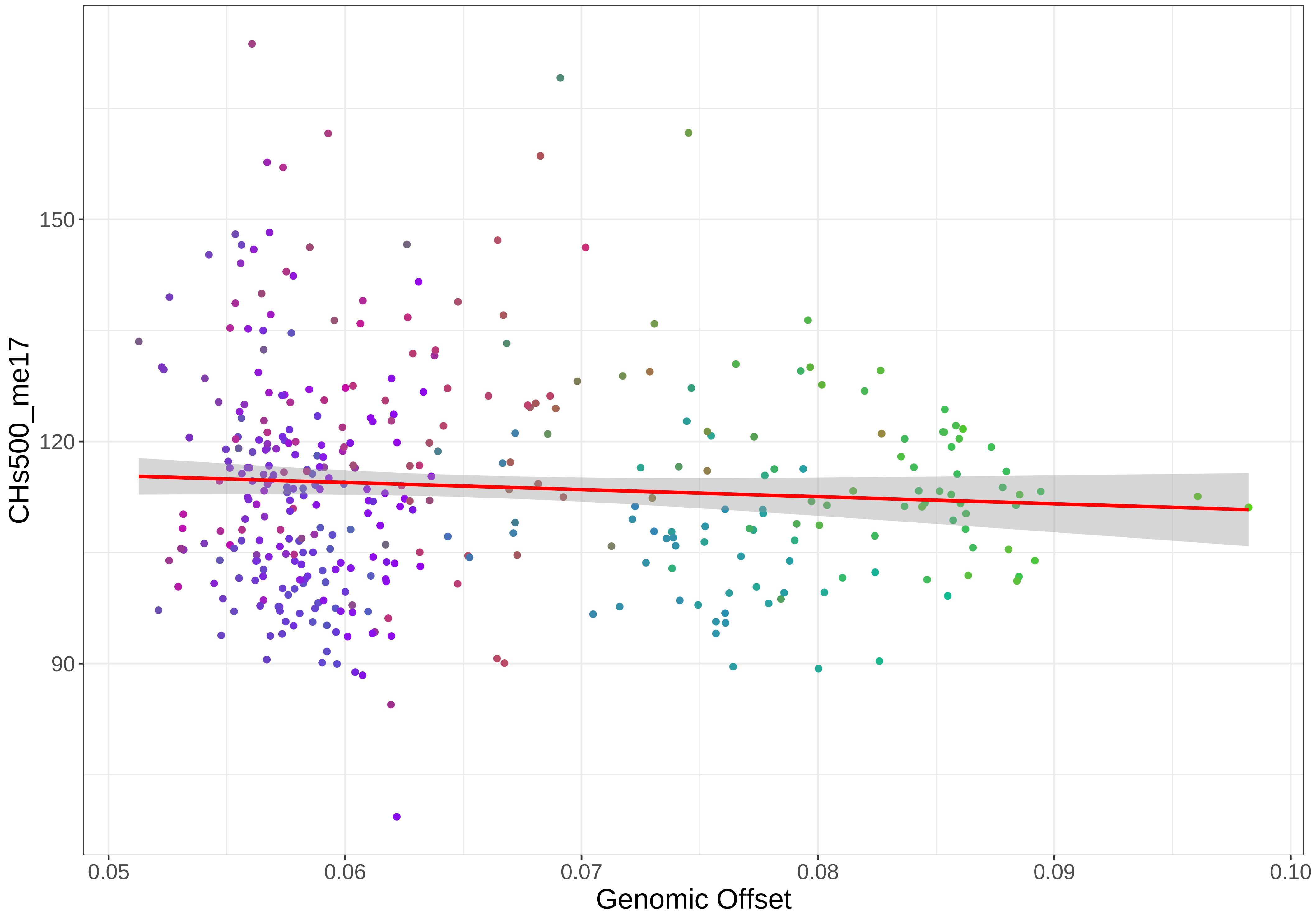

CHs500_me17
150
120
90
0.05
0.06
0.07
0.08
0.09
0.10
Genomic Offset

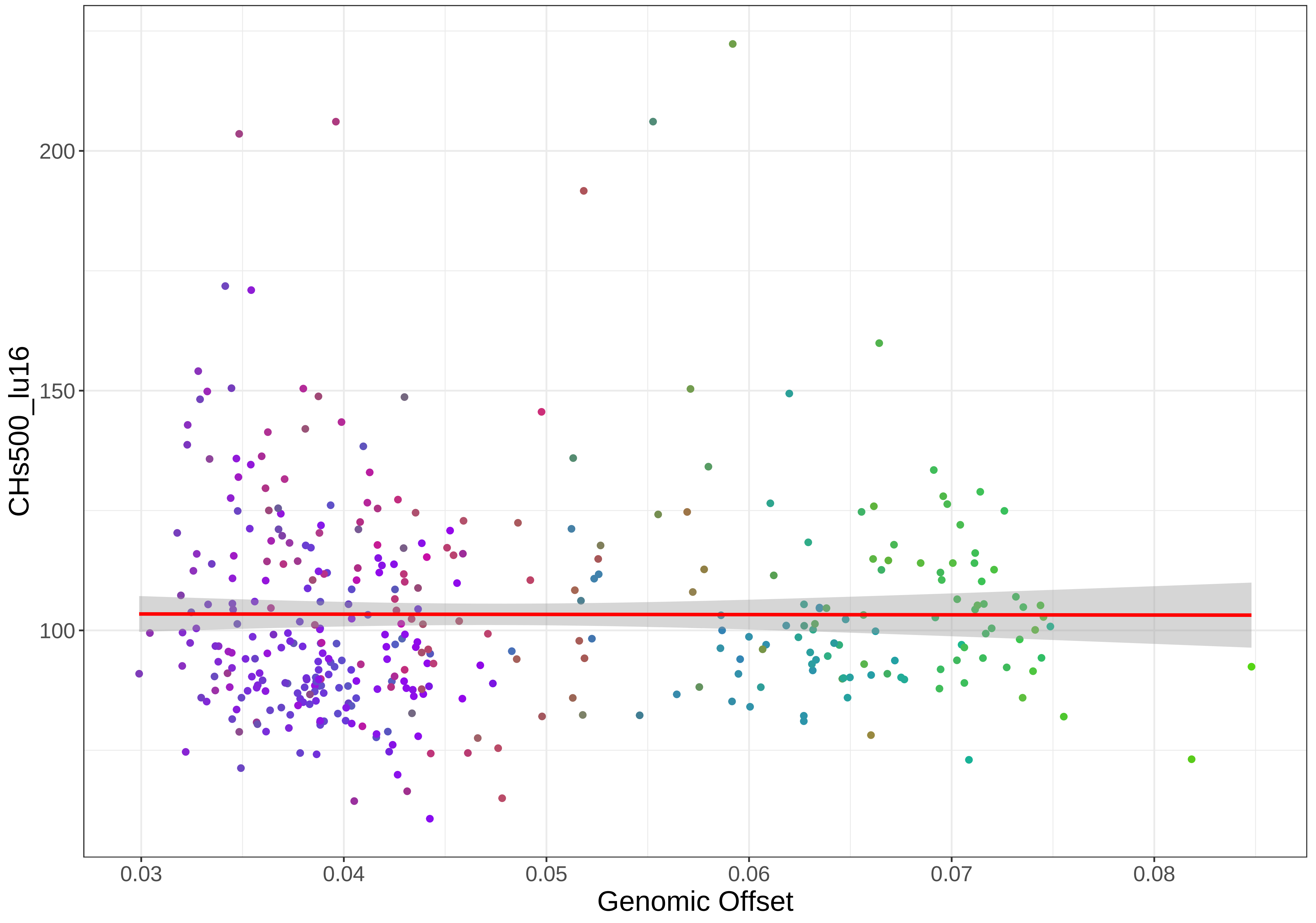

CHs500_lu16
200
150
100
0.03
0.04
0.05
0.06
0.07
0.08
Genomic Offset

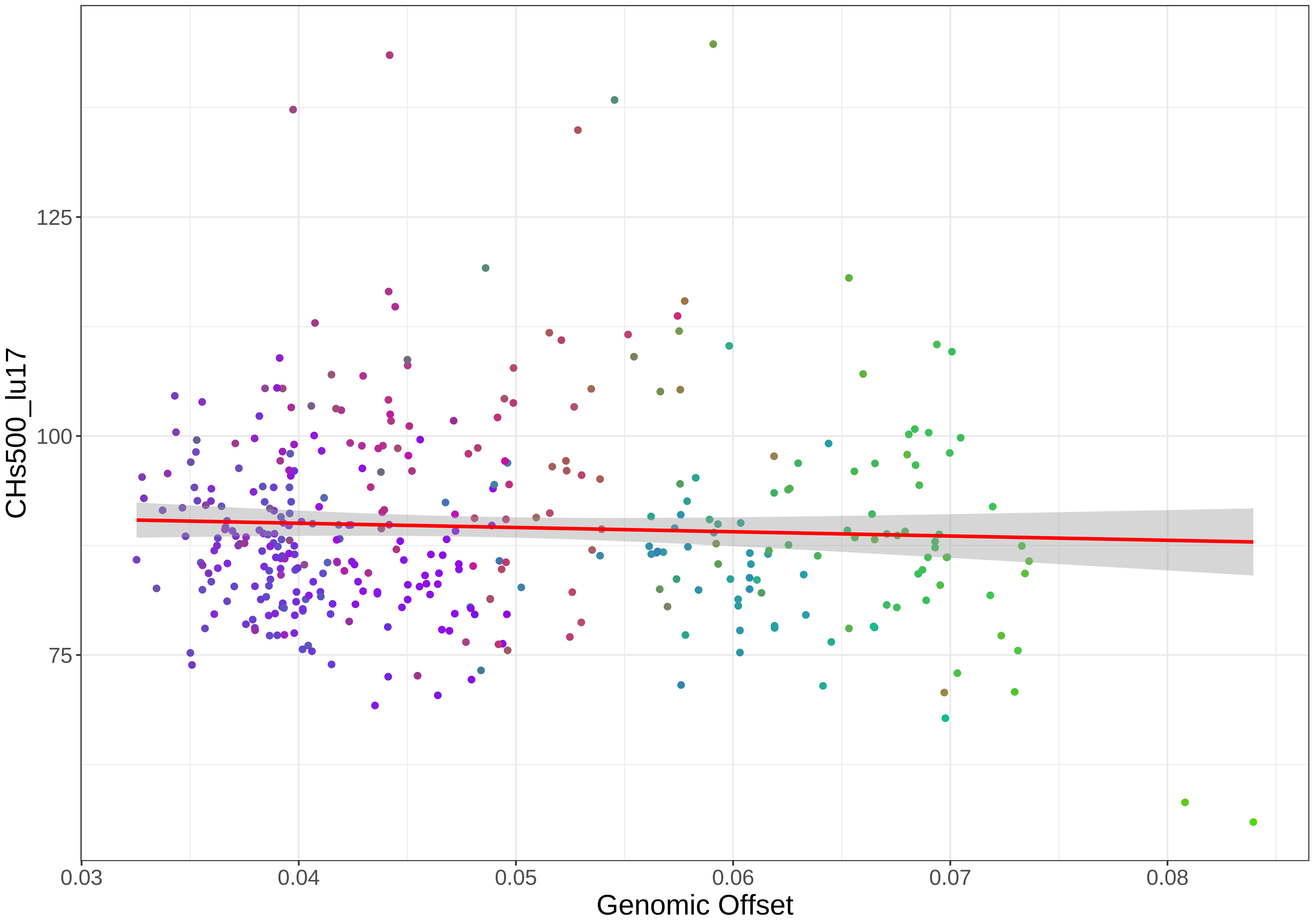

CHs500_lu17
125
100
75
0.03
0.04
0.05
0.06
0.07
0.08
Genomic Offset

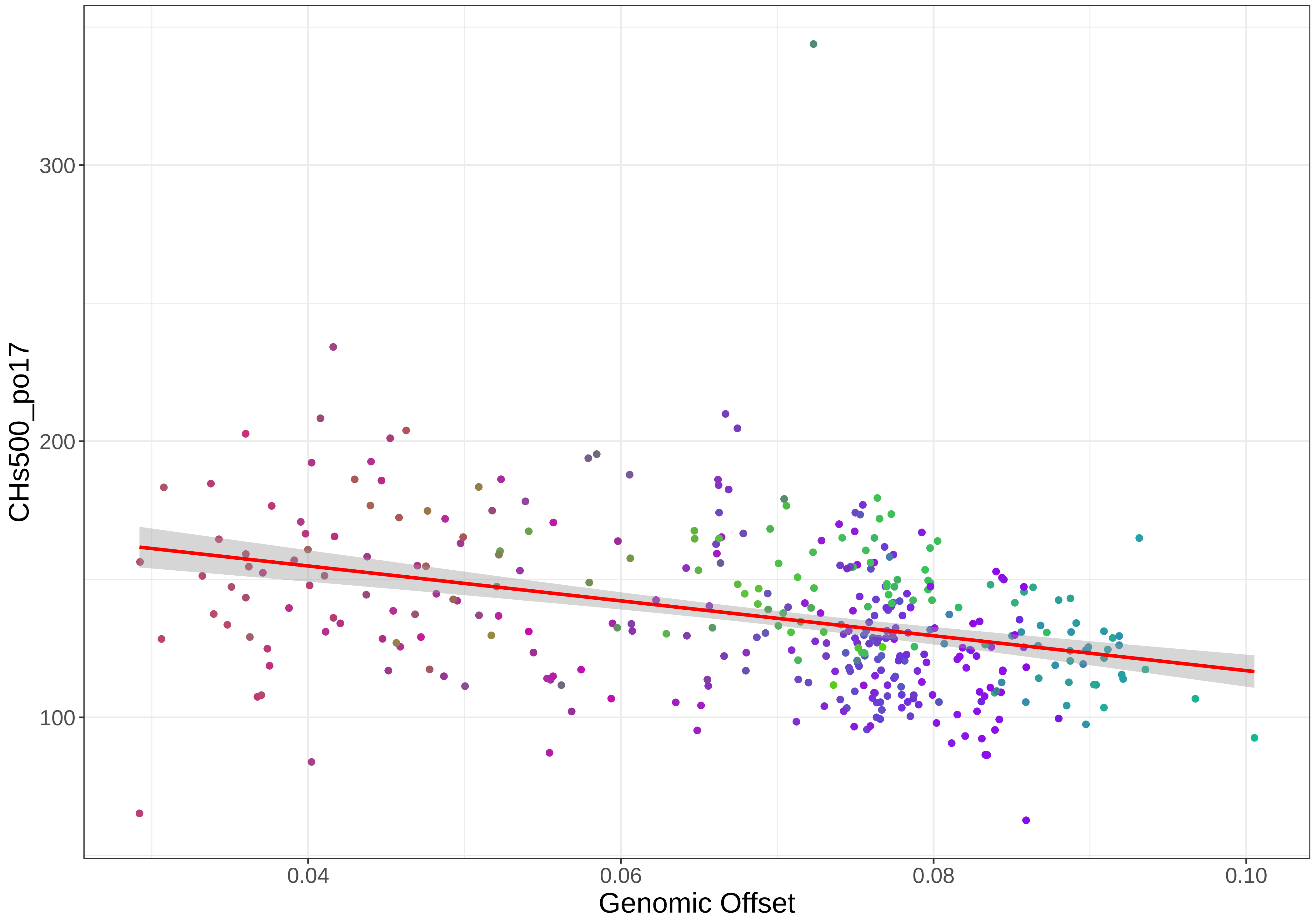

CHs500_po17
300
200
100
0.04
0.06
0.08
0.10
Genomic Offset

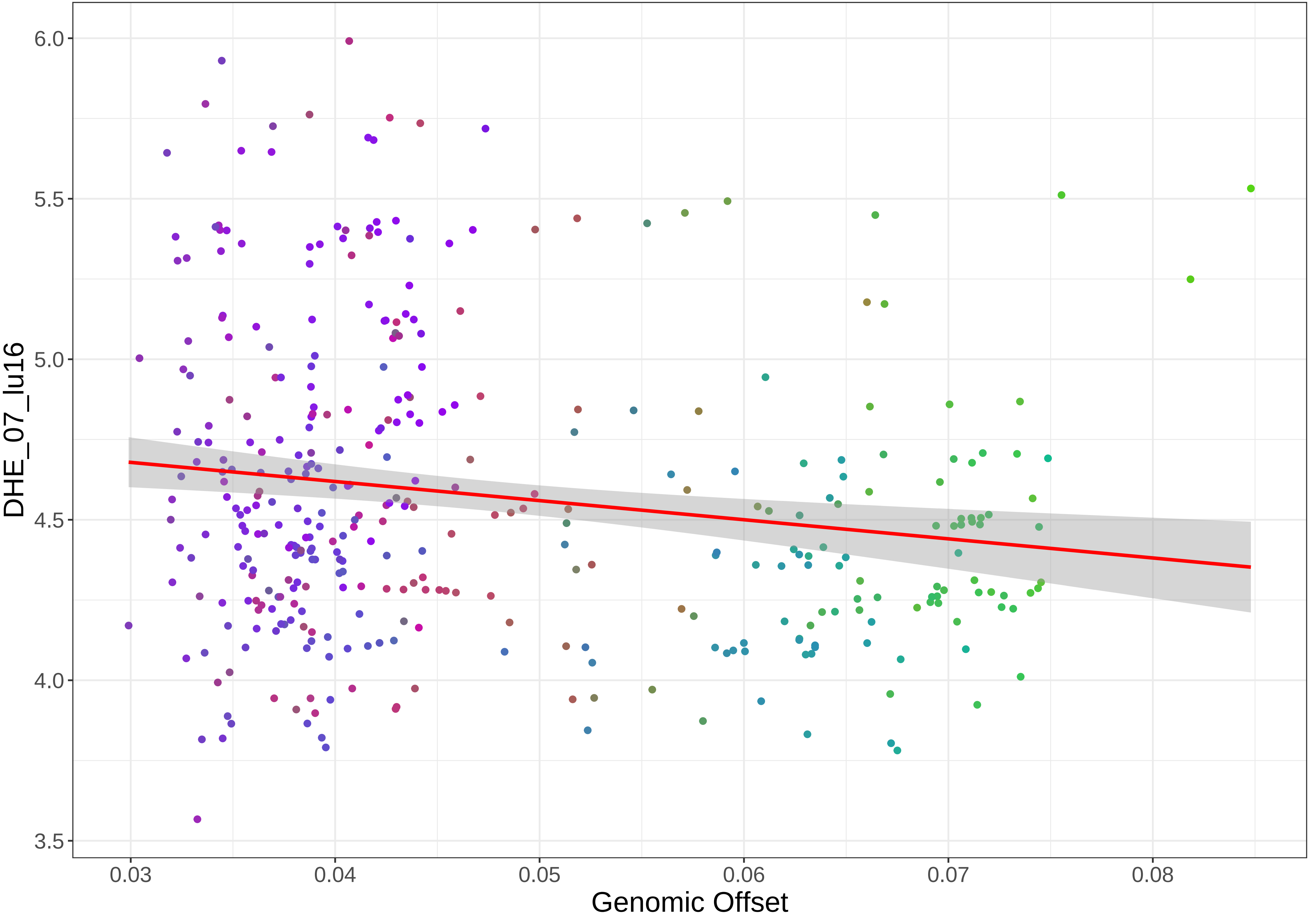

DHE_07_lu16
6.0
5.5
5.0
4.5
4.0
3.5
0.03
0.04
0.05
0.06
0.07
0.08
Genomic Offset

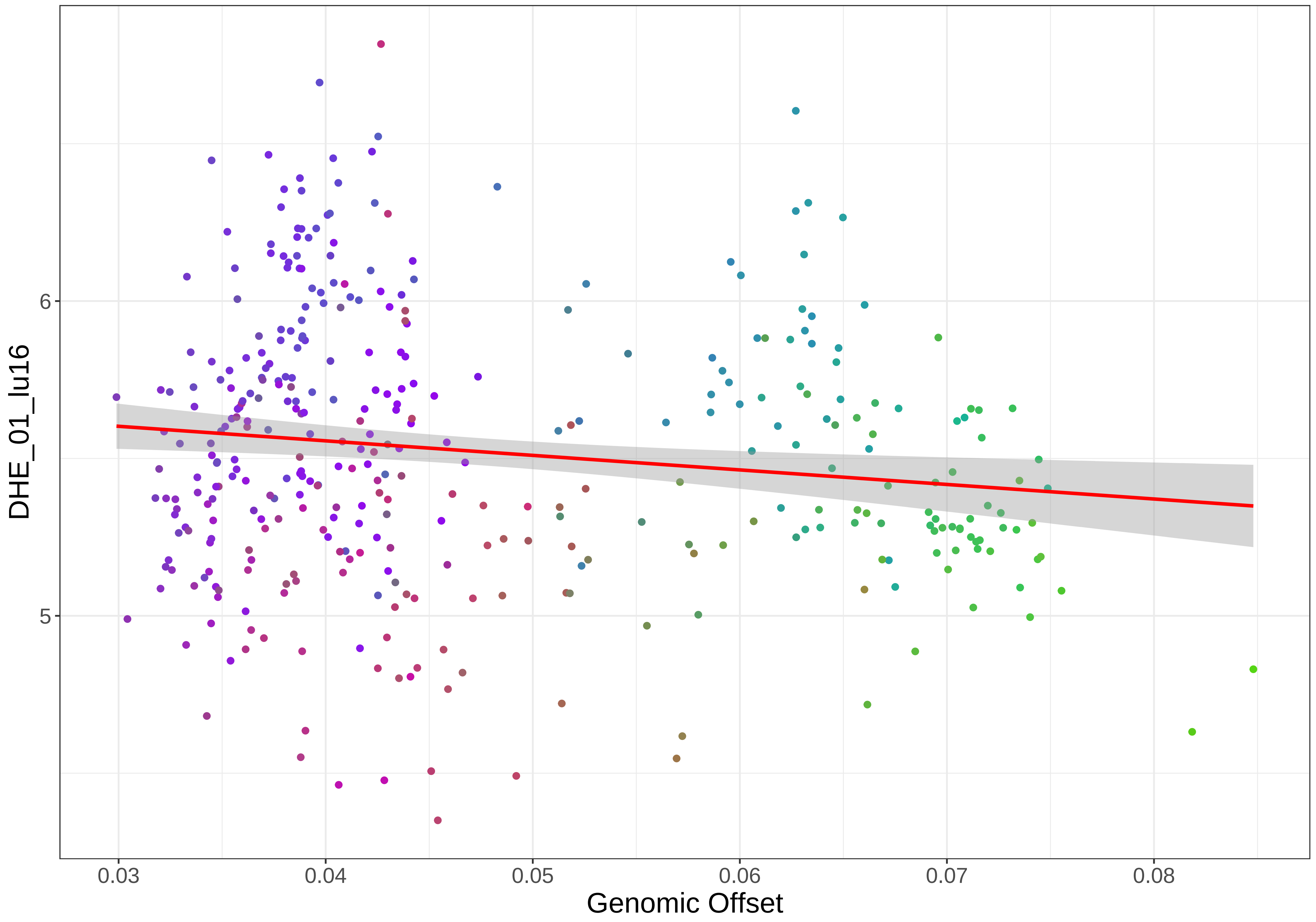

DHE_01_lu16
6
5
0.03
0.04
0.05
0.06
0.07
0.08
Genomic Offset

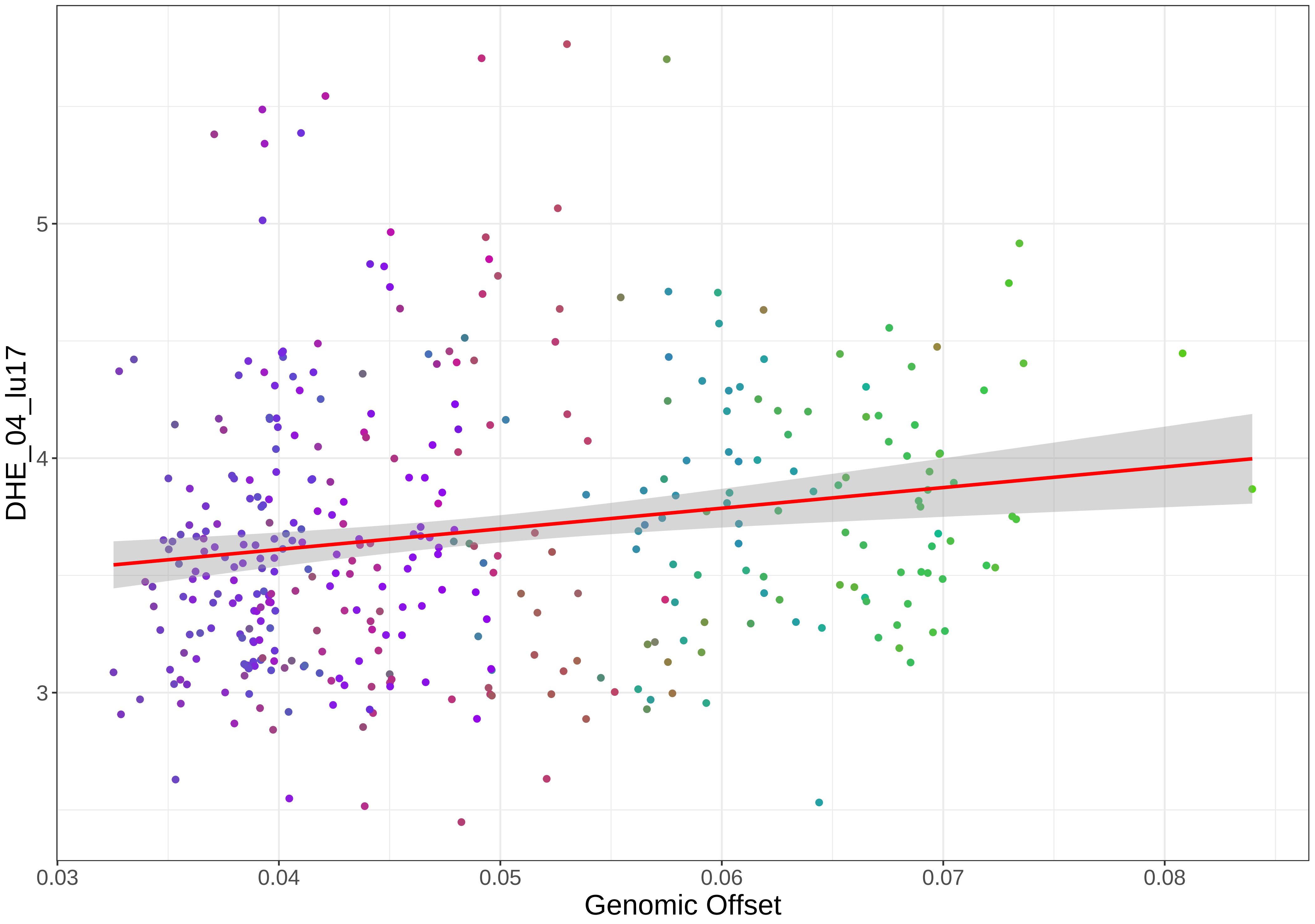
DHE_04_lu17
5
4
3
0.03
0.04
0.05
0.06
0.07
0.08
Genomic Offset

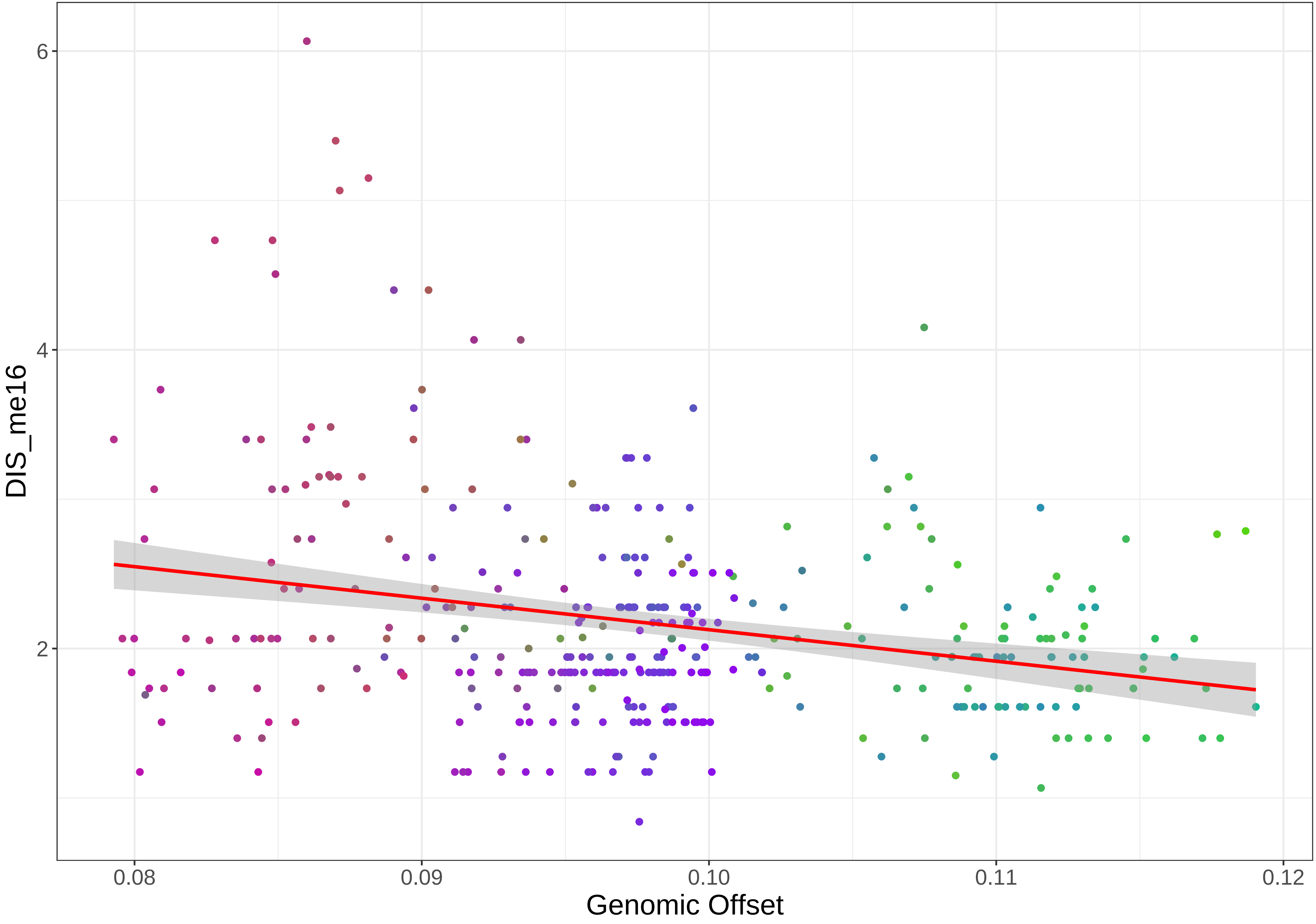

DIS_me16
6
4
2
0.08
0.09
0.10
0.11
0.12
Genomic Offset

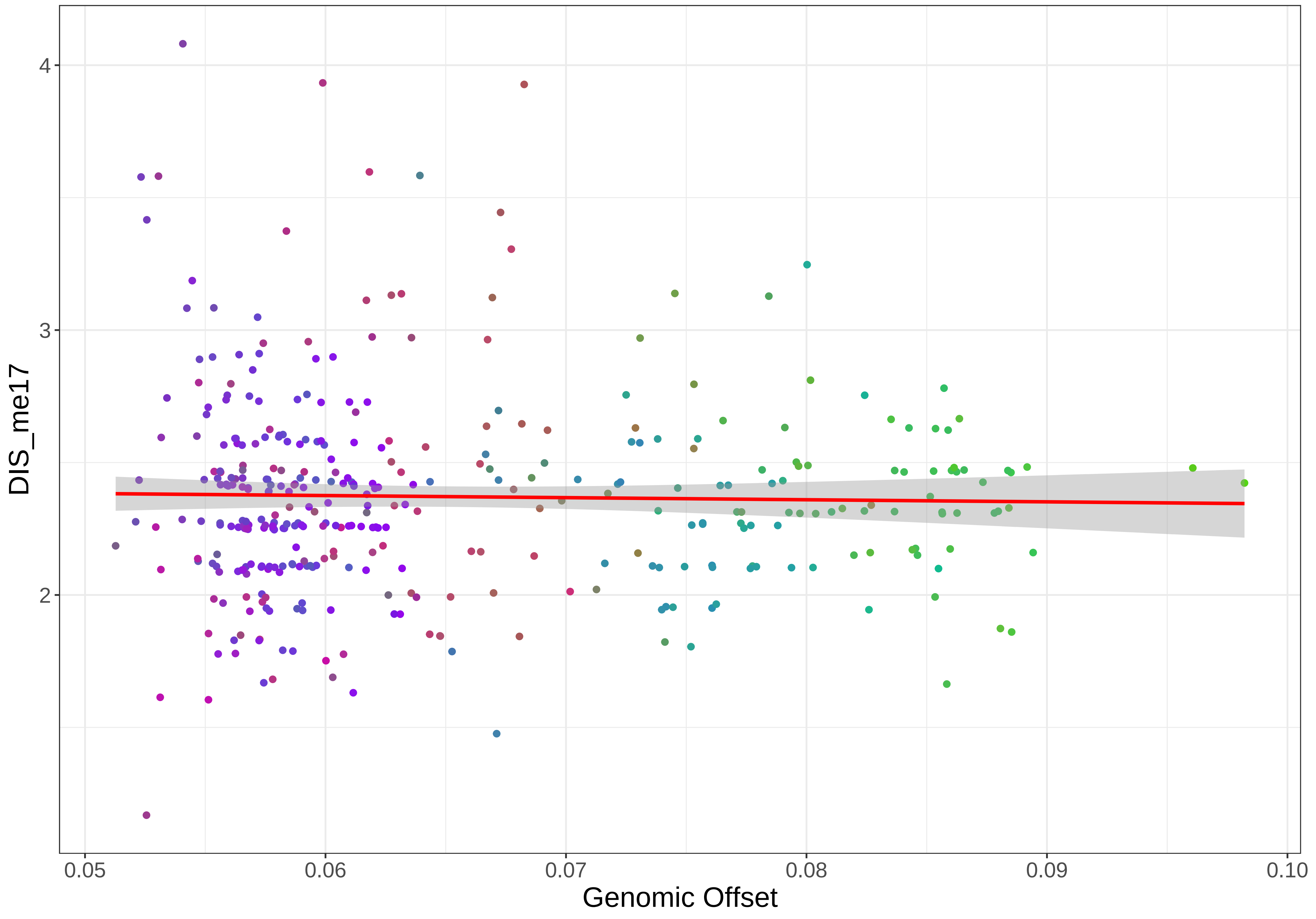
DIS_me17
4
3
2
0.05
0.06
0.07
0.08
0.09
0.10
Genomic Offset

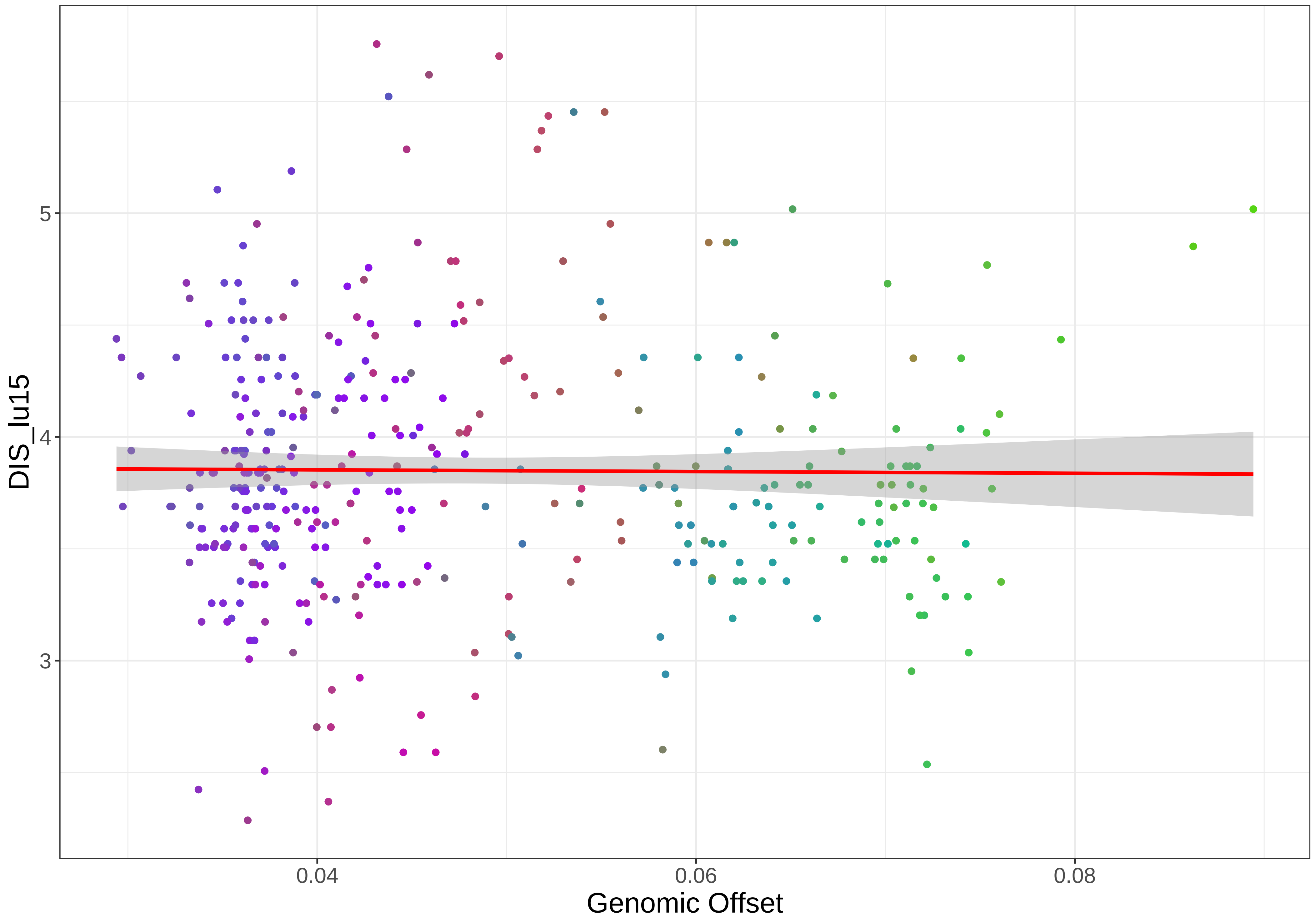

DIS_lu15
5
4
3
0.04
0.06
0.08
Genomic Offset

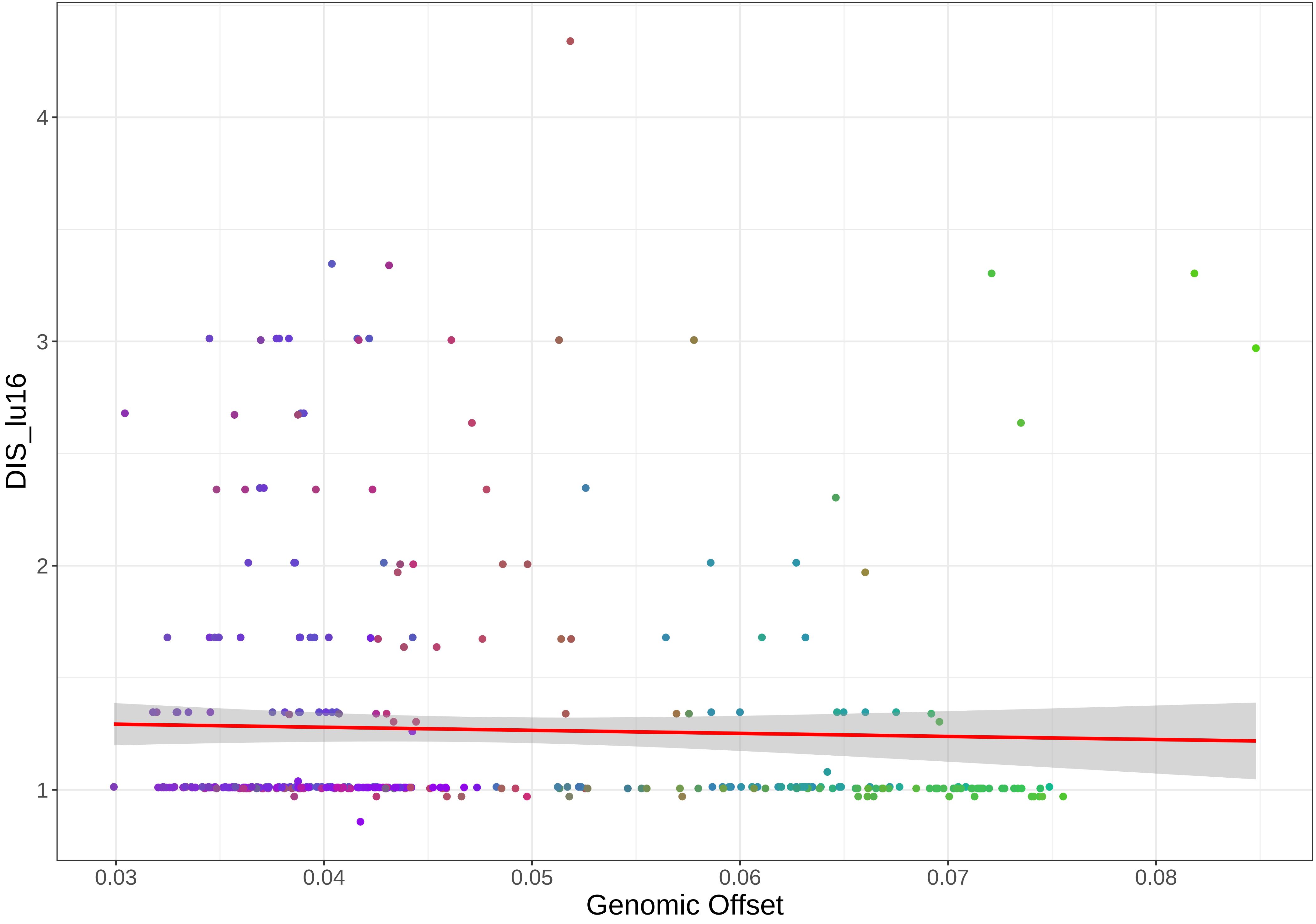
DIS_lu16
Genomic Offset
4
3
2
1
0.03
0.04
0.05
0.06
0.07
0.08

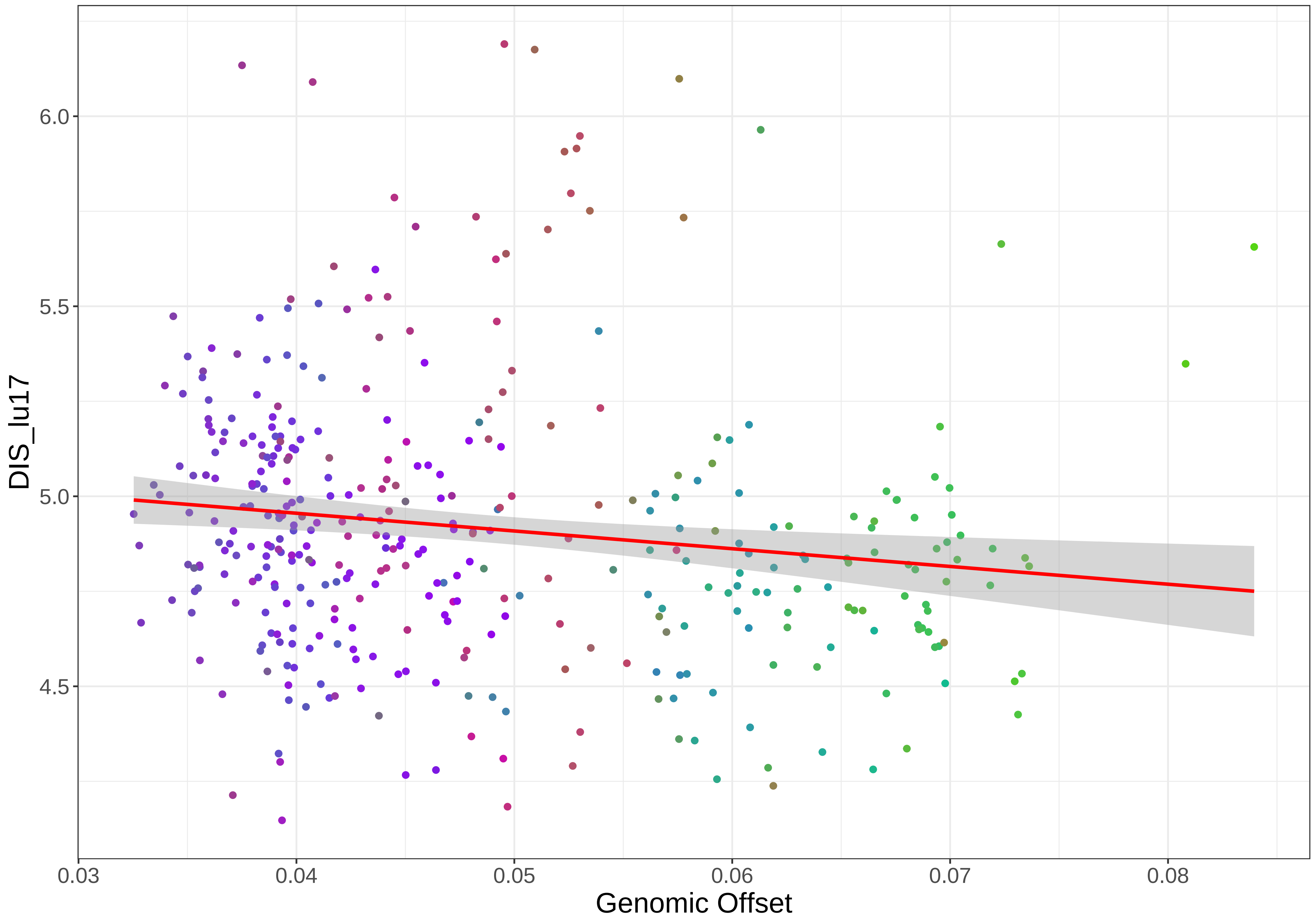

DIS_lu17
6.0
5.5
5.0
4.5
0.03
0.04
0.05
0.06
0.07
0.08
Genomic Offset

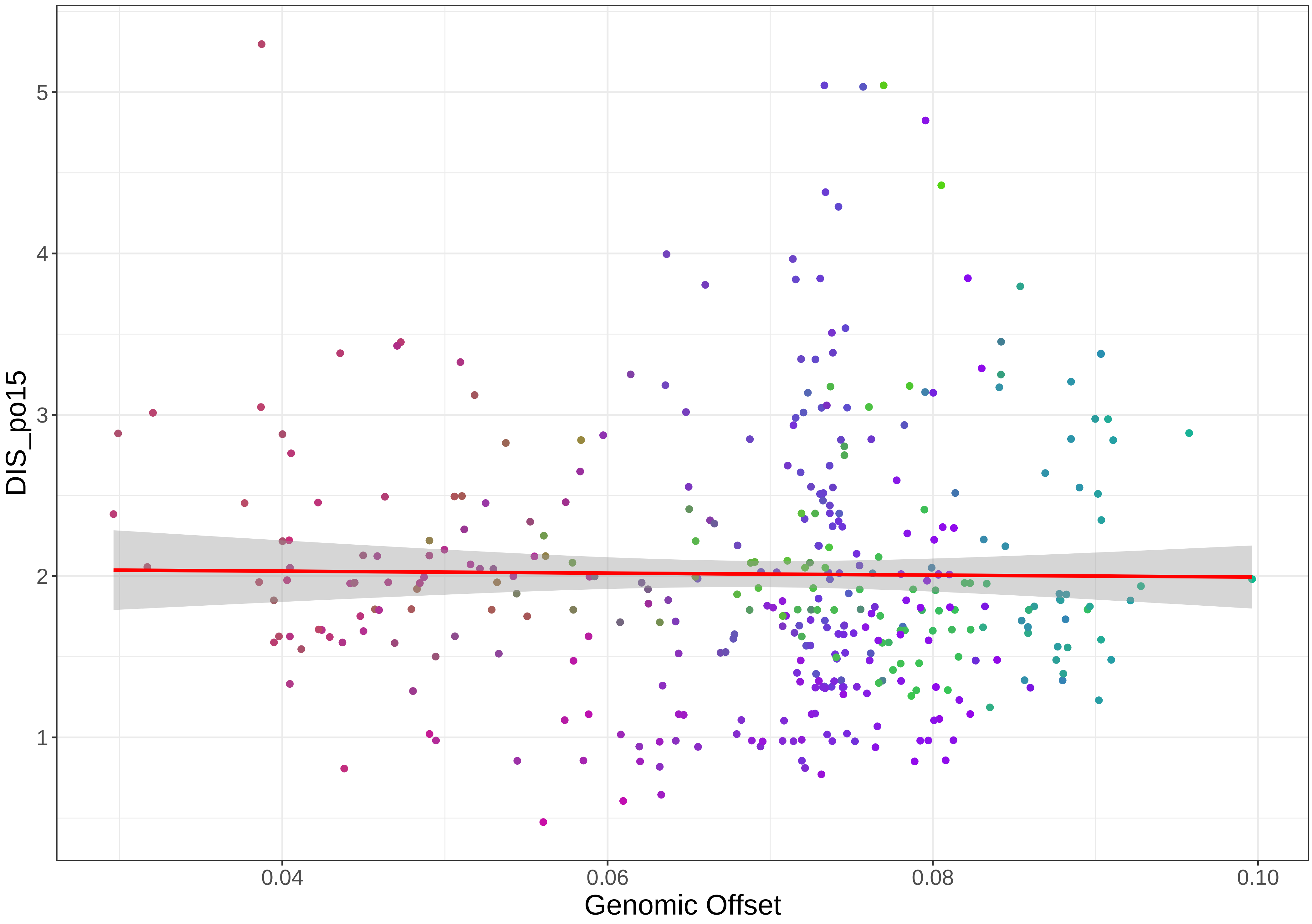
DIS_po15
Genomic Offset
1
2
3
4
5
0.04
0.06
0.08
0.10

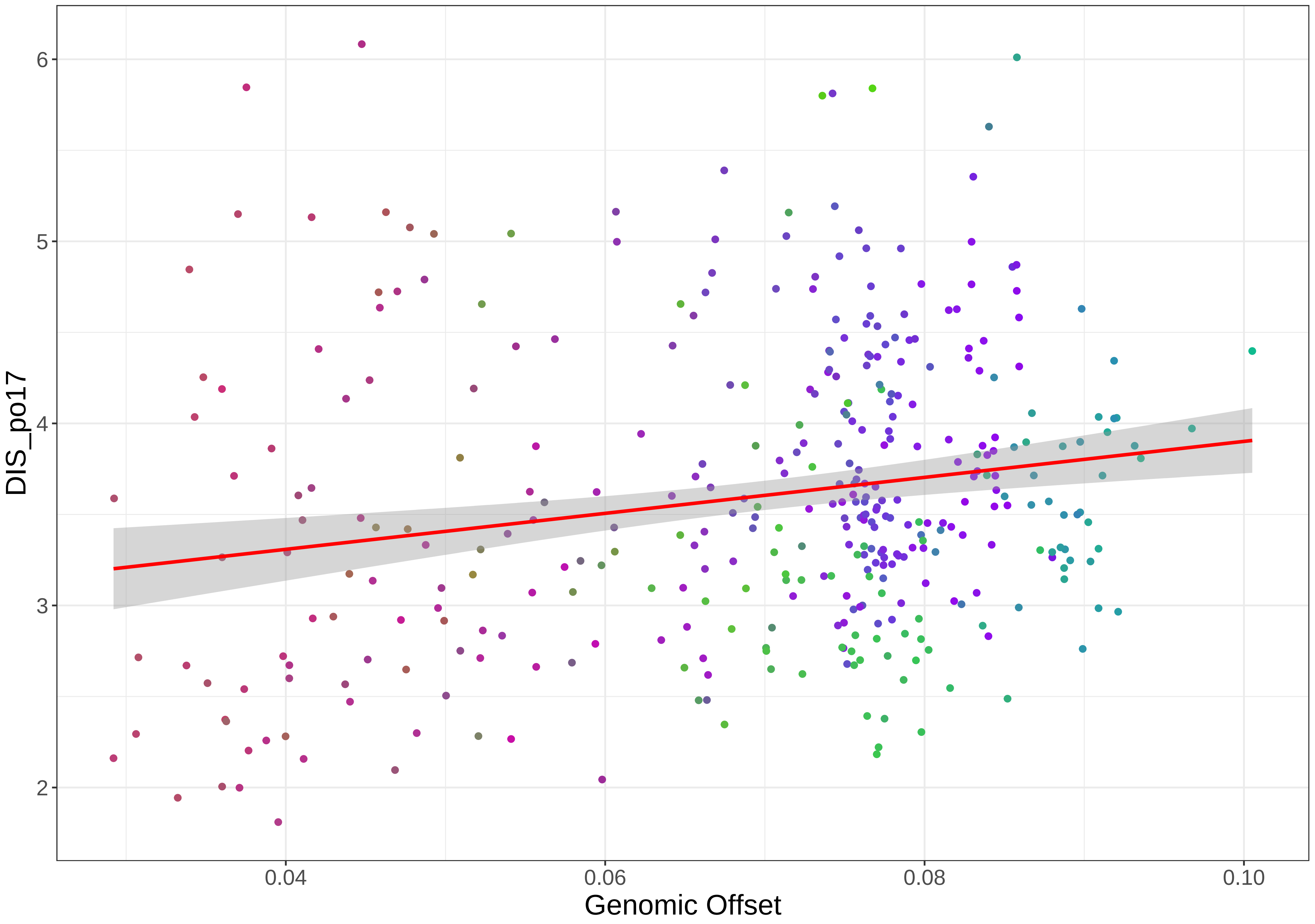
DIS_po17
6
5
4
3
2
0.04
0.06
0.08
0.10
Genomic Offset

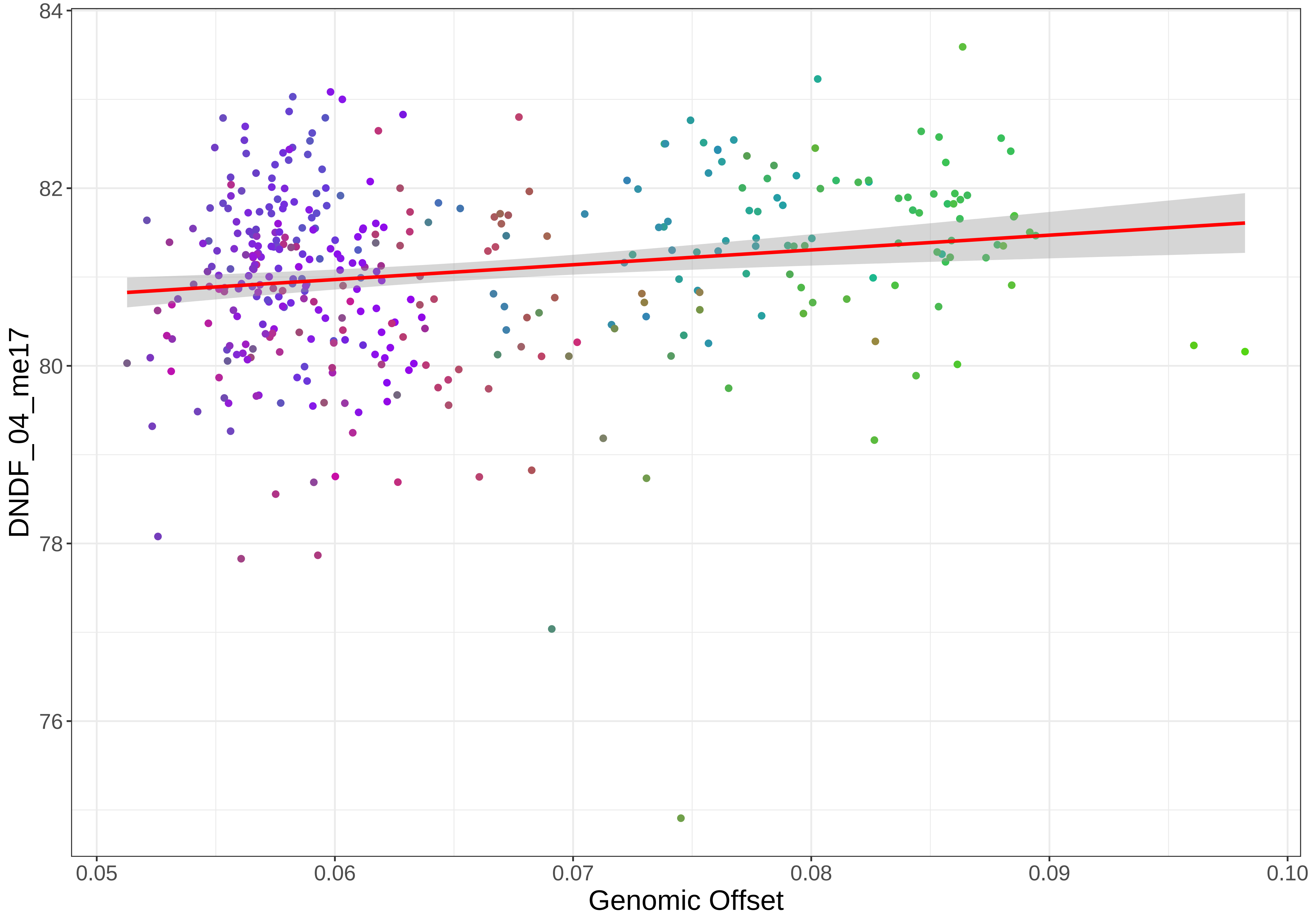
84
82
80
78
76
DNDF_04_me17
0.05
0.06
0.07
0.08
0.09
0.10
Genomic Offset

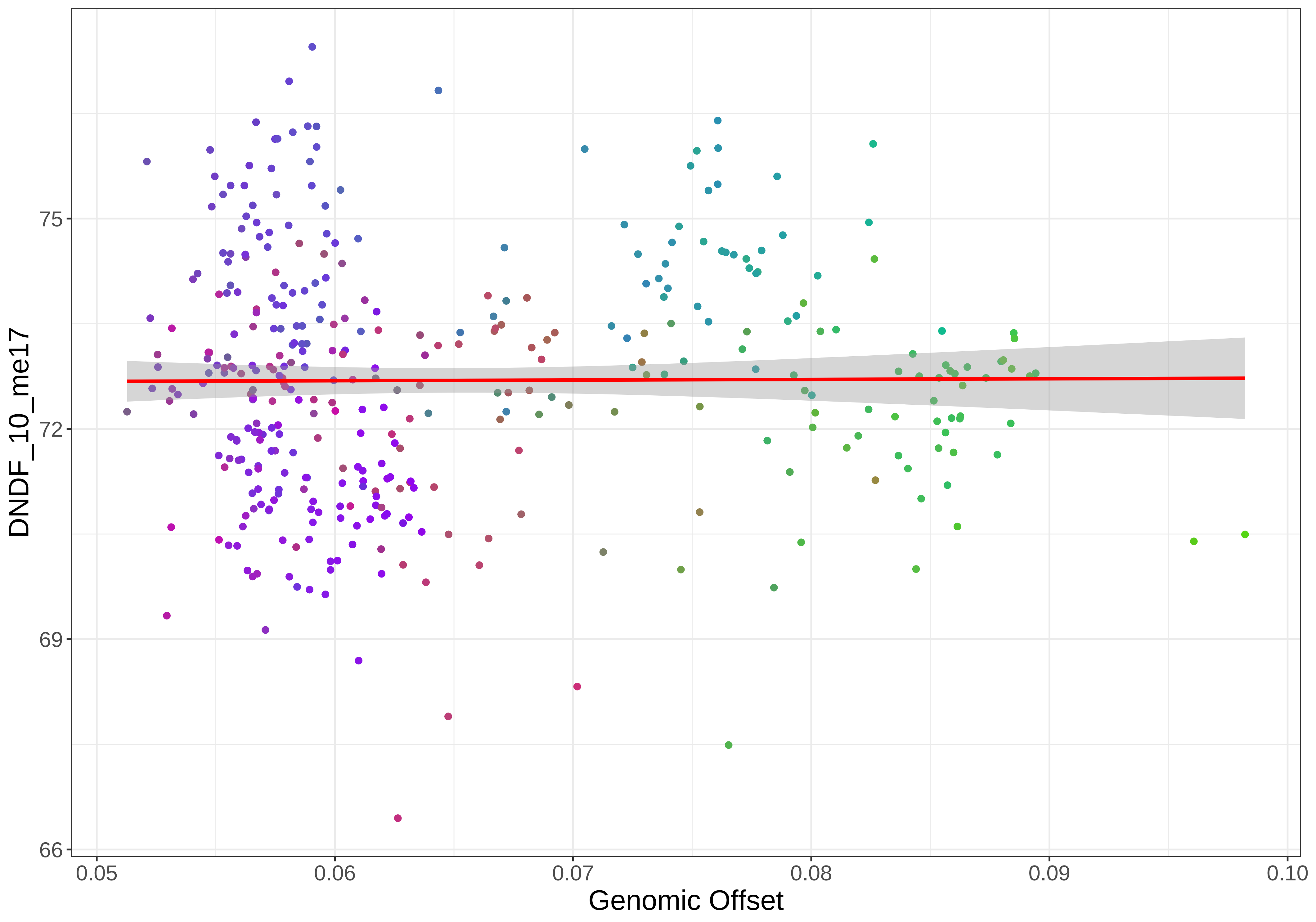

DNDF_10_me17
75
72
69
66
0.05
0.06
0.07
0.08
0.09
0.10
Genomic Offset

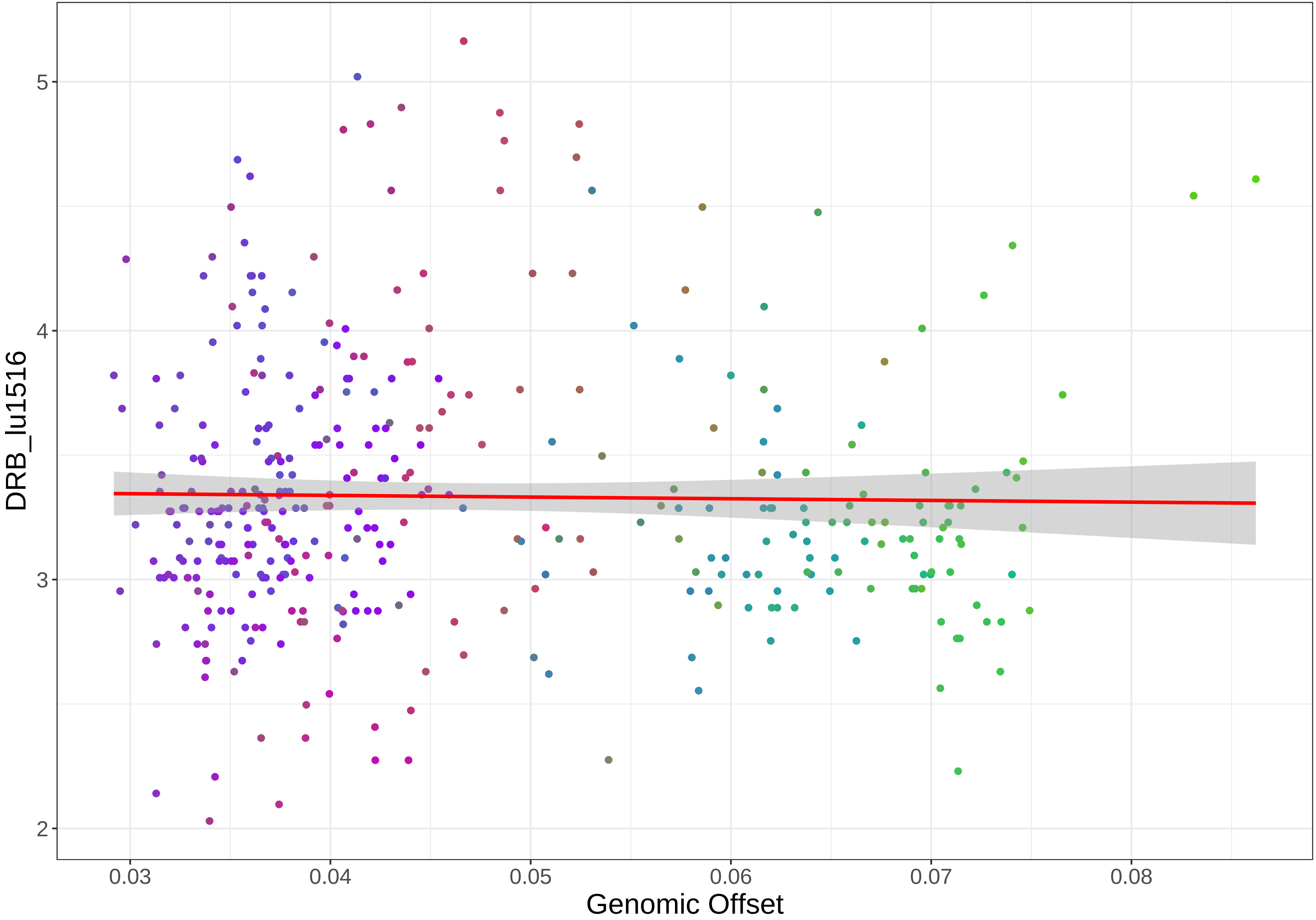

DRB_lu1516
Genomic Offset
5
4
3
2
0.03
0.04
0.05
0.06
0.07
0.08

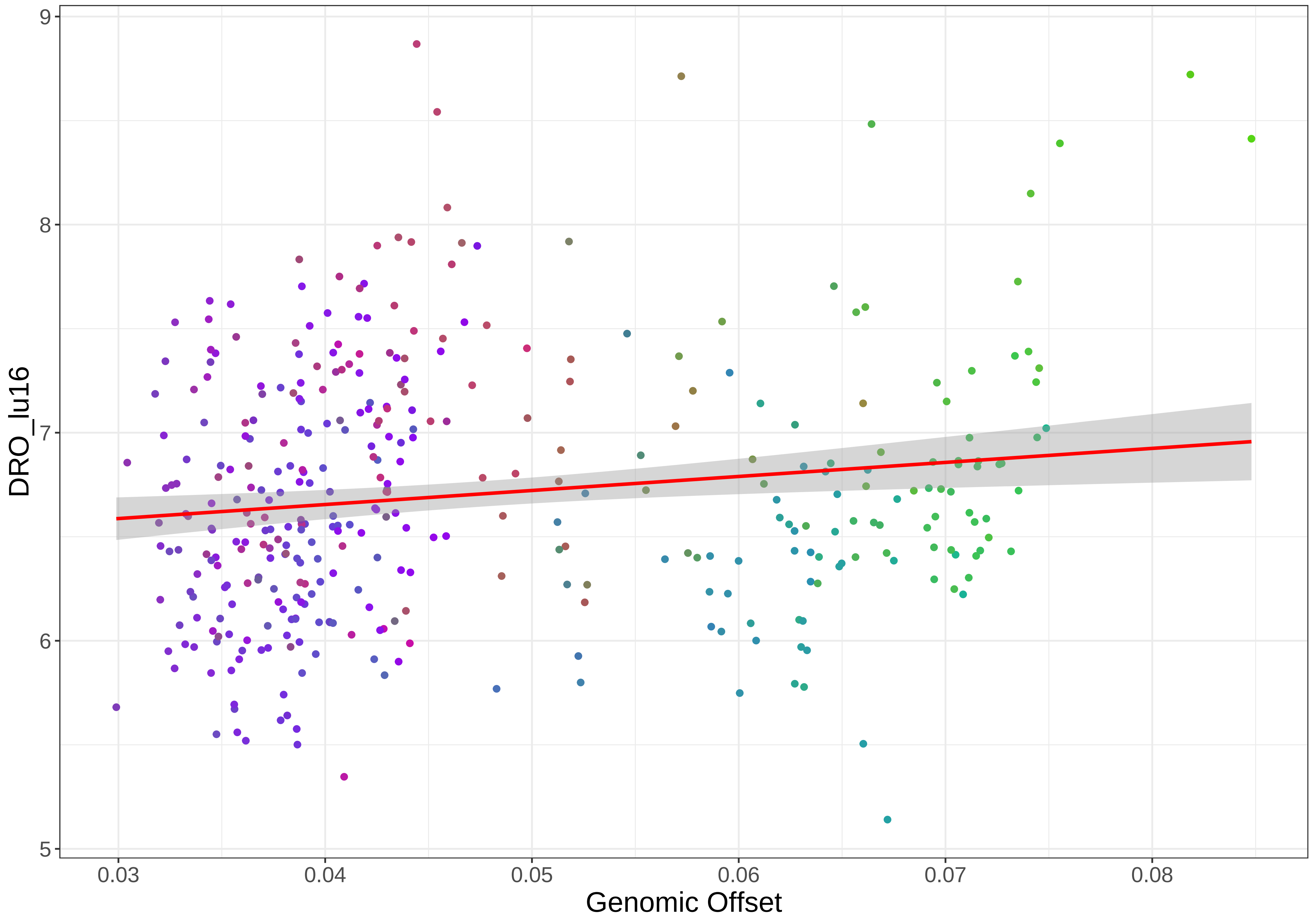

9
8
7
6
5
DRO_lu16
0.03
0.04
0.05
0.06
0.07
0.08
Genomic Offset

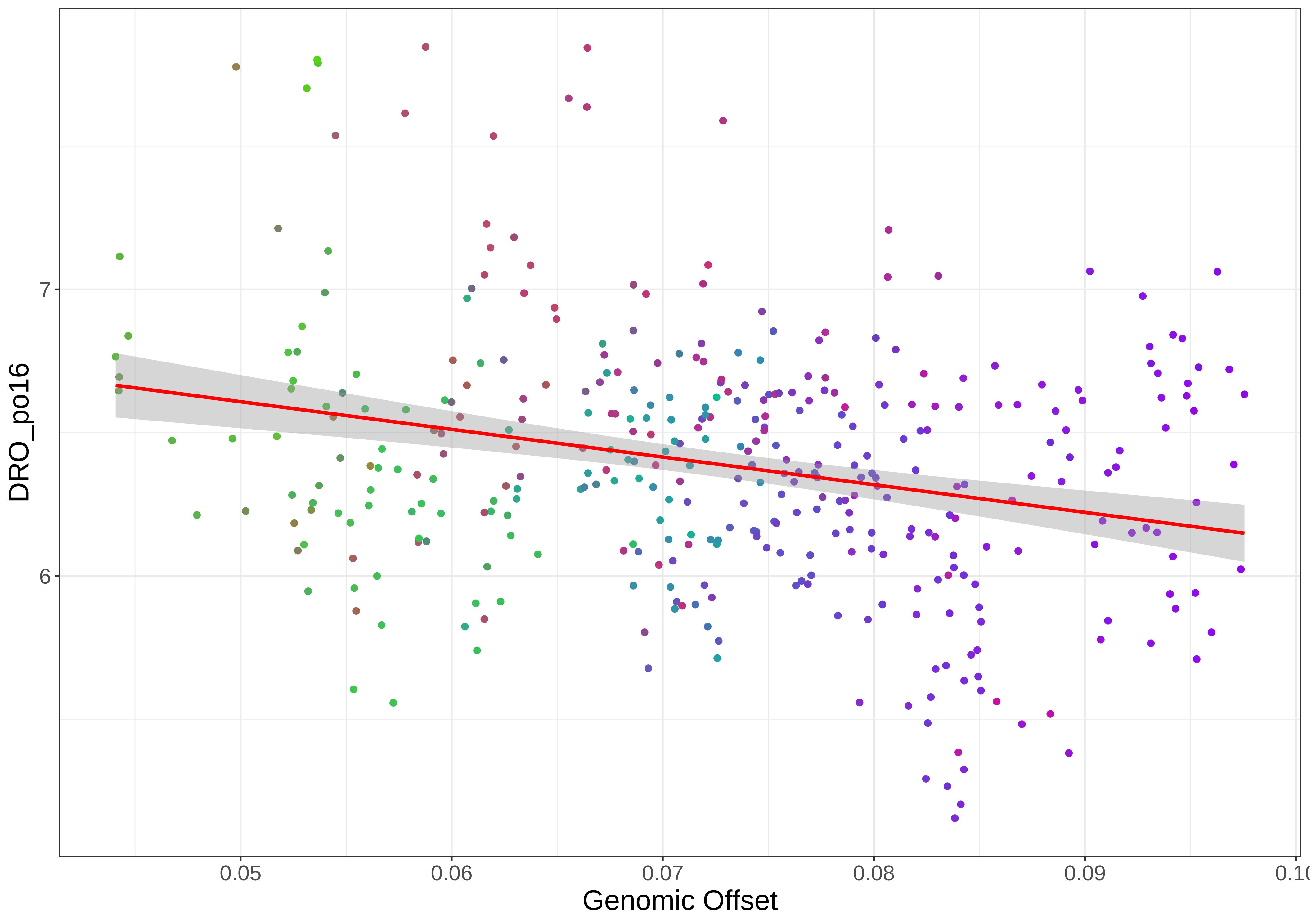

DRO_po16
7
6
0.05
0.06
0.07
0.08
0.09
Genomic Offset

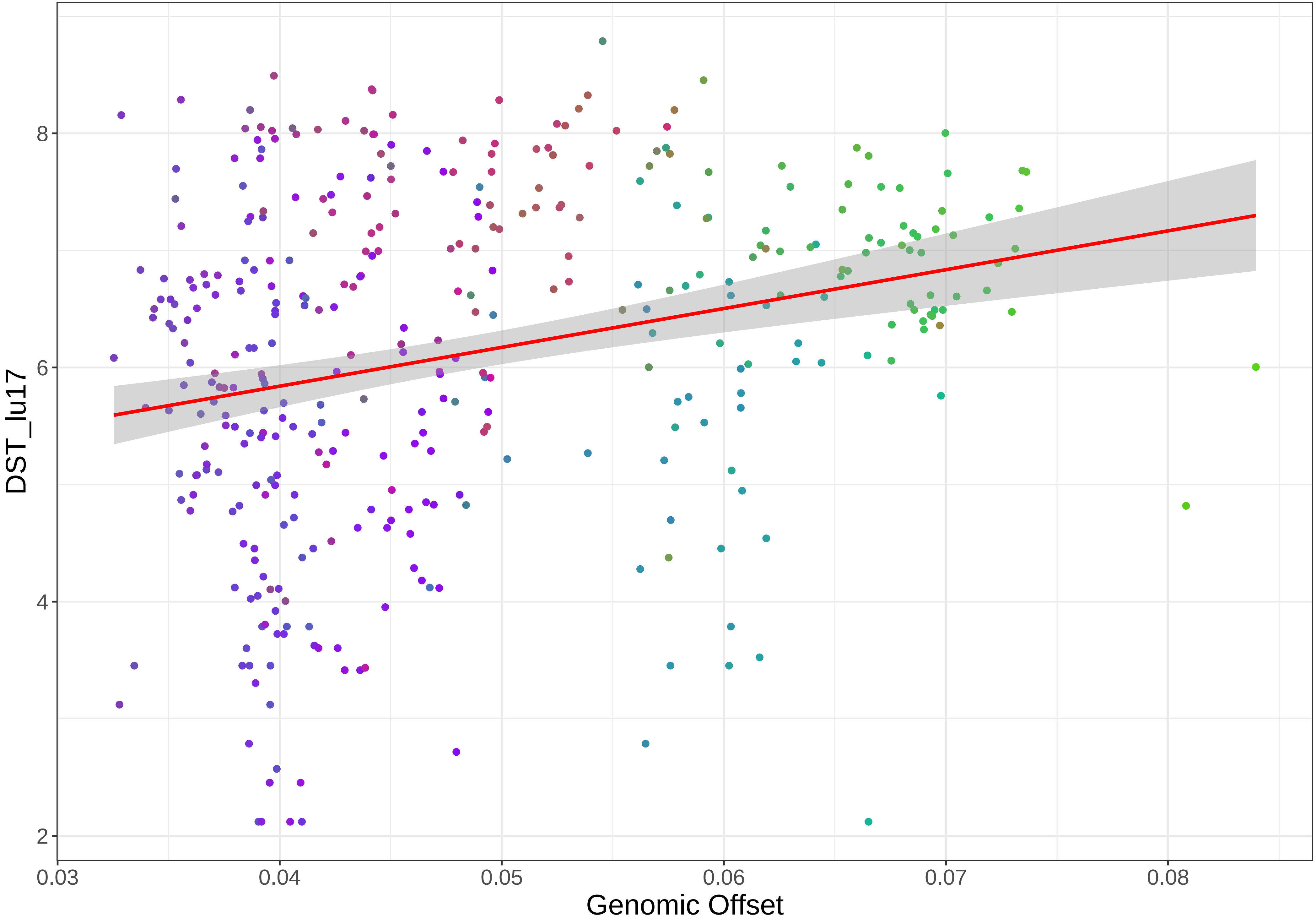
DST_lu17
8
6
4
2
0.03
0.04
0.05
0.06
0.07
0.08
Genomic Offset

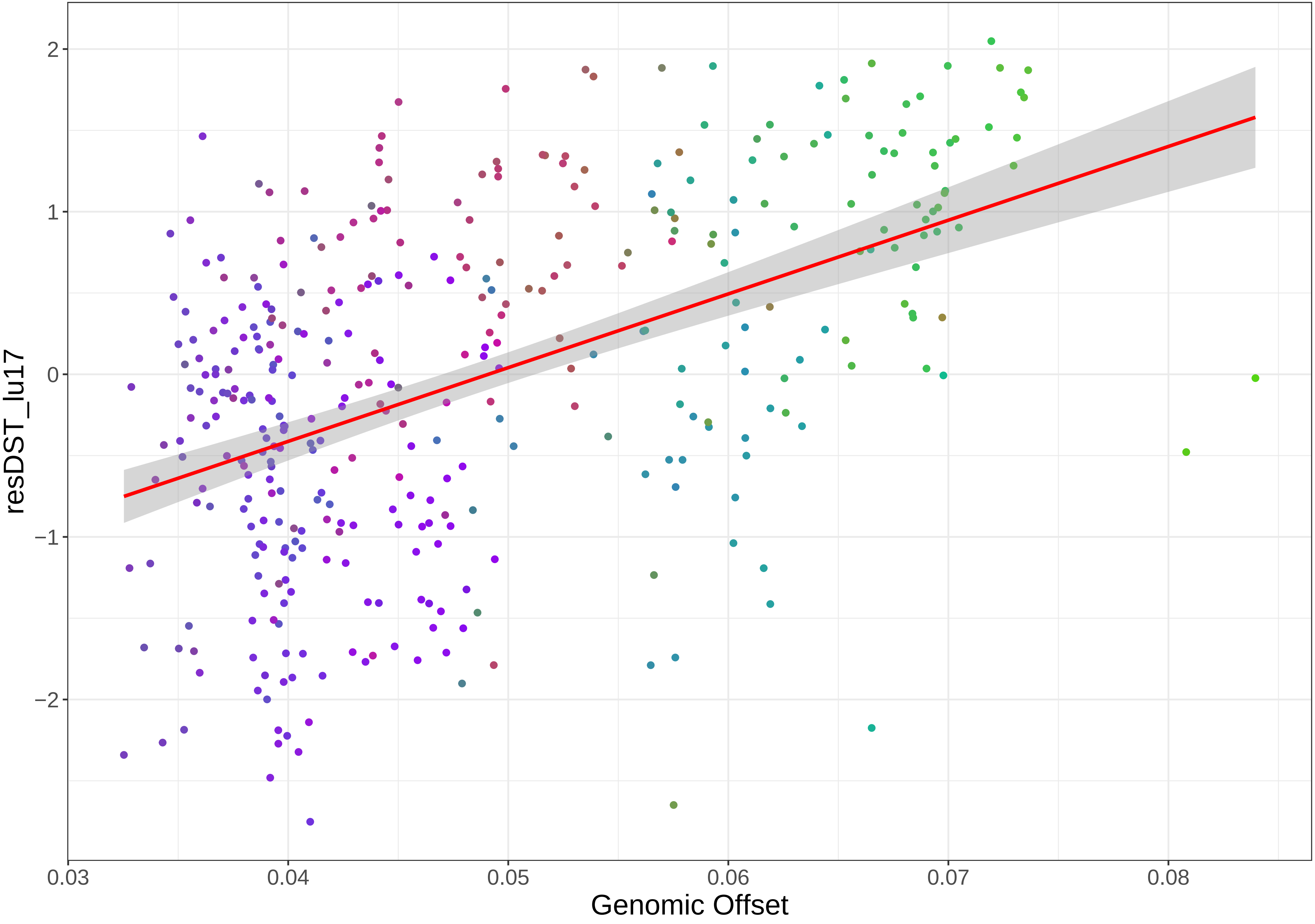
2
1
0
−1
−2
resDST_lu17
0.03
0.04
0.05
0.06
0.07
0.08
Genomic Offset

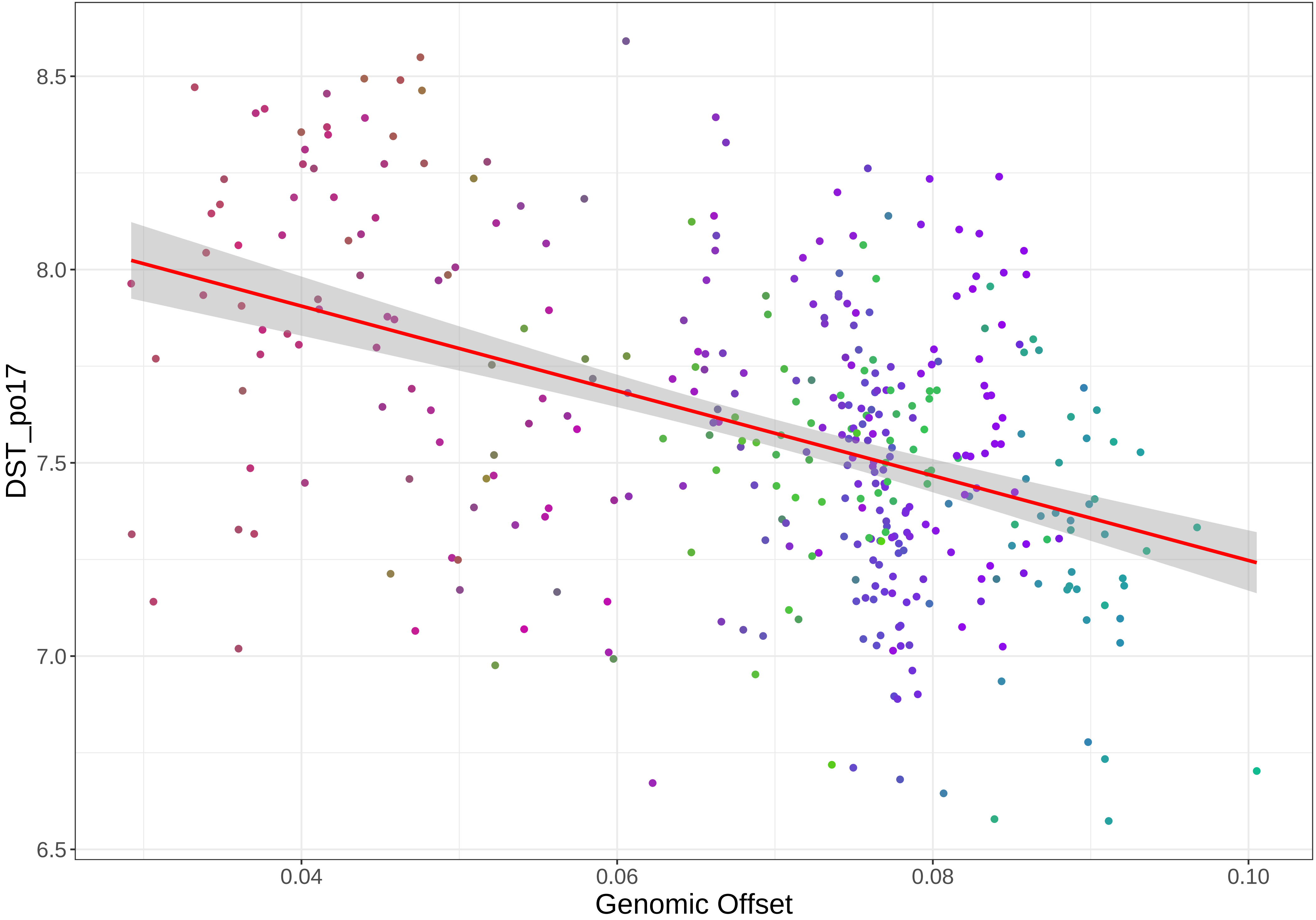
8.5
8.0
7.5
7.0
6.5
DST_po17
0.04
0.06
0.08
0.10
Genomic Offset

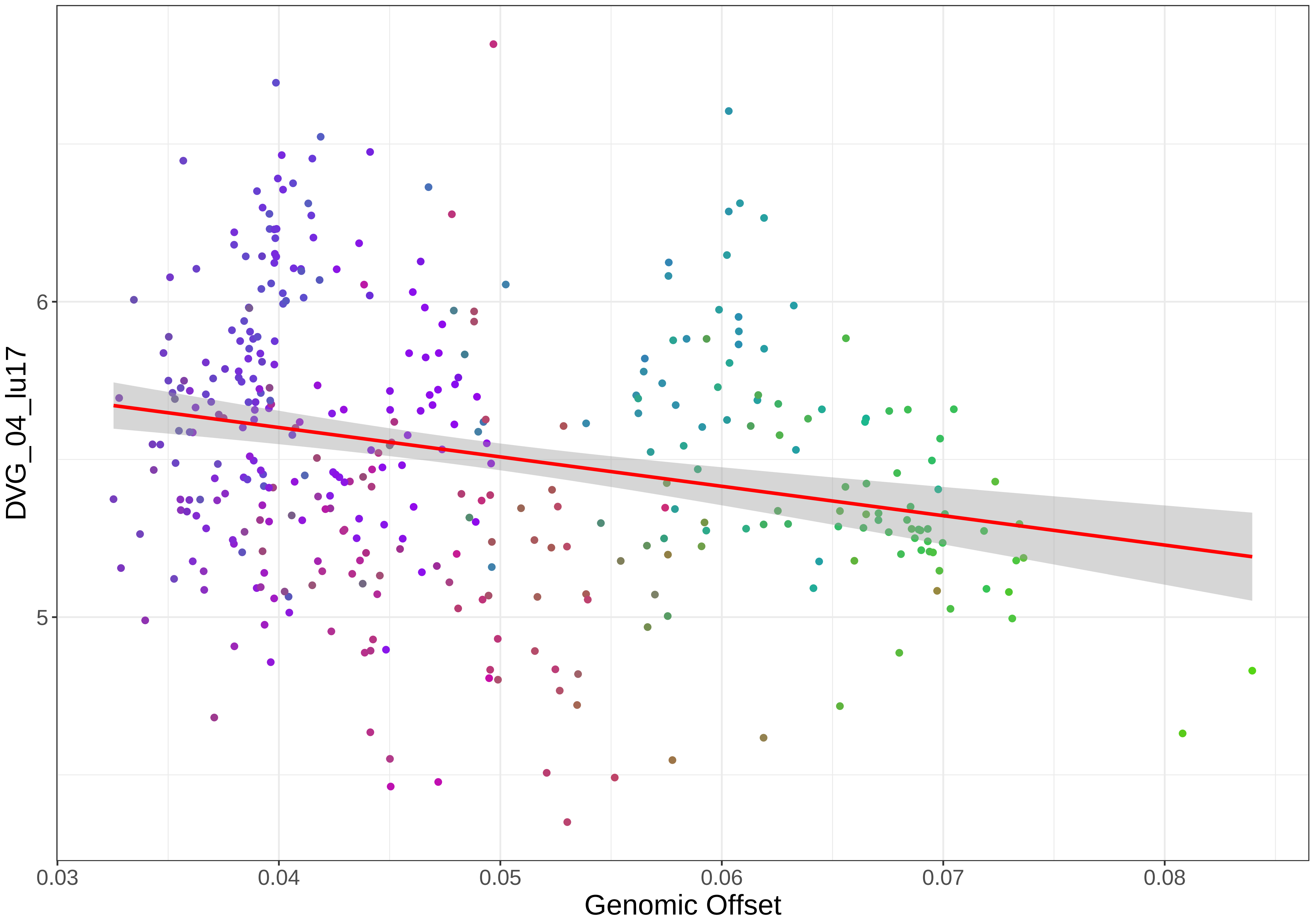

DVG_04_lu17
6
5
0.03
0.04
0.05
0.06
0.07
0.08
Genomic Offset

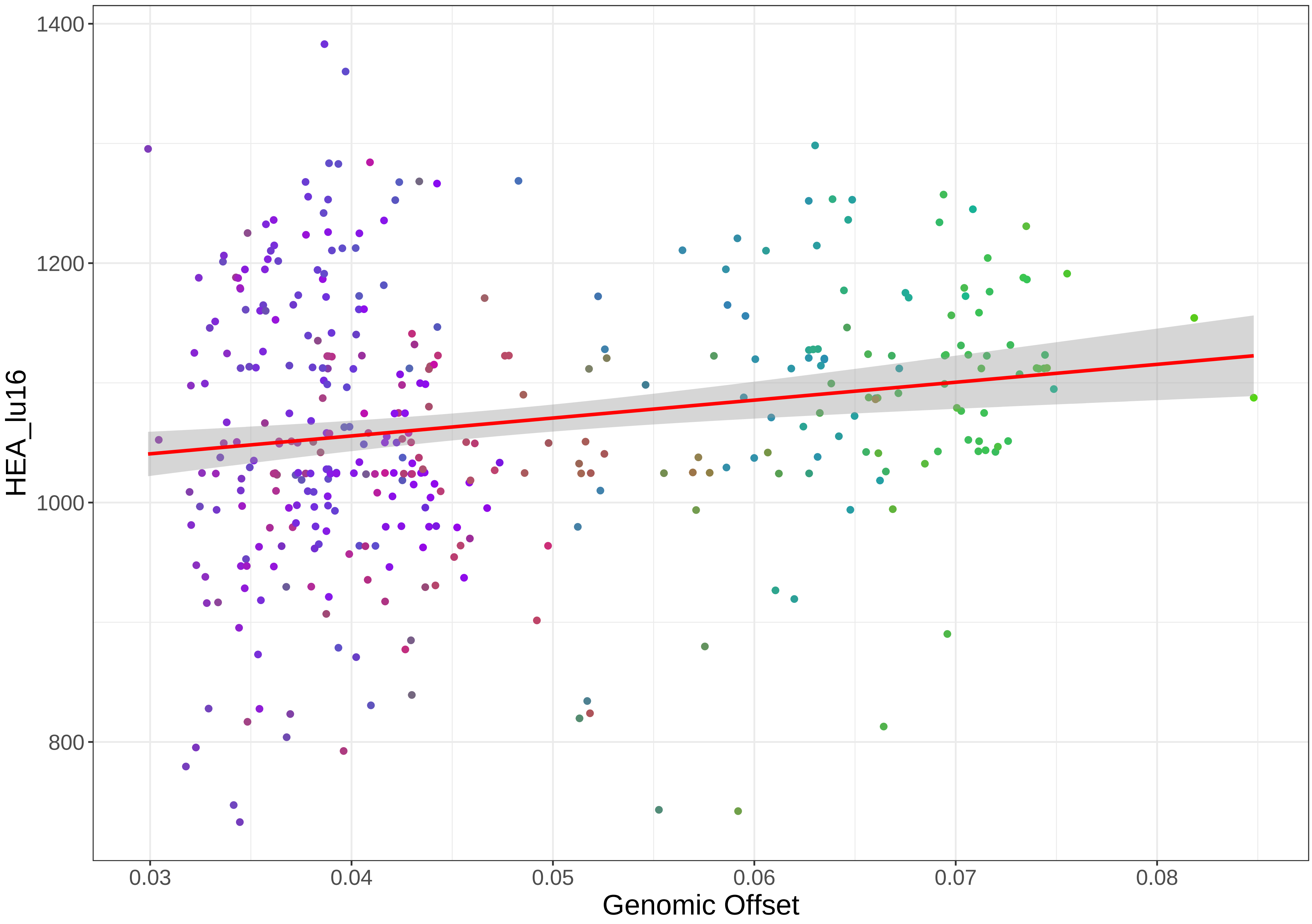

HEA_lu16
Genomic Offset
1400
1200
1000
800
0.03
0.04
0.05
0.06
0.07
0.08

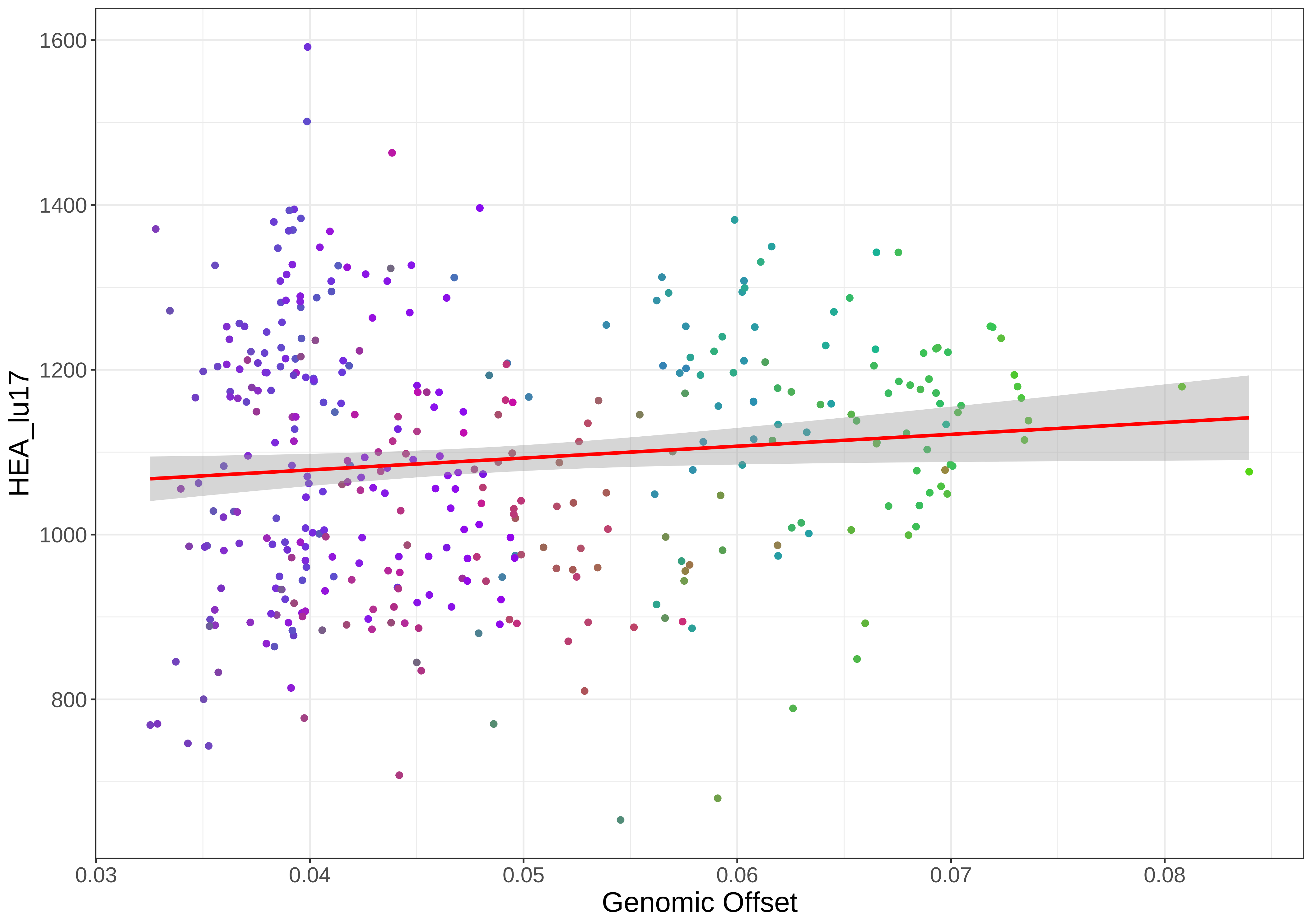

1600
1400
1200
1000
800
HEA_lu17
0.03
0.04
0.05
0.06
0.07
0.08
Genomic Offset

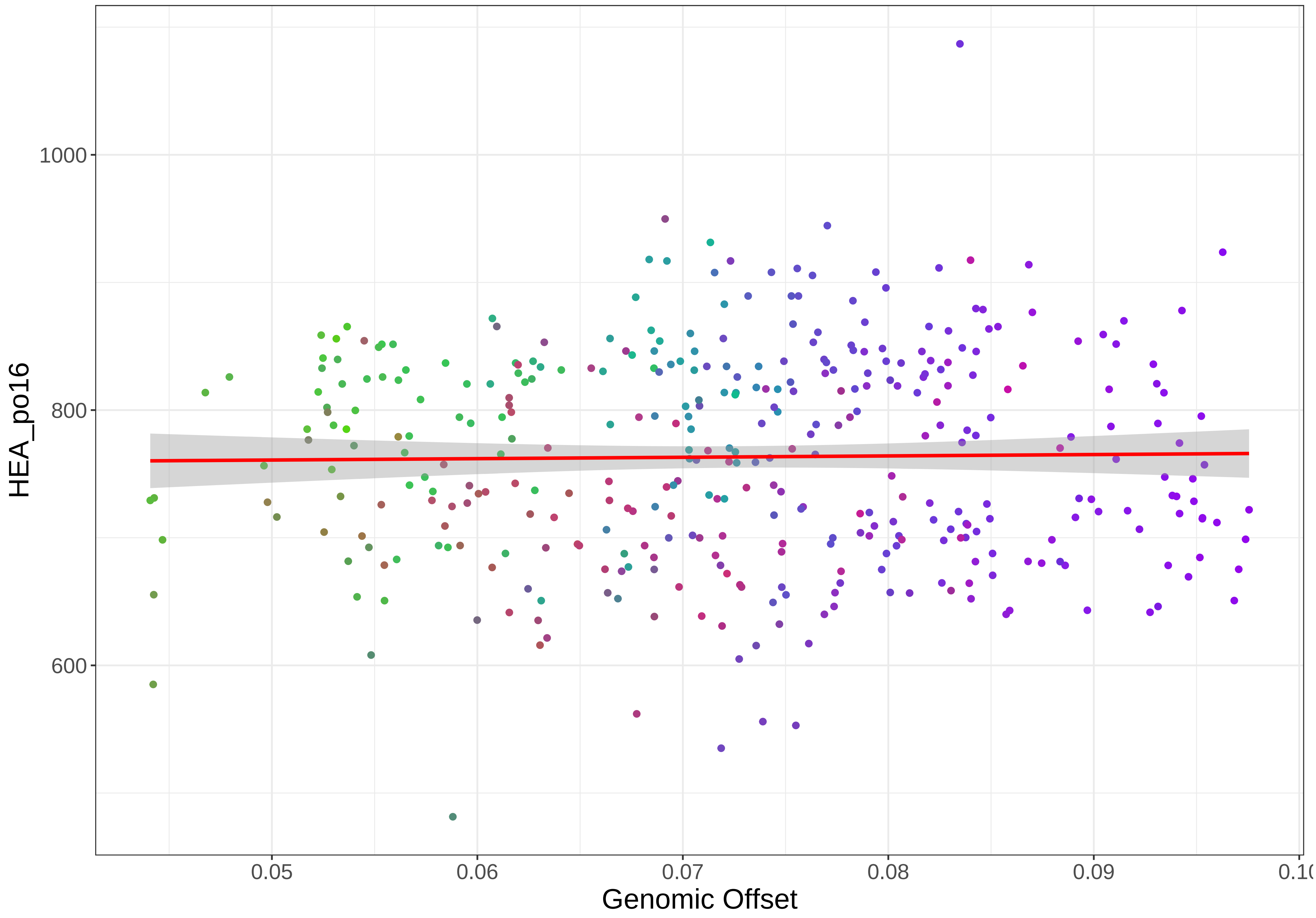

HEA_po16
1000
800
600
0.05
0.06
0.07
0.08
0.09
Genomic Offset

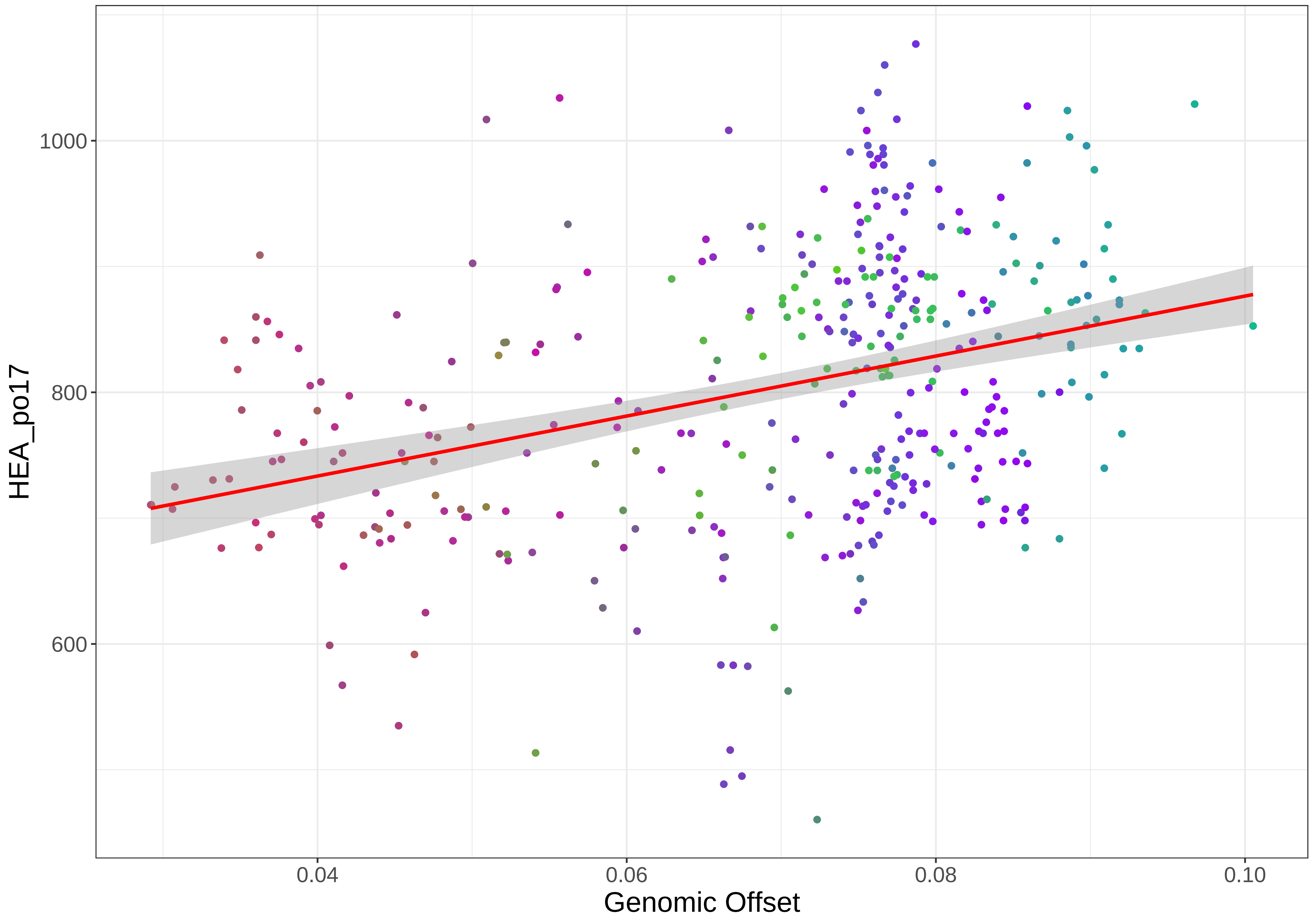

1000
800
600
HEA_po17
0.04
0.06
0.08
0.10
Genomic Offset

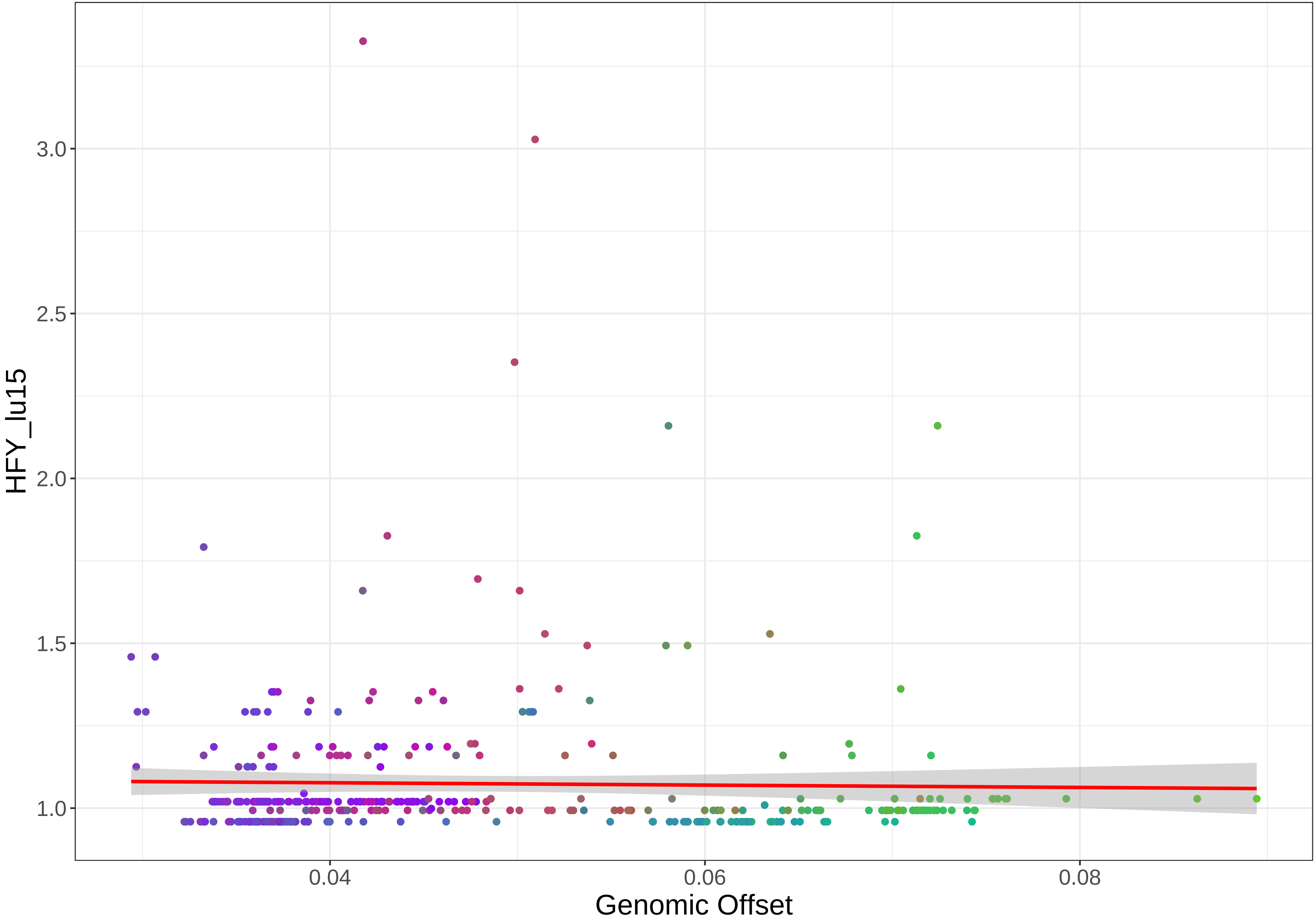

HFY_lu15
3.0
2.5
2.0
1.5
1.0
0.04
0.06
0.08
Genomic Offset

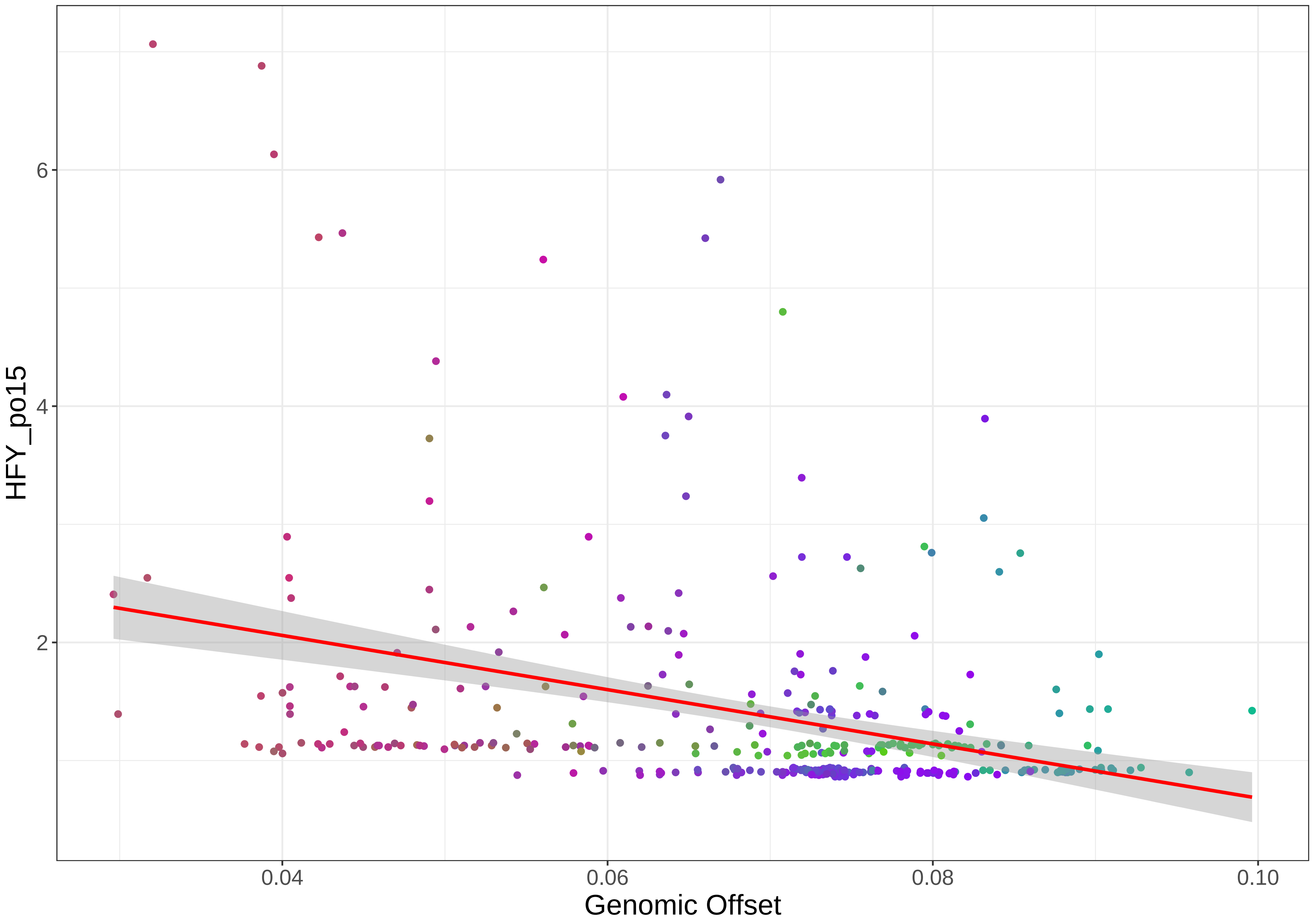

HFY_po15
6
4
2
0.04
0.06
0.08
0.10
Genomic Offset

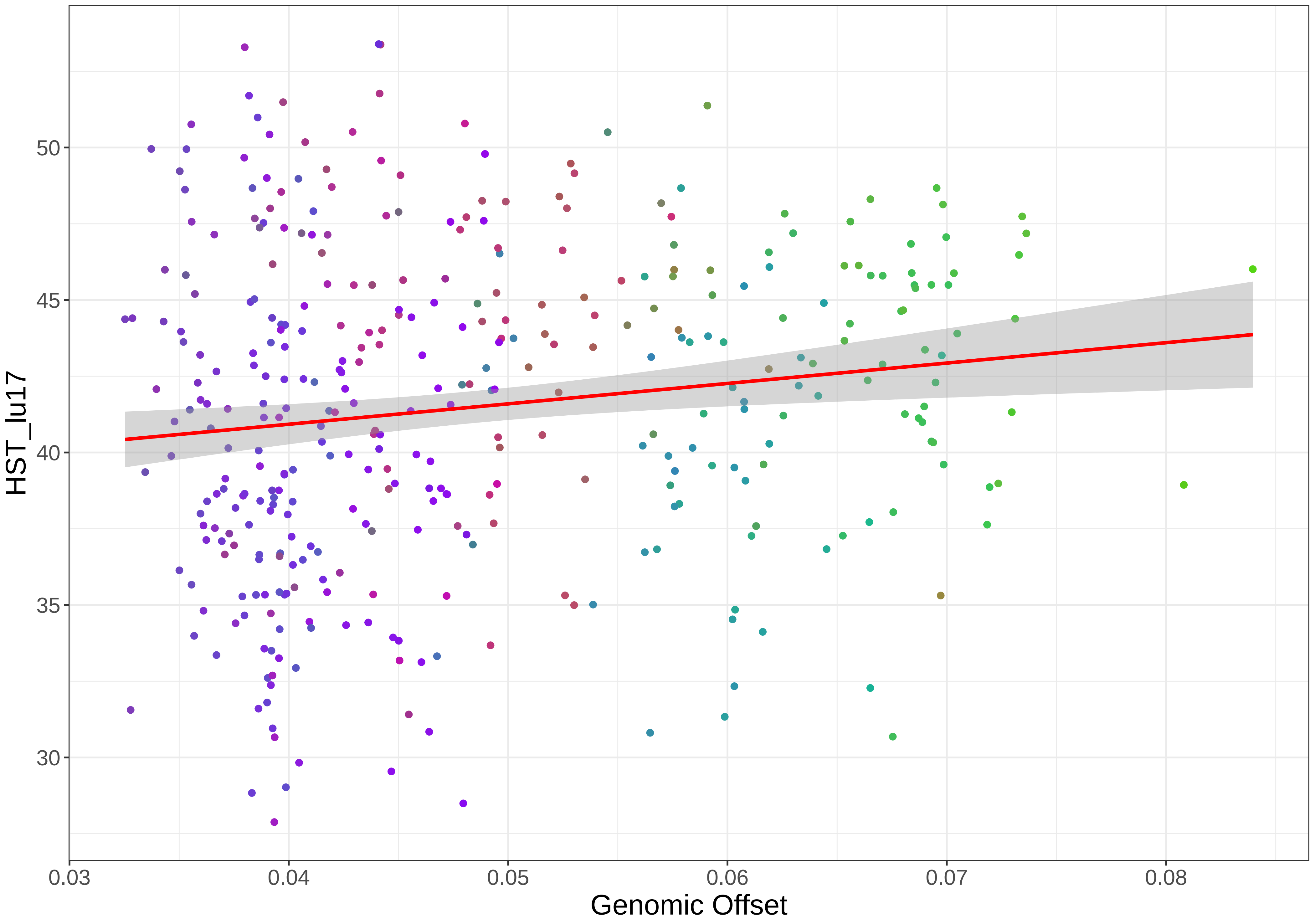

HST_lu17
Genomic Offset
50
45
40
35
30
0.03
0.04
0.05
0.06
0.07
0.08

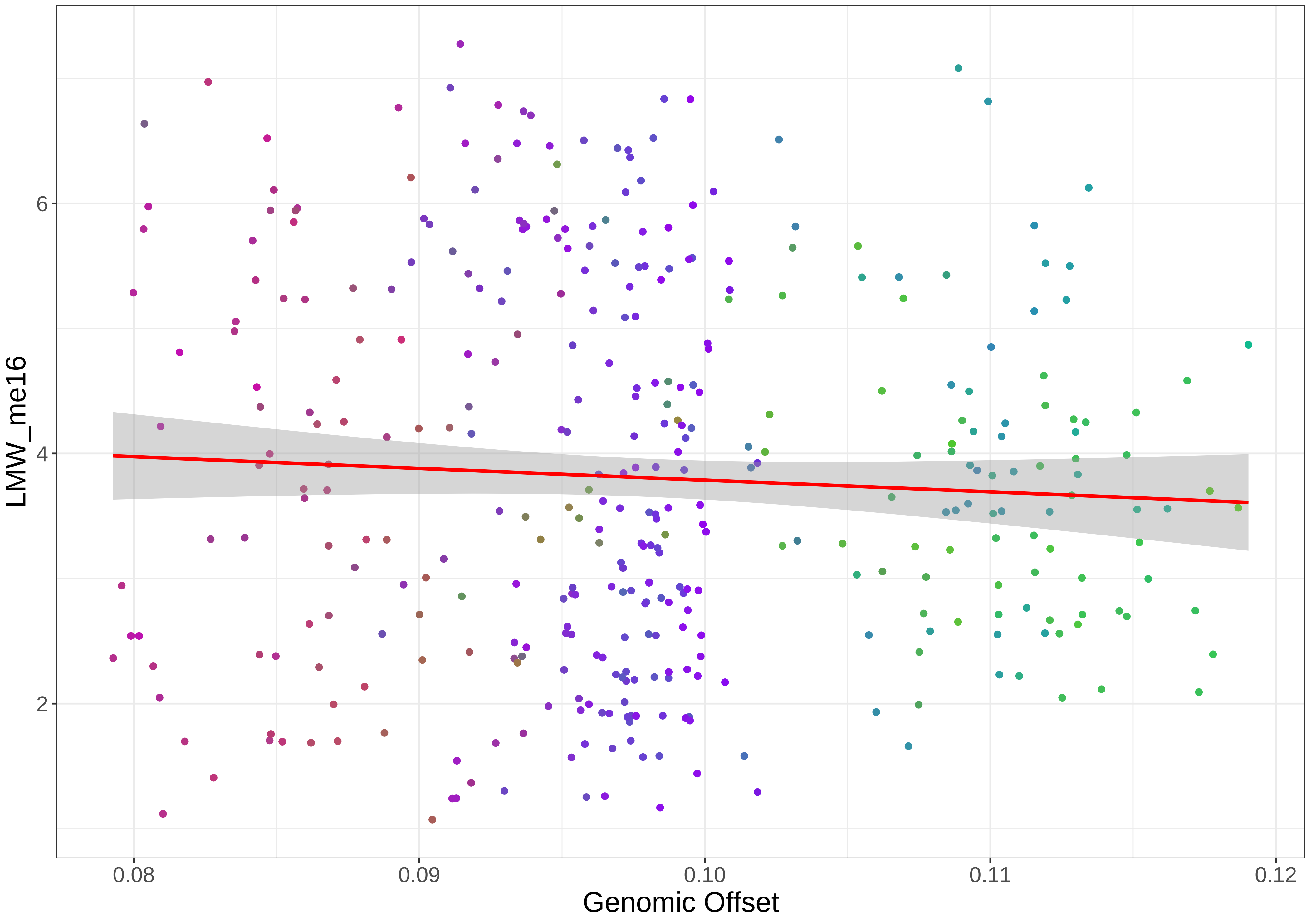

LMW_me16
6
4
2
0.08
0.09
0.10
0.11
0.12
Genomic Offset

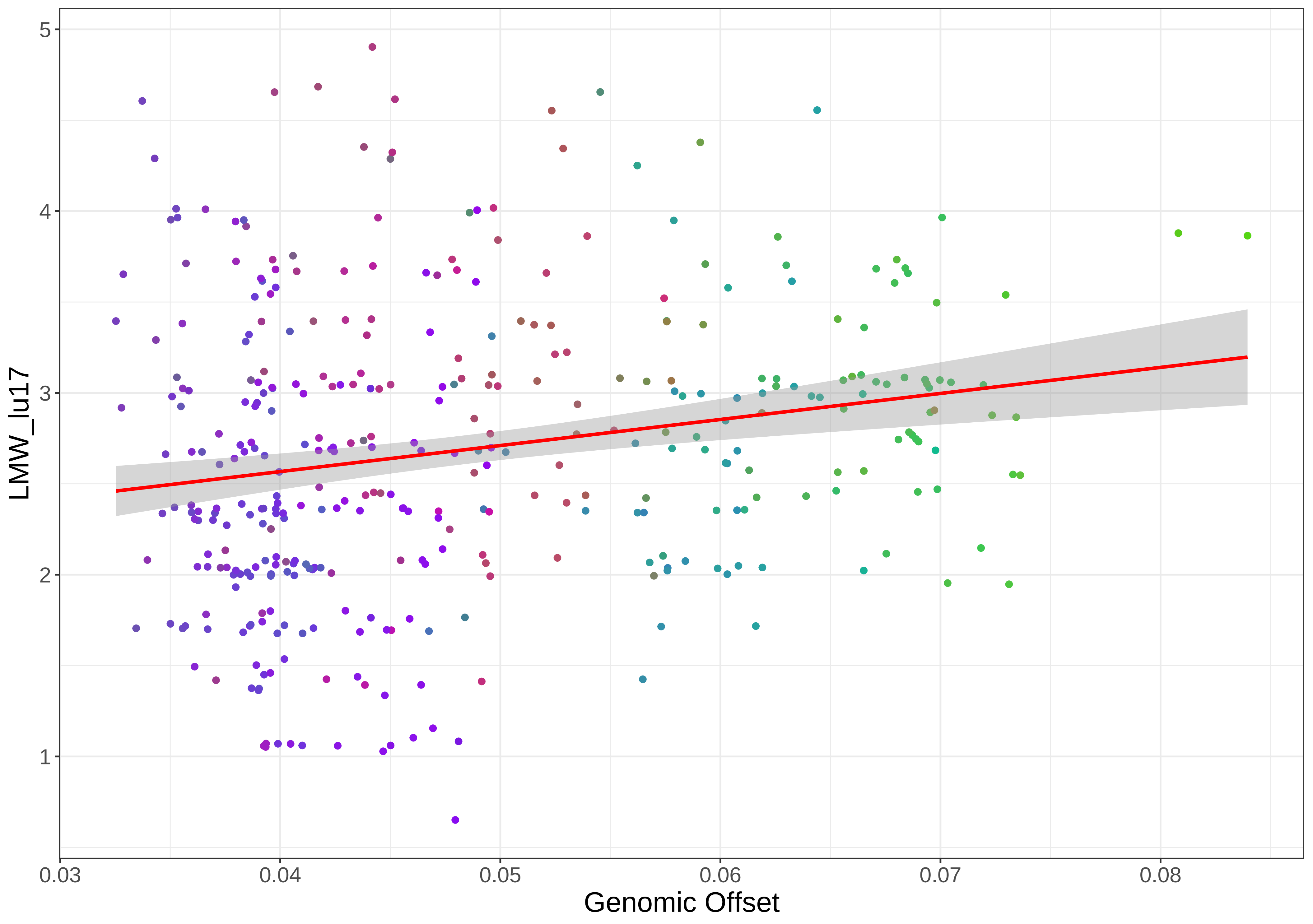

LMW_lu17
1
2
3
4
5
0.03
0.04
0.05
0.06
0.07
0.08
Genomic Offset

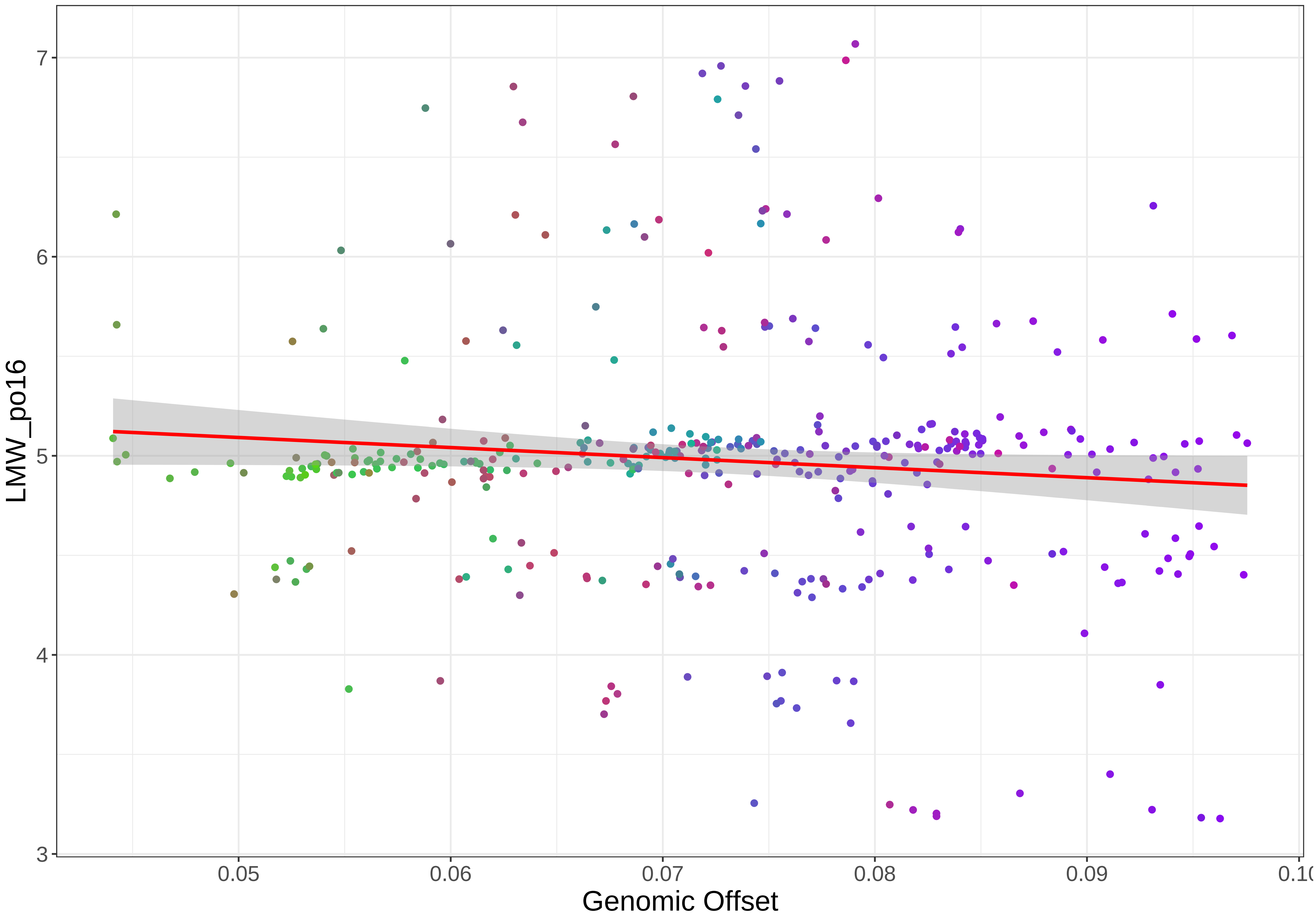
LMW_po16
Genomic Offset
3
4
5
6
7
0.05
0.06
0.07
0.08
0.09

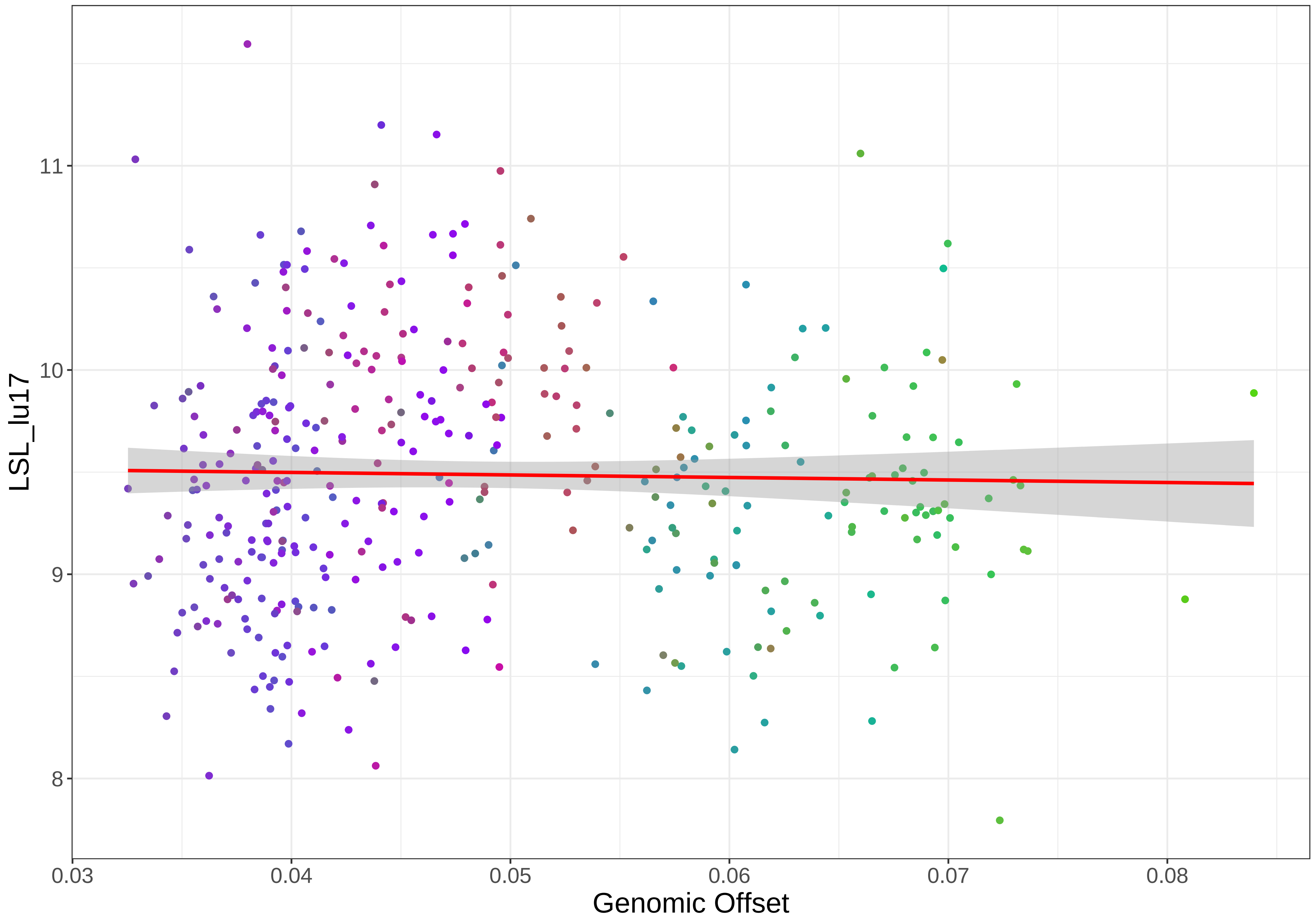
LSL_lu17
11
10
9
8
0.03
0.04
0.05
0.06
0.07
0.08
Genomic Offset

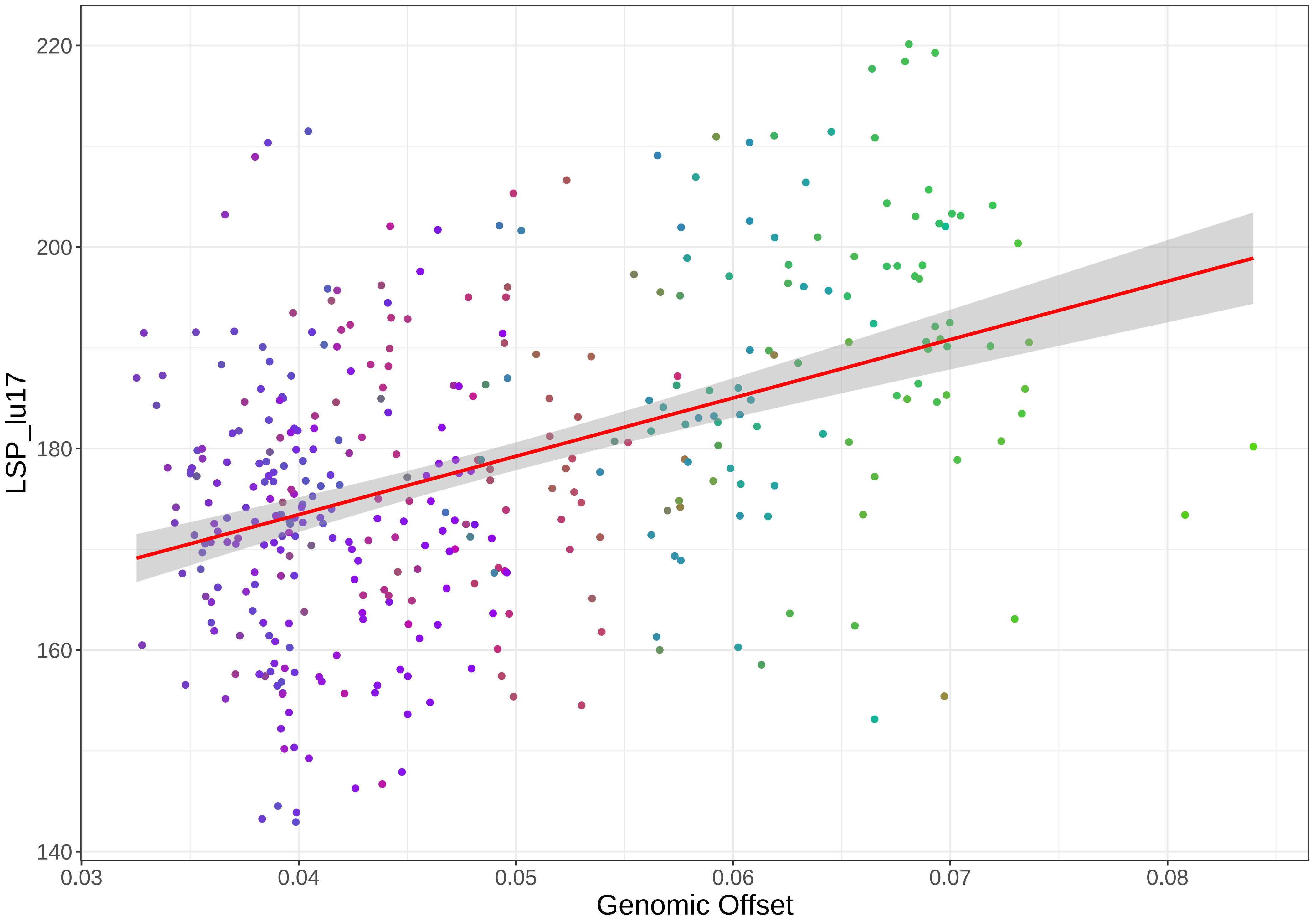

220
200
180
160
140
LSP_lu17
0.03
0.04
0.05
0.06
0.07
0.08
Genomic Offset

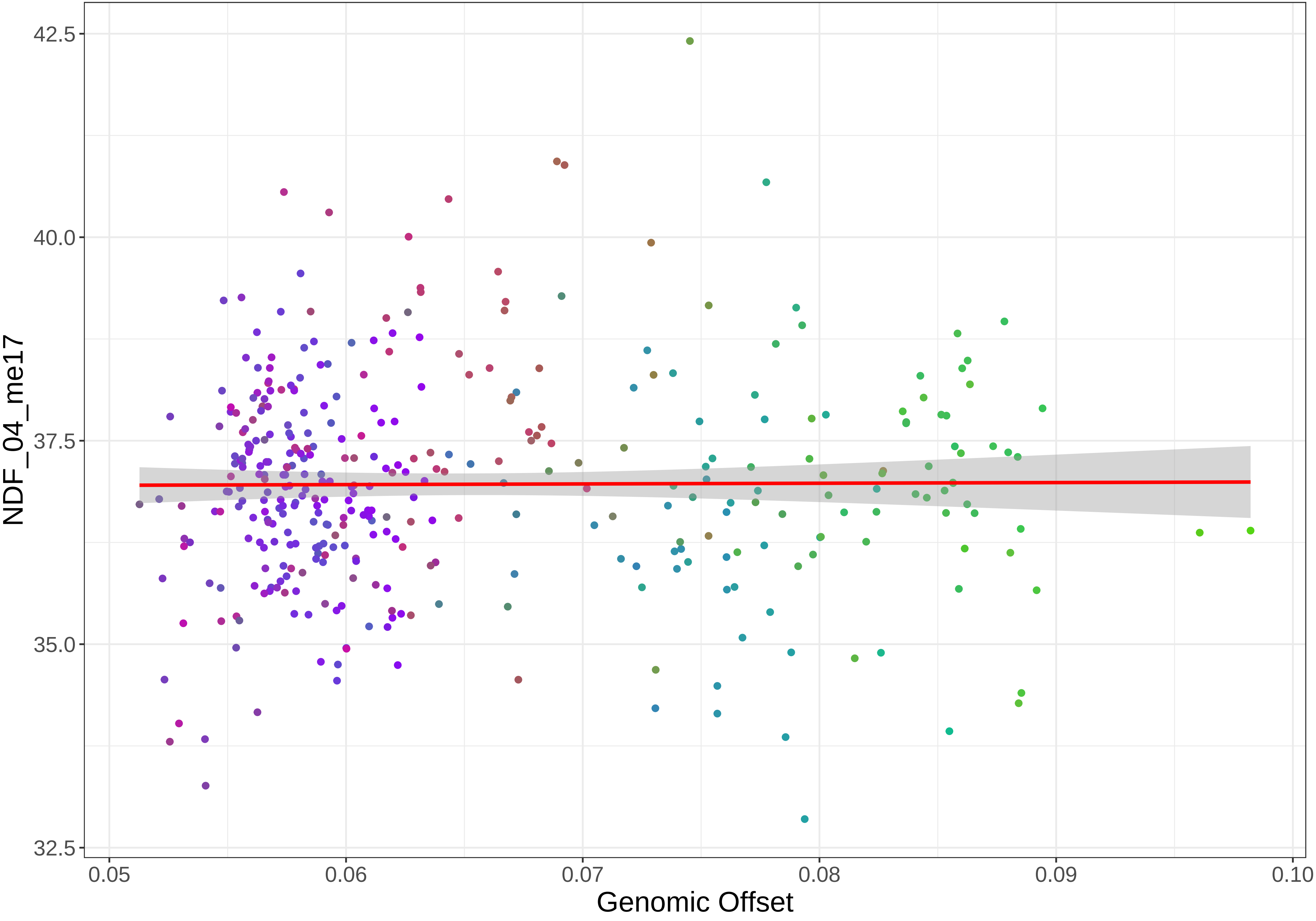
NDF_04_me17
42.5
40.0
37.5
35.0
32.5
0.05
0.06
0.07
0.08
0.09
0.10
Genomic Offset

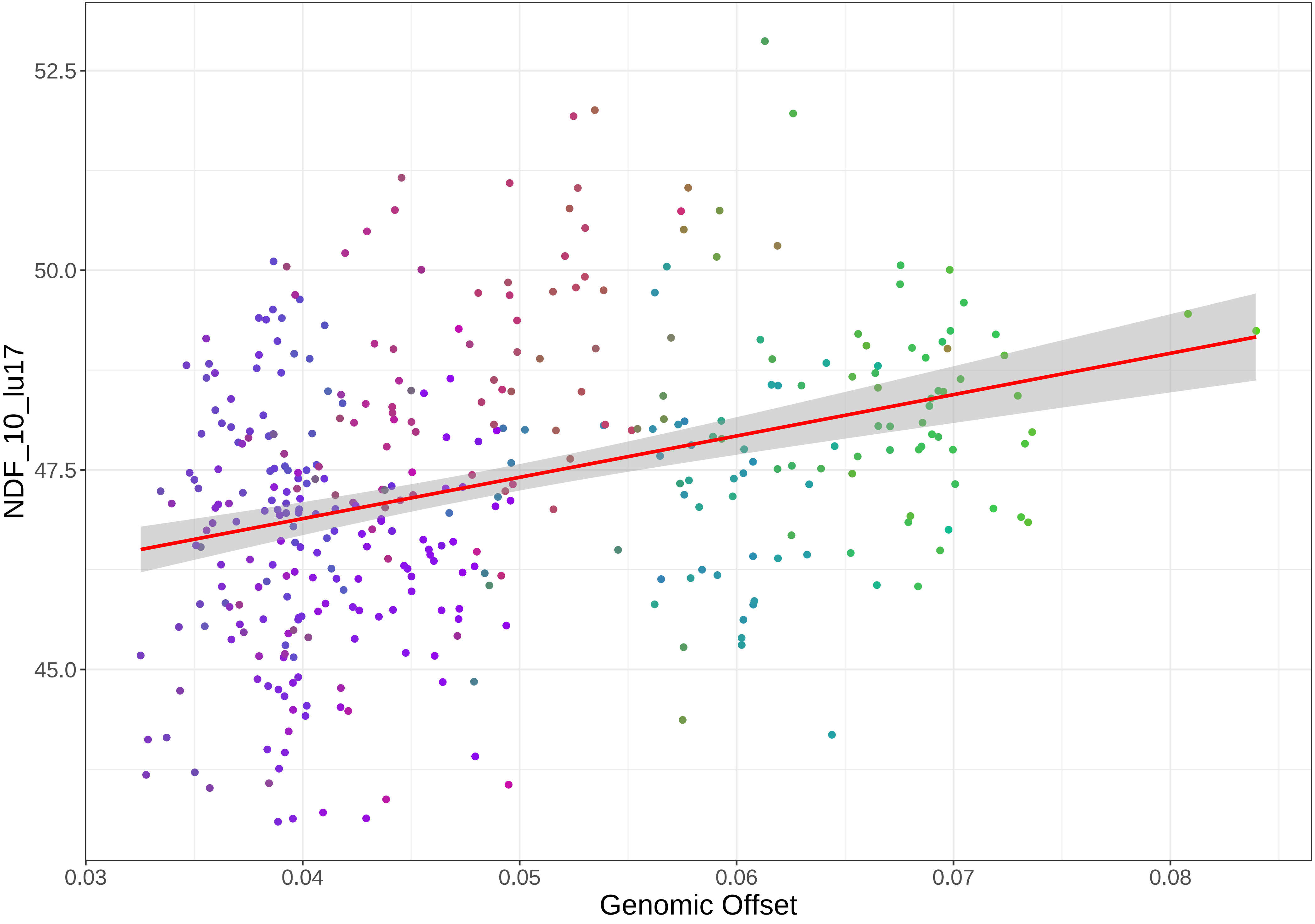

52.5
50.0
47.5
45.0
NDF_10_lu17
0.03
0.04
0.05
0.06
0.07
0.08
Genomic Offset

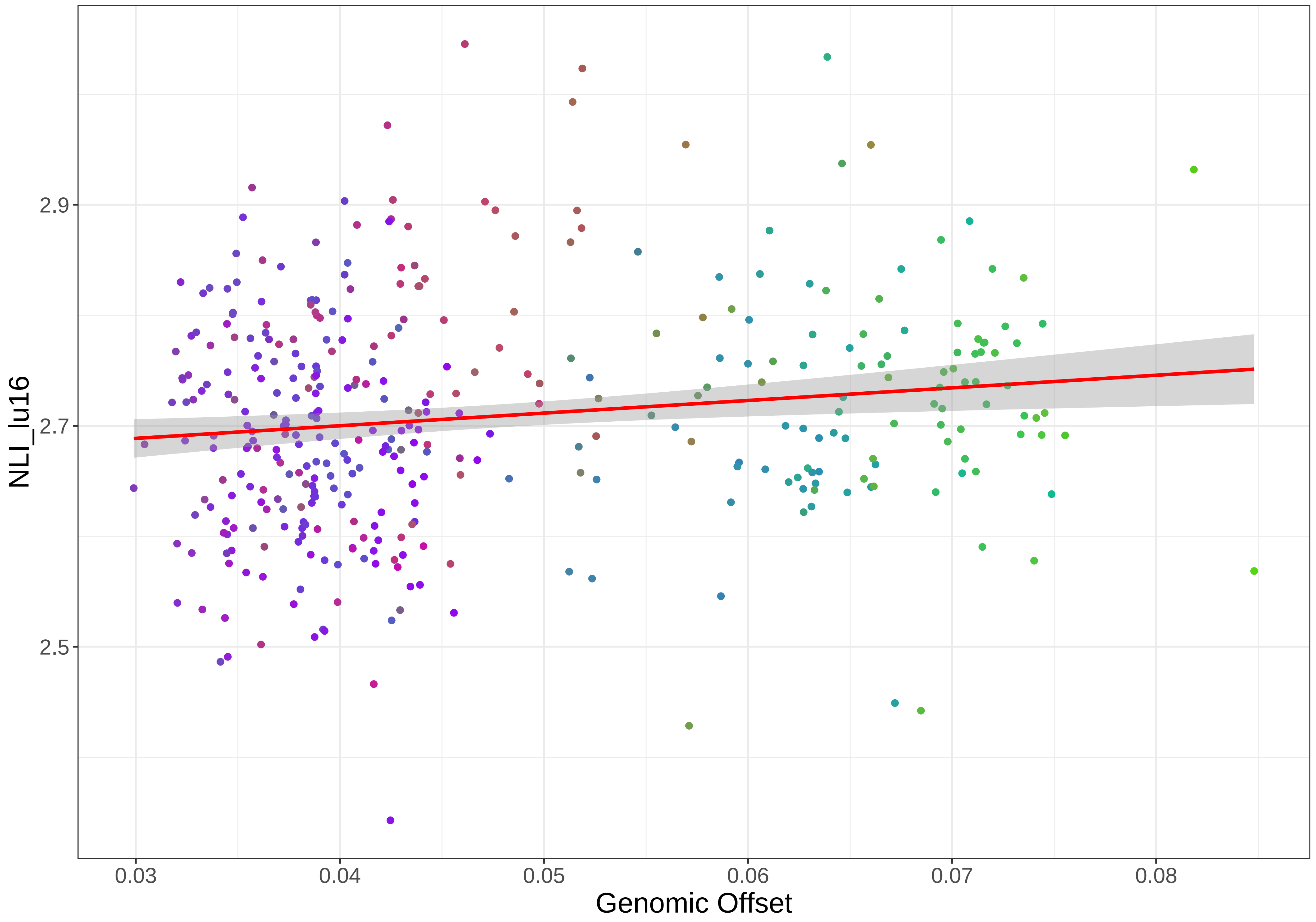

NLI_lu16
2.9
2.7
2.5
0.03
0.04
0.05
0.06
0.07
0.08
Genomic Offset

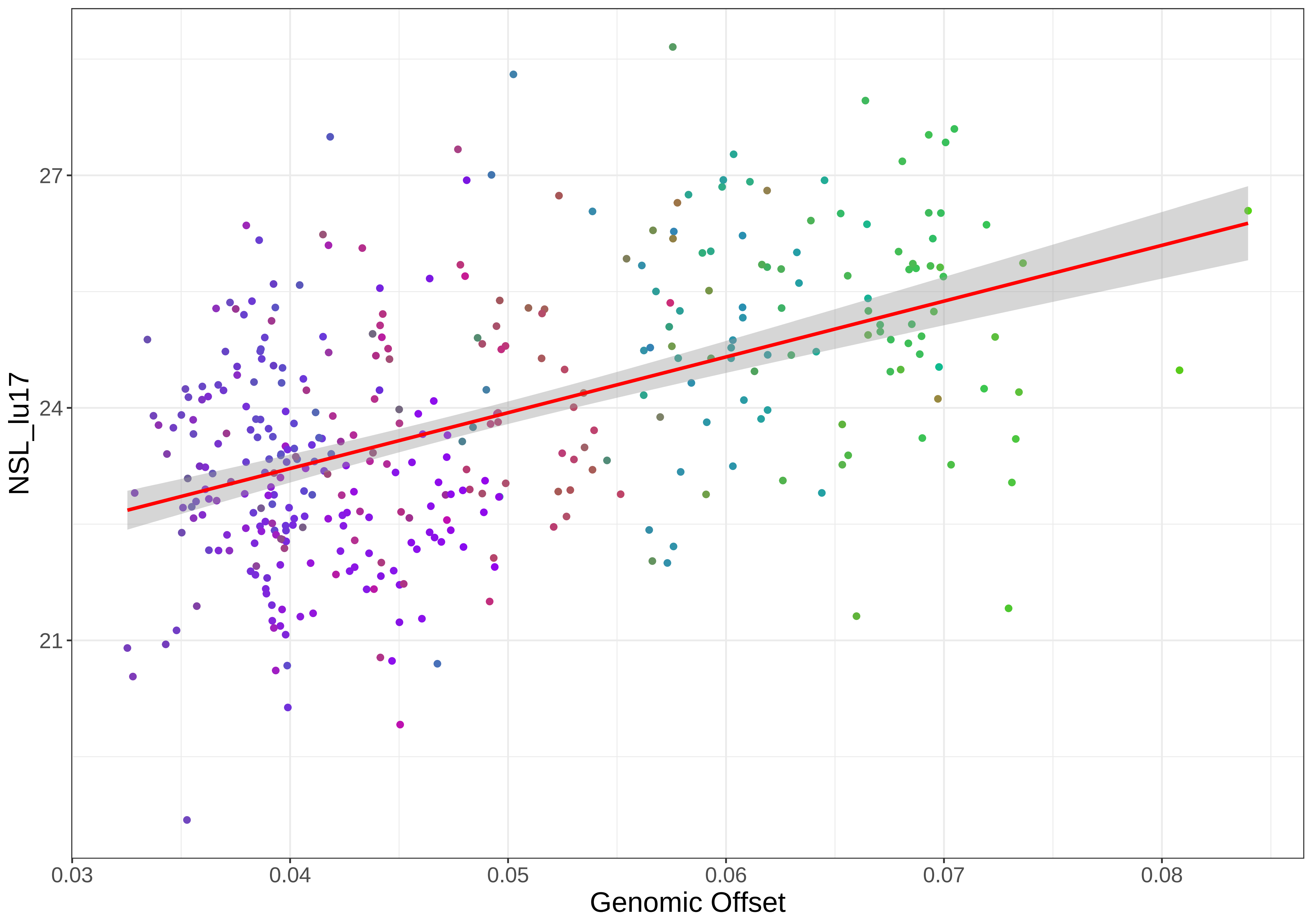
NSL_lu17
27
24
21
0.03
0.04
0.05
0.06
0.07
0.08
Genomic Offset

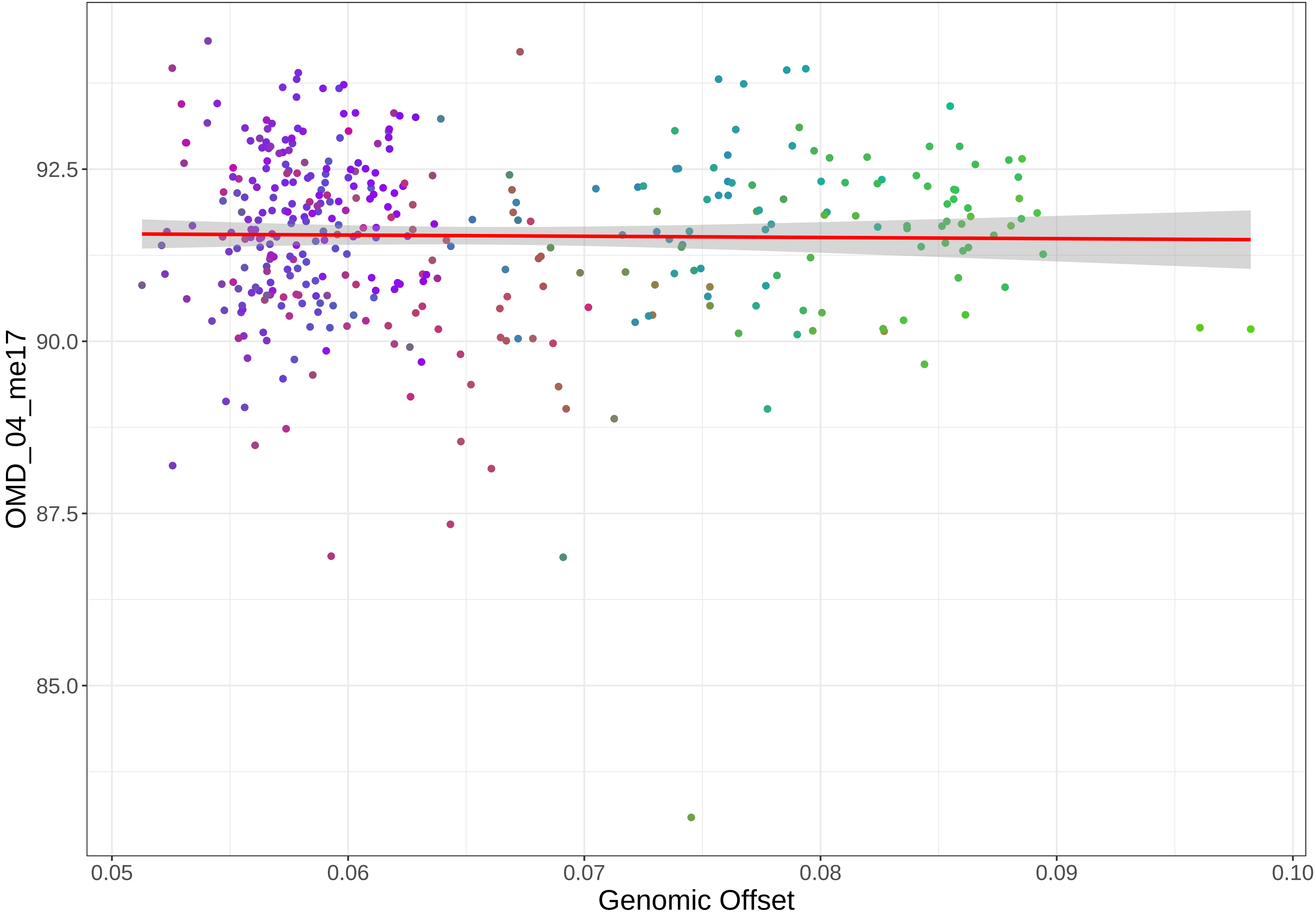

OMD_04_me17
92.5
90.0
87.5
85.0
0.05
0.06
0.07
0.08
0.09
0.10
Genomic Offset

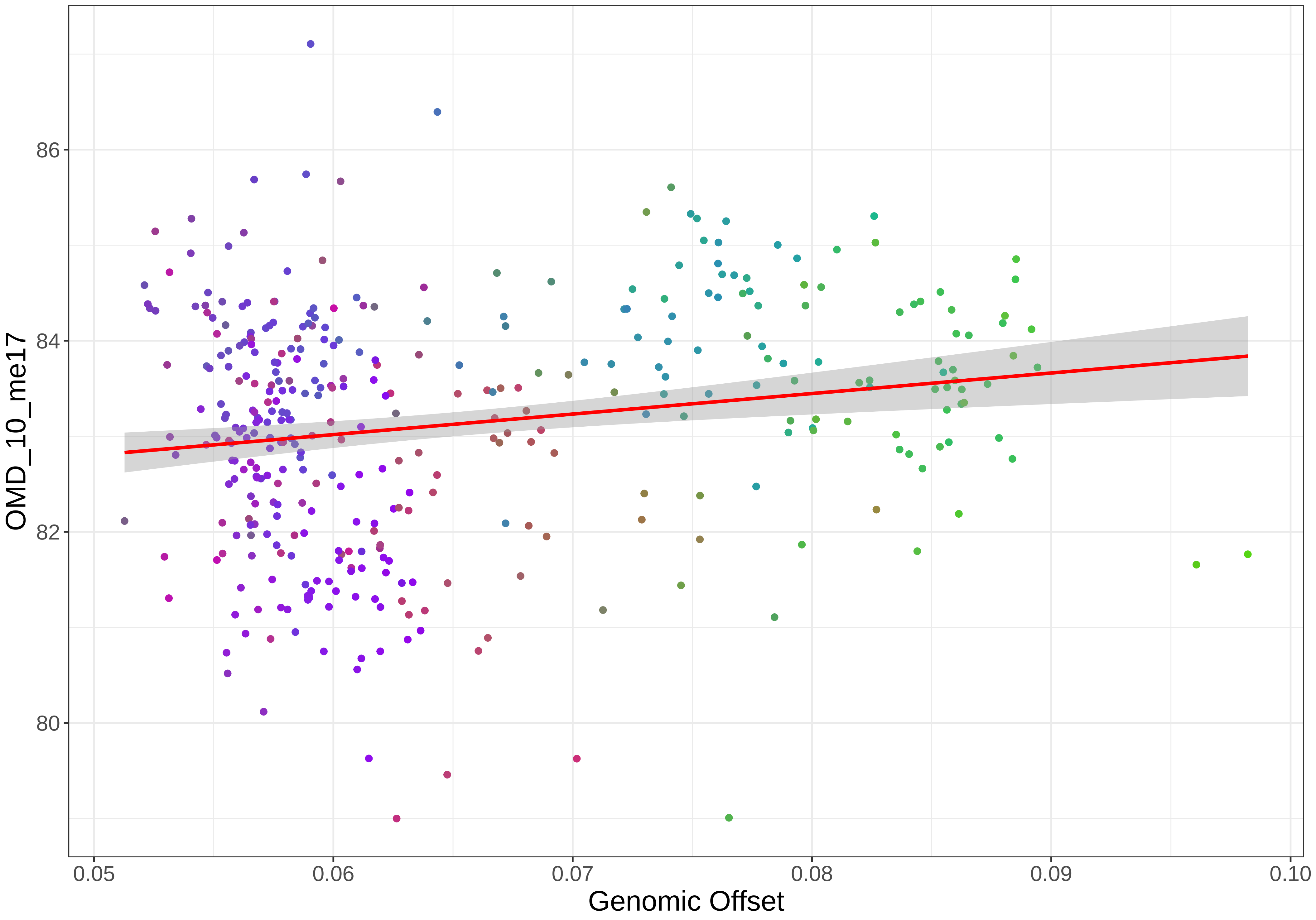

86
84
82
80
OMD_10_me17
0.05
0.06
0.07
0.08
0.09
0.10
Genomic Offset

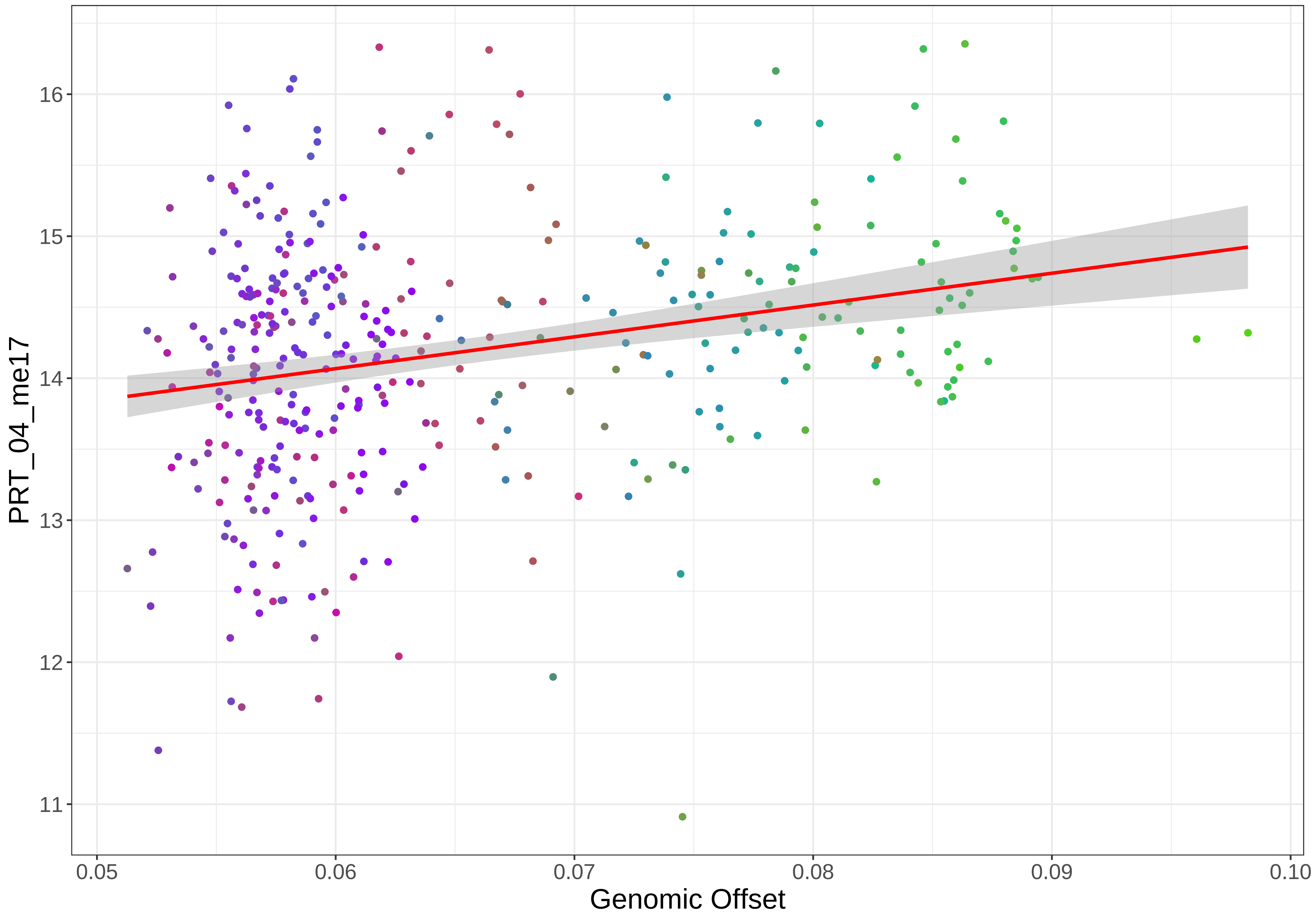

PRT_04_me17
16
15
14
13
12
11
0.05
0.06
0.07
0.08
0.09
0.10
Genomic Offset

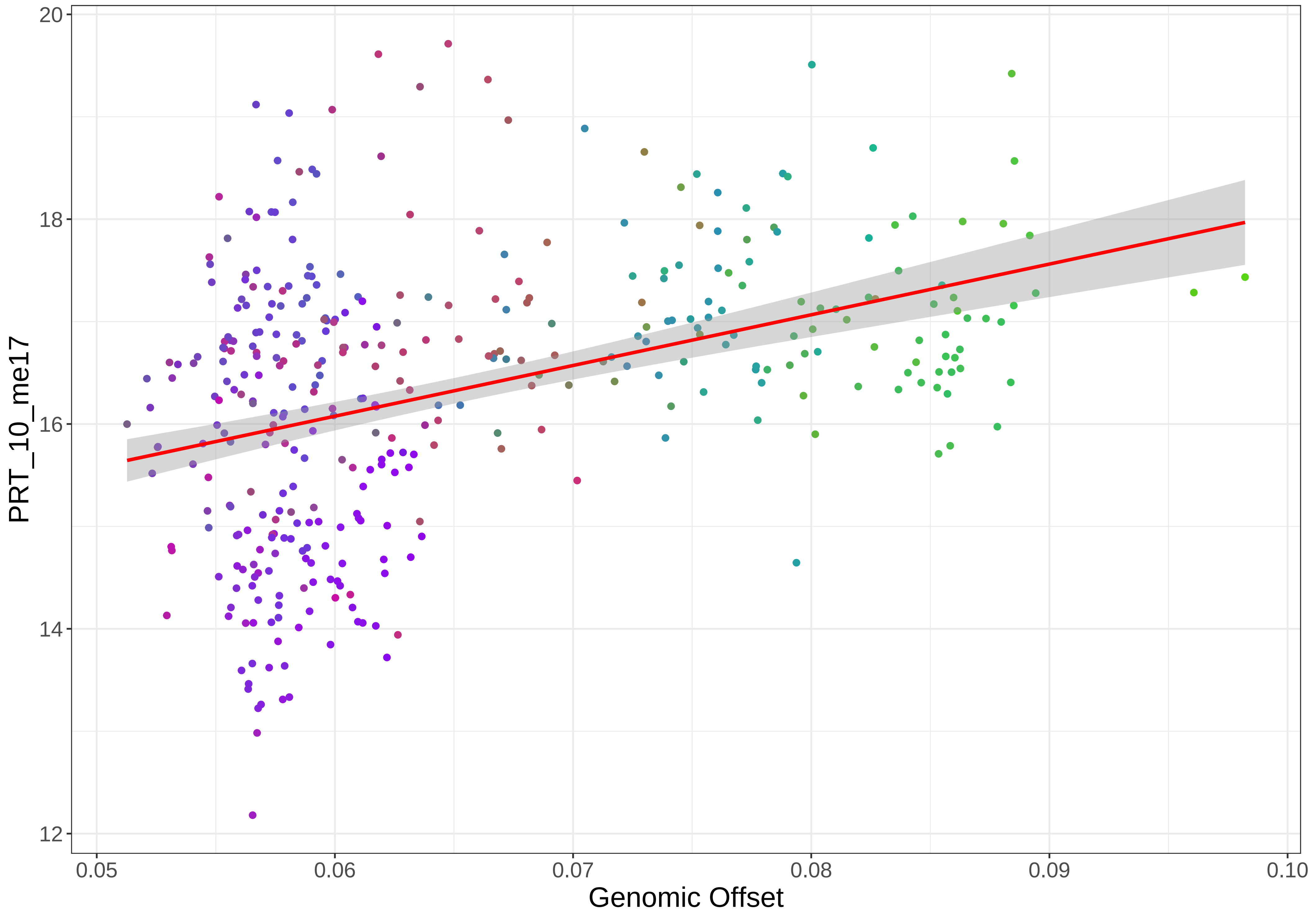

20
18
16
14
12
PRT_10_me17
0.05
0.06
0.07
0.08
0.09
0.10
Genomic Offset

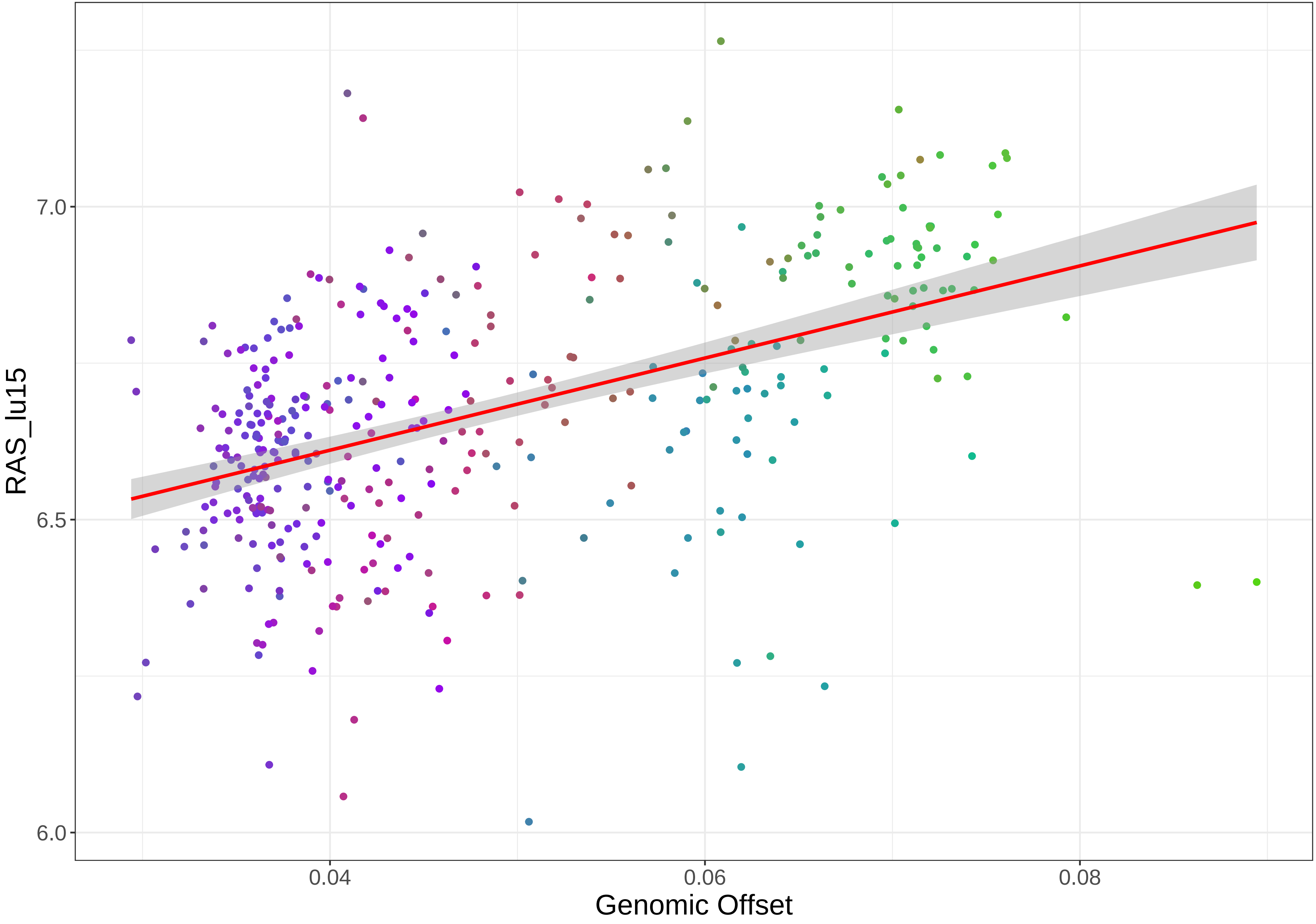
RAS_lu15
7.0
6.5
6.0
0.04
0.06
0.08
Genomic Offset

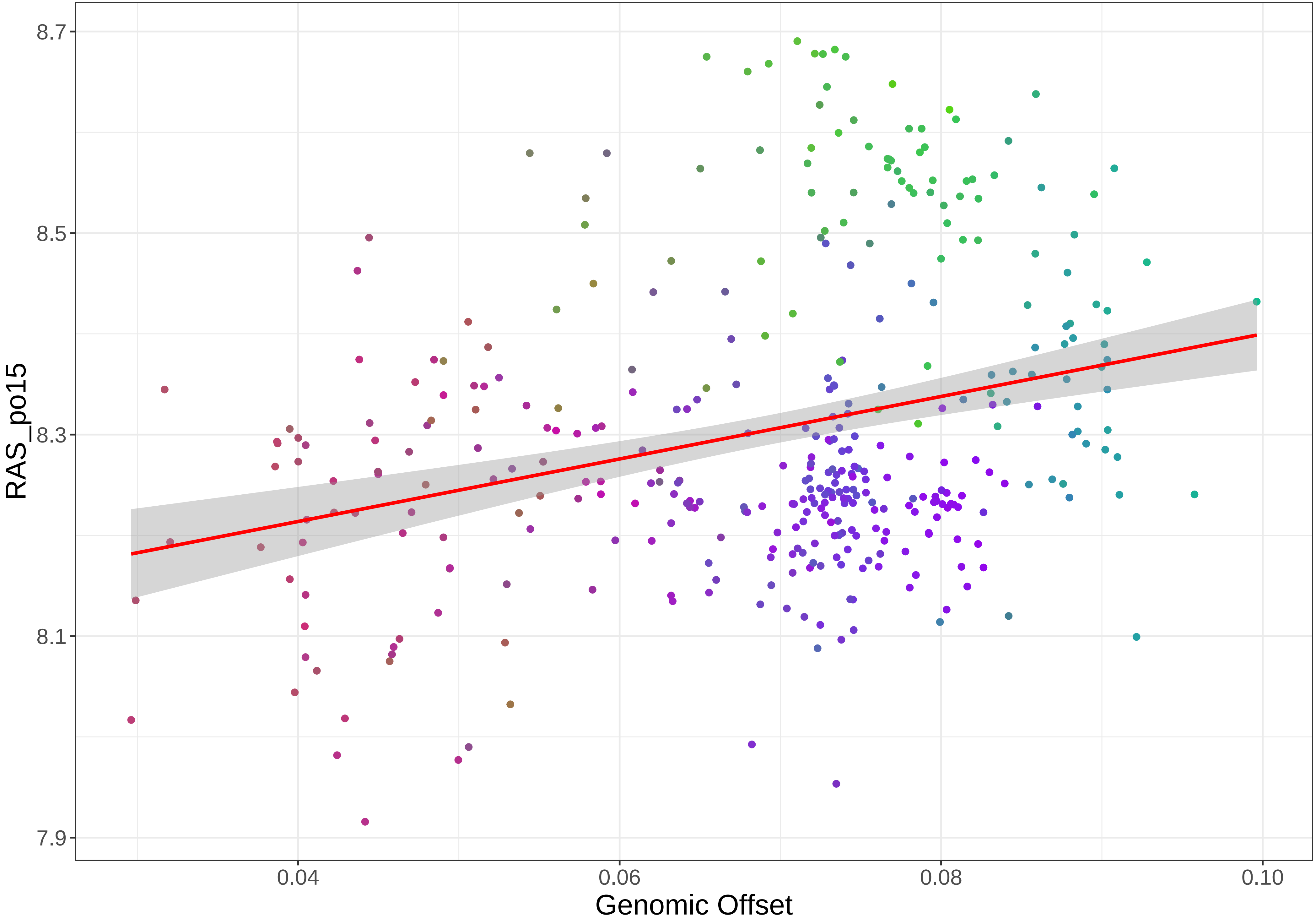

8.7
8.5
8.3
8.1
7.9
RAS_po15
0.04
0.06
0.08
0.10
Genomic Offset

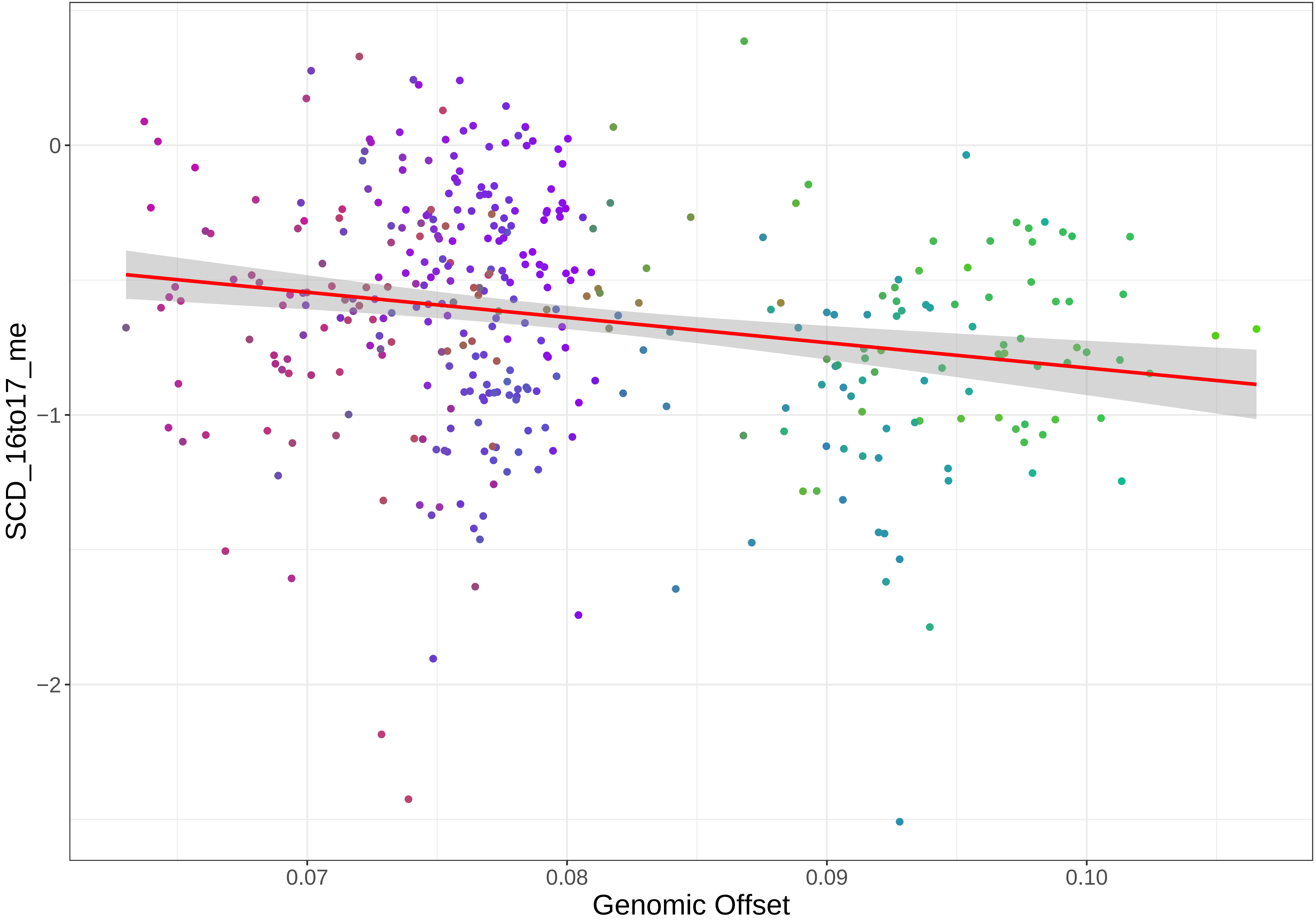

0
−1
−2
0.07
0.08
0.09
0.10
SCD_16to17_me
Genomic Offset

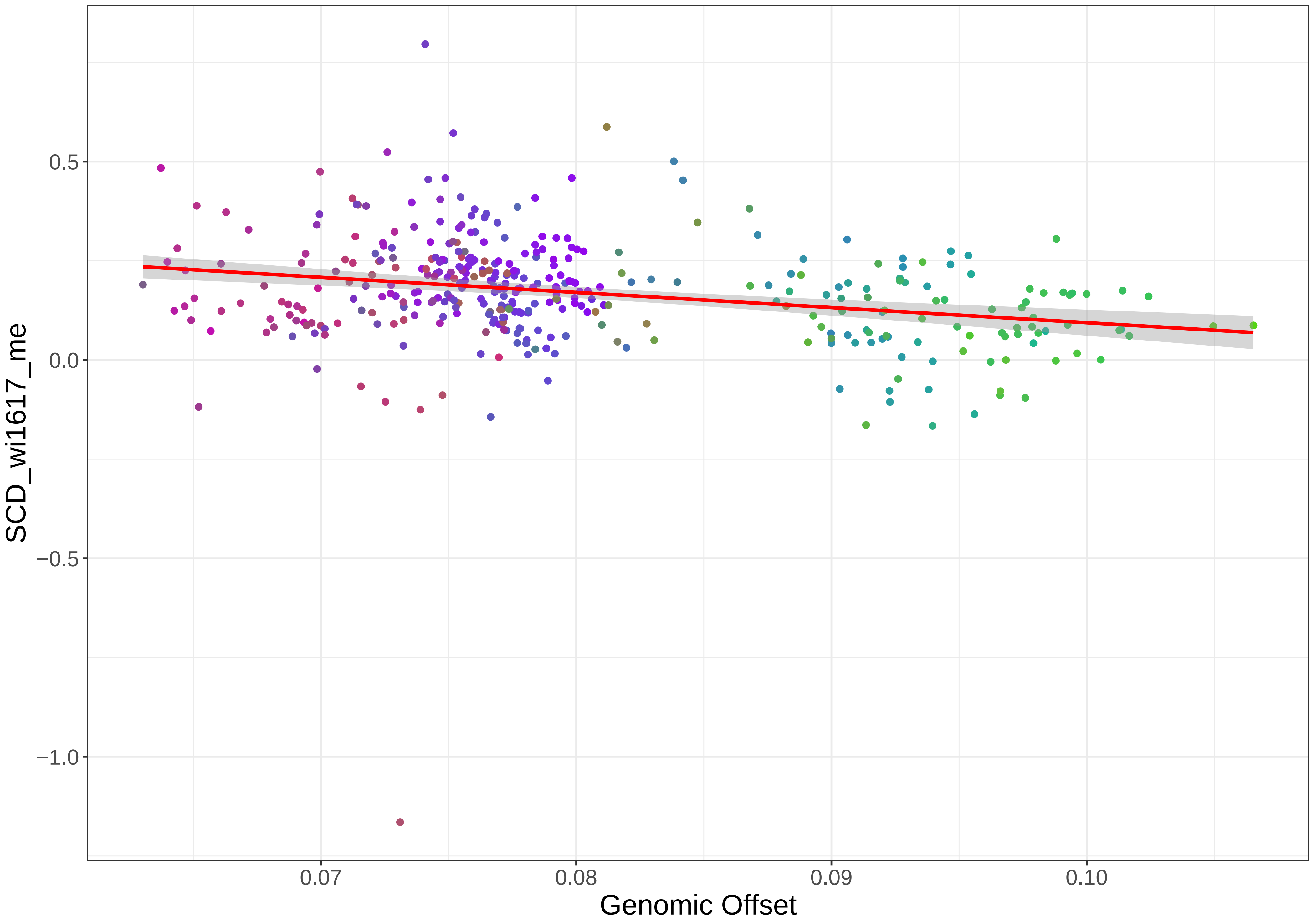
0.5
0.0
−0.5
−1.0
0.07
0.08
0.09
0.10
SCD_wi1617_me
Genomic Offset

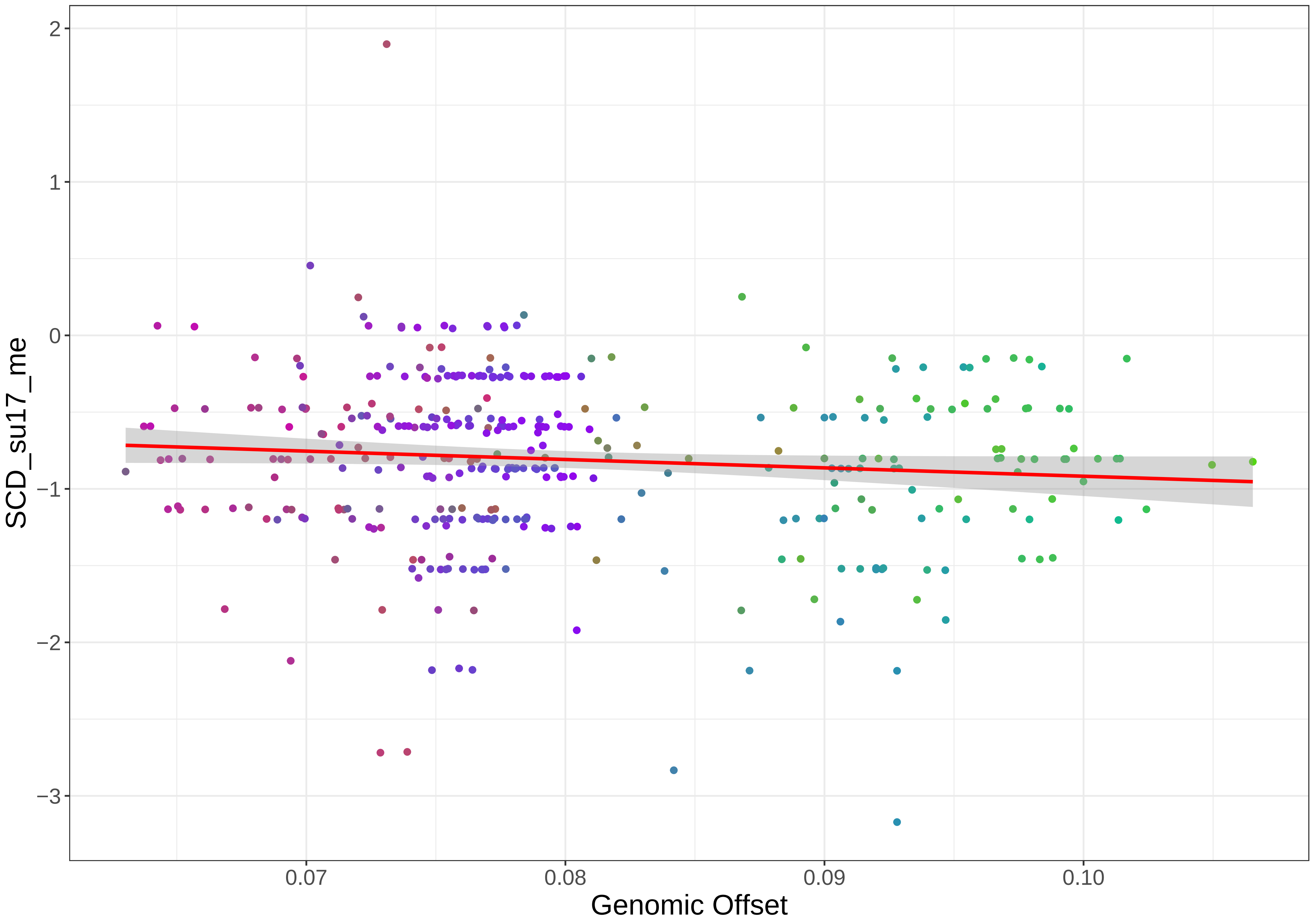

SCD_su17_me
2
1
0
−1
−2
−3
0.07
0.08
0.09
0.10
Genomic Offset

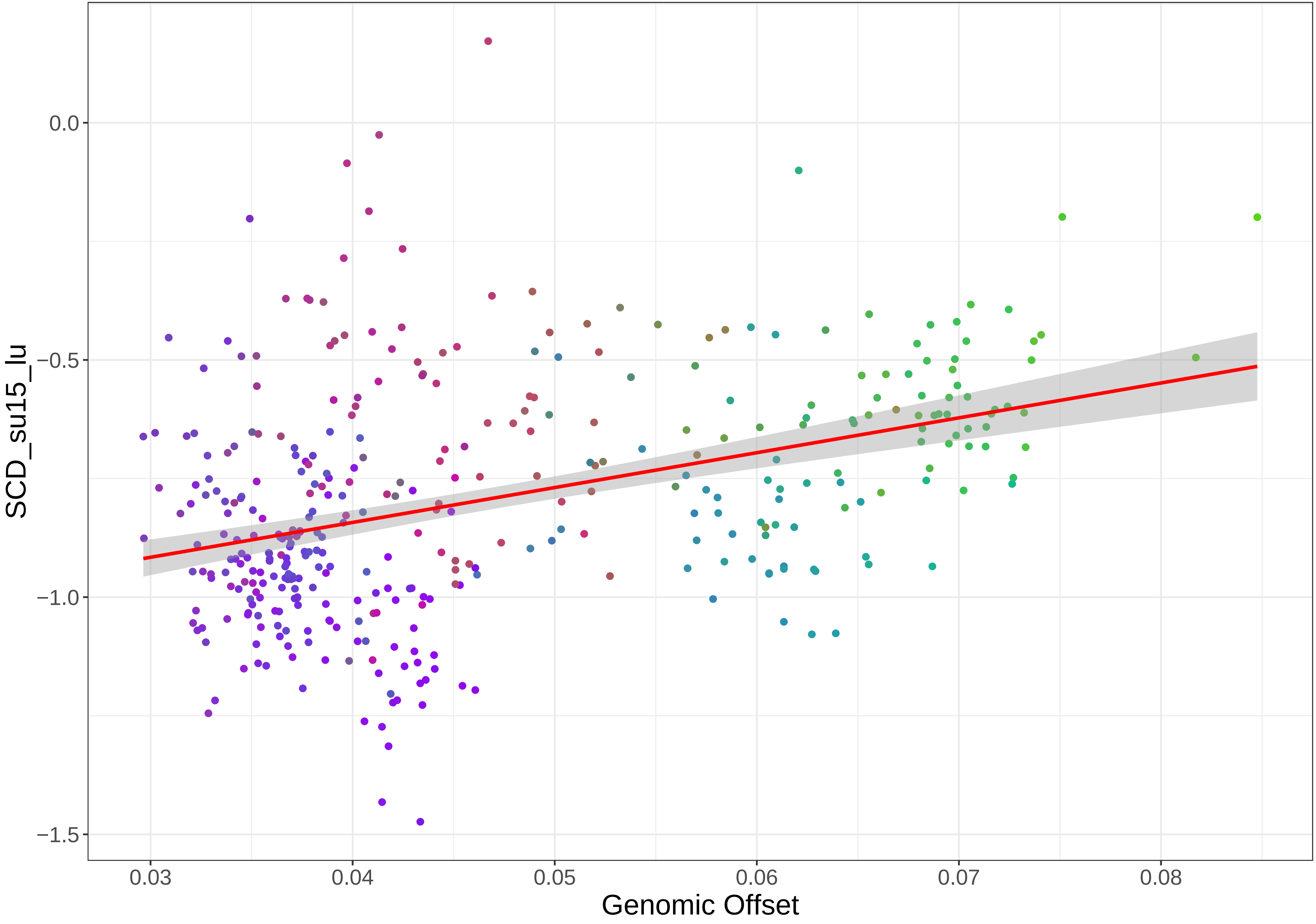

0.0
−0.5
−1.0
−1.5
0.03
0.04
0.05
0.06
0.07
0.08
SCD_su15_lu
Genomic Offset

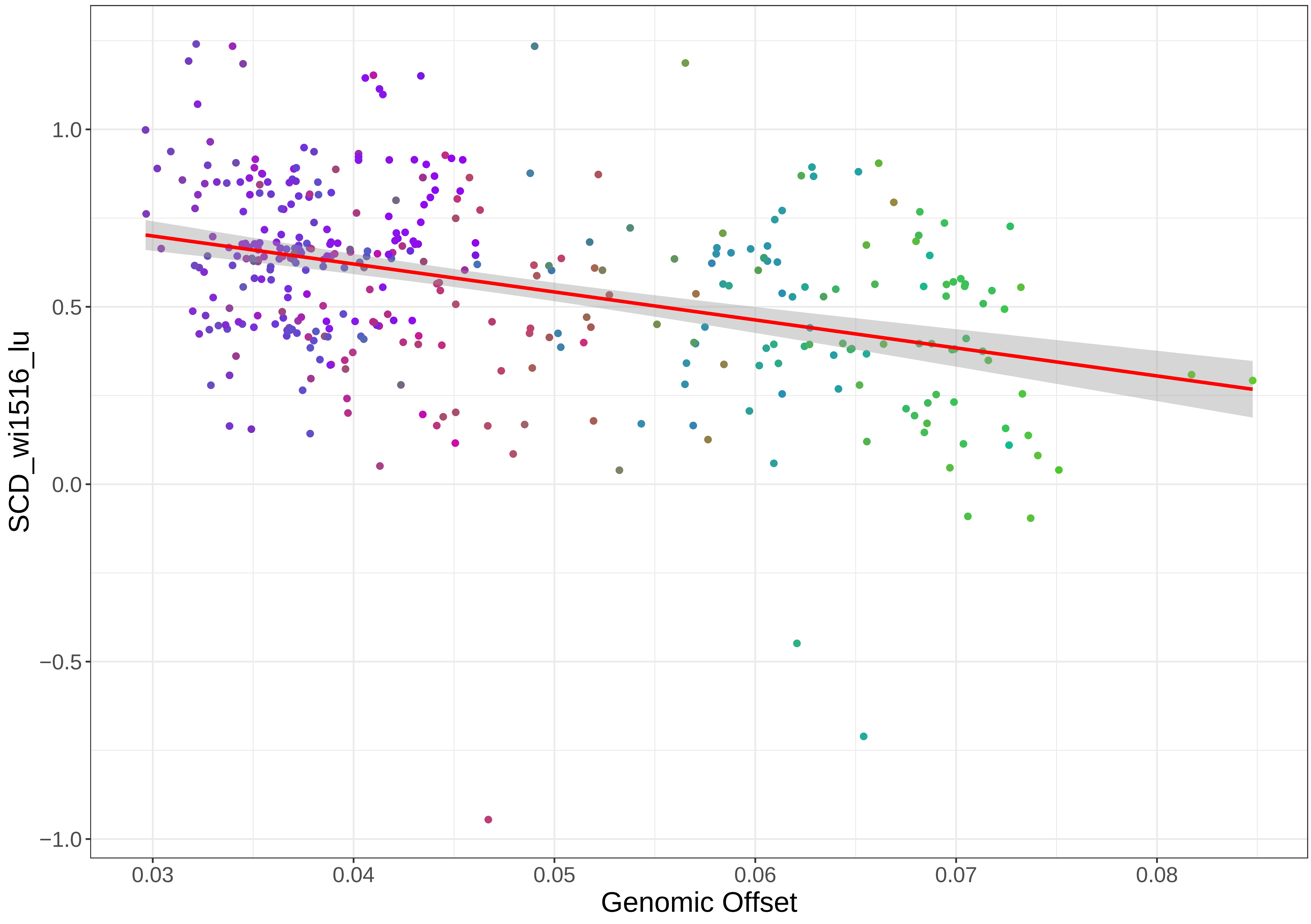

SCD_wi1516_lu
1.0
0.5
0.0
−0.5
−1.0
0.03
0.04
0.05
0.06
0.07
0.08
Genomic Offset

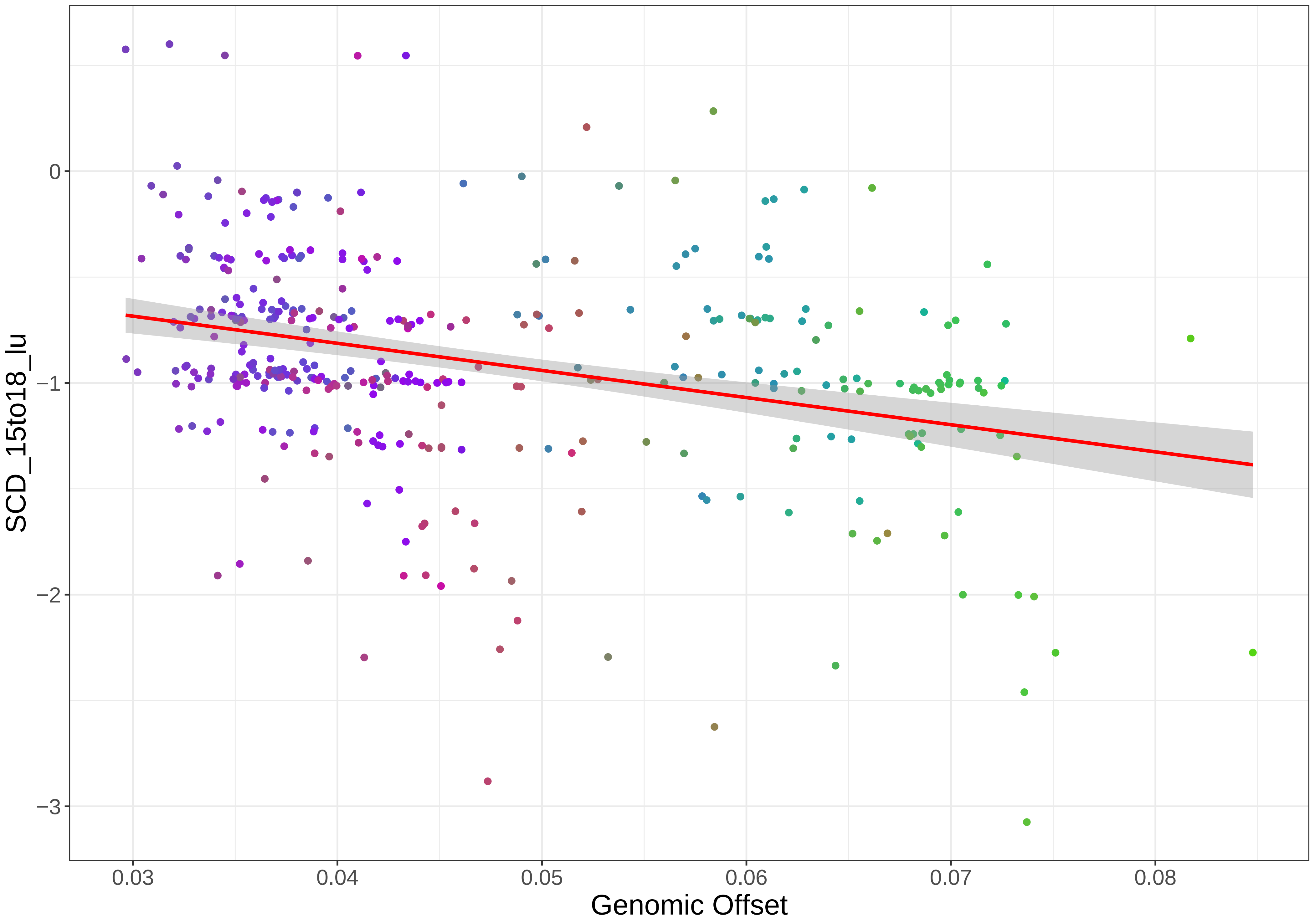

0
−1
−2
−3
0.03
0.04
0.05
0.06
0.07
0.08
Genomic Offset
SCD_15to18_lu

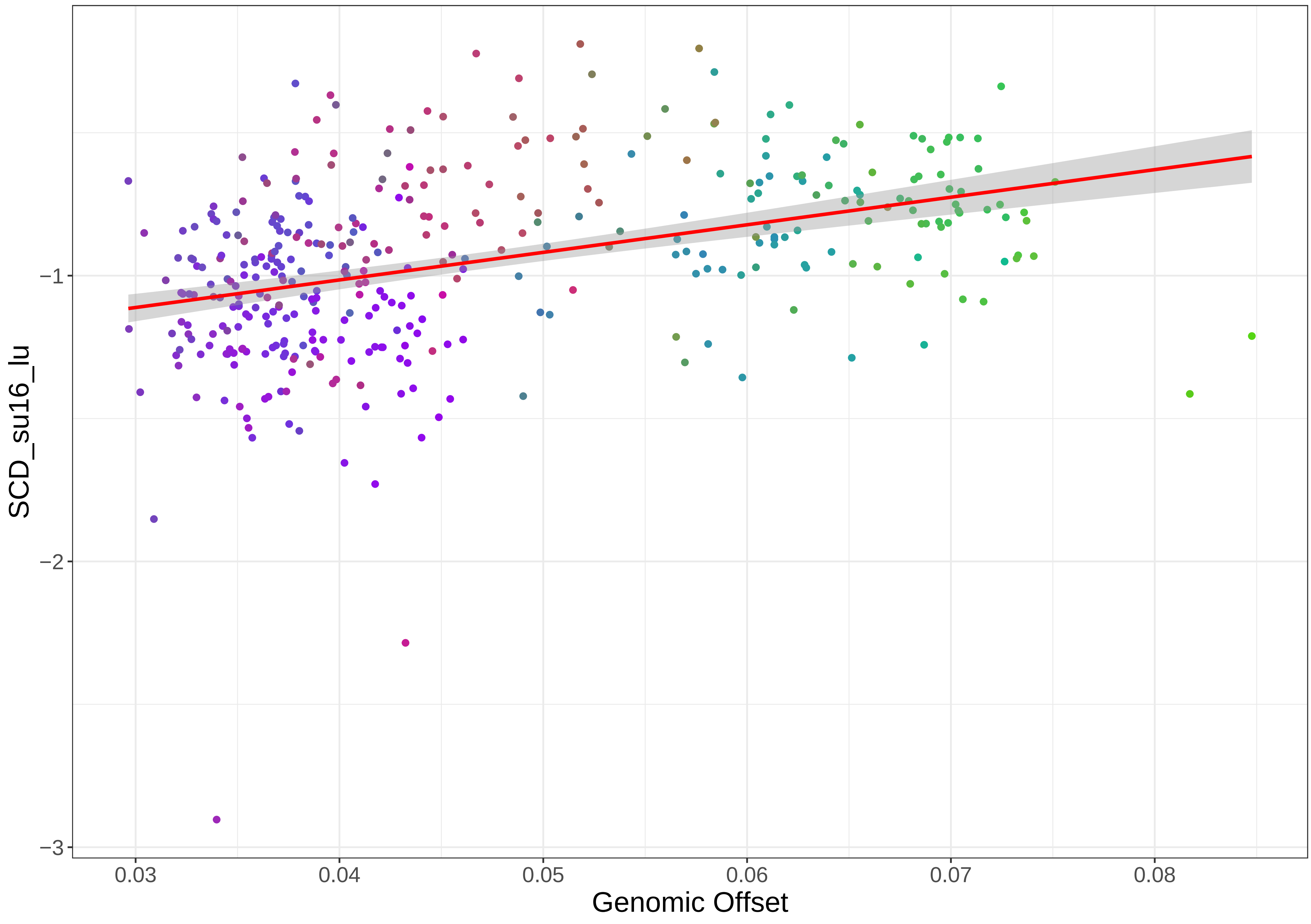

SCD_su16_lu
Genomic Offset
−1
−2
−3
0.03
0.04
0.05
0.06
0.07
0.08

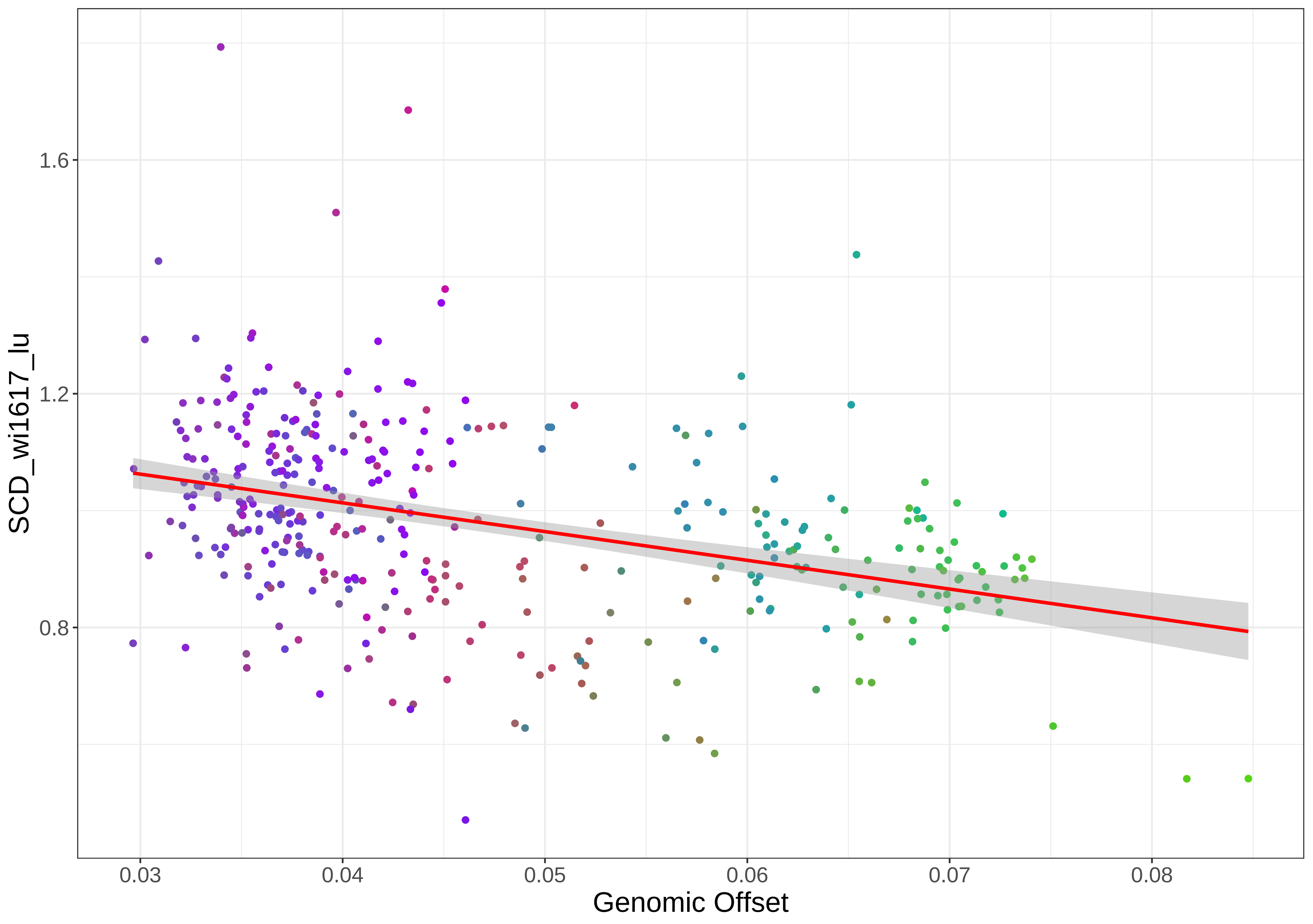

1.6
1.2
0.8
SCD_wi1617_lu
0.03
0.04
0.05
0.06
0.07
0.08
Genomic Offset

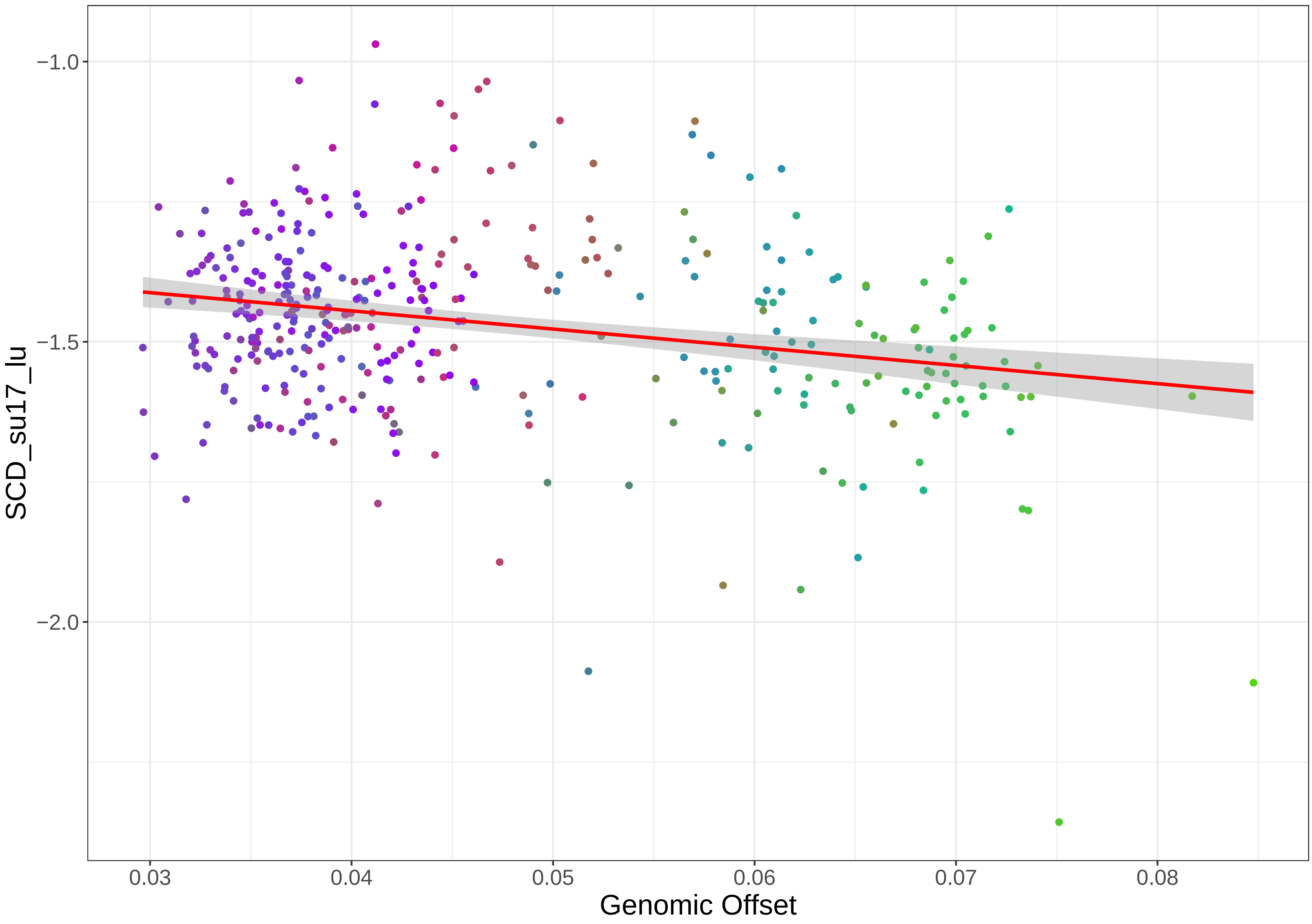

−1.0
−1.5
−2.0
SCD_su17_lu
0.03
0.04
0.05
0.06
0.07
0.08
Genomic Offset

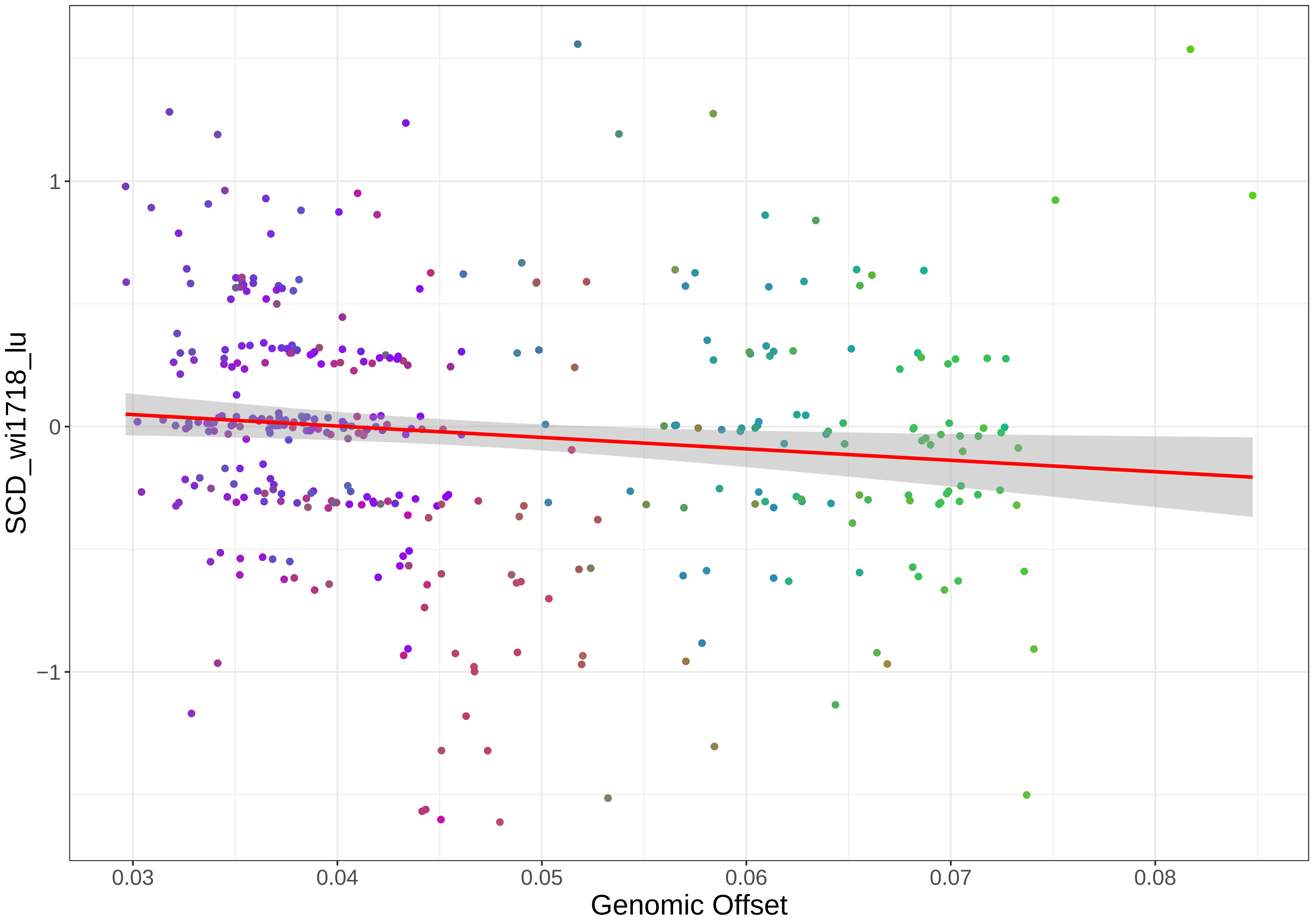

SCD_wi1718_lu
1
0
−1
0.03
0.04
0.05
0.06
0.07
0.08
Genomic Offset

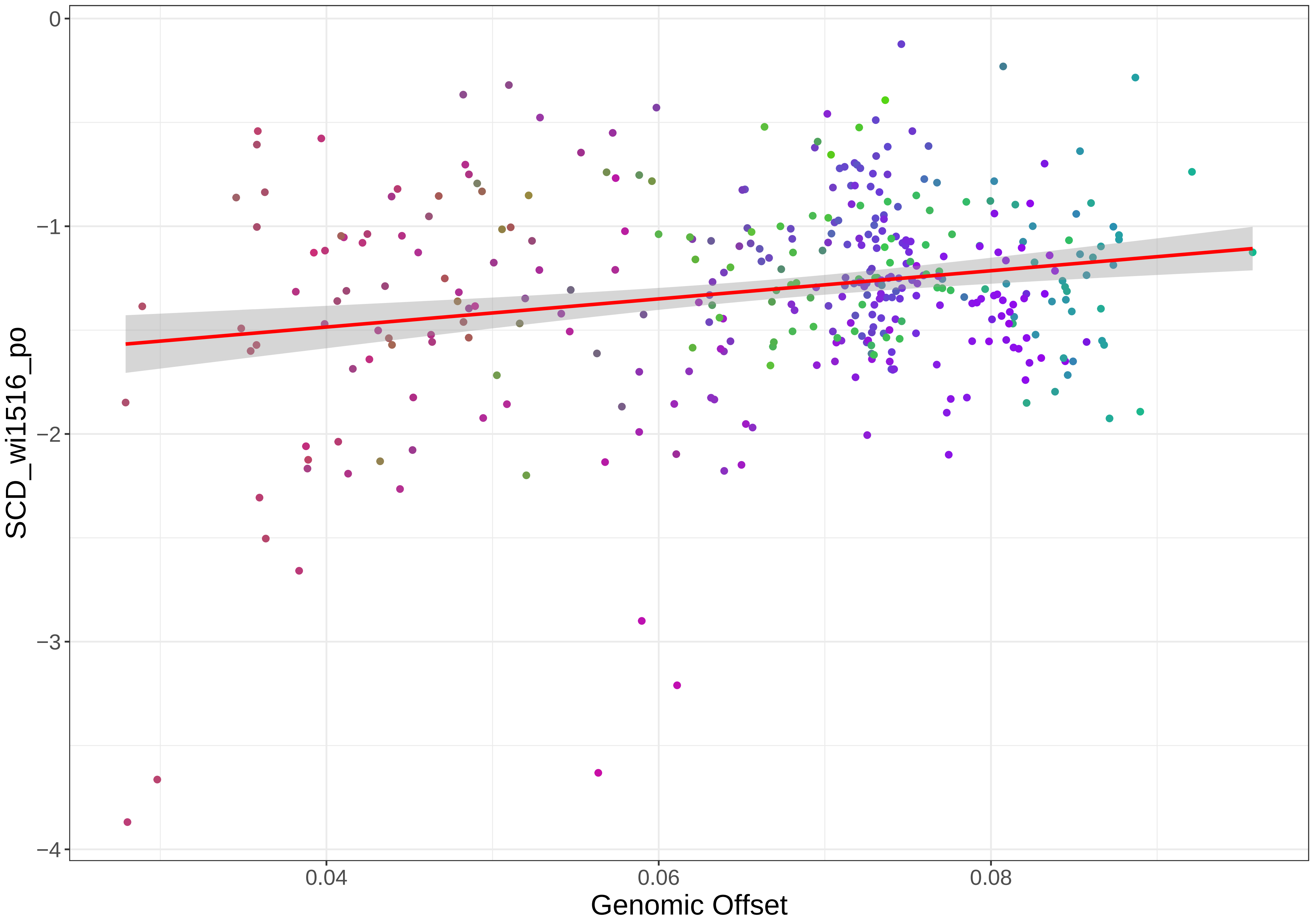

0
−1
−2
−3
−4
SCD_wi1516_po
0.04
0.06
0.08
Genomic Offset

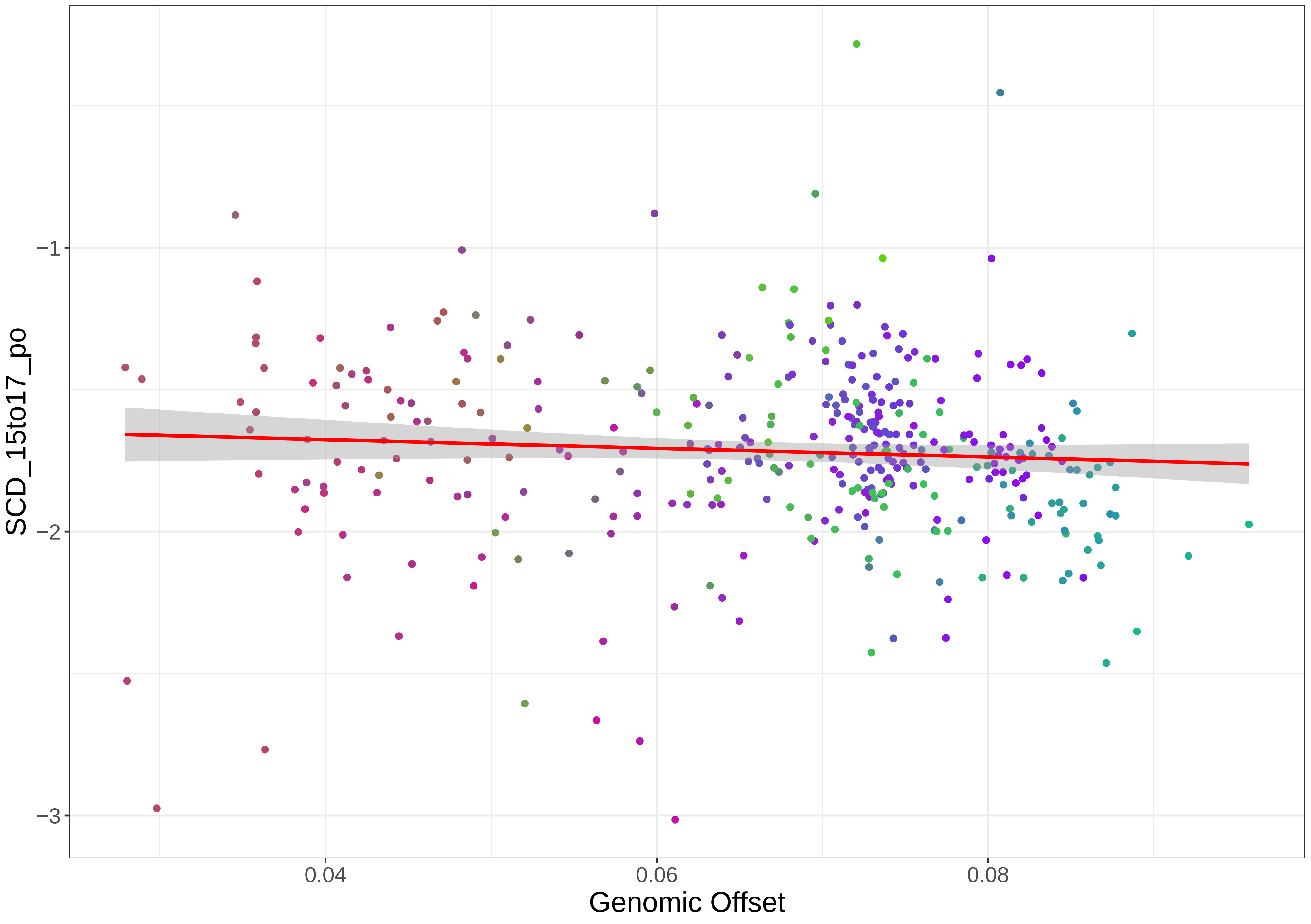
SCD_15to17_po
−1
−2
−3
0.04
0.06
0.08
Genomic Offset

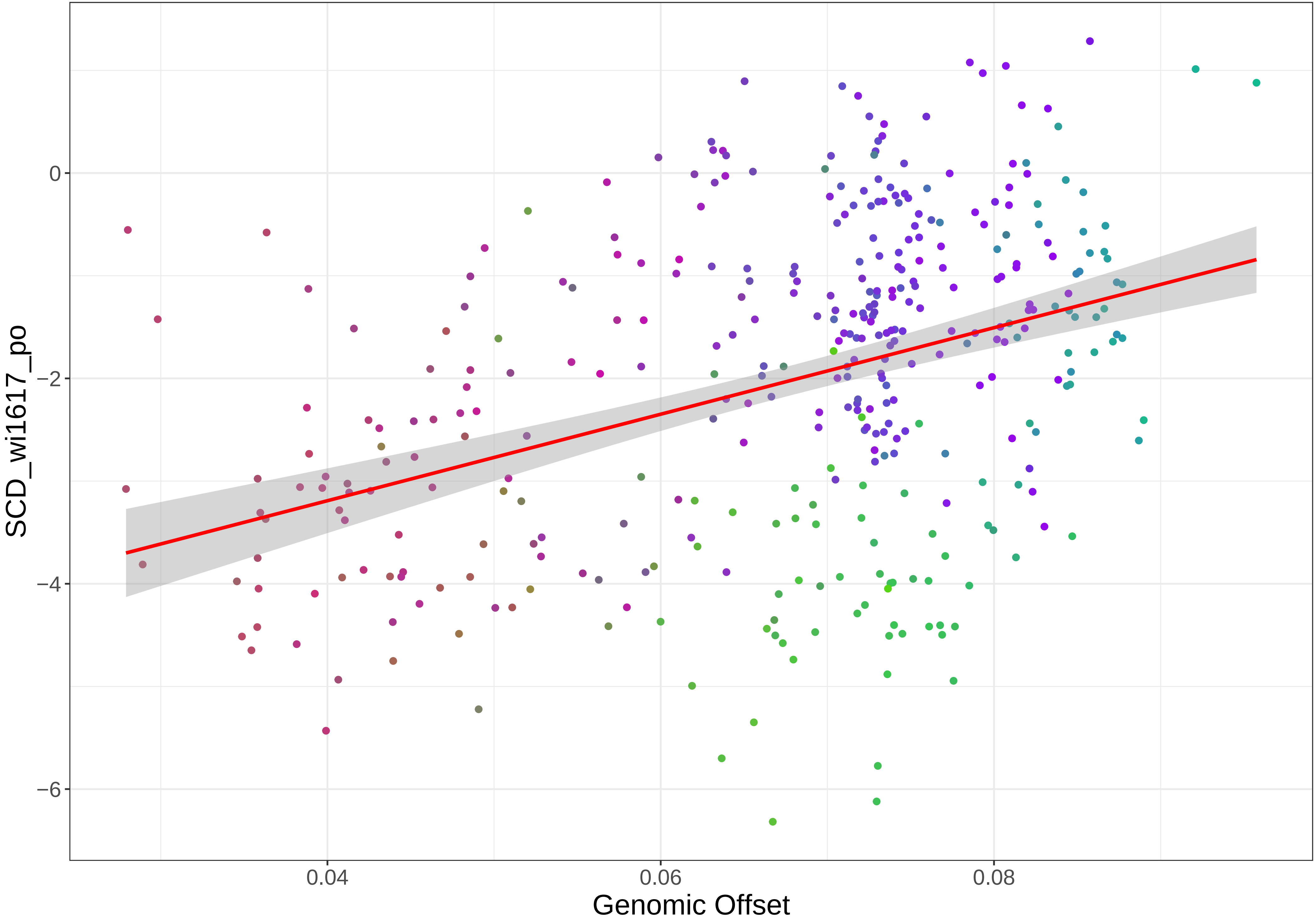

0
−2
−4
−6
0.04
0.06
0.08
Genomic Offset
SCD_wi1617_po

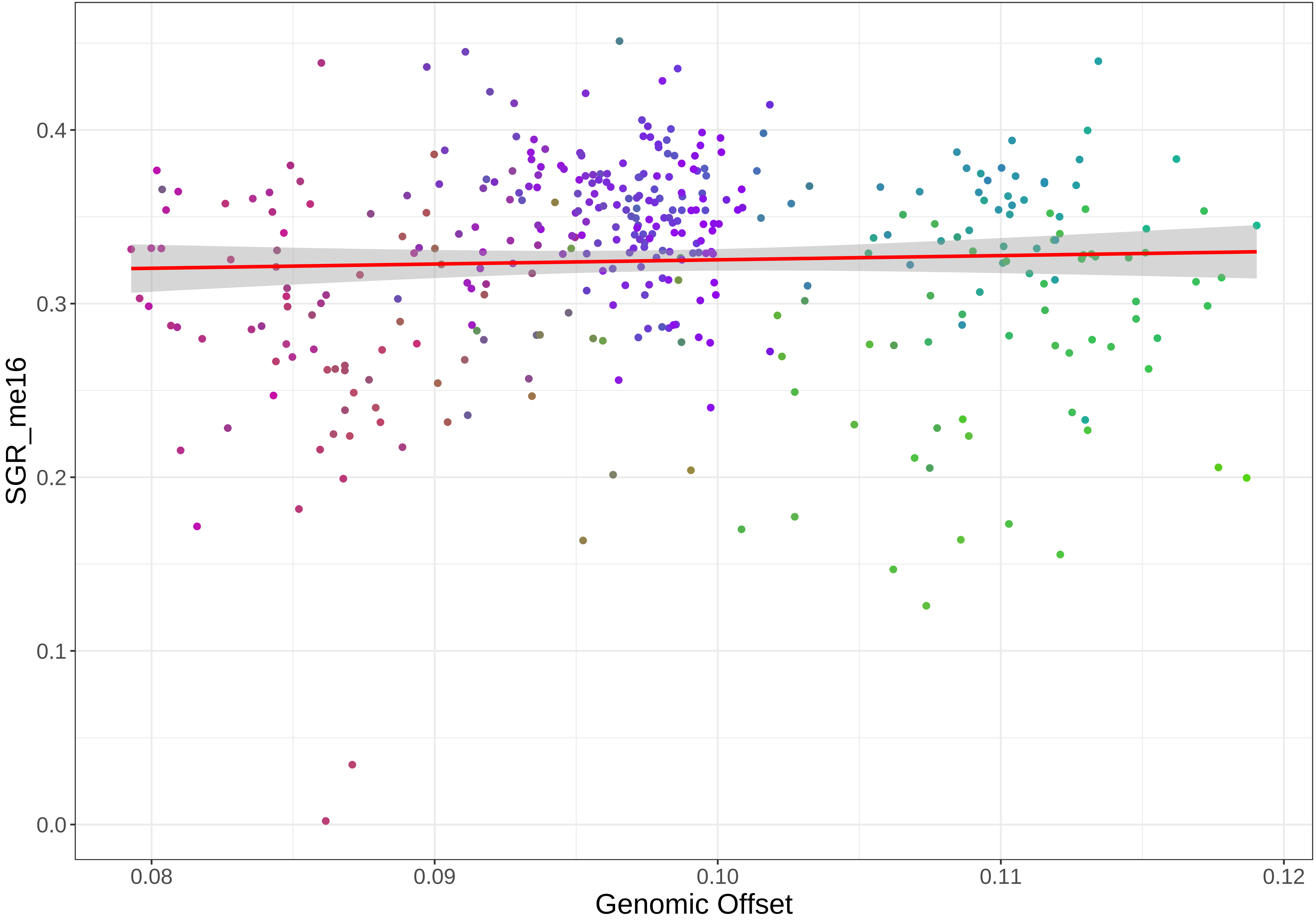

SGR_me16
0.0
0.1
0.2
0.3
0.4
0.08
0.09
0.10
0.11
0.12
Genomic Offset

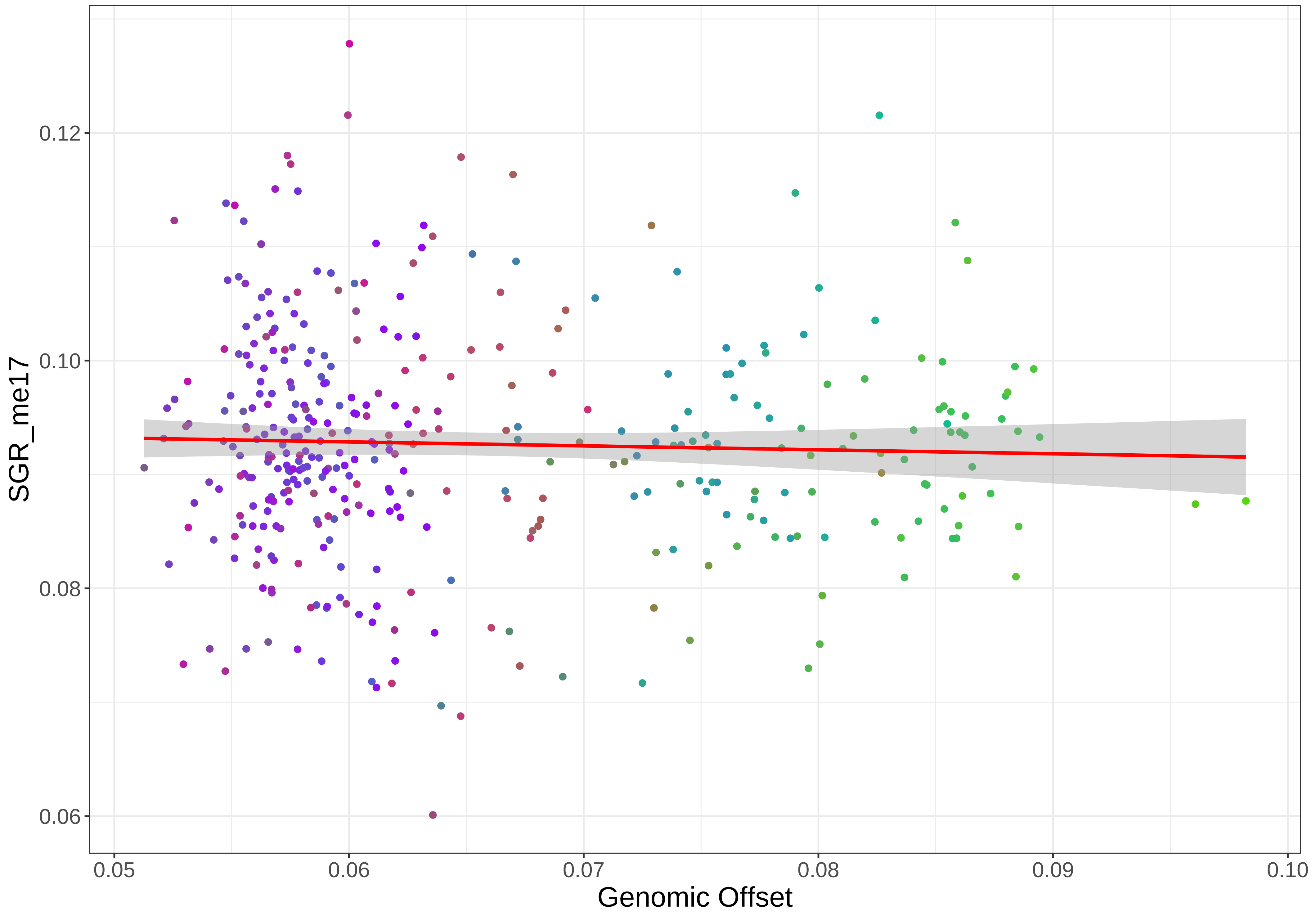

SGR_me17
0.06
0.08
0.10
0.12
0.05
0.06
0.07
0.08
0.09
0.10
Genomic Offset

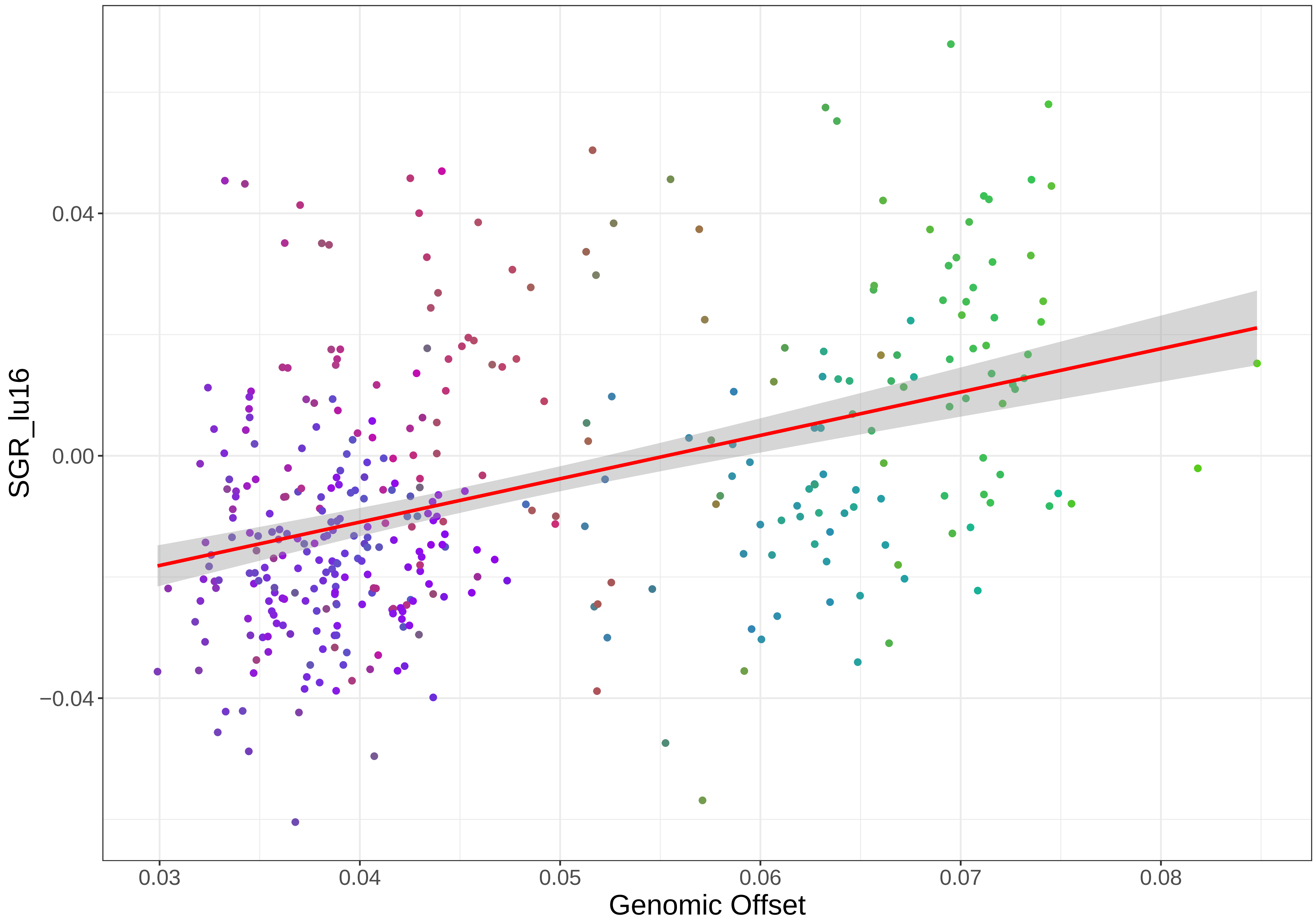

SGR_lu16
0.04
0.00
−0.04
0.03
0.04
0.05
0.06
0.07
0.08
Genomic Offset

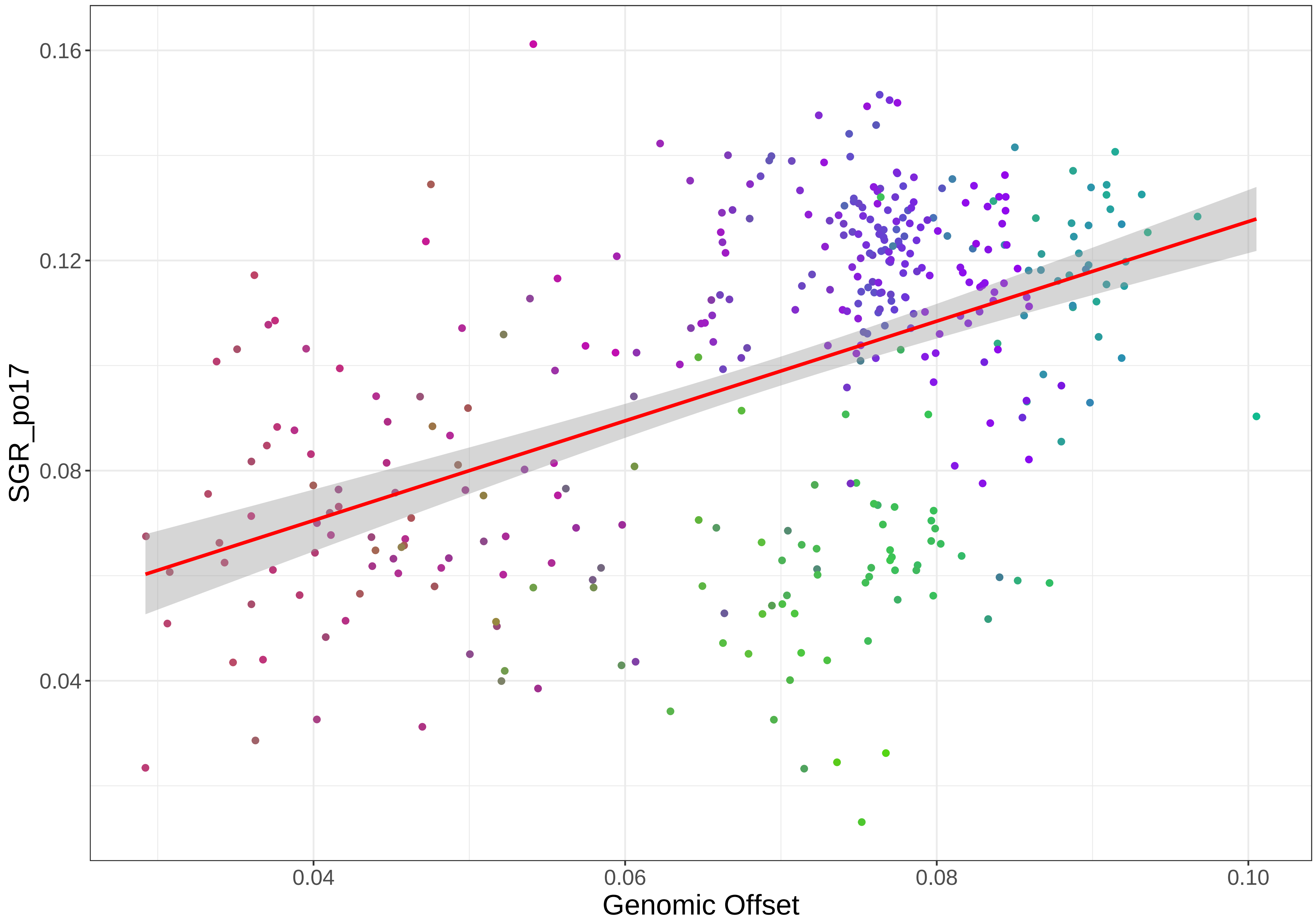

SGR_po17
0.16
0.12
0.08
0.04
0.04
0.06
0.08
0.10
Genomic Offset

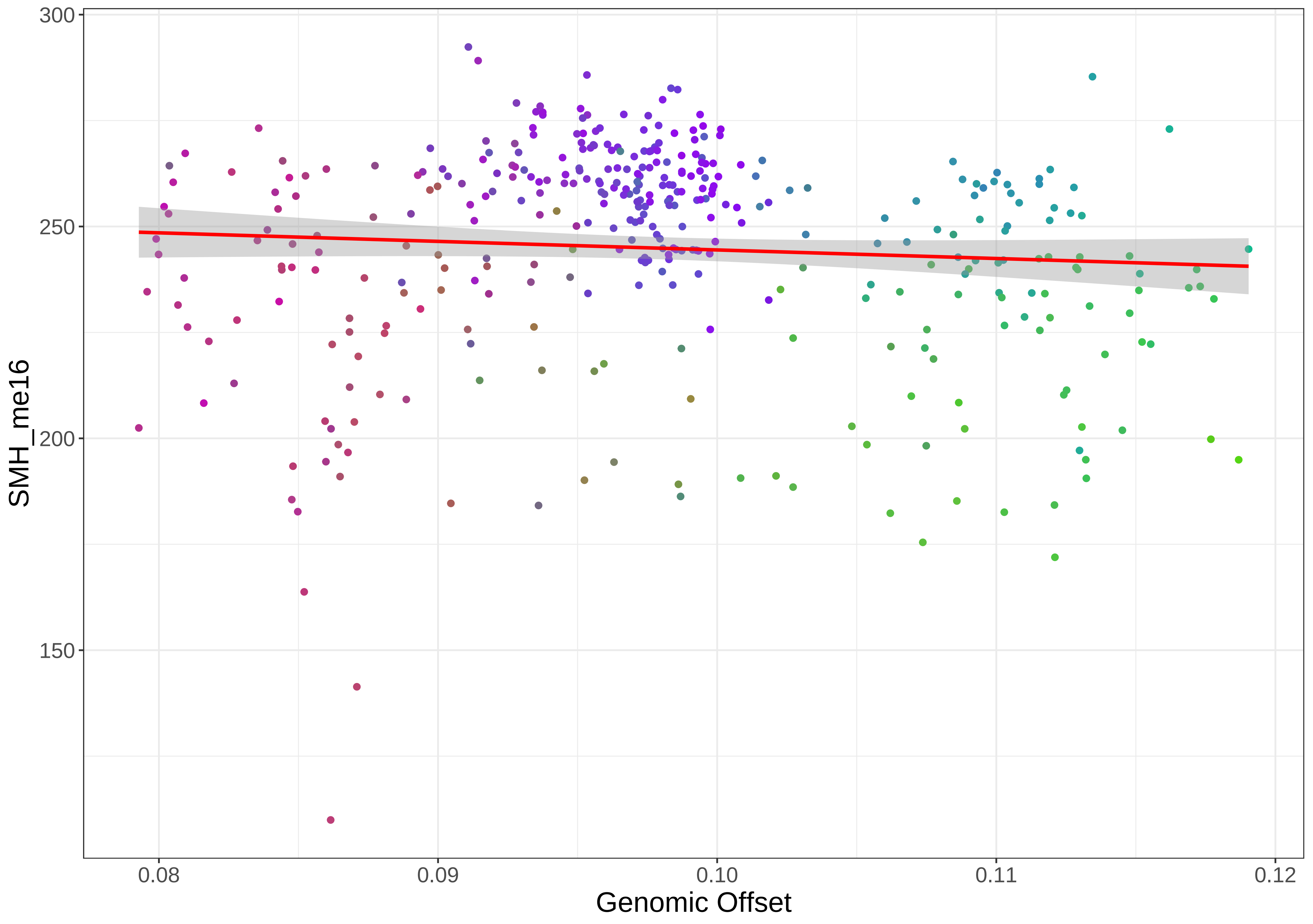

300
250
200
150
SMH_me16
0.08
0.09
0.10
0.11
0.12
Genomic Offset

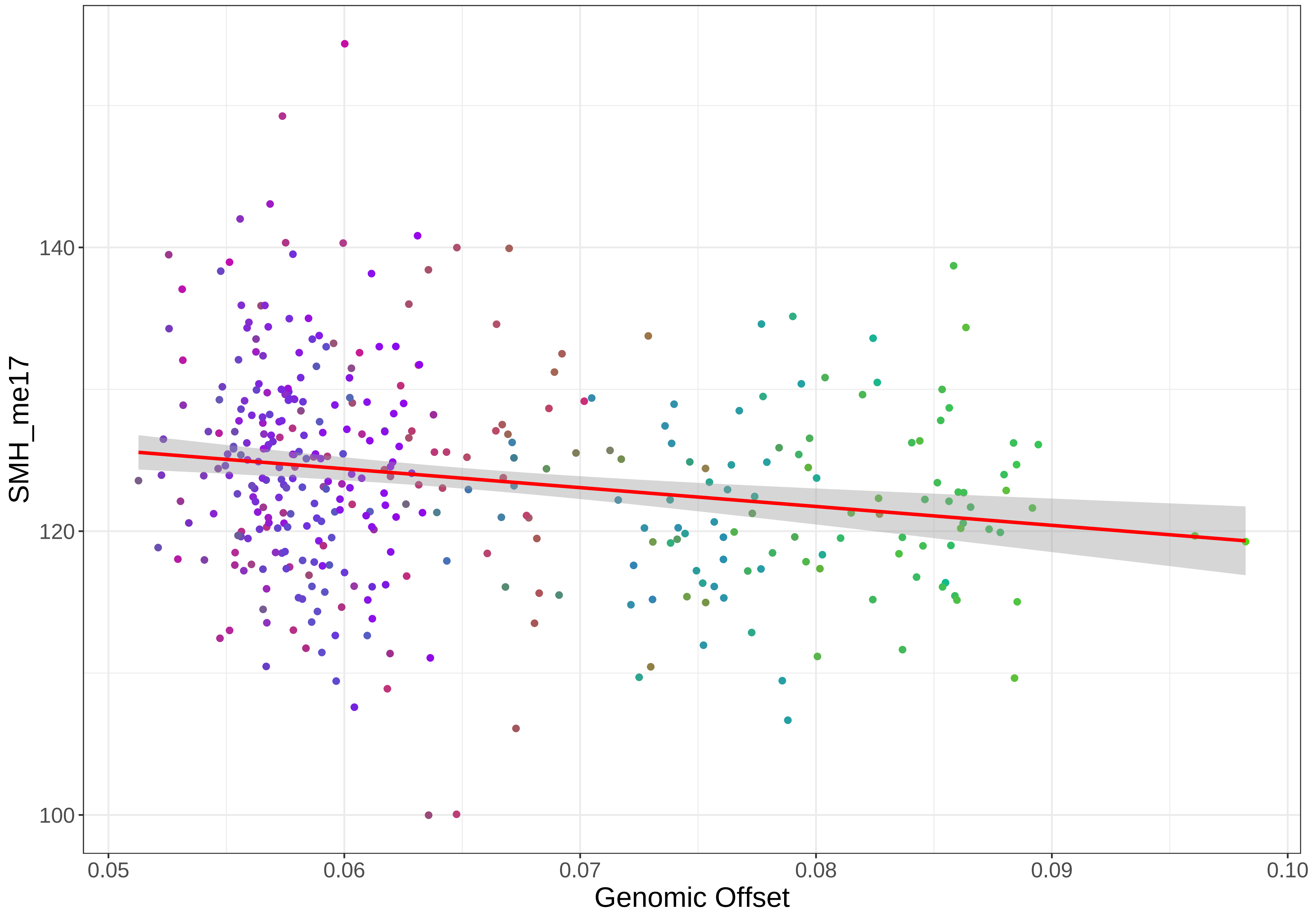
SMH_me17
140
120
100
0.05
0.06
0.07
0.08
0.09
0.10
Genomic Offset

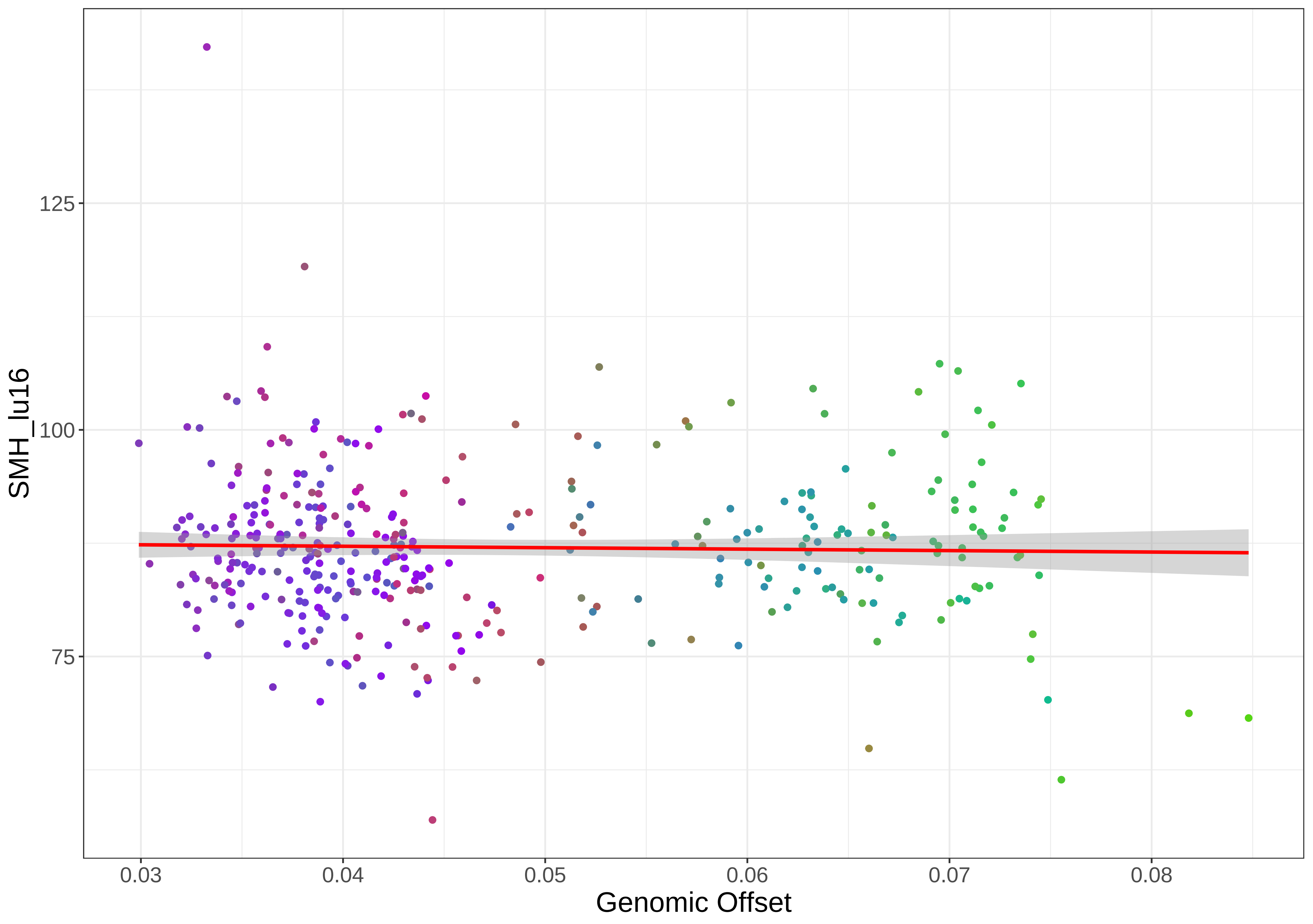

SMH_lu16
125
100
75
0.03
0.04
0.05
0.06
0.07
0.08
Genomic Offset

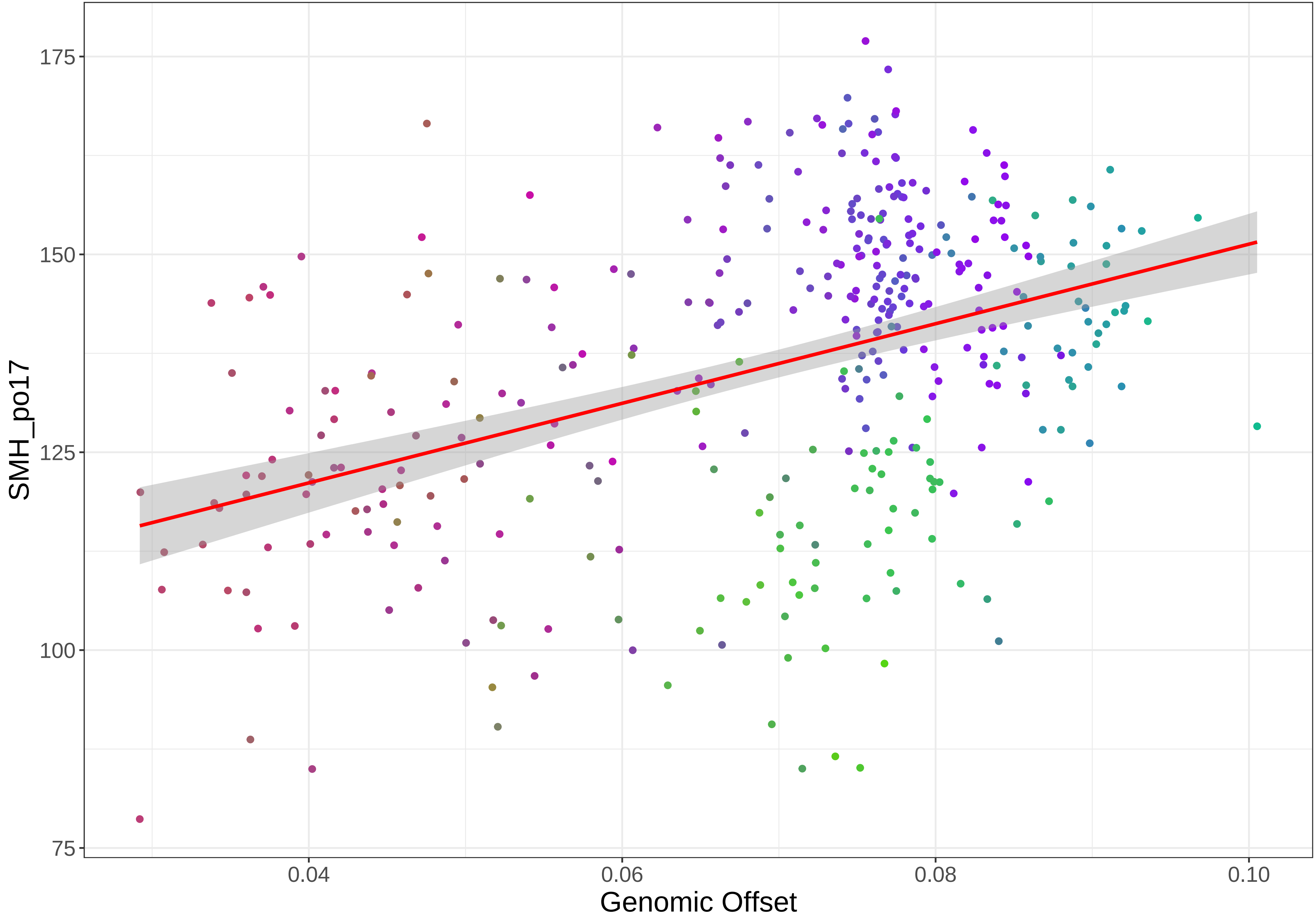

SMH_po17
175
150
125
100
75
0.04
0.06
0.08
0.10
Genomic Offset

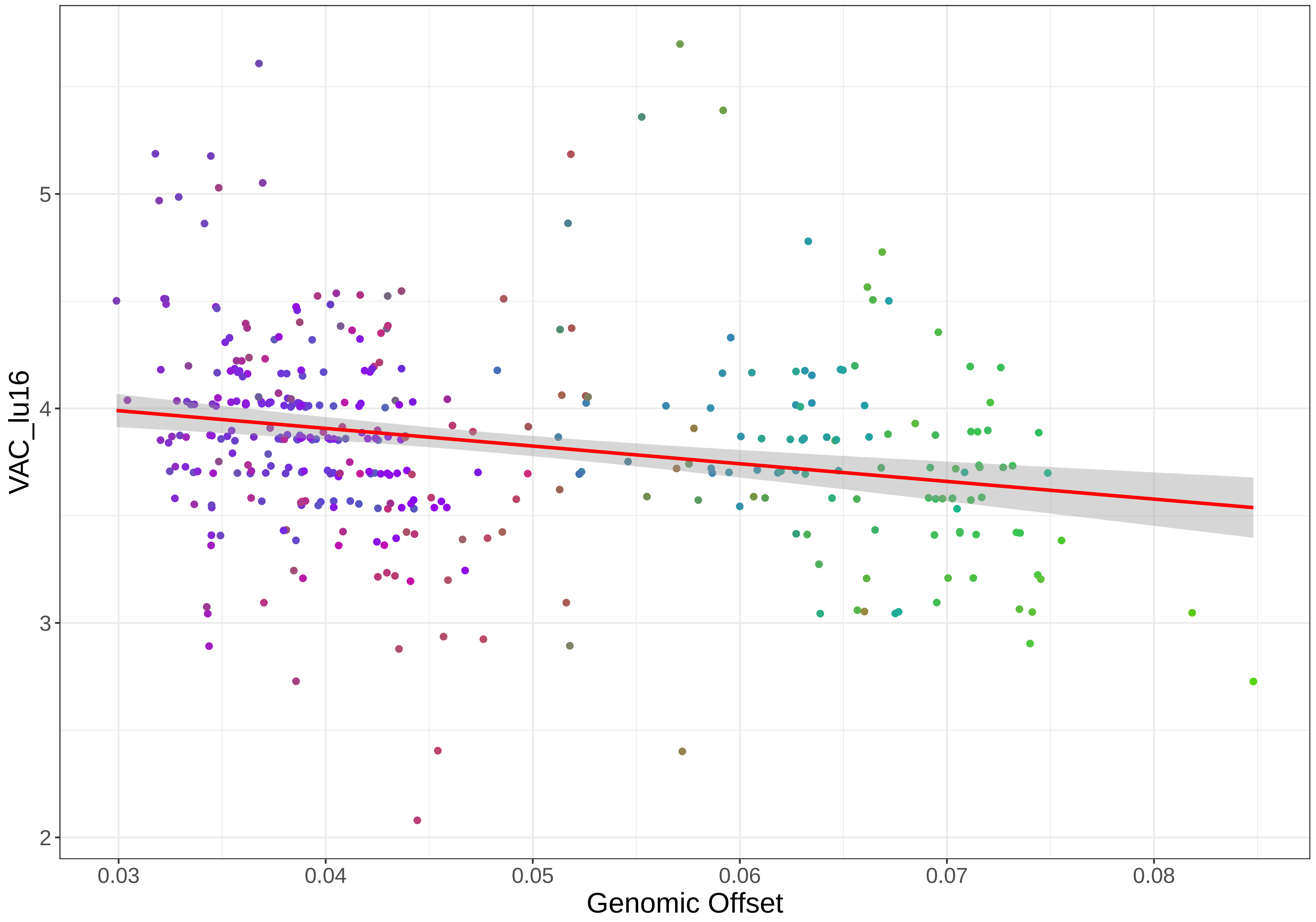

VAC_lu16
5
4
3
2
0.03
0.04
0.05
0.06
0.07
0.08
Genomic Offset

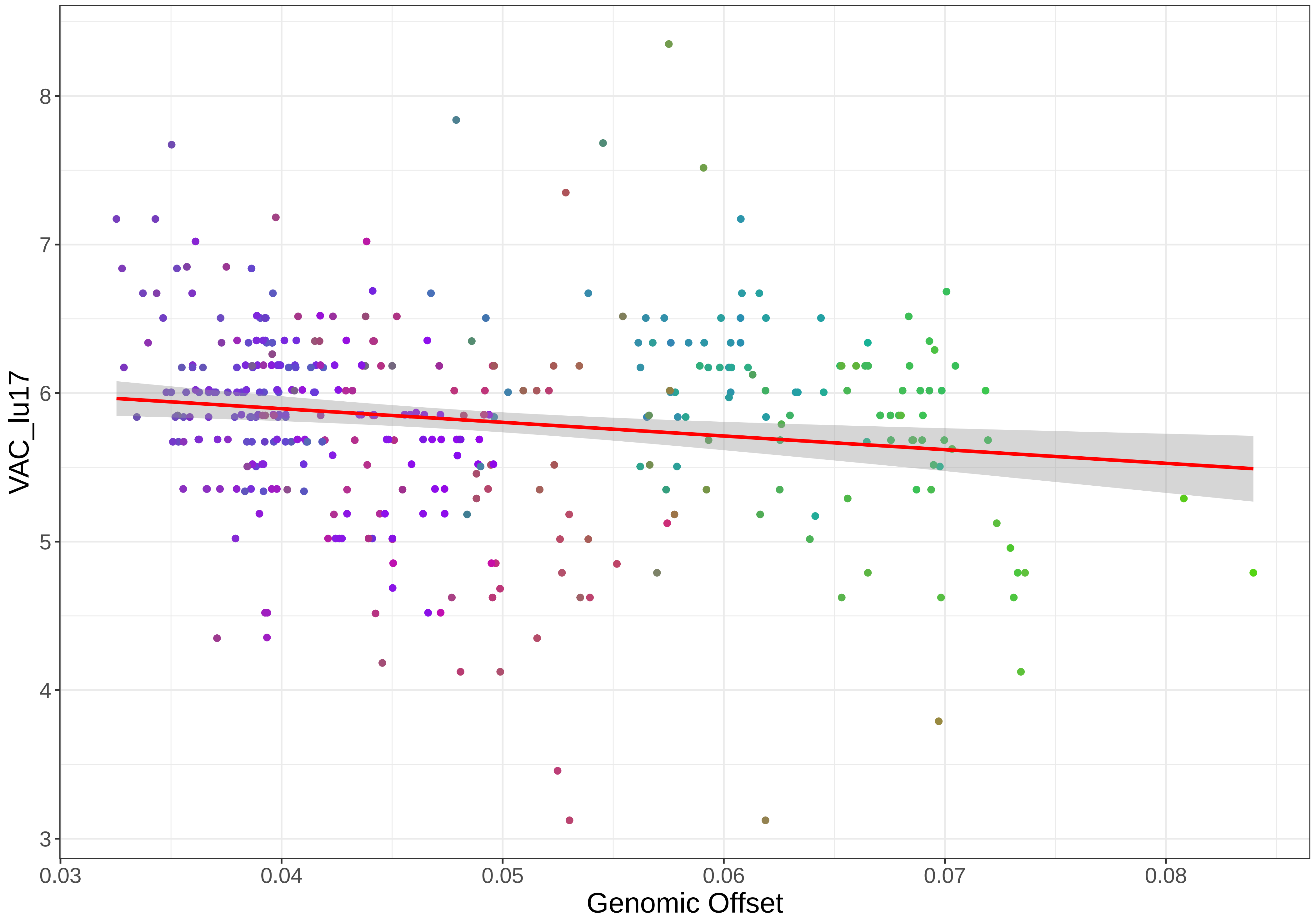
VAC_lu17
Genomic Offset
3
4
5
6
7
8
0.03
0.04
0.05
0.06
0.07
0.08

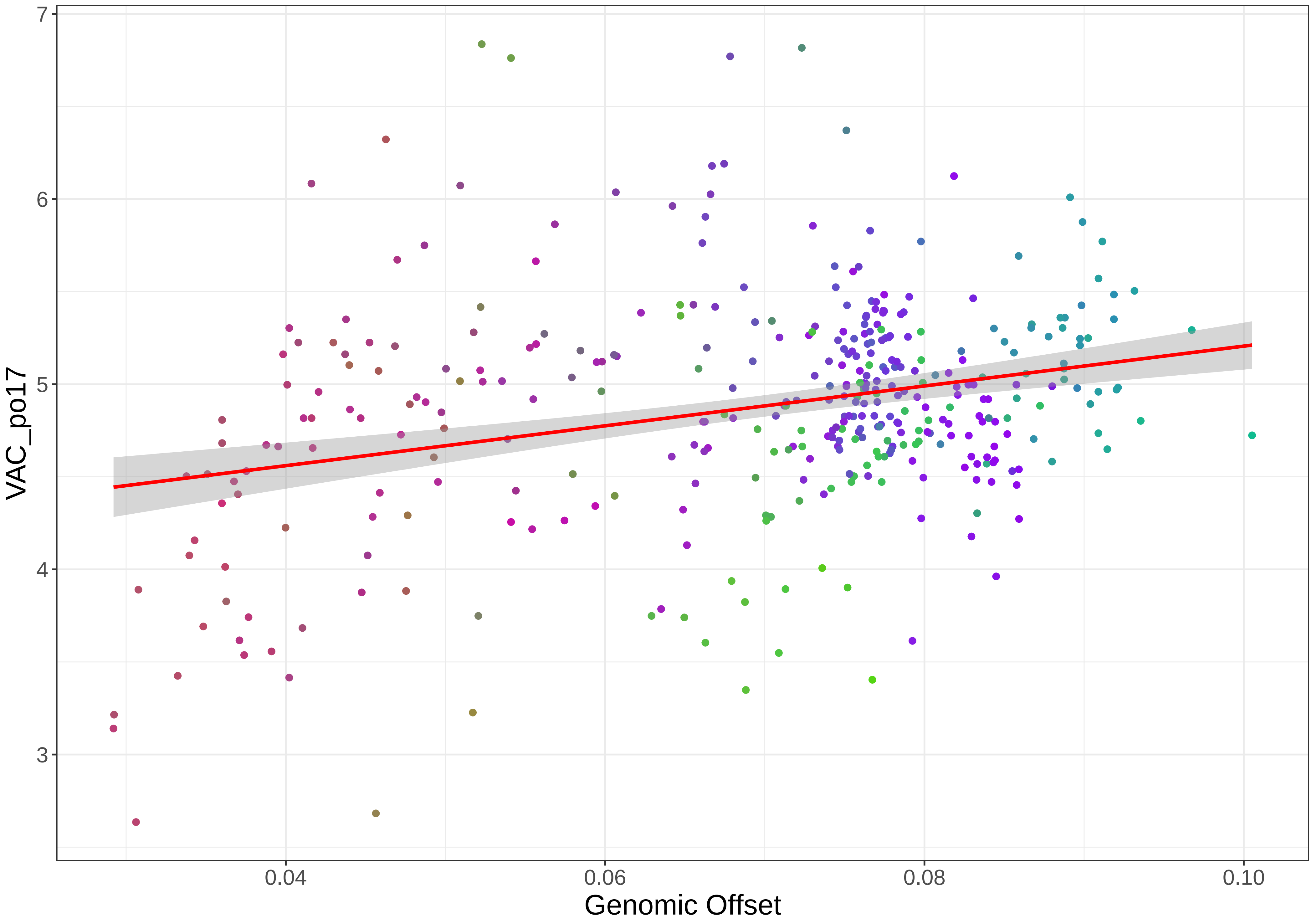

7
6
5
4
3
VAC_po17
0.04
0.06
0.08
0.10
Genomic Offset

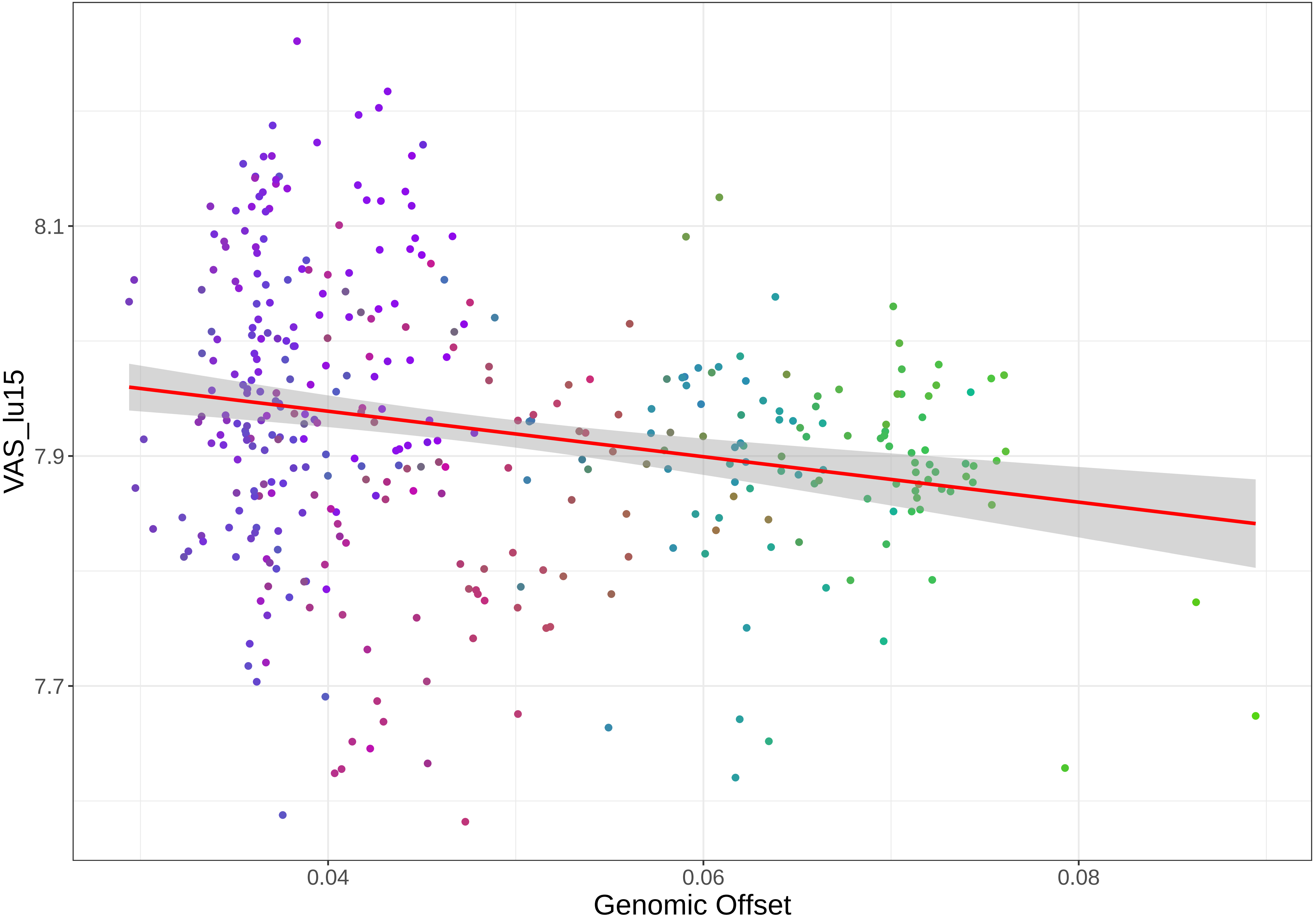

8.1
7.9
7.7
VAS_lu15
0.04
0.06
0.08
Genomic Offset

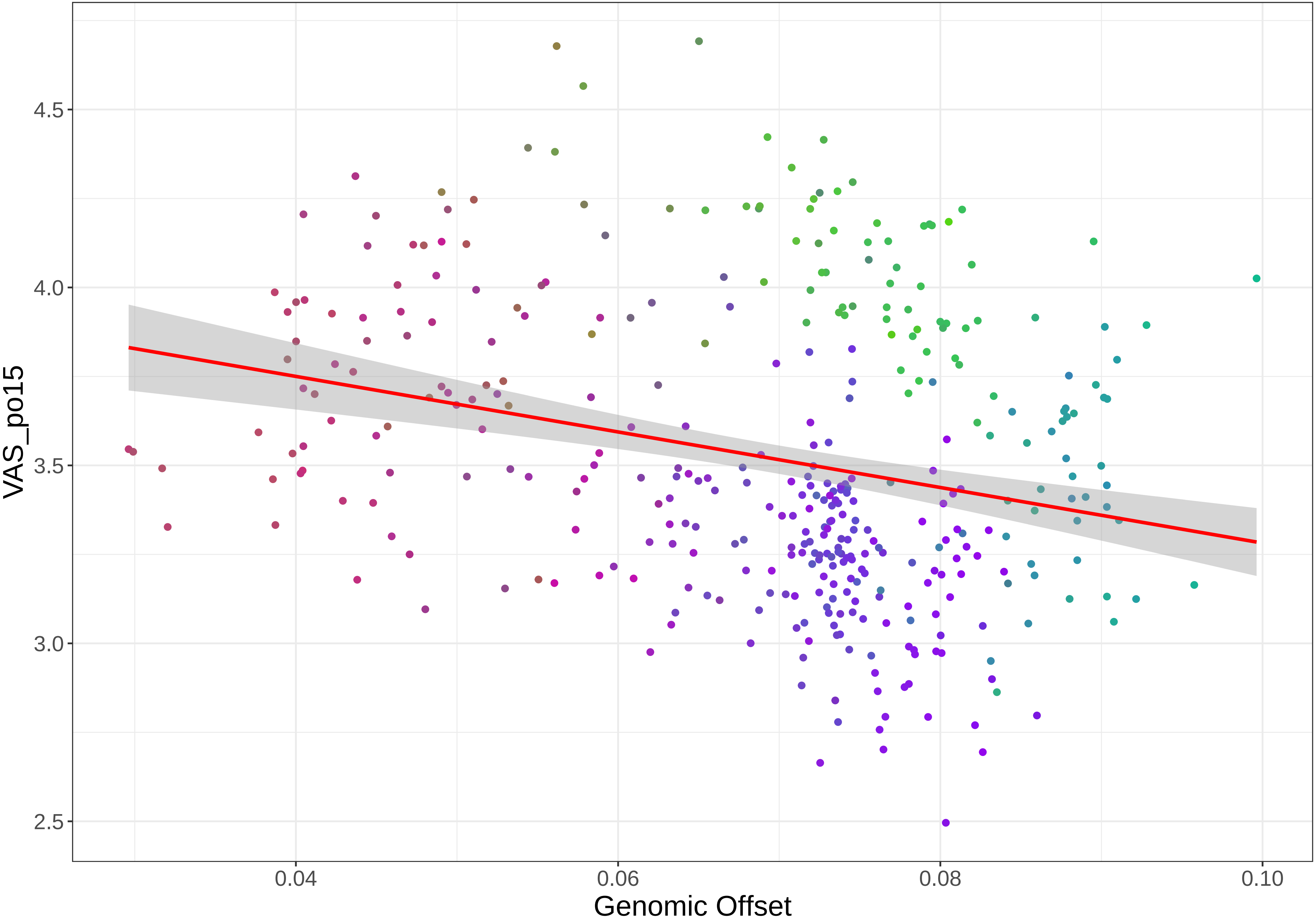

4.5
4.0
3.5
3.0
2.5
VAS_po15
0.04
0.06
0.08
0.10
Genomic Offset

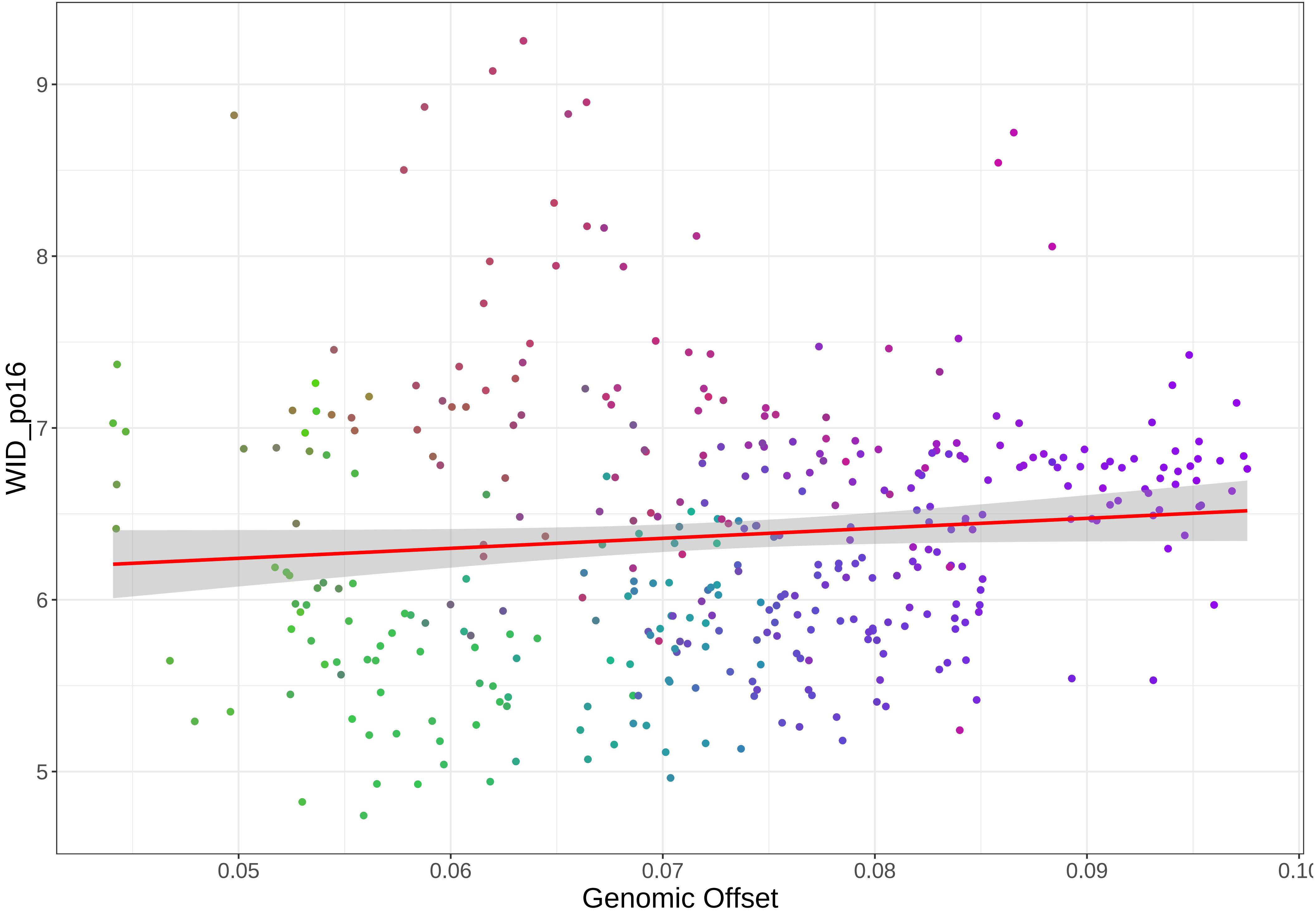

WID_po16
9
8
7
6
5
0.05
0.06
0.07
0.08
0.09
Genomic Offset

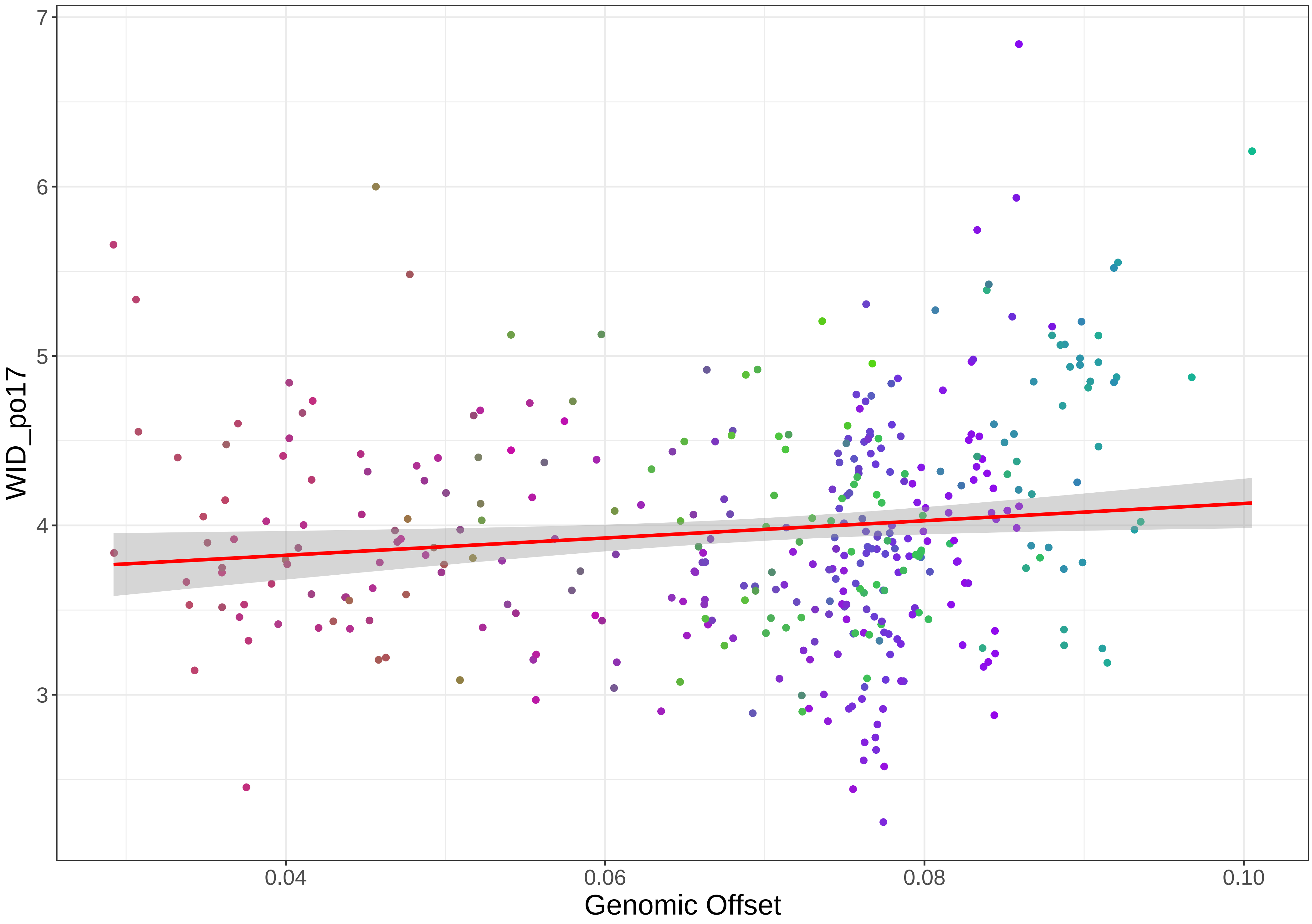
WID_po17
Genomic Offset
7
6
5
4
3
0.04
0.06
0.08
0.10

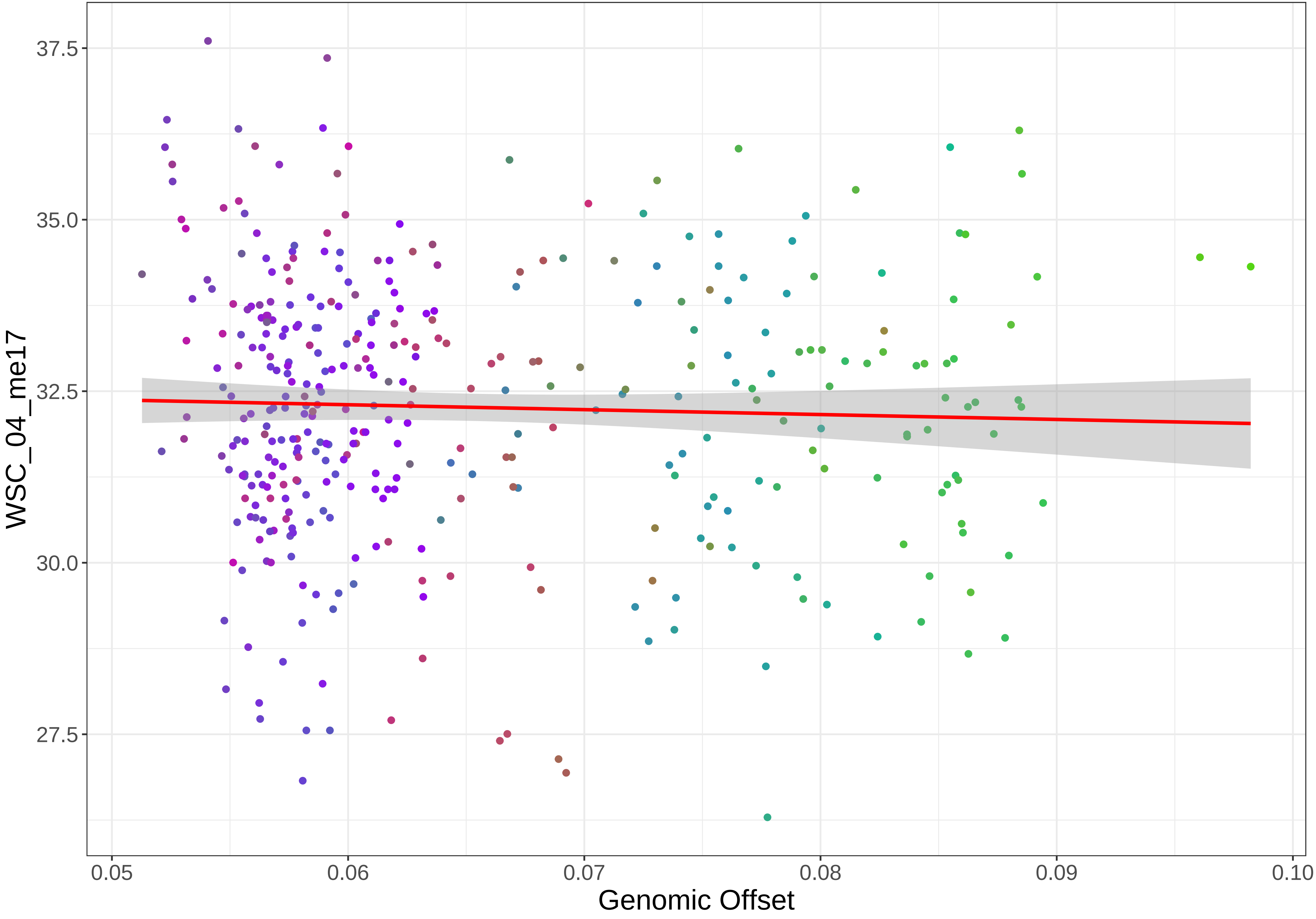
WSC_04_me17
37.5
35.0
32.5
30.0
27.5
0.05
0.06
0.07
0.08
0.09
0.10
Genomic Offset

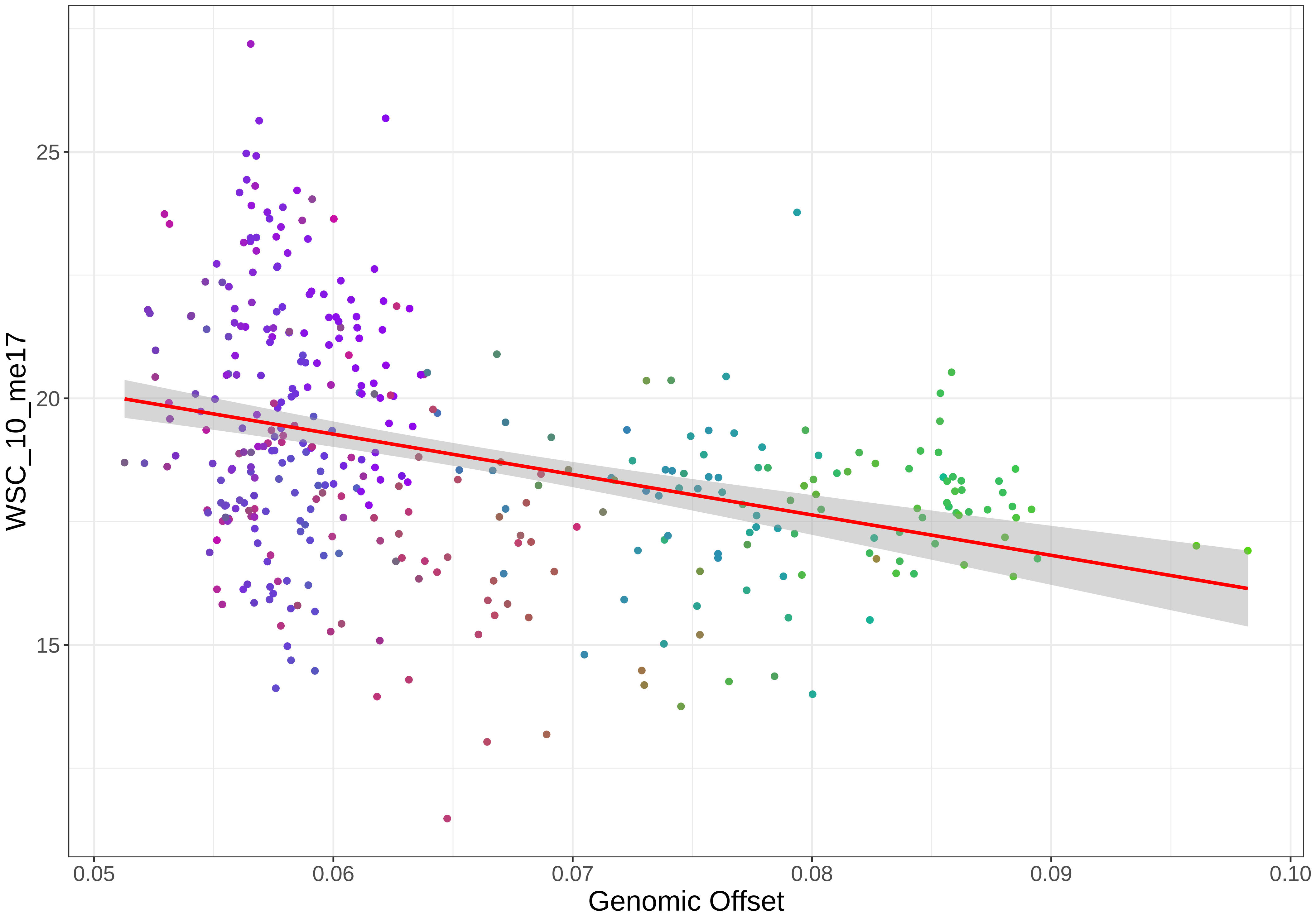

WSC_10_me17
25
20
15
0.05
0.06
0.07
0.08
0.09
0.10
Genomic Offset
